# A systems biology analysis of adrenergically stimulated adiponectin exocytosis in white adipocytes

**DOI:** 10.1101/2020.07.17.203703

**Authors:** William Lövfors, Christian Simonsson, Ali M. Komai, Elin Nyman, Charlotta S. Olofsson, Gunnar Cedersund

## Abstract

Circulating levels of the adipocyte hormone adiponectin are typically reduced in obesity and this deficiency has been linked to metabolic diseases. It is thus important to understand the mechanisms controlling adiponectin exocytosis. This understanding is hindered by the high complexity of the data and the underlying signaling network. To handle this complexity, we here analyze the data using systems biology mathematical modelling. Previously, we have developed a mathematical model for how different intracellular concentrations of Ca^2+^, cAMP and ATP affect adiponectin exocytosis (measured as increase in membrane capacitance). However, recent work has shown that adiponectin exocytosis is physiologically triggered via signaling pathways involving adrenergic β3 receptors (β_3_ARs). Therefore, we have herein developed a more comprehensive model that also includes adiponectin exocytosis stimulated by extracellularly applied epinephrine or the β_3_AR agonist CL 316,243. Our model can explain all previous patch-clamp data, as well as new data consisting of a combination of the intracellular mediators and extracellular adrenergic stimuli. Without changing the parameters, the model can accurately predict independent validation data with other combinations of patch-clamp pipette solutions and external stimuli. Finally, we use the model to perform new *in silico* experiments examining situations where corresponding wet lab experiments are difficult to perform. By this approach, we simulated adiponectin exocytosis in single cells, in response to the reduction of β_3_ARs that is observed in adipocytes from animals with obesity-induced diabetes. Our work brings us one step closer to understanding the intricate regulation of adiponectin exocytosis.

## INTRODUCTION

The white adipocyte hormone adiponectin is a major regulator of glucose and lipid homeostasis, with direct effects on the liver, muscle, the vasculature, and the pancreas (1, 2). Adiponectin levels are decreased in obese/type 2 diabetic individuals and preserved adiponectin levels are associated with a reduced risk of developing metabolic disease (3, 4).

Although much is known regarding the important pathophysiological roles of adiponectin, the mechanisms regulating its secretion from white adipocytes are less well studied. Our previous experimental work has defined a secretory pathway where adiponectin-containing vesicles are released in response to an elevation of intracellular cAMP and activation of Epac1 (*E*xchange *P*rotein directly *A*ctivated by *c*AMP, isoform 1). In addition, both Ca^2+^ and ATP have important roles for augmentation of adiponectin release as well as for maintenance of adiponectin secretion over extended time-periods (5, 6). Thus, the same mediators important for secretion of conventional hormones are involved in the regulation of adiponectin exocytosis (although hormone secretion is typically stimulated by an elevation of cytosolic Ca^2+^)(7). In later work, we showed that adiponectin secretion is physiologically stimulated via adrenergic signaling, chiefly involving beta3 adrenergic receptors (β_3_ARs). The same study demonstrates that adiponectin release is blunted in adipocytes from obese and diabetic mice, due to lower abundance of β_3_ARs in a state of *catecholamine resistance* (8). Clearly, the pathophysiological control of adiponectin secretion involves a highly complex signaling network of not yet fully understood interactions.

To deal with this high complexity, we have employed a systems biology approach combining experiments with mathematical modelling. We have previously developed a mathematical model for single-cell adiponectin exocytosis in 3T3-L1 adipocytes (9). In this work, changes in plasma membrane-capacitance is used to study exocytosis (10). Such measurements were utilized to model how vesicle exo- and endocytosis are affected by different combinations of intracellular Ca^2+^, cAMP, and ATP concentrations (9).The concentrations of intracellular mediators were altered by inclusion of different concentrations in the patch pipette attached to the cell. During the electrophysiological recordings, the pipette is in direct contact with the cell interior, allowing exchange between the pipette filling solution and the cell cytosol. We routinely measure adipocyte exocytosis in cultured adipocytes by this approach, and our studies have demonstrated that the cAMP-triggered and Ca^2+^-augmented capacitance increase in 3T3-L1 adipocytes largely represent secretion of adiponectin (5, 6, 8). Our model in (9) allowed us to extract detailed information contained in patch-clamp experiments and thus estimate the relative differences in adiponectin exo-and endocytosis rates. We could also correctly predict adiponectin secretion stimulated by cAMP in the presence or absence of Ca^2+^ or ATP. However, effects of extracellular cues, such as adrenergic stimulation, were not included in our previous model. This is an important shortcoming, since catecholamine stimulation constitutes a central stimulus-secretion pathway for adiponectin in normal physiological conditions.

In the current study, we have extended the model in Brännmark 2017 (9) to also include external, receptor-mediated, adrenergic stimulation. The resulting model can describe all previous capacitance data as well as new data of stimulation with epinephrine or the β_3_AR agonist CL 316,243 (CL). The model can explain data used for training, as well as accurately predict independent validation data. Simulations with the new and improved model allow us to better understand the mechanisms involved in the adrenergic control of adiponectin exocytosis.

## RESULTS

### Development of a new model including both pipette and external stimuli

By enlarging the pool of available data and evolving the mechanistic hypothesis, we expanded the model in (9) to include external receptor-mediated adrenergic stimulation (Fig. 1A). More specifically, we tested if the model could be improved to produce an acceptable agreement to the new expanded dataset, by doing several iterations of the lower branch in Fig. 1B. First, the original model was extended to include a more accurate representation of the pipette, where diffusion and the differences in volume between the pipette and intracellular compartments were taken into account. This was necessary because the external adrenergic stimulus present in the new data represents a more physiological way of elevating cytosolic cAMP, which differs from the previous studied effects of “clamping” cAMP levels by the pipette. With this more precise representation of the pipette introduced, adrenergic signaling was included in the model by adding the β_3_ARs and their effect on intracellular cAMP (Fig. 2).

**Figure 1.**
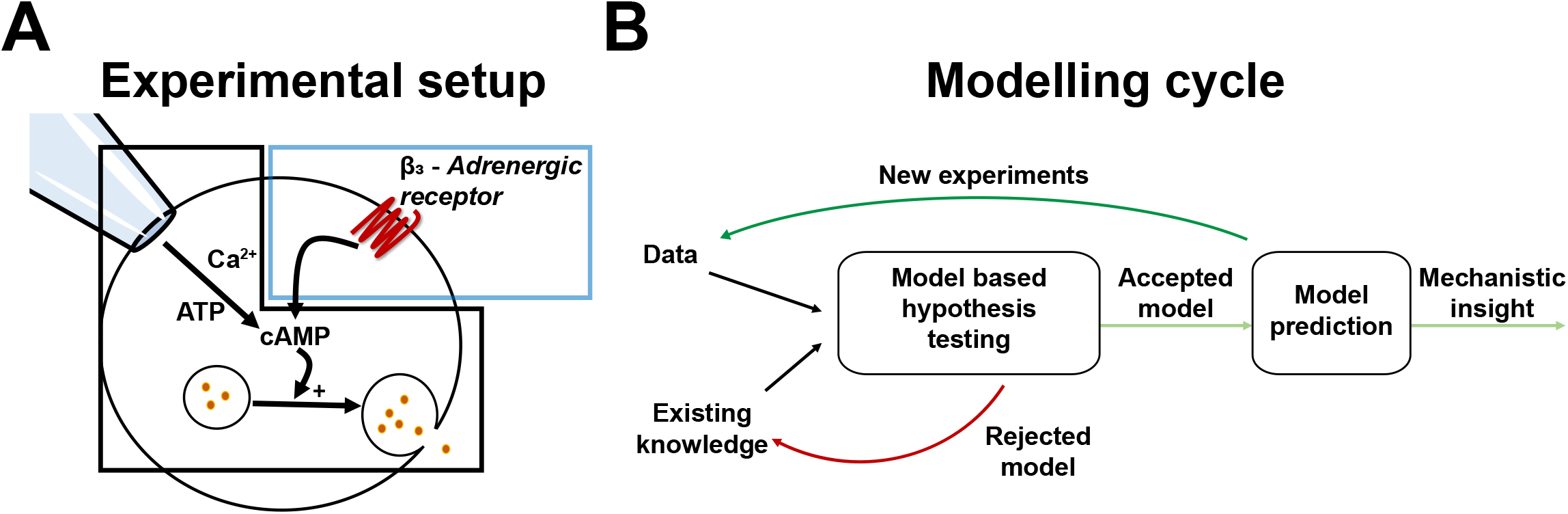
Overview of the experimental approach, and the modelling cycle. (A) The main variables of our previously published model of adiponectin exocytosis is contained within the black box, and the extension with adrenergic stimulus within the blue box. (B) A schematic overview of the model development cycle. Experimental data and existing knowledge are used to construct and test a hypothesis using mathematical simulations. If the model is rejected, the hypothesis must be revised. If it is not rejected, the model is used to predict new experiments which are later tested experimentally. A model that can accurately predict new experiments can be used to generate new insights into the detailed regulation of adiponectin vesicle release.

**Figure 2.**
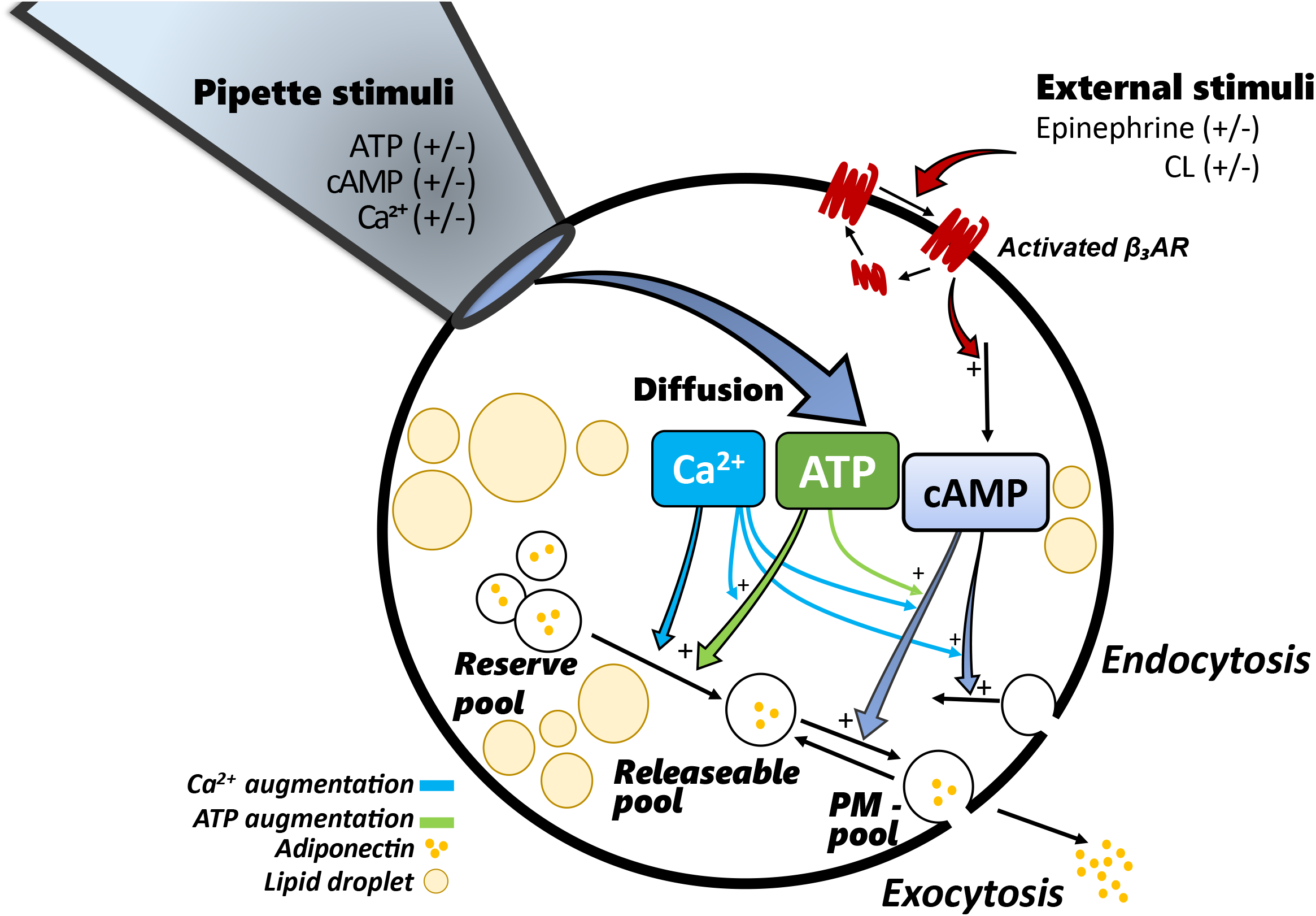
Detailed overview of our extended model. Stimulus in the model occurs either via the pipette solution (ATP, cAMP and Ca^2+^), or externally (epinephrine or CL). The pipette stimulus is allowed to diffuse between the pipette and the inside of the cell, while the external stimulus leads to activation of the receptor, which in turn leads to an increase in cytosolic cAMP. The transition of vesicles from the reserve pool to the releasable pool is stimulated by Ca^2+^, either alone or augmented with ATP. The fusion of adiponectin vesicles residing in the releasable pool is triggered by cAMP and augmented by both Ca^2+^and ATP. Vesicles in the PM-pool can either go back to the releasable pool or be exocytosed, independent of stimuli. Endocytosis is stimulated by cAMP and augmented by Ca^2+^. All equations and simulation files are available in Supplementary material.

### The model can describe all previous data as well as new data involving external adrenergic stimulations

Fig. 3 depicts the data used for developing the new model where each subfigure represents how a combination of extracellular (EC; adrenergic) and intracellular (IC; constituents of the pipette solution) input affects the cell. The new data including the response to external stimulation are shown in Fig. 3A-B, and the old data used in the development of the previous model (pipette stimulation only (9)) are shown in Fig. 3C-G (details are described in the legends). Adiponectin exocytosis is stimulated either by epinephrine (EPI) or CL with different concentrations of cAMP, ATP, and Ca^2+^ included in the pipette solution (IC). The experimental data points are denoted by black dots and error bars representing the mean and SEM values. The model simulations are shown as solid red lines for the best agreement with data (with optimal parameter set 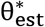), and the orange area represents the bounds of the model uncertainty. The model uncertainty was gathered by maximizing and minimizing the simulation in each experimentally measured time point, while still requiring a cost below the χ^2^-threshold. Clearly, the model agrees well with data, both for the EC and the IC stimulus data. This visual assessment of the good agreement is statistically supported by the fact that the model passes a 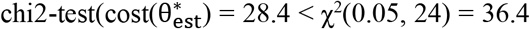; Materials and methods). As can be seen, the model uncertainties are reasonable and limited. In summary, the new model can describe all previous IC data as well as new data involving both IC and EC stimulation.

**Figure 3.**
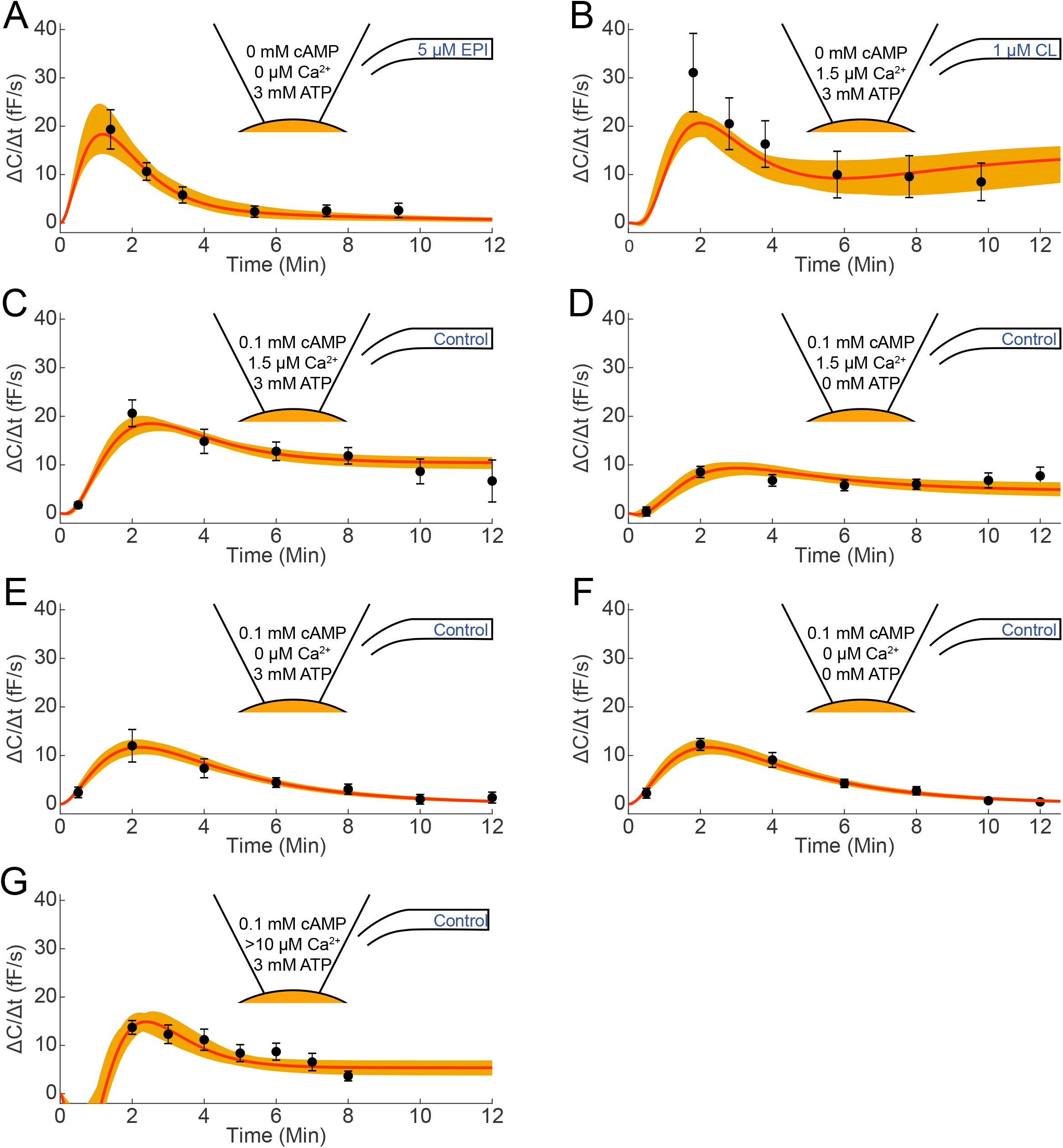
The model performance on estimation data, consisting of both intracellular pipette and extracellular adrenergic stimulation. (A-G) The black dots and error bars represent mean and SEM values of the experimental data, the red solid line represents the model simulation with the statistically best parameter set, and the orange area represents the model uncertainty. The exact IC and EC conditions are described within the symbolized pipette tip (different concentrations of cAMP, free Ca^2+^ and ATP) and within the symbol representing the glass capillary for the EC inlet (control, EPI or CL). The free [Ca^2+^] was calculated in (8). (A) n = 4 – 6, (B) n = 6 – 7, (C) n = 8, (D) n = 7, (E) n = 9, (F) n = 13, (G) n = 9 – 10

### The model is able to describe independent validation data involving other combinations of stimuli not used for model development

We next tested if the new model is able to correctly predict new independent validation data. The validation experiment consists of a combination of EC and IC inputs: 1 μM CL, and 3 mM ATP respectively (Fig. 4A). The prediction uncertainty was gathered in the same way as the model uncertainty, by maximizing and minimizing the prediction. Since the prediction has a low uncertainty (less than 30% at the peak value), it is meaningful to compare the prediction with experimental data, to test the quality of the model. This experimental data is shown in Fig. 4B. As can be seen, both quantitative and qualitative aspects are in agreement. At t = 2, simulations show a peak value between 11 and 20 fF/s, which is in excellent agreement with the experimental range of 13-20 fF/s. Note that it is not important that the ranges are identical, but only that they are overlapping. The agreement is also statistically confirmed by the fact that the model passes a χ^2^-test, with the optimal parameter set from the estimation 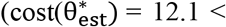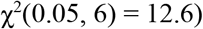.

**Figure 4.**
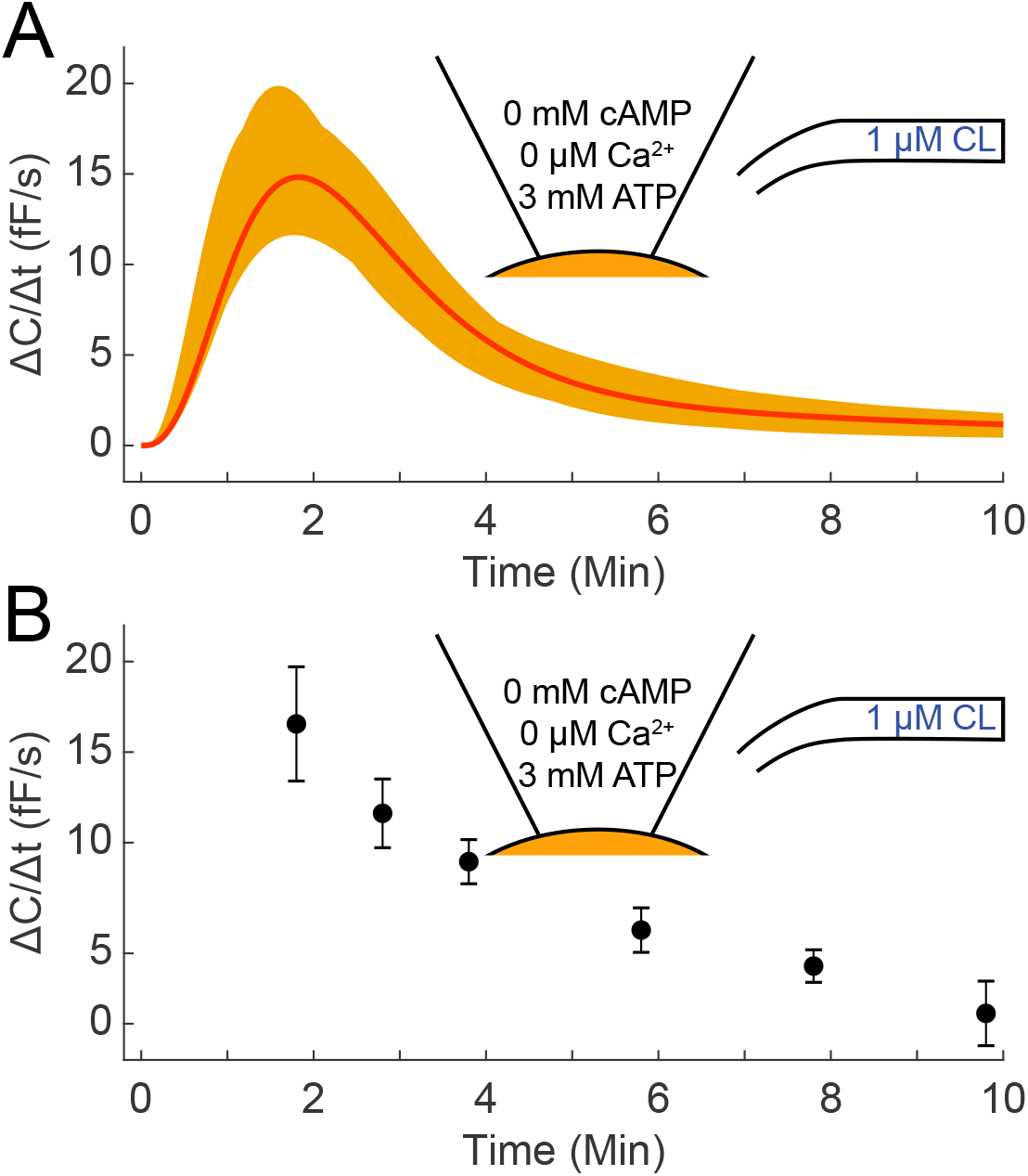
Independent model prediction and corresponding experimental data. (A) The stimulation with 1 μM CL EC in the presence of 3 mM ATP IC was predicted using the model. The solid red line represents the best parameter set with respect to the estimation data, and the orange area signifies the model prediction uncertainty. (B) The same experiment was performed experimentally, and the rate of exocytosis was measured at the indicated timepoints. Black dots and error bars denote mean and SEM values of the experimental data (n = 6 – 8).

### The model predicts a decline in peak secretion upon a decrease of β3 adrenergic receptors

The validated model can be used to predict the outcome of new experiments in single cells, given the mechanisms included in the model. To illustrate this capability, we simulated a new experiment that would be challenging to carry out experimentally. The simulated *in silico* experiment was inspired by published data demonstrating that adrenergically triggered adiponectin secretion (adiponectin collected during a 30 min static incubation) was almost completely abrogated in primary adipocytes isolated from obese and diabetic mice. This blunted secretion was associated with a 30% lower abundance of β_3_ARs (8), a condition we can also include in the model. Fig. 5A shows our own adiponectin secretion data from an experiment, where cultured 3T3-L1 adipocytes were genetically ablated for β_3_ARs (small interfering RNA reducing gene expression by ~60%) and stimulated with epinephrine during a 30 min static incubation. As can be seen, adiponectin release is increased two-fold in control (scramble) cells exposed to epinephrine whereas siRNA-treated adipocytes show a non-significant response. To experimentally investigate how reduced expression of β_3_ARs affects the molecular regulation of adiponectin exocytosis by applying single cell patch-clamp to siRNA-treated cells is demanding. We therefore used our model to examine the corresponding predictions *in silico*. The amount of β_3_AR was downregulated by 30% or 60%, thus corresponding to the two experiments described above, and the resulting exocytosis response was simulated. Fig. 5B shows the result, where the simulation consists of a combination of 5 μM EPI (EC) and 3 mM ATP (IC; no cAMP or Ca^2+^ included), for both control (orange area) and perturbed (blue striped areas) cells. As can be seen, the 30% and 60% downregulation of β_3_ARs is only translated into an approximate 20% and 47% decrease in peak exocytosis respectively; thus single-cell adiponectin exocytosis as understood by our model is significantly less affected than the experimental data on secretion indicates. As demonstrated in Fig. 5C, simulation with 1 μM CL EC, under the same IC conditions as in 5B, yielded similar results. Interestingly, as shown in Fig. 5D, inclusion of Ca^2+^ (IC) in the model partly rescues the reduced response (*c.f.* Fig. 5B and C). All ranges of peak inhibitions are given in Table 1.

**Table 1.** Model prediction of peak exocytosis inhibition. Summary of the predicted inhibition of adiponectin secretion upon a 30% and 60% reduction of β_3_ARs, using our new model for single cells. The three conditions I-III correspond to Figures 5B-D, and the quantifications are done for the parameter set resulting in the largest prediction, at the time point of the highest secretion.

| Experimental conditions |  | Inhibition (%) |  |
| --- | --- | --- | --- |
| | | $\beta_3$AR ↓ 30% | $\beta_3$AR ↓ 60% |
| I | 5 μM ADR, 3 mM ATP | 20 | 47 |
| II | 1 μM CL, 3 mM ATP | 21 | 48 |
| III | 1 μM CL, 3 mM ATP, 1.5 μM $\text{Ca}^{2+}$ | 9 | 24 |

**Figure 5.**
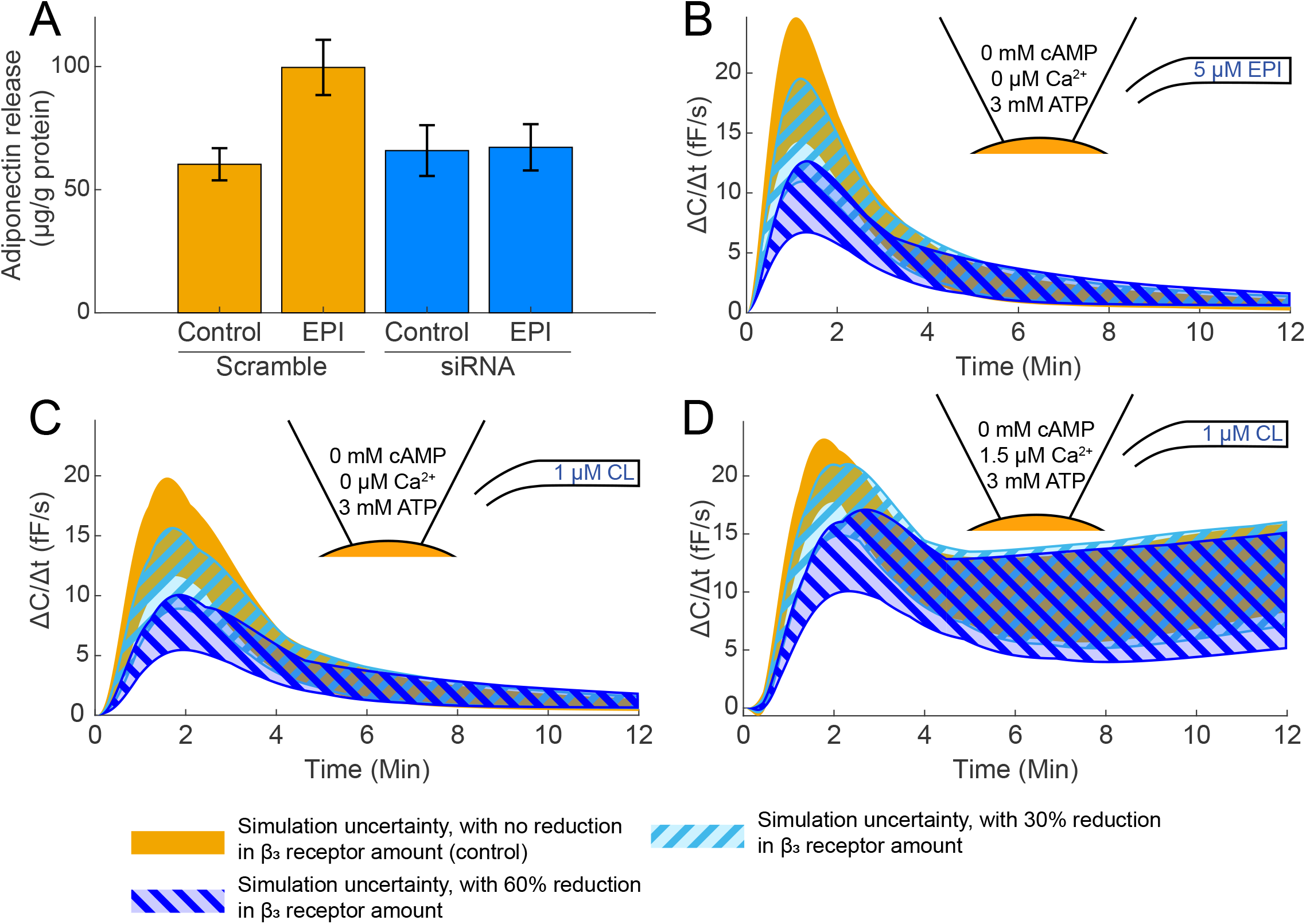
Model prediction of a 30 and 60% decrease in β3 adrenergic receptors. (A) Experimental data (mean and SEM values, n = 11) of absolute adiponectin secretion in control (scramble) and siRNA treated 3T3-L1 adipocytes, incubated with epinephrine through 30 minutes. Note that this data is produced from the experimental series in Fig. 7F of (5). (B-D) Model simulations upon downregulation of β_3_ARs. In all subfigures, the orange area corresponds to the prediction without reducing the amount of receptors and the striped areas represents the prediction with a reduction of the amount of β_3_ARs by 30% (cyan) and 60% (blue).

## DISCUSSION

The aim of this work was to develop a comprehensive mathematical model of adiponectin exocytosis that includes adrenergic stimulus, a chief signaling pathway involved in the physiological regulation of adiponectin secretion. To the best of our knowledge, no such model exists. The final model can explain a combination of experimental data and it can predict independent validation data as well as be used to simulate non-trivial predictions (alterations and measurements that are difficult to perform experimentally). Below we will discuss the usefulness of our model, as well as some of its limitations.

Although our mathematical model in (9) improved our understanding of the molecular regulation of adiponectin exocytosis, it was restricted to studying the role of intracellular cAMP, Ca^2+^ and ATP (5, 6). Our own recent research has shown that adiponectin exocytosis is triggered by catecholamine signaling via activation of β_3_ARs (8). In the current work, we have therefore developed a model that is able to describe the combination of internal and external (adrenergic) stimulus (Fig. 3), which thus brings us closer to a representative physiological model of adiponectin exocytosis. The model can describe both previous and new data (Fig. 3) and it can accurately predict independent validation data not used for estimating the parameter values (Fig. 4). This agreement with the data is supported by statistical tests showing that the data could have been generated by simulations with the model plus random noise.

We used the validated model to predict how adiponectin exocytosis would be affected if the abundance of β_3_ARs was reduced by 30%. This magnitude of decreased β_3_AR protein expression has been observed in primary subcutaneous adipocytes isolated from mice with obesity-induced diabetes (8). As shown in Fig. 5B and C (striped areas), the model yields a decrease in the exocytosis amplitude upon stimulation with either CL or EPI, when β_3_ARs are reduced. However, the rather slight decline in exocytosis amplitude stands in contrast to the complete abrogation of secretion observed in the “obese” adipocytes (8). Although the simulations and the data represent different conditions, i.e. a precise perturbation of single cell exocytosis vs. a more imprecise disease condition in a populations of cells, the observed differences argue that additional obesity-associated molecular disruptions need to be included in the model. One such mechanism could be the concomitant 30% reduction of Epac1 in the obese adipocytes; the cAMP-triggered adiponectin secretion occurs via activation of this signaling pathway (8). The obesity-associated reduced abundance of Epac1 likely contributes to the complete quenching of adiponectin secretion in the isolated cells. This is supported by that siRNA silencing of Epac1 in 3T3-L1 adipocytes abrogates EPI-stimulated adiponectin secretion (8). Moreover, we have recently demonstrated that gene expression of the exocytotic proteins VAMP (Vesicle-Associated Membrane Protein) 2 and 4 are reduced in visceral mouse adipocytes isolated from obese and diabetic mice (11). From this quick overview, it is apparent that several additional regulatory steps involved in the control of adiponectin exocytosis should be investigated further. Such future work, applying the types of predictions that we present here, will drive both further development of the mathematical model, and of associated experiments. In this integrated way, the model can be advanced and tested, to objectively refine and test our understanding of the adiponectin release mechanisms.

Diving further into our predictions and related experiments, Fig. 5 shows that even downregulating the β_3_AR expression by as much as 60% (blue striped area in Fig. 5B and C) does not abolish exocytosis to the extent shown for secretion in Fig. 5A. This is perhaps more perplexing than the difference between model and experimental data upon 30% downregulation of β_3_ARs, since the data in Fig. 5A represents an siRNA knockdown system, which can be regarded to be more “pure”, compared to a decreased expression owing to a pathophysiological condition. A plausible explanation for this second lack of agreement is that we compare responses in a population of cells during static conditions (secretion) with dynamic events in one single adipocyte (model). We cannot rule out the possibility that: 1) some adipocytes in Fig. 5A in fact release adiponectin in response to stimulation but that this population is too small to result in significantly elevated amounts of adiponectin in the medium; 2) adipocytes in the static incubations release substances that act on the cells in an autocrine or paracrine fashion to affect further secretion. Another alternative explanation is that a certain threshold level of β_3_ARs is required to maintain the functionality of the β_3_AR-cAMP-adiponectin signaling. Both cAMP and Ca^2+^ signaling is known to be contained to functional microdomains where signaling proteins and regulatory factors can interact (12, 13). A 30% decreased abundance of β_3_ARs is perhaps enough to critically disturb this organization. It should moreover be emphasized that the 60% siRNA knockdown represents gene expression and that we do not have data representing β_3_ARs protein level. Again, all of these mechanisms can and should be explored in future integrated experimental-modelling work.

As already described, Ca^2+^ alone is unable to stimulate exocytosis in 3T3-L1 adipocytes (5). In light of this, it may appear confusing that the elevation of intracellular Ca^2+^ in our model rescues the exocytosis response in adipocytes with reduced β_3_ARs (Fig. 5D and Table 1). However, our own data show that adiponectin secretion can be induced by the ionophore ionomycin (elevates intracellular Ca^2+^ to high levels) in adipocytes where endogenous cAMP levels are maintained (Musovic&Olofsson unpublished). We hypothesize that Ca^2+^, although unable to trigger exocytosis of adiponectin on its own, can act together with a non-stimulatory concentration of cAMP to induced secretion. It will be interesting to do joint experimental and modelling investigations to study this.

There are some technical details regarding the comparison with data that warrants further comments. For instance, some model simulations lie outside of the data uncertainty (c.f. Fig. 3B, t = 2, and Fig. 3G, t = {6,8}), but the agreement is still statistically acceptable. The model is trained to 47 data points, and to have a statistically acceptable agreement between model simulation and data, not all these data need to be in perfect agreement. Looking at Fig. 3B, the uncertainty in t = 2 is rather large and the cost of missing that data point is therefore small. In Fig. 3G, on the other hand, the uncertainty is rather small, and the cost of deviating the same distance from these data points is therefore higher. Since the model is statistically acceptable, there is no strong argument to improve the agreement to either of these data points. Moreover, to attain better agreements with the estimation data in Fig. 3, we would likely need to implement more detailed mechanisms. This could result in an overfitted model (i.e. a model that includes variations that are not representative of the “true” model structure), a poorer agreement with validation data (Fig. 4), and/or a higher uncertainty of the new predictions (Fig. 4A and Fig. 5). Another technical comment concerns the way we calculated these model uncertainties. The uncertainties were obtained by solving a modified optimization problem, suggested in (14), for each condition (minimization/maximization, experiment, and time point). The reformulated problem yielded wider estimations of the model uncertainties, compared to simply restarting the search for optimal parameter sets multiple times and saving all parameter sets (results not shown). In both these approaches, only parameter sets with sufficiently good agreement to data were used. We did not compare the reformulated problem against Markov Chain Monte-Carlo sampling (15) or prediction profile likelihood (16), but under the assumption that the reformulated problem is solved to optimality, our approach should yield at least an equally strong estimation of the uncertainties. Another aspect is that the uncertainties of the model simulations have different widths under the various experimental conditions; compare e.g. Fig. 3B to 3D. These different widths are explained by that the model uncertainty is largest in the datasets with the largest measurement uncertainty.

In conclusion, we here present the first model able to explain effects of both intracellular and extracellular cues on adiponectin exocytosis. The model is clearly useful for prediction of experimental data and shows promises to be valuable for gaining future mechanistic and molecular insight that would be difficult to obtain experimentally. In the more long-term perspective, these model-based insights can also be useful for medical applications. Obesity-associated diabetes is a common and costly disease, with severe complications, such as cardiovascular dysfunction, blindness, and kidney failure (17). In order to facilitate both prevention and treatment of diabetes, it is important to understand and study the mechanisms involved in metabolic regulation, as well as aberrations associated with disease development. Hypoadiponectinemia is one of several hallmarks in obesity-associated diabetes (4). We think that the work presented here has promising prospect to help solving a part of the puzzle, by enabling us to gain new knowledge about the pathophysiological regulation of adiponectin exocytosis. However, the progression into type 2 diabetes is the result of the breakdown of a complex interplay between several different organs, hormones, metabolites, and other factors. This complex interplay consists of several interacting subsystems, and both these subsystems and their interactions need to be understood, in order to fully comprehend the disease. The current model of adiponectin exocytosis cannot explain how adiponectin is controlled on an organ or whole-body level. An exciting way forward is to bring our model into the larger picture of the whole-body energy homeostasis, to both increase our understanding of normal physiology, and to acquire new knowledge of how adipose tissue dysfunction is connected to the development of type 2 diabetes and its associated comorbidities. Such advances can be done by combining our model with existing models at the molecular, organ and whole-body level (18–20).

## EXPERIMENTAL PROCEDURES

### Mathematical modelling

The model was constructed using ordinary differential equations (ODEs) in MATLAB and solved numerically using IQM tools, a continuation of SBtoolbox (21) available at https://iqmtools.intiquan.com, with Sundials CVODEs (22). The full model equations are given in the supplementary files. The performance of the model was evaluated using the objective function defined in Eq. 1, commonly referred to as the cost of the model.

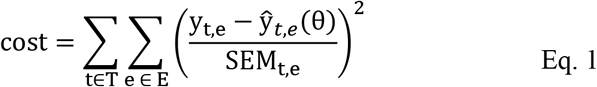

where y_t,e_ and SEM_t,e_ are the mean value and standard error of the mean (SEM) of the experimental data *e* at time *t*, T is the set of time points and E is the set of experiments, and 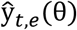 is the model simulation of experiment *e* at time *t*, depending on a set of parameters *θ*.

In order to evaluate the developed model, we use a χ^2^ -test which tests the null hypothesis that the experimental data have been generated by the model. In practice, the optimal cost given the optimal parameter set *θ\** is compared against the χ^2^ test statistic, with p = 0.05 and the degrees of freedom equal to the number of datapoints (minus one for the scaling parameter introduced in the new model). If the cost is larger than the χ^2^-limit, the model is rejected.

Due to the sparsity of the data, and prior knowledge about when the peak exocytosis should occur in the extracellularly stimulated experiments, we added constraints and rejected parameter sets that did not fulfil these conditions.

The optimal parameter set (*θ*\*) was found by using a serial combination of MathWorks particle swarm, simulated annealing, and fmincon solvers. This chain of solver was restarted multiple times and ran in parallel at the Swedish national supercomputing center (NSC). In the end, the current best estimation of the parameter set was used as a start guess for a combination of simulated annealing and fmincon, run multiple times at NSC.

The model uncertainty was estimated by maximizing and minimizing the model simulation in all available datapoints (i.e. for all timepoints and all experiments) independently. To constrain the uncertainty, the cost was required to be below the optimal cost plus the χ^2^-statistic with p = 0.05 and one degree of freedom. A description of the optimization problem is given in Eq. 2.

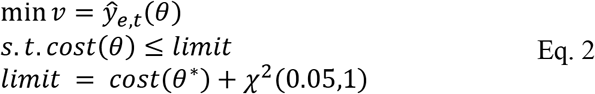

where 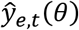 is the model simulation of experiment *e* and timepoint *t*, depending on the parameters *θ*. For finding the maximum simulation, the objective function of Eq. 3 was multiplied with −1 and solved as a minimization problem. In practice, the constraint was relaxed into the objective function and the formulation used is given in Eq. 3, where the penalty only is applied if *cost*(*θ*) > *limit*.

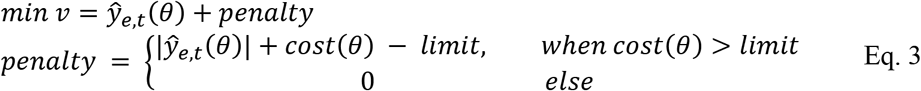

All subproblems were solved in parallel at NSC by using the current best estimation and running a chain of simulated annealing and fmincon repeated multiple times.

### Experimental methods

Patch-clamp measurements of exocytosis and adiponectin secretion experiments are described in detail in (9). In short, 3T3-L1 cells were grown in plastic petri dishes (Nunc, Denmark) or 12-well plates (Sarstedt) and differentiation was induced by addition of a differentiation cocktail (5). For electrophysiological recordings, the adipocytes were superfused with an extracellular solution (EC) containing (in mM): 140 NaCl, 3.6 KCl, 2 NaHCO3, 0.5 NaH2PO4, 0.5 MgSO4, 5 Hepes (pH 7.4 with NaOH), 2.6 CaCl2 supplemented with 5 mM glucose. The pipette-filling solutions contained (in mM): 125 potassium glutamate, 10 KCl, 10 NaCl, 1 MgCl2, 5 Hepes (pH 7.15 with KOH) and were supplemented with cAMP, ATP, and Ca2+ as indicated. Cells were voltage clamped at −70 mV and exocytosis was measured as increase in membrane capacitance in the standard whole-cell configuration, using an EPC-9 patch-clamp amplifier (HEKA Electronics, Lambrecht/Pfalz, Germany) and PatchMaster software. Exocytosis rates (ΔC/Δt) were measured by application of linear fits (OriginPro: OriginLab Corporation) at indicated time points. Secreted adiponectin was measured in EC collected from adipocytes incubated with indicated secretagogues during 30 min incubations at 32 °C. Secreted adiponectin (measured with mouse ELISA DuoSets; R&D Systems) was analyzed in relation to total protein content (Bradford protein assay). Small Interfering RNA experiments were performed as described in (8).

## Supporting information

All scripts used in this work

## DATA AVAILABILITY

All scripts and data used in this work are provided at https://github.com/willov/adiponectin-epi (DOI: 10.5281/zenodo.3979069), and is mirrored at https://gitlab.liu.se/ISBgroup/projects/adiponectin-epi.

## AUTHOR CONTRIBUTIONS

WL did most of the model-based analysis, and supervised the remaining modelling work done by CS. EN and GC supervised both WL and CS. Discussions on modelling details was done in regular meetings with WL, CS, EN, GC, and CSO. CSO supervised the experimental work performed by AMK. The manuscript was drafted by WL, CSO, and GC, with input from all authors. All authors read and approved of the final document.

