## Supplementary material for "A systems biology analysis of adrenergically stimulated adiponectin exocytosis in white adipocytes": All scripts used in this work: PlotFigures.m

```
%% Setup things
close all

cd('Scripts')

modelName='adr_endo';
fileDir='../Results/PPL/191220';
load('../Results/opt(28.3531).mat')
bestparam=optParam;

toEstimateOn={'Ca_ATP','Ca_noATP','noCa_ATP','noCa_noATP','highCa_ATP','EPI_ATP','CL_Ca'};
toValidateOn={'CL_ATP'};

[model, expData, estimation, validation, dgf]=Init(modelName, toEstimateOn, toValidateOn);

design=matlab.lang.makeValidName({'Ca+ATP';'Ca+noATP';'noCa+ATP';'noCa+noATP';'highCa+ATP';...
    'EPI+ATP';'CL+ATP';'CL+Ca'});
%   Ca,           ATP,         cAMP,   Epi, CL
stimulus=[...
    0.0015         3           0.1     0    0;   %'Ca+ATP'
    0.0015         0           0.1     0    0;   %'Ca+noATP'
    0              3           0.1     0    0;   %'noCa+ATP'
    0              0           0.1     0    0;   %'noCa+noATP'
    0.015          3           0.1     0    0;   %'highCa+ATP'
    0              3           0       5    0;   %'EPI+ATP'
    0              3           0       0    1;   %'CL+ATP'
    0.0015         3           0       0    1];   %'CL+Ca' 

experiments=table(stimulus(:,1:3), stimulus(:,4:end), 'VariableNames',{'Pipette','Agonist'},'RowNames',design);

%% Select what to plot
choice = input('What to Plot?  \n 1. Estimation  \n 2. Validation    \n 3. Prediction  \n Choice: ');

%% Load parameters
if choice ~=5
    costFun=@(param) CostFunction(param , model, expData, estimation, dgf);
    bestCost=costFun(optParam);
    sprintf('Cost: %.1f, dgf: %i, Chi2: %.1f', bestCost, dgf, chi2inv(0.95, dgf))
    
    if ~exist([fileDir '/allParams.mat'],'file')
        files=dir([fileDir '/**/*.mat']);
        nFiles=size(files,1);
        params=nan(nFiles,length(bestparam));
        for p=1:nFiles
            load([files(p).folder '/' files(p).name]);
             if round(costFun(optParam)*1e4)<=round((bestCost+chi2inv(0.95,1))*1e4)
                params(p,:)=optParam;
             else
                 disp('ERROR!!')
              end
            fprintf('Done loading %i of %i files\n',p,nFiles)
        end
        params(~any(~isnan(params),2),:)=[]; % Removes dummy parameter sets
        if ~exist(fileDir,'dir')
            mkdir(fileDir)
        end
    else
        load([fileDir '/allParams.mat'])
    end
   
    time=unique(expData{'Time',:});
end
%% Get uncertainty and plot
if choice == 1 % Estimation data
    [ boundry, bestSim] = MinMax(model, params, bestparam, expData, experiments(toEstimateOn,:));
    Plot_Exocytosis_Uncertainty (bestSim, boundry, expData)
elseif choice == 2 % Validation
    [ boundry, bestSim] = MinMax(model, params, bestparam, expData, experiments(toValidateOn,:));
    Plot_Exocytosis_Uncertainty (bestSim, boundry, expData)
    
elseif choice == 3 % Prediction
    exoExperiments=experiments({'EPI_ATP', 'CL_ATP', 'CL_Ca'},:);
    [boundry, bestSim] = MinMax(model, params, bestparam, expData, exoExperiments);
    boundry30 = MinMax(model, params, bestparam, expData, exoExperiments, 0.3);
    boundry60 = MinMax(model, params, bestparam, expData, exoExperiments, 0.6);
    
    for k = 1:length(exoExperiments.Properties.RowNames)
        subplot(2,2,k+1), Plot_Subplot(boundry, bestSim('Time',:), exoExperiments.Properties.RowNames(k), [], [])
        subplot(2,2,k+1), Plot_Subplot(boundry30, bestSim('Time',:), exoExperiments.Properties.RowNames(k), [], [66, 194, 245]/255)
        subplot(2,2,k+1), Plot_Subplot(boundry60, bestSim('Time',:), exoExperiments.Properties.RowNames(k), [], [0 0 1])
        axis([0 12 0 24.66])
        fprintf('----%s----\n',exoExperiments.Properties.RowNames{k})
        fprintf('Peak, Normal: %.2f\n',max(boundry{k,'Max'}));
        fprintf('Peak, Inhib30: %.2f\n',max(boundry30{k,'Max'}));
        fprintf('Peak, Inhib60: %.2f\n',max(boundry60{k,'Max'}));
        fprintf('Peak down, 30: %.2f\n',(1-max(boundry30{k,'Max'})/max(boundry{k,'Max'}))*100);
        fprintf('Peak down, 60: %.2f\n',(1-max(boundry60{k,'Max'})/max(boundry{k,'Max'}))*100);
    end
    
    load('fig5a-data.mat')
    subplot(2,2,1)
    b=bar([1 2 3 4],siRNA{'Mean',:});
    hold on
    
    b.FaceColor = 'flat';
    b.CData=[1 1 1; 1 1 1; 0.2 0.2  0.2; 0.2 0.2 0.2];
    errorbar([1 2 3 4],siRNA{'Mean',:},siRNA{'SEM',:},'marker','none','linestyle','none','color','k');
    set(gca, 'xticklabel', {'Control','Adr', 'Control','Adr'})
    set(gca, 'ytick',[0 0.5e-4 1e-4],'box','off')
end

cd('..')

%% Plotting functions
function [] = Plot_Exocytosis_Uncertainty(sim, boundry, expData)
design = sim.Properties.RowNames';
design(strcmp(design,'Time'))=[];
extra=7;
for e=design
    switch e{:}
        case 'Ca_ATP',            figure(1), subplot(4,2,3)
        case 'Ca_noATP',          figure(1), subplot(4,2,4)
        case 'noCa_ATP',          figure(1), subplot(4,2,5)
        case 'noCa_noATP',        figure(1), subplot(4,2,6)
        case 'highCa_ATP',        figure(1), subplot(4,2,7)
            
        case 'Ca_ATP_nocAMP',     figure(2), subplot(3,1,1)
        case 'noCa_ATP_nocAMP',   figure(2), subplot(3,1,2)
        case 'noCa_noATP_nocAMP', figure(2), subplot(3,1,3)
            
        case 'EPI_ATP',           figure(1), subplot(4,2,1)
        case 'CL_Ca',            figure(1), subplot(4,2,2)
            
        case 'CL_ATP'
            figure(1);
            hold on
            shadeColor=[0.95,0.65,0];
            Plotcolor=[1, 0.2, 0];
            simTime=sim{'Time','Measures'}./60;
            subplot(2,1,1)
            hold on
            title('Simulation')
            xx=[simTime fliplr(simTime) ];
            yy=[boundry{e,'Min'} fliplr(boundry{e,'Max'})];
            h=fill(xx,yy,shadeColor);
            set(h, 'EdgeColor','None');
            plot(simTime,sim{e,'Measures'},'color', Plotcolor , 'LineWidth' , 1.5 ) %Plot the best parameter set
            xlabel('Time (Min)')
            ylabel('\DeltaC/\Deltat (fF/s)')
            axis([-0.2 10 -2.5 20])
            box off
            
            subplot(2,1,2)
            errorbar(expData{'Time',e}./60,expData{'Mean',e},expData{'SEM',e},'ko', 'MarkerFaceColor','auto', 'MarkerFaceColor','auto')
            title('Experimental data','Interpreter', 'none')
            xlabel('Time (Min)')
            ylabel('\DeltaC/\Deltat (fF/s)')
            axis([-0.2 10 -2.5 20])
            box off
            sgtitle('External stimulus, CL no Ca')
            
            figure(15)
            tInd=ismember(simTime,expData{'Time',e}./60);
            y=sim{e,'Measures'};
            
            errorbar(expData{'Time',e}./60,expData{'Mean',e},expData{'SEM',e},'ko', 'MarkerFaceColor','auto', 'MarkerFaceColor','auto')
            hold on
            plot(simTime(tInd),y(tInd))
            
            cost= nansum((expData{'Mean',e}-y(tInd)).^2./expData{'SEM',e}.^2);
            dgf=length(expData{'Mean',e});
            sprintf('Prediction, CL_ATP, Cost: %.1f, dgf: %i, Chi2: %.1f', cost, dgf, chi2inv(0.95, dgf))
    
            
        otherwise,                figure(extra), extra=extra+1;
    end
    Plot_Subplot(boundry, sim, e, expData)
end

figure(1), SetSameAxes(1,[0,2],[],[],0)
end

function []=Plot_Subplot(boundry, bestSim,e, data, shadeColor)
hold on
if nargin<5 || isempty(shadeColor)
shadeColor=[0.95,0.65,0];
alpha=1;
else
    alpha=1;
end
bestColor=[1, 0.2, 0];
simTime=bestSim{'Time','Measures'}./60;
if any(strcmp(boundry.Properties.RowNames,e))
    xx=[simTime fliplr(simTime) ];
    yy=[boundry{e,'Min'} fliplr(boundry{e,'Max'})];
    h=fill(xx,yy,shadeColor);
    set(h, 'EdgeColor','None');
    set(h,'facealpha',alpha)
end
if any(strcmp(bestSim.Properties.RowNames,e))
    plot(simTime,bestSim{e,'Measures'},'color', bestColor , 'LineWidth' , 1.5 ) %Plot the best parameter set
end
if ~isempty(data) && any(strcmp(data.Properties.VariableNames,e))
    errorbar(data{'Time',e}./60,data{'Mean',e},data{'SEM',e},'ko', 'MarkerFaceColor','auto')
end

title(e,'Interpreter', 'none')
xlabel('Time (Min)')
ylabel('\DeltaC/\Deltat (fF/s)')
axis tight
box off
end

%% Estimating uncertainty
function [ boundry, sim] = MinMax(model, params, bestparam, expData, experiments, betaReduction)
if nargin<6
    betaReduction=[];
end
params=[params; bestparam];
for i = 1 : size(params,1)
    try
        intialconditions = SimulateSteadyState(params(i,:), model, 10000, betaReduction);
        [sim,~,~, maxcAMP]=SimulateExperiments(params(i,:), 0:1:720, intialconditions, model, expData, experiments); % Simulates the experiments.
        
        if i == 1
            minValues=sim.Measures;
            maxValues=minValues;
        end
        if maxcAMP<inf
            sim_values=sim.Measures;
            minValues(minValues>sim_values)=sim_values(minValues>sim_values);
            maxValues(maxValues<sim_values)=sim_values(maxValues<sim_values);
        end
        
    catch err
        disp('error in sim');
    end
    fprintf('Done with %i of %i parameter sets\n',i,size(params,1))
end
boundry=table(maxValues, minValues,'VariableNames',{'Max','Min'},'RowNames',sim.Properties.RowNames);
end
```
