## Supplementary material for "A systems biology analysis of adrenergically stimulated adiponectin exocytosis in white adipocytes": All scripts used in this work: readme.md

# A systems biology analysis of adrenergically stimulated adiponectin exocytosis in white adipocytes 

These files are used to generate the results and the plots in the paper published with the same title. 
To run the scripts, MATLAB 2018b or newer is necessary (for the function `sgtitle` used when plotting), and the toolboxes `global optimization toolbox` and `parallel computing toolbox` are needed to run the parameter estimation scripts. Furthermore, a valid c compiler is necessary. Check if MATLAB has one available by running `mex -setup`

Below is a list of the scripts and what they do. 
#### PlotFigures
Used to generate the figures in the paper. Some postprocessing was done in illustrator. 

#### RunOptimization and RunOptimizationPred
A top level script used to run multiple parameter estimations in parallel. 

#### runParallel and runParallelPred
Top level scripts used to run parameter estimation in parallel at the super computer center. 

#### EstimateParameters and EstimateParametersPred
The scripts that runs the actual parameter estimations. 

#### CostFunction and CostFunctionPred
Used to estimate the objective function value when either finding the optimal solution, or when finding the boundary of the predictions. 

#### SimulateExperiments
A function used to simulate corresponding biological experiments. 

#### SimulateSteadyState
A function use to simulate steady state before the experiment is started. 

#### adr_end.txt
The actual model file being used. All model equations are contained in this file.

#### expData
A MATLAB file containing the experimental data. 
