## Supplementary material for "A systems biology analysis of adrenergically stimulated adiponectin exocytosis in white adipocytes": All scripts used in this work: CostFunctionPred.m

```
function [ v, costTot ] = CostFunctionPred( param , model, expData,estimation, validation, dgf, p,t)

time=unique([0:1:max(expData{'Time',:}) expData{'Time',:}]);

SSsim_step=10000; % Resolution of the steady state simulation
experiments=estimation.Properties.RowNames';

try
    %% Check if cost < limit
    [intialconditions,SteadyState_cost] = SimulateSteadyState(param, model, SSsim_step);
    SS_cost=SteadyState_cost; 
    [sim,~,~, ~]=SimulateExperiments(param, time, intialconditions, model, expData, estimation); % Simulates the experiments.
    
    peakCost=0;
    cost=0;
    for e = experiments
        tInd=ismember(sim{'Time','Measures'},expData{'Time',e}); % Finds which time points to use when comparing simulation to experimental data
        y=sim{e,'Measures'};
        cost=cost+sum((y(tInd)-expData{'Mean',e}).^2./expData{'SEM',e}.^2); %
        
        if strcmp(e,'EPI_ATP')
            if  sim{e,'PeakTime'}<1.1 || sim{e,'PeakTime'}>1.7
                peakCost=peakCost+chi2inv(0.95,dgf)*(1+abs(1.4-sim{e,'PeakTime'}));
            end
        elseif strcmp(e,'CL_Ca')
            if  sim{e,'PeakTime'}<1.6 || sim{e,'PeakTime'}>2.0
                peakCost=peakCost+chi2inv(0.95,dgf)*(1+abs(1.8-sim{e,'PeakTime'}));
            end
        end
    end
    
    costTot = cost + SS_cost + peakCost; %
    
    
    if isnan(costTot)
        costTot=1e99;
    end
    
    
    %% check valid
    
    experiment=validation.Properties.RowNames';
    sim=SimulateExperiments(param, time, intialconditions, model, expData, validation); % Simulates the experiments.
    y=sim{experiment,'Measures'};
    
    tInd=ismember(sim{'Time','Measures'},t); % Finds which time points to use when comparing simulation to experimental data
    

    bestCost=28.3531;
    violation=costTot -(bestCost+chi2inv(0.95,1));
    if violation > 0
        v=p*y(tInd)+abs(y(tInd))+violation;
    else
        v=p*y(tInd);
    end
    
    
catch err
    disp(getReport(err))
    costTot=1e99;
end

end
```
