## Supplementary material for "A systems biology analysis of adrenergically stimulated adiponectin exocytosis in white adipocytes": All scripts used in this work: EstimateParameters.m

```
function EstimateParameters(worker,folderName, useBest, show_iter)
if nargin<1, worker=0; end
if nargin<2 || isempty(folderName), folderName='Temp'; end
if nargin<3, useBest=1; end
if nargin<4, show_iter=1; end

s=rng('shuffle');
s=rng(s.Seed+worker);

modelName='adr_endo';
allData={'Ca_ATP','Ca_noATP','noCa_ATP','noCa_noATP','highCa_ATP','EPI_ATP','CL_ATP','CL_Ca'};
toPredict={'CL_ATP'};
startTime=datestr(now,'yymmdd-HHMMSS');
toTrainOn = allData(~ismember(allData, toPredict));

[model, expData, estimation, validation, dgf, pNames, nParams, lb, ub]=Init(modelName, toTrainOn, toPredict);   % Sets necessary variables, depending on which model is being run.


folder=sprintf('../Results/%s/', folderName);
if ~exist(folder,'dir')
    mkdir(folder)
end
costFun=@(param) CostFunction( param , model, expData, estimation, dgf);


if show_iter
    saOpts = optimoptions(@simulannealbnd,'HybridFcn',@fmincon,'Display','iter'); % Simulated annealing optimization settings
    psOpts = optimoptions(@particleswarm,'Display','iter','useparallel',false);
else
    psOpts = optimoptions(@particleswarm,'Display','final','useparallel',false);
    saOpts = optimoptions(@simulannealbnd,'HybridFcn',@fmincon,'Display','final'); % Simulated annealing optimization settings
end
if ~useBest
    [optParamPS, minfunPS]=particleswarm(costFun, nParams, lb,ub,psOpts); % Optimize parameters using particle swarm.
    save(sprintf('%s/optPS(%.2f) - [%s], %s-%i.mat',folder, minfunPS, strjoin(toPredict), datestr(now,'yymmdd-HHMMSS'), s.Seed)  ,'optParamPS') % Saves the best parameters found.
else
    files=dir('../Results/**/opt*');
    if ~isempty(files)
        [~,ind]=natsortfiles({files.name});
        fileVar = load([files(ind(1)).folder '/' files(ind(1)).name]); %ascend or descend if min or max
        optParamPS=cell2mat(struct2cell(fileVar));
        minfunPS=costFun(optParamPS);
    else
        [optParamPS, minfunPS]=particleswarm(costFun, nParams, lb,ub,psOpts); % Optimize parameters using particle swarm.
    end
end
if minfunPS<100
    [optParam, minfun]=simulannealbnd(costFun, optParamPS, lb,ub,saOpts);
    save(sprintf('%s/opt(%.6f) - [%s], %s-%i.mat',folder, minfun, strjoin(toPredict),  startTime, s.Seed)  ,'optParam') % Saves the best parameters found.
end
if ~isempty(fid)
    fclose(fid)
end
end
```
