## Supplementary material for "A systems biology analysis of adrenergically stimulated adiponectin exocytosis in white adipocytes": All scripts used in this work: EstimateParametersPred.m

```
function EstimateParametersPred(worker,folderName, useBest, show_iter)

if nargin<1, worker=0; end
if nargin<2 || isempty(folderName), folderName='Temp'; end
if nargin<3, useBest=1; end
if nargin<4, show_iter=1; end

s=rng('shuffle');
s=rng(s.Seed+worker);

if nargin<3
    useBest = 1;
end
if show_iter
    saOpts = optimoptions(@simulannealbnd,'HybridFcn',@fmincon,'Display','iter'); % Simulated annealing optimization settings
    psOpts = optimoptions(@particleswarm,'Display','iter','useparallel',true);
else
    psOpts = optimoptions(@particleswarm,'Display','final','useparallel',true);
    saOpts = optimoptions(@simulannealbnd,'HybridFcn',@fmincon,'Display','final'); % Simulated annealing optimization settings
end

modelName='adr_endo';
toTrainOn={'Ca_ATP','Ca_noATP','noCa_ATP','noCa_noATP','highCa_ATP','EPI_ATP','CL_Ca'};
toPredict={'Ca_ATP','Ca_noATP','noCa_ATP','noCa_noATP','highCa_ATP','EPI_ATP','CL_Ca', 'CL_ATP'};

folder=sprintf('../Results/PPL/%s/', folderName);
if ~exist(folder,'dir')
    mkdir(folder)
end
Init(modelName);
bestCost = 28.3531;

for i = 1:100
%% Find the least searched experiment
for k = 1:length(toPredict)
    files=dir(sprintf('./Results/PPL/**/[%s*.mat', toPredict{1,k}));
    toPredict{2,k}=length(files);
end
toPredict=sortrows(toPredict',2)';
toPredict(2,:)=[];
e=toPredict(1);
[model, expData, estimation, validation, dgf, ~, ~, lb, ub]=Init(modelName, toTrainOn, e);   % Sets necessary variables, depending on which model is being run.
experimentTimes=expData{'Time',e};
experimentTimes(experimentTimes==0)=[];
%% Find the least searched time point
experimentTimes=[experimentTimes experimentTimes];
experimentTimes(2,:)=[-1*ones(1,length(experimentTimes)/2) ones(1,length(experimentTimes)/2)];
experimentTimes(2,length(experimentTimes)/2)=-1;

for k = 1:size(experimentTimes,2)
    
   if experimentTimes(2,k) == -1
       direction = 'max';
   elseif experimentTimes(2,k) == 1
       direction ='min';
   end
    
    files=dir(sprintf('../Results/PPL/**/[%s, %s, t%.2f]*.mat',e{:}, direction, experimentTimes(1,k)));
    experimentTimes(3,k)=length(files);
end
experimentTimes=sortrows(experimentTimes',3)';
experimentTimes(2:3,:)=[];
t=experimentTimes(1);

minFiles=length(dir(sprintf('../Results/PPL/**/[%s, min, t%.2f]*.mat',e{:}, experimentTimes(k))));
maxFiles=length(dir(sprintf('../Results/PPL/**/[%s, max, t%.2f]*.mat',e{:}, experimentTimes(k))));

if minFiles>maxFiles
    p=-1;
else
    p=1;
end
startTime=datestr(now,'yymmdd-HHMMSS');
costFun=@(param) CostFunctionPred( param , model, expData, estimation, validation, dgf, p,t);

if ~useBest
    [optParam, minfun]=particleswarm(costFun, nParams, lb,ub,psOpts); % Optimize parameters using particle swarm.
    
    if round(minfun*1e4)<=round((bestCost+chi2inv(0.95,1))*1e4)
        save(sprintf(saveStr,folder, e{:}, t,'opt', p*minfun,  startTime, s.Seed)  ,'optParam') % Saves the best parameters found.
    end
else
    if p==1
        saveStr='%s/[%s, min, t%.2f] %s(%.3f), %s-%i.mat';
        files=dir(sprintf('../Results/PPL/**/[%s, min, t%.2f]*.mat', e{:}, t));
        pos = 1;
    else
        saveStr='%s/[%s, max, t%.2f] %s(%.3f), %s-%i.mat';
        files=dir(sprintf('../Results/PPL/**/[%s, max, t%.2f]*.mat', e{:}, t));
        pos=length(files);
    end
    
    if ~isempty(files)
        [~,ind]=natsortfiles({files.name});
        files=files(ind);
        for n=0:length(files)-1 %Some numerical differences might yield nonvalid start guesses.
            load([files(pos+p*n).folder '/' files(pos+p*n).name]); %ascend or descend if min or max
            if exist('optParam','var')
                [~,cost]=costFun(optParam);
            else
                cost=inf;
            end
            if  round(cost*1e4)<=round((bestCost+chi2inv(0.95,1))*1e4)
                break;
            else
                if ~exist(strrep(files(pos+p*n).folder,'PPL','PPL_notValid'),'dir')
                    mkdir(strrep(files(pos+p*n).folder,'PPL','PPL_notValid'))
                end
                movefile([files(pos+p*n).folder '/' files(pos+p*n).name],...
                    [strrep(files(pos+p*n).folder,'PPL','PPL_notValid') '/' files(pos+p*n).name]);
            end
        end
    end
    if isempty(files)  || n>length(files)
        load('../Results/opt(28.3531).mat')
        fprintf('Missing opt, exp=%s, p=%i, t=%i\n',e{:},p,t)
    end
end
optParam=simulannealbnd(costFun, optParam, lb,ub,saOpts);
[minfun,cost]=costFun(optParam);
if round(cost*1e4)<=round((bestCost+chi2inv(0.95,1))*1e4)
    save(sprintf(saveStr,folder, e{:}, t,'opt', p*minfun,  startTime, s.Seed)  ,'optParam') % Saves the best parameters found.
else 
    disp('Not a valid solution') 
end
end


end
```
