## Supplementary material for "A systems biology analysis of adrenergically stimulated adiponectin exocytosis in white adipocytes": All scripts used in this work: Init.m

```
function [model, expData, estimation, validation, dgf, pNames, nParams, lb, ub]=Init(modelName, toEstimateOn, toValidateOn)
if ~exist('IQMsimulate','file')
    fprintf('\n\nTo use these scripts, a valid c-compiler is necessary. \nIf things fail, check if MATLAB is registering a valid compiler with the command "mex -setup"\n\nThe toolbox and modell will now be compiled. \nPress any key to continue.\n')
    pause;
    run('./Support/IQM Tools/installIQMtoolsInitial.m')
end

addpath(genpath('./Support'))
addpath('../Results');
IQMmakeMEXmodel(IQMmodel([modelName '.txt']))
%% Loading data-sets, to be used in objective function/Plotting

load expData
%% Model
design=matlab.lang.makeValidName({'Ca+ATP';'Ca+noATP';'noCa+ATP';'noCa+noATP';'highCa+ATP';...
    'EPI+ATP';'CL+ATP';'CL+Ca'});
%   Ca,           ATP,         cAMP,   Epi, CL
stimulus=[...
    0.0015         3           0.1     0    0;   %'Ca+ATP'
    0.0015         0           0.1     0    0;   %'Ca+noATP'
    0              3           0.1     0    0;   %'noCa+ATP'
    0              0           0.1     0    0;   %'noCa+noATP'
    0.015          3           0.1     0    0;   %'highCa+ATP'
    0              3           0       5    0;   %'EPI+ATP'
    0              3           0       0    1;   %'CL+ATP'
    0.0015         3           0       0    1];   %'CL+Ca' 

experimentalSetup=table(stimulus(:,1:3), stimulus(:,4:end), 'VariableNames',{'Pipette','Agonist'},'RowNames',design);

if nargin<1 || isempty(modelName)
    modelName='adr_endo'; % Name of model file.
end
if nargin<2
toEstimateOn={'Ca_ATP','Ca_noATP','noCa_ATP','noCa_noATP','highCa_ATP','EPI_ATP','CL_Ca'};
end
scaleExp={'Ca_ATP','Ca_noATP','noCa_ATP','noCa_noATP'};

if nargin<3
toValidateOn={'CL_ATP'};
end

expData(:,~ismember(expData.Properties.VariableNames,[toEstimateOn toValidateOn scaleExp(~ismember(scaleExp,toEstimateOn))]))=[];
estimation=experimentalSetup(toEstimateOn,:);
validation=experimentalSetup(toValidateOn,:);

model=str2func(modelName); % Sets the model as a function.

pNames=IQMparameters(model);
pNames(end-2:end)=[]; %Removing EPI and CL (not free params) and switch-parameter xpipette.
nParams=length(pNames); 

    
dgf=numel(expData{'Mean',toEstimateOn})-1-nParams;
if dgf<0, dgf=numel(expData{'Mean',toEstimateOn})-1; end 
ub=log(1e4)*ones(1,nParams-2); % removes Vpip and Vcell and basal cAMp/ATP/Ca2+. Sets the upper bounds of the parameter values (in log-space). 
lb=-ub; % Sets the lower bounds.

           % Vpip (volume)    vCell
ub=[ub log([60e-6            (214.6e-15)*0.8  ])]; 
lb=[lb log([20e-6            (10.8e-15)*0.8   ])]; 

if contains(modelName,'_hill')
    hillInd=~cellfun(@isempty,regexp(pNames,'^n[0-9]+$')); %Finds all names that begins with n, followed with numbers and then ends.
    ub(hillInd)=log(3);
    lb(hillInd)=log(1);
end

end
```
