## Supplementary material for "A systems biology analysis of adrenergically stimulated adiponectin exocytosis in white adipocytes": All scripts used in this work: RunOptimization.m

```
folderName=datestr(now,'yyyymmdd-HHMMSS');
Setup(folderName)
rng('shuffle')

parfor i=1:2500 %1:n, n = number of repeats
    random=randi([0,32767],1);
    EstimateParameters(random,folderName,1,1)
end
```
