## Supplementary material for "A systems biology analysis of adrenergically stimulated adiponectin exocytosis in white adipocytes": All scripts used in this work: RunOptimizationPred.m

```
folderName=datestr(now,'yyyymmdd-HHMMSS');
Setup(['PPL/' folderName])
rng('shuffle')

parfor i=1:100 %1:n, n = number of repeats
    random=randi([0,32767],1);
    EstimateParametersPred(random,folderName,1,1)
end
```
