## Supplementary material for "A systems biology analysis of adrenergically stimulated adiponectin exocytosis in white adipocytes": All scripts used in this work: runParallel.sh

```
#!/bin/bash
#
# Use 2 nodes, i.e. 2x16 cores, for 1 hour
#SBATCH -N 2 
#SBATCH -t 02:00:00

# Update the project if possible. 
#git pull

# Load the Matlab module
module load MATLAB/R2018b-nsc1 
module add gcc/8.2.0 

MATLAB='matlab -nodesktop -nodisplay -singleCompThread'
# The name of the Matlab script (without .m)
job=EstimateParameters
save_valid=0
useBest=1
now=$(date '+%Y%m%d-%H%M%S')
# Setup things first. 
${MATLAB} -r "Setup('${now}'); exit"

# Run main task. Note the explicit "exit" to exit Matlab.
for i in $(seq 1 256);do
  srun -N1 -n1  ${MATLAB} -r "${job}($RANDOM,'${now}',${useBest},${save_valid},0);exit" > ./Log/${job}_${i}.log &
done
wait # needed to prevent the script from exiting before all Matlab tasks are done

# End of script
```
