## Supplementary material for "A systems biology analysis of adrenergically stimulated adiponectin exocytosis in white adipocytes": All scripts used in this work: SetSameAxes.m

```
function []=SetSameAxes(figure, padding, xmin, xmax, ymin, ymax)
if nargin<1 || isempty(figure)
    figure=gcf;
end
if nargin<2
    padding=[0 0];
end

axs=findobj(figure,'type','axes');
xlim=[inf -inf];
ylim=[inf -inf];
for i=1:length(axs)
    xlim(1)=min(xlim(1), axs(i).XLim(1));
    xlim(2)=max(xlim(2), axs(i).XLim(2));
    
    ylim(1)=min(ylim(1), axs(i).YLim(1));
    ylim(2)=max(ylim(2), axs(i).YLim(2));
end

if nargin == 3 && ~isempty(xmin)
    xlim(1)=xmin;
end
if nargin == 4 && ~isempty(xmax)
    xlim(2)=xmax;
end
if nargin == 5 && ~isempty(ymin)
    ylim(1)=ymin;
end
if nargin == 6 && ~isempty(ymax)
    ylim(2)=ymax;
end
xlim(1)=xlim(1)-padding(1);
xlim(2)=xlim(2)+padding(1);

ylim(1)=ylim(1)-padding(2);
ylim(2)=ylim(2)+padding(2);


for i=1:length(axs)
    axs(i).XLim=xlim;
    axs(i).YLim=ylim;
end
end
```
