## Supplementary material for "A systems biology analysis of adrenergically stimulated adiponectin exocytosis in white adipocytes": All scripts used in this work: Setup.m

```
function []=Setup(folderName)
%% Setup path and toolboxes
if ~exist('IQMsimulate','file')
run('./Support/IQM Tools/installIQMtoolsInitial.m')
end


%% Create necessary folders and compile models
if ~exist('Log','dir')
    mkdir('Log')
end

if nargin<1 || isempty(folderName)
    folder='../Results/Temp/';
else
    folder=['../Results/' folderName '/'];
end
if ~exist(folder,'dir')
    mkdir(folder)
end
    
models=dir('*.txt');

for i =1:length(models)
    IQMmakeMEXmodel(IQMmodel(models(i).name))

end
end
```
