## Supplementary material for "A systems biology analysis of adrenergically stimulated adiponectin exocytosis in white adipocytes": All scripts used in this work: SimulateExperiments.m

```
function [ simulatedExperiments, simulations, scale, maxcAMP] = SimulateExperiments(param, times, ic, model, expData, experiments)

%% Initial setups

param=exp(param);

design=matlab.lang.makeValidName({'Ca+ATP';'Ca+noATP';'noCa+ATP';'noCa+noATP'}); %These are the ones used for scaling. 
%   Ca,           ATP,         cAMP,   Epi, CL
stimulus=[...
    0.0015         3           0.1     0    0;   %'Ca+ATP'
    0.0015         0           0.1     0    0;   %'Ca+noATP'
    0              3           0.1     0    0;   %'noCa+ATP'
    0              0           0.1     0    0];  %'noCa+noATP'
if nargin<6 || isempty(experiments)
    experiments=table(stimulus(:,1:3), stimulus(:,4:end), 'VariableNames',{'Pipette','Agonist'},'RowNames',design);
end

missing=[];
if any(~ismember(design,experiments.Properties.RowNames))
    missing=~ismember(design,experiments.Properties.RowNames);
    experiments(design(missing),:)=table(stimulus(missing,1:3), stimulus(missing,4:end));
end
if nargin==1
    times=expData{'Time',:}; %The last row in expdata is the time
end

scaleExp={'Ca_ATP','Ca_noATP','noCa_ATP','noCa_noATP'};
scaleTime=unique(expData{'Time',scaleExp});
simTimes=unique([0 times scaleTime]);


ic(end-2:end)=[];
ic(ismember(IQMstates(model),'Adiponectin'))=0;% Adiponection release is set to zero.

simulations=[];
simulatedExperiments=table(nan(height(experiments),length(simTimes)),'VariableNames',{'Measures'},'RowNames',experiments.Properties.RowNames);
simulatedExperiments.PeakTime=nan(height(simulatedExperiments),1);
maxcAMP=-1;
cAMPInd=strcmp(IQMstates(model),'cAMP');
%% Simulate experiments

for i=1:height(experiments)
    sim=model(simTimes,[ic experiments{i,'Pipette'}], [param experiments{i,'Agonist'}  1]);
    maxcAMP=max([maxcAMP; sim.statevalues(:,cAMPInd)]);
    simulations=[simulations sim]; % Collect the full simulation structure, but only for the time points requested.
    simulatedExperiments{i,'Measures'}=simulations(end).variablevalues(:,1)';
    simulatedExperiments.PeakTime(i)=sim.time(find(sim.variablevalues(:,1)==max(sim.variablevalues(:,1)),1))/60;
end


%% Do scaling
meanValues=expData{'Mean',scaleExp}';
sims=simulatedExperiments{scaleExp,'Measures'};
sims(:,~ismember(simTimes,scaleTime))=[];
sims=reshape(sims',1,numel(sims))';

sims(isnan(meanValues))=[];
meanValues(isnan(meanValues))=[];

scale=lscov(sims,meanValues);
if scale<0
    scale=1;
end

simulatedExperiments{:,'Measures'}=simulatedExperiments{:,'Measures'}.*scale;

%% Setup final output. 

simulatedExperiments(end+1,:)={simulations(end).time nan};
simulatedExperiments.Properties.RowNames{end}='Time';
tInd=ismember(simTimes,times);
simulatedExperiments.Measures(:,~tInd)=[]; %Removes extra timepoints only used for scaling.
simulatedExperiments(design(missing),:)=[]; % we might have added simulations not asked for because we needed them for scaling
simulations(end-sum(missing)+1:end)=[]; % See above. The number of missing is the amount we need to remove. 


end
```
