## Supplementary material for "A systems biology analysis of adrenergically stimulated adiponectin exocytosis in white adipocytes": All scripts used in this work: SimulateSteadyState.m

```
function [intialconditions,SteadyState_cost] = SimulateSteadyState(param, model, SSsim_step, betaReduction)

param=exp(param);
%% Simulate steadystate
simTimeEnd=300000;
if nargin<3
    simTime=[0 simTimeEnd-100 simTimeEnd];
else
    simTime = unique([0:SSsim_step:simTimeEnd simTimeEnd-100]);
end
SSstartGuess=param;
%                                EPI     CL   xPipette
SSstartGuess=[SSstartGuess      0       0     0         ];

if nargin > 3 && ~isempty(betaReduction)
   states=IQMstates(model);
   ic=IQMinitialconditions(model);
   ic(strcmp(states,'Beta'))=ic(strcmp(states,'Beta'))*(1-betaReduction);
else
    ic=[];
end
SSsim=model(simTime,ic,SSstartGuess); % Steady state simulation.

intialconditions=SSsim.statevalues(end,:);

%% Calculate steady state cost.
SteadyState_cost=0;
SSderv= abs((SSsim.statevalues(end,:)-SSsim.statevalues(end-1,:))./(SSsim.time(end) - SSsim.time(end-1)));
if length(IQMstates(model))== 14
    SSderv(11)=[];
elseif length(IQMstates(model))== 13
    SSderv(10)=[];
end

largeInd=SSderv > 1e-8;
if any(largeInd)
    SteadyState_cost= 200+sum((SSderv(largeInd)*1e-8).^2); 
end


end
```
