## Supplementary material for "A systems biology analysis of adrenergically stimulated adiponectin exocytosis in white adipocytes": All scripts used in this work: installation.txt

Installation of IQM Tools Suite
===============================

When you are reading this document, you already have successfully obtained IQM Tools Suite from somewhere.
Please perform the following steps to install it:

- Move the IQM Tools Suite folder to a location on your computer where you want to store it
- Start MATLAB
- When installing IQM Tools the first time, please read first the "IQMlite/installation.txt" file and handle as instructed,
  then read the "IQMpro/installation.txt" file and handle accordingly
- Change into the folder in which "IQMlite" and "IQMpro" folders are located
- Execute "installIQMtoolsInitial"

Note: The installation of IQM Tools does not save the MATLAB path. This means that everytime you start MATLAB you will need to 
      execute the "installIQMtools" script. This procedure has been chosen on purpose, for compliance and reproducibility reasons.
      The "installIQMtools" does the same as the "installIQMtoolsInitial" script, but does not compile the required libraries.
      It can be used instead of "installIQMtoolsInitial" script after this has been run at least once.
	  
 
