## Supplementary material for "A systems biology analysis of adrenergically stimulated adiponectin exocytosis in white adipocytes": All scripts used in this work: installIQMtools.m

```
% installIQMtools
% This script installs the LITE and PRO version of the IQM Suite of modeling tools.
% Basically it assumes that both packages are present in the folder and that 
% installIQMtoolsInitial has been executed previously to ensure that all libraries
% have been compiled for your system.
%
%       installIQMtools
%
% The function will check for a previous version of IQM Lite installed and
% in this case it will remove this previous version from the path and
% install the desired version.
%
% Additional IQM packages - if present, will be added as well. Each
% additional package is expected to have a name starting with "IQM" and to
% have an install.m file in its root folder.

% Try to install IQM Tools Lite
IQMliteInstalled = 0;
try
    cd IQMlite
    addpath(genpath(pwd));
    cd ..
    IQMliteInstalled = 1;
catch
    disp('Error installing IQM Tools Lite. You need to run the installation script from the folder it is located in.');
end

% Try to install IQM Tools Pro
if IQMliteInstalled,
    try
        cd IQMpro
        addpath(genpath(pwd));
        cd ..
    catch
        disp('Error installing IQM Tools Pro.');
    end
end

% Add additional packages if present
x = dir('IQM*');
x = x([x.isdir]);
x = {x.name};
x = setdiff(x,{'IQMlite'    'IQMpro'});
for k=1:length(x),
    cd(x{k});
    try
        addpath(genpath(pwd));
    catch
        disp(sprintf('Problems installing package "%s".',x{k}));
    end
    cd ..
end

addpath(pwd);
```
