## Supplementary material for "A systems biology analysis of adrenergically stimulated adiponectin exocytosis in white adipocytes": All scripts used in this work: compareFilesEqualIQM.m

```
function [result] = compareFilesEqualIQM(A,B,line)
% Compares if two text files have equal content. 
% A and B are paths to the files to be compared.
% line indicates the number of the line from which the comparison should
% start. As line separator char(10) is assumed.

    Ax = fileread(A);
    Bx = fileread(B);
    
    % Get from line-th line
    if line>1,
        ix = find(Ax==char(10)); Ax = Ax(ix(line-1)+1:end);
        ix = find(Bx==char(10)); Bx = Bx(ix(line-1)+1:end);
    end
    
    % Compare
    result = strcmp(Ax,Bx);
return
```
