## Supplementary material for "A systems biology analysis of adrenergically stimulated adiponectin exocytosis in white adipocytes": All scripts used in this work: mat2datasetIQM.m

```
function [output] = mat2datasetIQM(x,colNames)
% mat2datasetIQM: converts a double matrix to a dataset
% Function required to allow compatibility with pre R2013 versions of MATLAB
%
%   d = mat2datasetIQM(x)
%   d = mat2datasetIQM(x,colNames)
%
% x:        matlab matrix
% colNames: cell-array with columnnames

% <<<COPYRIGHTSTATEMENT - IQM TOOLS LITE>>>

if nargin == 1,
    colNames = {};
    for k=1:size(x,2),
        colNames{k} = sprintf('x%d',k);
    end
end

if length(colNames) ~= size(x,2),
    error('Incorrect number of column names.');
end

output = table();
for k=1:size(x,2),
    output.(colNames{k}) = x(:,k);
end

    
```
