## Supplementary material for "A systems biology analysis of adrenergically stimulated adiponectin exocytosis in white adipocytes": All scripts used in this work: strmatchIQM.m

```
function [I] = strmatchIQM(str,strarray,varargin)
% strmatchIQM: emulates the functionality of strmatch. strmatch will be
% removed from standard matlab soon (currently R2012a).
%
% I = strmatchIQM(STR, STRARRAY) looks through the rows of the character
% array or cell array of strings STRARRAY to find strings that begin
% with the string contained in STR, and returns the matching row indices.
% Any trailing space characters in STR or STRARRAY are ignored when
% matching. strmatch is fastest when STRARRAY is a character array.
% 
% I = strmatchIQM(STR, STRARRAY, 'exact') compares STR with each row of
% STRARRAY, looking for an exact match of the entire strings. Any
% trailing space characters in STR or STRARRAY are ignored when matching.

% <<<COPYRIGHTSTATEMENT - IQM TOOLS LITE>>>

% if verLessThan('matlab', '7.14'),
%     % strmatch exists
%     I = strmatch(str,strarray,varargin{:}); %#ok<*MATCH3>
% else
    % In newer versions "strmatch" might not exist - use a workaround instead

    % Check if strarray is a string array and if yes, convert to cell-array
    if ischar(strarray),
        strarraynew = {};
        for k=1:size(strarray,1),
            strarraynew{k} = strarray(k,:);
        end
        strarray = strarraynew;
    end
    
    if nargin==3,
        if strcmp(varargin{1},'exact'),
            I = find(strcmp(str,strarray));
            I = I(:);
        else
            error('Third input argument to strmatchIQM needs to be "exact" or not specified.');
        end
    else
        I = find(strncmp(str,strarray,length(str)));
        I = I(:);
    end
% end
        
if isempty(I),
    I = [];
end
```
