## Supplementary material for "A systems biology analysis of adrenergically stimulated adiponectin exocytosis in white adipocytes": All scripts used in this work: getComplianceModeIQM.m

```
function [ COMPLIANCE_OUTPUT_MODE ] = getComplianceModeIQM()
% This function returns value of the setting COMPLIANCE_OUTPUT_MODE in
% SETUP_PATHS_TOOLS_IQMLITE.   
%
% [SYNTAX]
% [ COMPLIANCE_OUTPUT_MODE ] = getComplianceModeIQM()
%
% [INPUT]
%
% [OUTPUT]
% COMPLIANCE_OUTPUT_MODE:     Value of COMPLIANCE_OUTPUT_MODE

% <<<COPYRIGHTSTATEMENT - IQM TOOLS PRO>>>

SETUP_PATHS_TOOLS_IQMLITE
```
