## Supplementary material for "A systems biology analysis of adrenergically stimulated adiponectin exocytosis in white adipocytes": All scripts used in this work: getFunctionCallInformationIQM.m

```
function [ results ] = getFunctionCallInformationIQM(FlagText,FlagLineNumbers,FlagCompleteNames)
% This function generates and returns information about the username, the
% current date and time and the stack of functions that have been called
% until this function. This function here is not included in the stack.
%
% For compliance reasons it is important to have time stamps etc. for all
% output - especially figures and tables.
% 
% This function allows to add this automatically when printing figures to
% file via IQMprintFigure and writing text to file via IQMwriteText2File.
%
% [SYNTAX]
% [ results ] = getFunctionCallInformationIQM()
% [ results ] = getFunctionCallInformationIQM(FlagText)
% [ results ] = getFunctionCallInformationIQM(FlagText,FlagLineNumbers)
% [ results ] = getFunctionCallInformationIQM(FlagText,FlagLineNumbers,FlagCompleteNames)
%
% [INPUT]
% FlagText:             =0: Return MATLAB structure with the information 
%                       =1: Return a cell-array with the information - in table format (default)
% FlagLineNumbers:      =0: Do not include line numbers (default)
%                       =1: Include line numbers
% FlagCompleteNames:    =0: Show filenames only (default)
%                       =1: Show complete filenames with absolute path
%
% [OUTPUT]
% results:  MATLAB structure with the following information:
%   results.username:           String with username who ran the function
%   results.data:               String with current date and time
%   results.callingSequence:    String with function and script names that
%                               have been called prior to this function.
%                               this function here is not included. Line
%                               number is included as well.

% <<<COPYRIGHTSTATEMENT - IQM TOOLS LITE>>>

% If COMPLIANCE_OUTPUT_MODE is on then Current_ROOT_file_IQMtools_compliance
% needs to be set to the name of the script in which the user executes a
% function that will lead (through several subscripts and functions) to an
% output by IQMprintFigure or IQMwriteText2File.
% The check requires global variables that are not accessible in a parfor
% loop. Thus we need to check if in parallel execution or not. And only if
% not then do the compliance check
if ~isInsideParForLoopIQM(),
    global Current_ROOT_file_IQMtools_compliance
    if getComplianceModeIQM(),
        if isempty(Current_ROOT_file_IQMtools_compliance),
            error(sprintf('Compliance mode is on. Please use the function "IQMinitializeCompliance" at the beginning of each main working script.\nDo not use "clear all" within a script - or rerun the "IQMinitializeCompliance" function after each call to "clear all".'));
        end
    end
else
    Current_ROOT_file_IQMtools_compliance = 'undefined due to parfor';
end

% Handle variable input arguments
if nargin<1,
    FlagText                = 1;
end
if nargin<2,
    FlagLineNumbers         = 0;
end
if nargin<3,
    FlagCompleteNames       = 0;
end

% Initialize results variable
results                     = [];

% Get basic information about user and date and time of function call
results.username            = usernameIQM();
results.date                = datestr(now,'yyyy-mmm-DD HH:MM');

% Get information about call sequence
x                           = dbstack('-completenames');

callingSequence             = {};
for k=length(x):-1:2,
    if FlagLineNumbers,
        callingSequence{end+1} = sprintf('%s (Line: %d)\n',x(k).file,x(k).line);
    else
        callingSequence{end+1} = sprintf('%s\n',x(k).file);
    end
end
% Adjust folder separators
results.callingSequence     = strrep(strtrim(callingSequence),'\','/');

% Check if Current_ROOT_file_IQMtools_compliance contents appear in 
% results.callingSequence. If not then add it. Only check for filename ...
% not necessarily with path - but with .m at the end and '/' before.
if getComplianceModeIQM(),
    [xp,xf,xe] = fileparts(Current_ROOT_file_IQMtools_compliance);
    checkName  = ['/' xf '.m'];
    ROOT_PRESENT = 0;
    for k=1:length(results.callingSequence),
        ix = strfind(results.callingSequence{k},checkName);
        if ~isempty(ix),
            ROOT_PRESENT = 1;
        end
    end
    if ~ROOT_PRESENT,
        % Root file defined by Current_ROOT_file_IQMtools_compliance not
        % present in calling sequence. This is due to manual or execution
        % of parts of the script only. Need to add the root file name
        if ~isInsideParForLoopIQM(),
            results.callingSequence = [{['Main analysis file: "' xf '.m"']} results.callingSequence];
        else
            results.callingSequence = [{'Main analysis file undefined due to execution in parfor loop'} results.callingSequence];
        end
    end
end

% Convert to table if desired
if FlagText,
    textTable               = {'<TR>' 'Username'            results.username                    };
    textTable(end+1,:)      = {'<TR>' 'Date of creation'    results.date                        };
    for k=1:length(results.callingSequence),
        if k==1,
                textTable(end+1,:)  = {'<TR>' 'Calling sequence'    results.callingSequence{k} };
        else
                textTable(end+1,:)  = {'<TR>' ' '                   results.callingSequence{k} };
        end
    end
    results = textTable;
end
```
