## Supplementary material for "A systems biology analysis of adrenergically stimulated adiponectin exocytosis in white adipocytes": All scripts used in this work: IQMconvertCellTable2ReportTable.m

```
function [text] = IQMconvertCellTable2ReportTable(cellTable,format,filename)
% This function allows to generate tables for display in the command window
% and also allows to write them out to a text file - both humanly readable
% and parse-able to subsequently reformat them for inclusion into a report.
%
% The input argument "cellTable" is a MATLAB cell-matrix that is allowed to
% use a certain syntax, defined below:
% 
% The first column of the cell-matrix can be used as a "control-column"
% and the first element in each row can contain any of the following <xy> tags:
%   <TT>: Table title (only first element of cellTable written out across whole table width)
%   <TH>: Table header (each cell of cellTable considered a table header element)
%   <TR>: Table row (each cell of cellTable considered a table data element)
%   <TF>: Table footer (only first element of cellTable written out across whole table width)
%   <HR>: Table separator (no content expected. Just introducing a break (e.g., horizontal line))
%   <TI>: Table information (only first element written out across the whole line. No separators top and bottom)
% 
% This control column can be missing and in this case the cellTable is just
% written out, considering each cell a cell in the table <TR>.
%
% Several tables can be present in the same cellTable. Each starts with
% <TT> or <TH>.
%
% Additional formatting in table cells:
% -------------------------------------
% * Line breaks ('\n') can be used in cell elements. For text output they
%   are replace by spaces (' '). For report outputs they are replace by
%   HTML tag '<br>'.
%
% [SYNTAX]
% [text] = IQMconvertCellTable2ReportTable(cellTable)
% [text] = IQMconvertCellTable2ReportTable(cellTable,format)
% [text] = IQMconvertCellTable2ReportTable(cellTable,format,filename)
%
% [INPUT]
% cellTable:    Cell-matrix with string and/or numeric entries and
%               potentially the control column for formatting 
% format:   	Control of output format
%           "text":     Will convert to text only without control tags.
%                       This can be used if information shoudl just be
%                       displayed to the user in the command window or exported
%                       to a file that will not be part of a report.
%           "report":   This will convert the table to text as well but
%                       additionally add some formatting that is needed to
%                       parse this table from a text file for later
%                       formatting into a report. (default).
% filename:     If a filename is provided then the table is exported to
%               filename.txt. (default: '');
%
% [OUTPUT]
% text:         Formatted string table output
% The text might also be written to a file.              

% <<<COPYRIGHTSTATEMENT - IQM TOOLS LITE>>>

% Handle variable input arguments
if nargin<2,
    format = 'report';
end
if nargin<3.
    filename = '';
end
format = lower(format);

% Check things
if isempty(strmatchIQM(format,{'report','text'},'exact')),
    error('Input argument "format" wrongly defined.');
end

% Convert all elements in cellTable to char 
for k=1:size(cellTable,1),
    for k2=1:size(cellTable,2),
        if isempty(cellTable{k,k2}),
            cellTable{k,k2} = '';
        end
        if isnumeric(cellTable{k,k2}),
            cellTable{k,k2} = num2str(cellTable{k,k2});
        end
    end
end

% Check line breaks in cellTable and replace by '<br>'
for k=1:size(cellTable,1),
    for k2=1:size(cellTable,2),
        cellTable{k,k2} = regexprep(cellTable{k,k2},'\n','<br>');
    end
end


% Check if control column present
CONTROL = 1;
if isnumeric(cellTable{1,1}),
    CONTROL = 0;
end
if isempty(intersect({'<TT>','<TH>','<TR>','<TF>','<HR>'},cellTable(:,1))),
    CONTROL = 0;
end

% If no control column then treat all rows as <TR>
if CONTROL,
    control_column = upper(cellTable(:,1));
    cellTable2     = cellTable(:,2:end);
else
    control_column = cell(size(cellTable,1),1);
    control_column(1:end) = {'<TR>'};
    cellTable2     = cellTable;
end
    
% Find maxcellTable lengths of strings in each <TR> and <TH> column
length_column_string = -Inf(1,size(cellTable2,2));
for k=1:size(cellTable2,2),
    for k2=1:size(cellTable2,1),
        if strcmp(control_column{k2},'<TR>') || strcmp(control_column{k2},'<TH>'),
            if length(cellTable2{k2,k}) > length_column_string(k),
                length_column_string(k) = length(cellTable2{k2,k});
            end
        end
    end
end

% Update table entries by postpadding with spaces to match
% length_column_string - do not postpad last column. 
for k=1:size(cellTable2,1),
    for k2=1:size(cellTable2,2),
        cellTable2{k,k2} = postFillCharIQM(cellTable2{k,k2},length_column_string(k2),' ');
    end
end

% Add separator in all but last column for <TH> and <TR> cells.
% Spaces in case of "text" and ' | in case of "report".
for k=1:size(cellTable2,1),
    if strcmp(control_column{k},'<TR>') || strcmp(control_column{k},'<TH>'),
        for k2=1:size(cellTable2,2)-1,
            if strcmp(format,'report'), separator = ' | '; else separator = ' | '; end
            cellTable2{k,k2} = [cellTable2{k,k2} separator];
        end
    end
end

% Get complete table width
WIDTH_TABLE = length(sprintf('%s',cellTable2{1,:}));
if strcmp(control_column{1,1},'<TT>'),
    WIDTH_TABLE = WIDTH_TABLE+3;
end

% If report then add control chars to first column in tabe
if strcmp(format,'report'),
    for k=1:size(cellTable2,1),
        cellTable2{k,1} = [control_column{k} '   ' cellTable2{k,1}];
    end
    addSpaces = '       ';
    addSpacesHR = '   ';
else
    addSpaces = '';
    addSpacesHR = '';
end    

% Convert to formatted text
text = '';
lengthTHRrow = 10;
for k=1:size(cellTable2,1),
    textadd = '';
    % Handle <TT>
    if strcmp(control_column{k},'<TT>'),
        if k>1,
            textadd = sprintf('\n\n%s\n%s%s\n\n',cellTable2{k,1},addSpaces,postFillCharIQM('',WIDTH_TABLE,'='));      
        else
            textadd = sprintf('%s\n%s%s\n\n',cellTable2{k,1},addSpaces,postFillCharIQM('',WIDTH_TABLE,'='));
        end
    end
    % Handle <TH>
    if strcmp(control_column{k},'<TH>'),
        textadd = '';
        for k2=1:size(cellTable2,2),
            textadd = sprintf('%s%s',textadd,cellTable2{k,k2});
        end
        lengthTHRrow = length(textadd)-length(addSpaces);
        textadd = sprintf('%s\n%s%s\n',textadd,addSpaces,postFillCharIQM('',lengthTHRrow,'-'));
    end
    % Handle <TR>
    if strcmp(control_column{k},'<TR>'),
        textadd = '';
        for k2=1:size(cellTable2,2),
            textadd = sprintf('%s%s',textadd,cellTable2{k,k2});
        end
        lengthTHRrow = length(textadd)-length(addSpaces);
        textadd = sprintf('%s\n',textadd);
    end
    % Handle <TF>
    if strcmp(control_column{k},'<TF>'),
        textadd = sprintf('%s%s\n%s\n',addSpaces,postFillCharIQM('',lengthTHRrow,'-'),cellTable2{k,1});
    end
    % Handle <HR>
    if strcmp(control_column{k},'<HR>'),
        textadd = sprintf('%s%s%s\n',strtrim(cellTable2{k,1}),addSpacesHR,postFillCharIQM('',lengthTHRrow,'-'));
    end
    % Handle <TI>
    if strcmp(control_column{k},'<TI>'),
        textadd = sprintf('%s\n',cellTable2{k,1});
    end
    
    % Combine text
    text = [text textadd];
end

% Remove tags if "format" is "text"
if strcmp(format,'text'),
    text = strrep(text,'<br>','    ');
    text = strrep(text,'<BR>','    ');
end

% Export results to file if filename defined
IQMwriteText2File(text,[strrep(filename,'.txt','') '.txt']);
```
