## Supplementary material for "A systems biology analysis of adrenergically stimulated adiponectin exocytosis in white adipocytes": All scripts used in this work: IQMinitializeCompliance.m

```
function [] = IQMinitializeCompliance(ScriptName)
% This function initializes the compliance check. Basically, it sets a
% global variable (Current_ROOT_file_IQMtools_compliance) to a value of
% "ScriptName". Additionally, this function sets the sed for random
% generators in MATLAB, to ensure that rerunning the same script leads to
% the same results.
%
% When the compliance mode is checked each analysis script needs to make a
% call to "IQMinitializeCompliance(ScriptName)" at the beginning of the
% script. The input argument needs to be the filename of the analysis
% script. This can include the absolute path or just be the filename.
%
% [SYNTAX]
% [] = IQMinitializeCompliance(ScriptName)
%
% [INPUT]
% ScriptName:   Filename (with or without path) to the analysis script.
%
% [OUTPUT]
% None

% <<<COPYRIGHTSTATEMENT - IQM TOOLS PRO>>>

% Clear global variable 
clear global Current_ROOT_file_IQMtools_compliance

% Initialize the global variable
global Current_ROOT_file_IQMtools_compliance

% Set its value
Current_ROOT_file_IQMtools_compliance = ScriptName;

% Remove extension (.m) if present 
Current_ROOT_file_IQMtools_compliance = strrep(Current_ROOT_file_IQMtools_compliance,'.m','');

% Check if a function above IQMinitializeCompliance is stored in dbstack
% ... if yes then assume this is the "root" script.
x           = dbstack();
rootFile    = strrep(x(end).file,'.m','');

if ~strcmp(rootFile,'IQMinitializeCompliance'),
    % There is at least one parent function to IQMinitializeCompliance
    % Check if Current_ROOT_file_IQMtools_compliance is the parent function of IQMinitializeCompliance 
    
    names       = {x.name};
    ix          = strmatchIQM('IQMinitializeCompliance',names,'exact');
    checkFile   = names{ix+1};
    
    [p,f,e] = fileparts(Current_ROOT_file_IQMtools_compliance);
    if ~strcmp(f,checkFile),
        error('Input argument to "IQMinitializeCompliance" does not match the filename of this script in which this command is run.');
    end
end

% Set seed
setseedIQM(123456)
```
