## Supplementary material for "A systems biology analysis of adrenergically stimulated adiponectin exocytosis in white adipocytes": All scripts used in this work: IQMoutputTable.m

```
function [text] = IQMoutputTable(xtable,xtitle,xfooter,filename)
% This function allows to convert a MATLAB table to a report table
% formatted to be compatible with IQReport.
%
% [SYNTAX]
% [text] = IQMoutputTable(xtable,xtitle,xfooter,filename)
%
% [INPUT]
% xtable:       MATLAB table
% xtitle:       String with the table caption (default: '')
% xfooter:      String with the table footer (default: '')
% filename:     If a filename is provided then the table is exported to
%               filename.txt. (default: '');
%
% [OUTPUT]
% The text is written to a file.              

% <<<COPYRIGHTSTATEMENT - IQM TOOLS LITE>>>

% Handle variable input arguments
if nargin<4,
    error('Please provide all input arguments.');
end

% Check table
if ~istable(xtable),
    error('First input argument is not a MATLAB table.');
end

% Get info
Ncols               = size(xtable,2)+1;
headerNames         = xtable.Properties.VariableNames;

% Generate table-content as cell-array
tableCells          = table2cell(xtable);

% Add <TR> info
tableRows           = cell(size(tableCells,1),1); 
tableRows(1:end)    = {'<TR>'};
tableCells          = [tableRows tableCells];

% Add <TH> info
tableCells          = [ [{'<TH>'} headerNames]; tableCells];

% Add <TT> info
tableTitle          = cell(1,Ncols);
tableTitle{1}       = '<TT>';
tableTitle{2}       = xtitle;
tableCells          = [tableTitle; tableCells];

% Add <TF> info
if ~isempty(xfooter),
    tableFooter         = cell(1,Ncols);
    tableFooter{1}      = '<TF>';
    tableFooter{2}      = xfooter;
    tableCells          = [tableCells; tableFooter];
end

% Convert to report table
IQMconvertCellTable2ReportTable(tableCells,'report',filename);
```
