## Supplementary material for "A systems biology analysis of adrenergically stimulated adiponectin exocytosis in white adipocytes": All scripts used in this work: diffIQM.m

```
function [status, res] = diffIQM(A,B)
% Comparison of two files using the diff command. 
% Wrapper to handle Unix and Windows

if isunix(),
    eval(sprintf('[status, res] = system(''diff %s %s'');', A,B));
else
    diffLocation = which('diff.exe');
    eval(sprintf('[status, res] = system(''"%s" %s %s'');', diffLocation,A,B))
end
```
