## Supplementary material for "A systems biology analysis of adrenergically stimulated adiponectin exocytosis in white adipocytes": All scripts used in this work: getDefaultIntegratorOptionsIQM.m

```
function [options_M,options_C] = getDefaultIntegratorOptionsIQM()
% getDefaultIntegratorOptionsIQM returns structures with default integrator 
% setting options for both MATLAB and CVODE simulation.

% MATLAB
options_M                     = odeset();
options_M.method              = 'ode23s';  
options_M.AbsTol              = 1e-6;
options_M.RelTol              = 1e-6;

% CVODE
options_C.method              = 'stiff';  % 'stiff' or 'nonstiff'
options_C.abstol              = 1e-6;
options_C.reltol              = 1e-6;
options_C.maxnumsteps         = 100000;   % max number of steps between two output points
```
