## Supplementary material for "A systems biology analysis of adrenergically stimulated adiponectin exocytosis in white adipocytes": All scripts used in this work: getSAStoolInfoIQM.m

```
function [PATH_SAS] = getSAStoolInfoIQM()
% Function loads and returns the content of SETUP_PATHS_TOOLS_IQMLITE already
% correct for the system. 

% Run the SETUP_PATHS_TOOLS_IQMLITE script
SETUP_PATHS_TOOLS_IQMLITE

if isunix,
    PATH_SAS        = PATH_SYSTEM_SAS_UNIX;
else
    PATH_SAS        = PATH_SYSTEM_SAS_WINDOWS;
end
```
