## Supplementary material for "A systems biology analysis of adrenergically stimulated adiponectin exocytosis in white adipocytes": All scripts used in this work: NovakTyson.m

```
% This script simply loads the novaktysonTestModelIQM.txt model that is included in this folder.
% The purpose of this is only remote support to allow people to easily generate a model.
%
% [SYNTAX]
% NovakTyson
%
% [INPUT]
% None
%
% [OUTPUT]
% None
%
% In the MATLAB workspace the model is made available as variable "iqm".

% <<<COPYRIGHTSTATEMENT - IQM TOOLS LITE>>>

iqm = IQMmodel('novaktysonTestModelIQM.txt')

disp('The Novak Tyson model has been loaded and is available as "iqm" in the MATLAB workspace.');
```
