## Supplementary material for "A systems biology analysis of adrenergically stimulated adiponectin exocytosis in white adipocytes": All scripts used in this work: isSymbolicpresentIQM.m

```
function [flag] = isSymbolicpresentIQM()
% isSymbolicpresentIQM: checks if the symbolic toolbox is present
%
% USAGE:
% ======
% [flag] = isSymbolicpresentIQM()
%
% Output Arguments:
% =================
% flag: 1 if present, 0 if not present

% <<<COPYRIGHTSTATEMENT - IQM TOOLS LITE>>>


flag = ~isempty(ver('symbolic'));
```
