## Supplementary material for "A systems biology analysis of adrenergically stimulated adiponectin exocytosis in white adipocytes": All scripts used in this work: lookforIQML.m

```
function [] = lookforIQML(text)
% lookforIQM: searches all m-files in the IQM Lite folder tree for the 
% text given as argument. It displays the filenames in which the text
% appears and additionally opens the files in the editor.

old = pwd;
cd([fileparts(which('installIQMlite')) '/..']);
try, recurseFolder('IQMlite',text); catch end
cd(old);
return

function recurseFolder(folder,text)
% change folder
cd(folder);
% check all m files in folder for the text
mfiles = dir('*.m');
for k = 1:length(mfiles),
    % read file
    content = fileread(strcat(mfiles(k).name));
    if strfind(content,text),
        disp(mfiles(k).name);
        edit(mfiles(k).name);
    else
%        edit(mfiles(k).name);
    end
end
% recurse in all subfolders
allfiles = dir;
for k = 1:length(allfiles),
    if ~strcmp(allfiles(k).name,'..') && ~strcmp(allfiles(k).name,'.') && allfiles(k).isdir == 1,
        recurseFolder(allfiles(k).name,text);
    end
end
% up we go
cd ..
return
```
