## Supplementary material for "A systems biology analysis of adrenergically stimulated adiponectin exocytosis in white adipocytes": All scripts used in this work: makePosSemiDefIQM.m

```
function [Xout] = makePosSemiDefIQM(X)
% Enforce matrix X to become a positive semidefinite matrix Xout.
% This is accomplished by setting the negative eigenvalues to 0.
% This function is mainly used in ensuring positive-semidefiniteness of
% covariance matrices. It turned out not to be enough to set the values to
% 0 so a slightly different approach has been chosen, documented below.

% <<<COPYRIGHTSTATEMENT - IQM TOOLS LITE>>>

THRES = 1e-10;

Xout = X;
try
    warning off
    % Make symmetric
    X                                           = 0.5*(X+X');
    % Eigenvalue decomposition
    [V,D]                                       = eig(X);
    xxx                                         = diag(D);
    % Set all eigenvalues below THRES to 0
    xxx(real(xxx)<THRES)                        = 0;
    % Set all elements of V below THRES (absolute value since it might be complex) to 0
    V(abs(V)<THRES)                             = 0;
    % Combine again to matrix and force it to be real
    D                                           = diag(xxx);
    Xout                                        = real(V*D*inv(V));
        
    warning on
catch
    disp('The covariance matrix seems to have an issue.');
end
```
