## Supplementary material for "A systems biology analysis of adrenergically stimulated adiponectin exocytosis in white adipocytes": All scripts used in this work: IQMconvert2pdf.m

```
function [] = IQMconvert2pdf(filename)
% Function converting a PS file to a PDF file.
% Now just a wrapper for ps2pdfIQM to allow the definition of just one
% filename.
%
% [SYNTAX]
% [] = IQMconvert2pdf(filename)
%
% [INPUT]
% filename:     filename to be used
% 
% [OUTPUT]

% <<<COPYRIGHTSTATEMENT - IQM TOOLS LITE>>>

if isempty(filename),
    return
end

% Remove extension from filename
filename = strrep(filename,'.ps','');
filename = strrep(filename,'.pdf','');
filename = strrep(filename,'.PS','');
filename = strrep(filename,'.PDF','');

if exist([filename '.ps']),
    ps2pdfIQM([filename '.ps'],[filename '.pdf']);
    
    % Generate the log information for PDF file creation (if COMPLIANCE_OUTPUT_MODE on)
    % ---------------------------------------------------------------------------------
    
    if getComplianceModeIQM(),
        % Get function call information as table
        results = getFunctionCallInformationIQM(1,1);
        
        % Add table title with filename
        addresults = cell(2,size(results,2));
        addresults{1,1} = '<TT>';
        addresults{1,2} = 'PDF file generation log';
        addresults{2,1} = '<TR>';
        addresults{2,2} = 'File (relative to calling function)';
        addresults{2,3} = [filename '.pdf'];
        results = [addresults; results];
        
        % Generate log file
        logfilename = [filename '.pdf.log'];
        logtext     = IQMconvertCellTable2ReportTable(results,'report');
        
        % Write out the logfile
        fid = fopen(logfilename,'w');
        fprintf(fid,'%s',logtext);
        fclose(fid);
        
        % Delete previous .ps.log file
        warning off
        delete([filename '.ps.log']);
        warning on
    end
end
```
