## Supplementary material for "A systems biology analysis of adrenergically stimulated adiponectin exocytosis in white adipocytes": All scripts used in this work: IQMgetcolors.m

```
function [colors,lines,dots,bwcolors] = IQMgetcolors()
% Help function to generate colors for plotting and for black and white
% plotting also line types.
%
% [SYNTAX]
% [colors,lines,dots,bwcolors] = IQMgetcolors()
%
% [INPUT]
% NONE
%
% [OUTPUT]
% colors: Nx3 matrix with colors to be used (see below in detail)
% lines:  cell-array with 48 unique line-styles
% dots:   cell-array with 12 unique dot-styles

% <<<COPYRIGHTSTATEMENT - IQM TOOLS LITE>>>


colors = [
    0         0.4470    0.7410
    0.8500    0.3250    0.0980
    0.9290    0.6940    0.1250
    0.4940    0.1840    0.5560
    0.4660    0.6740    0.1880
    0.3010    0.7450    0.9330
    0.6350    0.0780    0.1840
         0    0.2980    0.4940
    0.5667    0.2167    0.0653
    0.6193    0.4627    0.0833
    0.3293    0.1227    0.3707
    0.3107    0.4493    0.1253
    0.2007    0.4967    0.6220
    0.4233    0.0520    0.1227
    0.25 0.25 0.25
    0.6 0.2 0
    0.75 0.75 0
    0 0.75 0.75
    0 0.5 0
    0 0 1
    0.75 0 0.75
    1 0 0
    0.04 0.14 0.42
    0.31 0.31 0.31
    0.5 0.5 0.5
    0 0 0
    1 0.69 0.39
    0 1 0
    0 1 1
    1 0 1
    0.7 0.78 1
    1 1 0
    0.68 0.92 1
    0.85 0.7 1
    1 0.6 0.78
    ];


% colors = [0.87 0.49 0
%            0.17 0.51 0.34
% 		   0.08 0.16 0.55
% 		   0.48 0.06 0.89
% 		   0.85 0.16 0
% 		   0.68 0.47 0
% 		   0.04 0.52 0.78
% 		   0.25 0.25 0.25 
% 		   0.6 0.2 0
% 		   0.75 0.75 0
% 		   0 0.75 0.75
% 		   0 0.5 0
% 		   0 0 1
% 		   0.75 0 0.75
% 		   1 0 0
% 		   0.04 0.14 0.42
% 		   0.31 0.31 0.31
% 		   0.5 0.5 0.5
% 		   0 0 0
% 		   1 0.69 0.39
% 		   0 1 0
% 		   0 1 1
% 		   1 0 1
% 		   0.7 0.78 1
% 		   1 1 0
% 		   0.68 0.92 1
% 		   0.85 0.7 1
% 		   1 0.6 0.78];
      
lines = {'-' '--' ':' '-.' 'o-','x-','+-','*-','s-','d-','v-','^-','<-','>-','p-','h-',       'o--','x--','+--','*--','s--','d--','v--','^--','<--','>--','p--','h--',     'o-.','x-.','+-.','*-.','s-.','d-.','v-.','^-.','<-.','>-.','p-.','h-.',   'o:','x:','+:','*:','s:','d:','v:','^:','<:','>:','p:','h:'   };
      
dots = {'o','x','+','*','s','d','v','^','<','>','p','h'};      

bwcolors = [0 0 0; 0.33 0.33 0.33; 0.66 0.66 0.66];
      
```
