## Supplementary material for "A systems biology analysis of adrenergically stimulated adiponectin exocytosis in white adipocytes": All scripts used in this work: IQMprintFigure.m

```
function [] = IQMprintFigure(hfig, filename, format)
% Print figure "hfig" to file "filename". Supported formats: png, ps, jpg,
% and pdf. "ps" is default. Several figures can be appended in the same
% file by using format='ps'. Just run the function repeatedly with same
% filename. If format='png' or 'jpg', figures are not appended, but
% overwritten. 
%
% By default this function generates a .log file with the same name as the
% file at the same location with information about original text file name,
% username, date, and scripts and functions that where run to generate the
% information. The extension for this log file is '.log'
%
% Tips when using PS files:
%   1) If you want a PDF you need to generate it afterwards using the function: 
%             IQMconvert2pdf.
%   2) Dont use the PS format if you have transparency in your plots!
%   3) If you want to start a new file then remove the file first (by default 
%      plots are appended in PS mode). Files can be removed by: 
%             IQMstartNewPrintFigure
%
% [SYNTAX]
% [] = IQMprintFigure(hfig, filename)
% [] = IQMprintFigure(hfig, filename, format)
%
% [INPUT]
% hfig:         handle of figure to be printed
% filename:     filename to be used 
% format:       'ps', 'png', 'jpg' (default: 'ps')
% 
% [OUTPUT]
 
% <<<COPYRIGHTSTATEMENT - IQM TOOLS LITE>>>

% Handle variable input arguments
if nargin<2,
    filename = '';
end
if nargin<3,
    format = 'ps';
end

% Check filename
if isempty(filename),
    return
end

% Change format to lower
format = lower(format);

% Handle format settings
if strcmp(format,'ps'),
    imgformat   = 'psc2';
    append      = 1;
elseif strcmp(format,'png'),
    imgformat   = 'png';
    append      = 0;
elseif strcmp(format,'jpg'),
    imgformat   = 'jpeg95';
    append      = 0;
elseif strcmp(format,'pdf'),
    imgformat   = 'pdf';
    append      = 0;
else
    error('Currently unsupported image format');
end
        
% Setting figure properties for printing
if strcmp(imgformat,'png'),
    set(hfig,'PaperOrientation','portrait');
else
    set(hfig,'PaperOrientation','landscape');
end    
set(hfig,'PaperType','A4');
set(hfig,'PaperPositionMode', 'manual');
set(hfig,'PaperUnits', 'centimeters');
set(hfig,'PaperPosition', [0 0 29.7 21]);

% Determine path file and extension name
[path,file,ext] = fileparts(filename);
if isempty(path),
    path = '.';
end

% Create folder if not existing
warning off
mkdir(path);
warning on

% Change into path
oldPath = pwd;
cd(path);

% Print figure
if append,
    print(hfig,['-d',imgformat],file,'-append');
else
    print(hfig,['-d',imgformat],file);
end

% Generate the log information (if COMPLIANCE_OUTPUT_MODE on)
% -----------------------------------------------------------

if getComplianceModeIQM(),
    % Get function call information as table
    results = getFunctionCallInformationIQM(1,1);
    
    % Add table title with filename
    addresults = cell(2,size(results,2));
    addresults{1,1} = '<TT>';
    addresults{1,2} = 'Figure file generation log';
    addresults{2,1} = '<TR>';
    addresults{2,2} = 'File (relative to calling function)';
    addresults{2,3} = [path '/' file '.' format];
    results = [addresults; results];
    
    % Generate log file
    logfilename = [file '.' format '.log'];
    logtext     = IQMconvertCellTable2ReportTable(results,'report');
    
    % Write out the logfile
    fid = fopen(logfilename,'w');
    fprintf(fid,'%s',logtext);
    fclose(fid);
end

% Return to previous path
cd(oldPath);
```
