## Supplementary material for "A systems biology analysis of adrenergically stimulated adiponectin exocytosis in white adipocytes": All scripts used in this work: IQMstartNewPrintFigure.m

```
function [] = IQMstartNewPrintFigure(filename)
% Function starts a new file in which figures are to be printed.
% Basically this means, that it is checked if the file exists and if yes it is deleted.
% The function "IQMprintFigure" can then be used to print a figure into this file.
% The function "IQMconvert2pdf" can then be used to convert from PS to PDF (only unix).
% On Unix check is done for .PDF, on windows for .PS
%
% [SYNTAX]
% [] = IQMstartNewPrintFigure(filename)
%
% [INPUT]
% filename:     filename to be used
% 
% [OUTPUT]
 
% <<<COPYRIGHTSTATEMENT - IQM TOOLS LITE>>>

if isempty(filename),
    return
end

%%%%%%%%%%%%%%%%%%%%%%%%%%%%%%
% Create folder if it is not existing
%%%%%%%%%%%%%%%%%%%%%%%%%%%%%%
folder = fileparts(filename);
if ~isempty(folder),
    warning off
    mkdir(folder);
    warning on
end

%%%%%%%%%%%%%%%%%%%%%%%%%%%%%%
% If filename exists then delete it
%%%%%%%%%%%%%%%%%%%%%%%%%%%%%%
if ~isempty(filename),
    [path,file,ext] = fileparts(filename);
    filename_pdf = fullfile(path,[file '.pdf']);
    filename_ps  = fullfile(path,[file '.ps']);
    filename_pdf_log = fullfile(path,[file '.pdf.log']);
    filename_ps_log  = fullfile(path,[file '.ps.log']);

    warning off;
    try
        delete(filename_pdf);
    end
    
    try
        delete(filename_ps);
    end
    
    try
        delete(filename_pdf_log);
    end
    
    try
        delete(filename_ps_log);
    end
    
    warning on;
end
```
