## Supplementary material for "A systems biology analysis of adrenergically stimulated adiponectin exocytosis in white adipocytes": All scripts used in this work: IQMwriteText2File.m

```
function [] = IQMwriteText2File(text,filename)
% This function allows to write a given string (text) to a file (filename)
% Filename can include a path and all folders will be created if not yet
% existing.
% 
% By default this function generates a .log file with the same name as the
% file at the same location with information about original text file name,
% username, date, and scripts and functions that where run to generate the
% information. The extension for this log file is '.log'.
%
% [SYNTAX]
% [] = IQMwriteText2File(text,filename)
%
% [INPUT]
% text:     String to write to file
% filename: Filename (with path) to where to write the text
%
% [OUTPUT]
% File "filename" with given "text"

% <<<COPYRIGHTSTATEMENT - IQM TOOLS LITE>>>

% Check filename
filename = strrep(filename,'\','/');
[p,f,e] = fileparts(filename);
if isempty(p),
    p = '.';
end

% Only write out if filename defined
if isempty(f),
    return
end

% Create folder if not existing
warning off
mkdir(p);
warning on
    
% Write out the file
fid = fopen(filename,'w');
fprintf(fid,'%s',text);
fclose(fid);

% Generate the log information (if COMPLIANCE_OUTPUT_MODE on)
% -----------------------------------------------------------

if getComplianceModeIQM(),
    % Get function call information as table
    results = getFunctionCallInformationIQM(1,1);
    
    % Add table title with filename
    addresults = cell(2,size(results,2));
    addresults{1,1} = '<TT>';
    addresults{1,2} = 'Text file generation log';
    addresults{2,1} = '<TR>';
    addresults{2,2} = 'File (relative to calling function)';
    addresults{2,3} = [p '/' f e];
    results = [addresults; results];
    
    % Generate log file
    logfilename = [p '/' f e '.log'];
    logtext     = IQMconvertCellTable2ReportTable(results,'report');
    
    % Write out the logfile
    fid = fopen(logfilename,'w');
    fprintf(fid,'%s',logtext);
    fclose(fid);
end
```
