## Supplementary material for "A systems biology analysis of adrenergically stimulated adiponectin exocytosis in white adipocytes": All scripts used in this work: isInsideParForLoopIQM.m

```
function [output] = isInsideParForLoopIQM()
% This function checks if it is executed within a parfor loop.
%
% [SYNTAX]
% [output] = isInsideParForLoopIQM()
%
% [INPUT]
%
% [OUTPUT]
% output:   0 if not called within a parfor loop
%           1 if called within a parfor loop

% <<<COPYRIGHTSTATEMENT - IQM TOOLS LITE>>>

% Check if parallel toolbox available - if not then simply return 0
if ~exist('parpool'),
    output = 0;
    return
end

% Check
output = ~isempty(getCurrentTask());
```
