## Supplementary material for "A systems biology analysis of adrenergically stimulated adiponectin exocytosis in white adipocytes": All scripts used in this work: sizeParallelIQM.m

```
function [poolsize] = sizeParallelIQM()
% This function returns the size of the current pool of parallel
% processores reserved for MATLAB.
%
% [SYNTAX]
% [poolsize] = sizeParallelIQM()
%
% [INPUT]
%
% [OUTPUT]
% poolsize:     Number of parallel processors available

% <<<COPYRIGHTSTATEMENT - IQM TOOLS LITE>>>

if ~exist('parpool'),
    poolsize = 0;
    return
end

poolobj = gcp('nocreate'); % If no pool, do not create new one.
if isempty(poolobj)
    poolsize = 0;
else
    poolsize = poolobj.NumWorkers;
end
```
