## Supplementary material for "A systems biology analysis of adrenergically stimulated adiponectin exocytosis in white adipocytes": All scripts used in this work: startParallelIQM.m

```
function [killMATLABpool] = startParallelIQM(NRNODES)
% This function requests parallel nodes from the MATLAB parallel toolbox.
%
% Will only request parallel nodes if NRNODES>1.
%
% [SYNTAX]
% [killMATLABpool] = startParallelIQM()
% [killMATLABpool] = startParallelIQM(NRNODES)
%
% [INPUT]
% NRNODES:          Number of nodes requested. If not provided, then the
%                   default number is used, defined in SETUP_PATHS_TOOLS_IQMPRO
%
% [OUTPUT]
% killMATLABpool:   =0 if matlabpool active before call to this function
%                   =1 if matlabpool not active before call to this function
%                   Flag can be passed to stopParallelIQM

% <<<COPYRIGHTSTATEMENT - IQM TOOLS LITE>>>

% Check if parallel toolbox available - if not then simply return
if ~exist('parpool'),
    killMATLABpool = 0;
    return
end

% Evaluate SETUP_PATHS_TOOLS_IQMPRO to get name of MATLABPOOL profile and
% to obtain default number of nodes to connect to
SETUP_PATHS_TOOLS_IQMPRO

% Handle optional number of input arguments
if nargin==0,
    NRNODES = N_PROCESSORS_PAR;
end

% Handle the parallel nodes
killMATLABpool = 0;
if NRNODES > 1,
    if sizeParallelIQM() == 0,
        warning off
        eval(sprintf('parpool(''%s'',%d);',MATLABPOOL_PROFILE,NRNODES));
        warning on
        killMATLABpool = 1;
    end
end
```
