## Supplementary material for "A systems biology analysis of adrenergically stimulated adiponectin exocytosis in white adipocytes": All scripts used in this work: stopParallelIQM.m

```
function [] = stopParallelIQM(killMATLABpool)
% This function closes the connection to parallel nodes via the parallel 
% toolbox in MATLAB
%
% [SYNTAX]
% [] = stopParallelIQM()
% [] = stopParallelIQM(killMATLABpool)
%
% [INPUT]
% killMATLABpool: if 0 then do not close matlabpool.
%
% [OUTPUT]

% <<<COPYRIGHTSTATEMENT - IQM TOOLS LITE>>>

% Check if parallel toolbox available - if not then simply return
if ~exist('parpool'),
    return
end

% Handle variable input arguments
if nargin==0,
    killMATLABpool = 1;
end

% Close connection
if sizeParallelIQM()>0 && killMATLABpool==1,
    delete(gcp('nocreate'));
end
```
