## Supplementary material for "A systems biology analysis of adrenergically stimulated adiponectin exocytosis in white adipocytes": All scripts used in this work: setseedIQM.m

```
function [] = setseedIQM(seed)
% Set the default stream to defaultSeed. 
%
% [SYNTAX]
% [] = setseedIQM(defaultSeed)
%
% [INPUT]
% defaultSeed:   integer between 0 and 2^32.
%
% [OUTPUT]

% <<<COPYRIGHTSTATEMENT - IQM TOOLS LITE>>>

% Check that seed is an integer between 0 and 2^32
if (mod(seed,1)~=0) || seed<0 || seed>2^32,
    error('Seed must be an integer between 0 and 2^32');
end

% Get the version release
s = RandStream.create('mrg32k3a','Seed',seed);
RandStream.setGlobalStream(s);
```
