## Supplementary material for "A systems biology analysis of adrenergically stimulated adiponectin exocytosis in white adipocytes": All scripts used in this work: tempdirIQM.m

```
function [tmpdir] = tempdirIQM()
% tempdirIQM:    returns the path to the desired temp folder to be used by
% IQM Tools

% <<<COPYRIGHTSTATEMENT - IQM TOOLS LITE>>>

% just use the systems default temporary folder!
tmpdir = tempdir;
```
