## Supplementary material for "A systems biology analysis of adrenergically stimulated adiponectin exocytosis in white adipocytes": All scripts used in this work: tempnameIQM.m

```
function [tmpname] = tempnameIQM( input_args )
% tempnameIQM does the same as tempname in MATLAB but uses the IQM Tools own
% TEMP folder.

% <<<COPYRIGHTSTATEMENT - IQM TOOLS LITE>>>

[dummy, filename] = fileparts(tempname);
tmpname = [tempdirIQM,filename];
```
