## Supplementary material for "A systems biology analysis of adrenergically stimulated adiponectin exocytosis in white adipocytes": All scripts used in this work: usernameIQM.m

```
function [username] = usernameIQM()
% usernameIQM: Get the name of the current user

% <<<COPYRIGHTSTATEMENT - IQM TOOLS LITE>>>

if ispc,
    username = getenv('UserName');
else
    username = getenv('USER');
end
return
```
