## Supplementary material for "A systems biology analysis of adrenergically stimulated adiponectin exocytosis in white adipocytes": All scripts used in this work: display.m

```
function [] = display(dos)
% display: Displays information about IQMdosing object. This function is 
% called by MATLAB whenever an object is the result of a statement that
% is not terminated by a semicolon. 

% <<<COPYRIGHTSTATEMENT - IQM TOOLS LITE>>>

%%%%%%%%%%%%%%%%%%%%%%%%%%%%%%%%%%%%%%%%%%%%%%%
% COLLECT INFORMATION ABOUT THE EXPERIMENT
%%%%%%%%%%%%%%%%%%%%%%%%%%%%%%%%%%%%%%%%%%%%%%%
nrinputs = length(dos.inputs);
nameinputs = {dos.inputs.name};
typeinputs = {dos.inputs.type};

%%%%%%%%%%%%%%%%%%%%%%%%%%%%%%%%%%%%%%%%%%%%%%%
% DISPLAY INFORMATION
%%%%%%%%%%%%%%%%%%%%%%%%%%%%%%%%%%%%%%%%%%%%%%%
text = sprintf('\tIQMdosing\n\t===========\n');
text = sprintf('%s\tName:  %s\n',text,dos.name);
% text = sprintf('%s\tNotes: %s\n',text,dos.notes);
text = sprintf('%s\tNumber inputs:     %d',text,nrinputs);
disp(text);
for k=1:length(nameinputs),
    if length(dos.inputs(k).time) > 1,
        type2 = 'muliple';
    else 
        type2 = 'single';
    end
    if ~isempty(dos.inputs(k).Tlag),
        if ischar(dos.inputs(k).Tlag),
            disp(sprintf('\t\t%s: %s %s\t\t(LAG - NON NUMERIC)',dos.inputs(k).name,type2,dos.inputs(k).type));
        else
            disp(sprintf('\t\t%s: %s %s\t\t(LAG - NUMERIC)',dos.inputs(k).name,type2,dos.inputs(k).type));
        end
    else
        disp(sprintf('\t\t%s: %s %s',dos.inputs(k).name,type2,dos.inputs(k).type));
    end
end
```
