## Supplementary material for "A systems biology analysis of adrenergically stimulated adiponectin exocytosis in white adipocytes": All scripts used in this work: IQMdosing.m

```
function [dos] = IQMdosing(varargin)
% IQMdosing: creates an IQMdosing object defining a dosing schedule
%
% USAGE:
% ======
% [dos] = IQMdosing()             creates an empty IQMdosing object
% [dos] = IQMdosing(IQMstructure)  creates an IQMdosing object from a MATLAB
%                                   structure in the internal IQMdosing format
% [dos] = IQMdosing(dosin)        construction from a given IQMdosing object (dosin)
% [dos] = IQMdosing('file.dos')   converting a IQMdosing text description 
%                                   to an IQMdosing object.
%
% Output Arguments:
% =================
% dos: IQMdosing object 

% <<<COPYRIGHTSTATEMENT - IQM TOOLS LITE>>>

%%%%%%%%%%%%%%%%%%%%%%%%%%%%%%%%%%%%%%%%%%%%%%%
% CHECK THE TYPE OF THE INPUT
%%%%%%%%%%%%%%%%%%%%%%%%%%%%%%%%%%%%%%%%%%%%%%%
if nargin==0,
    inputType = 'empty';
elseif nargin == 1 || nargin == 2,
    if isIQMdosing(varargin{1}),
        inputType = 'IQMdosing';
        dosInput = varargin{1};
    elseif isstruct(varargin{1}),
        inputType = 'IQMstructure';
        IQMstructure = varargin{1};
    elseif ischar(varargin{1}),
        % check if '.dos' given as extension. If yes, then import text description
        filename = varargin{1};
        if ~isempty(strfind(filename,'.dos')),
            inputType = 'TextDosFile';
        elseif strcmp('DosingAsTextString', varargin{2}),
            inputType = varargin{2};
        else
            error('Input argument of unknown type');
        end
    else 
        error('Input argument of unknown type');
    end
else
    error('Wrong number of input arguments.');
end

%%%%%%%%%%%%%%%%%%%%%%%%%%%%%%%%%%%%%%%%%%%%%%%
% CONSTRUCT THE IQMdosing OBJECT 
%%%%%%%%%%%%%%%%%%%%%%%%%%%%%%%%%%%%%%%%%%%%%%%
if strcmp('empty',inputType),
    % Create empty IQMstructure
    % parameters substructure
    parametersStruct = struct('name',{},'value',{},'notes',{});
    % inputs substructure
    inputsStruct = struct('name',{},'type',{},'time',{},'Tlag',{},'D',{},'parameters',parametersStruct,'TlagNotes',{},'notes',{});
    % Create IQMstructure
    IQMstructure = struct('type','IQMdosing','name','unnamed_dosing','notes','no notes','inputs',inputsStruct);
    % construct the dosing object
    dos = class(IQMstructure,'IQMdosing');
elseif strcmp('IQMdosing',inputType),
    % copy the object
    dos = dosInput;
elseif strcmp('IQMstructure',inputType),
    % check if the given structure is a IQMstructure 
    if isfield(IQMstructure,'type'),
        if ~strcmp(IQMstructure.type,'IQMdosing'),
            error('Given structure is not a valid internal IQMdosing structure.');
        end
    else
        error('Given structure is not a valid internal IQMdosing structure.');
    end
    % construct the dosing object
    dos = class(IQMstructure,'IQMdosing');
elseif strcmp('TextDosFile',inputType),
    % check if a file with given filename exists
    [path,filename,ext] = fileparts(filename);
    filename = fullfile(path, [filename '.dos']); 
    if ~exist(filename),
        error(sprintf('Dosing file, "%s", does not exist.', filename));
    end
    % If file exists then first load it
    dosText = fileread(filename);
    % then convert it to IQMstructure
    [IQMstructure, errorMsg] = convertTextToDosIQM(dosText);
    % Check if error occurred while importing the dosing description
    if ~isempty(errorMsg),
        error(errorMsg);
    end
    % construct the dosing object
    dos = class(IQMstructure,'IQMdosing');
elseif strcmp('DosingAsTextString', inputType),
    dosText = varargin{1};
    % then convert text dosing to IQMstructure
    [IQMstructure, errorMsg] = convertTextToDosIQM(dosText);
    % Check if error occurred while importing the dosing definition
    if ~isempty(errorMsg),
        error(errorMsg);
    end
    % construct the dosing object
    dos = class(IQMstructure,'IQMdosing');
else
    error('Wrong input arguments.');
end

% Cycle through all inputs in the dosing scheme and expand the dose and
% paramter to time length if needed. but only if dose or parameter is a
% scalar and time a vector.
ds = struct(dos);
for k=1:length(ds.inputs),
    if ~strcmp(ds.inputs(k).type,'BOLUS'),
        NTIME = length(ds.inputs(k).time);
        NDOSE = length(ds.inputs(k).D);
        NPARA = length(ds.inputs(k).parameters.value);
        if NDOSE==1 && NTIME>1,
            ds.inputs(k).D = ds.inputs(k).D*ones(1,NTIME);
        end
        if NPARA==1 && NTIME>1,
            ds.inputs(k).parameters.value = ds.inputs(k).parameters.value*ones(1,NTIME);
        end
    end
end

%% Check
for k=1:length(ds.inputs),
    if ~strcmp(ds.inputs(k).type,'BOLUS'),
        NTIME = length(ds.inputs(k).time);
        NDOSE = length(ds.inputs(k).D);
        NPARA = length(ds.inputs(k).parameters.value);
        if NDOSE ~= NTIME,
            error('Different lengths of time and dose vector definitions in dosing object. Scalar dose is expanded automatically to length of dose vector.');
        end
        if NPARA ~= NTIME,
            error('Different lengths of time and parameter vector definitions in dosing object. Scalar parameter is expanded automatically to length of dose vector.');
        end
    end
end

%% Create dosing object for output
dos = class(ds,'IQMdosing');
```
