## Supplementary material for "A systems biology analysis of adrenergically stimulated adiponectin exocytosis in white adipocytes": All scripts used in this work: IQMstruct.m

```
function [IQMstructure] = IQMstruct(dos)
% IQMstruct: This function returns the internal data structure
% of an IQMdosing object
%
% USAGE:
% ======
% [IQMstructure] = IQMstruct(dos) 
%
% dos: IQMdosing object 
%
% Output Arguments:
% =================
% IQMstructure: internal data structure of the IQMdosing object

% <<<COPYRIGHTSTATEMENT - IQM TOOLS LITE>>>


%%%%%%%%%%%%%%%%%%%%%%%%%%%%%%%%%%%%%%%%%%%%%%%
% VERY SIMPLE FUNCTION
%%%%%%%%%%%%%%%%%%%%%%%%%%%%%%%%%%%%%%%%%%%%%%%
IQMstructure = struct(dos);
```
