## Supplementary material for "A systems biology analysis of adrenergically stimulated adiponectin exocytosis in white adipocytes": All scripts used in this work: display.m

```
function [] = display(exp)
% display: Displays information about IQMexperiment object. This function is 
% called by MATLAB whenever an object is the result of a statement that
% is not terminated by a semicolon. 

% <<<COPYRIGHTSTATEMENT - IQM TOOLS LITE>>>


%%%%%%%%%%%%%%%%%%%%%%%%%%%%%%%%%%%%%%%%%%%%%%%
% COLLECT INFORMATION ABOUT THE EXPERIMENT
%%%%%%%%%%%%%%%%%%%%%%%%%%%%%%%%%%%%%%%%%%%%%%%
name = exp.name;
notes = exp.notes;
nrparamicsettings = length(exp.paramicsettings);
nrparameterchanges = length(exp.parameterchanges);
nrstatechanges = length(exp.stateevents);

%%%%%%%%%%%%%%%%%%%%%%%%%%%%%%%%%%%%%%%%%%%%%%%
% DISPLAY INFORMATION
%%%%%%%%%%%%%%%%%%%%%%%%%%%%%%%%%%%%%%%%%%%%%%%
text = sprintf('\tIQMexperiment\n\t============\n');
text = sprintf('%s\tName:  %s\n',text,name);
text = sprintf('%s\tNumber Param+IC Settings: %d\n',text,nrparamicsettings);
text = sprintf('%s\tNumber Param Changes:     %d\n',text,nrparameterchanges);
text = sprintf('%s\tNumber State Changes:     %d\n',text,nrstatechanges);
disp(text);
```
