## Supplementary material for "A systems biology analysis of adrenergically stimulated adiponectin exocytosis in white adipocytes": All scripts used in this work: IQMexperiment.m

```
function [exp] = IQMexperiment(varargin)
% IQMexperiment: creates an experiment object defining experiment settings 
%
% USAGE:
% ======
% [exp] = IQMexperiment()                creates an empty IQMexperiment object
% [exp] = IQMexperiment(IQMstructure)     creates an IQMexperiment object from a MATLAB
%                                       structure in the internal experiment format
% [exp] = IQMexperiment(expin)           construction from a given IQMexperiment object (expin)
% [exp] = IQMexperiment('file.exp')      converting a experiment text description 
%                                       to an IQMexperiment object.
% [exp] = IQMexperiment('file.exp',path2paramset)  converting a experiment text description 
%                                       to an IQMexperiment object. When
%                                       "activeSet" and/or "parameterSet"
%                                       definitions are used then the path
%                                       to the root folder of these
%                                       definitions needs to be provided.
%                                       Otherwise: ERROR.
%
% Output Arguments:
% =================
% exp: IQMexperiment object 

% <<<COPYRIGHTSTATEMENT - IQM TOOLS LITE>>>

flag = 0;
path2paramset = '';

%%%%%%%%%%%%%%%%%%%%%%%%%%%%%%%%%%%%%%%%%%%%%%%
% CHECK THE TYPE OF THE INPUT
%%%%%%%%%%%%%%%%%%%%%%%%%%%%%%%%%%%%%%%%%%%%%%%
if nargin==0,
    inputType = 'empty';
elseif nargin == 1 || nargin == 2,
    if strcmp('IQMexperiment',class(varargin{1})),
        inputType = 'IQMexperiment';
        expInput = varargin{1};
    elseif isstruct(varargin{1}),
        inputType = 'IQMstructure';
        IQMstructure = varargin{1};
    elseif ischar(varargin{1}),
        % check if '.txt' given as extension. If yes, then import text
        % description
        filename = varargin{1};
        if ~isempty(strfind(filename,'.exp')),
            inputType = 'TextExpFile';
            if nargin == 2,
                path2paramset = varargin{2};
            end
        elseif nargin == 2,
            if strcmp('ExperimentAsTextString', varargin{2}),
                inputType = varargin{2};
            end
        else
            error('Input argument of unknown type');
        end
    else 
        error('Input argument of unknown type');
    end
else
    error('Wrong number of input arguments.');
end

%%%%%%%%%%%%%%%%%%%%%%%%%%%%%%%%%%%%%%%%%%%%%%%
% CONSTRUCT THE IQMexperiment OBJECT 
%%%%%%%%%%%%%%%%%%%%%%%%%%%%%%%%%%%%%%%%%%%%%%%
if strcmp('empty',inputType),
    % Create empty IQMstructure
    % parameter settings substructure
    paramicsettingsStruct = struct('name',{},'formula',{},'notes',{},'icflag',{});
    % parameter settings substructure
    parameterchangesStruct = struct('name',{},'formula',{},'notes',{});
    % event assignment substructure
    eventassignmentStruct = struct('variable',{},'formula',{});
    % state events substructure
    stateeventsStruct = struct('name',{},'trigger',{},'assignment',eventassignmentStruct,'notes',{});
    % Create IQMstructure
    IQMstructure = struct('name','unnamed_experiment','notes','no notes','paramicsettings',paramicsettingsStruct,'parameterchanges',parameterchangesStruct,'stateevents',stateeventsStruct);
    % construct the model object
    exp = class(IQMstructure,'IQMexperiment');
elseif strcmp('IQMexperiment',inputType),
    % copy the model object
    exp = expInput;
elseif strcmp('IQMstructure',inputType),
    % check if the given structure is a IQMstructure (only check the
    % top-level fields)
    checkFields = {'name','notes','paramicsettings','parameterchanges','stateevents'};
    for k = 1:length(checkFields),
        if ~isfield(IQMstructure,checkFields{k}),
            error('Given structure is not a valid internal IQMexperiment structure.');
        end
    end
    % construct the model object
    exp = class(IQMstructure,'IQMexperiment');
elseif strcmp('TextExpFile',inputType),
    % check if a file with given filename exists
    [path,filename,ext] = fileparts(filename);
    if isempty(path2paramset),
        filename = fullfile(path, [filename '.exp']); 
    else
        filename = fullfile(path2paramset,path, [filename '.exp']); 
    end
    if ~exist(filename),
        errorMsg = sprintf('Experiment file, "%s", does not exist.', filename);
        error(errorMsg);
    end
    % If file exists then first load it
    expText = fileread(filename);
	% take commented lines out of the experiment description
	expText = regexprep(expText,'\n%[^\n]*','');
    %%%%%%%%%%%%%%%%%%%%%%%%%%%%%%%%%%%%%%%%%%%%%%%%
    % CHECK IF "activeSet" and/or "parameterSet" 
    % definitions are used. If yes, then preprocess
    % the experiment text.
    %%%%%%%%%%%%%%%%%%%%%%%%%%%%%%%%%%%%%%%%%%%%%%%%
    expText = checkProcessActiveSetParameterSetIQM(expText,path2paramset);
    % then convert it to IQMstructure
    [IQMstructure, errorMsg] = convertTextToExpIQM(expText);
    % Check if error occurred while importing the SBML model
    if ~isempty(errorMsg),
        error(errorMsg);
    end
    % construct the model object
    exp = class(IQMstructure,'IQMexperiment');
elseif strcmp('ExperimentAsTextString', inputType),
    expText = varargin{1};
	% take commented lines out of the experiment description
	expText = regexprep(expText,'\n%[^\n]*','');
    %%%%%%%%%%%%%%%%%%%%%%%%%%%%%%%%%%%%%%%%%%%%%%%%
    % CHECK IF "activeSet" and/or "parameterSet" 
    % definitions are used. If yes, then preprocess
    % the experiment text.
    %%%%%%%%%%%%%%%%%%%%%%%%%%%%%%%%%%%%%%%%%%%%%%%%
    expText = checkProcessActiveSetParameterSetIQM(expText,path2paramset);
    % then convert text experiment to IQMstructure
    [IQMstructure, errorMsg] = convertTextToExpIQM(expText);
    % Check if error occurred while importing the SBML model
    if ~isempty(errorMsg),
        error(errorMsg);
    end
    % construct the model object
    exp = class(IQMstructure,'IQMexperiment');
else
    error('Wrong input arguments.');
end
return
```
