## Supplementary material for "A systems biology analysis of adrenergically stimulated adiponectin exocytosis in white adipocytes": All scripts used in this work: display.m

```
function [] = display(measurement)
% display: Displays information about IQMmeasurement. This function is 
% called by MATLAB whenever an object is the result of a statement that
% is not terminated by a semicolon. 

% <<<COPYRIGHTSTATEMENT - IQM TOOLS LITE>>>


%%%%%%%%%%%%%%%%%%%%%%%%%%%%%%%%%%%%%%%%%%%%%%%
% COLLECT INFORMATION ABOUT THE DATA OBJECT
%%%%%%%%%%%%%%%%%%%%%%%%%%%%%%%%%%%%%%%%%%%%%%%
name = measurement.name;
notes = measurement.notes;
numbermeasurements = length(measurement.data);
numbertimesteps = length(measurement.time);
errorboundFlag = 0;
allMeasurementsDone = 1;
for k = 1:length(measurement.data),
    % check if errorbounds are present
    if ~isempty(measurement.data(k).maxvalues) && max(isnan(measurement.data(k).maxvalues))~=1,
        errorboundFlag = 1;
    end
    % check if NaN present in data
    if ~isempty(find(isnan(measurement.data(k).values))),
        allMeasurementsDone = 0;
    end
end

%%%%%%%%%%%%%%%%%%%%%%%%%%%%%%%%%%%%%%%%%%%%%%%
% DISPLAY INFORMATION
%%%%%%%%%%%%%%%%%%%%%%%%%%%%%%%%%%%%%%%%%%%%%%%
text = sprintf('\tIQMmeasurement\n\t=============\n');
text = sprintf('%s\tName: %s\n',text,name);
text = sprintf('%s\tMeasured components:\t%d\n',text,numbermeasurements);
text = sprintf('%s\tNumber time points: \t%d\n',text,numbertimesteps);
if errorboundFlag == 1,
    text = sprintf('%s\tError bound information present at least for one measurement.\n',text);
end
if allMeasurementsDone == 1,
    text = sprintf('%s\tMeasurements for all time points present\n',text);
else
    text = sprintf('%s\tMeasurements not present for all time points\n',text);
end
disp(text);
```
