## Supplementary material for "A systems biology analysis of adrenergically stimulated adiponectin exocytosis in white adipocytes": All scripts used in this work: IQMmeasurement.m

```
function [output] = IQMmeasurement(varargin)
% IQMmeasurement: creates an object that is intended to contain 
% measurement data - from real experiments or insilico computations
%
% USAGE:
% ======
% [output] = IQMmeasurement()                creates an empty IQMmeasurement object
% [output] = IQMmeasurement(struct)          creates an IQMmeasurement object from
%                                           a MATLAB structure in the internal 
%                                           measurement object format
% [output] = IQMmeasurement(inputobject)     construction from a given IQMmeasurement object 
% [output] = IQMmeasurement('filename.xls')  converting an excel file to 
%                                           an IQMmeasurement object.
% [output] = IQMmeasurement('filename.csv')  converting an CSV (comma
%                                           separated value) file to  
%                                           an IQMmeasurement object
%
% Output Arguments:
% =================
% output: IQMmeasurement object 
%         In the case that several sets of measurement data are defined in 
%         an Excel file (on separate sheets) the output will be cell-array
%         in which the elements are IQMmeasurement objects.

% <<<COPYRIGHTSTATEMENT - IQM TOOLS LITE>>>


%%%%%%%%%%%%%%%%%%%%%%%%%%%%%%%%%%%%%%%%%%%%%%%
% CHECK THE TYPE OF THE INPUT
%%%%%%%%%%%%%%%%%%%%%%%%%%%%%%%%%%%%%%%%%%%%%%%
if nargin==0,
    inputType = 'empty';
elseif nargin == 1,
    if strcmp('IQMmeasurement',class(varargin{1})),
        inputType = 'IQMmeasurement';
        measurementinput = varargin{1};
    elseif strcmp('struct',class(varargin{1})),
        inputType = 'measurementstructure';
        measurementstructure = varargin{1};
    elseif strcmp('char',class(varargin{1})),
        % assume filenamec given as input argument
        % check if '.xls' given as extension. If yes, then import measurement from
        % excel file format
        filename = varargin{1};
        if ~isempty(strfind(lower(filename),'.xls')),
            inputType = 'XLSmeasurementFile';
        elseif ~isempty(strfind(lower(filename),'.csv')),
            inputType = 'CSVmeasurementFile';
        else
            error('Unknown filetype!');
        end
    else 
        error('Input argument of unknown type');
    end
else
    error('Wrong number of input arguments.');
end

%%%%%%%%%%%%%%%%%%%%%%%%%%%%%%%%%%%%%%%%%%%%%%%
% CONSTRUCT THE IQMMODEL OBJECT 
%%%%%%%%%%%%%%%%%%%%%%%%%%%%%%%%%%%%%%%%%%%%%%%
if strcmp('empty',inputType),
    % Create empty measurement structure
    % data substructure
    datastructure = struct('name',{},'notes',{},'values',{},'maxvalues',{},'minvalues',{});
    % Create measurement structure
    measurementsStruct = struct('name','untitled','notes','','time',[],'data',datastructure);
    output = class(measurementsStruct,'IQMmeasurement');
elseif strcmp('IQMmeasurement',inputType),
    % copy the model object
    output = measurementinput;
elseif strcmp('measurementstructure',inputType),
    % check if the given structure is a measurement structure (only check the
    % top-level fields)
    checkFields = {'name','notes','time','data'};
    for k = 1:length(checkFields),
        if ~isfield(measurementstructure,checkFields{k}),
            errorMsg = sprintf('Given structure is not a valid internal IQMmeasurement structure.');
            error(errorMsg);
        end
    end
    % construct the data object
    output = class(measurementstructure,'IQMmeasurement');
elseif strcmp('XLSmeasurementFile',inputType),
    % check if a file with given filename exists
    [path,filename,ext] = fileparts(filename);
    filename = fullfile(path, [filename '.xls']); 
    if ~exist(filename),
        errorMsg = sprintf('XLS measurement file, "%s", does not exist.', filename);
        error(errorMsg);
    end
    % If file exists then import it
    [outputstructures,errorMsg] = IQMimportXLSmeasurement(filename);
    % Check if error occurred while importing the XLS data
    if length(errorMsg) ~= 0,
        error(errorMsg);
    end
    % construct the data objects (one for each sheet)
    if length(outputstructures) == 1,
        output = class(outputstructures{1},'IQMmeasurement');
    else
        output = {};
        for k = 1:length(outputstructures),
            output{k} = class(outputstructures{k},'IQMmeasurement');
        end
    end
elseif strcmp('CSVmeasurementFile',inputType),
    % check if a file with given filename exists
    [path,filename,ext] = fileparts(filename);
    filename = fullfile(path, [filename '.csv']); 
    if ~exist(filename),
        errorMsg = sprintf('CSV measurement file, "%s", does not exist.', filename);
        error(errorMsg);
    end
    % If file exists then import it
    [measurementstructure,errorMsg] = IQMimportCSVmeasurement(filename);
    % Check if error occurred while importing the CSV data
    if length(errorMsg) ~= 0,
        error(errorMsg);
    end
    % construct the measurement object
    output = class(measurementstructure,'IQMmeasurement');
end
return
```
