## Supplementary material for "A systems biology analysis of adrenergically stimulated adiponectin exocytosis in white adipocytes": All scripts used in this work: IQMstruct.m

```
function [datastructure] = IQMstruct(iqmmeasurement)
% IQMstruct: This function returns the internal data structure
% of an IQMmeasurement object
%
% USAGE:
% ======
% [datastructure] = IQMstruct(iqmmeasurement) 
%
% iqmmeasurement: IQMmeasurement data object 
%
% Output Arguments:
% =================
% datastructure: internal data structure of the IQMmeasurement object

% <<<COPYRIGHTSTATEMENT - IQM TOOLS LITE>>>


%%%%%%%%%%%%%%%%%%%%%%%%%%%%%%%%%%%%%%%%%%%%%%%
% VERY SIMPLE FUNCTION
%%%%%%%%%%%%%%%%%%%%%%%%%%%%%%%%%%%%%%%%%%%%%%%
datastructure = struct(iqmmeasurement);
```
