## Supplementary material for "A systems biology analysis of adrenergically stimulated adiponectin exocytosis in white adipocytes": All scripts used in this work: display.m

```
function [] = display(iqm)
% display: Displays information about IQMmodel. This function is 
% called by MATLAB whenever an object is the result of a statement that
% is not terminated by a semicolon. 

% <<<COPYRIGHTSTATEMENT - IQM TOOLS LITE>>>

%%%%%%%%%%%%%%%%%%%%%%%%%%%%%%%%%%%%%%%%%%%%%%%
% COLLECT INFORMATION ABOUT THE MODEL
%%%%%%%%%%%%%%%%%%%%%%%%%%%%%%%%%%%%%%%%%%%%%%%
numberStates = length(iqm.states);
numberVariables = length(iqm.variables);
numberParameters = length(iqm.parameters);
numberReactions = length(iqm.reactions);
numberFunctions = length(iqm.functions);
numberEvents = length(iqm.events);
functionsMATLABpresent = ~isempty(iqm.functionsMATLAB);
delaysPresent = usedelayIQM(iqm);
numberARs = length(iqm.algebraic);
fastReactionsPresent = usefastIQM(iqm);
nnICsPresent = ~hasonlynumericICsIQM(iqm);
nrInputs = length(iqm.inputs);
nrOutputs = length(iqm.outputs);

%%%%%%%%%%%%%%%%%%%%%%%%%%%%%%%%%%%%%%%%%%%%%%%
% DISPLAY INFORMATION
%%%%%%%%%%%%%%%%%%%%%%%%%%%%%%%%%%%%%%%%%%%%%%%
text = sprintf('\tIQMmodel\n\t=======\n');
text = sprintf('%s\tName: %s\n',text,iqm.name);
text = sprintf('%s\tNumber States:\t\t\t%d\n',text,numberStates);
text = sprintf('%s\tNumber Variables:\t\t%d\n',text,numberVariables);
text = sprintf('%s\tNumber Parameters:\t\t%d\n',text,numberParameters);
text = sprintf('%s\tNumber Reactions:\t\t%d\n',text,numberReactions);
text = sprintf('%s\tNumber Functions:\t\t%d\n',text,numberFunctions);
if numberEvents > 0,
    text = sprintf('%s\tNumber Events:\t\t\t%d\n',text,numberEvents);
end
if numberARs > 0,
    text = sprintf('%s\tNumber Algebraic Rules:\t%d\n',text,numberARs);
end
if nrInputs > 0,
    text = sprintf('%s\tNumber Inputs:\t\t\t%d\n',text,nrInputs);
end
if nrOutputs > 0,
    text = sprintf('%s\tNumber Outputs:\t\t\t%d\n',text,nrOutputs);
end
if functionsMATLABpresent,
    text = sprintf('%s\tMATLAB functions present\n',text);
end
if delaysPresent,
    text = sprintf('%s\tDelay function(s) present',text);
end
if fastReactionsPresent,
    text = sprintf('%s\tFast reactions are present in the model:\n\t\tPlease note that this information is ONLY taken into account\n\t\tduring simulation using IQMsimulate. NO other function will\n\t\tconsider fast reactions.\n',text);
end
if nnICsPresent,
    text = sprintf('%s\tNon-numeric initial conditions are present in the model.\n\t\tPlease check the documentation for further information.\n',text);
end    
if numberARs > 0,
    text = sprintf('%s\tAlgebraic rules present in the model.\n\t\tYou should not use ANY other function on this model than IQMsimulate.\n',text);
end
disp(text);
```
