## Supplementary material for "A systems biology analysis of adrenergically stimulated adiponectin exocytosis in white adipocytes": All scripts used in this work: IQMmodel.m

```
function [model] = IQMmodel(varargin)
% IQMmodel: creates a model object 
%
% [SYNTAX]
% [model] = IQMmodel()                
% [model] = IQMmodel(model)    
% [model] = IQMmodel(model,nocheck) 
% [model] = IQMmodel(model,nocheck,useSBMLnames) 
% [model] = IQMmodel(model,nocheck,useSBMLnames,stripSBMLinfo) 
%
% [INPUT]
% model:            Can have multiple meanings
%                   - If it is an IQMmodel structure (struct(model)) then this structure
%                     is converted to an IQMmodel
%                   - If it is a filename (file.txt, file.txtbc, file.xml) then the
%                     file is interpreted as a valid IQMmodel text file (txt or txtbc) 
%                     or as an SBML file (.xml) and imported accordingly to an IQMmodel object.
%                     If no extension is specified, then SBML format is assumed.
%                   - If it is an IQMmodel, then this model is just passed through and 
%                     returned as output argument.
% nocheck:          0: standard behavior. Consistency of models is checked and errors 
%                      are shown if something is wrong.
%                   1: Consistency is not checked. Instead of errors warnings are shown.
%                      This functionality is mainly thought for the import of (in)completely 
%                      defined SBML models. It does not require that a model is fully defined 
%                      in terms of parameter values, and rate equations, etc. 
%                   (default: 0)
% useSBMLnames:     0: During SBML import, use the element IDs as names of model components
%                   1: During SBML import, use the element NAMEs as names of model components
%                      This is needed for cases where the SBML modeling software in which the model
%                      was created uses random SBML IDs, which translates to unreadable models.
%                      This is the case for CellDesigner and Simbiology (e.g.). The names in SBML models
%                      are not required to be unique. This is taken care off by making them unique through 
%                      appending indices (_1, _2, ...).
%                   (default: 0)
% stripSBMLinfo:    0: keep additional information for SBML conversion in the model after import
%                   1: remove additional information from SBML import
%                   (default: 0)
%
% [OUTPUT]
% model:            IQMmodel object 

% <<<COPYRIGHTSTATEMENT - IQM TOOLS LITE>>>

% Handle optional input flags
nocheck         = 0;
useSBMLnames    = 0;
stripSBMLinfo   = 0;
if nargin > 1,
    nocheck = varargin{2};
end
if nargin > 2,
    useSBMLnames = varargin{3};
end
if nargin > 3,
    stripSBMLinfo = varargin{4};
end

% Check something
if ~isnumeric(nocheck),
    error('Problem with IQMmodel');
end

% CHECK THE TYPE OF THE INPUT
if nargin==0,
    inputType = 'empty';
end

if nargin >0,
    if strcmp('IQMmodel',class(varargin{1})),
        inputType = 'IQMmodel';
        iqmInput = varargin{1};
    elseif isstruct(varargin{1}),
        inputType = 'IQMstructure';
        IQMstructure = varargin{1};
    elseif ischar(varargin{1}),
        % check if '.txt' or '.txtbc' given as extension. If yes, then import text
        % model, otherwise assume an SBML model is to be imported.
        filename = varargin{1};
        if ~isempty(strfind(filename,'.txtbc')),
            inputType = 'TextBCModelFile';
        elseif ~isempty(strfind(filename,'.txt')),
            inputType = 'TextModelFile';
%         elseif strcmp('ModelAsTextString', flag),
%             modelText = varargin{1};
%             inputType = flag;
%         elseif strcmp('ModelAsTextBCString', flag),
%             modelText = varargin{1};
%             inputType = flag;
        else
            inputType = 'SBMLmodelFile';
        end        
    else 
        error('Input argument of unknown type');
    end
end


% model.inputs.name:       input name
% model.inputs.factors:    cell-array with input factors
% model.inputs.terms:      cell-array with complete input string (for
%                          simpler removing)
% model.inputs.stateindex: vector with stateindices to which the 
%                          input is applied
% model.inputs.parindex:   index of the INPUT* parameter definition in
%                          the IQMmodel (used to remove it when
%                          parameters are written to (e.g.) an MLXTRAN
%                          file).   
% model.outputs.name:      output name
% model.outputs.formula:   output formula
% model.outputs.notes:     output notes
% model.outputs.varindex:  index of output in model variables


%%%%%%%%%%%%%%%%%%%%%%%%%%%%%%%%%%%%%%%%%%%%%%%
% CONSTRUCT THE IQMMODEL OBJECT 
%%%%%%%%%%%%%%%%%%%%%%%%%%%%%%%%%%%%%%%%%%%%%%%
if strcmp('empty',inputType),
    % Create empty IQMstructure
    % inputs substructure
    inputsStruct = struct('name',{},'factors',{},'terms',{},'stateindex',{},'parindex',{});
    % outputs substructure
    outputsStruct = struct('name',{},'formula',{},'notes',{},'varindex',{});
    % functions substructure
    functionsStruct = struct('name',{},'arguments',{},'formula',{},'notes',{});
    % states substructure
    statesStruct = struct('name',{},'initialCondition',{},'ODE',{},'type',{},'compartment',{},'unittype',{},'notes',{});
    % algebraic substructure
    algebraicStruct = struct('name',{},'formula',{},'initialCondition',{},'type',{},'compartment',{},'unittype',{},'notes',{});
    % parameters substructure
    parametersStruct = struct('name',{},'value',{},'type',{},'compartment',{},'unittype',{},'notes',{});
    % variables substructure
    variablesStruct = struct('name',{},'formula',{},'type',{},'compartment',{},'unittype',{},'notes',{});
    % reactions substructure
    reactionsStruct = struct('name',{},'formula',{},'notes',{},'reversible',{},'fast',{});
    % event assignment substructure
    eventassignmentStruct = struct('variable',{},'formula',{});
    % event substructure
    eventStruct = struct('name',{},'trigger',{},'assignment',eventassignmentStruct,'notes',{});
    % Create IQMstructure
    IQMstructure = struct('name','unnamed_model','notes','','functions',functionsStruct,'states',statesStruct,'algebraic',algebraicStruct,'parameters',parametersStruct,'variables',variablesStruct,'reactions',reactionsStruct,'events',eventStruct,'functionsMATLAB','','inputs',inputsStruct,'outputs',outputsStruct);
    % construct the model object
    model = class(IQMstructure,'IQMmodel');
elseif strcmp('IQMmodel',inputType),
    % copy the model object
    model = iqmInput;
elseif strcmp('IQMstructure',inputType),
    % check if given structure contains inputs and outputs. If not then add these 
    if ~isfield(IQMstructure,'inputs'),
        IQMstructure.inputs  = struct('name',{},'factors',{},'terms',{},'stateindex',{},'parindex',{});
    end
    if ~isfield(IQMstructure,'outputs'),
        IQMstructure.outputs = struct('name',{},'formula',{},'notes',{},'varindex',{});
    end
    % check if the given structure is a IQMstructure (only check the
    % top-level fields)
    checkFields = {'name','notes','functions','states','algebraic','parameters','variables','reactions','events','functionsMATLAB'};
    for k = 1:length(checkFields),
        if ~isfield(IQMstructure,checkFields{k}),
            errorMsg = sprintf('Given structure is not a valid internal IQMmodel structure.');
            error(errorMsg);
        end
    end
    % construct the model object
    model = class(IQMstructure,'IQMmodel');
elseif strcmp('SBMLmodelFile',inputType),
    % check if a file with given filename exists
    [path,filename,ext] = fileparts(filename);
    filename = fullfile(path, [filename '.xml']); 
    test = dir(filename);
    if isempty(test),
        errorMsg = sprintf('SBML model, "%s", does not exist in given folder.', filename);
        error(errorMsg);
    end
    
    % If file exists then import it
    [IQMstructure,errorMsg] = importSBMLIQM(filename,useSBMLnames);
    if nocheck == 0,
        if ~isempty(errorMsg), error(errorMsg);  end
    else
        if ~isempty(errorMsg), warning on; warning(errorMsg);  end
    end
    
    % Remove SBML information if requested
    if stripSBMLinfo,
        for k = 1:length(IQMstructure.states),
            IQMstructure.states(k).type = '';
            IQMstructure.states(k).compartment = '';
            IQMstructure.states(k).unittype = '';
        end
        for k = 1:length(IQMstructure.variables),
            IQMstructure.variables(k).type = '';
            IQMstructure.variables(k).compartment = '';
            IQMstructure.variables(k).unittype = '';
        end
        for k = 1:length(IQMstructure.parameters),
            IQMstructure.parameters(k).type = '';
            IQMstructure.parameters(k).compartment = '';
            IQMstructure.parameters(k).unittype = '';
        end
    end
    
    
    
    % construct the model object
    model = class(IQMstructure,'IQMmodel');
elseif strcmp('TextModelFile',inputType),
    % check if a file with given filename exists
    [path,filename,ext] = fileparts(filename);
    filename = fullfile(path, [filename '.txt']); 
    if ~exist(filename),
        errorMsg = sprintf('TEXT model, "%s", does not exist.', filename);
        error(errorMsg);
    end
    % If file exists then first load it
    modelText = fileread(filename);  
    % then convert it to IQMstructure
    [IQMstructure, errorMsg] = convertTextToModelIQM(modelText);
    % Check if error occurred while importing the SBML model
    if ~isempty(errorMsg),
        error(errorMsg);
    end
    % handle arrays for TEXT models
    IQMstructure = convertTEXTArrayDefIQM(IQMstructure);
    % construct the model object
    model = class(IQMstructure,'IQMmodel');
elseif strcmp('TextBCModelFile',inputType),
    % check if a file with given filename exists
    [path,filename,ext] = fileparts(filename);
    filename = fullfile(path, [filename '.txtbc']); 
    if ~exist(filename),
        errorMsg = sprintf('TEXTBC model, "%s", does not exist.', filename);
        error(errorMsg);
    end
    % If file exists then first load it
    modelText = fileread(filename);
    % then convert it to IQMstructure
    [IQMstructure, errorMsg] = convertTextBCToModelIQM(modelText);
    % Check if error occurred while importing the SBML model
    if ~isempty(errorMsg),
        error(errorMsg);
    end
    % construct the model object
    model = class(IQMstructure,'IQMmodel');
elseif strcmp('ModelAsTextString', inputType),
    % convert model text to IQMstructure
    [IQMstructure, errorMsg] = convertTextToModelIQM(modelText);
    % Check if error occurred while importing the SBML model
    if ~isempty(errorMsg),
        error(errorMsg);
    end
    % construct the model object
    model = class(IQMstructure,'IQMmodel');
elseif strcmp('ModelAsTextBCString', inputType),
    % then convert model text to IQMstructure
    [IQMstructure, errorMsg] = convertTextBCToModelIQM(modelText);
    % Check if error occurred while importing the SBML model
    if ~isempty(errorMsg),
        error(errorMsg);
    end
    % construct the model object
    model = class(IQMstructure,'IQMmodel');
else
    error('Wrong input arguments.');
end
%%%%%%%%%%%%%%%%%%%%%%%%%%%%%%%%%%%%%%%%%%
% Handle inputs and outputs (add info about them to the structure and
% eventually add parameters for inputs)
%%%%%%%%%%%%%%%%%%%%%%%%%%%%%%%%%%%%%%%%%%
model = inputoutputIQMmodelParsingIQM(model);
return

%%%%%%%%%%%%%%%%%%%%%%%%%%%%%%%%%%%%%%%%%%%%%%%%%%%%%%%%%%%%%%%%%%%%%%%%%%
% DELETE WHITESPACES IN STRINGS
%%%%%%%%%%%%%%%%%%%%%%%%%%%%%%%%%%%%%%%%%%%%%%%%%%%%%%%%%%%%%%%%%%%%%%%%%%
% Useful for taking away whitespaces in kineticLaw formulas, as
% seen in some example models
function [outputString] = removeWhiteSpace(inputString)
outputString = strrep(inputString,' ','');
% return
return
```
