## Supplementary material for "A systems biology analysis of adrenergically stimulated adiponectin exocytosis in white adipocytes": All scripts used in this work: IQMstruct.m

```
function [IQMstructure] = IQMstruct(iqm)
% IQMstruct: This function returns the internal data structure
% of an IQMmodel
%
% USAGE:
% ======
% [IQMstructure] = IQMstruct(IQMmodel) 
%
% IQMmodel: IQMmodel 
%
% Output Arguments:
% =================
% IQMstructure: internal data structure of the IQMmodel

% <<<COPYRIGHTSTATEMENT - IQM TOOLS LITE>>>


%%%%%%%%%%%%%%%%%%%%%%%%%%%%%%%%%%%%%%%%%%%%%%%
% VERY SIMPLE FUNCTION
%%%%%%%%%%%%%%%%%%%%%%%%%%%%%%%%%%%%%%%%%%%%%%%
IQMstructure = struct(iqm);
```
