## Supplementary material for "A systems biology analysis of adrenergically stimulated adiponectin exocytosis in white adipocytes": All scripts used in this work: IQMcreateDOSfile.m

```
function IQMcreateDOSfile(varargin)
% IQMcreateDOSfile: creates a *.dos file with the dosing text description
%
% USAGE:
% ======
% [] = IQMcreateDOSfile()         
% [] = IQMcreateDOSfile(filename)         
% [] = IQMcreateDOSfile(dos)         
% [] = IQMcreateDOSfile(dos,filename)
%
% dos: IQMdosing object to convert to a textfile description
% filename: filename for the created textfile 
%
% If dos is undefined, then an empty IQMdosing textfile will be created.
%
% DEFAULT VALUES:
% ===============
% dos: the IQMdosing object to be exported as DOS file. 
% filename: constructed from the IQMdosing object name

% <<<COPYRIGHTSTATEMENT - IQM TOOLS LITE>>>

%%%%%%%%%%%%%%%%%%%%%%%%%%%%%%%%%%%%%%%%%%%%%%%
% PROCESS VARIABLE INPUT ARGUMENTS
%%%%%%%%%%%%%%%%%%%%%%%%%%%%%%%%%%%%%%%%%%%%%%%
if nargin == 0,
    % create empty IQMdosing file with the name "unnamed.dos"
    dos = IQMdosing();
    filename = 'unnamed.dos';
elseif nargin == 1,
    % check if first input argument dosing object or filename
    if isIQMdosing(varargin{1}),
        % dosing object given
        dos = varargin{1};
        % if no filename provided then use the name of the IQMdosing
        % object as filename but remove unwanted characters first
        ds = struct(dos);
        functionName = regexprep(ds.name,'\W','');
        filename = strcat(functionName,'.dos');
    else
        % filename given?
        if ~ischar(varargin{1}),
            error('Wrong input argument.');
        end
        filename = strcat(varargin{1},'.dos');
        dos = IQMdosing();
    end
elseif nargin == 2,
    dos = varargin{1};
    filename = strcat(varargin{2},'.dos');
else
    error('Incorrect number of input arguments.');
end

%%%%%%%%%%%%%%%%%%%%%%%%%%%%%%%%%%%%%%%%%%%%%%%
% CHECK INPUT
%%%%%%%%%%%%%%%%%%%%%%%%%%%%%%%%%%%%%%%%%%%%%%%
if ~isIQMdosing(dos),
    error('No IQMdosing object as first argument.');
end

%%%%%%%%%%%%%%%%%%%%%%%%%%%%%%%%%%%%%%%%%%%%%%%
% CONVERT DOSING TO TEXT
%%%%%%%%%%%%%%%%%%%%%%%%%%%%%%%%%%%%%%%%%%%%%%%
[dosTextStructure] = convertDosToTextIQM(dos);
[completeText] = setPartsToCompleteTextDosIQM(dosTextStructure);

%%%%%%%%%%%%%%%%%%%%%%%%%%%%%%%%%%%%%%%%%%%%%%%
% WRITE THE FILE
%%%%%%%%%%%%%%%%%%%%%%%%%%%%%%%%%%%%%%%%%%%%%%%
fid = fopen(filename,'w');
fwrite(fid,completeText);
fclose(fid);
return
```
