## Supplementary material for "A systems biology analysis of adrenergically stimulated adiponectin exocytosis in white adipocytes": All scripts used in this work: IQMcreateDOSING.m

```
function [dosing] = IQMcreateDOSING(type,dose,time,parametervalue,tlagvalue)
% IQMcreateDOSING: Creates a dosing scheme as desired based on inputs
%
% USAGE:
% ======
% [dosing] = IQMcreateDOSING(type)         
% [dosing] = IQMcreateDOSING(type,dose,time,parametervalue)         
% [dosing] = IQMcreateDOSING(type,dose,time,parametervalue,tlagvalue)         
%
% type:             cell-array with input types in the order as inputs will be defined
%                   possible types: "INFUSION", "ABSORPTION0", "ABSORPTION1", "BOLUS"
% dose:             cell-array with dose vectors for each defined input - if scalar then same dose will be used for each dosing time
% time:             cell-array with time vectors for each defined input
% parametervalue:   cell-array with TINF, ka, TK0 values - if scalar then
%                   same value assumed for each dosing time. Empty cell entry if bolus.
% tlagvalue:        cell-array with lag time for each input definition
%
% dosing:           produced dosing scheme
%
% If only type is provided, then all other entries are set to "zero".

% <<<COPYRIGHTSTATEMENT - IQM TOOLS LITE>>>

% Create empty dosing scheme
ds = struct(IQMdosing);

% Check type first (needed in second step)
if ~iscell(type), type = {type}; end

% Variable input arguments
if nargin == 1,
    dose            = cell(1,length(type)); dose(1:end) = {0};
    time            = cell(1,length(type)); time(1:end) = {0};
    parametervalue  = cell(1,length(type)); parametervalue(strmatchIQM('INFUSION',type)) = {1}; parametervalue(strmatchIQM('ABSORPTION0',type)) = {1}; parametervalue(strmatchIQM('ABSORPTION1',type)) = {1};
    tlagvalue       = [];
elseif nargin == 4,
    tlagvalue = [];
elseif nargin == 5,
    if ~iscell(tlagvalue), tlagvalue = {tlagvalue}; end
else
    error('Incorrect number of input arguments.');
end

%%%%%%%%%%%%%%%%%%%%%%%%%%%%%%%%%%%%%%%%%%%%%%%
% Check inputs and convert to cells if needed
%%%%%%%%%%%%%%%%%%%%%%%%%%%%%%%%%%%%%%%%%%%%%%%
if ~iscell(dose), dose = {dose}; end
if ~iscell(time), time = {time}; end
if ~iscell(parametervalue), parametervalue = {parametervalue}; end

%%%%%%%%%%%%%%%%%%%%%%%%%%%%%%%%%%%%%%%%%%%%%%%
% Update name fields
%%%%%%%%%%%%%%%%%%%%%%%%%%%%%%%%%%%%%%%%%%%%%%%
for k=1:length(type),
    ds.inputs(k).name = sprintf('INPUT%d',k);
end

%%%%%%%%%%%%%%%%%%%%%%%%%%%%%%%%%%%%%%%%%%%%%%%
% Update type fields
%%%%%%%%%%%%%%%%%%%%%%%%%%%%%%%%%%%%%%%%%%%%%%%
for k=1:length(type),
    ds.inputs(k).type = type{k};
end

%%%%%%%%%%%%%%%%%%%%%%%%%%%%%%%%%%%%%%%%%%%%%%%
% Update time fields
%%%%%%%%%%%%%%%%%%%%%%%%%%%%%%%%%%%%%%%%%%%%%%%
if length(time) ~= length(type),
    error('Number of entries in time cell-array different from number in type cell-array');
end
for k=1:length(time),
    ds.inputs(k).time = time{k};
end

%%%%%%%%%%%%%%%%%%%%%%%%%%%%%%%%%%%%%%%%%%%%%%%
% Update dose fields
%%%%%%%%%%%%%%%%%%%%%%%%%%%%%%%%%%%%%%%%%%%%%%%
if length(dose) ~= length(type),
    error('Number of entries in dose cell-array different from number in type cell-array');
end
for k=1:length(dose),
    dosek = dose{k};
    if length(dosek) == 1,
        ds.inputs(k).D = dosek*ones(1,length(ds.inputs(k).time));
    else
        if length(dosek) ~= length(ds.inputs(k).time),
            error('Different lengths of time and dose vectors for input %d',k);
        else
            ds.inputs(k).D = dosek;
        end
    end
end

%%%%%%%%%%%%%%%%%%%%%%%%%%%%%%%%%%%%%%%%%%%%%%%
% Update parameters fields
% Only handle if type not BOLUS
%%%%%%%%%%%%%%%%%%%%%%%%%%%%%%%%%%%%%%%%%%%%%%%
if length(parametervalue) ~= length(type),
    error('Number of entries in parametervalue cell-array different from number in type cell-array');
end
for k=1:length(type),
    if ~strcmp(type{k},'BOLUS'),
        pvk = parametervalue{k};
        if strcmp(type{k},'INFUSION'),
            ds.inputs(k).parameters.name = 'Tinf';
        elseif strcmp(type{k},'ABSORPTION1'),
            ds.inputs(k).parameters.name = 'ka';
        elseif strcmp(type{k},'ABSORPTION0'),
            ds.inputs(k).parameters.name = 'Tk0';
        end            
        ds.inputs(k).parameters.notes = '';
        
        if length(pvk) == 1,
            ds.inputs(k).parameters.value = pvk*ones(1,length(ds.inputs(k).time));
        else
            if length(pvk) ~= length(ds.inputs(k).time),
                error('Different lengths of parameter values and dose vectors for input %d',k);
            else
                ds.inputs(k).parameters.value = pvk;
            end
        end
    end
end

%%%%%%%%%%%%%%%%%%%%%%%%%%%%%%%%%%%%%%%%%%%%%%%
% Update Tlag fields
%%%%%%%%%%%%%%%%%%%%%%%%%%%%%%%%%%%%%%%%%%%%%%%
if ~isempty(tlagvalue),
    if length(tlagvalue) ~= length(type),
        error('Number of entries in tlagvalue cell-array different from number in type cell-array');
    end
    for k=1:length(tlagvalue),
        ds.inputs(k).Tlag = tlagvalue{k};
    end
% else
%     for k=1:length(tlagvalue),
%         ds.inputs(k).Tlag = 0;
%     end
end    

%%%%%%%%%%%%%%%%%%%%%%%%%%%%%%%%%%%%%%%%%%%%%%%
% Create dosing scheme
%%%%%%%%%%%%%%%%%%%%%%%%%%%%%%%%%%%%%%%%%%%%%%%
dosing = IQMdosing(ds);
```
