## Supplementary material for "A systems biology analysis of adrenergically stimulated adiponectin exocytosis in white adipocytes": All scripts used in this work: IQMdoseupdatevalue.m

```
function [dosingchanged] = IQMdoseupdatevalue(dosing,inputname,value)
% IQMdoseupdatevalue: Allows to update the dosing amount for a given
%   input, defined in an IQMdosing object.
%
% USAGE:
% ======
% [dosingchanged] = IQMdoseupdatevalue(dosing,inputname,value)         
%
% dosing: IQMdosing object 
% inputname: name of the input for which to change the dosing amount
% value: value to set the dosing amount to (only scalar value allowed, used
%   for all dosing instances in case of multiple doses)
%
% Output Arguments:
% =================
% dosingchanged: updated IQMdosing object with changed dosing amount

% <<<COPYRIGHTSTATEMENT - IQM TOOLS LITE>>>

%%%%%%%%%%%%%%%%%%%%%%%%%%%%%%%%%%%%%%%%%%%%%%%
% Get dosing structure
%%%%%%%%%%%%%%%%%%%%%%%%%%%%%%%%%%%%%%%%%%%%%%%
ds = struct(dosing);

%%%%%%%%%%%%%%%%%%%%%%%%%%%%%%%%%%%%%%%%%%%%%%%
% Find input name
%%%%%%%%%%%%%%%%%%%%%%%%%%%%%%%%%%%%%%%%%%%%%%%
index = strmatchIQM(inputname,{ds.inputs.name},'exact');

%%%%%%%%%%%%%%%%%%%%%%%%%%%%%%%%%%%%%%%%%%%%%%%
% Change Dosing amount if index found
%%%%%%%%%%%%%%%%%%%%%%%%%%%%%%%%%%%%%%%%%%%%%%%
if isempty(index),
    disp(['Input ' inputname ' does not exist in the dosing schedule.']);
else
    % update dosing amount with new value
    ds.inputs(index).D = value*ones(1,length(ds.inputs(index).time));
end

%%%%%%%%%%%%%%%%%%%%%%%%%%%%%%%%%%%%%%%%%%%%%%%
% DONE => Construct output
%%%%%%%%%%%%%%%%%%%%%%%%%%%%%%%%%%%%%%%%%%%%%%%
dosingchanged = IQMdosing(ds);
```
