## Supplementary material for "A systems biology analysis of adrenergically stimulated adiponectin exocytosis in white adipocytes": All scripts used in this work: IQMmergemoddos.m

```
function [simmodel] = IQMmergemoddos(model,dosing)
% IQMmergemoddos: Based on a model and dosing object, a new IQMmodel is
% generated, that implements the defined dosings. Multiple dosing schedules
% are realized by updating the parameters of subsequent dosings using
% events.
%
% This function is useful to simulate single dosing schedules. However, if
% you want to run trial simulations, which change the dosing amounts (or
% other things) between simulations, this function is not the best way to
% go, since you would need to apply these changes to the dosing schedule,
% run this function, recompile your model, etc.
%
% USAGE:
% ======
% [simmodel] = IQMmergemoddos(model,dosing) 
%
% model: IQMmodel
% dosing: IQMdosing object
%
% Output Arguments:
% =================
% simmodel: Simulation model, implementing the dosing scheme, defined in
%   the model and the IQMdosing object.

% <<<COPYRIGHTSTATEMENT - IQM TOOLS LITE>>>

%%%%%%%%%%%%%%%%%%%%%%%%%%%%%%%%%%%%%%%%%%%%%%%
% Check input
%%%%%%%%%%%%%%%%%%%%%%%%%%%%%%%%%%%%%%%%%%%%%%%
if ~isIQMmodel(model),
    error('First input argument is not an IQMmodel.');
end
if ~isIQMdosing(dosing),
    error('Second input argument is not an IQMdosing object.');
end

%%%%%%%%%%%%%%%%%%%%%%%%%%%%%%%%%%%%%%%%%%%%%%%
% Get model augmented with dosing related 
% components and the experiment description,
% implementing the dosing applications
%%%%%%%%%%%%%%%%%%%%%%%%%%%%%%%%%%%%%%%%%%%%%%%
[moddos,experiment] = mergemoddosIQM(model,dosing);

%%%%%%%%%%%%%%%%%%%%%%%%%%%%%%%%%%%%%%%%%%%%%%%
% Combine new model with experiment to get the
% simulation model
%%%%%%%%%%%%%%%%%%%%%%%%%%%%%%%%%%%%%%%%%%%%%%%
simmodel = IQMmergemodexp(moddos,experiment);
```
