## Supplementary material for "A systems biology analysis of adrenergically stimulated adiponectin exocytosis in white adipocytes": All scripts used in this work: mergemoddosIQM.m

```
function [moddos,expdos] = mergemoddosIQM(model,dosing)
% mergemoddosIQM: This function takes an IQMmodel and an IQMdosing
% scheme as inputs. It adds necessary elements to the model that are
% required to implement the different dosing inputs (bolus, infusion,
% absorption0, absorption1) for simulation purposes. MLXTRAN and
% NONMEM will not be handled using this function. 
%
% The dose parameters are set to zero, independently of the dosing values
% given in the dosing settings.
%
% The second output argument "expdos" is an experiment description,
% which contains all state-events that are necessary to realize the dosing
% applications defined in the dosing object. 
%
% USAGE:
% ======
% [moddos,expdos] = mergemoddosIQM(model,dosing) 
%
% model: IQMmodel
% dosing: IQMdosing object
%
% Output Arguments:
% =================
% moddos: IQMmodel with added components to implement the different types of
%  inputs.
% expdos: IQMexperiment containing all information to simulate the dosing
%  schedule, defined in the IQMdosing object.

% <<<COPYRIGHTSTATEMENT - IQM TOOLS LITE>>>

%%%%%%%%%%%%%%%%%%%%%%%%%%%%%%%%%%%%%%%%%%%%%%%
% Check input
%%%%%%%%%%%%%%%%%%%%%%%%%%%%%%%%%%%%%%%%%%%%%%%
if ~isIQMmodel(model),
    error('First input argument is not an IQMmodel.');
end
if ~isIQMdosing(dosing),
    error('Second input argument is not an IQMdosing object.');
end

%%%%%%%%%%%%%%%%%%%%%%%%%%%%%%%%%%%%%%%%%%%%%%%
% Get model, experiment and dosing structures
%%%%%%%%%%%%%%%%%%%%%%%%%%%%%%%%%%%%%%%%%%%%%%%
ms = struct(model);
ds = struct(dosing);
mds = ms;               % initialize new model structure
eds = struct(IQMexperiment());
eds.name = ds.name;
eds.notes = ds.notes;

%%%%%%%%%%%%%%%%%%%%%%%%%%%%%%%%%%%%%%%%%%%%%%%
% Get the inputinfo structure
% This structure contains now all necessary
% information about all inputs that need to be 
% implemented.
%%%%%%%%%%%%%%%%%%%%%%%%%%%%%%%%%%%%%%%%%%%%%%%    
inputinfo = getmoddosinputinfoIQM(model,dosing);

%%%%%%%%%%%%%%%%%%%%%%%%%%%%%%%%%%%%%%%%%%%%%%%
% Run through the inputinfo struct and modify the model
% Additionally update the experiment structure ...
%%%%%%%%%%%%%%%%%%%%%%%%%%%%%%%%%%%%%%%%%%%%%%%
for k=1:length(inputinfo),
    % Handle the different types differently
    switch inputinfo(k).type,
        case 'BOLUS',
            [mds,eds] = handleBolus(ms,mds,eds,inputinfo(k));
        case 'INFUSION',
            [mds,eds] = handleInfusion(ms,mds,eds,inputinfo(k));
        case 'ABSORPTION1',
            [mds,eds] = handleAbsorption1(ms,mds,eds,inputinfo(k));
        case 'ABSORPTION0',
            [mds,eds] = handleAbsorption0(ms,mds,eds,inputinfo(k));
    end
end

%%%%%%%%%%%%%%%%%%%%%%%%%%%%%%%%%%%%%%%%%%%%%%%
% Delete the inputs implemented into the model
% from the input field structure in the model
%%%%%%%%%%%%%%%%%%%%%%%%%%%%%%%%%%%%%%%%%%%%%%%
removeInputIndex = [];
for k=1:length(mds.inputs),
    iname = mds.inputs(k).name;
    % check if the name exists as parameter in the model (then dont remove it)
    if isempty(strmatchIQM(iname,{mds.parameters.name},'exact')),
        % input parameter removed => remove input definition
        removeInputIndex(end+1) = k;
    end
end
if ~isempty(removeInputIndex),
    mds.inputs(removeInputIndex) = [];
end

%%%%%%%%%%%%%%%%%%%%%%%%%%%%%%%%%%%%%%%%%%%%%%%
% Construct output model and experimet
%%%%%%%%%%%%%%%%%%%%%%%%%%%%%%%%%%%%%%%%%%%%%%%
moddos = IQMmodel(mds);
expdos = IQMexperiment(eds);

%%%%%%%%%%%%%%%%%%%%%%%%%%%%%%%%%%%%%%%%%%%%%%%
% Return
%%%%%%%%%%%%%%%%%%%%%%%%%%%%%%%%%%%%%%%%%%%%%%%
return

%%%%%%%%%%%%%%%%%%%%%%%%%%%%%%%%%%%%%%%%%%%%%%%%%%%%%%%%%%%%%%%%%%%%%%%%%%%
% handle BOLUS input
%%%%%%%%%%%%%%%%%%%%%%%%%%%%%%%%%%%%%%%%%%%%%%%%%%%%%%%%%%%%%%%%%%%%%%%%%%%
% also add dosing time and amount names to the inputinfo structure
function [mds,eds] = handleBolus(ms,mds,eds,inputinfo)
% Get input name
iname = inputinfo.name;
% Cycle through all input different input terms
for k=1:length(inputinfo.terms),
    % a) Determine the input variable name and other names to be used in the model
    ivarname = getoutname(lower(iname), length(inputinfo.terms), k);
    iDosename = ['Dose' lower(iname)];
    iTimename = ['Time' lower(iname)];
    iBolusTimename = ['DeltaT' lower(iname)];
    iTlagname = ['Tlag' lower(iname)];
    % b) Replace ODE INPUT definition with input variable
    term = inputinfo.terms{k};
    stateindex = inputinfo.stateindex(k);
    ODE = mds.states(stateindex).ODE;
    ODE = strrep(ODE,term,['+' ivarname]);
    mds.states(stateindex).ODE = ODE;
    % b2) Handle Tlag 
    if k==1,    % only add Tlag parameters once!
        % Tlag (only if defined in dosing schedule)
        % If defined numerically, then as parameter, non-numerically as
        % variable ...
        if ~isempty(inputinfo.Tlag) && isnumeric(inputinfo.Tlag),
            % Add as parameter
            mds.parameters(end+1).name = iTlagname;
            mds.parameters(end).value = inputinfo.Tlag;
            mds.parameters(end).type = '';
            mds.parameters(end).compartment = '';
            mds.parameters(end).unittype = '';
            mds.parameters(end).notes = sprintf('%s Dosing lag time, used in variable ''%s'' (BOLUS).',inputinfo.TlagNotes,ivarname);
        elseif ~isempty(inputinfo.Tlag) && ischar(inputinfo.Tlag),
            % Add as variable
            mds.variables(end+1).name = iTlagname;
            mds.variables(end).formula = inputinfo.Tlag;
            mds.variables(end).type = '';
            mds.variables(end).compartment = '';
            mds.variables(end).unittype = '';
            mds.variables(end).notes = sprintf('%s Dosing lag time, used in variable ''%s'' (BOLUS).',inputinfo.TlagNotes,ivarname);
        end
    end
    % c) Create the input variable (differently, depending if Tlag is
    % defined or not)
    if ~isempty(inputinfo.Tlag),    
        vformula = sprintf('%s*%s/%s * piecewiseIQM(1,andIQM(ge(time,%s+%s),lt(time,%s+%s+%s)),0)',inputinfo.factors{k},iDosename,iBolusTimename,iTimename,iTlagname,iTimename,iTlagname,iBolusTimename);
    else
        vformula = sprintf('%s*%s/%s * piecewiseIQM(1,andIQM(ge(time,%s),lt(time,%s+%s)),0)',inputinfo.factors{k},iDosename,iBolusTimename,iTimename,iTimename,iBolusTimename);
    end        
    % d) Add the variable to the moddos model
    mds.variables(end+1).name = ivarname;
    mds.variables(end).formula = vformula;
    mds.variables(end).type = '';
    mds.variables(end).compartment = '';
    mds.variables(end).unittype = '';
    mds.variables(end).notes = sprintf(' Input variable realizing ''%s'' on state ''%s'' (BOLUS).',iname,ms.states(stateindex).name);
    % e) Remove the INPUT* parameter (will change the parindex fields but
    % doesn't matter, since this input parameters are not used anymore ...
    % if a dosing input is defined for this INPUT* element).
    index = strmatchIQM(iname,{mds.parameters.name},'exact');
    mds.parameters(index) = [];
    % f) Need to define an initial set of parameters used in the vformula.
    % Single dosing events can be implemented in this way. In the case of
    % multiple dosing events only the first will be implemented in the
    % model. The remaining ones will be handled using an outside loop in a
    % higher level function.
    if k==1,    % only add Dose and Time parameters once! (also the bolus deltaT)
        % Dose
        mds.parameters(end+1).name = iDosename;
        mds.parameters(end).value = 0; %inputinfo.D(1);
        mds.parameters(end).type = '';
        mds.parameters(end).compartment = '';
        mds.parameters(end).unittype = '';
        mds.parameters(end).notes = sprintf(' Dose amount for INPUT ''%s'' (BOLUS).', iname);
        % Time
        mds.parameters(end+1).name = iTimename;
        mds.parameters(end).value = inputinfo.time(1);
        mds.parameters(end).type = '';
        mds.parameters(end).compartment = '';
        mds.parameters(end).unittype = '';
        mds.parameters(end).notes = sprintf(' Dosing instant for INPUT ''%s'' (BOLUS).', iname);
        % iBolusTimename
        mds.parameters(end+1).name = iBolusTimename;
        mds.parameters(end).value = 0.0001; % Assume 1e-4 of time units (should be fast enough to realistically implement a BOLUS)
        mds.parameters(end).type = '';
        mds.parameters(end).compartment = '';
        mds.parameters(end).unittype = '';
        mds.parameters(end).notes = sprintf(' Very small dosing duration for INPUT ''%s'' (BOLUS).', iname);
        % Add the necessary information to the experiment structure (only time
        % and dosing amounts can change between subsequent dosing instants
[eds] = buildexpstruct(eds,iDosename,iTimename,iTlagname,inputinfo,iname,'',[]);
    end
end
return

%%%%%%%%%%%%%%%%%%%%%%%%%%%%%%%%%%%%%%%%%%%%%%%%%%%%%%%%%%%%%%%%%%%%%%%%%%%
% handle Infusion input
%%%%%%%%%%%%%%%%%%%%%%%%%%%%%%%%%%%%%%%%%%%%%%%%%%%%%%%%%%%%%%%%%%%%%%%%%%%
% also add dosing time and amount names to the inputinfo structure
function [mds,eds] = handleInfusion(ms,mds,eds,inputinfo)
% Get input name
iname = inputinfo.name;
% Cycle through all input different input terms
for k=1:length(inputinfo.terms),
    % Check if infusion time or infusion rate
    paramname = inputinfo.parameters.name;
    if strcmp(paramname,'Tinf'),
        TinfFlag = 1;
    else
        TinfFlag = 0;
    end
    % a) Determine the input variable name and other names to be used in the model
    ivarname = getoutname(lower(iname), length(inputinfo.terms), k);
    iDosename = ['Dose' lower(iname)];
    if TinfFlag,
        iParamNameInfusion = ['Tinf' lower(iname)];
    else
        iParamNameInfusion = ['Rate' lower(iname)];
    end
    iTimename = ['Time' lower(iname)];
    iTlagname = ['Tlag' lower(iname)];
    % b) Replace ODE INPUT definition with input variable
    term = inputinfo.terms{k};
    stateindex = inputinfo.stateindex(k);
    ODE = mds.states(stateindex).ODE;
    ODE = strrep(ODE,term,['+' ivarname]);
    mds.states(stateindex).ODE = ODE;
    % b2) Handle Tlag 
    if k==1,    % only add Tlag parameters once!
        % Tlag (only if defined in dosing schedule)
        % If defined numerically, then as parameter, non-numerically as
        % variable ...
        if ~isempty(inputinfo.Tlag) && isnumeric(inputinfo.Tlag),
            % Add as parameter
            mds.parameters(end+1).name = iTlagname;
            mds.parameters(end).value = inputinfo.Tlag;
            mds.parameters(end).type = '';
            mds.parameters(end).compartment = '';
            mds.parameters(end).unittype = '';
            mds.parameters(end).notes = sprintf('%s Dosing lag time, used in variable ''%s'' (INFUSION).',inputinfo.TlagNotes,ivarname);
        elseif ~isempty(inputinfo.Tlag) && ischar(inputinfo.Tlag),
            % Add as variable
            mds.variables(end+1).name = iTlagname;
            mds.variables(end).formula = inputinfo.Tlag;
            mds.variables(end).type = '';
            mds.variables(end).compartment = '';
            mds.variables(end).unittype = '';
            mds.variables(end).notes = sprintf('%s Dosing lag time, used in variable ''%s'' (INFUSION).',inputinfo.TlagNotes,ivarname);
        end
    end
    % c) Create the input variable (differently, depending if Tlag is
    % defined or not)
    if TinfFlag,
        % Implement infusion time
        if ~isempty(inputinfo.Tlag),
            vformula = sprintf('%s*%s/%s * piecewiseIQM(1,andIQM(ge(time,%s+%s),lt(time,%s+%s+%s)),0)',inputinfo.factors{k},iDosename,iParamNameInfusion,iTimename,iTlagname,iTimename,iTlagname,iParamNameInfusion);
        else
            vformula = sprintf('%s*%s/%s * piecewiseIQM(1,andIQM(ge(time,%s),lt(time,%s+%s)),0)',inputinfo.factors{k},iDosename,iParamNameInfusion,iTimename,iTimename,iParamNameInfusion);
        end
    else
        % Implement infusion Rate
        if ~isempty(inputinfo.Tlag),
            vformula = sprintf('%s*%s * piecewiseIQM(1,andIQM(ge(time,%s+%s),lt(time,%s+%s+%s/%s)),0)',inputinfo.factors{k},iParamNameInfusion,iTimename,iTlagname,iTimename,iTlagname,iDosename,iParamNameInfusion);
        else
            vformula = sprintf('%s*%s * piecewiseIQM(1,andIQM(ge(time,%s),lt(time,%s+%s/%s)),0)',inputinfo.factors{k},iParamNameInfusion,iTimename,iTimename,iDosename,iParamNameInfusion);
        end
    end
    % d) Add the variable to the moddos model
    mds.variables(end+1).name = ivarname;
    mds.variables(end).formula = vformula;
    mds.variables(end).type = '';
    mds.variables(end).compartment = '';
    mds.variables(end).unittype = '';
    mds.variables(end).notes = sprintf(' Input variable realizing ''%s'' on state ''%s'' (INFUSION).',iname,ms.states(stateindex).name);
    % e) Remove the INPUT* parameter (will change the parindex fields but
    % doesn't matter, since this input parameters are not used anymore ...
    % if a dosing input is defined for this INPUT* element).
    index = strmatchIQM(iname,{mds.parameters.name},'exact');
    mds.parameters(index) = [];
    % f) Need to define an initial set of parameters used in the vformula.
    % Single dosing events can be implemented in this way. In the case of
    % multiple dosing events the parameters will be changed using events.
    if k==1,    % only add Dose and Time parameters once!
        % Dose
        mds.parameters(end+1).name = iDosename;
        mds.parameters(end).value = 0;%inputinfo.D(1);
        mds.parameters(end).type = '';
        mds.parameters(end).compartment = '';
        mds.parameters(end).unittype = '';
        mds.parameters(end).notes = sprintf(' Dose amount for INPUT ''%s'' (INFUSION).', iname);
        % Time
        mds.parameters(end+1).name = iTimename;
        mds.parameters(end).value = inputinfo.time(1);
        mds.parameters(end).type = '';
        mds.parameters(end).compartment = '';
        mds.parameters(end).unittype = '';
        mds.parameters(end).notes = sprintf(' Dosing instant for INPUT ''%s'' (INFUSION).', iname);
        % Rate
        mds.parameters(end+1).name = iParamNameInfusion;
        mds.parameters(end).value = inputinfo.parameters(1).value(1);
        mds.parameters(end).type = '';
        mds.parameters(end).compartment = '';
        mds.parameters(end).unittype = '';
        if TinfFlag,
            mds.parameters(end).notes = sprintf(' Dosing infusion time, used in variable ''%s'' (INFUSION).', ivarname);
        else
            mds.parameters(end).notes = sprintf(' Dosing infusion rate, used in variable ''%s'' (INFUSION).', ivarname);
        end
        % Add the necessary information to the experiment structure (only time
        % and dosing amounts can change between subsequent dosing instants
[eds] = buildexpstruct(eds,iDosename,iTimename,iTlagname,inputinfo,iname,iParamNameInfusion,inputinfo.parameters(1).value);
%         % Additionally set the Rate/Tinf parameter in the experiment
%         eds = addparamicsetting(eds,iParamNameInfusion,num2str(inputinfo.parameters(1).value),0,'');
    end
end
return

%%%%%%%%%%%%%%%%%%%%%%%%%%%%%%%%%%%%%%%%%%%%%%%%%%%%%%%%%%%%%%%%%%%%%%%%%%%
% handle Absorption1 input
%%%%%%%%%%%%%%%%%%%%%%%%%%%%%%%%%%%%%%%%%%%%%%%%%%%%%%%%%%%%%%%%%%%%%%%%%%%
% also add dosing time and amount names to the inputinfo structure
function [mds,eds] = handleAbsorption1(ms,mds,eds,inputinfo)
% Get input name
iname = inputinfo.name;
% Cycle through all input different input terms
for k=1:length(inputinfo.terms),
    % a) Determine the input variable name and other names to be used in the model
    ivarname = getoutname(lower(iname), length(inputinfo.terms), k);
    iCompname = ['Comp_' ivarname];
    iabsorptvarname =['vAbsorption_' ivarname];
    iDosename = ['Dose' lower(iname)];
    iTimename = ['Time' lower(iname)];
    iBolusTimename = ['DeltaT' lower(iname)];    

    iKaname   = ['ka' lower(iname)];
    iTlagname = ['Tlag' lower(iname)];
    
    % b) Replace ODE INPUT definition with ABSORPTION input variable
    % (difference to all the other dosing application types).
    term = inputinfo.terms{k};
    stateindex = inputinfo.stateindex(k);
    ODE = mds.states(stateindex).ODE;
    ODE = strrep(ODE,term,['+' iabsorptvarname]);
    mds.states(stateindex).ODE = ODE;
    % b2) Handle Tlag 
    if k==1,    % only add Tlag parameters once!
        % Tlag (only if defined in dosing schedule)
        % If defined numerically, then as parameter, non-numerically as
        % variable ...
        if ~isempty(inputinfo.Tlag) && isnumeric(inputinfo.Tlag),
            % Add as parameter
            mds.parameters(end+1).name = iTlagname;
            mds.parameters(end).value = inputinfo.Tlag;
            mds.parameters(end).type = '';
            mds.parameters(end).compartment = '';
            mds.parameters(end).unittype = '';
            mds.parameters(end).notes = sprintf('%s Dosing lag time, used in variable ''%s'' (ABSORPTION1).',inputinfo.TlagNotes,ivarname);
        elseif ~isempty(inputinfo.Tlag) && ischar(inputinfo.Tlag),
            % Add as variable
            mds.variables(end+1).name = iTlagname;
            mds.variables(end).formula = inputinfo.Tlag;
            mds.variables(end).type = '';
            mds.variables(end).compartment = '';
            mds.variables(end).unittype = '';
            mds.variables(end).notes = sprintf('%s Dosing lag time, used in variable ''%s'' (ABSORPTION1).',inputinfo.TlagNotes,ivarname);
        end
    end
    % c) Add a dosing compartment for the ABSORPTION1 input
    mds.states(end+1).name = iCompname;
    mds.states(end).initialCondition = 0;
    mds.states(end).ODE = [ivarname '-' iabsorptvarname];
    mds.states(end).type = '';
    mds.states(end).compartment = '';
    mds.states(end).unittype = '';
    mds.states(end).notes = sprintf(' Dosing compartment for input ''%s'' (ABSORPTION1).',iname);
    % d) Add a variable to define the absorption equation
    mds.variables(end+1).name = iabsorptvarname;
    mds.variables(end).formula = [iKaname '*' iCompname];
    mds.variables(end).type = '';
    mds.variables(end).compartment = '';
    mds.variables(end).unittype = '';
    mds.variables(end).notes = sprintf(' Absorption rate for input ''%s'' (ABSORPTION1).',iname);
    % e) Create the input variable for application of the bolus to the
    % dosing compartment (use same approach as for BOLUS applications, this
    % means, a small application time of 0.0001).
    if ~isempty(inputinfo.Tlag),    
        vformula = sprintf('%s*%s/%s * piecewiseIQM(1,andIQM(ge(time,%s+%s),lt(time,%s+%s+%s)),0)',inputinfo.factors{k},iDosename,iBolusTimename,iTimename,iTlagname,iTimename,iTlagname,iBolusTimename);
    else
        vformula = sprintf('%s*%s/%s * piecewiseIQM(1,andIQM(ge(time,%s),lt(time,%s+%s)),0)',inputinfo.factors{k},iDosename,iBolusTimename,iTimename,iTimename,iBolusTimename);
    end        
    % f) Add the variable to the moddos model
    mds.variables(end+1).name = ivarname;
    mds.variables(end).formula = vformula;
    mds.variables(end).type = '';
    mds.variables(end).compartment = '';
    mds.variables(end).unittype = '';
    mds.variables(end).notes = sprintf(' Input variable realizing a BOLUS on state ''%s'', implementing input ''%s'' (ABSORPTION1).',iCompname,iname);
    % f) Remove the INPUT* parameter (will change the parindex fields but
    % doesn't matter, since this input parameters are not used anymore ...
    % if a dosing input is defined for this INPUT* element).
    index = strmatchIQM(iname,{mds.parameters.name},'exact');
    mds.parameters(index) = [];
    % f) Need to define an initial set of parameters used in the vformula.
    % Single dosing events can be implemented in this way. In the case of
    % multiple dosing events only the first will be implemented in the
    % model. The remaining ones will be handled using an outside loop in a
    % higher level function.
    if k==1,    % only add Dose and Time parameters once! also bolus deltaT
        % Dose
        mds.parameters(end+1).name = iDosename;
        mds.parameters(end).value = 0;%inputinfo.D(1);
        mds.parameters(end).type = '';
        mds.parameters(end).compartment = '';
        mds.parameters(end).unittype = '';
        mds.parameters(end).notes = sprintf(' Dose amount for INPUT ''%s'' (ABSORPTION1).', iname);
        % Time
        mds.parameters(end+1).name = iTimename;
        mds.parameters(end).value = inputinfo.time(1);
        mds.parameters(end).type = '';
        mds.parameters(end).compartment = '';
        mds.parameters(end).unittype = '';
        mds.parameters(end).notes = sprintf(' Dosing instant for INPUT ''%s'' (ABSORPTION1).', iname);
        % iBolusTimename
        mds.parameters(end+1).name = iBolusTimename;
        mds.parameters(end).value = 0.0001; % Assume 1e-4 of time units (should be fast enough to realistically implement a BOLUS)
        mds.parameters(end).type = '';
        mds.parameters(end).compartment = '';
        mds.parameters(end).unittype = '';
        mds.parameters(end).notes = sprintf(' Very small dosing duration (implementing a BOLUS into dosing compartment) for INPUT ''%s'' (ABSORPTION1).', iname);
        % Ka
        mds.parameters(end+1).name = iKaname;
        mds.parameters(end).value = inputinfo.parameters(1).value(1);
        mds.parameters(end).type = '';
        mds.parameters(end).compartment = '';
        mds.parameters(end).unittype = '';
        mds.parameters(end).notes = sprintf(' Dose absorption rate constant, used in variable ''%s'' (ABSORPTION1).', iabsorptvarname);
        % Add the necessary information to the experiment structure (only time
        % and dosing amounts can change between subsequent dosing instants
[eds] = buildexpstruct(eds,iDosename,iTimename,iTlagname,inputinfo,iname,iKaname,inputinfo.parameters(1).value);
%         % Additionally set the ka rate parameters in the experiment
%         eds = addparamicsetting(eds,iKaname,num2str(inputinfo.parameters(1).value),0,'');
    end
end
return

%%%%%%%%%%%%%%%%%%%%%%%%%%%%%%%%%%%%%%%%%%%%%%%%%%%%%%%%%%%%%%%%%%%%%%%%%%%
% handle Absorption0 input
%%%%%%%%%%%%%%%%%%%%%%%%%%%%%%%%%%%%%%%%%%%%%%%%%%%%%%%%%%%%%%%%%%%%%%%%%%%
% also add dosing time and amount names to the inputinfo structure
function [mds,eds] = handleAbsorption0(ms,mds,eds,inputinfo)
% Get input name
iname = inputinfo.name;
% Cycle through all input different input terms
for k=1:length(inputinfo.terms),
    % a) Determine the input variable name and other names to be used in the model
    ivarname = getoutname(lower(iname), length(inputinfo.terms), k);
    iDosename = ['Dose' lower(iname)];
    iTimename = ['Time' lower(iname)];
    iTk0name = ['Tk0' lower(iname)];
    iTlagname = ['Tlag' lower(iname)];
    % b) Replace ODE INPUT definition with input variable
    term = inputinfo.terms{k};
    stateindex = inputinfo.stateindex(k);
    ODE = mds.states(stateindex).ODE;
    ODE = strrep(ODE,term,['+' ivarname]);
    mds.states(stateindex).ODE = ODE;
    % b2) Handle Tlag 
    if k==1,    % only add Tlag parameters once!
        % Tlag (only if defined in dosing schedule)
        % If defined numerically, then as parameter, non-numerically as
        % variable ...
        if ~isempty(inputinfo.Tlag) && isnumeric(inputinfo.Tlag),
            % Add as parameter
            mds.parameters(end+1).name = iTlagname;
            mds.parameters(end).value = inputinfo.Tlag;
            mds.parameters(end).type = '';
            mds.parameters(end).compartment = '';
            mds.parameters(end).unittype = '';
            mds.parameters(end).notes = sprintf('%s Dosing lag time, used in variable ''%s'' (ABSORPTION0).',inputinfo.TlagNotes,ivarname);
        elseif ~isempty(inputinfo.Tlag) && ischar(inputinfo.Tlag),
            % Add as variable
            mds.variables(end+1).name = iTlagname;
            mds.variables(end).formula = inputinfo.Tlag;
            mds.variables(end).type = '';
            mds.variables(end).compartment = '';
            mds.variables(end).unittype = '';
            mds.variables(end).notes = sprintf('%s Dosing lag time, used in variable ''%s'' (ABSORPTION0).',inputinfo.TlagNotes,ivarname);
        end
    end
    % c) Create the input variable (differently, depending if Tlag is
    % defined or not)
    if ~isempty(inputinfo.Tlag),    
        vformula = sprintf('%s*%s/%s * piecewiseIQM(1,andIQM(ge(time,%s+%s),lt(time,%s+%s+%s)),0)',inputinfo.factors{k},iDosename,iTk0name,iTimename,iTlagname,iTimename,iTlagname,iTk0name);
    else
        vformula = sprintf('%s*%s/%s * piecewiseIQM(1,andIQM(ge(time,%s),lt(time,%s+%s)),0)',inputinfo.factors{k},iDosename,iTk0name,iTimename,iTimename,iTk0name);
    end        
    % d) Add the variable to the moddos model
    mds.variables(end+1).name = ivarname;
    mds.variables(end).formula = vformula;
    mds.variables(end).type = '';
    mds.variables(end).compartment = '';
    mds.variables(end).unittype = '';
    mds.variables(end).notes = sprintf(' Input variable realizing ''%s'' on state ''%s'' (ABSORPTION0).',iname,ms.states(stateindex).name);
    % e) Remove the INPUT* parameter (will change the parindex fields but
    % doesn't matter, since this input parameters are not used anymore ...
    % if a dosing input is defined for this INPUT* element).
    index = strmatchIQM(iname,{mds.parameters.name},'exact');
    mds.parameters(index) = [];
    % f) Need to define an initial set of parameters used in the vformula.
    % Single dosing events can be implemented in this way. In the case of
    % multiple dosing events only the first will be implemented in the
    % model. The remaining ones will be handled using an outside loop in a
    % higher level function.
    if k==1,    % only add Dose and Time parameters once!
        % Dose
        mds.parameters(end+1).name = iDosename;
        mds.parameters(end).value = 0;%inputinfo.D(1);
        mds.parameters(end).type = '';
        mds.parameters(end).compartment = '';
        mds.parameters(end).unittype = '';
        mds.parameters(end).notes = sprintf(' Dose amount for INPUT ''%s'' (ABSORPTION0).', iname);
        % Time
        mds.parameters(end+1).name = iTimename;
        mds.parameters(end).value = inputinfo.time(1);
        mds.parameters(end).type = '';
        mds.parameters(end).compartment = '';
        mds.parameters(end).unittype = '';
        mds.parameters(end).notes = sprintf(' Dosing instant for INPUT ''%s'' (ABSORPTION0).', iname);
        % Tk0
        mds.parameters(end+1).name = iTk0name;
        mds.parameters(end).value = inputinfo.parameters(1).value(1);
        mds.parameters(end).type = '';
        mds.parameters(end).compartment = '';
        mds.parameters(end).unittype = '';
        mds.parameters(end).notes = sprintf('%s Dosing duration, used in variable ''%s'' (ABSORPTION0).', inputinfo.parameters(1).notes,ivarname);
        % Add the necessary information to the experiment structure (only time
        % and dosing amounts can change between subsequent dosing instants
[eds] = buildexpstruct(eds,iDosename,iTimename,iTlagname,inputinfo,ivarname,iTk0name,inputinfo.parameters(1).value);
%         % Additionally set the ka rate parameters in the experiment
%         eds = addparamicsetting(eds,iTk0name,num2str(inputinfo.parameters(1).value),0,'');
    end
end
return

%%%%%%%%%%%%%%%%%%%%%%%%%%%%%%%%%%%%%%%%%%%%%%%%%%%%%%%%%%%%%%%%%%%%%%%%%%%
% help function to get right names
%%%%%%%%%%%%%%%%%%%%%%%%%%%%%%%%%%%%%%%%%%%%%%%%%%%%%%%%%%%%%%%%%%%%%%%%%%%
function outname = getoutname(base, kmax, k)
if kmax == 1,
    outname = base;
else
    outname = [base '_' sprintf('%d',k)];
end
return

%%%%%%%%%%%%%%%%%%%%%%%%%%%%%%%%%%%%%%%%%%%%%%%%%%%
% Handle the experiment description for simulation
%%%%%%%%%%%%%%%%%%%%%%%%%%%%%%%%%%%%%%%%%%%%%%%%%%%
% The first dosing instant is going to be put into the parameter and ic
% settings section (same as already in the model). The subsequent
% dosings in case of multiple doses is put into the event section.
% parameter section. This function is the same for all dosing types. So no
% need to write it 4 times.
function [eds] = buildexpstruct(eds,iDosename,iTimename,iTlagname,inputinfo,ivarname,iParName,iParValue)
eds = addparamicsetting(eds,iDosename,num2str(inputinfo.D(1)),0,'');
eds = addparamicsetting(eds,iTimename,num2str(inputinfo.time(1)),0,'');
% Same with the additional dosing parameter if defined
if ~isempty(iParName),
    eds = addparamicsetting(eds,iParName,num2str(iParValue(1)),0,'');
end

% Tlag (only if defined in dosing schedule) added for each input instant
if ~isempty(inputinfo.Tlag),
    eds = addparamicsetting(eds,iTlagname,num2str(inputinfo.Tlag),0,'');
end

% subsequent dosings in the events (Time and Dose needs to be
% updated by events and also the dosing parameter, if defined)
if length(inputinfo.time) > 1,
    % multiple dosings => get time vector and dose vector (without
    % first dosing instance)
    timevec = inputinfo.time(2:end);
    Dvec = inputinfo.D(2:end);
    if ~isempty(iParValue),
        parvec = iParValue(2:end);
    else
        parvec = [];
    end
    for k2=1:length(timevec),
        % add the events
        name = sprintf('%s_event_%d',ivarname,k2);
        if ~isempty(parvec),
            eds = addstateevent(eds,name,timevec(k2),{iDosename,iTimename,iParName},{num2str(Dvec(k2)),num2str(timevec(k2)),num2str(parvec(k2))},'');
        else
            eds = addstateevent(eds,name,timevec(k2),{iDosename,iTimename},{num2str(Dvec(k2)),num2str(timevec(k2))},'');
        end            
    end
end
return

%%%%%%%%%%%%%%%%%%%%%%%%%%%%%%%%%%%%%%%%%%%%%%%%%%%%%%%%%%%%%%%%%%%%%%%%%%%
% add parameter or ic to experiment
%%%%%%%%%%%%%%%%%%%%%%%%%%%%%%%%%%%%%%%%%%%%%%%%%%%%%%%%%%%%%%%%%%%%%%%%%%%
function [eds] = addparamicsetting(eds,name,formula,icflag,notes)
    eds.paramicsettings(end+1).name = name;
    eds.paramicsettings(end).formula = formula;
    eds.paramicsettings(end).icflag = icflag;
    eds.paramicsettings(end).notes = notes;
return

%%%%%%%%%%%%%%%%%%%%%%%%%%%%%%%%%%%%%%%%%%%%%%%%%%%%%%%%%%%%%%%%%%%%%%%%%%%
% add stateevents to experiment
%%%%%%%%%%%%%%%%%%%%%%%%%%%%%%%%%%%%%%%%%%%%%%%%%%%%%%%%%%%%%%%%%%%%%%%%%%%
function [eds] = addstateevent(eds,name,time,compnames,compformulas,notes)
    eds.stateevents(end+1).name = name;
    eds.stateevents(end).trigger = ['ge(time,' num2str(time) ')'];
    for k=1:length(compnames),
        eds.stateevents(end).assignment(k).variable = compnames{k};
        eds.stateevents(end).assignment(k).formula = compformulas{k};
    end
    eds.stateevents(end).notes = notes;
return
```
