## Supplementary material for "A systems biology analysis of adrenergically stimulated adiponectin exocytosis in white adipocytes": All scripts used in this work: IQMcreateEXPfile.m

```
function IQMcreateEXPfile(varargin)
% IQMcreateEXPfile: creates a *.exp file with the experiments text description
%
% USAGE:
% ======
% [] = IQMcreateEXPfile()         
% [] = IQMcreateEXPfile(filename)         
% [] = IQMcreateEXPfile(exp)         
% [] = IQMcreateEXPfile(exp,filename)
%
% exp: IQMexperiment object to convert to a textfile description
% filename: filename for the created textfile 
%
% If exp is undefined, then an empty experiment file will be created. 
%
% DEFAULT VALUES:
% ===============
% exp: the experiment to be exported as EXP file. 
% filename: constructed from the experiments name

% <<<COPYRIGHTSTATEMENT - IQM TOOLS LITE>>>


%%%%%%%%%%%%%%%%%%%%%%%%%%%%%%%%%%%%%%%%%%%%%%%
% PROCESS VARIABLE INPUT ARGUMENTS
%%%%%%%%%%%%%%%%%%%%%%%%%%%%%%%%%%%%%%%%%%%%%%%
if nargin == 0,
    % create empty experiment file with the name "unnamed.exp"
    exp = IQMexperiment();
    filename = 'unnamed.exp';
elseif nargin == 1,
    % check if first input argument experiment or filename
    if isIQMexperiment(varargin{1}),
        % experiment given
        exp = varargin{1};
        % if no filename provided then use the name of the IQMexperiment object as filename
        % remove unwanted characters first
        es = struct(exp);
%         functionName = regexprep(es.name,'\W','');
        functionName = es.name;
        filename = strcat(functionName,'.exp');
    else
        % filename given?
        if ~ischar(varargin{1}),
            error('Wrong input argument.');
        end
        filename = strcat(varargin{1},'.exp');
        exp = IQMexperiment();
    end
elseif nargin == 2,
    exp = varargin{1};
    filename = strcat(varargin{2},'.exp');
else
    error('Incorrect number of input arguments.');
end

%%%%%%%%%%%%%%%%%%%%%%%%%%%%%%%%%%%%%%%%%%%%%%%
% CHECK INPUT
%%%%%%%%%%%%%%%%%%%%%%%%%%%%%%%%%%%%%%%%%%%%%%%
if ~isIQMexperiment(exp),
    error('No IQMexperiment as first argument.');
end

%%%%%%%%%%%%%%%%%%%%%%%%%%%%%%%%%%%%%%%%%%%%%%%
% CONVERT MODEL TO TEXT
%%%%%%%%%%%%%%%%%%%%%%%%%%%%%%%%%%%%%%%%%%%%%%%
[expTextStructure] = convertExpToTextIQM(exp);
[completeText] = setPartsToCompleteTextExpIQM(expTextStructure);

%%%%%%%%%%%%%%%%%%%%%%%%%%%%%%%%%%%%%%%%%%%%%%%
% WRITE THE FILE
%%%%%%%%%%%%%%%%%%%%%%%%%%%%%%%%%%%%%%%%%%%%%%%
fid = fopen(filename,'w');
fwrite(fid,completeText);
fclose(fid);
return
```
