## Supplementary material for "A systems biology analysis of adrenergically stimulated adiponectin exocytosis in white adipocytes": All scripts used in this work: IQMmergemodexp.m

```
function [expmodel] = IQMmergemodexp(model, experiment)
% IQMmergemodexp: combines an experiment with a model, and produces a "merged 
% model", as output. The original model and experiment arguments can be
% given as textfiles, or IQMexperiment objects. The output model is an
% IQMmodel.
%
% DESCRIPTIONS + SYNTAX:
% ======================
% To fill in!
%
% USAGE:
% ======
% [expmodel] = IQMmergemodexp(model, experiment)        
% [expmodel] = IQMmergemodexp(model, experimentfile)        
%
% model: IQMmodel 
% experiment: An IQMexperiment object describing an experiment that should be done
%             with the model
% experimentfile: String with the name of an experiment file
%
% Output Arguments:
% =================
% Merged model containing the original model and the experimental settings.

% <<<COPYRIGHTSTATEMENT - IQM TOOLS LITE>>>


expmodel = [];
time = 0;   % per default time=0 is assumed

%%%%%%%%%%%%%%%%%%%%%%%%%%%%%%%%%
% Check input arguments 
%%%%%%%%%%%%%%%%%%%%%%%%%%%%%%%%%
if ~isIQMmodel(model),
    error('The first input argument needs to be an IQMmodel.');
else
    modelstruct = IQMstruct(model);
end
if ~isIQMexperiment(experiment),
    % Then see if this file exists
    if (exist(experiment, 'file') == 2)
       % try open this experiment file 
       try
          experiment = IQMexperiment(experiment); 
       catch exception
           error('The second input argument needs to be either a IQMexperiment or an experiment text file.');
       end
    else    
        error('The second input argument needs to be either a IQMexperiment or an experiment text file.');
    end
end
expstruct = IQMstruct(experiment);

%%%%%%%%%%%%%%%%%%%%%%%%%%%%%%%%%
% Initialize the output model structure
%%%%%%%%%%%%%%%%%%%%%%%%%%%%%%%%%
newmodstruct = modelstruct;                    

%%%%%%%%%%%%%%%%%%%%%%%%%%%%%%%%%
% Define all initial conditions and parameters in the workspace. Do not
% define VARIABLES and REACTIONS! FIRST DEFINE PARAMETERS ... then ICs,
% since ICs can depend on parameters.
%%%%%%%%%%%%%%%%%%%%%%%%%%%%%%%%%
for kkkloopnotuseinmodel = 1:length(newmodstruct.parameters),
    if ~isempty(newmodstruct.parameters(kkkloopnotuseinmodel).value),
        eval([newmodstruct.parameters(kkkloopnotuseinmodel).name '=' num2str(newmodstruct.parameters(kkkloopnotuseinmodel).value) ';']);
    else
        error('Value for parameter ''%s'' is undefined.',newmodstruct.parameters(kkkloopnotuseinmodel).name);
    end
end
for kkkloopnotuseinmodel = 1:length(newmodstruct.states),
    if ~isempty(newmodstruct.states(kkkloopnotuseinmodel).initialCondition),
        eval([newmodstruct.states(kkkloopnotuseinmodel).name '=' num2str(newmodstruct.states(kkkloopnotuseinmodel).initialCondition) ';']);
    else
        error('Initial condition for state ''%s'' is undefined.',newmodstruct.states(kkkloopnotuseinmodel).name);
    end
end

%%%%%%%%%%%%%%%%%%%%%%%%%%%%%%%%%
% Define all MODEL FUNCTIONS as inline objects to be used in the
% determination of initial conditions and initial parameter settings
%%%%%%%%%%%%%%%%%%%%%%%%%%%%%%%%%
for kkkloopnotuseinmodel = 1:length(newmodstruct.functions),
    eval(sprintf('%s = @(%s)%s;',newmodstruct.functions(kkkloopnotuseinmodel).name,newmodstruct.functions(kkkloopnotuseinmodel).arguments,newmodstruct.functions(kkkloopnotuseinmodel).formula));
end


%%%%%%%%%%%%%%%%%%%%%%%%%%%%%%%%%
% Now just evaluate the paramicsettings sequentially 
% and add the new models to the structure
%%%%%%%%%%%%%%%%%%%%%%%%%%%%%%%%%
for kkkloopnotuseinmodel = 1:length(expstruct.paramicsettings)
    if ~isempty(expstruct.paramicsettings(kkkloopnotuseinmodel).formula),
        try
            % get value and assign value to variable in workspace
            valuenotuseinmodel = eval(expstruct.paramicsettings(kkkloopnotuseinmodel).formula);
            eval([expstruct.paramicsettings(kkkloopnotuseinmodel).name '=' num2str(valuenotuseinmodel) ';']);
        catch
            if expstruct.paramicsettings(kkkloopnotuseinmodel).icflag==1,
                text = sprintf('Error in initial condition setting for state ''%s'':\n\n"%s"',expstruct.paramicsettings(kkkloopnotuseinmodel).name,lasterr);
                text = sprintf('%s\n\nNote: Only a models states and parameters can be used in mathematical\nexpressions for the initial parameter and state settings.',text);
                error(text);
            else
                text = sprintf('Error in initial condition setting for parameter ''%s'':\n\n"%s"',expstruct.paramicsettings(kkkloopnotuseinmodel).name,lasterr);
                text = sprintf('%s\n\nOnly a models states and parameters can be used in mathematical\nexpressions for the initial parameter and state settings.',text);
                error(text);
            end
        end
    else
        error('Formula for initial condition for state ''%s'' is undefined: %s',expstruct.paramicsettings(kkkloopnotuseinmodel).name,lasterr);
    end
    % value determined (valuenotuseinmodel). Add it to the model structure
    if expstruct.paramicsettings(kkkloopnotuseinmodel).icflag==1,
        % if initial condition then search states
        indexnotuseinmodel = strmatchIQM(expstruct.paramicsettings(kkkloopnotuseinmodel).name,{newmodstruct.states.name},'exact');
        if isempty(indexnotuseinmodel),
            error('Initial condition for ''%s'' defined in experiment but state does not exist in the model.',expstruct.paramicsettings(kkkloopnotuseinmodel).name);
        end 
        newmodstruct.states(indexnotuseinmodel).initialCondition = valuenotuseinmodel;
        newmodstruct.states(indexnotuseinmodel).notes = expstruct.paramicsettings(kkkloopnotuseinmodel).notes;
    else
        % if not initial condition then search in parameters 
        indexnotuseinmodel = strmatchIQM(expstruct.paramicsettings(kkkloopnotuseinmodel).name,{newmodstruct.parameters.name},'exact');
        if ~isempty(indexnotuseinmodel),
            % only update value if parameter appears in the model (help
            % variables are handled fine).
            newmodstruct.parameters(indexnotuseinmodel).value = valuenotuseinmodel;
            newmodstruct.parameters(indexnotuseinmodel).notes = expstruct.paramicsettings(kkkloopnotuseinmodel).notes;
        end
    end
end

%%%%%%%%%%%%%%%%%%%%%%%%%%%%%%%%%
% Add the parameter changes to the model
%%%%%%%%%%%%%%%%%%%%%%%%%%%%%%%%%
% Copy old variables to be able to add the new variables (parameters) first
oldvariables = newmodstruct.variables;
modelstructempty = struct(IQMmodel());
newmodstruct.variables = modelstructempty.variables;
% Add new variables as first in the list
for k = 1:length(expstruct.parameterchanges),
    % check that no parameter is defined twice
    if ~isempty(strmatchIQM(expstruct.parameterchanges(k).name,{expstruct.paramicsettings.name},'exact')),
        error('Parameter ''%s'' is defined twice (settings and changes).',expstruct.parameterchanges(k).name);
    end
    % delete this parameter from the model parameters
    index = strmatchIQM(expstruct.parameterchanges(k).name,{newmodstruct.parameters.name},'exact');
    if isempty(index),
        error('Parameter ''%s'' not present in the model but changed in the experiment.',expstruct.parameterchanges(k).name);
    end
    newmodstruct.parameters(index) = [];
    newmodstruct.variables(end+1).name = expstruct.parameterchanges(k).name;
    newmodstruct.variables(end).formula = expstruct.parameterchanges(k).formula;
    newmodstruct.variables(end).notes = expstruct.parameterchanges(k).notes;
    newmodstruct.variables(end).type = '';
    newmodstruct.variables(end).compartment = '';
    newmodstruct.variables(end).unittype = '';
end
% Add old variables at the end
for k = 1:length(oldvariables),
    newmodstruct.variables(end+1).name = oldvariables(k).name;
    newmodstruct.variables(end).formula = oldvariables(k).formula;
    newmodstruct.variables(end).notes = oldvariables(k).notes;
    newmodstruct.variables(end).type = oldvariables(k).type;
    newmodstruct.variables(end).compartment = oldvariables(k).compartment;
    newmodstruct.variables(end).unittype = oldvariables(k).unittype;    
end

%%%%%%%%%%%%%%%%%%%%%%%%%%%%%%%%%
% Add experiment events to the model
%%%%%%%%%%%%%%%%%%%%%%%%%%%%%%%%%
for k = 1:length(expstruct.stateevents)
    newmodstruct.events(end+1) = expstruct.stateevents(k);
end

%%%%%%%%%%%%%%%%%%%%%%%%%%%%%%%%%
% return the experiment model
%%%%%%%%%%%%%%%%%%%%%%%%%%%%%%%%%
expmodel = IQMmodel(newmodstruct);
return


%%%%%%%%%%%%%%%%%%%%%%%%%%%%%%%%%%%%%%%%%%%%%%%%%%%%%%%%%%%%%%%%%%%%%%%%%%%
%%% The "find index" function
%%%%%%%%%%%%%%%%%%%%%%%%%%%%%%%%%%%%%%%%%%%%%%%%%%%%%%%%%%%%%%%%%%%%%%%%%%%
function [index] = find_index(name_array, comp_string)
k = 0;
l = length(name_array);
index = inf;
while k < l
    k = k+1;
    if strcmp(comp_string, name_array(k))
        index = k;
    end
end
if index == inf,
    error('The experiment description contains the element ''%s'' that is not present in the model.',comp_string);
end
return
```
