## Supplementary material for "A systems biology analysis of adrenergically stimulated adiponectin exocytosis in white adipocytes": All scripts used in this work: IQMexportCSVmeasurement.m

```
function [] = IQMexportCSVmeasurement(varargin)
% IQMexportCSVmeasurement
% Exports an IQMmeasurement object to a CSV (comma separated value) file.
% The format of the written CSV file is explained in the user's reference
% manual and example files can be found in the IQMlite/examples folder.
% 
% USAGE:
% ======
% [] = IQMexportCSVmeasurement(measurement)
% [] = IQMexportCSVmeasurement(measurement,filename)
%
% measurement: IQMmeasurement object containing the data
% filename:    desired filename for CSV file. The extension '.csv' is not
%              required.
%
% DEFAULT VALUES:
% ===============
% filename: constructed from the data objects name

% <<<COPYRIGHTSTATEMENT - IQM TOOLS LITE>>>


%%%%%%%%%%%%%%%%%%%%%%%%%%%%%%%%%%%%%%%%%%%%%%%
% PROCESS VARIABLE INPUT ARGUMENTS
%%%%%%%%%%%%%%%%%%%%%%%%%%%%%%%%%%%%%%%%%%%%%%%
if nargin == 1,
    measurement = varargin{1};
    % convert object to structure
    measurement = struct(measurement);    
    % if no filename provided then use the name of the IQMmeasurement object 
    % as filename. Just delete all the special characters.
    filename = measurement.name;   % white spaces
elseif nargin == 2,
    measurement = varargin{1};
    % convert object to structure
    measurement = struct(measurement);    
    % extract filename from input arguments to skip eventual extension
    [PATHSTR,filename,EXT] = fileparts(varargin{2});
else
    error('Incorrect number of input arguments.');
end

%%%%%%%%%%%%%%%%%%%%%%%%%%%%%%%%%%%%%%%%%%%%%%%
% CHECK IF OBJECT CONTAINS MEASUREMENTS
%%%%%%%%%%%%%%%%%%%%%%%%%%%%%%%%%%%%%%%%%%%%%%%
if length(measurement.data) == 0,
    error('The object does not contain any measurements');
end

%%%%%%%%%%%%%%%%%%%%%%%%%%%%%%%%%%%%%%%%%%%%%%%
% Create the file 
%%%%%%%%%%%%%%%%%%%%%%%%%%%%%%%%%%%%%%%%%%%%%%%
fid = fopen(strcat(filename,'.csv'),'w');

%%%%%%%%%%%%%%%%%%%%%%%%%%%%%%%%%%%%%%%%%%%%%%%
% Write header
%%%%%%%%%%%%%%%%%%%%%%%%%%%%%%%%%%%%%%%%%%%%%%%
fprintf(fid,'%% Measurement file generated: %s\n\n',date);

%%%%%%%%%%%%%%%%%%%%%%%%%%%%%%%%%%%%%%%%%%%%%%%
% NAME
%%%%%%%%%%%%%%%%%%%%%%%%%%%%%%%%%%%%%%%%%%%%%%%
fprintf(fid,'[Name]\n');
fprintf(fid,'%s\n',measurement.name);
fprintf(fid,'\n');

%%%%%%%%%%%%%%%%%%%%%%%%%%%%%%%%%%%%%%%%%%%%%%%
% NOTES
%%%%%%%%%%%%%%%%%%%%%%%%%%%%%%%%%%%%%%%%%%%%%%%
fprintf(fid,'[Notes]\n');
fprintf(fid,'%s\n',measurement.notes);
fprintf(fid,'\n');

%%%%%%%%%%%%%%%%%%%%%%%%%%%%%%%%%%%%%%%%%%%%%%%
% COMPONENTS
%%%%%%%%%%%%%%%%%%%%%%%%%%%%%%%%%%%%%%%%%%%%%%%
fprintf(fid,'[Components]\n');
text = 'time,';
for k=1:length(measurement.data),
    text = sprintf('%s%s,',text,measurement.data(k).name);
    % add error bound names
    if ~isempty(measurement.data(k).maxvalues) && max(isnan(measurement.data(k).maxvalues))~=1,
        text = sprintf('%s%s+,',text,measurement.data(k).name);
    end
    if ~isempty(measurement.data(k).minvalues) && max(isnan(measurement.data(k).minvalues))~=1,
        text = sprintf('%s%s-,',text,measurement.data(k).name);
    end
end
text = text(1:end-1);
fprintf(fid,'%s\n',text);
fprintf(fid,'\n');

%%%%%%%%%%%%%%%%%%%%%%%%%%%%%%%%%%%%%%%%%%%%%%%
% COMPONENTNOTES
%%%%%%%%%%%%%%%%%%%%%%%%%%%%%%%%%%%%%%%%%%%%%%%
fprintf(fid,'[Componentnotes]\n');
for k=1:length(measurement.data),
    if ~isempty(measurement.data(k).notes),
        notes = regexprep(measurement.data(k).notes,'\n',' ');
        notes = regexprep(notes,'  ',' ');
        notes = regexprep(notes,'  ',' ');
        if ~isempty(measurement.data(k).name),
            fprintf(fid,'%s: %s\n',measurement.data(k).name,notes);
        else
            fprintf(fid,'%s: %s\n',measurement.data(k).formula,notes);
        end
    end
end
fprintf(fid,'\n');

%%%%%%%%%%%%%%%%%%%%%%%%%%%%%%%%%%%%%%%%%%%%%%%
% Values
%%%%%%%%%%%%%%%%%%%%%%%%%%%%%%%%%%%%%%%%%%%%%%%
fprintf(fid,'[Values]\n');
experimentData = measurement.time(:);
for k = 1:length(measurement.data),
    experimentData(:,end+1) = measurement.data(k).values(:);
    % add error bound values
    if ~isempty(measurement.data(k).maxvalues) && max(isnan(measurement.data(k).maxvalues))~=1,
        experimentData(:,end+1) = measurement.data(k).maxvalues(:);
    end
    if ~isempty(measurement.data(k).minvalues) && max(isnan(measurement.data(k).minvalues))~=1,
        experimentData(:,end+1) = measurement.data(k).minvalues(:);
    end
end
dataString = '';
for k = 1:size(experimentData,1),
    % Change for accurate printing of low values
    %dataString = sprintf('%f,',experimentData(k,:));
    dataString = sprintf('%14.12g,',experimentData(k,:));
    fprintf(fid,'%s\n',dataString(1:end-1));
end

%%%%%%%%%%%%%%%%%%%%%%%%%%%%%%%%%%%%%%%%%%%%%%%
% Close the file
%%%%%%%%%%%%%%%%%%%%%%%%%%%%%%%%%%%%%%%%%%%%%%%
fclose(fid);
return
```
