## Supplementary material for "A systems biology analysis of adrenergically stimulated adiponectin exocytosis in white adipocytes": All scripts used in this work: IQMexportXLSmeasurement.m

```
function [] = IQMexportXLSmeasurement(varargin)
% IQMexportXLSmeasurement
% Exports an IQMmeasurement object to an XLS (excel) file.
% The format of the written XLS file is explained in the user's reference
% manual and example files can be found in the IQMlite/examples folder.
%
% USAGE:
% ======
% [] = IQMexportXLSmeasurement(measurement)
% [] = IQMexportXLSmeasurement(measurement,filename)
% [] = IQMexportXLSmeasurement(measurement,filename,sheet)
%
% measurement: IQMmeasurement object containing the data
% filename:    desired filename for XLS file. The extension '.xls' is not
%              required.
% sheet:       number of the sheet in the Excel file to which the data
%              should be written.
%
% DEFAULT VALUES:
% ===============
% filename: constructed from the data objects name

% <<<COPYRIGHTSTATEMENT - IQM TOOLS LITE>>>


%%%%%%%%%%%%%%%%%%%%%%%%%%%%%%%%%%%%%%%%%%%%%%%
% PROCESS VARIABLE INPUT ARGUMENTS
%%%%%%%%%%%%%%%%%%%%%%%%%%%%%%%%%%%%%%%%%%%%%%%
sheet = 1;
if nargin == 1,
    measurement = varargin{1};
    % convert object to structure
    measurement = struct(measurement);
    % if no filename provided then use the name of the IQMmeasurement object
    % as filename. Just delete all the special characters.
    filename = regexprep(measurement.name,'\s','');   % white spaces
    filename = regexprep(filename,'\W','');    % other
elseif nargin == 2,
    measurement = varargin{1};
    % convert object to structure
    measurement = struct(measurement);
    % extract filename from input arguments to skip eventual extension
    [PATHSTR,filename,EXT] = fileparts(varargin{2});
elseif nargin == 3,
    measurement = varargin{1};
    % convert object to structure
    measurement = struct(measurement);
    % extract filename from input arguments to skip eventual extension
    [PATHSTR,filename,EXT] = fileparts(varargin{2});
    sheet = varargin{3};
else
    error('Incorrect number of input arguments.');
end

%%%%%%%%%%%%%%%%%%%%%%%%%%%%%%%%%%%%%%%%%%%%%%%
% CHECK IF OBJECT CONTAINS MEASUREMENTS
%%%%%%%%%%%%%%%%%%%%%%%%%%%%%%%%%%%%%%%%%%%%%%%
numberComponents = length(measurement.data);
if numberComponents == 0,
    error('The object does not contain any measurements');
end

%%%%%%%%%%%%%%%%%%%%%%%%%%%%%%%%%%%%%%%%%%%%%%%
% Construct Cell-Array Matrix of correct size
%%%%%%%%%%%%%%%%%%%%%%%%%%%%%%%%%%%%%%%%%%%%%%%
RAW = NaN*ones(4+length(measurement.time),numberComponents+2);
RAW = num2cell(RAW);

%%%%%%%%%%%%%%%%%%%%%%%%%%%%%%%%%%%%%%%%%%%%%%%
% NAME
%%%%%%%%%%%%%%%%%%%%%%%%%%%%%%%%%%%%%%%%%%%%%%%
RAW{1,1} = 'Name';
RAW{1,2} = measurement.name;

%%%%%%%%%%%%%%%%%%%%%%%%%%%%%%%%%%%%%%%%%%%%%%%
% NOTES
%%%%%%%%%%%%%%%%%%%%%%%%%%%%%%%%%%%%%%%%%%%%%%%
% It seems that matlab requires the entry sizes to be max 911.
% So lets split them up. Make sure we truncate at a space.
RAW{2,1} = 'Notes';
offset = 0;
if length(measurement.notes) <= 911,
    RAW{2,2} = measurement.notes;
else
    notes = measurement.notes;
    while length(notes) > 911,
        index = 911;
        while double(notes(index)) ~= 32,
            index = index - 1;
        end
        RAW{2+offset,2} = notes(1:index-1);
        offset = offset + 1;
        notes = notes(index+1:end);
    end
    RAW{2+offset,2} = notes;
end

%%%%%%%%%%%%%%%%%%%%%%%%%%%%%%%%%%%%%%%%%%%%%%%
% COMPONENTS
%%%%%%%%%%%%%%%%%%%%%%%%%%%%%%%%%%%%%%%%%%%%%%%
componentsall = {};
RAW{4+offset,1} = 'Components';
RAW{4+offset,2} = 'time';
index = 3;
for k=1:numberComponents,
    RAW{4+offset,index} = measurement.data(k).name;
    componentsall{end+1} = measurement.data(k).name;
    index = index+1;
    % add error bound names
    if ~isempty(measurement.data(k).maxvalues) && max(isnan(measurement.data(k).maxvalues))~=1,
        RAW{4+offset,index} = sprintf('%s+',measurement.data(k).name);
        componentsall{end+1} = sprintf('%s+',measurement.data(k).name);
        index = index + 1;
    end
    if ~isempty(measurement.data(k).minvalues) && max(isnan(measurement.data(k).minvalues))~=1,
        RAW{4+offset,index} = sprintf('%s-',measurement.data(k).name);
        componentsall{end+1} = sprintf('%s-',measurement.data(k).name);
        index = index + 1;
    end
end

%%%%%%%%%%%%%%%%%%%%%%%%%%%%%%%%%%%%%%%%%%%%%%%
% COMPONENTNOTES
%%%%%%%%%%%%%%%%%%%%%%%%%%%%%%%%%%%%%%%%%%%%%%%
RAW{3+offset,1} = 'Componentnotes';
for k=1:numberComponents,
    index = strmatchIQM(measurement.data(k).name,componentsall,'exact');
    RAW{3+offset,index+2} = measurement.data(k).notes;
end

%%%%%%%%%%%%%%%%%%%%%%%%%%%%%%%%%%%%%%%%%%%%%%%
% VALUES
%%%%%%%%%%%%%%%%%%%%%%%%%%%%%%%%%%%%%%%%%%%%%%%
RAW{5+offset,1} = 'Values';
RAW(5+offset:5+offset+length(measurement.time)-1,2) = num2cell(measurement.time);
index = 3;
for k=1:numberComponents,
    RAW(5+offset:5+offset+length(measurement.data(k).values)-1,index) = num2cell(measurement.data(k).values);
    index = index + 1;
    if ~isempty(measurement.data(k).maxvalues) && max(isnan(measurement.data(k).maxvalues))~=1,
        RAW(5+offset:5+offset+length(measurement.data(k).values)-1,index) = num2cell(measurement.data(k).maxvalues);
        index = index + 1;
    end
    if ~isempty(measurement.data(k).minvalues) && max(isnan(measurement.data(k).minvalues))~=1,
        RAW(5+offset:5+offset+length(measurement.data(k).values)-1,index) = num2cell(measurement.data(k).minvalues);
        index = index + 1;
    end
end

%%%%%%%%%%%%%%%%%%%%%%%%%%%%%%%%%%%%%%%%%%%%%%%
% WRITE TO FILE
%%%%%%%%%%%%%%%%%%%%%%%%%%%%%%%%%%%%%%%%%%%%%%%
[success,message] = xlswrite(strcat(filename,'.xls'),RAW,sheet);
return
```
