## Supplementary material for "A systems biology analysis of adrenergically stimulated adiponectin exocytosis in white adipocytes": All scripts used in this work: IQMexportXLSmeasurements.m

```
function [] = IQMexportXLSmeasurements(measurements,filename)
% IQMexportXLSmeasurement
% Exports several IQMmeasurement objects to the same XLS (excel) file.
% Each measurement will be added to a separate sheet in the file.
%
% USAGE:
% ======
% [] = IQMexportXLSmeasurement(measurements,filename)
%
% measurements: A cell-array in which all the elements are IQMmeasurement objects.
% filename:     Desired filename for XLS file. The extension '.xls' is not
%               required.

% <<<COPYRIGHTSTATEMENT - IQM TOOLS LITE>>>

[PATHSTR,filename,EXT] = fileparts(filename);

%%%%%%%%%%%%%%%%%%%%%%%%%%%%%%%%%%%%%%%%%%%%%%%
% CHECK IF MULTIPLE MEASUREMENTS
%%%%%%%%%%%%%%%%%%%%%%%%%%%%%%%%%%%%%%%%%%%%%%%
if ~iscell(measurements),
    input = {measurements};
else
    input = measurements;
end

%%%%%%%%%%%%%%%%%%%%%%%%%%%%%%%%%%%%%%%%%%%%%%%
% DELETE FILE WITH FILENAME IF PRESENT
%%%%%%%%%%%%%%%%%%%%%%%%%%%%%%%%%%%%%%%%%%%%%%%
warning off;
delete(strcat(filename,'.xls'));
warning on;

%%%%%%%%%%%%%%%%%%%%%%%%%%%%%%%%%%%%%%%%%%%%%%%
% LOOP THROUGH THE MEASUREMENTS AND EXPORT
%%%%%%%%%%%%%%%%%%%%%%%%%%%%%%%%%%%%%%%%%%%%%%%
for sheet=1:length(input),
    measurement = input{sheet};
    IQMexportXLSmeasurement(measurement,strcat(filename,'.xls'),sheet);
end
return
```
