## Supplementary material for "A systems biology analysis of adrenergically stimulated adiponectin exocytosis in white adipocytes": All scripts used in this work: IQMimportCSVmeasurement.m

```
function [measurementstructure,errorMsg] = IQMimportCSVmeasurement(filename)
% IQMimportCSVmeasurement
% Imports experimental measurement data stored in an CSV (comma separated 
% value) file into the measurement structure used by the IQMmeasurement
% object. Please note that a special format of the CSV measurement is
% required. This format is explained in the user's reference manual and
% example files can be found in the IQMlite/examples folder.
% 
% filename: name of the .csv file containing the measurement
%
% measurementstructure: measurement structure used by IQMmeasurement object 
%                      (empty if error occurred)
% errorMsg: string containing possible error messages. 

% <<<COPYRIGHTSTATEMENT - IQM TOOLS LITE>>>


%%%%%%%%%%%%%%%%%%%%%%%%%%%%%%%%%%%%%%%%%%%%%%%
% INITIALIZATION
%%%%%%%%%%%%%%%%%%%%%%%%%%%%%%%%%%%%%%%%%%%%%%%
errorMsg = '';
measurementstructure = struct(IQMmeasurement());

%%%%%%%%%%%%%%%%%%%%%%%%%%%%%%%%%%%%%%%%%%%%%%%
% Open the file and read all its content 
% Skip the comments
%%%%%%%%%%%%%%%%%%%%%%%%%%%%%%%%%%%%%%%%%%%%%%%
content = sprintf('\n%s',fileread(filename));
content = regexprep(content,'\n%[^\n]*','');

%%%%%%%%%%%%%%%%%%%%%%%%%%%%%%%%%%%%%%%%%%%%%%%
% Extract the pieces
%%%%%%%%%%%%%%%%%%%%%%%%%%%%%%%%%%%%%%%%%%%%%%%
% Find the starts of the different view data
nameStart = regexp(content,'\[Name\]');
notesStart = regexp(content,'\[Notes\]');
componentsStart = regexp(content,'\[Components\]');
componentnotesStart = regexp(content,'\[Componentnotes\]');
valuesStart = regexp(content,'\[Values\]');
if isempty(nameStart) || isempty(notesStart) || isempty(componentsStart) || isempty(componentnotesStart) || isempty(valuesStart),
    errorMsg = sprintf('%sAt least one of the identifiers in the measurement file is missing or misspelled.\n',errorMsg);
end
% Cut out the different pieces and assign them to the modelTextStructure structure
nameraw = strtrim(content(nameStart+length('[Name]'):notesStart-1));
notesraw = strtrim(content(notesStart+length('[Notes]'):componentsStart-1));
componentsraw = strtrim(content(componentsStart+length('[Components]'):componentnotesStart-1));
componentnotesraw = strtrim(content(componentnotesStart+length('[Componentnotes]'):valuesStart-1));
valuesraw = strtrim(content(valuesStart+length('[Values]'):end));

%%%%%%%%%%%%%%%%%%%%%%%%%%%%%%%%%%%%%%%%%%%%%%%
% Get name
%%%%%%%%%%%%%%%%%%%%%%%%%%%%%%%%%%%%%%%%%%%%%%%
if isempty(nameraw),
    measurementstructure.name = 'untitled';
else
    % just remove special signs that are not allowed for file names.
    measurementstructure.name = regexprep(nameraw,'\W','');
end

%%%%%%%%%%%%%%%%%%%%%%%%%%%%%%%%%%%%%%%%%%%%%%%
% Get notes
%%%%%%%%%%%%%%%%%%%%%%%%%%%%%%%%%%%%%%%%%%%%%%%
measurementstructure.notes = notesraw;

%%%%%%%%%%%%%%%%%%%%%%%%%%%%%%%%%%%%%%%%%%%%%%%
% Get components - check also for timeindex
%%%%%%%%%%%%%%%%%%%%%%%%%%%%%%%%%%%%%%%%%%%%%%%
% explode the components
components = explodePCIQM(componentsraw);
% remove spaces from comment names
for k = 1:length(components),
    components = regexprep(components,' ','');
end
% find 'time' component
timeindex = strmatchIQM('time',components,'exact');
% initialize help structure for min max values (error bounds) + components
errorbounddata = struct('name',{},'type',{},'indexvalues',{});
componentdata = struct('name',{},'indexvalues',{});
% fill in component names/formulas in structure
for k=1:length(components),
    if k ~= timeindex,
        % check if componentname defines an upper or lower bound
        if ~isempty(regexp(components{k},'[+]')),
            % component defines an upper bound
            errorbounddata(end+1).name = regexprep(components{k},'\W','');
            errorbounddata(end).type = 'max';
            errorbounddata(end).indexvalues = k;
        elseif ~isempty(regexp(components{k},'[-]')),
            % component defines a lower bound
            errorbounddata(end+1).name = regexprep(components{k},'\W','');
            errorbounddata(end).type = 'min';
            errorbounddata(end).indexvalues = k;
        else
            measurementstructure.data(end+1).name = components{k};
            componentdata(end+1).name = components{k};
            componentdata(end).indexvalues = k;
        end
    end
end
    
%%%%%%%%%%%%%%%%%%%%%%%%%%%%%%%%%%%%%%%%%%%%%%%
% Get componentnotes
%%%%%%%%%%%%%%%%%%%%%%%%%%%%%%%%%%%%%%%%%%%%%%%    
% parse the componentnotes
componentnotesall = regexp(componentnotesraw,'([^\n]*)','tokens');
for k = 1:length(componentnotesall),
    componentnotesk = componentnotesall{k}{1};
    % get componentname
    index = strfind(componentnotesk,':');
    componentname = componentnotesk(1:index(1)-1);
    componentnotes = componentnotesk(index(1)+1:end);
    index = strmatchIQM(componentname,{componentdata.name},'exact');
    if isempty(index),
        errorMsg = sprintf('%sNote for component ''%s'' defined but the component does not exist.\n',errorMsg,componentname);
    else
        if ~isempty(componentnotes),
            measurementstructure.data(index).notes = strtrim(componentnotes);
        end
    end
end

%%%%%%%%%%%%%%%%%%%%%%%%%%%%%%%%%%%%%%%%%%%%%%%
% Get values and time
%%%%%%%%%%%%%%%%%%%%%%%%%%%%%%%%%%%%%%%%%%%%%%%
% delete spaces
values = regexprep(valuesraw,' ','');
% replace ",," events by ",NaN,"
values = regexprep(values,',,',',NaN,');
values = regexprep(values,',,',',NaN,');
% replace empty elements at the end of each row with ",NaN"
values = regexprep(values,',\n',',NaN\n');
valuesmatrix = eval(sprintf('[%s]',values));
% assign the time vector into the structure
measurementstructure.time = valuesmatrix(:,timeindex);
% assign the measurement data into the structure
for k=1:length(componentdata),
    measurementstructure.data(k).values = valuesmatrix(:,componentdata(k).indexvalues);
end
% assign the error bound data if present (and corresponding component
% present too ... otherwise warning).
for k=1:length(errorbounddata),
    indexcomponent = strmatchIQM(errorbounddata(k).name,{componentdata.name},'exact');
    if isempty(indexcomponent),
        warning('Component ''%s'' has given error bound but does not exist in the data file.',errorbounddata(k).name);
    else
        if strcmp(errorbounddata(k).type,'max'),
            measurementstructure.data(indexcomponent).maxvalues = valuesmatrix(:,errorbounddata(k).indexvalues);
        else
            measurementstructure.data(indexcomponent).minvalues = valuesmatrix(:,errorbounddata(k).indexvalues);
        end
    end
end

%%%%%%%%%%%%%%%%%%%%%%%%%%%%%%%%%%%%%%%%%%%%%%%
% Fill the nonavailable errorbounds with NaN
%%%%%%%%%%%%%%%%%%%%%%%%%%%%%%%%%%%%%%%%%%%%%%%
for k=1:length(measurementstructure.data),
    if isempty(measurementstructure.data(k).minvalues) || isempty(measurementstructure.data(k).maxvalues),
        measurementstructure.data(k).maxvalues = NaN(size(measurementstructure.data(k).values));
        measurementstructure.data(k).minvalues = NaN(size(measurementstructure.data(k).values));
    end
end
return
```
