## Supplementary material for "A systems biology analysis of adrenergically stimulated adiponectin exocytosis in white adipocytes": All scripts used in this work: IQMimportXLSmeasurement.m

```
function [output,errorMsg] = IQMimportXLSmeasurement(filename)
% IQMimportXLSmeasurement
% Imports experimental measurement data stored in an XLS Excel file
% into the measurementstructure used by the IQMmeasurement object.
% Please note that a special format of the excel data is required. 
% This format is explained in the user's reference manual and example 
% files can be found in the IQMlite/examples folder.
% 
% filename: name of the .xls file containing the data
%
% output: cell-array with data structures used by
%         IQMmeasurement object  (empty if error occurred).
%         One element of the cell-array corresponds to one
%         sheet in the excel file.
% errorMsg: string containing possible error messages. 

% <<<COPYRIGHTSTATEMENT - IQM TOOLS LITE>>>


%%%%%%%%%%%%%%%%%%%%%%%%%%%%%%%%%%%%%%%%%%%%%%%
% INITIALIZATION
%%%%%%%%%%%%%%%%%%%%%%%%%%%%%%%%%%%%%%%%%%%%%%%
errorMsg = '';
measurementstructure = struct(IQMmeasurement());

%%%%%%%%%%%%%%%%%%%%%%%%%%%%%%%%%%%%%%%%%%%%%%%
% Determine number of sheets
%%%%%%%%%%%%%%%%%%%%%%%%%%%%%%%%%%%%%%%%%%%%%%%
[typ, dsc] = xlsfinfo(filename);
nrsheets = length(dsc);

%%%%%%%%%%%%%%%%%%%%%%%%%%%%%%%%%%%%%%%%%%%%%%%
% Process one sheet at a time
%%%%%%%%%%%%%%%%%%%%%%%%%%%%%%%%%%%%%%%%%%%%%%%
output = {};
for sheet=1:nrsheets,
    % Read the current sheet
    warning off;
    [NUMERIC,TXT,RAW] = xlsread(filename,dsc{sheet});
    warning on;
    % Check if current sheet is a valid measurement sheet
    % to be that it requires 'Name' in A1.
    if iscell(RAW),
        testSheet = RAW{1,1};
        if ischar(testSheet),
            if strcmp(strtrim(lower(RAW{1,1})),'name'),
                % Sheet is valid (probably ;))
                % Process the raw information and fill in the measurement data
                % structure
                [output{end+1}, errorMsg] = processData(RAW,sheet,errorMsg);
            end
        end
    end
end
return

%%%%%%%%%%%%%%%%%%%%%%%%%%%%%%%%%%%%%%%%%%%%%%%%%%%%%%%%%%%%%%%%%%%%%%%%%%%
% Process RAW Data
%%%%%%%%%%%%%%%%%%%%%%%%%%%%%%%%%%%%%%%%%%%%%%%%%%%%%%%%%%%%%%%%%%%%%%%%%%%
function [measurementstructure,errorMsg] = processData(RAW,sheet,errorMsg)
% initialize empty measurement structure
measurementstructure = struct(IQMmeasurement());
% get size of RAW matrix
[nrows, ncols] = size(RAW);
% each identifier needs to appear but only once!
% furthermore, the identifieres need to appear in the correct order!
rowName = 0;
rowNotes = 0;
rowComponentnotes = 0;
rowComponents = 0;
rowValues = 0;
%%%%%%%%%%%%%%%%%%%%%%%%%%%%%%%%%%%%%%%%%%%%%%%
% get rows of identifiers and check the order
%%%%%%%%%%%%%%%%%%%%%%%%%%%%%%%%%%%%%%%%%%%%%%%
for row=1:nrows,
    if ~isnan(RAW{row,1}),
        if strcmp(strtrim(lower(RAW{row,1})),'name'),
            rowName = row;
            if rowNotes+rowComponents+rowComponentnotes+rowValues ~= 0,
                errorMsg = sprintf('%sIdentifier ''Name'' in sheet %d does not come in correct order.\n',errorMsg,sheet);
            end
        end
        if strcmp(strtrim(lower(RAW{row,1})),'notes'),
            rowNotes = row;
            if rowComponents+rowComponentnotes+rowValues ~= 0,
                errorMsg = sprintf('%sIdentifier ''Notes'' in sheet %d does not come in correct order.\n',errorMsg,sheet);
            end
        end
        if strcmp(strtrim(lower(RAW{row,1})),'componentnotes'),
            rowComponentnotes = row;
            if rowComponents + rowValues ~= 0,
                errorMsg = sprintf('%sIdentifier ''Componentnotes'' in sheet %d does not come in correct order.\n',errorMsg,sheet);
            end
        end
        if strcmp(strtrim(lower(RAW{row,1})),'components'),
            rowComponents = row;
            if rowValues ~= 0,
                errorMsg = sprintf('%sIdentifier ''Components'' in sheet %d does not come in correct order.\n',errorMsg,sheet);
            end
        end
        if strcmp(strtrim(lower(RAW{row,1})),'values'),
            rowValues = row;
        end
        % check if all identifiers found then break the loop
        if rowName*rowNotes*rowComponents*rowComponentnotes*rowValues ~= 0,
            break;
        end
    end
end
% check if all identifiers present
if rowName*rowNotes*rowComponents*rowComponentnotes*rowValues == 0,
    errorMsg = sprintf('%sAt least one identifier is missing in in sheet %d.\n',errorMsg,sheet);
end

%%%%%%%%%%%%%%%%%%%%%%%%%%%%%%%%%%%%%%%%%%%%%%%
% Name
%%%%%%%%%%%%%%%%%%%%%%%%%%%%%%%%%%%%%%%%%%%%%%%
% Name only one line but several columns
name = '';
for col=2:ncols,
    if ~isnan(RAW{rowName,col}),
        if ischar(RAW{rowName,col}),
            name = sprintf('%s %s',name,RAW{rowName,col});
        end
    else
        break;
    end
end
measurementstructure.name = strtrim(name);

%%%%%%%%%%%%%%%%%%%%%%%%%%%%%%%%%%%%%%%%%%%%%%%
% Notes
%%%%%%%%%%%%%%%%%%%%%%%%%%%%%%%%%%%%%%%%%%%%%%%
% Notes several lines and several columns
notes = '';
spaceYes = 1;
for row=rowNotes:rowComponentnotes-1,
    for col=2:ncols,
        if ~isnan(RAW{row,col}),
            if ischar(RAW{row,col}),
                if spaceYes,
                    notes = sprintf('%s %s',notes,strtrim(RAW{row,col}));
                else
                    notes = sprintf('%s%s',notes,strtrim(RAW{row,col}));
                    spaceYes = 1;
                end
            end
        end
    end
    notes = sprintf('%s\n',strtrim(notes)); 
    spaceYes = 0;
end
measurementstructure.notes = notes;

%%%%%%%%%%%%%%%%%%%%%%%%%%%%%%%%%%%%%%%%%%%%%%%
% Components
%%%%%%%%%%%%%%%%%%%%%%%%%%%%%%%%%%%%%%%%%%%%%%%
components = {};
for col = 2:ncols,
    if ~isnan(RAW{rowComponents,col}),
        components{end+1} = regexprep(RAW{rowComponents,col},' ','');
    else
        break;
    end
end
% find 'time' component
timeindex = strmatchIQM('time',components,'exact');
% initialize help structure for min max values (error bounds) + components
errorbounddata = struct('name',{},'type',{},'indexvalues',{});
componentdata = struct('name',{},'indexvalues',{});
% fill in component names/formulas in structure
for k=1:length(components),
    if k ~= timeindex,
        % check if componentname defines an upper or lower bound
        if ~isempty(regexp(components{k},'[+]')),
            % component defines an upper bound
            errorbounddata(end+1).name = regexprep(components{k},'\W','');
            errorbounddata(end).type = 'max';
            errorbounddata(end).indexvalues = k;
        elseif ~isempty(regexp(components{k},'[-]')),
            % component defines a lower bound
            errorbounddata(end+1).name = regexprep(components{k},'\W','');
            errorbounddata(end).type = 'min';
            errorbounddata(end).indexvalues = k;
        else
            measurementstructure.data(end+1).name = components{k};
            componentdata(end+1).name = components{k};
            componentdata(end).indexvalues = k;
        end
    end
end

%%%%%%%%%%%%%%%%%%%%%%%%%%%%%%%%%%%%%%%%%%%%%%%
% Componentnotes
%%%%%%%%%%%%%%%%%%%%%%%%%%%%%%%%%%%%%%%%%%%%%%%
index = 1;
for k = 1:length(componentdata)
    col = componentdata(k).indexvalues+1;
    if ~isnan(RAW{rowComponentnotes,col}),
        if ischar(RAW{rowComponentnotes,col}),
            componentnotes = RAW{rowComponentnotes,col};
            measurementstructure.data(k).notes = strtrim(componentnotes);
        end
    else
        measurementstructure.data(k).notes = '';
    end
end

%%%%%%%%%%%%%%%%%%%%%%%%%%%%%%%%%%%%%%%%%%%%%%%
% Values and errorbounds
%%%%%%%%%%%%%%%%%%%%%%%%%%%%%%%%%%%%%%%%%%%%%%%
% get values matrix
try
    valuesmatrix = cell2mat(RAW(rowValues:nrows,2:length(components)+1));
catch
    error('Please check the datatypes in the values section of the Excel file.');
end
time = valuesmatrix(:,timeindex);
% check for first occurrence of NaN in time vector ... then cut off.
indexNaN = find(isnan(time)==1);
if isempty(indexNaN),
    numbertimesteps = length(time);
else
    numbertimesteps = indexNaN(1)-1;
end
measurementstructure.time = time(1:numbertimesteps);
% assign the measurement data into the structure
% assign the measurement data into the structure
for k=1:length(componentdata),
    measurementstructure.data(k).values = valuesmatrix(1:numbertimesteps,componentdata(k).indexvalues);
end
% assign the error bound data if present (and corresponding component
% present too ... otherwise warning).
for k=1:length(errorbounddata),
    indexcomponent = strmatchIQM(errorbounddata(k).name,{componentdata.name},'exact');
    if isempty(indexcomponent),
        warning('Component ''%s'' has given error bound but does not exist in the data file.',errorbounddata(k).name);
    else
        if strcmp(errorbounddata(k).type,'max'),
            measurementstructure.data(indexcomponent).maxvalues = valuesmatrix(1:numbertimesteps,errorbounddata(k).indexvalues);
        else
            measurementstructure.data(indexcomponent).minvalues = valuesmatrix(1:numbertimesteps,errorbounddata(k).indexvalues);
        end
    end
end

%%%%%%%%%%%%%%%%%%%%%%%%%%%%%%%%%%%%%%%%%%%%%%%
% Fill the nonavailable errorbounds with NaN
%%%%%%%%%%%%%%%%%%%%%%%%%%%%%%%%%%%%%%%%%%%%%%%
for k=1:length(measurementstructure.data),
    if isempty(measurementstructure.data(k).minvalues) || isempty(measurementstructure.data(k).maxvalues),
        measurementstructure.data(k).maxvalues = NaN(size(measurementstructure.data(k).values));
        measurementstructure.data(k).minvalues = NaN(size(measurementstructure.data(k).values));
    end
end
return
```
