## Supplementary material for "A systems biology analysis of adrenergically stimulated adiponectin exocytosis in white adipocytes": All scripts used in this work: IQMmeasurementdata.m

```
function [time, componentNames, values, minvalues, maxvalues] = IQMmeasurementdata(measurement)
% IQMmeasurementdata
% This functions allows to extract information from an IQMmeasurement structure
% 
% USAGE:
% ======
% [time, componentNames] = IQMmeasurementdata(measurement)
% [time, componentNames, values] = IQMmeasurementdata(measurement)
% [time, componentNames, values,minvalues,maxvalues] = IQMmeasurementdata(measurement)
% 
% measurement: IQMmeasurement object
%
% Output Arguments:
% =================
% time: time vector of all measurement instants
% componentNames: cell-array containing the names of the measured components
% values: matrix containing all the measurements of the components.
%   One row per time instant and one column per measured component.
%   The ordering of the columns corresponds to the ordering of the names in
%   the "componentNames" output variable. Non measured elements are set to
%   NaN (not a number).
% minvalues: matrix containing all the min values for the measured components.
%   One row per time instant and one column per measured component.
%   The ordering of the columns corresponds to the ordering of the names in
%   the "componentNames" output variable. Non measured elements are set to
%   NaN (not a number).
% maxvalues: matrix containing all the max values for the measured components.
%   One row per time instant and one column per measured component.
%   The ordering of the columns corresponds to the ordering of the names in
%   the "componentNames" output variable. Non measured elements are set to
%   NaN (not a number).

% <<<COPYRIGHTSTATEMENT - IQM TOOLS LITE>>>

if ~isIQMmeasurement(measurement),
    error('Input argument is not an IQMmeasurement.');
end
measurement = struct(measurement);

%%%%%%%%%%%%%%%%%%%%%%%%%%%%%%%%%%%%%%%%%%%%%%%
% CHECK IF MODEL CONTAINS MEASUREMENTS
%%%%%%%%%%%%%%%%%%%%%%%%%%%%%%%%%%%%%%%%%%%%%%%
if length(measurement.data) == 0,
    error('The model does not contain any measurements');
end

%%%%%%%%%%%%%%%%%%%%%%%%%%%%%%%%%%%%%%%%%%%%%%%
% GET INFORMATION
%%%%%%%%%%%%%%%%%%%%%%%%%%%%%%%%%%%%%%%%%%%%%%%
% get time vector
time = measurement.time;
% get component names
componentNames = {};
values = [];
minvalues = [];
maxvalues = [];
for k=1:length(measurement.data),
    componentNames{end+1} = measurement.data(k).name;
    values(:,end+1) = measurement.data(k).values;
    minvalues(:,end+1) = measurement.data(k).minvalues;
    maxvalues(:,end+1) = measurement.data(k).maxvalues;
end
return
```
