## Supplementary material for "A systems biology analysis of adrenergically stimulated adiponectin exocytosis in white adipocytes": All scripts used in this work: IQMvisualizemeasurement.m

```
function [varargout] = IQMvisualizemeasurement(measurement)
% IQMvisualizemeasurement
% Function allowing to visualize the content of an IQMmeasurement object. 
% The function just prepares the data. Display is then realized using
% the IQMplot function.
% 
% USAGE:
% ======
% [] = IQMvisualizemeasurement(measurement)
% [output] = IQMvisualizemeasurement(measurement)
%
% measurement: IQMmeasurement object containing the data
%
% Output Arguments:
% =================
% If an output argument is specified, the data are not plotted, but a
% structure is returned that can be used as input argument for IQMplot to
% show the data.

% <<<COPYRIGHTSTATEMENT - IQM TOOLS LITE>>>

% convert IQMmeasurement object to struct
measurement = IQMstruct(measurement);

%%%%%%%%%%%%%%%%%%%%%%%%%%%%%%%%%%%%%%%%%%%%%%%
% CHECK IF MODEL CONTAINS MEASUREMENTS
%%%%%%%%%%%%%%%%%%%%%%%%%%%%%%%%%%%%%%%%%%%%%%%
if length(measurement.data) == 0,
    error('The IQMmeasurement object does not contain any measurements.');
end

%%%%%%%%%%%%%%%%%%%%%%%%%%%%%%%%%%%%%%%%%%%%%%%
% GET TIME VECTOR
%%%%%%%%%%%%%%%%%%%%%%%%%%%%%%%%%%%%%%%%%%%%%%%
time = measurement.time;

%%%%%%%%%%%%%%%%%%%%%%%%%%%%%%%%%%%%%%%%%%%%%%%
% PROCESS THE DATA INTO A MATRIX
%%%%%%%%%%%%%%%%%%%%%%%%%%%%%%%%%%%%%%%%%%%%%%%
% also construct dataNames and legendtext data
dataNames = {};
dataMatrix = NaN*ones(length(measurement.time),length(measurement.data));
timeMatrix = dataMatrix;
dataErrorindices = [];
dataMinvalues = [];
dataMaxvalues = [];

legendtext = {};
for k = 1:length(measurement.data),
    name = measurement.data(k).name;
    dataNames{k} = name;
    if ~isempty(measurement.data(k).notes),
        legendtext{k} = sprintf('%s (%s)',name,measurement.data(k).notes);
    else
        legendtext{k} = sprintf('%s',name);
    end
    dataComponent = measurement.data(k).values;
    timeComponent = measurement.time;
    dataComponent = dataComponent;
    timeComponent = timeComponent;
    dataMatrix(1:length(dataComponent),k) = dataComponent;
    timeMatrix(1:length(timeComponent),k) = timeComponent;

    % Process error bounds if present (only if both are present)
    if ~isempty(measurement.data(k).maxvalues) && ~isempty(measurement.data(k).minvalues),
        if length(measurement.data(k).minvalues) ~= length(measurement.data(k).maxvalues),
            warning('Measurement ''%s'' does have different numbers of max and min bounds.',measurement.data(k).name);
        else
            dataErrorindices(end+1) = k;
            dataMinvalues(1:length(measurement.data(k).minvalues),end+1) = measurement.data(k).minvalues;
            dataMaxvalues(1:length(measurement.data(k).maxvalues),end+1) = measurement.data(k).maxvalues;
        end
    end
end

%%%%%%%%%%%%%%%%%%%%%%%%%%%%%%%%%%%%%%%%%%%%%%%
% Handle variable output arguments
%%%%%%%%%%%%%%%%%%%%%%%%%%%%%%%%%%%%%%%%%%%%%%%
% Keep the commented part as long as it works!
if nargout==0,
    % plot
%    IQMplot(timeMatrix,dataMatrix,dataNames,dataErrorindices,dataMinvalues,dataMaxvalues,legendtext,'*:',measurement.name);
    IQMplot(timeComponent,dataMatrix,dataNames,dataErrorindices,dataMinvalues,dataMaxvalues,legendtext,'*',measurement.name);
elseif nargout == 1,
%    varargout{1} = createdatastructIQMplotIQM(timeMatrix,dataMatrix,dataNames,dataErrorindices,dataMinvalues,dataMaxvalues,legendtext,'*:',measurement.name);
    varargout{1} = createdatastructIQMplotIQM(timeComponent,dataMatrix,dataNames,dataErrorindices,dataMinvalues,dataMaxvalues,legendtext,'*',measurement.name);
else
    error('Incorrect number of output arguments.');
end
return
```
