## Supplementary material for "A systems biology analysis of adrenergically stimulated adiponectin exocytosis in white adipocytes": All scripts used in this work: isIQMmeasurement.m

```
function [output] = isIQMmeasurement(input)
% isIQMmeasurement: check if input argument is an IQMmeasurement.

% <<<COPYRIGHTSTATEMENT - IQM TOOLS LITE>>>

output = strcmp(class(input),'IQMmeasurement');
```
