## Supplementary material for "A systems biology analysis of adrenergically stimulated adiponectin exocytosis in white adipocytes": All scripts used in this work: IQMalgebraic.m

```
function [names,formulas] = IQMalgebraic(model)
% IQMalgebraic: Returns information about the algebraic equations in a model.
%
% USAGE:
% ======
% [names,formulas] = IQMalgebraic(model)
%
% model: IQMmodel or m-file ODE description of model
%
% Output Arguments:
% =================
% names: cell-array with names of the variables that are determined using
% algebraic equations.
% formulas: cell-array with right hand side formula for the algebraic rules

% <<<COPYRIGHTSTATEMENT - IQM TOOLS LITE>>>

%%%%%%%%%%%%%%%%%%%%%%%%%%%%%%%%%%%%%%%%%%%%%%%
% PROCESS IQMMODEL OR ODE FILE
%%%%%%%%%%%%%%%%%%%%%%%%%%%%%%%%%%%%%%%%%%%%%%%
if isIQMmodel(model),
    iqm = IQMstruct(model);
    names = {iqm.algebraic.name};
    formulas = {iqm.algebraic.formula};
else
    names = feval(model,'algebraic');
    formulas = {};
end
names = names(:);
formulas = formulas(:);
return
```
