## Supplementary material for "A systems biology analysis of adrenergically stimulated adiponectin exocytosis in white adipocytes": All scripts used in this work: IQMcalcICvector.m

```
function [output] = IQMcalcICvector(model,varargin)
% IQMcalcICvector: This function determines an IC vector for models with
% non-numeric initial conditions. These can depend on other initial
% conditions and on the model parameters. If no new initial conditions or
% parameters are specified as input arguments to the function the nominal
% values are used that are stored in the model. Otherwise the information
% in the additional input arguments is used to determine the new initial
% conditions. For models with only numeric initial conditions either the
% ones stored in the model are returned or the ICs provided as input
% argument are returned. The definition of a parameter vector as input
% argument will not have an effect in this case.
%
% USAGE:
% ======
% [output] = IQMcalcICvector(model)
% [output] = IQMcalcICvector(model,IC)
% [output] = IQMcalcICvector(model,IC,parametervector)
%
% model: IQMmodel, ODE, or MEX file model description
% IC: cell-array with statenames for which to give back the initial
%   conditions.
% parametervector: vector with initial conditions to update the model with
%
% Output Arguments:
% =================
% output: determined numeric initial condition vector

% <<<COPYRIGHTSTATEMENT - IQM TOOLS LITE>>>

%%%%%%%%%%%%%%%%%%%%%%%%%%%%%%%%%%%%%%%%%%%%%%%
% EXTRACT SOME INFO FROM THE MODEL
%%%%%%%%%%%%%%%%%%%%%%%%%%%%%%%%%%%%%%%%%%%%%%%
% get the models ICs and statenames
if isIQMmodel(model),
    ms = struct(model);
    if hasonlynumericICsIQM(model),
        mics = [ms.states.initialCondition];
    else
        mics = {ms.states.initialCondition};
    end
    sn = {ms.states.name};    
else
    % assume model is an ODE or MEX file
    % We need to handle ODE and MEX files differently, since ODE files can use 
    % algebraic expressions, but MEX files can not and this translates to different lengths in the IC vector
    
    % Handling MEX models
    mics = feval(model);
    sn = feval(model,'states'); 
    
    % Append some info if using ODE models and algebraic equations present
    try
        % only working for ODE models -  in case of MEX models an error will occurr
        an = feval(model,'algebraic');
        for ii = 1:length(an)
            sn{end+ii} = an{ii};
        end
    catch
        % do nothing in case of MEX models
    end
end
% get the models parameter values
[pn,pv] = IQMparameters(model);
[vn,vf] = IQMvariables(model);

%%%%%%%%%%%%%%%%%%%%%%%%%%%%%%%%%%%%%%%%%%%%%%%
% HANDLE VARIABLE INPUT ARGUMENTS
%%%%%%%%%%%%%%%%%%%%%%%%%%%%%%%%%%%%%%%%%%%%%%%
if nargin == 1,
    IC = [];
    % use the models nominal parameters
    parametervector = pv;
elseif nargin == 2,
    IC = varargin{1};
    % use the models nominal parameters
    parametervector = pv;
elseif nargin == 3,
    IC = varargin{1};
    parametervector = varargin{2};
else
    error('Incorrect number of input arguments.');
end

%%%%%%%%%%%%%%%%%%%%%%%%%%%%%%%%%%%%%%%%%%%%%%%
% CHECK INPUTS (IC and parametervector)
%%%%%%%%%%%%%%%%%%%%%%%%%%%%%%%%%%%%%%%%%%%%%%%
if ~isempty(IC) && length(IC) ~= length(mics),
    error('Length of initial condition vector does not fit the number of states in the model.');
end
if ~isempty(parametervector) && length(parametervector) ~= length(pv),
    error('Length of parameter vector does not fit the number of parameters in the model.');
end
if isempty(parametervector),
    parametervector = pv;
end

%%%%%%%%%%%%%%%%%%%%%%%%%%%%%%%%%%%%%%%%%%%%%%%
% HANDLE MODEL WITH NUMERIC ICs
%%%%%%%%%%%%%%%%%%%%%%%%%%%%%%%%%%%%%%%%%%%%%%%
if hasonlynumericICsIQM(model),
    if isempty(IC),
        % return the nominal initial condition vector in the model
        output = mics(:);
        return
    else
        % return the initial condition vector provided as input argument
        output = IC(:);
        return
    end
end

%%%%%%%%%%%%%%%%%%%%%%%%%%%%%%%%%%%%%%%%%%%%%%%
% HANDLE MODEL WITH NON NUMERIC ICs
%%%%%%%%%%%%%%%%%%%%%%%%%%%%%%%%%%%%%%%%%%%%%%%
% mics is a cell-array, IC a double vector either empty or with new values.
% the ones corresponding to non-numeric ICs are neglected. 
% 1) update the mics vector by merging with the IC vector
if ~isempty(IC),
    for k=1:length(mics),
        % do not change the non-numeric ICs
        if isnumeric(mics{k}),
            mics{k} = IC(k);
        end
    end
end
% 2) From now on we only need to handle mics and parametervector. For 
% uniqueness "parametervector" is renamed to "pv_1a2b3c" and pn to 
% "pn_1a2b3c", etc.
pv_1a2b3c = parametervector;
pn_1a2b3c = pn;
vn_1a2b3c = vn;
vf_1a2b3c = vf;
mics_1a2b3c = mics;
sn_1a2b3c = sn;
% 3) define all model parameters (nominal or provided) in the workspace
for k_1a2b3c = 1:length(pv_1a2b3c),
    eval(sprintf('%s = pv_1a2b3c(k_1a2b3c);',pn_1a2b3c{k_1a2b3c}));
end
allDefined = 0;
ic_1a2b3c = [];
count = 0;
while ~allDefined && count<10,
    count = count + 1;
    % 4) define all model variables in the workspace
    for k_1a2b3c = 1:length(vn_1a2b3c),
        try
            eval(sprintf('%s = %s;',vn_1a2b3c{k_1a2b3c},vf_1a2b3c{k_1a2b3c}));
            allDefined = 1;            
        catch
            allDefined = 0;
        end
    end
    % 5) determine the output initial condition vector by evaluating all cells
    % in mics (in the order of mics).

    for k_1a2b3c = 1:length(mics_1a2b3c),
        if isnumeric(mics_1a2b3c{k_1a2b3c}),
            % if ic entry is numeric then copy it in ic_1a2b3c and
            % define the corresponding state in the workspace
            ic_1a2b3c(k_1a2b3c) = mics_1a2b3c{k_1a2b3c};
            eval(sprintf('%s = ic_1a2b3c(end);',sn_1a2b3c{k_1a2b3c}));
        else
            % if ic entry is non-numeric then evaluate this entry, store it in
            % the ic_1a2b3c vector and define the corresponding state in the
            % workspace
            try
                ic_1a2b3c(k_1a2b3c) = eval(mics_1a2b3c{k_1a2b3c});
                eval(sprintf('%s = ic_1a2b3c(end);',sn_1a2b3c{k_1a2b3c}));
                allDefined = 1;
            catch
                allDefined = 0;                
                %             error('Problem with non-numeric initial condition for state ''%s''.\nNOTE: non-numeric ICs are evaluated in a certain order:\n  1) TEXT-models:   order given by the ordering of the ODEs.\n  2) TEXTBC-models: order given by the ordering of the initial conditions.',sn_1a2b3c{k_1a2b3c});
            end
        end
    end
end
if ~allDefined,
    error('Trouble finding numeric initial conditions.');
end
% Its done, just return the calculated ic vector
output = ic_1a2b3c;
end
```
