## Supplementary material for "A systems biology analysis of adrenergically stimulated adiponectin exocytosis in white adipocytes": All scripts used in this work: IQMconvert2MA.m

```
function [MAstructure] = IQMconvert2MA(model)
% IQMconvert2MA: Takes an IQMmodel and if possible returns the stoichiometric
% matrix, the numeric kinetic parameters, and the initial conditions. The
% IQMmodel is not allowed to contain variables, functions, events, or
% functionsMATLAB. All right hand sides of the ODEs have to be defined in
% terms of reaction names. All reactions need to be irreversible and mass action type! 
%
% Please note that the function is NOT parsing the reaction kinetics to
% check if they are correct mass action kinetic rate laws. The user needs
% to make sure that this is the case. Examples of correct expressions are:
%   Reaction = k * species1 * species2
%   Reaction = species1 * species2 * k
%   Reaction = k * species1
%   Reaction = 3.141592 * species1^2
%   Reaction = species1^2
% The kinetic rate constant is then simply determined by setting all
% species concentrations to 1 and evaluating the rate expressions.
%
% USAGE:
% ======
% [MAstructure] = IQMconvert2MA(model) 
%
% model: IQMmodel 
%
% Output Arguments:
% =================
% MAstructure: structure containing information about the MA model
%          MAstructure.N: stoichiometric matrix
%          MAstructure.L: reactant stoichiometric matrix
%          MAstructure.kineticParameters: vector containing the kinetic
%               parameters for each reaction
%          MAstructure.initialConditions: vector of initial conditions for all
%               species
%          MAstructure.species: cell-array with species names
%          MAstructure.reactions: cell-array with reaction names

% <<<COPYRIGHTSTATEMENT - IQM TOOLS LITE>>>

%%%%%%%%%%%%%%%%%%%%%%%%%%%%%%%%%%%%%%%%%%%%%%%
% HANDLE NON-NUMERIC INITIAL CONDITIONS
%%%%%%%%%%%%%%%%%%%%%%%%%%%%%%%%%%%%%%%%%%%%%%%
% just by replacing them
model = IQMconvertNonNum2NumIC(model);

%%%%%%%%%%%%%%%%%%%%%%%%%%%%%%%%%%
% Process the model
%%%%%%%%%%%%%%%%%%%%%%%%%%%%%%%%%%
% make all reactions irreversible (if necessary)
modelirr = IQMmakeirreversible(model);
% take away parenthesis around backward reaction expressions
ms = struct(modelirr);
for k = 1:length(ms.reactions),
    ms.reactions(k).formula = strrep(ms.reactions(k).formula,'(','');
    ms.reactions(k).formula = strrep(ms.reactions(k).formula,')','');
end
model = IQMmodel(ms);
IQMstructure = IQMstruct(model);

%%%%%%%%%%%%%%%%%%%%%%%%%%%%%%%%%%%%%%%%%%%%%%%
% CHECK IF NON DESIRED ELEMENTS ARE PRESENT IN THE MODEL
%%%%%%%%%%%%%%%%%%%%%%%%%%%%%%%%%%%%%%%%%%%%%%%
% undesired: 
%            - functions
%            - variables
%            - events
%            - functionsMATLAB
%            - states defined by rule
% check functions, variables, events, functionsMATLAB
if ~isempty(IQMstructure.functions),
    error('The model contains functions. This is not allowed when converting it to an MA representation.');
end
if ~isempty(IQMstructure.variables),
    error('The model contains variables. This is not allowed when converting it to an MA representation.');
end
if ~isempty(IQMstructure.events),
    error('The model contains events. This is not allowed when converting it to an MA representation.');
end
if ~isempty(IQMstructure.functionsMATLAB),
    error('The model contains MATLAB functions. This is not allowed when converting it to an MA representation.');
end

%%%%%%%%%%%%%%%%%%%%%%%%%%%%%%%%%%%%%%%%%%%%%%%
% CHECK FOR REVERSIBLE REACTIONS (NOT ALLOWED)
%%%%%%%%%%%%%%%%%%%%%%%%%%%%%%%%%%%%%%%%%%%%%%%
[reactionNames, reactionFormulas] = IQMreactions(model);

%%%%%%%%%%%%%%%%%%%%%%%%%%%%%%%%%%%%%%%%%%%%%%%
% GET THE (REACTANT) STOICHIOMETRIC MATRIX 
%%%%%%%%%%%%%%%%%%%%%%%%%%%%%%%%%%%%%%%%%%%%%%%
% also check if states are defined by rules, that is, not by summation of
% reaction terms. easily done by checking the stoichiometric matrix and 
% comparing number of components in it and number of all states in the
% model.
allStates = IQMstates(model);
[N,componentNames,reactionNames,reversibleFlag] = IQMstoichiometry(model);
if length(allStates) ~= length(componentNames),
    error('Not all ODEs are expressed in terms of reactions. The full stoichiometric matrix can thus not be determined.');
end
L = IQMreactantstoichiometry(model);

%%%%%%%%%%%%%%%%%%%%%%%%%%%%%%%%%%%%%%%%%%%%%%%
% GET THE KINETIC PARAMETERS
%%%%%%%%%%%%%%%%%%%%%%%%%%%%%%%%%%%%%%%%%%%%%%%
% do it very easy ... just set all concentrations to 1 and evaluate the
% rate expressions. Under the required format for the rate expressions this
% gives the kinetic parameters.
[x,y,z,a,kineticParameters] = IQMreactions(model,ones(1,length(allStates)));

%%%%%%%%%%%%%%%%%%%%%%%%%%%%%%%%%%%%%%%%%%%%%%%
% GET THE INITIAL CONDITIONS
%%%%%%%%%%%%%%%%%%%%%%%%%%%%%%%%%%%%%%%%%%%%%%%
initialconditions = IQMinitialconditions(model);

%%%%%%%%%%%%%%%%%%%%%%%%%%%%%%%%%%%%%%%%%%%%%%%
% FORMAT OUTPUT VALUES
%%%%%%%%%%%%%%%%%%%%%%%%%%%%%%%%%%%%%%%%%%%%%%%
MAstructure.N = N;
MAstructure.L = L;
MAstructure.kineticParameters = kineticParameters(:);
MAstructure.initialConditions = initialconditions(:);
MAstructure.species = allStates;
MAstructure.reactions = {IQMstructure.reactions.name}';
%%%%%%%%%%%%%%%%%%%%%%%%%%%%%%%%%%%%%%%%%%%%%%%
% RETURN
%%%%%%%%%%%%%%%%%%%%%%%%%%%%%%%%%%%%%%%%%%%%%%%
return
```
