## Supplementary material for "A systems biology analysis of adrenergically stimulated adiponectin exocytosis in white adipocytes": All scripts used in this work: IQMconvertNonNum2NumIC.m

```
function [newmodel] = IQMconvertNonNum2NumIC(model)
% IQMconvertNonNum2NumIC: This function converts non-numeric iniital
% conditions to numeric initial conditions in the model and returns the
% updated model. Models with numeric ICs are unaffected.
%
% USAGE:
% ======
% [newmodel] = IQMconvertNonNum2NumIC(model)
%
% model: IQMmodel
%
% Output Arguments:
% =================
% newmodel: IQMmodel where non-numeric ICs have been replaced by numeric ICs

% <<<COPYRIGHTSTATEMENT - IQM TOOLS LITE>>>

%%%%%%%%%%%%%%%%%%%%%%%%%%%%%%%%%%%%%%%%%%%%%%%
% CHECK
%%%%%%%%%%%%%%%%%%%%%%%%%%%%%%%%%%%%%%%%%%%%%%%
if ~isIQMmodel(model),
    error('Input argument needs to be an IQMmodel.');
end

%%%%%%%%%%%%%%%%%%%%%%%%%%%%%%%%%%%%%%%%%%%%%%%
% NUMERIC ICs ONLY
%%%%%%%%%%%%%%%%%%%%%%%%%%%%%%%%%%%%%%%%%%%%%%%
if hasonlynumericICsIQM(model),
    newmodel = model;
    return
end

%%%%%%%%%%%%%%%%%%%%%%%%%%%%%%%%%%%%%%%%%%%%%%%
% NON-NUMERIC ICs
%%%%%%%%%%%%%%%%%%%%%%%%%%%%%%%%%%%%%%%%%%%%%%%
numICs = IQMcalcICvector(model);
ms = struct(model);
for k=1:length(ms.states),
    ms.states(k).initialCondition = numICs(k);
end
newmodel = IQMmodel(ms);
```
