## Supplementary material for "A systems biology analysis of adrenergically stimulated adiponectin exocytosis in white adipocytes": All scripts used in this work: IQMcreateODEfile.m

```
function IQMcreateODEfile(varargin)
% IQMcreateODEfile: creates an m-file that can be simulated by 
% using standard MATLAB integrators (ODE45, ODE23s, etc.)
%
% USAGE:
% ======
% [] = IQMcreateODEfile(iqm)         
% [] = IQMcreateODEfile(iqm,filename)
% [] = IQMcreateODEfile(iqm,filename,dataFileFlag)
% [] = IQMcreateODEfile(iqm,filename,dataFileFlag,eventFlag)
% [] = IQMcreateODEfile(iqm,filename,dataFileFlag,eventFlag,augmKernelName)
%
% iqm: IQMmodel to convert to an ODE file description
% filename: filename for the created ODE file (without extension)
% dataFileFlag: =1: creates an additional file allowing to determine
%                   variable and reaction rate values for given state values 
%                   and time vector.
%               =0: doe not create the additional file
% eventFlag: =1: creates 2 additional files for the handling of events in the model. 
%            =0: does not create these two files.
%            THE EVENT FILES ARE ONLY CREATED IN CASE THAT EVENTS ARE
%            PRESENT IN THE MODEL
% augmKernelName: needed for the simulation of fast reactions. It is the
%           name of the global variable in which the mass matrix is
%           defined. The name comes from the fact that the null space of
%           the fast stoichiometric matrix is contained in the mass matrix.
%            
% DEFAULT VALUES:
% ===============
% filename: constructed from the models name
% dataFileFlag: 0
% eventFlag: 0
% augmKernelName: ''

% <<<COPYRIGHTSTATEMENT - IQM TOOLS LITE>>>

%%%%%%%%%%%%%%%%%%%%%%%%%%%%%%%%%%%%%%%%%%%%%%%
% GENERATE DELAY BASE NAME
%%%%%%%%%%%%%%%%%%%%%%%%%%%%%%%%%%%%%%%%%%%%%%%
[dummy,delaybase] = fileparts(tempname);
delaybase = char([double('delaybase_') double(delaybase(1:8))]);

%%%%%%%%%%%%%%%%%%%%%%%%%%%%%%%%%%%%%%%%%%%%%%%
% PROCESS VARIABLE INPUT ARGUMENTS
%%%%%%%%%%%%%%%%%%%%%%%%%%%%%%%%%%%%%%%%%%%%%%%
dataFileFlag = 0;
eventFlag = 0;
augmKernelName = '';
tempFlag = 0;
if nargin == 1,
    iqm = varargin{1};
    % convert object to structure
    iqmstruct = struct(iqm);
    % if no filename provided then use the name of the IQMmodel as filename
    % remove white spaces from the name.
    functionName = strrep(iqmstruct.name,' ','');
    functionName = strrep(functionName,'-','');
    functionName = strrep(functionName,'+','');
    functionName = strrep(functionName,'*','');
    functionName = strrep(functionName,'/','');
    filename = strcat(functionName,'.m');
elseif nargin == 2,
    iqm = varargin{1};
    % convert object to structure
    iqmstruct = struct(iqm);    
    % extract filename from input argument number 2 - needed since 
    % IQMsimulate gives a whole path as filename
    [PATHSTR,functionName,EXT] = fileparts(varargin{2});
    if isempty(PATHSTR),
        filename = strcat(functionName,'.m');
    else
        filename = strcat(PATHSTR,'/',functionName,'.m');
    end
elseif nargin == 3,
    iqm = varargin{1};
    % convert object to structure
    iqmstruct = struct(iqm);    
    % extract filename from input argument number 2 - needed since 
    % IQMsimulate gives a whole path as filename
    [PATHSTR,functionName,EXT] = fileparts(varargin{2});
    if isempty(PATHSTR),
        filename = strcat(functionName,'.m');
    else
        filename = strcat(PATHSTR,'/',functionName,'.m');
    end
    dataFileFlag = varargin{3};
elseif nargin == 4,
    iqm = varargin{1};
    % convert object to structure
    iqmstruct = struct(iqm);    
    % extract filename from input argument number 2 - needed since 
    % IQMsimulate gives a whole path as filename
    [PATHSTR,functionName,EXT] = fileparts(varargin{2});
    if isempty(PATHSTR),
        filename = strcat(functionName,'.m');
    else
        filename = strcat(PATHSTR,'/',functionName,'.m');
    end
    dataFileFlag = varargin{3};
    eventFlag = varargin{4};
elseif nargin == 5,
    iqm = varargin{1};
    % convert object to structure
    iqmstruct = struct(iqm);    
    % extract filename from input argument number 2 - needed since 
    % IQMsimulate gives a whole path as filename
    [PATHSTR,functionName,EXT] = fileparts(varargin{2});
    if isempty(PATHSTR),
        filename = strcat(functionName,'.m');
    else
        filename = strcat(PATHSTR,'/',functionName,'.m');
    end
    dataFileFlag = varargin{3};
    eventFlag = varargin{4};   
    augmKernelName = varargin{5};
elseif nargin == 6,
    iqm = varargin{1};
    % convert object to structure
    iqmstruct = struct(iqm);    
    % extract filename from input argument number 2 - needed since 
    % IQMsimulate gives a whole path as filename
    [PATHSTR,functionName,EXT] = fileparts(varargin{2});
    if isempty(PATHSTR),
        filename = strcat(functionName,'.m');
    else
        filename = strcat(PATHSTR,'/',functionName,'.m');
    end
    dataFileFlag = varargin{3};
    eventFlag = varargin{4};   
    augmKernelName = varargin{5};
    tempFlag = varargin{6}; % set only by IQMcreateTempODEfile to avoid event error message ... its annoying
else
    error('Incorrect number of input arguments.');
end

%%%%%%%%%%%%%%%%%%%%%%%%%%%%%%%%%%%%%%%%%%%%%%%
% MAKE augmKernelName a global variable
%%%%%%%%%%%%%%%%%%%%%%%%%%%%%%%%%%%%%%%%%%%%%%%
if ~isempty(augmKernelName),
    eval(sprintf('global %s',augmKernelName));
end

%%%%%%%%%%%%%%%%%%%%%%%%%%%%%%%%%%%%%%%%%%%%%%%
% CHECK IF MODEL CONTAINS STATES
%%%%%%%%%%%%%%%%%%%%%%%%%%%%%%%%%%%%%%%%%%%%%%%
if isempty(iqmstruct.states),
    error('The model does not contain any states');
end

%%%%%%%%%%%%%%%%%%%%%%%%%%%%%%%%%%%%%%%%%%%%%%%
% HANDLE THE EVENT FLAG
%%%%%%%%%%%%%%%%%%%%%%%%%%%%%%%%%%%%%%%%%%%%%%%
% if the eventflag is set to a non-zero value AND at least on event is
% present in the model an event function file is created that has to be 
% used when calling the integrator. additionally a function is created 
% that determines the new values to which the states have to be set. 
if eventFlag == 1 && ~isempty(iqmstruct.events),
    filenameEvent = strrep(filename,functionName,strcat('event_',functionName));
    filenameEventAssignment = strrep(filename,functionName,strcat('event_assignment_',functionName));
    createEventFunctionIQM(iqm,filenameEvent,delaybase);
    createEventAssignmentFunctionIQM(iqm,filenameEventAssignment,delaybase);
end

%%%%%%%%%%%%%%%%%%%%%%%%%%%%%%%%%%%%%%%%%%%%%%%
% HANDLE THE DATAFILE FLAG
%%%%%%%%%%%%%%%%%%%%%%%%%%%%%%%%%%%%%%%%%%%%%%%
if dataFileFlag == 1,
    filenameDataFile = strrep(filename,functionName,strcat('datafile_',functionName));
    createSimulationDataFunctionIQM(iqm,filenameDataFile,delaybase);
end

%%%%%%%%%%%%%%%%%%%%%%%%%%%%%%%%%%%%%%%%%%%%%%%
% WRITE WARNING MESSAGE IN CASE EVENT FLAG IS NOT SET AND EVENTS ARE PRESENT
%%%%%%%%%%%%%%%%%%%%%%%%%%%%%%%%%%%%%%%%%%%%%%%
if eventFlag == 0 && ~isempty(iqmstruct.events) && tempFlag ~= 1,
    if length(iqmstruct.events) == 1,
        disp(sprintf('Warning: 1 event is present in the model.\nFor this analysis it is not taken into account.'));
    else
        disp(sprintf('Warning: %d events are present in the model.\nFor this analysis they are not taken into account.',length(iqmstruct.events)));
    end
end

%%%%%%%%%%%%%%%%%%%%%%%%%%%%%%%%%%%%%%%%%%%%%%%
% OPEN FILE FOR WRITING
%%%%%%%%%%%%%%%%%%%%%%%%%%%%%%%%%%%%%%%%%%%%%%%
fid = fopen(filename,'w');

%%%%%%%%%%%%%%%%%%%%%%%%%%%%%%%%%%%%%%%%%%%%%%%
% WRITE FUNCTION DEFINITION
%%%%%%%%%%%%%%%%%%%%%%%%%%%%%%%%%%%%%%%%%%%%%%%
fprintf(fid,'function [output] = %s(varargin)\n',functionName);

%%%%%%%%%%%%%%%%%%%%%%%%%%%%%%%%%%%%%%%%%%%%%%%
% WRITE THE HEADER
%%%%%%%%%%%%%%%%%%%%%%%%%%%%%%%%%%%%%%%%%%%%%%%
fprintf(fid,'%%%%%%%%%%%%%%%%%%%%%%%%%%%%%%%%%%%%%%%%%%%%%%%%%%%%%%%%%%%%%%%%%%%%%%%%%%%%%%%%%%%%%%%%%%%%%%%%%%%%%%%%%%%%%%%%%%%%%%%%%%%%%%%%%%%%%%%%%%%%%%%%%%%%\n');
fprintf(fid,'%% %s\n',iqmstruct.name);
fprintf(fid,'%% Generated: %s\n',datestr(now));
fprintf(fid,'%% \n');
fprintf(fid,'%% [output] = %s() => output = initial conditions in column vector\n',functionName);
fprintf(fid,'%% [output] = %s(''states'') => output = state names in cell-array\n',functionName);
fprintf(fid,'%% [output] = %s(''algebraic'') => output = algebraic variable names in cell-array\n',functionName);
fprintf(fid,'%% [output] = %s(''parameters'') => output = parameter names in cell-array\n',functionName);
fprintf(fid,'%% [output] = %s(''parametervalues'') => output = parameter values in column vector\n',functionName);
fprintf(fid,'%% [output] = %s(''variablenames'') => output = variable names in cell-array\n',functionName);
fprintf(fid,'%% [output] = %s(''variableformulas'') => output = variable formulas in cell-array\n',functionName);
fprintf(fid,'%% [output] = %s(time,statevector) => output = time derivatives in column vector\n',functionName);
fprintf(fid,'%% \n');
fprintf(fid,'%% State names and ordering:\n');
fprintf(fid,'%% \n');
for k = 1:length(iqmstruct.states),
    fprintf(fid,'%% statevector(%d): %s\n',k,iqmstruct.states(k).name);
end
fprintf(fid,'%% \n');
if ~isempty(iqmstruct.algebraic),
    didWriteStart = 0;
    offset = length(iqmstruct.states);
    offset2 = 1;
    for k = 1:length(iqmstruct.algebraic),
        if ~isempty(iqmstruct.algebraic(k).name) && didWriteStart == 0,    
            fprintf(fid,'%% Algebraic variable names and ordering (included in state and IC vector):\n');
            fprintf(fid,'%% \n'); 
            didWriteStart = 1;
        end
        if ~isempty(iqmstruct.algebraic(k).name),
            fprintf(fid,'%% statevector(%d): %s\n',offset+offset2,iqmstruct.algebraic(k).name);
            offset2 = offset2 + 1;
        end
    end
    if didWriteStart == 1,
        fprintf(fid,'%% \n');
    end
end
fprintf(fid,'%%%%%%%%%%%%%%%%%%%%%%%%%%%%%%%%%%%%%%%%%%%%%%%%%%%%%%%%%%%%%%%%%%%%%%%%%%%%%%%%%%%%%%%%%%%%%%%%%%%%%%%%%%%%%%%%%%%%%%%%%%%%%%%%%%%%%%%%%%%%%%%%%%%%\n');
fprintf(fid,'\n');
fprintf(fid,'global time\n');
if ~isempty(augmKernelName),
    fprintf(fid,'global %s\n',augmKernelName);
end 
fprintf(fid,'parameterValuesNew = [];\n\n');

%%%%%%%%%%%%%%%%%%%%%%%%%%%%%%%%%%%%%%%%%%%%%%%
% HANDLE VARARGINS
%%%%%%%%%%%%%%%%%%%%%%%%%%%%%%%%%%%%%%%%%%%%%%%
fprintf(fid,'%%%%%%%%%%%%%%%%%%%%%%%%%%%%%%%%%%%%%%%%%%%%%%%%%%%%%%%%%%%%%%%%%%%%%%%%%%%%%%%%%%%%%%%%%%%%%%\n');
fprintf(fid,'%% HANDLE VARIABLE INPUT ARGUMENTS\n');
fprintf(fid,'%%%%%%%%%%%%%%%%%%%%%%%%%%%%%%%%%%%%%%%%%%%%%%%%%%%%%%%%%%%%%%%%%%%%%%%%%%%%%%%%%%%%%%%%%%%%%%\n');
fprintf(fid,'if nargin == 0,\n');
    % Return initial conditions for state variables
    % If all initial conditions are numeric then a vector is defined.
    % Otherwise a cell-array with the corresponding entries. An additional
    % outside function then needs to be used to calculate the correct ICs
    % from the IC cellarray, IC values, and parameter values.
    fprintf(fid,'\t%% Return initial conditions of the state variables (and possibly algebraic variables)\n');
    if hasonlynumericICsIQM(iqm),
        icvector = [iqmstruct.states.initialCondition iqmstruct.algebraic.initialCondition];
        if ~isempty(icvector),
            fprintf(fid,'\toutput = [');
        else
            fprintf(fid,'\toutput = [];\n');
        end
        count = 0;
        for k = 1:length(icvector)-1,
            fprintf(fid,'%1.6g, ',icvector(k));
            count = count + 1;
            if count == 10, fprintf(fid,'...\n\t\t'); count = 0; end
        end
        if ~isempty(icvector),
            fprintf(fid,'%1.6g];\n',icvector(end));
        end
        fprintf(fid,'\toutput = output(:);\n');
    else
        % the model contains non-numeric initial conditions
        % get the ICs as cell-array
        icvector = {iqmstruct.states.initialCondition iqmstruct.algebraic.initialCondition};
        if ~isempty(icvector),
            fprintf(fid,'\toutput = {');
        else
            fprintf(fid,'\toutput = {};\n');
        end
        count = 0;
        for k = 1:length(icvector)-1,
            if isnumeric(icvector{k}),
                fprintf(fid,'%g, ',icvector{k});
            else
                fprintf(fid,'''%s'', ',strtrim(icvector{k}));
            end
            count = count + 1;
            if count == 10, fprintf(fid,'...\n\t\t'); count = 0; end
        end
        if ~isempty(icvector),
            if isnumeric(icvector{end}),
                fprintf(fid,'%g};\n',icvector{end});
            else
                fprintf(fid,'''%s''};\n',strtrim(icvector{end}));
            end
        end
    end
    fprintf(fid,'\treturn\n');
fprintf(fid,'elseif nargin == 1,\n');
	fprintf(fid,'\tif strcmp(varargin{1},''states''),\n');
        fprintf(fid,'\t\t%% Return state names in cell-array\n');
        if ~isempty(iqmstruct.states),
            fprintf(fid,'\t\toutput = {');
        else
            fprintf(fid,'\t\toutput = {};\n');
        end 
        count = 0;
        for k = 1:length(iqmstruct.states)-1,
            fprintf(fid,'''%s'', ',iqmstruct.states(k).name);
            count = count + 1;
            if count == 10, fprintf(fid,'...\n\t\t\t'); count = 0; end
        end
        if ~isempty(iqmstruct.states),
            fprintf(fid,'''%s''};\n',iqmstruct.states(end).name);
        end
	fprintf(fid,'\telseif strcmp(varargin{1},''algebraic''),\n');
        fprintf(fid,'\t\t%% Return algebraic variable names in cell-array\n');
        if ~isempty(iqmstruct.algebraic),
            fprintf(fid,'\t\toutput = {');
        else
            fprintf(fid,'\t\toutput = {};\n');
        end 
        count = 0;
        for k = 1:length(iqmstruct.algebraic)-1,
            if ~isempty(iqmstruct.algebraic(k).name),
                fprintf(fid,'''%s'', ',iqmstruct.algebraic(k).name);
                count = count + 1;
                if count == 10, fprintf(fid,'...\n\t\t\t'); count = 0; end
            end
        end
        if ~isempty(iqmstruct.algebraic),
            if ~isempty(iqmstruct.algebraic(end).name),
                fprintf(fid,'''%s''};\n',iqmstruct.algebraic(end).name);
            else
                fprintf(fid,'};\n');
            end
        end
    fprintf(fid,'\telseif strcmp(varargin{1},''parameters''),\n');
        fprintf(fid,'\t\t%% Return parameter names in cell-array\n');
        if ~isempty(iqmstruct.parameters),
            fprintf(fid,'\t\toutput = {');
        else
            fprintf(fid,'\t\toutput = {};\n');
        end 
        count = 0;
        for k = 1:length(iqmstruct.parameters)-1,
            fprintf(fid,'''%s'', ',iqmstruct.parameters(k).name);
            count = count + 1;
            if count == 10, fprintf(fid,'...\n\t\t\t'); count = 0; end
        end
        if ~isempty(iqmstruct.parameters),
            fprintf(fid,'''%s''};\n',iqmstruct.parameters(length(iqmstruct.parameters)).name);
        end
    fprintf(fid,'\telseif strcmp(varargin{1},''parametervalues''),\n');
        fprintf(fid,'\t\t%% Return parameter values in column vector\n');
        if ~isempty(iqmstruct.parameters),
            fprintf(fid,'\t\toutput = [');
        else
            fprintf(fid,'\t\toutput = [];\n');
        end 
        count = 0;
        for k = 1:length(iqmstruct.parameters)-1,
            fprintf(fid,'%1.6g, ',iqmstruct.parameters(k).value);
            count = count + 1;
            if count == 10, fprintf(fid,'...\n\t\t\t'); count = 0; end
        end
        if ~isempty(iqmstruct.parameters),
            fprintf(fid,'%1.6g];\n',iqmstruct.parameters(length(iqmstruct.parameters)).value);
        end
        
        
    fprintf(fid,'\telseif strcmp(varargin{1},''variablenames''),\n');
        fprintf(fid,'\t\t%% Return variable names in cell-array\n');
        if ~isempty(iqmstruct.variables),
            fprintf(fid,'\t\toutput = {');
        else
            fprintf(fid,'\t\toutput = {};\n');
        end 
        count = 0;
        for k = 1:length(iqmstruct.variables)-1,
            fprintf(fid,'''%s'', ',iqmstruct.variables(k).name);
            count = count + 1;
            if count == 10, fprintf(fid,'...\n\t\t\t'); count = 0; end
        end
        if ~isempty(iqmstruct.variables),
            fprintf(fid,'''%s''};\n',iqmstruct.variables(length(iqmstruct.variables)).name);
        end
    fprintf(fid,'\telseif strcmp(varargin{1},''variableformulas''),\n');
        fprintf(fid,'\t\t%% Return variable formulas in cell-array\n');
        if ~isempty(iqmstruct.variables),
            fprintf(fid,'\t\toutput = {');
        else
            fprintf(fid,'\t\toutput = {};\n');
        end 
        count = 0;
        for k = 1:length(iqmstruct.variables)-1,
            fprintf(fid,'''%s'', ',iqmstruct.variables(k).formula);
            count = count + 1;
            if count == 10, fprintf(fid,'...\n\t\t\t'); count = 0; end
        end
        if ~isempty(iqmstruct.variables),
            fprintf(fid,'''%s''};\n',iqmstruct.variables(length(iqmstruct.variables)).formula);
        end
        
        
    fprintf(fid,'\telse\n');
        fprintf(fid,'\t\terror(''Wrong input arguments! Please read the help text to the ODE file.'');\n');
    fprintf(fid,'\tend\n');
    fprintf(fid,'\toutput = output(:);\n');
    fprintf(fid,'\treturn\n');
fprintf(fid,'elseif nargin == 2,\n');
    fprintf(fid,'\ttime = varargin{1};\n');
    fprintf(fid,'\tstatevector = varargin{2};\n');
fprintf(fid,'elseif nargin == 3,\n');
    fprintf(fid,'\ttime = varargin{1};\n');
    fprintf(fid,'\tstatevector = varargin{2};\n');
    fprintf(fid,'\tparameterValuesNew = varargin{3};\n');
    fprintf(fid,'\tif length(parameterValuesNew) ~= %d,\n',length(iqmstruct.parameters));
    fprintf(fid,'\t\tparameterValuesNew = [];\n');
    fprintf(fid,'\tend\n');
fprintf(fid,'elseif nargin == 4,\n');
    fprintf(fid,'\ttime = varargin{1};\n');
    fprintf(fid,'\tstatevector = varargin{2};\n');
    fprintf(fid,'\tparameterValuesNew = varargin{4};\n');
fprintf(fid,'else\n');
    fprintf(fid,'\terror(''Wrong input arguments! Please read the help text to the ODE file.'');\n');
fprintf(fid,'end\n');
fprintf(fid,'\n');

%%%%%%%%%%%%%%%%%%%%%%%%%%%%%%%%%%%%%%%%%%%%%%%
% INITIALIZE THE STATES
%%%%%%%%%%%%%%%%%%%%%%%%%%%%%%%%%%%%%%%%%%%%%%%
fprintf(fid,'%%%%%%%%%%%%%%%%%%%%%%%%%%%%%%%%%%%%%%%%%%%%%%%%%%%%%%%%%%%%%%%%%%%%%%%%%%%%%%%%%%%%%%%%%%%%%%\n');
fprintf(fid,'%% STATES\n');
fprintf(fid,'%%%%%%%%%%%%%%%%%%%%%%%%%%%%%%%%%%%%%%%%%%%%%%%%%%%%%%%%%%%%%%%%%%%%%%%%%%%%%%%%%%%%%%%%%%%%%%\n');
for k = 1:length(iqmstruct.states)
    fprintf(fid,'%s = statevector(%d);\n',iqmstruct.states(k).name,k);
end
fprintf(fid,'\n');

%%%%%%%%%%%%%%%%%%%%%%%%%%%%%%%%%%%%%%%%%%%%%%%
% INITIALIZE THE ALGEBRAIC VARIABLES
%%%%%%%%%%%%%%%%%%%%%%%%%%%%%%%%%%%%%%%%%%%%%%%
if ~isempty(iqmstruct.algebraic),
    didWriteStart = 0;
    offset = length(iqmstruct.states);
    offset2 = 1;
    for k = 1:length(iqmstruct.algebraic)
        if ~isempty(iqmstruct.algebraic(k).name) && didWriteStart == 0,
            fprintf(fid,'%%%%%%%%%%%%%%%%%%%%%%%%%%%%%%%%%%%%%%%%%%%%%%%%%%%%%%%%%%%%%%%%%%%%%%%%%%%%%%%%%%%%%%%%%%%%%%\n');
            fprintf(fid,'%% ALGEBRAIC VARIABLES\n');
            fprintf(fid,'%%%%%%%%%%%%%%%%%%%%%%%%%%%%%%%%%%%%%%%%%%%%%%%%%%%%%%%%%%%%%%%%%%%%%%%%%%%%%%%%%%%%%%%%%%%%%%\n');
            didWriteStart = 1;
        end
        if ~isempty(iqmstruct.algebraic(k).name),
            fprintf(fid,'%s = statevector(%d);\n',iqmstruct.algebraic(k).name,offset+offset2);
            offset2 = offset2 + 1;
        end
    end
    if didWriteStart == 1,
        fprintf(fid,'\n');
    end
end

%%%%%%%%%%%%%%%%%%%%%%%%%%%%%%%%%%%%%%%%%%%%%%%
% INITIALIZE THE PARAMETES
%%%%%%%%%%%%%%%%%%%%%%%%%%%%%%%%%%%%%%%%%%%%%%%
if ~isempty(iqmstruct.parameters),
    fprintf(fid,'%%%%%%%%%%%%%%%%%%%%%%%%%%%%%%%%%%%%%%%%%%%%%%%%%%%%%%%%%%%%%%%%%%%%%%%%%%%%%%%%%%%%%%%%%%%%%%\n');
    fprintf(fid,'%% PARAMETERS\n');
    fprintf(fid,'%%%%%%%%%%%%%%%%%%%%%%%%%%%%%%%%%%%%%%%%%%%%%%%%%%%%%%%%%%%%%%%%%%%%%%%%%%%%%%%%%%%%%%%%%%%%%%\n');
    fprintf(fid,'if isempty(parameterValuesNew),\n');
    for k = 1:length(iqmstruct.parameters),
        fprintf(fid,'\t%s = %1.6g;\n',iqmstruct.parameters(k).name,iqmstruct.parameters(k).value);
    end
    fprintf(fid,'else\n');
    for k = 1:length(iqmstruct.parameters),
        fprintf(fid,'\t%s = parameterValuesNew(%d);\n',iqmstruct.parameters(k).name,k);
    end
    fprintf(fid,'end\n');
    fprintf(fid,'\n');
end

%%%%%%%%%%%%%%%%%%%%%%%%%%%%%%%%%%%%%%%%%%%%%%%
% INITIALIZE THE VARIABLES
%%%%%%%%%%%%%%%%%%%%%%%%%%%%%%%%%%%%%%%%%%%%%%%
if ~isempty(iqmstruct.variables),
    fprintf(fid,'%%%%%%%%%%%%%%%%%%%%%%%%%%%%%%%%%%%%%%%%%%%%%%%%%%%%%%%%%%%%%%%%%%%%%%%%%%%%%%%%%%%%%%%%%%%%%%\n');
    fprintf(fid,'%% VARIABLES\n');
    fprintf(fid,'%%%%%%%%%%%%%%%%%%%%%%%%%%%%%%%%%%%%%%%%%%%%%%%%%%%%%%%%%%%%%%%%%%%%%%%%%%%%%%%%%%%%%%%%%%%%%%\n');
    for k = 1:length(iqmstruct.variables),
        delayname = [delaybase '_var_' sprintf('%d',k)];
        fprintf(fid,'%s = %s;\n',iqmstruct.variables(k).name,processFormulaIQM(iqmstruct.variables(k).formula,delayname));
    end
    fprintf(fid,'\n');
end

%%%%%%%%%%%%%%%%%%%%%%%%%%%%%%%%%%%%%%%%%%%%%%%
% INITIALIZE THE REACTIONS
%%%%%%%%%%%%%%%%%%%%%%%%%%%%%%%%%%%%%%%%%%%%%%%
if ~isempty(iqmstruct.reactions),
    fprintf(fid,'%%%%%%%%%%%%%%%%%%%%%%%%%%%%%%%%%%%%%%%%%%%%%%%%%%%%%%%%%%%%%%%%%%%%%%%%%%%%%%%%%%%%%%%%%%%%%%\n');
    fprintf(fid,'%% REACTION KINETICS \n');
    fprintf(fid,'%%%%%%%%%%%%%%%%%%%%%%%%%%%%%%%%%%%%%%%%%%%%%%%%%%%%%%%%%%%%%%%%%%%%%%%%%%%%%%%%%%%%%%%%%%%%%%\n');
    for k = 1:length(iqmstruct.reactions),
        delayname = [delaybase '_reac_' sprintf('%d',k)];
        fprintf(fid,'%s = %s;\n',iqmstruct.reactions(k).name,processFormulaIQM(iqmstruct.reactions(k).formula,delayname));
    end
    fprintf(fid,'\n');
end

%%%%%%%%%%%%%%%%%%%%%%%%%%%%%%%%%%%%%%%%%%%%%%%
% DETERMINE ALSO EVENTASSIGNMENTS (NEEDED TO BE ABLE TO HANDEL DELAYS IN
% EVENT ASSIGNMENTS (for delayed events)
%%%%%%%%%%%%%%%%%%%%%%%%%%%%%%%%%%%%%%%%%%%%%%%
if ~isempty(iqmstruct.events),
    fprintf(fid,'%%%%%%%%%%%%%%%%%%%%%%%%%%%%%%%%%%%%%%%%%%%%%%%%%%%%%%%%%%%%%%%%%%%%%%%%%%%%%%%%%%%%%%%%%%%%%%\n');
    fprintf(fid,'%% EVENT ASSIGNMENTS AND TRIGGERS ... to handle delayed events\n');
    fprintf(fid,'%%%%%%%%%%%%%%%%%%%%%%%%%%%%%%%%%%%%%%%%%%%%%%%%%%%%%%%%%%%%%%%%%%%%%%%%%%%%%%%%%%%%%%%%%%%%%%\n');
    for k = 1:length(iqmstruct.events),
        delayname = [delaybase '_eventtrigger_' sprintf('%d',k)];        
        fprintf(fid,'eventtrigger_%d = %s;\n',k,processFormulaIQM(iqmstruct.events(k).trigger,delayname));
        for k2 = 1:length(iqmstruct.events(k).assignment),
            delayname = [delaybase '_eventassign_' sprintf('%d',k) '_' sprintf('%d',k2)];        
            fprintf(fid,'eventassign_%d_%d = %s;\n',k,k2,processFormulaIQM(iqmstruct.events(k).assignment(k2).formula,delayname));
        end
    end
    fprintf(fid,'\n');
end

%%%%%%%%%%%%%%%%%%%%%%%%%%%%%%%%%%%%%%%%%%%%%%%
% WRITE THE ODES
%%%%%%%%%%%%%%%%%%%%%%%%%%%%%%%%%%%%%%%%%%%%%%%
fprintf(fid,'%%%%%%%%%%%%%%%%%%%%%%%%%%%%%%%%%%%%%%%%%%%%%%%%%%%%%%%%%%%%%%%%%%%%%%%%%%%%%%%%%%%%%%%%%%%%%%\n');
fprintf(fid,'%% DIFFERENTIAL EQUATIONS\n');
fprintf(fid,'%%%%%%%%%%%%%%%%%%%%%%%%%%%%%%%%%%%%%%%%%%%%%%%%%%%%%%%%%%%%%%%%%%%%%%%%%%%%%%%%%%%%%%%%%%%%%%\n');
for k = 1:length(iqmstruct.states),   
    delayname = [delaybase '_ode_' sprintf('%d',k)];
    fprintf(fid,'%s_dot = %s;\n',iqmstruct.states(k).name,processFormulaIQM(iqmstruct.states(k).ODE,delayname));
end
fprintf(fid,'\n');

%%%%%%%%%%%%%%%%%%%%%%%%%%%%%%%%%%%%%%%%%%%%%%%
% WRITE THE ALGEBRAIC RULES
%%%%%%%%%%%%%%%%%%%%%%%%%%%%%%%%%%%%%%%%%%%%%%%
if ~isempty(iqmstruct.algebraic),
    fprintf(fid,'%%%%%%%%%%%%%%%%%%%%%%%%%%%%%%%%%%%%%%%%%%%%%%%%%%%%%%%%%%%%%%%%%%%%%%%%%%%%%%%%%%%%%%%%%%%%%%\n');
    fprintf(fid,'%% ALGEBRAIC RULES\n');
    fprintf(fid,'%%%%%%%%%%%%%%%%%%%%%%%%%%%%%%%%%%%%%%%%%%%%%%%%%%%%%%%%%%%%%%%%%%%%%%%%%%%%%%%%%%%%%%%%%%%%%%\n');
    for k = 1:length(iqmstruct.algebraic)
        fprintf(fid,'AlgebraicRule_%d = %s;\n',k,iqmstruct.algebraic(k).formula);
    end
    fprintf(fid,'\n');
end

%%%%%%%%%%%%%%%%%%%%%%%%%%%%%%%%%%%%%%%%%%%%%%%
% WRITE RETURN VALUES
%%%%%%%%%%%%%%%%%%%%%%%%%%%%%%%%%%%%%%%%%%%%%%%
fprintf(fid,'%%%%%%%%%%%%%%%%%%%%%%%%%%%%%%%%%%%%%%%%%%%%%%%%%%%%%%%%%%%%%%%%%%%%%%%%%%%%%%%%%%%%%%%%%%%%%%\n');
fprintf(fid,'%% RETURN VALUES\n');
fprintf(fid,'%%%%%%%%%%%%%%%%%%%%%%%%%%%%%%%%%%%%%%%%%%%%%%%%%%%%%%%%%%%%%%%%%%%%%%%%%%%%%%%%%%%%%%%%%%%%%%\n');
if isempty(augmKernelName),
    fprintf(fid,'%% STATE ODEs\n');
    for k = 1:length(iqmstruct.states),
        fprintf(fid,'output(%d) = %s_dot;\n',k,iqmstruct.states(k).name);
    end
    if ~isempty(iqmstruct.algebraic),
        fprintf(fid,'%% ARs\n');
        offset = length(iqmstruct.states);
        for k = 1:length(iqmstruct.algebraic),
            fprintf(fid,'output(%d) = AlgebraicRule_%d;\n',k+offset,k);
        end
    end
else
    fprintf(fid,'%% STATE ODEs ... HANDLING FAST REACTIONS\n');
    for k = 1:length(iqmstruct.states),
        fprintf(fid,'states(%d) = %s_dot;\n',k,iqmstruct.states(k).name);
    end
    fprintf(fid,'%% MULTIPLY WITH AUGMENTED KERNEL\n');
    fprintf(fid,'output = %s*states(:);\n',augmKernelName);
    if ~isempty(iqmstruct.algebraic),
        fprintf(fid,'%% ARs\n');
        offset = eval(sprintf('size(%s,1);',augmKernelName));
        for k = 1:length(iqmstruct.algebraic),
            fprintf(fid,'output(%d) = AlgebraicRule_%d;\n',k+offset,k);
        end
    end    
end
fprintf(fid,'%% return a column vector \n');
% Return a column vector
fprintf(fid,'output = output(:);\n');
fprintf(fid,'return\n');
fprintf(fid,'\n');
fprintf(fid,'\n');

%%%%%%%%%%%%%%%%%%%%%%%%%%%%%%%%%%%%%%%%%%%%%%%
% WRITE THE FUNCTIONS
%%%%%%%%%%%%%%%%%%%%%%%%%%%%%%%%%%%%%%%%%%%%%%%
if ~isempty(iqmstruct.functions),
    fprintf(fid,'%%%%%%%%%%%%%%%%%%%%%%%%%%%%%%%%%%%%%%%%%%%%%%%%%%%%%%%%%%%%%%%%%%%%%%%%%%%%%%%%%%%%%%%%%%%%%%\n');
    fprintf(fid,'%% FUNCTIONS\n');
    fprintf(fid,'%%%%%%%%%%%%%%%%%%%%%%%%%%%%%%%%%%%%%%%%%%%%%%%%%%%%%%%%%%%%%%%%%%%%%%%%%%%%%%%%%%%%%%%%%%%%%%\n');
    for k = 1:length(iqmstruct.functions),
        fprintf(fid,'function [result] = %s(%s)\n',iqmstruct.functions(k).name,iqmstruct.functions(k).arguments);
        fprintf(fid,'global time\n');
        delayname = [delaybase '_func_' sprintf('%d',k)];        
        fprintf(fid,'result = %s;\n',processFormulaIQM(iqmstruct.functions(k).formula,delayname));
        fprintf(fid,'return\n');
        fprintf(fid,'\n');
    end
    fprintf(fid,'\n');
end

%%%%%%%%%%%%%%%%%%%%%%%%%%%%%%%%%%%%%%%%%%%%%%%
% WRITE THE MATLAB FUNCTIONS
%%%%%%%%%%%%%%%%%%%%%%%%%%%%%%%%%%%%%%%%%%%%%%%
if ~isempty(iqmstruct.functionsMATLAB),
    fprintf(fid,'%%%%%%%%%%%%%%%%%%%%%%%%%%%%%%%%%%%%%%%%%%%%%%%%%%%%%%%%%%%%%%%%%%%%%%%%%%%%%%%%%%%%%%%%%%%%%%\n');
    fprintf(fid,'%% MATLAB FUNCTIONS\n');
    fprintf(fid,'%%%%%%%%%%%%%%%%%%%%%%%%%%%%%%%%%%%%%%%%%%%%%%%%%%%%%%%%%%%%%%%%%%%%%%%%%%%%%%%%%%%%%%%%%%%%%%\n');
    delayname = [delaybase '_funcmatlab'];
    fprintf(fid,'%s',processFormulaIQM(iqmstruct.functionsMATLAB,delayname));
    fprintf(fid,'\n');
end

%%%%%%%%%%%%%%%%%%%%%%%%%%%%%%%%%%%%%%%%%%%%%%%
% CLOSE THE FILE
%%%%%%%%%%%%%%%%%%%%%%%%%%%%%%%%%%%%%%%%%%%%%%%
status = fclose(fid);
rehash
% Return
return
```
