## Supplementary material for "A systems biology analysis of adrenergically stimulated adiponectin exocytosis in white adipocytes": All scripts used in this work: IQMcreateTempODEfile.m

```
function [ODEfctname, ODEfilefullpath, DATAfctname, EVENTfctname, EVENTASSGNfctname] = IQMcreateTempODEfile(varargin)
% IQMcreateTempODEfile: creates a temporary ODE file, same as
% IQMcreateODEfile, but the file is written in the systems temporary
% directory. Creates also datafile and eventhandling files if wanted.
% No filename is passed, but a temporary filename is chosen automatically
% and returned back as output argument.
%
% USAGE:
% ======
% [ODEfctname, ODEfilefullpath] = IQMcreateTempODEfile(iqm)
% [ODEfctname, ODEfilefullpath, DATAfctname] = IQMcreateTempODEfile(iqm,dataFileFlag)
% [ODEfctname, ODEfilefullpath, DATAfctname, EVENTfctname, EVENTASSGNfctname] = IQMcreateTempODEfile(iqm,dataFileFlag,eventFlag)
% [ODEfctname, ODEfilefullpath, DATAfctname, EVENTfctname, EVENTASSGNfctname] = IQMcreateTempODEfile(iqm,dataFileFlag,eventFlag,augmKernelName)
%
% iqm: IQMmodel to convert to an ODE file description
% dataFileFlag: =1: creates an additional file allowing to determine
%                   variable and reaction rate values for given state values 
%                   and time vector.
%               =0: doe not create the additional file
% eventFlag: =1: creates 2 additional files for the handling of events in the model. 
%            =0: does not create these two files.
%            THE EVENT FILES ARE ONLY CREATED IN CASE THAT EVENTS ARE
%            PRESENT IN THE MODEL            
%
% DEFAULT VALUES:
% ===============
% dataFileFlag: 0
% eventFlag: 0
%
% OUTPUT ARGUMENTS:
% =================
% ODEfctname: name of the function (filename without .m extension)
% DATAfctname: name of the datafile function (filename without .m extension)
% EVENTfctname: name of the event function (filename without .m extension)
% EVENTASSGNfctname: name of the event assignment function (filename without .m extension)
% ODEfilefullpath: complete path to the created ODE file (including .m extension)

% <<<COPYRIGHTSTATEMENT - IQM TOOLS LITE>>>


%%%%%%%%%%%%%%%%%%%%%%%%%%%%%%%%%%%%%%%%%%%%%%%
% TEMPDIR 
%%%%%%%%%%%%%%%%%%%%%%%%%%%%%%%%%%%%%%%%%%%%%%%
addpath(tempdirIQM);   % add it to the path

%%%%%%%%%%%%%%%%%%%%%%%%%%%%%%%%%%%%%%%%%%%%%%%
% PROCESS VARIABLE INPUT ARGUMENTS
%%%%%%%%%%%%%%%%%%%%%%%%%%%%%%%%%%%%%%%%%%%%%%%
eventFlag = 0;
dataFileFlag = 0;
augmKernelName = '';
if nargin == 1,
    iqm = varargin{1};
elseif nargin == 2,
    iqm = varargin{1};
    dataFileFlag = varargin{2};
elseif nargin == 3,
    iqm = varargin{1};
    dataFileFlag = varargin{2};
    eventFlag = varargin{3};
elseif nargin == 4,
    iqm = varargin{1};
    dataFileFlag = varargin{2};
    eventFlag = varargin{3};
    augmKernelName = varargin{4};
else
    error('Incorrect number of input arguments.');
end

%%%%%%%%%%%%%%%%%%%%%%%%%%%%%%%%%%%%%%%%%%%%%%%
% GET TEMP FILE NAME
%%%%%%%%%%%%%%%%%%%%%%%%%%%%%%%%%%%%%%%%%%%%%%%
% Get temporary filenme
tempfilefullpath = tempnameIQM;
[pathstr,tempfilename,ext] = fileparts(tempfilefullpath);     

%%%%%%%%%%%%%%%%%%%%%%%%%%%%%%%%%%%%%%%%%%%%%%%
% CREATE THE FILES
%%%%%%%%%%%%%%%%%%%%%%%%%%%%%%%%%%%%%%%%%%%%%%%
tempFlag = 1; % to avoid the event warning when creating files for models with events w/o setting the event flag
IQMcreateODEfile(iqm,tempfilefullpath,dataFileFlag,eventFlag,augmKernelName,tempFlag);

%%%%%%%%%%%%%%%%%%%%%%%%%%%%%%%%%%%%%%%%%%%%%%%
% CREATE THE FILE NAMES
%%%%%%%%%%%%%%%%%%%%%%%%%%%%%%%%%%%%%%%%%%%%%%%
ODEfctname = tempfilename;
if dataFileFlag == 1,
    DATAfctname = strcat('datafile_',ODEfctname);
else 
    DATAfctname = '';
end
if eventFlag == 1,
    EVENTfctname = strcat('event_',ODEfctname);
    EVENTASSGNfctname = strcat('event_assignment_',ODEfctname);
else
    EVENTfctname = '';
    EVENTASSGNfctname = '';
end
ODEfilefullpath = strcat(tempfilefullpath,'.m');
```
