## Supplementary material for "A systems biology analysis of adrenergically stimulated adiponectin exocytosis in white adipocytes": All scripts used in this work: IQMcreateTEXTBCfile.m

```
function IQMcreateTEXTBCfile(varargin)
% IQMcreateTEXTBCfile: creates a *.txtbc file with the models text
% description in a biochemically oriented format
%
% USAGE:
% ======
% [] = IQMcreateTEXTBCfile(iqm)         
% [] = IQMcreateTEXTBCfile(iqm,filename)
%
% iqm: IQMmodel to convert to a textfile description
% filename: filename for the created textfile (.txtbc)
%
% DEFAULT VALUES:
% ===============
% filename: constructed from the models name

% <<<COPYRIGHTSTATEMENT - IQM TOOLS LITE>>>

%%%%%%%%%%%%%%%%%%%%%%%%%%%%%%%%%%%%%%%%%%%%%%%
% PROCESS VARIABLE INPUT ARGUMENTS
%%%%%%%%%%%%%%%%%%%%%%%%%%%%%%%%%%%%%%%%%%%%%%%
if nargin == 1,
    iqm = varargin{1};
    % convert object to structure
    ms = struct(iqm);    
    % if no filename provided then use the name of the IQMmodel as filename
    % remove unwanted characters first
    functionName = regexprep(ms.name,'\W','');
    filename = strcat(functionName,'.txtbc');
elseif nargin == 2,
    iqm = varargin{1};
    filename = strcat(varargin{2},'.txtbc');
else
    error('Incorrect number of input arguments.');
end

%%%%%%%%%%%%%%%%%%%%%%%%%%%%%%%%%%%%%%%%%%%%%%%
% CONVERT MODEL TO TEXT
%%%%%%%%%%%%%%%%%%%%%%%%%%%%%%%%%%%%%%%%%%%%%%%
[modelTextBCStructure] = convertModelToTextBCIQM(iqm);
[completeTextBC] = setPartsToCompleteTextBCIQM(modelTextBCStructure);

%%%%%%%%%%%%%%%%%%%%%%%%%%%%%%%%%%%%%%%%%%%%%%%
% WRITE THE FILE
%%%%%%%%%%%%%%%%%%%%%%%%%%%%%%%%%%%%%%%%%%%%%%%
fid = fopen(filename,'w');
fwrite(fid,completeTextBC);
fclose(fid);
return
```
