## Supplementary material for "A systems biology analysis of adrenergically stimulated adiponectin exocytosis in white adipocytes": All scripts used in this work: IQMcreateTEXTBCmodel.m

```
function [] = IQMcreateTEXTBCmodel(name,varargin)
% IQMcreateTEXTBCmodel: Creates a new TEXTBC model, saves the corresponding
% file in the current directory and opens it in the editor, if opening is
% desired.
%
% USAGE:
% ======
% IQMcreateTEXTBCmodel(name)
% IQMcreateTEXTBCmodel(name,openFlag)
%
% name: filename of the model and model name
% openFlag: decides if the created model is automatically opened in the
%           editor or not. (=0: do not open, =1: do open)
%
% DEFAULT VALUES:
% ===============
% openFlag: 1 (do open)

% <<<COPYRIGHTSTATEMENT - IQM TOOLS LITE>>>

%%%%%%%%%%%%%%%%%%%%%%%%%%%%%%%%%%%%%%%%%%%%%%%
% HANDLE VARIABE INPUT ARGUMENTS
%%%%%%%%%%%%%%%%%%%%%%%%%%%%%%%%%%%%%%%%%%%%%%%
openFlag = 1;
if nargin == 2,
    openFlag = varargin{1};
end

%%%%%%%%%%%%%%%%%%%%%%%%%%%%%%%%%%%%%%%%%%%%%%%
% CREATE NEW MODEL
%%%%%%%%%%%%%%%%%%%%%%%%%%%%%%%%%%%%%%%%%%%%%%%
model = IQMmodel();
ms = struct(model);
ms.name = name;
model = IQMmodel(ms);
IQMcreateTEXTBCfile(model,name);

%%%%%%%%%%%%%%%%%%%%%%%%%%%%%%%%%%%%%%%%%%%%%%%
% OPEN IF DESIRED
%%%%%%%%%%%%%%%%%%%%%%%%%%%%%%%%%%%%%%%%%%%%%%%
if openFlag,
    edit(strcat(name,'.txtbc'));
end

return
```
