## Supplementary material for "A systems biology analysis of adrenergically stimulated adiponectin exocytosis in white adipocytes": All scripts used in this work: IQMcreateTEXTfile.m

```
function IQMcreateTEXTfile(varargin)
% IQMcreateTEXTfile: creates a *.txt file with the models text description
% based on ordinary differential equations
%
% USAGE:
% ======
% [] = IQMcreateTEXTfile(iqm)         
% [] = IQMcreateTEXTfile(iqm,filename)
%
% iqm: IQMmodel to convert to a textfile description
% filename: filename for the created textfile 
%
% DEFAULT VALUES:
% ===============
% filename: constructed from the models name

% <<<COPYRIGHTSTATEMENT - IQM TOOLS LITE>>>


%%%%%%%%%%%%%%%%%%%%%%%%%%%%%%%%%%%%%%%%%%%%%%%
% PROCESS VARIABLE INPUT ARGUMENTS
%%%%%%%%%%%%%%%%%%%%%%%%%%%%%%%%%%%%%%%%%%%%%%%
if nargin == 1,
    iqm = varargin{1};
    % convert object to structure
    ms = struct(iqm);    
    % if no filename provided then use the name of the IQMmodel as filename
    % remove unwanted characters first
    functionName = regexprep(ms.name,'\W','');
    filename = strcat(functionName,'.txt');
elseif nargin == 2,
    iqm = varargin{1};
    filename = strcat(varargin{2},'.txt');
else
    error('Incorrect number of input arguments.');
end

%%%%%%%%%%%%%%%%%%%%%%%%%%%%%%%%%%%%%%%%%%%%%%%
% CONVERT MODEL TO TEXT
%%%%%%%%%%%%%%%%%%%%%%%%%%%%%%%%%%%%%%%%%%%%%%%
[modelTextStructure] = convertModelToTextIQM(iqm);
[completeText] = setPartsToCompleteTextIQM(modelTextStructure);

%%%%%%%%%%%%%%%%%%%%%%%%%%%%%%%%%%%%%%%%%%%%%%%
% WRITE THE FILE
%%%%%%%%%%%%%%%%%%%%%%%%%%%%%%%%%%%%%%%%%%%%%%%
fid = fopen(filename,'w');
fwrite(fid,completeText);
fclose(fid);
return
```
