## Supplementary material for "A systems biology analysis of adrenergically stimulated adiponectin exocytosis in white adipocytes": All scripts used in this work: IQMdeleteTempODEfile.m

```
function [] = IQMdeleteTempODEfile(fullpathfilename)
% This function takes as input argument the full path to an ODE file that
% is to be deleted. It also tries to delete the datafile_* and the event
% files that belong to this ODE file. Warnings are switched of not to enoy
% the user with unecessary warnings when a file is not present.

% <<<COPYRIGHTSTATEMENT - IQM TOOLS LITE>>>


% initialize simulation results

[pathstr,filename,ext] = fileparts(fullpathfilename);
ODEfilename = strcat(pathstr,'/',filename,'.m');
DATAfilename = strcat(pathstr,'/','datafile_',filename,'.m');
EVENTfilename = strcat(pathstr,'/','event_',filename,'.m');
EVENTASSIGNfilename = strcat(pathstr,'/','event_assignment_',filename,'.m');
warning off;  % toggle warnings off as not all files might exist
delete(ODEfilename);
delete(DATAfilename);
delete(EVENTfilename);
delete(EVENTASSIGNfilename);
warning on;  % toggle warnings on again
```
