## Supplementary material for "A systems biology analysis of adrenergically stimulated adiponectin exocytosis in white adipocytes": All scripts used in this work: IQMevents.m

```
function [names,triggers,variables,formulas] = IQMevents(model)
% IQMevents: Returns information about the events in an IQMmodel.
%
% USAGE:
% ======
% [names,triggers,variables,formulas] = IQMevents(model)
%
% model: IQMmodel (function can not be used on M-file model)
%
% Output Arguments:
% =================
% names: cell-array with models event names
% triggers: cell-array with triggers for the events
% variables: cell-array with the variables affected by the events
% formulas: cell-array with the formulas how to affect the variables if an
%   event is fired
%
% The ordering of the elements in the cell-arrays is important. The i-th
% elements in the output variables belongs to the i-th event.

% <<<COPYRIGHTSTATEMENT - IQM TOOLS LITE>>>

%%%%%%%%%%%%%%%%%%%%%%%%%%%%%%%%%%%%%%%%%%%%%%%
% PROCESS IQMMODEL OR ODE FILE
%%%%%%%%%%%%%%%%%%%%%%%%%%%%%%%%%%%%%%%%%%%%%%%
if strcmp('IQMmodel',class(model)),
    iqm = IQMstruct(model);
    names = {};
    triggers = {};
    variables = {};
    formulas = {};
    for k = 1:length(iqm.events),
        names{k} = iqm.events(k).name;
        triggers{k} = iqm.events(k).trigger;
        variables{k} = {iqm.events(k).assignment.variable};
        formulas{k} = {iqm.events(k).assignment.formula};
    end
else
    error('The function can only be used on IQMmodels, not on M-file ODE models');
end
names = names(:);
triggers = triggers(:);
return
```
