## Supplementary material for "A systems biology analysis of adrenergically stimulated adiponectin exocytosis in white adipocytes": All scripts used in this work: IQMexportSBML.m

```
function [] = IQMexportSBML(iqm, filename)
% IQMexportSBML 
% exports an IQMmodel to an SBML Level 2 Version 1 model using the OutputSBML function
% from the SBML toolbox. Non-numeric ICs are not handled but converted to
% numeric ICs. 
%
% IMPORTANT:
% ==========
% To export a model, the user is required to provide correct information in 
% the "type,compartment,unittype" fields of the internal IQMmodel data structure.
%
% This information can be added by directly editing the models structure,
% or by providing the information in the model text representation.
%
% 1) Editing the structure
% ------------------------
% The model structure is obtained by typing "modelstruct = IQMstruct(iqmmodel)"
% The 'states', 'algebraic', 'parameters', and 'variables' fields of the structure have the 
% following subfields that need to be set to correct values.
%   subfields       needed content          comment
%   'type':      	'isSpecie'              in case the state/parameter/variable in
%                                           the IQMmodel is an SBML specie
%                   'isParameter'           in case the state/parameter/variable in
%                                           the IQMmodel is an SBML
%                                           parameter
%                   'isCompartment'         in case the state/parameter/variable in
%                                           the IQMmodel is an SBML compartment
%
%   'compartment':  specie compartment      if type='isSpecie' it defines the name of the
%                                           compartment the specie is located in.
%                   outside compartment     if type='isCompartment' it defined the name 
%                                           of the compartment outside
%                   ''                      if type='isParameter' or type='isCompartment' 
%                                           and there is no outside compartment or 
%                                           type='isSpecie' and there is no
%                                           compartment defined.
%
%   'unittype':     'amount'                if type='isSpecie' and the
%   species units 
%                                           are in 'amount'
%                   'concentration'         if type='isSpecie' and the species units 
%                                           are in 'concentration'
%                   ''                      if type is not 'isSpecie'                   
%
% 2) Providing the information in the model text description, by IQMedit,
%    IQMeditBC or directly by editing text files
% ----------------------------------------------------------------------
% For each state, parameter, and variable the additional information has
% the following syntax:
%
% {type:compartment:unittype}
%
% where 'type', 'compartment', and 'unittype' is defined as above.
% This argument needs to be placed in each definition of a state,
% algebraic, parameter, and variable just before the optional comment.
%
% Example:
% --------
% An example model that has been correctly prepared for SBML export 
% is defined in the file IQMlite/examples/exampleSBMLexport.txt
%
% USAGE:
% ======
% [] = IQMexportSBML(model)
% [] = IQMexportSBML(model,filename)
%
% model:    IQMmodel to export to SBML
% filename: Name of the xml file to export to (optional: dialog window opens)

% <<<COPYRIGHTSTATEMENT - IQM TOOLS LITE>>>


%%%%%%%%%%%%%%%%%%%%%%%%%%%%%%%%%%%%%%%%%%%%%%%
% HANDLE NON NUMERIC ICs
%%%%%%%%%%%%%%%%%%%%%%%%%%%%%%%%%%%%%%%%%%%%%%%
if ~hasonlynumericICsIQM(iqm),
    iqm = IQMconvertNonNum2NumIC(iqm);
    disp('Model contains non-numeric initial conditions. For the SBML export');
    disp('these are converted to numeric ones using the IQMconvertNonNum2NumIC function.');
end


% skipSBMLpreset = 0;
% if nargin == 2,
%     skipSBMLpreset = varargin{1};
% end

%%%%%%%%%%%%%%%%%%%%%%%%%%%%%%%%%%%%%%%%%%%%%%%
% PREPROCESS and CHECK IQMmodel (fill type, compartment, unittype fields if possible)
%%%%%%%%%%%%%%%%%%%%%%%%%%%%%%%%%%%%%%%%%%%%%%%
% if ~skipSBMLpreset,
    iqm = IQMmakeSBMLpresets(iqm);
% end

% test wether all name fields are filled and are unique
namesOK = checkNamesInIQMmodel(iqm, 1);
if ~namesOK,
    errorMessage{1} = 'SBML export aborted because of errors concerning the names used in the given IQMmodel. For further information use "checkNamesInIQMmodel" function!';
    messageOutput(errorMessage, 1);
    return    
end
% if test shows that model isn't SBML compliant abort export
[sbmlCompliant, onlyCompartmentError] = testSBMLsettings(iqm, 1);
if ~sbmlCompliant,
    if onlyCompartmentError,
        iqm = addDefaultCompartment(iqm);
    else
        errorMessage{1} = 'SBML export aborted because of errors concerning the SBML compliance. For further information use "testSBMLsettings" function!';
        messageOutput(errorMessage, 1);
        return
    end
end

% convert object to structure
ms = struct(iqm);  

%% Handle the division by compartments if amount and revert that
tempfile = tempname;
IQMcreateTEXTfile(iqm,tempfile);
contents = fileread([tempfile '.txt']);
% Get the species and compartment names for amount species
speciesNames = {};
compartmentNames = {};
for k=1:length(ms.states),
    if strcmp(ms.states(k).type,'isSpecie') && strcmp(ms.states(k).unittype,'amount'),
        speciesNames{end+1} = ms.states(k).name;
        compartmentNames{end+1} = ms.states(k).compartment;
    end
end
for k=1:length(ms.parameters),
    if strcmp(ms.parameters(k).type,'isSpecie') && strcmp(ms.parameters(k).unittype,'amount'),
        speciesNames{end+1} = ms.parameters(k).name;
        compartmentNames{end+1} = ms.parameters(k).compartment;
    end
end
for k=1:length(ms.variables),
    if strcmp(ms.variables(k).type,'isSpecie') && strcmp(ms.variables(k).unittype,'amount'),
        speciesNames{end+1} = ms.variables(k).name;
        compartmentNames{end+1} = ms.variables(k).compartment;
    end
end
% Cycle through the species and remove the division by compartment if amount
for k=1:length(speciesNames),
    old = ['(' speciesNames{k} '/' compartmentNames{k} ')'];
    new = speciesNames{k};
    contents = strrep(contents,old,new);
end
% Save file again
fid = fopen([tempfile '.txt'],'w');
fprintf(fid,'%s',contents);
fclose(fid);
% Load changed model
iqm = IQMmodel([tempfile '.txt']);
% Get structure again
ms = struct(iqm);
% Delete tempfile
delete([tempfile '.txt']);

%%


%%%%%%%%%%%%%%%%%%%%%%%%%%%%%%%%%%%%%%%%%%%%%%%
% CHECK IF MODEL CONTAINS MATLAB FUNCTIONS
%%%%%%%%%%%%%%%%%%%%%%%%%%%%%%%%%%%%%%%%%%%%%%%
if ~isempty(ms.functionsMATLAB),
    error('Model contains MATLAB functions. Such functions are not supported in SBML.');
end

%%%%%%%%%%%%%%%%%%%%%%%%%%%%%%%%%%%%%%%%%%%%%%%
% DETERMINE STOICHIOMETRIX MATRIX and related stuff (to determine if
% boundary species or species)
%%%%%%%%%%%%%%%%%%%%%%%%%%%%%%%%%%%%%%%%%%%%%%%
[StoichiometricMatrix,SBMLspeciesNames,SBMLreactionNames,SBMLreversibleFlag] = IQMstoichiometry(iqm,1,1);

%%%%%%%%%%%%%%%%%%%%%%%%%%%%%%%%%%%%%%%%%%%%%%%
% CREATE AN EMPTY SBML MATLAB STRUCTURE
%%%%%%%%%%%%%%%%%%%%%%%%%%%%%%%%%%%%%%%%%%%%%%%
% Create empty SBMLstructure
% functionDefinition substructure
functionDefinitionStruct = struct('typecode', {}, 'metaid', '', 'notes', {}, 'annotation', {}, 'name', {}, 'id', {}, 'math', {});
% unitDefinition substructure
unitDefinitionStruct = struct('typecode',{}, 'metaid', '','notes',{},'annotation',{}, 'name', {}, 'id', {}, 'unit', {});
% compartment substructure
compartmentStruct = struct('typecode', {}, 'metaid', '', 'notes', {}, 'annotation', {}, 'name', {}, 'id', {}, 'spatialDimensions', {}, 'size', {}, 'units', {}, 'outside', {}, 'constant', {}, 'isSetSize', {}, 'isSetVolume', {});
% species substructure
speciesStruct = struct('typecode', {}, 'metaid', '', 'notes', {}, 'annotation', {}, 'name', {},'id', {}, 'compartment', {}, 'initialAmount', {}, 'initialConcentration', {}, 'substanceUnits', {}, 'spatialSizeUnits', {}, 'hasOnlySubstanceUnits', {}, 'boundaryCondition', {}, 'charge', {}, 'constant', {}, 'isSetInitialAmount', {}, 'isSetInitialConcentration', {}, 'isSetCharge', {});
% parameter substructure
parameterStruct = struct('typecode', {}, 'metaid', '', 'notes', {}, 'annotation', {}, 'name', {}, 'id', {}, 'value', {}, 'units', {}, 'constant', {}, 'isSetValue', {});
% rule substructure
ruleStruct = struct('typecode', {}, 'metaid', '', 'notes', {}, 'annotation', {}, 'formula', {}, 'variable', {}, 'species', {}, 'compartment', {}, 'name', {}, 'units', {});
% reaction substructure
reactantStruct = struct('typecode', {}, 'metaid', '', 'notes', {}, 'annotation', {}, 'species', {}, 'stoichiometry', {}, 'denominator', {}, 'stoichiometryMath', {});
productStruct = struct('typecode', {}, 'metaid', '', 'notes', {}, 'annotation', {}, 'species', {}, 'stoichiometry', {}, 'denominator', {}, 'stoichiometryMath', {});
modifierStruct = struct('typecode', {}, 'metaid', '', 'notes', {}, 'annotation', {}, 'species', {});
kineticLawParameterStruct = struct('typecode', {}, 'metaid', '', 'notes', {}, 'annotation', {}, 'name', {}, 'id', {}, 'value', {}, 'units', {}, 'constant', {}, 'isSetValue', {});
kineticLawStruct = struct('typecode', {}, 'metaid', '', 'notes', {}, 'annotation', {}, 'formula', {}, 'math', {}, 'parameter', kineticLawParameterStruct, 'timeUnits', {}, 'substanceUnits', {});
reactionStruct = struct('typecode', {}, 'metaid', '', 'notes', {}, 'annotation', {}, 'name', {}, 'id', {}, 'reactant', reactantStruct, 'product', productStruct, 'modifier', modifierStruct, 'kineticLaw', kineticLawStruct, 'reversible', {}, 'fast', {}, 'isSetFast', {});
eventStruct = struct('typecode', {}, 'metaid', '', 'notes', {}, 'annotation', {}, 'name', {}, 'id', {}, 'trigger', {}, 'delay', {}, 'timeUnits', {}, 'eventAssignment', {});
namespaceStruct = struct('prefix', '', 'uri', 'http://www.sbml.org/sbml/level2');
% Create IQMstructure
SBMLstructure = struct('typecode', 'SBML_MODEL',...
                       'metaid', '', ...
                       'notes', '', ...
                       'annotation', '', ...
                       'SBML_level', 2, ...
                       'SBML_version', 1, ...
                       'name', '', ...
                       'id', '', ...
                       'functionDefinition', functionDefinitionStruct, ...
                       'unitDefinition', unitDefinitionStruct, ...
                       'compartment', compartmentStruct, ...
                       'species', speciesStruct, ...
                       'parameter', parameterStruct, ...
                       'rule', ruleStruct, ...
                       'reaction', reactionStruct, ...
                       'event', eventStruct, ...
                       'time_symbol', 'time', ...
                       'delay_symbol', '', ...
                       'namespaces', namespaceStruct);

%%%%%%%%%%%%%%%%%%%%%%%%%%%%%%%%%%%%%%%%%%%%%%%
% SOME VARIABLES
%%%%%%%%%%%%%%%%%%%%%%%%%%%%%%%%%%%%%%%%%%%%%%%                       
ruleIndex = 1;
parameterIndex = 1;
compartmentIndex = 1;
specieIndex = 1;
                       
%%%%%%%%%%%%%%%%%%%%%%%%%%%%%%%%%%%%%%%%%%%%%%%
% COPY NAME AND NOTES
%%%%%%%%%%%%%%%%%%%%%%%%%%%%%%%%%%%%%%%%%%%%%%%                       
SBMLstructure.name = ms.name;
SBMLstructure.notes = convert2SBMLNotes(ms.notes);

%%%%%%%%%%%%%%%%%%%%%%%%%%%%%%%%%%%%%%%%%%%%%%%
% PROCESS COMPARTMENTS
%%%%%%%%%%%%%%%%%%%%%%%%%%%%%%%%%%%%%%%%%%%%%%%                       
% compartments can be defined as parameters, variables or states. 
% If the field 'type' is set to 'isCompartment' it indicates a compartment. 
% check parameters for compartments
for k = 1:length(ms.parameters),
    if strcmp(ms.parameters(k).type,'isCompartment'),
        % current parameter determines compartment size
        % always use spatial dimensions 3
        SBMLstructure.compartment(compartmentIndex).typecode = 'SBML_COMPARTMENT';
        SBMLstructure.compartment(compartmentIndex).metaid = '';
        SBMLstructure.compartment(compartmentIndex).notes = convert2SBMLNotes(ms.parameters(k).notes);
        SBMLstructure.compartment(compartmentIndex).annotation = '';
        SBMLstructure.compartment(compartmentIndex).name = ms.parameters(k).name;
        SBMLstructure.compartment(compartmentIndex).id = ms.parameters(k).name;
        SBMLstructure.compartment(compartmentIndex).spatialDimensions = 3;
        SBMLstructure.compartment(compartmentIndex).size = ms.parameters(k).value;
        SBMLstructure.compartment(compartmentIndex).units = '';
        SBMLstructure.compartment(compartmentIndex).outside = ms.parameters(k).compartment;
        SBMLstructure.compartment(compartmentIndex).constant = 1;
        SBMLstructure.compartment(compartmentIndex).isSetSize = 1;
        SBMLstructure.compartment(compartmentIndex).isSetVolume = 1;
        compartmentIndex = compartmentIndex + 1;
    end
end
% check states for compartments
for k = 1:length(ms.states),
    if strcmp(ms.states(k).type,'isCompartment'),
        % current state determines compartment size
        % set size to initial condition of state
        SBMLstructure.compartment(compartmentIndex).typecode = 'SBML_COMPARTMENT';
        SBMLstructure.compartment(compartmentIndex).metaid = '';
        SBMLstructure.compartment(compartmentIndex).notes = convert2SBMLNotes(ms.states(k).notes);
        SBMLstructure.compartment(compartmentIndex).annotation = '';
        SBMLstructure.compartment(compartmentIndex).name = ms.states(k).name;
        SBMLstructure.compartment(compartmentIndex).id = ms.states(k).name;
        SBMLstructure.compartment(compartmentIndex).spatialDimensions = 3;
        SBMLstructure.compartment(compartmentIndex).size = ms.states(k).initialCondition;
        SBMLstructure.compartment(compartmentIndex).units = '';
        SBMLstructure.compartment(compartmentIndex).outside = ms.states(k).compartment;
        SBMLstructure.compartment(compartmentIndex).constant = 0;
        SBMLstructure.compartment(compartmentIndex).isSetSize = 1;
        SBMLstructure.compartment(compartmentIndex).isSetVolume = 1;
        compartmentIndex = compartmentIndex + 1;
        % Compartment defined by a rule. Include this rule in the structure
        SBMLstructure.rule(ruleIndex).typecode = 'SBML_RATE_RULE';
        SBMLstructure.rule(ruleIndex).metaid = '';
        SBMLstructure.rule(ruleIndex).notes = convert2SBMLNotes(ms.states(k).notes);
        SBMLstructure.rule(ruleIndex).annotation = '';
        SBMLstructure.rule(ruleIndex).formula = replaceMATLABexpressions(ms.states(k).ODE);
        SBMLstructure.rule(ruleIndex).variable = ms.states(k).name;
        SBMLstructure.rule(ruleIndex).species = '';
        SBMLstructure.rule(ruleIndex).compartment = '';
        SBMLstructure.rule(ruleIndex).name = '';
        SBMLstructure.rule(ruleIndex).units = '';
        ruleIndex = ruleIndex + 1;        
    end
end

%%%%%%%%%%%%%%%%%%%%%%%%%%%%%%%%%%%%%%%%%%%%%%%
% PROCESS PARAMETERS
%%%%%%%%%%%%%%%%%%%%%%%%%%%%%%%%%%%%%%%%%%%%%%% 
% parameters can be defined as parameters, variables or states. If the
% field 'type' is set to 'isParameter' it indicates an SBML parameter.  
% we only will have global parameters in the SBML model.
% check parameters for parameters
for k = 1:length(ms.parameters),
    if strcmp(ms.parameters(k).type,'isParameter'),
        % current parameter determines compartment size
        SBMLstructure.parameter(parameterIndex).typecode = 'SBML_PARAMETER';
        SBMLstructure.parameter(parameterIndex).metaid = '';
        SBMLstructure.parameter(parameterIndex).notes = convert2SBMLNotes(ms.parameters(k).notes);
        SBMLstructure.parameter(parameterIndex).annotation = '';
        SBMLstructure.parameter(parameterIndex).name = ms.parameters(k).name;
        SBMLstructure.parameter(parameterIndex).id = ms.parameters(k).name;
        SBMLstructure.parameter(parameterIndex).value = ms.parameters(k).value;
        SBMLstructure.parameter(parameterIndex).units = '';
        SBMLstructure.parameter(parameterIndex).constant = 1;
        SBMLstructure.parameter(parameterIndex).isSetValue = 1;
        parameterIndex = parameterIndex + 1;
    end
end
% check states for parameters
for k = 1:length(ms.states),
    if strcmp(ms.states(k).type,'isParameter'),
        % current state determines compartment size
        SBMLstructure.parameter(parameterIndex).typecode = 'SBML_PARAMETER';
        SBMLstructure.parameter(parameterIndex).metaid = '';
        SBMLstructure.parameter(parameterIndex).notes = convert2SBMLNotes(ms.states(k).notes);
        SBMLstructure.parameter(parameterIndex).annotation = '';
        SBMLstructure.parameter(parameterIndex).name = ms.states(k).name;
        SBMLstructure.parameter(parameterIndex).id = ms.states(k).name;
        SBMLstructure.parameter(parameterIndex).value = ms.states(k).initialCondition;
        SBMLstructure.parameter(parameterIndex).units = '';
        SBMLstructure.parameter(parameterIndex).constant = 0;
        SBMLstructure.parameter(parameterIndex).isSetValue = 1;
        parameterIndex = parameterIndex + 1;        
        % Parameter defined by a rule. Include this rule in the structure
        SBMLstructure.rule(ruleIndex).typecode = 'SBML_RATE_RULE';
        SBMLstructure.rule(ruleIndex).metaid = '';
        SBMLstructure.rule(ruleIndex).notes = convert2SBMLNotes(ms.states(k).notes);
        SBMLstructure.rule(ruleIndex).annotation = '';
        SBMLstructure.rule(ruleIndex).formula = replaceMATLABexpressions(ms.states(k).ODE);
        SBMLstructure.rule(ruleIndex).variable = ms.states(k).name;
        SBMLstructure.rule(ruleIndex).species = '';
        SBMLstructure.rule(ruleIndex).compartment = '';
        SBMLstructure.rule(ruleIndex).name = '';
        SBMLstructure.rule(ruleIndex).units = '';
        ruleIndex = ruleIndex + 1;        
    end
end            

%%%%%%%%%%%%%%%%%%%%%%%%%%%%%%%%%%%%%%%%%%%%%%%
% PROCESS FUNCTIONS
%%%%%%%%%%%%%%%%%%%%%%%%%%%%%%%%%%%%%%%%%%%%%%% 
% add functions
for k = 1:length(ms.functions),
    SBMLstructure.functionDefinition(k).typecode = 'SBML_FUNCTION_DEFINITION';
    SBMLstructure.functionDefinition(k).metaid = '';
    SBMLstructure.functionDefinition(k).notes = convert2SBMLNotes(ms.functions(k).notes);
    SBMLstructure.functionDefinition(k).annotation = '';
    SBMLstructure.functionDefinition(k).name = ms.functions(k).name;
    SBMLstructure.functionDefinition(k).id = ms.functions(k).name;
    SBMLstructure.functionDefinition(k).math = sprintf('lambda(%s,%s)',ms.functions(k).arguments,replaceMATLABexpressions(ms.functions(k).formula));
end

%%%%%%%%%%%%%%%%%%%%%%%%%%%%%%%%%%%%%%%%%%%%%%%
% PROCESS CONSTANT (BOUNDARY) SPECIES
%%%%%%%%%%%%%%%%%%%%%%%%%%%%%%%%%%%%%%%%%%%%%%% 
% constant species can only be defined as parameters.
% If the field 'type' of a parameter is set to 'isSpecie' we assume that
% this indicates a constant boundary species.
for k = 1:length(ms.parameters),
    if strcmp(ms.parameters(k).type,'isSpecie'),
        % Determine if amount or concentration
        if strcmp(ms.parameters(k).unittype,'amount'), 
            isAmount = 1;
            isConcentration = 0;
            hasOnlySubstanceUnits = 1;
        elseif strcmp(ms.parameters(k).unittype,'concentration'),
            isAmount = 0;
            isConcentration = 1;
            hasOnlySubstanceUnits = 0;
        end
        % import constant boundary species
        SBMLstructure.species(specieIndex).typecode = 'SBML_SPECIES';
        SBMLstructure.species(specieIndex).metaid = '';
        SBMLstructure.species(specieIndex).notes = convert2SBMLNotes(ms.parameters(k).notes);
        SBMLstructure.species(specieIndex).annotation = '';
        SBMLstructure.species(specieIndex).name = ms.parameters(k).name;
        SBMLstructure.species(specieIndex).id = ms.parameters(k).name;
        SBMLstructure.species(specieIndex).compartment = ms.parameters(k).compartment;
        SBMLstructure.species(specieIndex).initialAmount = ms.parameters(k).value;
        SBMLstructure.species(specieIndex).initialConcentration = ms.parameters(k).value;
        SBMLstructure.species(specieIndex).substanceUnits = '';         % no units!
        SBMLstructure.species(specieIndex).spatialSizeUnits = '';       % no units!
        SBMLstructure.species(specieIndex).hasOnlySubstanceUnits = hasOnlySubstanceUnits;
        SBMLstructure.species(specieIndex).boundaryCondition = 1;       % boundary 
        SBMLstructure.species(specieIndex).charge = 0;                  
        SBMLstructure.species(specieIndex).constant = 1;                % species is set constant
        SBMLstructure.species(specieIndex).isSetInitialAmount = isAmount;
        SBMLstructure.species(specieIndex).isSetInitialConcentration = isConcentration;
        SBMLstructure.species(specieIndex).isSetCharge = 0;             % charge not supported
        specieIndex = specieIndex + 1;
    end
end

%%%%%%%%%%%%%%%%%%%%%%%%%%%%%%%%%%%%%%%%%%%%%%%
% PROCESS BOUNDARY SPECIES
%%%%%%%%%%%%%%%%%%%%%%%%%%%%%%%%%%%%%%%%%%%%%%% 
% boundary species can be defined variables or states (not as parameters,
% since then they become constant boundary species).
% The difference between species and boundary species is here that species
% are only defined by reactions, while boundary species are defined by
% rules. The variable "SBMLspeciesNames" contains the names of all states
% that are defined only by reactions.
% now check states (boundary if isSpecie and not in the SBMLspeciesNames list)
for k = 1:length(ms.states),
    if strcmp(ms.states(k).type,'isSpecie') && isempty(strmatchIQM(ms.states(k).name,SBMLspeciesNames)),
        % Determine if amount or concentration
        if strcmp(ms.states(k).unittype,'amount') 
            isAmount = 1;
            isConcentration = 0;
            hasOnlySubstanceUnits = 1;
        elseif strcmp(ms.states(k).unittype,'concentration') 
            isAmount = 0;
            isConcentration = 1;
            hasOnlySubstanceUnits = 0;
        end
        % import boundary species
        SBMLstructure.species(specieIndex).typecode = 'SBML_SPECIES';
        SBMLstructure.species(specieIndex).metaid = '';
        SBMLstructure.species(specieIndex).notes = convert2SBMLNotes(ms.states(k).notes);
        SBMLstructure.species(specieIndex).annotation = '';
        SBMLstructure.species(specieIndex).name = ms.states(k).name;
        SBMLstructure.species(specieIndex).id = ms.states(k).name;
        SBMLstructure.species(specieIndex).compartment = ms.states(k).compartment;
        SBMLstructure.species(specieIndex).initialAmount = ms.states(k).initialCondition;
        SBMLstructure.species(specieIndex).initialConcentration = ms.states(k).initialCondition;
        SBMLstructure.species(specieIndex).substanceUnits = '';         % no units!
        SBMLstructure.species(specieIndex).spatialSizeUnits = '';       % no units!
        SBMLstructure.species(specieIndex).hasOnlySubstanceUnits = hasOnlySubstanceUnits;
        SBMLstructure.species(specieIndex).boundaryCondition = 1;       % boundary 
        SBMLstructure.species(specieIndex).charge = 0;                  
        SBMLstructure.species(specieIndex).constant = 0;                % species is not set constant
        SBMLstructure.species(specieIndex).isSetInitialAmount = isAmount;
        SBMLstructure.species(specieIndex).isSetInitialConcentration = isConcentration;
        SBMLstructure.species(specieIndex).isSetCharge = 0;             % charge not supported
        specieIndex = specieIndex + 1;
        % Boundary species defined by a rule. Include this rule in the structure
        SBMLstructure.rule(ruleIndex).typecode = 'SBML_RATE_RULE';
        SBMLstructure.rule(ruleIndex).metaid = '';
        SBMLstructure.rule(ruleIndex).notes = convert2SBMLNotes(ms.states(k).notes);
        SBMLstructure.rule(ruleIndex).annotation = '';
        SBMLstructure.rule(ruleIndex).formula = replaceMATLABexpressions(ms.states(k).ODE);
        SBMLstructure.rule(ruleIndex).variable = ms.states(k).name;
        SBMLstructure.rule(ruleIndex).species = '';
        SBMLstructure.rule(ruleIndex).compartment = '';
        SBMLstructure.rule(ruleIndex).name = '';
        SBMLstructure.rule(ruleIndex).units = '';
        ruleIndex = ruleIndex + 1;
    end
end

%%%%%%%%%%%%%%%%%%%%%%%%%%%%%%%%%%%%%%%%%%%%%%%
% PROCESS SPECIES
%%%%%%%%%%%%%%%%%%%%%%%%%%%%%%%%%%%%%%%%%%%%%%% 
% we assume that species are only defined as states and determined by
% reactions
% check simple species defined by reactions
% save the species name and the species ODE for use when processing the
% reactions
speciesReactions = [];
for k = 1:length(ms.states),
    if strcmp(ms.states(k).type,'isSpecie') && ~isempty(strmatchIQM(ms.states(k).name,SBMLspeciesNames)),
        % Determine if amount or concentration
        if strcmp(ms.states(k).unittype,'amount') 
            isAmount = 1;
            isConcentration = 0;
            hasOnlySubstanceUnits = 1;
        elseif strcmp(ms.states(k).unittype,'concentration') 
            isAmount = 0;
            isConcentration = 1;
            hasOnlySubstanceUnits = 0;
        end
        % import constant species
        SBMLstructure.species(specieIndex).typecode = 'SBML_SPECIES';
        SBMLstructure.species(specieIndex).metaid = '';
        SBMLstructure.species(specieIndex).notes = convert2SBMLNotes(ms.states(k).notes);
        SBMLstructure.species(specieIndex).annotation = '';
        SBMLstructure.species(specieIndex).name = ms.states(k).name;
        SBMLstructure.species(specieIndex).id = ms.states(k).name;
        SBMLstructure.species(specieIndex).compartment = ms.states(k).compartment;
        SBMLstructure.species(specieIndex).initialAmount = ms.states(k).initialCondition;
        SBMLstructure.species(specieIndex).initialConcentration = ms.states(k).initialCondition;
        SBMLstructure.species(specieIndex).substanceUnits = '';         % no units!
        SBMLstructure.species(specieIndex).spatialSizeUnits = '';       % no units!
        SBMLstructure.species(specieIndex).hasOnlySubstanceUnits = hasOnlySubstanceUnits;
        SBMLstructure.species(specieIndex).boundaryCondition = 0;       % not boundary 
        SBMLstructure.species(specieIndex).charge = 0;                  
        SBMLstructure.species(specieIndex).constant = 0;                % species is not set constant
        SBMLstructure.species(specieIndex).isSetInitialAmount = isAmount;
        SBMLstructure.species(specieIndex).isSetInitialConcentration = isConcentration;
        SBMLstructure.species(specieIndex).isSetCharge = 0;             % charge not supported
        specieIndex = specieIndex + 1;
    end
end

%%%%%%%%%%%%%%%%%%%%%%%%%%%%%%%%%%%%%%%%%%%%%%%
% PROCESS ASSIGNMENT RULES ... NEED TO BE ORDERED ... THUS HERE
%%%%%%%%%%%%%%%%%%%%%%%%%%%%%%%%%%%%%%%%%%%%%%%
for k = 1:length(ms.variables),
    if strcmp(ms.variables(k).type,'isCompartment'),
        % current variable determines compartment size
        % set size to 1 (not important, since not used)
        SBMLstructure.compartment(compartmentIndex).typecode = 'SBML_COMPARTMENT';
        SBMLstructure.compartment(compartmentIndex).metaid = '';
        SBMLstructure.compartment(compartmentIndex).notes = convert2SBMLNotes(ms.variables(k).notes);
        SBMLstructure.compartment(compartmentIndex).annotation = '';
        SBMLstructure.compartment(compartmentIndex).name = ms.variables(k).name;
        SBMLstructure.compartment(compartmentIndex).id = ms.variables(k).name;
        SBMLstructure.compartment(compartmentIndex).spatialDimensions = 3;
        SBMLstructure.compartment(compartmentIndex).size = 1;
        SBMLstructure.compartment(compartmentIndex).units = '';
        SBMLstructure.compartment(compartmentIndex).outside = ms.variables(k).compartment;
        SBMLstructure.compartment(compartmentIndex).constant = 0;
        SBMLstructure.compartment(compartmentIndex).isSetSize = 1;
        SBMLstructure.compartment(compartmentIndex).isSetVolume = 1;
        compartmentIndex = compartmentIndex + 1;
        % Compartment defined by a rule. Include this rule in the structure
        SBMLstructure.rule(ruleIndex).typecode = 'SBML_ASSIGNMENT_RULE';
        SBMLstructure.rule(ruleIndex).metaid = '';
        SBMLstructure.rule(ruleIndex).notes = convert2SBMLNotes(ms.variables(k).notes);
        SBMLstructure.rule(ruleIndex).annotation = '';
        SBMLstructure.rule(ruleIndex).formula = replaceMATLABexpressions(ms.variables(k).formula);
        SBMLstructure.rule(ruleIndex).variable = ms.variables(k).name;
        SBMLstructure.rule(ruleIndex).species = '';
        SBMLstructure.rule(ruleIndex).compartment = '';
        SBMLstructure.rule(ruleIndex).name = '';
        SBMLstructure.rule(ruleIndex).units = '';
        ruleIndex = ruleIndex + 1;
    end
    if strcmp(ms.variables(k).type,'isParameter'),
        % current variable determines compartment size
        % value set to zero (not needed)
        SBMLstructure.parameter(parameterIndex).typecode = 'SBML_PARAMETER';
        SBMLstructure.parameter(parameterIndex).metaid = '';
        SBMLstructure.parameter(parameterIndex).notes = convert2SBMLNotes(ms.variables(k).notes);
        SBMLstructure.parameter(parameterIndex).annotation = '';
        SBMLstructure.parameter(parameterIndex).name = ms.variables(k).name;
        SBMLstructure.parameter(parameterIndex).id = ms.variables(k).name;
        SBMLstructure.parameter(parameterIndex).value = 0;  % no value needed (assignment rule)
        SBMLstructure.parameter(parameterIndex).units = '';
        SBMLstructure.parameter(parameterIndex).constant = 0;
        SBMLstructure.parameter(parameterIndex).isSetValue = 1;
        parameterIndex = parameterIndex + 1;        
        % Parameter defined by a rule. Include this rule in the structure
        SBMLstructure.rule(ruleIndex).typecode = 'SBML_ASSIGNMENT_RULE';
        SBMLstructure.rule(ruleIndex).metaid = '';
        SBMLstructure.rule(ruleIndex).notes = convert2SBMLNotes(ms.variables(k).notes);
        SBMLstructure.rule(ruleIndex).annotation = '';
        SBMLstructure.rule(ruleIndex).formula = replaceMATLABexpressions(ms.variables(k).formula);
        SBMLstructure.rule(ruleIndex).variable = ms.variables(k).name;
        SBMLstructure.rule(ruleIndex).species = '';
        SBMLstructure.rule(ruleIndex).compartment = '';
        SBMLstructure.rule(ruleIndex).name = '';
        SBMLstructure.rule(ruleIndex).units = '';
        ruleIndex = ruleIndex + 1;
    end
    if strcmp(ms.variables(k).type,'isSpecie'),
        % Determine if amount or concentration
        if strcmp(ms.variables(k).unittype,'amount'), 
            isAmount = 1;
            isConcentration = 0;
            hasOnlySubstanceUnits = 1;
        elseif strcmp(ms.variables(k).unittype,'concentration'), 
            isAmount = 0;
            isConcentration = 1;
            hasOnlySubstanceUnits = 0;
        end
        % import boundary species
        SBMLstructure.species(specieIndex).typecode = 'SBML_SPECIES';
        SBMLstructure.species(specieIndex).metaid = '';
        SBMLstructure.species(specieIndex).notes = convert2SBMLNotes(ms.variables(k).notes);
        SBMLstructure.species(specieIndex).annotation = '';
        SBMLstructure.species(specieIndex).name = ms.variables(k).name;
        SBMLstructure.species(specieIndex).id = ms.variables(k).name;
        SBMLstructure.species(specieIndex).compartment = ms.variables(k).compartment;
        SBMLstructure.species(specieIndex).initialAmount = 0;  % not set (assignment rule)
        SBMLstructure.species(specieIndex).initialConcentration = 0; % not set (assignment rule)
        SBMLstructure.species(specieIndex).substanceUnits = '';         % no units!
        SBMLstructure.species(specieIndex).spatialSizeUnits = '';       % no units!
        SBMLstructure.species(specieIndex).hasOnlySubstanceUnits = hasOnlySubstanceUnits;
        SBMLstructure.species(specieIndex).boundaryCondition = 1;       % boundary 
        SBMLstructure.species(specieIndex).charge = 0;                  
        SBMLstructure.species(specieIndex).constant = 0;                % species is not set constant
        SBMLstructure.species(specieIndex).isSetInitialAmount = isAmount;
        SBMLstructure.species(specieIndex).isSetInitialConcentration = isConcentration;
        SBMLstructure.species(specieIndex).isSetCharge = 0;             % charge not supported
        specieIndex = specieIndex + 1;
        % Boundary species defined by a rule. Include this rule in the structure
        SBMLstructure.rule(ruleIndex).typecode = 'SBML_ASSIGNMENT_RULE';
        SBMLstructure.rule(ruleIndex).metaid = '';
        SBMLstructure.rule(ruleIndex).notes = convert2SBMLNotes(ms.variables(k).notes);
        SBMLstructure.rule(ruleIndex).annotation = '';
        SBMLstructure.rule(ruleIndex).formula = replaceMATLABexpressions(ms.variables(k).formula);
        SBMLstructure.rule(ruleIndex).variable = ms.variables(k).name;
        SBMLstructure.rule(ruleIndex).species = '';
        SBMLstructure.rule(ruleIndex).compartment = '';
        SBMLstructure.rule(ruleIndex).name = '';
        SBMLstructure.rule(ruleIndex).units = '';
        ruleIndex = ruleIndex + 1;
    end
end

%%%%%%%%%%%%%%%%%%%%%%%%%%%%%%%%%%%%%%%%%%%%%%%
% PROCESS REACTIONS
%%%%%%%%%%%%%%%%%%%%%%%%%%%%%%%%%%%%%%%%%%%%%%%
% Before processing the reactions we are checking for moiety conservations
% and reversing them into the stoichiometric matrix (to be able to draw the
% full picture)
[StoichiometricMatrix,SBMLspeciesNames] = reverseMoietyConservationsIQM(iqm, 1);
for k1 = 1:length(ms.reactions),
    reactionName = ms.reactions(k1).name;
    reactionFormula = ms.reactions(k1).formula;
    % get needed information about reaction
    [reactant, product, modifier] = getReactionInfo(reactionName,StoichiometricMatrix,SBMLspeciesNames,SBMLreactionNames,SBMLreversibleFlag,iqm,ms);
    % add reaction to the structure
    SBMLstructure.reaction(k1).typecode = 'SBML_REACTION';
    SBMLstructure.reaction(k1).metaid = '';
    SBMLstructure.reaction(k1).notes = convert2SBMLNotes(ms.reactions(k1).notes);
    SBMLstructure.reaction(k1).annotation = '';
    SBMLstructure.reaction(k1).name = reactionName;
    SBMLstructure.reaction(k1).id = reactionName;
    
    SBMLstructure.reaction(k1).reactant = reactant;
    SBMLstructure.reaction(k1).product = product;
    SBMLstructure.reaction(k1).modifier = modifier; 
    
    SBMLstructure.reaction(k1).kineticLaw.typecode = 'SBML_KINETIC_LAW';
    SBMLstructure.reaction(k1).kineticLaw.metaid = '';
    SBMLstructure.reaction(k1).kineticLaw.notes = '';
    SBMLstructure.reaction(k1).kineticLaw.annotation = '';
    SBMLstructure.reaction(k1).kineticLaw.formula = replaceMATLABexpressions(reactionFormula);
    SBMLstructure.reaction(k1).kineticLaw.math = replaceMATLABexpressions(reactionFormula);
    SBMLstructure.reaction(k1).kineticLaw.parameter = kineticLawParameterStruct;    % no local parameters
    
    SBMLstructure.reaction(k1).kineticLaw.timeUnits = '';
    SBMLstructure.reaction(k1).kineticLaw.substanceUnits = '';
    
    SBMLstructure.reaction(k1).reversible = ms.reactions(k1).reversible;
    
    % handle set the fast flag
    if ms.reactions(k1).fast == 0,
        SBMLstructure.reaction(k1).fast = 0;
        SBMLstructure.reaction(k1).isSetFast = 0;
    else
        SBMLstructure.reaction(k1).fast = 1;
        SBMLstructure.reaction(k1).isSetFast = 1;
    end        
end

%%%%%%%%%%%%%%%%%%%%%%%%%%%%%%%%%%%%%%%%%%%%%%%
% LEAVE UNITDEFINITIONS EMPTY
%%%%%%%%%%%%%%%%%%%%%%%%%%%%%%%%%%%%%%%%%%%%%%%                       

%%%%%%%%%%%%%%%%%%%%%%%%%%%%%%%%%%%%%%%%%%%%%%%
% Process Events
%%%%%%%%%%%%%%%%%%%%%%%%%%%%%%%%%%%%%%%%%%%%%%%                       
for k1=1:length(ms.events),
    SBMLstructure.event(k1).typecode = 'SBML_EVENT';
    SBMLstructure.event(k1).metaid = '';
    SBMLstructure.event(k1).notes = convert2SBMLNotes(ms.events(k1).notes);
    SBMLstructure.event(k1).annotation = '';
    SBMLstructure.event(k1).name = ms.events(k1).name;
    SBMLstructure.event(k1).id = ms.events(k1).name;
    SBMLstructure.event(k1).trigger = replaceMATLABexpressions(ms.events(k1).trigger);
    SBMLstructure.event(k1).delay = '0'; % always zero
    SBMLstructure.event(k1).timeUnits = '';
    for k2 = 1:length(ms.events(k1).assignment),
        SBMLstructure.event(k1).eventAssignment(k2).typecode = 'SBML_EVENT_ASSIGNMENT';
        SBMLstructure.event(k1).eventAssignment(k2).metaid = '';
        SBMLstructure.event(k1).eventAssignment(k2).notes = '';
        SBMLstructure.event(k1).eventAssignment(k2).annotation = '';
        SBMLstructure.event(k1).eventAssignment(k2).variable = ms.events(k1).assignment(k2).variable;
        SBMLstructure.event(k1).eventAssignment(k2).math =  replaceMATLABexpressions(ms.events(k1).assignment(k2).formula);
    end
end

%%%%%%%%%%%%%%%%%%%%%%%%%%%%%%%%%%%%%%%%%%%%%%%
% Process Algebraic Rules
%%%%%%%%%%%%%%%%%%%%%%%%%%%%%%%%%%%%%%%%%%%%%%%                       
% algebraic rules are just added. names NEED to be present otherwise it
% makes no sense
for k=1:length(ms.algebraic),
    name = ms.algebraic(k).name;
    formula = ms.algebraic(k).formula;
    initialCondition = ms.algebraic(k).initialCondition;
    type = ms.algebraic(k).type;
    compartment = ms.algebraic(k).compartment;
    unittype = ms.algebraic(k).unittype;
    notes = ms.algebraic(k).notes;
    % add the rule
    SBMLstructure.rule(end+1).typecode = 'SBML_ALGEBRAIC_RULE';
    SBMLstructure.rule(end).metaid = '';
    SBMLstructure.rule(end).notes = convert2SBMLNotes(notes);
    SBMLstructure.rule(end).annotation = '';
    SBMLstructure.rule(end).formula = formula;
    SBMLstructure.rule(end).variable = '';
    SBMLstructure.rule(end).species = '';
    SBMLstructure.rule(end).compartment = '';
    SBMLstructure.rule(end).name = '';
    SBMLstructure.rule(end).units = '';
    % add name as specie, parameter, or compartment
    if strcmp(type,'isSpecie'),
        SBMLstructure.species(end+1).typecode = 'SBML_SPECIES';
        SBMLstructure.species(end).metaid = '';
        SBMLstructure.species(end).notes = convert2SBMLNotes(notes);
        SBMLstructure.species(end).annotation = '';
        SBMLstructure.species(end).name = name;
        SBMLstructure.species(end).id = name;
        SBMLstructure.species(end).compartment = compartment;
        SBMLstructure.species(end).initialAmount = initialCondition;
        SBMLstructure.species(end).initialConcentration = initialCondition;
        SBMLstructure.species(end).substanceUnits = '';         % no units!
        SBMLstructure.species(end).spatialSizeUnits = '';       % no units!
        SBMLstructure.species(end).hasOnlySubstanceUnits = strcmp(unittype,'amount');
        SBMLstructure.species(end).boundaryCondition = 1;       % set boundary cond (det. by rule) 
        SBMLstructure.species(end).charge = 0;                  
        SBMLstructure.species(end).constant = 0;                % species is not set constant
        SBMLstructure.species(end).isSetInitialAmount = strcmp(unittype,'amount');
        SBMLstructure.species(end).isSetInitialConcentration = strcmp(unittype,'concentration');
        SBMLstructure.species(end).isSetCharge = 0;             % charge not supported       
    end
    if strcmp(type,'isParameter'),
        SBMLstructure.parameter(end+1).typecode = 'SBML_PARAMETER';
        SBMLstructure.parameter(end).metaid = '';
        SBMLstructure.parameter(end).notes = convert2SBMLNotes(notes);
        SBMLstructure.parameter(end).annotation = '';
        SBMLstructure.parameter(end).name = name;
        SBMLstructure.parameter(end).id = name;
        SBMLstructure.parameter(end).value = initialCondition;
        SBMLstructure.parameter(end).units = '';
        SBMLstructure.parameter(end).constant = 0;
        SBMLstructure.parameter(end).isSetValue = 1;
    end
    if strcmp(type,'isCompartment'),
        SBMLstructure.compartment(end+1).typecode = 'SBML_COMPARTMENT';
        SBMLstructure.compartment(end).metaid = '';
        SBMLstructure.compartment(end).notes = convert2SBMLNotes(notes);
        SBMLstructure.compartment(end).annotation = '';
        SBMLstructure.compartment(end).name = name;
        SBMLstructure.compartment(end).id = name;
        SBMLstructure.compartment(end).spatialDimensions = 3;
        SBMLstructure.compartment(end).size = initialCondition;
        SBMLstructure.compartment(end).units = '';
        SBMLstructure.compartment(end).outside = compartment;
        SBMLstructure.compartment(end).constant = 0;
        SBMLstructure.compartment(end).isSetSize = 1;
        SBMLstructure.compartment(end).isSetVolume = 1;
    end    
end

%%%%%%%%%%%%%%%%%%%%%%%%%%%%%%%%%%%%%%%%%%%%%%%
% WRITE SBML FILE
%%%%%%%%%%%%%%%%%%%%%%%%%%%%%%%%%%%%%%%%%%%%%%%                       
result = SBMLstructure;
try 
    if nargin==1,
        OutputSBML(SBMLstructure);
    else
        OutputSBML(SBMLstructure,strrep(filename,'.xml',''));
    end
catch
    if ~strcmp(lasterr, 'Cannot copy filename'),
        warning(sprintf('Error occurred: %s\n',lasterr));
    end
end
return


%%%%%%%%%%%%%%%%%%%%%%%%%%%%%%%%%%%%%%%%%%%%%%%
% DETERMINE REACTION INFORMATION
%%%%%%%%%%%%%%%%%%%%%%%%%%%%%%%%%%%%%%%%%%%%%%%                       
% we look through all ODEs where the gioven reaction is involved in.
% we determine reversibility, reactants, products, and modifiers.
function [reactant, product, modifier] = getReactionInfo(reactionName,StoichiometricMatrix,SBMLspeciesNames,SBMLreactionNames,SBMLreversibleFlag,iqm,ms)
    % initialize the structures
    reactant = struct('typecode', {}, 'metaid', {}, 'notes', {}, 'annotation', {}, 'species', {}, 'stoichiometry', {}, 'denominator', {}, 'stoichiometryMath', {});
    product = struct('typecode', {}, 'metaid', {}, 'notes', {}, 'annotation', {}, 'species', {}, 'stoichiometry', {}, 'denominator', {}, 'stoichiometryMath', {});
    modifier = struct('typecode', {}, 'metaid', {}, 'notes', {}, 'annotation', {}, 'species', {});
    % get index of current reaction
    reactionIndex = strmatchIQM(reactionName,SBMLreactionNames,'exact');
    % get corresponding stoichiometries
    [stoichRows, stoichCols] = size(StoichiometricMatrix);
    if (stoichCols >= reactionIndex),
        Nreaction = StoichiometricMatrix(:,reactionIndex);
    else
        % for the current reaction there's no stoichiometry
        % or products, reactants or modifierer found
        return
    end
    % get reactant indices (Nreaction < 0)
    reactantIndices = find(Nreaction < 0);
    % get product indices (Nreaction > 0)
    productIndices = find(Nreaction > 0);
    % add reactants
    for k = 1:length(reactantIndices),
        reactant(k).typecode = 'SBML_SPECIES_REFERENCE';
        reactant(k).metaid = '';
        reactant(k).notes = '';
        reactant(k).annotation = '';
        reactant(k).species = SBMLspeciesNames{reactantIndices(k)};
        reactant(k).stoichiometry = abs(Nreaction(reactantIndices(k)));
        reactant(k).denominator = 1;
        reactant(k).stoichiometryMath = '';
    end
    % add products
    for k = 1:length(productIndices),
        product(k).typecode = 'SBML_SPECIES_REFERENCE';
        product(k).metaid = '';
        product(k).notes = '';
        product(k).annotation = '';
        product(k).species = SBMLspeciesNames{productIndices(k)};
        product(k).stoichiometry = Nreaction(productIndices(k));
        product(k).denominator = 1;
        product(k).stoichiometryMath = '';
    end
    % search for modifiers among the non reactants and non products and add them
    specieListPossible = SBMLspeciesNames(find(Nreaction == 0));
    modifierList = {};
    modifierList = search4Modifiers(modifierList,ms.reactions(reactionIndex).formula,specieListPossible,{ms.variables.name},iqm,ms);
    modifierList = unique(modifierList);
    for k = 1:length(modifierList),
        modifier(k).typecode = 'SBML_MODIFIER_SPECIES_REFERENCE';
        modifier(k).metaid = '';
        modifier(k).notes = '';
        modifier(k).annotation = '';
        modifier(k).species = modifierList{k};
    end
return

%%%%%%%%%%%%%%%%%%%%%%%%%%%%%%%%%%%%%%%%%%%%%%%
% DETERMINE REACTION INFORMATION
%%%%%%%%%%%%%%%%%%%%%%%%%%%%%%%%%%%%%%%%%%%%%%%                       
% we go through all reactions and eventually variables to determine the
% modifiers (recursively)
function [modifierList] = search4Modifiers(modifierList,formula,specieListPossible,variableNames,iqm,ms)
    % search in current formula for modifiers
    formula = strcat('#',formula,'#');
    for k=1:length(specieListPossible),
        if ~isempty(regexp(formula,strcat('\W',specieListPossible{k},'\W'))),
            modifierList{end+1} = specieListPossible{k};
        end
    end
    % search in current formula for variable names
    searchVariablesIndices = [];
    for k=1:length(variableNames),
        if ~isempty(regexp(formula,strcat('\W',variableNames{k},'\W'))),
            searchVariablesIndices(end+1) = k;
        end
    end
    % loop through variable names and search for modifiers
    for k=1:length(searchVariablesIndices),
        formula = ms.variables(searchVariablesIndices(k)).formula;
        modifierList = search4Modifiers(modifierList,formula,specieListPossible,variableNames,iqm,ms);
    end
return

%%%%%%%%%%%%%%%%%%%%%%%%%%%%%%%%%%%%%%%%%%%%%%%%%%%%%%%%%%%%%%%%%%%%%%%%%%
% EXCHANGE MATLAB FUNCTION NAMES TO MATHML FUNCTION NAMES IN ALL FORMULAS
%%%%%%%%%%%%%%%%%%%%%%%%%%%%%%%%%%%%%%%%%%%%%%%%%%%%%%%%%%%%%%%%%%%%%%%%%%
% MathML functions can have different names than MATLAB function names
% we need to exchange them in the formula strings. The term 'MathML
% expression' relates to what is required by OutputSBML!
function [formula] = replaceMATLABexpressions(formula)
% MathML expressions 
MathMLexpressions = {'arccos(','arcsin(','arctan(','ceiling(','ln(','log(10,',...
    'pow(','arccos(','arccosh(','arccot(','arccoth(','arccsc(','arccsch(',...
    'arcsec(','arcsech(','arcsin(','arcsinh(','arctan(','arctanh(',...
    'exponentiale','log(','piecewise(','leq(','geq(','and(','or('};
% Corresponding MATLAB expressions
MATLABexpressions = {'acos(','asin(','atan(','ceil(','log(','log10(',...
    'power(','acos(','acosh(','acot(','acoth(','acsc(','acsch(',...
    'asec(','asech(','asin(','asinh(','atan(','atanh('...
    'exp(1)','logbase(','piecewiseIQM(','le(','ge(','andIQM(','orIQM('};
% do the simple replace
for k = 1:length(MathMLexpressions),
    formula = strrep(formula,MATLABexpressions{k},MathMLexpressions{k});
end
return


    
    
    
```
