## Supplementary material for "A systems biology analysis of adrenergically stimulated adiponectin exocytosis in white adipocytes": All scripts used in this work: IQMfunctions.m

```
function [names,formulas,arguments] = IQMfunctions(model)
% IQMfunctions: Returns information about the functions in an IQMmodel.
%
% USAGE:
% ======
% [names,formulas,arguments] = IQMfunctions(model)
%
% model: IQMmodel (function can not be used on M-file model)
%
% Output Arguments:
% =================
% names: cell-array with models function names
% formulas: cell-array with formuas for the functions
% arguments: cell-array with arguments for the functions

% <<<COPYRIGHTSTATEMENT - IQM TOOLS LITE>>>

%%%%%%%%%%%%%%%%%%%%%%%%%%%%%%%%%%%%%%%%%%%%%%%
% PROCESS IQMMODEL OR ODE FILE
%%%%%%%%%%%%%%%%%%%%%%%%%%%%%%%%%%%%%%%%%%%%%%%
if strcmp('IQMmodel',class(model)),
    iqm = IQMstruct(model);
    if ~isempty(iqm.functions),
        names = {iqm.functions.name};
        formulas = {iqm.functions.formula};
        arguments = {iqm.functions.arguments};
    else
        names = {};
        formulas = {};
        arguments = {};
    end
else
    error('The function can only be used on IQMmodels, not on M-file ODE models');
end
names = names(:);
formulas = formulas(:);
arguments = arguments(:);
return
```
