## Supplementary material for "A systems biology analysis of adrenergically stimulated adiponectin exocytosis in white adipocytes": All scripts used in this work: IQMfunctionsMATLAB.m

```
function [functionsMATLAB] = IQMfunctionsMATLAB(model)
% IQMfunctionsMATLAB: Returns information about the MATLAB functions in an
% IQMmodel. 
%
% USAGE:
% ======
% [functionsMATLAB] = IQMfunctionsMATLAB(model)
%
% model: IQMmodel (function can not be used on M-file model)
%
% Output Arguments:
% =================
% functionsMATLAB: string containing eventual MATLAB functions defined in
%   the model.

% <<<COPYRIGHTSTATEMENT - IQM TOOLS LITE>>>

%%%%%%%%%%%%%%%%%%%%%%%%%%%%%%%%%%%%%%%%%%%%%%%
% PROCESS IQMMODEL OR ODE FILE
%%%%%%%%%%%%%%%%%%%%%%%%%%%%%%%%%%%%%%%%%%%%%%%
if strcmp('IQMmodel',class(model)),
    iqm = IQMstruct(model);
    functionsMATLAB = iqm.functionsMATLAB;
else
    error('The function can only be used on IQMmodels, not on M-file ODE models');
end
return
```
