## Supplementary material for "A systems biology analysis of adrenergically stimulated adiponectin exocytosis in white adipocytes": All scripts used in this work: IQMinitialconditions.m

```
function [output] = IQMinitialconditions(model,varargin)
% IQMinitialconditions: This functions does the following:
%   1) Returns the initial conditions, stored in a model. For models with
%      numeric initial conditions, these are returned in a vector. For
%      models with non-numeric initial conditions the ICs are returned in a
%      cell-array.
%   2) Same as point 1) but for selected states.
%   3) Changes the initial conditions in an IQMmodel. The non-numeric ICs
%      are unaffected. Only the numeric ones are changed. Despite that, a
%      full numeric IC vector needs to be provided, in which the values
%      corresponding to the non-numeric ICs are arbitrary and have no
%      meaning.
%
% USAGE:
% ======
% [ICs] = IQMinitialconditions(model)
% [ICs] = IQMinitialconditions(model,statenames) 
% [model] = IQMinitialconditions(model,values) 
% [model] = IQMinitialconditions(model,statenames,values) 
%
% model: IQMmodel, ODE, or MEX file model description (in the two last cases
%   the initial conditions can't be set!)
% statenames: cell-array with statenames for which to give back the initial
%   conditions.
% values: vector with initial conditions to update the model with
%
% Output Arguments:
% =================
% ICs: initial conditions (vector or cell-array)
% model: IQMmodel with changed initial conditions

% <<<COPYRIGHTSTATEMENT - IQM TOOLS LITE>>>

%%%%%%%%%%%%%%%%%%%%%%%%%%%%%%%%%%%%%%%%%%%%%%%
% EXTRACT SOME INFO FROM THE MODEL
%%%%%%%%%%%%%%%%%%%%%%%%%%%%%%%%%%%%%%%%%%%%%%%
% get the models ICs and statenames
if isIQMmodel(model),
    ms = struct(model);
    if hasonlynumericICsIQM(model),
        ICs = [ms.states.initialCondition];
    else
        ICs = {ms.states.initialCondition};
    end
    sn = {ms.states.name};
else
    % assume model is an ODE or MEX file
    ICs = feval(model);
    sn = feval(model,'states');
end

%%%%%%%%%%%%%%%%%%%%%%%%%%%%%%%%%%%%%%%%%%%%%%%
% HANDLE VARIABLE INPUT ARGUMENTS
%%%%%%%%%%%%%%%%%%%%%%%%%%%%%%%%%%%%%%%%%%%%%%%
if nargin == 1,
    % simply return the initial conditions for all states
    output = ICs;
elseif nargin == 2,
    % check if values to be changed or ICs for certain states returned
    if isnumeric(varargin{1}),
        % change values
        values = varargin{1};
        % Check if IQMmodel
        if ~isIQMmodel(model),
            error('Initial conditions can only be changed for IQMmodels.');
        end
        ms = struct(model);
        % check length of values vector
        if length(values) ~= length(ms.states),
            error('The length of the values vector does not match the number of states in the model.');
        end
        % Check if numeric ICs
        if hasonlynumericICsIQM(model),
            % model has only numeric ICs => simple update
            cellvalues = mat2cell(values(:)',1,ones(1,length(values)));
            [ms.states.initialCondition] = deal(cellvalues{:});
        else
            % model has non-numeric ICs => update only the numeric values
            for k=1:length(ms.states),
                if isnumeric(ms.states(k).initialCondition),
                    ms.states(k).initialCondition = values(k);
                end
            end
            disp('Please note that only the numeric ICs in the model have been set to the new values.');
        end            

        % Return the IQMmodel with new initial values
        output = IQMmodel(ms);
    elseif ischar(varargin{1}) || iscell(varargin{1}),
        % return specific state values
        statenames = varargin{1};
        if ischar(statenames),
            statenames = {statenames};
        end
        % check if numeric or non-numeric
        if iscell(ICs),
            % non-numeric
            output = {};
            for k=1:length(statenames),
                index = strmatchIQM(statenames{k},sn,'exact');
                if isempty(index),
                    error('State ''%s'' does not exist in the model.\n',statenames{k});
                end
                output{k} = ICs{index};
            end
        else
            % numeric
            output = [];
            for k=1:length(statenames),
                index = strmatchIQM(statenames{k},sn,'exact');
                if isempty(index),
                    error('State ''%s'' does not exist in the model.\n',statenames{k});
                end
                output(k) = ICs(index);
            end
        end
    else
        error('Incorrect second input argument.');
    end
elseif nargin == 3,
    states = varargin{1};
    newvalues = varargin{2};
    if ischar(states),
        states = {states};
    end
    % check if given states exists in model and do the changes
    modelstruct = IQMstruct(model);
    for k0 = 1:length(states),
        state = states{k0};
        newvalue = newvalues(k0);
        stateIndex = strmatchIQM(state,{modelstruct.states.name},'exact');
        if isempty(stateIndex),
            error(sprintf('State ''%s'' does not exist in model.\n',state));
        end
        if length(stateIndex) > 1,
            error(sprintf('State ''%s'' is defined %d times.\n',state,length(stateIndex)));
        end
        % update state with new value
        modelstruct.states(stateIndex).initialCondition = newvalue;
    end
    model = IQMmodel(modelstruct);
    output = model;    
else
    error('Incorrect number of input arguments.');
end
```
