## Supplementary material for "A systems biology analysis of adrenergically stimulated adiponectin exocytosis in white adipocytes": All scripts used in this work: IQMmodelnotes.m

```
function notes = IQMmodelnotes(model)
% IQMmodelnotes: displays the notes in model.
%
% USAGE:
% ======
% [notes] = IQMmodelnotes(model)
%
% model: IQMmodel description of model
%
% Output Arguments:
% =================
% notes: notes, stored in the model

% <<<COPYRIGHTSTATEMENT - IQM TOOLS LITE>>>

ms = struct(model);
notes = ms.notes;
```
