## Supplementary material for "A systems biology analysis of adrenergically stimulated adiponectin exocytosis in white adipocytes": All scripts used in this work: IQMoverwriteICs.m

```
function [newmodel] = IQMoverwriteICs(model,icvector)
% IQMoverwriteICs: This function sets new initial conditions, specified as
% input argument. If the input model contains non-numeric initial
% conditions, these are overwritten (in contrast, the function
% IQMinitialconditions only overwrites the ICs that are defined by numerical
% values).
%
% USAGE:
% ======
% [newmodel] = IQMoverwriteICs(model,icvector)
%
% model: IQMmodel
% icvector: numeric vector, defining the new initial conditions for the model
%
% Output Arguments:
% =================
% newmodel: IQMmodel with new initial conditions

% <<<COPYRIGHTSTATEMENT - IQM TOOLS LITE>>>

%%%%%%%%%%%%%%%%%%%%%%%%%%%%%%%%%%%%%%%%%%%%%%%
% CHECK
%%%%%%%%%%%%%%%%%%%%%%%%%%%%%%%%%%%%%%%%%%%%%%%
if ~isIQMmodel(model),
    error('First input argument needs to be an IQMmodel.');
end
if length(icvector) ~= length(IQMinitialconditions(model)),
    error('Provided initial condition vector does not have the correct length.');
end
    
%%%%%%%%%%%%%%%%%%%%%%%%%%%%%%%%%%%%%%%%%%%%%%%
% UPDATE MODEL WITH ICVECTOR
%%%%%%%%%%%%%%%%%%%%%%%%%%%%%%%%%%%%%%%%%%%%%%%
newmodel = IQMconvertNonNum2NumIC(model);            % convert to numeric
newmodel = IQMinitialconditions(newmodel,icvector);  % then update ics
```
