## Supplementary material for "A systems biology analysis of adrenergically stimulated adiponectin exocytosis in white adipocytes": All scripts used in this work: IQMparameters.m

```
function [varargout] = IQMparameters(model,varargin)
% IQMparameters: Returns information about the parameters in a model, can
% also be used to set a parameter value.
%
% USAGE:
% ======
% [names,values] = IQMparameters(model)
% [values] = IQMparameters(model,parameters)
% [model] = IQMparameters(model,parameters,newvalues)  (not working for ODE
% file model)
%
% model: IQMmodel or m-file ODE description of model
% parameters: Name of the parameter for which the value is to be returned
%   or for which a new value is to be set. Alternatively ''parameters'' can
%   be a cell array, containing several parameter names.
% newvalues: Vector of new parameter values for parameters defined by
%   'parameters'
%
% Output Arguments:
% =================
% names: cell-array with models parameter names
% values: vector with parameter values
% model: if parameter value changed then this is the new model

% <<<COPYRIGHTSTATEMENT - IQM TOOLS LITE>>>

errorMsg = '';

%%%%%%%%%%%%%%%%%%%%%%%%%%%%%%%%%%%%%%%%%%%%%%%
% PROCESS IQMMODEL OR ODE FILE
%%%%%%%%%%%%%%%%%%%%%%%%%%%%%%%%%%%%%%%%%%%%%%%
if strcmp('IQMmodel',class(model)),
    iqm = IQMstruct(model);
    if ~isempty(iqm.parameters),
        names = {iqm.parameters.name};
        values = [iqm.parameters.value];
    else
        names = {};
        values = {};
    end
else
    if nargin > 2,
        error('Parameters can not be changed when model is given as ODE file.');
    end
    names = feval(model,'parameters');
    values = feval(model,'parametervalues');
end
names = names(:);
values = values(:);

% return values if information requested
if nargin == 1,
    varargout{1} = names;
    varargout{2} = values;
elseif nargin == 2,
    parameters = varargin{1};
    if ischar(parameters),
        parameters = {parameters};
    end
    % check if given parameters exists in model and do the changes
    outputvalues = [];
    for k0 = 1:length(parameters),
        parameter = parameters{k0};
        parameterIndex = strmatchIQM(parameter,names,'exact');
        if isempty(parameterIndex),
            errorMsg = sprintf('%sParameter ''%s'' does not exist in model.\n',errorMsg,parameter);
        elseif length(parameterIndex) > 1,
            errorMsg = sprintf('%sParameter ''%s'' is defined %d times.\n',errorMsg,parameter,length(parameterIndex));
        else
            outputvalues = [outputvalues values(parameterIndex)];
        end
    end
    varargout{1} = outputvalues(:);
elseif nargin == 3,
    parameters = varargin{1};
    newvalues = varargin{2};
    if ischar(parameters),
        parameters = {parameters};
    end
    % check if given parameters exists in model and do the changes
    modelstruct = IQMstruct(model);
    for k0 = 1:length(parameters),
        parameter = parameters{k0};
        newvalue = newvalues(k0);
        parameterIndex = strmatchIQM(parameter,{modelstruct.parameters.name},'exact');
        if isempty(parameterIndex),
            errorMsg = sprintf('%sParameter ''%s'' does not exist in model.\n',errorMsg,parameter);
        end
        if length(parameterIndex) > 1,
            errorMsg = sprintf('%sParameter ''%s'' is defined %d times.\n',errorMsg,parameter,length(parameterIndex));
        end
        % update parameter with new value
        modelstruct.parameters(parameterIndex).value = newvalue;
    end
    model = IQMmodel(modelstruct);
    varargout{1} = model;
else
    error('Incorrect number of input arguments.');
end
if ~isempty(errorMsg),
    error(errorMsg);
end
return
```
