## Supplementary material for "A systems biology analysis of adrenergically stimulated adiponectin exocytosis in white adipocytes": All scripts used in this work: IQMreactions.m

```
function [names,formulas,reversibleFlag,fastFlag,varargout] = IQMreactions(model,varargin)
% IQMreactions: Returns information about the reactions in an IQMmodel.
% If a state vector is passed additionally, the corresponding reaction
% rates are returned also.
%
% USAGE:
% ======
% [names,formulas,reversibleFlag,fastFlag] = IQMreactions(model)
% [names,formulas,reversibleFlag,fastFlag,reactionrates] = IQMreactions(model,statevector)
% [names,formulas,reversibleFlag,fastFlag,reactionrates] = IQMreactions(model,time,statevector)
%
% model: IQMmodel (function can not be used on M-file model)
% statevector: vector with corresponding statevalues
% time: time instant of the statevector (only needed for time variant
% systems)
%
% DEFAULT VALUES:
% ===============
% statevector: not needed 
% time: 0  (makes no difference for time invariant systems)
%
% Output Arguments:
% =================
% names: cell-array with models reaction names
% formulas: cell-array with formuas for the kinetic laws of the reactions
% reversibleFlag: vector with same number of entries as reactions. An entry
%   of 0 indicates an irreversible reaction, an entry of 1 indicates a
%   reversible reaction. The ordering is the same as the reaction names.
% fastFlag: vector with same number of entries as reactions. An entry
%   of 1 indicates a fast reaction, an entry of 0 indicates a
%   reaction that is determined by its kinetic rate law.
% reactionrates: the reaction rates at the given state and time

% <<<COPYRIGHTSTATEMENT - IQM TOOLS LITE>>>

%%%%%%%%%%%%%%%%%%%%%%%%%%%%%%%%%%%%%%%%%%%%%%%
% PROCESS IQMMODEL OR ODE FILE
%%%%%%%%%%%%%%%%%%%%%%%%%%%%%%%%%%%%%%%%%%%%%%%
if strcmp('IQMmodel',class(model)),
    iqm = IQMstruct(model);
    if ~isempty(iqm.reactions),
        names = {iqm.reactions.name};
        formulas = {iqm.reactions.formula};
        reversibleFlag = [iqm.reactions.reversible];
        fastFlag = [iqm.reactions.fast];
    else 
        names = {};
        formulas = {};
        reversibleFlag = [];
        fastFlag = [];
    end
else
    error('The function can only be used on IQMmodels, not on M-file ODE models');
end
names = names(:);
formulas = formulas(:);
reversibleFlag = reversibleFlag(:);
fastFlag = fastFlag(:);

% check if the reaction rates need to be determined:
if nargin == 2,    
    statevector = varargin{1};
    time = 0;
    % create data file (using IQMcreateTempODEfile function)
    [ODEfctname, ODEfilefullpath, DATAfctname] = IQMcreateTempODEfile(model,1);
    data = feval(DATAfctname, time, statevector);
    % delete all temporary m files
    IQMdeleteTempODEfile(ODEfilefullpath);
    varargout{1} = data.reactionvalues';
elseif nargin == 3,
    time = varargin{1};
    statevector = varargin{2};
    % create data file (using IQMcreateTempODEfile function)
    [ODEfctname, ODEfilefullpath, DATAfctname] = IQMcreateTempODEfile(model,1);
    data = feval(DATAfctname, time, statevector);
    % delete all temporary m files
    IQMdeleteTempODEfile(ODEfilefullpath);
    varargout{1} = data.reactionvalues';
end
return
```
