## Supplementary material for "A systems biology analysis of adrenergically stimulated adiponectin exocytosis in white adipocytes": All scripts used in this work: IQMstates.m

```
function [names,ODEs,initialConditions] = IQMstates(model)
% IQMstates: Returns information about the states in a model.
%
% USAGE:
% ======
% [names,ODEs,initialConditions] = IQMstates(model)
%
% model: IQMmodel or m-file ODE description of model
%
% Output Arguments:
% =================
% names: cell-array with models state names
% ODEs: cell-array with right hand side formula for the states ODE
%       This output variable is empty if the model is defined by an ODE
%       file, and non-empty in case the model is defined as an IQMmodel
% initialConditions: vector with initial conditions for states

% <<<COPYRIGHTSTATEMENT - IQM TOOLS LITE>>>

%%%%%%%%%%%%%%%%%%%%%%%%%%%%%%%%%%%%%%%%%%%%%%%
% PROCESS IQMMODEL OR ODE FILE
%%%%%%%%%%%%%%%%%%%%%%%%%%%%%%%%%%%%%%%%%%%%%%%
if strcmp('IQMmodel',class(model)),
    iqm = IQMstruct(model);
    names = {iqm.states.name};
    ODEs = {iqm.states.ODE};
    initialConditions = IQMinitialconditions(model);
else
    names = feval(model,'states');
    ODEs = {};
    initialConditions = IQMinitialconditions(model);
end
names = names(:);
ODEs = ODEs(:);
initialConditions = initialConditions(:);
return
```
