## Supplementary material for "A systems biology analysis of adrenergically stimulated adiponectin exocytosis in white adipocytes": All scripts used in this work: IQMvariables.m

```
function [names,formulas,varargout] = IQMvariables(model,varargin)
% IQMvariables: Returns information about the variables in an IQMmodel.
% If a state vector is passed additionally, the corresponding values of the
% variables are returned also.
%
% USAGE:
% ======
% [names,formulas] = IQMvariables(model)
% [names,formulas,variablevalues] = IQMvariables(model,statevector)
% [names,formulas,variablevalues] = IQMvariables(model,time,statevector)
%
% model: IQMmodel (function can not be used on M-file model)
% statevector: vector with corresponding statevalues
% time: time instant of the statevector (only needed for time variant
% systems)
%
% DEFAULT VALUES:
% ===============
% statevector: not needed 
% time: 0  (makes no difference for time invariant systems)
%
% Output Arguments:
% =================
% names: cell-array with models variable names
% formulas: cell-array with formuals for the variables
% variablevalues: the values of the variables in the model for the given state and time

% <<<COPYRIGHTSTATEMENT - IQM TOOLS LITE>>>

%%%%%%%%%%%%%%%%%%%%%%%%%%%%%%%%%%%%%%%%%%%%%%%
% PROCESS IQMMODEL OR ODE FILE
%%%%%%%%%%%%%%%%%%%%%%%%%%%%%%%%%%%%%%%%%%%%%%%
if strcmp('IQMmodel',class(model)),
    iqm = IQMstruct(model);
    if ~isempty(iqm.variables),
        names = {iqm.variables.name};
        formulas = {iqm.variables.formula};
    else
        names = {};
        formulas = {};
    end
else
    if nargin>1,
        error('On ODE models this function can only return variable names and formulas.');
    end
    names = feval(model,'variablenames');
    formulas = feval(model,'variableformulas');
end
names = names(:);
formulas = formulas(:);

% check if the variable values need to be determined:
if nargin == 2,    
    statevector = varargin{1};
    time = 0;
    % create data file (using IQMcreateTempODEfile function)
    [ODEfctname, ODEfilefullpath, DATAfctname] = IQMcreateTempODEfile(model,1);
    data = feval(DATAfctname, time, statevector);
    % delete all temporary m files
    IQMdeleteTempODEfile(ODEfilefullpath);
    varargout{1} = data.variablevalues';
elseif nargin == 3,
    time = varargin{1};
    statevector = varargin{2};
    % create data file (using IQMcreateTempODEfile function)
    [ODEfctname, ODEfilefullpath, DATAfctname] = IQMcreateTempODEfile(model,1);
    data = feval(DATAfctname, time, statevector);
    % delete all temporary m files
    IQMdeleteTempODEfile(ODEfilefullpath);
    varargout{1} = data.variablevalues';
end
return
```
