## Supplementary material for "A systems biology analysis of adrenergically stimulated adiponectin exocytosis in white adipocytes": All scripts used in this work: Contents.m

```
% IQM Tools Lite
% Version 1.2.2 (R2015B) 02.01.2017
% 
% IQMlite
% =======
% 	Contents                                - This help text
% 	SETUP_PATHS_TOOLS_IQMLITE               - Setup script for IQMlite - please customize according to your needs
% 	installIQMlite                          - This script installs the IQMlite package of the IQM Suite of modeling tools
% 
% 	IQMnewscript                            - Creates a new script with a short guide on how to  initialize IQM Tools.
%     
% 	IQMinitializeCompliance                 - Initializes the compliance check when compliance mode is activated in SETUP_PATHS_TOOLS_IQMLITE
%                       
% Model definition and handling
% =============================
% 	IQMmodel                                - Creates a model object 
% 
%   IQMeditBC                               - GUI for editing IQMmodels in a more biochemically oriented format
% 	IQMedit                                 - GUI for editing IQMmodels
%         
% 	IQMalgebraic                            - Returns information about the algebraic equations in a model.
% 	IQMcalcICvector                         - Determines an IC vector for models with  non-numeric initial conditions. 
% 	IQMconvert2MA                           - Takes an IQMmodel and if possible returns the stoichiometric matrix, etc.
% 	IQMconvertNonNum2NumIC                  - Converts non-numeric iniital conditions to numeric initial conditions in the model.
% 	IQMcreateODEfile                        - Creates an m-file that can be simulated by using standard MATLAB integrators (ODE45, ODE23s, etc.)
% 	IQMcreateTEXTBCfile                     - Creates a *.txtbc file with the models text  description in a biochemically oriented format
% 	IQMcreateTEXTBCmodel                    - Creates a new TEXTBC model, saves the corresponding file in the current directory and opens it in the editor, if opening is  desired.
% 	IQMcreateTEXTfile                       - creates a *.txt file with the models text description based on ordinary differential equations
% 	IQMcreateTEXTmodel                      - Creates a new TEXT model, saves the corresponding file in the current directory and opens it in the editor, if opening is  desired.
% 	IQMcreateTempODEfile                    - Creates a temporary ODE file, same as IQMcreateODEfile, but the file is written in the systems temporary  directory
% 	
%   IQMevents                               - Returns information about the events in an IQMmodel.
% 	IQMexportSBML                           - Exports an IQMmodel to an SBML Level 2 Version 1 model using the OutputSBML function 
% 	IQMfunctions                            - Returns information about the functions in an IQMmodel.
% 	IQMfunctionsMATLAB                      - Returns information about the MATLAB functions in an IQMmodel.
% 	IQMinitialconditions                    - Returns or sets  initial conditions in a model
% 	IQMmodelnotes                           - Displays the notes of the model.
% 	IQMoverwriteICs                         - Sets new initial conditions - overwrites non-numerical ones
% 	IQMparameters                           - Returns information about the parameters in a model, can  also be used to set a parameter value.
% 	IQMreactions                            - Returns information about the reactions in an IQMmodel. 
% 	IQMstates                               - Returns information about the states in a model.
% 	IQMvariables                            - Returns information about the variables in an IQMmodel.  
% 	
%   isIQMmodel                              - Checks if provided input argument is an IQMmodel.
% 	
% 	cleanmodelIQM                           - Remove unused reactions, variables and parameters from a model
% 	reactionindexIQM                        - Returns the number of the reaction 'reactionname' in model  'model'. If the reaction does not exist then [] is returned.
% 	stateindexIQM                           - Returns the number of the state 'statename' in model  'model'. If the state does not exist then [] is returned.
% 	variableindexIQM                        - Returns the number of the variable 'variablename' in model  'model'. If the variable does not exist then [] is returned.
% 	hasmoietyconservationsIQM               - Checks if the model contains moiety  conservations.
% 	hasonlynumericICsIQM                    - Checks if the model contains only numeric initial  conditions. The model can be an IQMmodel or an ODE or MEX file model.
% 	isparameterIQM                          - Checks if "name" is a parameter in the provided model. 
% 	isreactionIQM                           - Checks if "name" is a reaction in the provided model. 
% 	isstateIQM                              - Checks if "name" is a state in the provided model.  
% 	isvariableIQM                           - Checks if "name" is a variable in the provided model.  
% 	usealgebraicIQM                         - Checks if an IQMmodel contains algebraic rules.
% 	usedelayIQM                             - Checks if an IQMmodel contains the delayIQM function.
% 	useeventIQM                             - Checks if an IQMmodel contains events.
% 	usefastIQM                              - Checks if an IQMmodel contains "fast" reactions.
% 
% Experiment creation and handling
% ================================
% 	IQMexperiment                           - Creates an experiment object defining experiment settings
% 	
%   IQMcreateEXPfile                        - Creates a *.exp file with the experiments text description
% 	IQMmergemodexp                          - Combines an experiment with a model, and produces a "merged model", as output
% 	
%   isIQMexperiment                         - Checks if provided input argument is an IQMexperiment
% 
% Measurement creation and handling
% =================================
%   IQMmeasurement                          - Creates an object that is intended to contain measurement data - from real experiments or insilico computations
% 	
%   IQMexportCSVmeasurement                 - Exports an IQMmeasurement object to a CSV (comma separated value) file. 
% 	IQMexportXLSmeasurement                 - Exports an IQMmeasurement object to an XLS (excel) file.  
% 	IQMexportXLSmeasurements                - Exports several IQMmeasurement objects to the same XLS (excel) file. 
% 	IQMimportCSVmeasurement                 - Imports experimental measurement data stored in an CSV (comma separated value) file
% 	IQMimportXLSmeasurement                 - Imports experimental measurement data stored in an XLS Excel file 
% 	IQMmeasurementdata                      - This functions allows to extract information from an IQMmeasurement structure
% 	IQMvisualizemeasurement                 - Function allowing to visualize the content of an IQMmeasurement object.  
% 	
%   isIQMmeasurement                        - Checks if provided input argument is an IQMmeasurement
% 
% Dosing creation and handling    
% ============================
% 	IQMdosing                               - Creates an IQMdosing object defining a dosing schedule
% 
%   IQMcreateDOSING                         - Creates a dosing scheme as desired based on inputs
% 	IQMcreateDOSfile                        - Creates a *.dos file with the dosing text description
% 	IQMdoseupdatevalue                      - Allows to update the dosing amount for a given input, defined in an IQMdosing object.
% 
%   isIQMdosing                             - Checks if provided input argument is an IQMdosing
% 
% 	IQMmergemoddos                          - Based on a model and dosing object, a new IQMmodel is generated, that implements the defined dosings. 
%   mergemoddosIQM                          - Similar to IQMmergemoddos but only preparing the model for taking dosings of this type, not implementing them in the model.
% 
% Analysis and simulation
% =======================
% simulation
% 	IQMinsilicoexp                          - IQMinsilicoexp runs an in-silico experiment.
% 	IQMsimdosing                            - Simulates the application of a dosing schedule to a model that has been prepared for it by mergemoddosIQM
% 	IQMsimulate                             - Simulation of models and simulation of models and dosing schemes.
% 	IQMstochsim                             - Stochastic simulation of IQMmodels that only contain mass action kinetics. 
% 
%   IQMjacobian                             - Determines the Jacobian of a given IQMmodel
% 	IQMmakeirreversible                     - Makes reactions irreversible
% 	IQMmoietyconservations                  - Determines the moitey conservations
% 	IQMreactantstoichiometry                - Determines the stoichiometric coefficients of only the reactants
% 	IQMreducemodel                          - Reduces an IQMmodel by identifying algebraic relations between time dependent variable
% 	IQMsteadystate                          - Determines a steady-state of an IQMmodel 
% 	IQMstoichiometry                        - Determines the stoichiometric matrix for the given model
% 
% behaviorlocalization
% 	IQMlocbehavcomp                         - Determines the importance of components in the given biochemical system in the creation of an observed complex behavior
% 	IQMlocbehavinteract                     - Determines the importance of direct interactions between components (states) in the given biochemical system in the creation of complex behavior
% 	IQMlocbehavinteract2                    - Similar to IQMlocbehavinteract (see help text)
% 
% globalparametersensitivity
% 	IQMsensglobalfast                       - Global sensitivity analysis method: Extended FAST method
% 	IQMsensglobalprcc                       - Global sensitivity analysis method: PRCC method
% 	IQMsensglobalsobol                      - Global sensitivity analysis method: SOBOL's method
% 	IQMsensglobalwals                       - Global sensitivity analysis method: WALS (weighted average of local  sensitivities)
% 
% localparametersensitivity
% 	IQMmca                                  - Metabolic control analysis  
% 	IQMsensamplitude                        - Function evaluating local first order amplitude sensitivities.
% 	IQMsensdataosc                          - Function allowing to generate data that subsequently can be used for different kinds of parametric sensitivity analyses. 
% 	IQMsensdataoscevents                    - Function allowing to generate data that subsequently can be used for different kinds of parametric sensitivity analyses. 
% 	IQMsensdatastat                         - Function allowing to generate steady-state data that subsequently can be used for different kinds of parametric sensitivity analyses.
% 	IQMsensperiod                           - Function evaluating local first order period sensitivities.
% 	IQMsensstat                             - Function evaluating local first order parameter sensitivities of the steady-state values of states and reaction rates.
% 
% modelreduction
% 	IQMprepredreac                          - Prepares a model for subsequent reduction of reaction rate expressions. 
% 	IQMredallreac                           - Wrapper for the IQMredreac function allowing to easily go through a whole model and reduce all reactions that are possible to reduce.
% 	IQMredreac                              - Reduction of single reaction expressions with the goal of reducing complexity and number of parameters. 
% 
% Tools
% =====
% dataset
%   Datasets used in IQM Tools use the MATLAB "table" type.
%     
% 	IQMdataset2wide                         - Expands a dataset from a row based format to a column based format. 
% 	IQMexportCSVdataset                     - Exports a MATLAB table as standard comma separated (CSV) datafile. 
% 	IQMgetdatasetHeader                     - Returns the column names of the dataset in a cell-array.  
% 	IQMloadCSVdataset                       - Loads a standard CSV datafile with header as a TABLE object into MATLAB. 
% 	IQMloadNONCSVdataset                    - Loads a NON CSV datafile with header as a dataset into MATLAB. 
% 	IQMresampleDataset                      - Resamples a dataset. The structure and the number of subjects is preserved. Useful for bootstrapping. 
% 	IQMsas7bdat2csv                         - Converts a sas7bdat file into a CSV file (requires SAS installation)
% 	ixdataIQM                               - Is a wrapper that supports easier subsetting of data.  
% 	subsetIQM                               - Allows to subset a dataset or table. Both numeric and cell-array with string columns are handled.
% 
% optimization
% 	SSmIQM                                  - Global optimization algorithm for MINLP's based on Scatter Search.
% 	fSSmIQM                                 - Global optimization algorithm for MINLP's based on Scatter Search ("fast" version).
% 	isresIQM                                - Stochastic Ranking for Constrained Evolutionary Minimization.
% 	pswarmIQM                               - Particle swarm pattern search algorithm for global optimization.
% 	simannealingIQM                         - Minimization by simulated annealing
% 	simplexIQM                              - Downhill Simplex Method in Multidimensions
% 
% plots
% 	IQMbarplotErrors                        - Grouped bar plot with error bars, customizable.
% 	IQMplotCovarianceCat                    - Plots the covariance relationship between a list of continuous variables and a list of categorical variables 
% 	IQMplotHistogram                        - Plots histograms 
% 	IQMplotQQ                               - QQ plot for provided input data.
% 	IQMplotXY                               - Plots pairwise Ydata vs. Xdata. 
% 	IQMplotfacetgrid                        - Plots a matrix version of a trellis plot.
% 	IQMplotfill                             - Will fill a region with a color between upper and lower bounds provided 
% 	IQMplotpairwiseCorr                     - Plots the pairwise correlation between variables passed in columns of a matrix or passed as a dataset.
% 	IQMplottrellis                          - Plots a Trellis plot. Plenty of options available.
% 	IQMplot                                 - IQMplot - plots given data.
%   IQMplot2                                - IQMplot2 - plots bar diagrams for given data.
%   IQMplotCatCat                           - This function plots a "bubble" plot for categorical data to assess correlations
%   IQMplotKM                               - Kaplan-Meier plot
% 
% functions
%   These functions can be used on IQMmodels but on command line as well. When used in models then these are also working when converting the IQMmodel to C-code
%     
% 	andIQM                                  - Is used instead of the MATLAB "and" function, allowing more than two input arguments
% 	delayIQM                                - Realizes a time delay of "tau" time units (hande with care)
% 	indexmaxIQM                             - Searches for the maximum input argument and returns the index of the max.
% 	maxIQM                                  - Is used instead of the MATLAB "max" function, allowing more than two scalar input arguments
% 	minIQM                                  - Is used instead of the MATLAB "min" function, allowing more than two scalar input arguments
% 	multiplyIQM                             - Implementing the multiply MathML function
% 	orIQM                                   - Is used instead of the MATLAB "or" function, allowing more than two input arguments
% 	piecewiseIQM                            - Implements support for the SBML / MATHML piecewise operator.
% 	piecewiseSmoothIQM                      - Implements a smoothing function between two values y0 and y1
% 	piecewiseT0IQM                          - Is similar to the piecewiseIQM function (see help text)
% 	xorIQM                                  - Is used instead of the MATLAB "or" function, allowing more than two input arguments
% 
% interpolation
% 	interp0IQM                              - Zero order interpolation function (lookup table)
% 	interp1IQM                              - Linear interpolation function (lookup table)  
% 	interpcsIQM                             - Cubic spline interpolation function (lookup table)  
% 	interpcseIQM                            - Cubic spline interpolation with endpoints
% 	interpcseSlopeIQM                       - Cubic spline interpolation with endpoints, returning the derivative at the considered point.
% 	interpcsexIQM                           - Cubic spline interpolation with endpoints 
% 	interpcsexSlopeIQM                      - Cubic spline interpolation with endpoints 
% 
% signal
% 	centeredfftIQM                          - Uses the fft function of MATLAB to determine a two sided spectrum 
% 	positivefftIQM                          - Uses the fft function of MATLAB to determine a one sided spectrum 
% 	resampleIQM                             - Resamples time series x1, which is sampled at the time instances t1 to time series x2 using a sampling defined by t2.
% 	xcorrIQM                                - Compute correlation R_xy of X and Y for various lags k:
% 
% smoothing
% 	binnedmeanIQM                           - Calcuate binned means
% 	binnedmedianIQM                         - Calcuate binned medians
% 	binnedquantilesIQM                      - Calcuate binned quantiles
% 	movingAverageIQM                        - Compute moving averages over a range
% 	movingMedianIQM                         - Compute moving medians over a range
% 	movingQuantileIQM                       - Compute moving quantiles over a range
%   smoothIQM                               - loess, lowess smoother
%
% solvers
% 	fsolveIQM                               - Attempts to solve equations of the form FUN(X)=0 with Newton iteration
% 
% statistic
% 	boxplotIQM                              - Plot a boxplot for given data
% 	centerIQM                               - Center by subtracting means
% 	clusteringIQM                           - Performs UPGMA on distance matrix and produces a denddrogram plot.
% 	kurtosisIQM                             - Returns the kurtosis over the first non-singleton dimension. 
% 	mvnrndIQM                               - Draw n random d-dimensional vectors from a multivariate Gaussian distribution with mean mu and covariance matrix Sigma.
% 	nanmeanIQM                              - Determines the mean of x along the dimension dim by treating NaNs as missing values.
% 	nanmedianIQM                            - Determines the median of x along the dimension dim by treating NaNs as  missing values.
% 	nanstdIQM                               - Determines the std of x along the dimension dim by treating NaNs as missing values.
% 	pdistIQM                                - Determines the distance matrix for a set of points whose coordinates are given as row-vectors in the data matrix.
% 	prctileIQM                              - prctileIQM calculates the percentiles of histograms and sample arrays.
% 	princompIQM                             - Compute principal components of X
% 	qqplotIQM                               - Perform a QQ-plot (quantile plot). Only for comparison to standard normal  distribution.
% 	quantileIQM                             - quantileIQM calculates the quantiles of histograms and sample arrays.
% 	regressIQM                              - Multiple Linear Regression using Least Squares Fit of y on X  with the model y = X * beta + e.
% 	sumsqIQM                                - Sum of squares of elements along dimension dim.
% 	swtestIQM                               - Shapiro-Wilk parametric hypothesis test of composite normality.
% 	zscoreIQM                               - Compute the z-score of each element of X relative to the data  in the columns of X.
% 
% distributions
% 	betacdfIQM                              - Cumulative density function of the Beta distribution
% 	betainvIQM                              - Quantile function of the Beta distribution
% 	betapdfIQM                              - Probability density function of the Beta distribution
% 	chi2cdfIQM                              - Cumulative density function of the chi-square distribution
% 	chi2invIQM                              - Quantile function of the chi-square distribution
% 	chi2pdfIQM                              - Probability density function of the chi-square distribution
% 	fcdfIQM                                 - Cumulative density function of the f distribution.
% 	finvIQM                                 - Quantile function of the f distribution
% 	fpdfIQM                                 - Probability density function of the f distribution
% 	gamcdfIQM                               - Cumulative density function of the Gamma distribution
% 	gaminvIQM                               - Quantile function of the Gamma distribution
% 	gampdfIQM                               - Probability density function of the Gamma distribution
% 	normcdfIQM                              - Cumulative densitiy function of the normal distribution
% 	norminvIQM                              - Quantile function of the normal distribution
% 	normpdfIQM                              - Probability density function of the normal distribution
% 	stdnormalcdfIQM                         - Cumulative density function of the standard normal distribution
% 	stdnormalinvIQM                         - Quantile function of the standard normal distribution
% 	stdnormalpdfIQM                         - Probability density function of the standard normal distribution
% 	tcdfIQM                                 - Cumulative density function of the t distribution
% 	tinvIQM                                 - Quantile function of the t distribution
% 	tpdfIQM                                 - Probability density function of the t distribution
% 
% symbolic
%   Function that do symbolic computations require the symbolic toolbox to be present
%     
% 	IQMsymjacobian                          - Determines a symbolic Jacobian and the derivative of the ODE right hand side with respect to given parameters. 
%     
% kineticlaws
%   These kinetic laws can be used in IQMmodels, allowing for more informative coding and less complex formulas.
%     
% 	kin_allosteric_inihib_empirical_rev     - Allosteric inhibition (reversible) kinetics
% 	kin_allosteric_inihib_mwc_irr           - Allosteric inhibition (irreversible) kinetics
% 	kin_catalytic_activation_irr            - Catalytic activation (irreversible) kinetics
% 	kin_catalytic_activation_rev            - Catalytic activation (reversible) kinetics
% 	kin_comp_inihib_irr                     - Competitive inhibition (irreversible) kinetics
% 	kin_comp_inihib_rev                     - Competitive inhibition (reversible) kinetics
% 	kin_constantflux                        - Competitive inhibition (reversible) kinetics
% 	kin_degradation                         - Linear degradation kinetics
% 	kin_hill_1_modifier_rev                 - Reversible Hill type kinetics with one modifier
% 	kin_hill_2_modifiers_rev                - Reversible Hill type kinetics with one modifier
% 	kin_hill_cooperativity_irr              - Hill type (irreversible) kinetics
% 	kin_hill_rev                            - Hill type (reversible) kinetics
% 	kin_hyperbolic_modifier_irr             - Hyperbolic modifier (irreversible) kinetics
% 	kin_hyperbolic_modifier_rev             - Hyperbolic modifier (reversible) kinetics
% 	kin_iso_uni_uni_rev                     - enzyme isomerization product inhibition
% 	kin_mass_action_irr                     - Mass action (irreversible) kinetics
% 	kin_mass_action_rev                     - Mass action (reversible) kinetics
% 	kin_michaelis_menten_irr                - Michaelis Menten (irreversible) kinetics
% 	kin_michaelis_menten_rev                - Michaelis Menten (reversible) kinetics
% 	kin_mixed_activation_irr                - Mixed activation irreversible
% 	kin_mixed_activation_rev                - Mixed activation reversible
% 	kin_mixed_inihib_irr                    - Mixed inhibition (irreversible) kinetics
% 	kin_mixed_inihib_rev                    - Mixed inhibition (reversible) kinetics
% 	kin_noncomp_inihib_irr                  - Noncompetitive inhibition (irreversible) kinetics
% 	kin_noncomp_inihib_rev                  - Noncompetitive inhibition (reversible) kinetics
% 	kin_ordered_bi_bi_rev                   - Ordered bi-bi reversible
% 	kin_ordered_bi_uni_rev                  - Ordered bi-uni reversible
% 	kin_ordered_uni_bi_rev                  - Ordered uni-bi reversible
% 	kin_ping_pong_bi_bi_rev                 - Ping pong bi bi kinetics (reversible)
% 	kin_specific_activation_irr             - Specific activation (irreversible) kinetics
% 	kin_specific_activation_rev             - Specific activation (reversible) kinetics
% 	kin_substrate_activation_irr            - Substrate activation (irreversible) kinetics
% 	kin_substrate_inihib_irr                - Substrate inhibition (irreversible) kinetics
% 	kin_substrate_inihib_rev                - Substrate inhibition (reversible) kinetics
% 	kin_uncomp_inihib_irr                   - Uncompetitive inhibition (irreversible) kinetics
% 	kin_uncomp_inihib_rev                   - Uncompetitive inhibition (reversible) kinetics
% 	kin_uni_uni_rev                         - Uni uni reversible kinetics
% 
% survival
%   IQMcoxExt 								- Parameter estimation for the extended Cox model
%   IQMcoxPH								- Parameter estimation for the Cox proportional hazards model
%   IQMcoxStratPH 							- Parameter estimation for the Stratified Cox proportional hazards model
%   IQMhr 									- Hazard ratio estimate for Cox models
%   IQMkm 									- Kaplan-Meier curves estimation and plot (see also IQMplotKM)
%   IQMphTest 								- Schoenfeld residuals test for the proportional hazards assumption
%   IQMxext 								- Extension to time varying covariates (input to IQMcoxExt)
%    
% Auxiliary
% =========
% 	getDefaultIntegratorOptionsIQM          - Inside this function/file default values for integrator options are set. This function also retrieves them
% 	lookforIQML                             - Allows to search IQM Lite folders recursively for a given text
% 	setseedIQM                              - Set the default stream to defaultSeed
% 	usernameIQM                             - Returns the username of the current user
%   compareFilesEqualIQM 					- Compare two text files for equality
%
% output
% 	IQMconvert2pdf                          - Converts a PS file to a PDF file.  
% 	IQMgetcolors                            - Generate colors for plotting and for black and white plotting also line types.
% 	IQMprintFigure                          - Print a MATLAB figure to png, ps, jpg, or pdf.
% 	IQMstartNewPrintFigure                  - Function starts a new file in which figures are to be printed.  
% 	IQMwriteText2File                       - Allows to write a given string (text) to a file (filename) 
% 
% parallel
%   These functions require the presence of the parallel toolbox in MATLAB
%     
% 	startParallelIQM                        - Requests parallel nodes from the MATLAB parallel toolbox.
% 	stopParallelIQM                         - Closes the connection to parallel nodes via the parallel    toolbox in MATLAB

% <<<COPYRIGHTSTATEMENT - IQM TOOLS LITE>>>
```
