## Supplementary material for "A systems biology analysis of adrenergically stimulated adiponectin exocytosis in white adipocytes": All scripts used in this work: installation.txt

Installation of IQM Tools Lite
==============================

When you are reading this document, you already have successfully obtained IQM Tools Lite from somewhere.
Please perform the following steps to install it:

- Move the IQMlite folder to a location on your computer where you would like to store it
- Start MATLAB 
- Type "mex -setup" to check if you have a C compiler installed for MATLAB. If not, please read the information from this command and install a C compiler.
- Change into the IQMlite folder
- Open the file "SETUP_PATHS_TOOLS_IQMLITE.m" in the editor, read its documentation and modify setup accordingly (if needed). Then save and close the file
- Run "installIQMlite.m" script
- If you have the parallel toolbox for MATLAB, please do the following setting
    - Home->parallel->parallel preferences deselect the checkbox "Automatically create a parallel pool" 

Note: The installation of IQM Tools Lite does not save the MATLAB path. This means that everytime you start MATLAB you will need to 
      execute the "installIQMlite" script. This procedure has been chosen on purpose, for compliance and reproducibility reasons.
