## Supplementary material for "A systems biology analysis of adrenergically stimulated adiponectin exocytosis in white adipocytes": All scripts used in this work: installIQMlite.m

```
function [] = installIQMlite()
% installIQMlite
% This script installs the LITE version of the IQM Suite of modeling tools.
%
%       installIQMlite
%
% The function will check for a previous version of IQM Lite installed and
% in this case it will remove this previous version from the path and
% install the desired version.

% <<<COPYRIGHTSTATEMENT - IQM TOOLS LITE>>>

% Check correct starting folder
currentDir      = pwd;
installIQMliteDir    = fileparts(which('installIQMlite.m'));
if ~strcmp(currentDir,installIQMliteDir),
    error('Run the ''installIQMlite'' script from the folder where it is located.');
end

% Check that correct local path (network paths are not allowed)
if strcmp(currentDir(1:2),'\\'),
    error(sprintf('The installation can not be run from a network path (\\\\...).\nPlease run the installation from a local path.'));
end

% Check if IQM Lite already installed
IGMLiteVer = ver('IQMlite');
if length(IGMLiteVer) >= 1,
    % Reset the path
    restoredefaultpath();
end

% Add IQM Lite to the path 
addpath(genpath(pwd));
addpath(tempdirIQM)

% Compile and install the needed packages 
% Compilation is done for Unix AND Windows systems
PATH_IQMLITE = pwd();
try
    % interpcseIQM
    cd(fileparts(which('interpcseIQM.m')));
    mex interpcseIQM.c
    mex interpcseSlopeIQM.c
catch, end
cd(PATH_IQMLITE)
try
    % isrsort
    cd(fileparts(which('isrsort.c')));
    mex isrsort.c
catch, end
cd(PATH_IQMLITE)

% Install SBML tools if path defined
if isunix,
    % Get path definitions
    SETUP_PATHS_TOOLS_IQMLITE
    if ~isempty(PATH_SYSTEM_SBML_UNIX),
        try
            oldpath = pwd();
            cd(PATH_SYSTEM_SBML_UNIX);           
            addpath(genpath(pwd));
            cd(oldpath);
        catch
            disp('Problems detected with path to SBML tools. Please have a look at the file "SETUP_PATHS_TOOLS_IQMLITE.m".');
        end
    end
end

% Message
disp(' ');
disp('IQM Tools Lite');
disp('	- Developer: IntiQuan GmbH');
disp('	- Installation completed');
disp(' ');
disp(' ');

% Output license information, etc.
disp('License IQM Tools Lite:');
disp(' ');
disp(sprintf('This program is Free Open Source Software: you can redistribute it and/or modify '));
disp(sprintf('it under the terms of the GNU General Public License as published by '));
disp(sprintf('the Free Software Foundation, either version 3 of the License, or '));
disp(sprintf('(at your option) any later version. '));
disp(sprintf(' '));
disp(sprintf('This program is distributed in the hope that it will be useful, '));
disp(sprintf('but WITHOUT ANY WARRANTY; without even the implied warranty of '));
disp(sprintf('MERCHANTABILITY or FITNESS FOR A PARTICULAR PURPOSE. See the '));
disp(sprintf('GNU General Public License for more details. '));
disp(sprintf(' '));
disp(sprintf('You should have received a copy of the GNU General Public License '));
disp(sprintf('along with this program. If not, see <http://www.gnu.org/licenses/>.'));
disp(sprintf(' '));
disp(sprintf(' '));
disp(sprintf(' '));


try
    % Check version number (>=R2013B required)
    if verLessThan('matlab','8.2.0'),
        warning('The dataset import/export functions in IQM Tools Lite require >=MATLAB R2013B.');
    end
end
```
