## Supplementary material for "A systems biology analysis of adrenergically stimulated adiponectin exocytosis in white adipocytes": All scripts used in this work: release.txt

Release Notes IQM Tools Lite
============================

Version 1.2.2 (02.01.2017)
--------------------------
- Minor big fixes
- Improved graphical output
- Survival analysis functions

Version 1.2.1 (30.04.2016)
------------------------
- Minor big fixes

Version 1.2 (13.04.2016)
------------------------
- Allow non-numeric lag time definitions in IQMdosing schemes
- Ensured compatibility with MONOLIX 4.3.3 
- Ensured better compatibility with Mac computers
- Added "bubble plots" function to allow for correlation assesment of categorical variables
- Added loess/lowess/etc. smoother

Version 1.1 (10.02.2016)
------------------------
- Minor bug fixes
- Update of IQMdosing to fix automatically lenght of the parameter field
- Added functionality in dosing scenarios to allow estimation of 0th order absorption time

Version 1.0 (09.11.2015)
------------------------
- First full release of IQM Lite - the follow-up of SBPOP and SBTOOLBOX2
- Many improvements with respect to SBPOP (stability, usability, documentation, functionality)
- Where SBPOP used the "dataset" type, IQM Tools uses the "table" type
- IQM Tools does not require the presence of the stats toolbox 
- Parallel computation is used in several functions if the parallel toolbox is available
- Recommended use of MATLAB >= R2015b (>=R2013B is required)
- Removed support of SBML Level 1
- Added support of SBML Level 2, Version 2,3, and 4
- Implemented "compliance mode" for time and information stamps on figures, tables, text
