## Supplementary material for "A systems biology analysis of adrenergically stimulated adiponectin exocytosis in white adipocytes": All scripts used in this work: SETUP_PATHS_TOOLS_IQMLITE.m

```
% In this file you need to provide the names and potentially the paths to
% the executables for tools, such as SAS, libSBML, etc.
% If executables in the system path, then only the names of the executables 
% are needed.
%
% It is possible to define paths for Windows and Unix independently,
% allowing to use the package on different systems without the need to
% re-edit the paths. If a tool is not available on a system, then just
% leave the path empty.

% SAS
PATH_SYSTEM_SAS_WINDOWS             = '';
PATH_SYSTEM_SAS_UNIX                = '';

% SBML import and export
% In this path the functions TranslateSBML and OutputSBML need to be located.
% On windows you do not need to define anything, since it is part of IQM Tools Lite.
% On other operating systems you need to install it yourself and then provide here the path.
PATH_SYSTEM_SBML_WINDOWS             = ''; % Leave empty
PATH_SYSTEM_SBML_UNIX                = ''; % Need to define

% Set compliance mode
% In the "compliance mode" each figure and text that is exported using the
% functions "IQMprintFigure" and "IQMwriteText2File" will be annotated with
% a log file that specifies the username, the location of the created file,
% the scripts and functions used to generate this file, its size and the
% date of creation.
% If this mode is enabled, then a global variable
% "Current_ROOT_file_IQMtools_compliance" has to be set in each root script for
% the analysis - this is needed since when working interactively, the root
% script name is unknown. The information gathering function will check
% that this name is defined and if not in the list of functions that will
% lead to the output generation it will add it. If undefined, there will be
% an error, explaining that this variable needs to be set.
COMPLIANCE_OUTPUT_MODE               = 0;  % 1=enabled, 0=off
```
