## Supplementary material for "A systems biology analysis of adrenergically stimulated adiponectin exocytosis in white adipocytes": All scripts used in this work: IQMjacobian.m

```
function [jacobian,statenames] = IQMjacobian(varargin)
% IQMjacobian: determines the Jacobian of a given IQMmodel or an ODE file 
% description. This function should only be used for time-invariant systems.
% If used for time-variant systems an error will occur.
%
% USAGE:
% ======
% [jacobian] = IQMjacobian(model,state)         
% [jacobian] = IQMjacobian(model,state,delta)         
% [jacobian,statenames] = IQMjacobian(model,state)         
% [jacobian,statenames] = IQMjacobian(model,state,delta)         
%
% model: IQMmodel or ODE file model description
% state: state at which to determine the Jacobian
% delta: stepsize used for numerical differentiation. in case of nonzero
%        states the applied change is relative, in case of a zero value the
%        applied change is absolute.
%
% DEFAULT VALUES:
% ===============
% delta: 1e-4
%
% Output Arguments:
% =================
% jacobian: Jacobian of the model at the given state
% statenames: cell-array with the names of the states of the model

% <<<COPYRIGHTSTATEMENT - IQM TOOLS LITE>>>

%%%%%%%%%%%%%%%%%%%%%%%%%%%%%%%%%%%%%%%%%%%%%%%
% GLOBAL VARIABLE
%%%%%%%%%%%%%%%%%%%%%%%%%%%%%%%%%%%%%%%%%%%%%%%
global ODEfctname

%%%%%%%%%%%%%%%%%%%%%%%%%%%%%%%%%%%%%%%%%%%%%%%
% PARAMETERS
%%%%%%%%%%%%%%%%%%%%%%%%%%%%%%%%%%%%%%%%%%%%%%%
% Default stepsize for numerical differentiation
DEFAULT_DELTA = 1e-4;

%%%%%%%%%%%%%%%%%%%%%%%%%%%%%%%%%%%%%%%%%%%%%%%
% CHECK IF IQMMODEL OR FILENAME
%%%%%%%%%%%%%%%%%%%%%%%%%%%%%%%%%%%%%%%%%%%%%%%
if isIQMmodel(varargin{1}),
    % IQMmodel
    iqm = varargin{1};
    % check delays and events
    if usedelayIQM(iqm),
        error('The model contains delays. These can not be handled by this function.');
    end
    % Create temporary ODE file
    [ODEfctname, ODEfilefullpath] = IQMcreateTempODEfile(iqm);    
else
    % ODEfctname of ODE file
    ODEfctname = varargin{1};
    % Check if file exists
    if exist(strcat(ODEfctname,'.m')) ~= 2,
        error('ODE file could not be found.');
    end
end


%%%%%%%%%%%%%%%%%%%%%%%%%%%%%%%%%%%%%%%%%%%%%%%
% PROCESS VARIABLE INPUT ARGUMENTS
%%%%%%%%%%%%%%%%%%%%%%%%%%%%%%%%%%%%%%%%%%%%%%%
if nargin == 2,
    state = varargin{2};
    delta = DEFAULT_DELTA;
elseif nargin == 3,
    state = varargin{2};
    delta = varargin{3};
else
    error('Wrong number of input arguments');
end
% Check if correct number of elements in "state".
teststate = feval(ODEfctname);
if length(state) ~= length(teststate),
    error('Number of elements in provided state-vector does not match the number of states in the model.');
end

%%%%%%%%%%%%%%%%%%%%%%%%%%%%%%%%%%%%%%%%%%%%
% GET JACOBIAN AT GIVEN STATE
%%%%%%%%%%%%%%%%%%%%%%%%%%%%%%%%%%%%%%%%%%%%
jacobian = calculateJacobian(state,delta);

%%%%%%%%%%%%%%%%%%%%%%%%%%%%%%%%%%%%%%%%%%%%
% GET THE STATE NAMES
%%%%%%%%%%%%%%%%%%%%%%%%%%%%%%%%%%%%%%%%%%%%
statenames = feval(ODEfctname,'states');

%%%%%%%%%%%%%%%%%%%%%%%%%%%%%%%%%%%%%%%%%%%%%%%
% DELETE FILE IF IQMMODEL
%%%%%%%%%%%%%%%%%%%%%%%%%%%%%%%%%%%%%%%%%%%%%%%
if strcmp('IQMmodel',class(varargin{1})),
    IQMdeleteTempODEfile(ODEfilefullpath);
end
return

%%%%%%%%%%%%%%%%%%%%%%%%%%%%%%%%%%%%%%%%%%%%%%%%%%%%%%%%%%%%%%%%%%%%%%%%%%
% Calculate Jacobian 
%%%%%%%%%%%%%%%%%%%%%%%%%%%%%%%%%%%%%%%%%%%%%%%%%%%%%%%%%%%%%%%%%%%%%%%%%%
function [jacobian] = calculateJacobian(state,delta)
n = length(state);          % size of system
jacobian = zeros(n,n);      % initialize jacobian variable
% determine the Jacobian by numerical differentiation
for k = 1:n,                
    statedn = state;
    stateup = state; 
    if stateup(k) == 0,
        stateup(k) = stateup(k) + delta;
        jacobian(:,k) = (model(stateup)'-model(statedn)')/delta;
    else
        stateup(k) = stateup(k)*(1+delta);
        jacobian(:,k) = (model(stateup)'-model(statedn)')/(delta*stateup(k));
    end
end
return 

%%%%%%%%%%%%%%%%%%%%%%%%%%%%%%%%%%%%%%%%%%%%%%%%%%%%%%%%%%%%%%%%%%%%%%%%%%
% MODEL
%%%%%%%%%%%%%%%%%%%%%%%%%%%%%%%%%%%%%%%%%%%%%%%%%%%%%%%%%%%%%%%%%%%%%%%%%%
function [xdot] = model(x)
    global ODEfctname
    xdot = feval(ODEfctname,0,x);
return
```
