## Supplementary material for "A systems biology analysis of adrenergically stimulated adiponectin exocytosis in white adipocytes": All scripts used in this work: IQMmakeirreversible.m

```
function [output] = IQMmakeirreversible(model)
% IQMmakeirreversible
% For certain analysis methods it is useful that all reaction rate
% expressions in a model are irreversible. This function takes an IQMmodel
% as input and replaces all reversible reactions by irreversible ones. This
% is done by splitting up reversible reaction rates into one forward and one
% backward rate. For this to be possible it is IMPORTANT that the reaction
% rate expressions that are marked as reversible in the model are defined
% in the following format:
%
%     ReactionRate = RateForward - RateReverse
%
% where "ReactionRate" can be any reaction rate name, and the terms
% "RateForward" and "RateReverse" can be any mathematical expressions,
% involving parameters, states, functions, and/or variables.
% However, it is IMPORTANT that there is only one minus sign in the top
% level of the formula. Minus signs in eventual parentheses are not taken
% into account for the parsing and are therefor, of course, allowed to be
% present.
%
% Examples: R1 = Rf1 - Rr1
%           R2 = k1*A - k2*B
%           R3 = (k1-k2*A) - (k3+k4*B)
%
% This function requires that each reaction only appears once in each ODE.
% Example:    d/dt(A) = -R1 + R2        is OK
%             d/dt(B) = -R1 + R2 + R1   is not OK 
% 
% Replace last ODE by:    d/dt(B) = R2   or do the irreversible making manually.
%
% USAGE:
% ======
% [output] = IQMmakeirreversible(model)
%
% model: IQMmodel for which to convert reversible reactions to irreversible ones.
%
% Output Arguments:
% =================
% output: converted model

% <<<COPYRIGHTSTATEMENT - IQM TOOLS LITE>>>

%%%%%%%%%%%%%%%%%%%%%%%%%%%%%%%%%%%%%%%%%%%%%%%
% CHECK IF IQMmodel
%%%%%%%%%%%%%%%%%%%%%%%%%%%%%%%%%%%%%%%%%%%%%%%
if ~strcmp('IQMmodel',class(model)),
    error('Function only defined for IQMmodels.');
end

%%%%%%%%%%%%%%%%%%%%%%%%%%%%%%%%%%%%%%%%%%%%%%%
% SAVE ORIGINAL MODEL FOR LATER USE
%%%%%%%%%%%%%%%%%%%%%%%%%%%%%%%%%%%%%%%%%%%%%%%
originalmodel = model;

%%%%%%%%%%%%%%%%%%%%%%%%%%%%%%%%%%%%%%%%%%%%%%%
% CHECK THAT ALL ODE EXPRESSIONS ARE DEFINED VIA REACTION TERMS
%%%%%%%%%%%%%%%%%%%%%%%%%%%%%%%%%%%%%%%%%%%%%%%
[Ntest,componentsTest] = IQMstoichiometry(model,1);  % set rawFlag to 1 
componentsModel = IQMstates(model);
if length(componentsTest) ~= length(componentsModel),
    error('Not all ODEs seem to be constructed by reaction terms. Therefor the full stoichiometric information is not possible to determine.');
end

%%%%%%%%%%%%%%%%%%%%%%%%%%%%%%%%%%%%%%%%%%%%%%%
% GET NAMES AND DATA OF REVERSIBLE REACTIONS
%%%%%%%%%%%%%%%%%%%%%%%%%%%%%%%%%%%%%%%%%%%%%%%
[names,formulas,reversibleFlag,fastFlag] = IQMreactions(model);
reversibleReactionsAll = reversibleFlag;
% check if reversible reactions are present
if sum(reversibleFlag) == 0,
    output = model;
    disp('The model does not contain any reversible reactions.');
    return
end
% get indices of reversible reactions
reversibleIndices = find(reversibleFlag ~= 0);
reactionsStore = [];
for k = 1:length(reversibleIndices),
    reactionsStore(k).name = names{reversibleIndices(k)};
    reactionsStore(k).formula = formulas{reversibleIndices(k)};
    reactionsStore(k).fastFlag = fastFlag(reversibleIndices(k));
end

%%%%%%%%%%%%%%%%%%%%%%%%%%%%%%%%%%%%%%%%%%%%%%%
% CHECK SYNTAX OF REACTION RATES (R = Rf-Rr)
% AND SPLIT THEM UP INTO Rf and Rr
%%%%%%%%%%%%%%%%%%%%%%%%%%%%%%%%%%%%%%%%%%%%%%%
errorReversibleRateDefinitions = '';
for k = 1:length(reactionsStore),
    irreversibleRates = explodePCIQM(reactionsStore(k).formula,'-');
    if length(irreversibleRates) ~= 2,
        % Need to have two parts that are separated by a '-' sign. (Forward
        % first, then reverse reaction kinetics).
        errorReversibleRateDefinitions = sprintf('%sError in rate definition of reaction rate ''%s''. It does not seem to be reversible.\n', errorReversibleRateDefinitions, reactionsStore(k).name);
    else
        % Seems fine ... save the different parts
        reactionsStore(k).forwardRate = irreversibleRates{1};
        reactionsStore(k).reverseRate = irreversibleRates{2};        
    end
end
if ~isempty(errorReversibleRateDefinitions),
    error(errorReversibleRateDefinitions);
end

%%%%%%%%%%%%%%%%%%%%%%%%%%%%%%%%%%%%%%%%%%%%%%%
% DELETE REVERSIBLE RATES AND ADD IRREVERSIBLE ONES
%%%%%%%%%%%%%%%%%%%%%%%%%%%%%%%%%%%%%%%%%%%%%%%
for k = 1:length(reactionsStore),
    model = deletereactionratesIQM(model, reactionsStore(k).name);
    forwardReactionName = strcat(reactionsStore(k).name,'_forward');
    reverseReactionName = strcat(reactionsStore(k).name,'_reverse');
    reversibleFlag = 0;
    notes = '';
    model = addreactionrateIQM(model, forwardReactionName, reactionsStore(k).forwardRate, notes, reversibleFlag, reactionsStore(k).fastFlag);
    model = addreactionrateIQM(model, reverseReactionName, reactionsStore(k).reverseRate, notes, reversibleFlag, reactionsStore(k).fastFlag);
end

%%%%%%%%%%%%%%%%%%%%%%%%%%%%%%%%%%%%%%%%%%%%%%%
% GET AND SAVE COMPARTMENT INFORMATION (original model)
%%%%%%%%%%%%%%%%%%%%%%%%%%%%%%%%%%%%%%%%%%%%%%%
[allComponentNames,allODEs] = IQMstates(originalmodel);
compartmentInformation = {}; % Strings ordered in the same way as states in the model
% check if a scaling by the compartment volume is done.
% in this case the expected syntax is 
% ODE = ("reactionterms")/compartmentvolume
% the compartment information is saved and used later to restore the ODEs
for k = 1:length(allODEs),     
    ODE = strtrim(allODEs{k});
    numberOpenParentheses = length(find(ODE == '('));
    numberClosedParentheses = length(find(ODE == ')'));
    % all eventual errorneous cases are caught in the beginning of this
    % function by calling IQMstoichiometry
    if ODE(1) == '(',
        % if the first character is an open parenthesis then assume that 
        % this is due to a adjustement to compartment sizes. here we only
        % need to keep the compartment information.
        % cut out the content of the parentheses
        closeParenthesis = find(ODE == ')');
        compartmentInformation{k} = ODE(closeParenthesis+1:end);
    else
        compartmentInformation{k} = '';
    end
end

%%%%%%%%%%%%%%%%%%%%%%%%%%%%%%%%%%%%%%%%%%%%%%%
% UPDATE RIGHT HAND SIDE OF THE ODES
%%%%%%%%%%%%%%%%%%%%%%%%%%%%%%%%%%%%%%%%%%%%%%%
% get model structure
modelstruct = IQMstruct(model);
% determine stoichiometric information of original model to 
% see where and with which stoichiometric coefficient the reversible
% reactions come into play. The stoichiometry function takes care of
% compartment factors by itself 
N = IQMstoichiometry(originalmodel,1); % set rawFlag to 1
% get info about non elementary reactions and delete them from the
% stoichiometric matrix (will be taken care off later)
L = IQMreactantstoichiometry(originalmodel,1); % reactant stoichiometry
Lnone = L+N.*(N<0);
if sum(sum(abs(Lnone))) ~= 0,
    error('Same reaction appears in at least one ODE more than once. Please see the help text to IQMmakeirreversible.');
end
% Rnone = N.*(Lnone~=0)+L.*(Lnone>0);
% % now delete
% N = N.*(Lnone == 0).*(Rnone == 0);

% get the submatrix of N corresponding only to the irreversible reactions
% reversibleIndices is same as determined above and irreversibleIndices can
% be determined by
irreversibleIndices = setdiff(1:length(IQMreactions(originalmodel)),reversibleIndices);
Nirrev = N(:,irreversibleIndices);
% get the submatrix of N corresponding only to the reversible reactions
% reversibleIndices is same as determined above
Nrev = N(:,reversibleIndices);
% clear all ODEs
for k = 1:length(modelstruct.states),
    modelstruct.states(k).ODE = '';
end
% cycle through the columns of Nirrev (irreversible reaction rates) and
% construct the corresponding ODEs
reactionNamesOriginal = IQMreactions(originalmodel);
irrevReactionNames = reactionNamesOriginal(irreversibleIndices);
for k1 = 1:size(Nirrev,2),
    % get indices of state number where to add the terms
    addTermsIndices = find(Nirrev(:,k1) ~= 0);
    for k2 = 1:length(addTermsIndices),
        % construct expression
        stoichCoeff = Nirrev(addTermsIndices(k2),k1);
        if stoichCoeff > 0,
            if stoichCoeff ~= 1,
                addTerm = sprintf('+%g*%s',abs(stoichCoeff),irrevReactionNames{k1});
            else
                addTerm = sprintf('+%s',irrevReactionNames{k1});
            end
        elseif stoichCoeff < 0,
            if stoichCoeff ~= -1,
                addTerm = sprintf('-%g*%s',abs(stoichCoeff),irrevReactionNames{k1});
            else
                addTerm = sprintf('-%s',irrevReactionNames{k1});
            end
        else
            error('This can not happen :)');
        end
        % add the new term to the corresponding ODE
        modelstruct.states(addTermsIndices(k2)).ODE = strcat(modelstruct.states(addTermsIndices(k2)).ODE, addTerm);
    end
end
% cycle trough the columns of Nrev (reversible reaction rates) and add the
% corresponding terms to the corresponding ODE expressions
for k1 = 1:size(Nrev,2),
    % get indices of state number where to add the terms
    addTermsIndices = find(Nrev(:,k1) ~= 0);
    for k2 = 1:length(addTermsIndices),
        % construct expression
        stoichCoeff = Nrev(addTermsIndices(k2),k1);
        if stoichCoeff > 0,
            if stoichCoeff ~= 1,
                addTerm = sprintf('+%g*%s_forward-%g*%s_reverse',abs(stoichCoeff),reactionsStore(k1).name,abs(stoichCoeff),reactionsStore(k1).name);
            else
                addTerm = sprintf('+%s_forward-%s_reverse',reactionsStore(k1).name,reactionsStore(k1).name);
            end                
        elseif stoichCoeff < 0,
            if stoichCoeff ~= -1,
                addTerm = sprintf('-%g*%s_forward+%g*%s_reverse',abs(stoichCoeff),reactionsStore(k1).name,abs(stoichCoeff),reactionsStore(k1).name);
            else
                addTerm = sprintf('-%s_forward+%s_reverse',reactionsStore(k1).name,reactionsStore(k1).name);
            end
        else
            error('This can not happen :)');
        end
        % add the new term to the corresponding ODE
        modelstruct.states(addTermsIndices(k2)).ODE = strcat(modelstruct.states(addTermsIndices(k2)).ODE, addTerm);
    end
end

% % handle non elementary reactions
% if sum(sum(abs(Lnone))) ~= 0,
%     % non elementary reactions are present ... need to add those reactions
%     % to the ODEs
%     for k = 1:size(N,1),
%         Lrow = Lnone(k,:);
%         Rrow = Rnone(k,:);
%         if sum(Lrow) ~= 0,
%             for k2 = 1:length(Lrow),
%                 if Lrow(k2) ~= 0,
%                     if ~reversibleReactionsAll(k2),
%                         modelstruct.states(k).ODE = sprintf('%s-%g*%s',modelstruct.states(k).ODE, Lrow(k2), reactionNamesOriginal{k2});
%                     else
%                         addTerm = sprintf('-%g*%s_forward+%g*%s_reverse',Lrow(k2),reactionNamesOriginal{k2},Rrow(k2),reactionNamesOriginal{k2});
%                         modelstruct.states(k).ODE = sprintf('%s%s',modelstruct.states(k).ODE, addTerm);
%                     end
%                 end
%             end
%         end
%         if sum(Rrow) ~= 0,
%             for k2 = 1:length(Rrow),
%                 if Rrow(k2) ~= 0,
%                     if ~reversibleReactionsAll(k2),
%                         modelstruct.states(k).ODE = sprintf('%s+%g*%s',modelstruct.states(k).ODE, Rrow(k2), reactionNamesOriginal{k2});
%                     else
%                         addTerm = sprintf('-%g*%s_forward+%g*%s_reverse',Rrow(k2),reactionNamesOriginal{k2},Lrow(k2),reactionNamesOriginal{k2});
%                         modelstruct.states(k).ODE = sprintf('%s%s',modelstruct.states(k).ODE, addTerm);
%                     end
%                 end
%             end
%         end
%     end
% end

%%%%%%%%%%%%%%%%%%%%%%%%%%%%%%%%%%%%%%%%%%%%%%%
% ADD EVENTUAL COMPARTMENT INFORMATION TO ODEs
%%%%%%%%%%%%%%%%%%%%%%%%%%%%%%%%%%%%%%%%%%%%%%%
for k = 1:length(compartmentInformation),
    if ~isempty(compartmentInformation{k}),
        modelstruct.states(k).ODE = strcat('(',modelstruct.states(k).ODE,')',compartmentInformation{k});
    end
end

% convert structure to model again
model = IQMmodel(modelstruct);

%%%%%%%%%%%%%%%%%%%%%%%%%%%%%%%%%%%%%%%%%%%%%%%
% OUTPUT THE MODEL
%%%%%%%%%%%%%%%%%%%%%%%%%%%%%%%%%%%%%%%%%%%%%%%
output = model;
return
```
