## Supplementary material for "A systems biology analysis of adrenergically stimulated adiponectin exocytosis in white adipocytes": All scripts used in this work: IQMmoietyconservations.m

```
function [varargout] = IQMmoietyconservations(varargin)
% IQMmoietyconservations: determines the moitey conservations that are
% present in a model. The model can be specified as IQMmodel or as ODE file.
%
% USAGE:
% ======
% [] = IQMmoietyconservations(model)         
% [] = IQMmoietyconservations(model,stateValues)         
% [] = IQMmoietyconservations(model,stateValues,tol)         
% [formulas,constants] = IQMmoietyconservations(model,stateValues,tol)         
% [depVarIndex, depVarConstant, depVarFactor, message] = IQMmoietyconservations(model)         
% [depVarIndex, depVarConstant, depVarFactor, message] = IQMmoietyconservations(model,stateValues)         
% [depVarIndex, depVarConstant, depVarFactor, message] = IQMmoietyconservations(model,stateValues,tol)         
%
% model: IQMmodel or ODE file model description
% stateValues: values of the state variables for which to perform the
%   analysis
% tol: Tolerance for the determination of the number of algebraic
% 	relationships in the model. If the method fails than this
%   might be due to a wrong rank computation and in this case
%   the tolerance should be increased. If set, the same tolerance
%   setting is used when determining the indices of the
%   dependent variables.
%
%
% DEFAULT VALUES:
% ===============
% stateValues: if not given, the state stored in the model is used to
%   determine the moiety conservations. It is a good idea to a systems
%   steady-state for the computation of the conservations
%
% Output Arguments:
% =================
% depVarIndex: index of dependent states
% depVarConstant: Sum of states taking part in conservation
% depVarFactor: Factor for states - non-zero values for the ones that take
%   part in conservation
% message: Any message that otherwise is printed to the MATLAB window
% formulas: A cell-array containing the equations of the found algebraic
%           relations

% <<<COPYRIGHTSTATEMENT - IQM TOOLS LITE>>>

global message
message = '';

%%%%%%%%%%%%%%%%%%%%%%%%%%%%%%%%%%%%%%%%%%%%%%%
% CHECK IF IQMMODEL OR FILENAME
%%%%%%%%%%%%%%%%%%%%%%%%%%%%%%%%%%%%%%%%%%%%%%%
if strcmp('IQMmodel',class(varargin{1})),
    % IQMmodel
    iqm = varargin{1};
    % Create temporary ODE file
    [ODEfctname, ODEfilefullpath] = IQMcreateTempODEfile(iqm);    
else
    % ODEfctname of ODE file
    ODEfctname = varargin{1};
    % set iqm to empty!
    iqm = [];
    % Check if file exists
    if exist(strcat(ODEfctname,'.m')) ~= 2,
        error('ODE file could not be found.');
    end
end

%%%%%%%%%%%%%%%%%%%%%%%%%%%%%%%%%%%%%%%%%%%%%%%
% HANDLE VARIABLE INPUT ARGUMENTS
%%%%%%%%%%%%%%%%%%%%%%%%%%%%%%%%%%%%%%%%%%%%%%%
if nargin == 1,
    % handle numeric and non-numeric ICs
    stateValues = IQMcalcICvector(ODEfctname);
    tol = 0;
elseif nargin == 2,
    stateValues = varargin{2};
    stateValues = stateValues(:);
    tol = 0;
elseif nargin == 3,
    stateValues = varargin{2};
    stateValues = stateValues(:);
    tol = varargin{3};
else
    error('Incorrect number of input arguments.');
end

%%%%%%%%%%%%%%%%%%%%%%%%%%%%%%%%%%%%%%%%%%%%%%%
% DETERMINE THE MOIETY CONSERVATIONS
%%%%%%%%%%%%%%%%%%%%%%%%%%%%%%%%%%%%%%%%%%%%%%%
[depVarIndex, depVarConstant, depVarFactor] = getMoietyConservations(iqm,ODEfctname,stateValues,tol);
% invert order of moiety conservations (to be able to evaluate them
% sequentially in case of model reduction)
depVarIndex(1:end) = depVarIndex(end:-1:1);
depVarConstant(1:end) = depVarConstant(end:-1:1);
depVarFactor(1:end,:) = depVarFactor(end:-1:1,:);

%%%%%%%%%%%%%%%%%%%%%%%%%%%%%%%%%%%%%%%%%%%%%%%
% CONSTRUCT RESULT MESSAGE
%%%%%%%%%%%%%%%%%%%%%%%%%%%%%%%%%%%%%%%%%%%%%%%
MCarray = {};
MCconstants = [];
if ~isempty(depVarIndex),
    message = sprintf('%s\nFollowing moitey conservations have been found:\n',message);
    variableNames = feval(ODEfctname,'states');
    for k1 = 1:length(depVarIndex),
        MCarray{k1} = sprintf('%s = moietyConserv_%d',variableNames{depVarIndex(k1)},k1);
        MCconstants(k1) = depVarConstant(k1);
        message = sprintf('%s\n%s = %g',message,variableNames{depVarIndex(k1)},depVarConstant(k1));
        for k2 = 1:length(depVarFactor(k1,:)),
            if abs(depVarFactor(k1,k2)) > 0,
                % Here a check against 0 can be done since zerocheck
                % already done and small values set to identically zero.
                if depVarFactor(k1,k2) > 0
                    element = sprintf('+%g*',depVarFactor(k1,k2));
                else
                    element = sprintf('-%g*',abs(depVarFactor(k1,k2)));
                end
                MCarray{k1} = sprintf('%s%s%s',MCarray{k1},element,variableNames{k2});
                message = sprintf('%s%s%s',message,element,variableNames{k2});
            end
        end
    end
    message = sprintf('%s\n',message);
else
    message = sprintf('%sNo moiety conservations have been found.\n',message);
end
message = sprintf('%s\nIf this result does not seem correct, do the analysis\nusing a different initial condition and/or tolerance setting.',message);

%%%%%%%%%%%%%%%%%%%%%%%%%%%%%%%%%%%%%%%%%%%%%%%
% PROCESS VARIABLE OUTPUT ARGUMENTS
%%%%%%%%%%%%%%%%%%%%%%%%%%%%%%%%%%%%%%%%%%%%%%%
if nargout == 0,
    % no output argument => simply display the results
    disp(message);
elseif nargout == 2,
    varargout{1} = MCarray;
    varargout{2} = MCconstants;
elseif nargout == 4,
    varargout{1} = depVarIndex;
    varargout{2} = depVarConstant;
    varargout{3} = depVarFactor;
    varargout{4} = message;
else
    error('Incorrect number of output arguments');
end

%%%%%%%%%%%%%%%%%%%%%%%%%%%%%%%%%%%%%%%%%%%%%%%
% DELETE FILE IF IQMMODEL
%%%%%%%%%%%%%%%%%%%%%%%%%%%%%%%%%%%%%%%%%%%%%%%
if strcmp('IQMmodel',class(varargin{1})),
    IQMdeleteTempODEfile(ODEfilefullpath);
end
return

%%%%%%%%%%%%%%%%%%%%%%%%%%%%%%%%%%%%%%%%%%%%%%%%%%%%%%%%%%%%%%%%%%%%%%%%%%
% GET THE MOIETY CONSERVATIONS
%%%%%%%%%%%%%%%%%%%%%%%%%%%%%%%%%%%%%%%%%%%%%%%%%%%%%%%%%%%%%%%%%%%%%%%%%%
% the function works either based on the jacobian or if it is possible on
% the stoichiometric matrix. the latter is only possible if the model is
% given as IQMmodel and the odes are fully described in terms of reaction
% rates.
function [depVarIndex, depVarConstant, depVarFactor] = getMoietyConservations(iqm,ODEfctname,stateValues,tol)
global message;
% initialize return variables
depVarIndex = [];
depVarConstant = [];
depVarFactor = [];
% get initial condition 
initialCondition = stateValues;
% check if model given as IQMmodel:
if ~isempty(iqm),
    % check if stoichiometric matrix gives full information
    N = IQMstoichiometry(iqm,0,1); % no rawFlag but silentFlag!
    if length(initialCondition) == size(N,1),
        % yes, full information available
        useStoich = 1;
    else
        useStoich = 0;
    end
else
    useStoich = 0;
end
% handle stoich and jac differently
if useStoich == 0,
    % use Jacobian!
    % Get jacobian and determine its rankdeficiency - use default deltaX for
    % numerical differentiation
    J = IQMjacobian(ODEfctname,initialCondition);

% handle ARs by reducing J
nrARs = length(feval(ODEfctname,'algebraic'));
J = J(1:end-nrARs, 1:end-nrARs);
  
    if tol == 0,
        d = length(J) - rank(J);        
    else
        d = length(J) - rank(J,tol);
    end
else
    % use stoichiometry
    if tol == 0,
        X = rref([N,eye(size(N,1))]);
    else
        X = rref([N,eye(size(N,1))],tol);
    end        
    Nx = X(:,1:size(N,2));
    d = length(find(sum(abs(Nx)')'==0));
end
if d > 0,
    % print out a message
    message = sprintf('%sThe system has a rank deficiency of %d. This might be due to moiety or\nother conservations in the model, to the excessive use of zero initial\nconditions and/or integrating behavior of the system. If this does\nnot seem to be correct, please check the initial conditions and/or\nset a different value for the tolerance "tol" in the options.\n',message,d);
    % The systems contains algebraic relationships (moiety conservations)
    % Handle stoich and jac differently
    if useStoich == 0,
        message = sprintf('%s\nThe algebraic relationships are determined using the Jacobian.\nThis is numerically not very stable and you need eventually\nto change tolerance "tol" settings.\n',message);
        % Use the singular value decomposition of the Jacobian to determine
        % the algebraic relationships
        [U,S,V] = svd(J);
        U0 = U(:,length(J)-d+1:length(J));
        if tol == 0,
            factors = rref(U0');
        else
            factors = rref(U0',tol);
        end
    else
        message = sprintf('%s\nThe algebraic relationships are determined using the Stoichiometric matrix.\n',message);
        % Use rref of the Stoichiometric Matrix
        factors = X(end-d+1:end,size(N,2)+1:end);
    end    
    % set smaller values than tolerance to zero
    factors(find(abs(factors)<tol)) = 0;
% handle ARs by reducing initialConditions
nrARs = length(feval(ODEfctname,'algebraic'));
iCs = initialCondition(1:end-nrARs);    
    constantValues = factors*iCs;
    % Process the result and get the algebraic equations
    for k = 1:size(factors,1),
        relation = [factors(k,:) constantValues(k)];
        % find largest element (abs) in the first n-1 elements
        [value,index] = max(abs(relation(1:length(relation)-1)));
        % normalize the relation
        relation = relation./relation(index);
        % the index of the largest element is the index of a dependent variable
        depVarIndex = [depVarIndex index];
        % the constant value is given by the last element in relation
        depVarConstant = [depVarConstant; relation(length(relation))];
        % the factor vector is constructed as follows:
        factorVector = -relation(1:length(relation)-1); factorVector(index) = 0;
        depVarFactor = [depVarFactor; factorVector];
    end
end
return
```
