## Supplementary material for "A systems biology analysis of adrenergically stimulated adiponectin exocytosis in white adipocytes": All scripts used in this work: IQMreducemodel.m

```
function [reducedmodel] = IQMreducemodel(varargin)
% IQMreducemodel: Reduces an IQMmodel by identifying algebraic relations 
% between time dependent variable, defined by differential equations, and 
% deleting the dependent variables. This function first checks if the 
% IQMmodel contains algebraic relations between the state variables. If 
% there are, it lets the user decide which variables to choose as dependent
% ones and to replace the ODEs by algebraic relations.
%
% Due to moiety conservations algebraic relations are often present in 
% biological systems. However, for several analysis methods models are 
% required to be non-singular. This function helps to avoid this problem.
%
% USAGE:
% ======
% [reducedmodel] = IQMreducemodel(model)         
% [reducedmodel] = IQMreducemodel(model,tol)         
%
% model: IQMmodel model to reduce
% tol: tolerance to be used to determine moiety conservations
%
% DEFAULT VALUES:
% ===============
% tol: same as for IQMmoietyconservations
%
% Output Arguments:
% =================
% reducedmodel: Reduced IQMmodel. If no algebraic relations were present in
%       the model, the initial model is returned unchanged.

% <<<COPYRIGHTSTATEMENT - IQM TOOLS LITE>>>

%%%%%%%%%%%%%%%%%%%%%%%%%%%%%%%%%%%%%%%%%%%%%%%
% NEEDED GLOBAL VARIABLES
%%%%%%%%%%%%%%%%%%%%%%%%%%%%%%%%%%%%%%%%%%%%%%%
global ODEfctname 

%%%%%%%%%%%%%%%%%%%%%%%%%%%%%%%%%%%%%%%%%%%%%%%
% CHECK IF IQMMODEL OR FILENAME
%%%%%%%%%%%%%%%%%%%%%%%%%%%%%%%%%%%%%%%%%%%%%%%
if strcmp('IQMmodel',class(varargin{1})),
    % IQMmodel
    iqm = varargin{1};
    % Create temporary ODE file
    [ODEfctname, ODEfilefullpath] = IQMcreateTempODEfile(iqm);    
else
    error('This function can not be applied to ODE files.');
end

if nargin == 1,
    tol = 0;
elseif nargin == 2,
    tol = varargin{2};
else
    error('Incorrect number of input arguments.');
end

%%%%%%%%%%%%%%%%%%%%%%%%%%%%%%%%%%%%%%%%%%%%%%%
% CHECK AND REDUCE THE MODEL
%%%%%%%%%%%%%%%%%%%%%%%%%%%%%%%%%%%%%%%%%%%%%%%
reducedmodel = reduceModel(iqm,tol);

%%%%%%%%%%%%%%%%%%%%%%%%%%%%%%%%%%%%%%%%%%%%%%%
% DELETE FILE 
%%%%%%%%%%%%%%%%%%%%%%%%%%%%%%%%%%%%%%%%%%%%%%%
IQMdeleteTempODEfile(ODEfilefullpath);
return

%%%%%%%%%%%%%%%%%%%%%%%%%%%%%%%%%%%%%%%%%%%%%%%%%%%%%%%%%%%%%%%%%%%%%%%%%%
% CHECK AND REDUCE THE MODEL
%%%%%%%%%%%%%%%%%%%%%%%%%%%%%%%%%%%%%%%%%%%%%%%%%%%%%%%%%%%%%%%%%%%%%%%%%%
function [iqmReduced] = reduceModel(iqm,tol)
% needed global variables
global ODEfctname

%%%%%%%%%%%%%%%%%%%%%%%%%%%%%%%%%%%%%%%%%%%%%%%
% DETERMINE THE MOIETY CONSERVATIONS
%%%%%%%%%%%%%%%%%%%%%%%%%%%%%%%%%%%%%%%%%%%%%%%
% Get initial conditions (numeric and non-numeric ICs)
initialCondition = IQMcalcICvector(ODEfctname);
[depVarIndex, depVarConstant, depVarFactor, message] = IQMmoietyconservations(iqm,initialCondition,tol);

%%%%%%%%%%%%%%%%%%%%%%%%%%%%%%%%%%%%%%%%%%%%%%%
% CHECK IF REDUCTION NECESSARY
%%%%%%%%%%%%%%%%%%%%%%%%%%%%%%%%%%%%%%%%%%%%%%%
if isempty(depVarIndex),
    % System is non-singular - return the original system
    disp('Model is non-singular. Nothing is done.');
    iqmReduced = iqm; 
    return
end

%%%%%%%%%%%%%%%%%%%%%%%%%%%%%%%%%%%%%%%%%%%%%%%
% REMOVE NON-NUMERIC ICs (if present)
%%%%%%%%%%%%%%%%%%%%%%%%%%%%%%%%%%%%%%%%%%%%%%%
% print a warning if non-numeric initial conditions (reduction will be
% correct but non-numeric ICs have been replaced by numeric ones).
if ~hasonlynumericICsIQM(iqm),
    numICs = IQMcalcICvector(iqm);
    ms = struct(iqm);
    for k=1:length(ms.states),
        ms.states(k).initialCondition = numICs(k);
    end
    iqm = IQMmodel(ms);
end

%%%%%%%%%%%%%%%%%%%%%%%%%%%%%%%%%%%%%%%%%%%%%%%
% CHECK THE CONSERVATION EQUATIONS 
%%%%%%%%%%%%%%%%%%%%%%%%%%%%%%%%%%%%%%%%%%%%%%%
% To be able to include them as variables in an IQMmodel it is important
% that the moietyconservations are independent of each other. No algebraic
% loop is allowed.
% for k1 = 1:size(depVarFactor,1),
%     for k2 = 1:size(depVarFactor,1),
%         if k1 ~= k2,
%            if ~isempty(intersect(find(depVarFactor(k1,:)~=0),find(depVarFactor(k2,:)~=0))),
%                error(sprintf('The model can not be reduced due to dependency among the conservations.\nIn the future the toolbox will support these kind of things, but not today.\nSorry!'));
%            end
%         end
%     end
% end

%%%%%%%%%%%%%%%%%%%%%%%%%%%%%%%%%%%%%%%%%%%%%%%
% CONVERSION OF THE MOIETY CONSERVATIONS
%%%%%%%%%%%%%%%%%%%%%%%%%%%%%%%%%%%%%%%%%%%%%%%
% need to convert these results to a different format.
% result format: dependentVariable = constantTerm + factor1*variable1 + ... + factorn*variablen
% needed format: constantTerm = dependentVariable + NewFactor1*variable1 -
% ... + NewFactorn*variablen
% => change sign in depVarFactor and add a 1 in the places corresponding to
% the dependent variables.
depVarFactor = -depVarFactor;
for k = 1:length(depVarIndex),
    depVarFactor(k,depVarIndex(k)) = 1;
end

%%%%%%%%%%%%%%%%%%%%%%%%%%%%%%%%%%%%%%%%%%%%%%%
% PROCESS SYSTEM
%%%%%%%%%%%%%%%%%%%%%%%%%%%%%%%%%%%%%%%%%%%%%%%
% System is singular - process the algebraic relations
d = length(depVarIndex);

%%%%%%%%%%%%%%%%%%%%%%%%%%%%%%%%%%%%%%%%%%%%%%%
% DISPLAY ALL ALGEBRAIC RELATIONS
%%%%%%%%%%%%%%%%%%%%%%%%%%%%%%%%%%%%%%%%%%%%%%%
disp(message);

%%%%%%%%%%%%%%%%%%%%%%%%%%%%%%%%%%%%%%%%%%%%%%%
% DECIDE WHICH STATES TO REMOVE (USER)
%%%%%%%%%%%%%%%%%%%%%%%%%%%%%%%%%%%%%%%%%%%%%%%
text = sprintf('You need to decide, for each algebraic relation, which state to remove from the model.\n');
disp(text);
% handle the case when events are present in the model (the states that are
% changed by events can not be removed). If this leads to no reducable
% states an error message will be displayed.
iqms = struct(iqm);
eventvariables = {};
eventstateindices = [];
if ~isempty(iqms.events),
    for k=1:length(iqms.events),
        eventvariables = {eventvariables{:} iqms.events(k).assignment(1:end).variable};
    end
    eventvariables = unique(eventvariables);
    % check which of these eventvariables are states 
    eventstateindices = stateindexIQM(iqm,eventvariables); 
    % remove the elements that are -1 (no states) ... the rest are indices
    % of states that are changed by events
    eventstateindices(eventstateindices<0) = [];
    % mark the elements that correspond to state variables that have
    % a zero right hand side of the ODE (these become parameters and thus
    % can be changed by events).
    for k=1:length(eventstateindices),
        if iqms.states(eventstateindices(k)).ODE == '0',
            eventstateindices(k) = -1;
        end
    end
    % and remove them
    eventstateindices(eventstateindices<0) = [];
    % eventstateindices now contains the indices of the states that are not
    % allowed to be removed from the set of ODEs. simply because these can
    % be changed by events and this is only possible if they are kept as
    % states.
end
statesRemove = [];
variableNames = feval(ODEfctname,'states');
% cycle through all the algebraic relations, display them and let the
% user decide which state to remove.
for k1 = 1:length(depVarConstant),
    % delete the eventstateindices states from the states that are removable
    removableStates = setdiff(find(depVarFactor(k1,:) ~= 0), eventstateindices);
    % remove the already removed states
    indices = setdiff(removableStates,statesRemove);
    % check if at least one state removable
    if isempty(indices),
        error(sprintf('There is a moiety conservation but none of the states can be removed.\nThis is most certainly due to the fact that at least one event is present\nin the model that changes the states involved in this moiety conservation.\nDeleting the states would lead to an incorrect event behavior.'));
    end
    text = sprintf('Relation %d: %s',k1,getTextAlgebraicRelation(depVarConstant(k1),depVarFactor(k1,:)));
    disp(text);
    text = sprintf('Choose the number of the state that should be removed (1-%d).',length(indices));
    disp(text);
    text = '';
    for k2 = 1:length(indices),
        text = sprintf('%s (%d)%s  ',text,k2,variableNames{indices(k2)});
    end
    disp(text);
    indexRemove = input('State to remove: ');
    if indexRemove > length(indices) || indexRemove < 1,
        error('Wrong input!');
    end
    statesRemove = [statesRemove indices(indexRemove)];
end

%%%%%%%%%%%%%%%%%%%%%%%%%%%%%%%%%%%%%%%%%%%%%%%
% CONSTRUCT REDUCED MODEL
%%%%%%%%%%%%%%%%%%%%%%%%%%%%%%%%%%%%%%%%%%%%%%%
% get the structure of the IQMmodel
modelStructure = IQMstruct(iqm); % will contain the reduced information
fullModelStructure = IQMstruct(iqm); % contains the full information (needed only for updating the additional information fields of the variables)
% create new "states" entry and copy only the states that are not
% deleted into it
states = [];
statesCount = 1;
for k1 = 1:length(modelStructure.states),
    if sum(ismember(statesRemove,k1)) == 0,
        % copy this state information to new model
        states(statesCount).name = modelStructure.states(k1).name;
        states(statesCount).initialCondition = modelStructure.states(k1).initialCondition;
        states(statesCount).ODE = modelStructure.states(k1).ODE;
        states(statesCount).type = modelStructure.states(k1).type;
        states(statesCount).compartment = modelStructure.states(k1).compartment;
        states(statesCount).unittype = modelStructure.states(k1).unittype;
        states(statesCount).notes = modelStructure.states(k1).notes;
        statesCount = statesCount + 1;
    end
end
% Include the state information in the model structure
modelStructure.states = states;
% Now add the algebraic relations as variables.
% It is important to add the relations before all other variables!
% Since they might be used in the other variables!
variables = [];
variableCount = 1;
parameterCount = length(modelStructure.parameters)+1;
% First add the algebraic relations
for k1 = 1:length(depVarConstant),
    % Construct variable formula corresponding to algebraic relation
    % Get the constant term
    constantTerm = depVarConstant(k1);
    % Scale the factor vector with -1 and set reduced element to zero
    factor = depVarFactor(k1,:);
    scaleFactor = 1/abs(factor(statesRemove(k1)));
    factor = -factor*scaleFactor;
    factor(statesRemove(k1)) = 0;
    % Scale the constant term
    constantTerm = constantTerm*scaleFactor;
    % get the variableName
    variable = variableNames{statesRemove(k1)};
    % Construct the formula string
    moietyConservationName = sprintf('conserv%d',k1);
    formula = sprintf('%s',moietyConservationName);
    % CHECK factor ... if all zero then add MC thing as parameter not as
    % variable!
    if sum(abs(factor)) ~= 0,
        for k2 = 1:length(factor),
            if factor(k2) > 0,
                % here 0 can be used (zerocheck already done)
                formula = sprintf('%s+%g*%s',formula,abs(factor(k2)),variableNames{k2});
            elseif factor(k2) < 0,
                % here 0 can be used (zerocheck already done)
                formula = sprintf('%s-%g*%s',formula,abs(factor(k2)),variableNames{k2});
            end
        end
        % Add algebraic relation as variable
        variables(variableCount).name = variable;
        variables(variableCount).formula = formula;
        variables(variableCount).notes = 'Removed algebraic relation';
        variables(variableCount).type = fullModelStructure.states(statesRemove(k1)).type;
        variables(variableCount).compartment = fullModelStructure.states(statesRemove(k1)).compartment;
        variables(variableCount).unittype = fullModelStructure.states(statesRemove(k1)).unittype;
        variableCount = variableCount + 1;
        % add the constant term as a parameter
        modelStructure.parameters(parameterCount).name = moietyConservationName;
        modelStructure.parameters(parameterCount).value = str2num(sprintf('%g',constantTerm));
        moietyConservationNotes = sprintf('Moiety conservation for: %s',getTextAlgebraicRelation(depVarConstant(variableCount-1),depVarFactor(variableCount-1,:)));
        moietyConservationNotes = moietyConservationNotes(1:strfind(moietyConservationNotes,'=')-1);
        modelStructure.parameters(parameterCount).type = 'isParameter';
        modelStructure.parameters(parameterCount).compartment = '';
        modelStructure.parameters(parameterCount).unittype = '';
        modelStructure.parameters(parameterCount).notes = moietyConservationNotes;
        parameterCount = parameterCount + 1;
    else
        % add the constant term as a parameter
        modelStructure.parameters(parameterCount).name = variable;
        modelStructure.parameters(parameterCount).value = str2num(sprintf('%g',constantTerm));
        moietyConservationNotes = sprintf('Moiety conservation for: %s',variable);
        modelStructure.parameters(parameterCount).type = fullModelStructure.states(statesRemove(k1)).type;
        modelStructure.parameters(parameterCount).compartment = fullModelStructure.states(statesRemove(k1)).compartment;
        modelStructure.parameters(parameterCount).unittype = fullModelStructure.states(statesRemove(k1)).unittype;
        modelStructure.parameters(parameterCount).notes = moietyConservationNotes;
        parameterCount = parameterCount + 1;        
    end
end
% Now add the other variables
for k = 1:length(fullModelStructure.variables),
    variables(variableCount).name = fullModelStructure.variables(k).name;
    variables(variableCount).formula = fullModelStructure.variables(k).formula;
    variables(variableCount).type = fullModelStructure.variables(k).type;
    variables(variableCount).compartment = fullModelStructure.variables(k).compartment;
    variables(variableCount).unittype = fullModelStructure.variables(k).unittype;
    variables(variableCount).notes = fullModelStructure.variables(k).notes;
    variableCount = variableCount + 1;
end
% Include the variable information in the model structure
modelStructure.variables = variables;
% Create IQMmodel with reduced model structure
iqmReduced = IQMmodel(modelStructure);

if ~hasonlynumericICsIQM(iqm),
    disp('Please note: The model contains non-numeric initial conditions. In the reduced model');
    disp('these have been replaced by corresponding numeric initial conditions.');
end
return

%%%%%%%%%%%%%%%%%%%%%%%%%%%%%%%%%%%%%%%%%%%%%%%%%%%%%%%%%%%%%%%%%%%%%%%%%%
% DISPLAY THE ALGEBRAIC RELATIONS
%%%%%%%%%%%%%%%%%%%%%%%%%%%%%%%%%%%%%%%%%%%%%%%%%%%%%%%%%%%%%%%%%%%%%%%%%%
function [text] = getTextAlgebraicRelation(depVarConstant,depVarFactor)
    % needed global variables
    global ODEfctname
    % get variable names from ODE file
    variableNames = feval(ODEfctname,'states');
    text = sprintf(' = %g',depVarConstant);
    for k2 = 1:length(depVarFactor),
        if abs(depVarFactor(k2)) > 0,    % here 0 can be used (zerocheck already done)
            if depVarFactor(k2) > 0
                element = sprintf('+ %g',depVarFactor(k2));
            else 
                element = sprintf('- %g',abs(depVarFactor(k2)));
            end
            text = sprintf('%s %s %s',element,variableNames{k2},text);
        end
    end
return
```
