## Supplementary material for "A systems biology analysis of adrenergically stimulated adiponectin exocytosis in white adipocytes": All scripts used in this work: IQMsteadystate.m

```
function [varargout] = IQMsteadystate(varargin)
% IQMsteadystate: determines a steady-state of an IQMmodel or an ODE file 
% description created by IQMcreateODEfile. This function should only be used 
% for time-independent systems. Otherwise an error will
% occur. The function is able to deal with models having a singular 
% Jacobian, e.g., due to moity conservations.
%
% USAGE:
% ======
% [steadystate,residual,message] = IQMsteadystate(model)         
% [steadystate,residual,message] = IQMsteadystate(model,initialCondition)         
% [steadystate,residual,message] = IQMsteadystate(model,initialCondition,OPTIONS)
% [steadystate,residual] = IQMsteadystate(model)         
% [steadystate,residual] = IQMsteadystate(model,initialCondition)         
% [steadystate,residual] = IQMsteadystate(model,initialCondition,OPTIONS)
% [steadystate] = IQMsteadystate(model)         
% [steadystate] = IQMsteadystate(model,initialCondition)         
% [steadystate] = IQMsteadystate(model,initialCondition,OPTIONS)
%                                   
% model: IQMmodel model or name of ODE file (without .m suffix)
% initialCondition: Array with initial conditions around which to start the
%   search for a steady-state. If not given the initial conditions or given
%   as an empty vector the initial values are taken from the IQMmodel.
% OPTIONS: structure containing options
%          OPTIONS.TolFun: Tolerance for max element in function evaluation
%          OPTIONS.tol: Tolerance for the determination of the number of algebraic
%               relationships in the model. If the method fails than this
%               might be due to a wrong rank computation and in this case
%               the tolerance should be increased. If set, the same tolerance
%               setting is used when determining the indices of the
%               dependent variables.
%          OPTIONS.MaxIter: Maximum number of iterations
%          OPTIONS.Delta: Step length for numerical differentiation to obtain 
%               the Jacobian.
%
% DEFAULT VALUES:
% ===============
% initialCondition: the initial conditions stored in the model
% OPTIONS.TolFun:   1e-11
% OPTIONS.tol:      standard MATLAB tolerance: s = svd(A); tol = max(size(A))*eps(max(s));
% OPTIONS.MaxIter:  1000
% OPTIONS.Delta:    1e-6
%
% Output Arguments:
% =================
% steadystate: array with steadystate values
% residual: 2-norm of the derivatives at the found steady-state. residual
%           is optional
% message: message about the steady state not able to find and/or message about 
%          found algebraic relationships / moiety conservations.
%          Message as output argument is optional. If omitted, the message
%          will be displayed in the Matlab window.

% <<<COPYRIGHTSTATEMENT - IQM TOOLS LITE>>>

%%%%%%%%%%%%%%%%%%%%%%%%%%%%%%%%%%%%%%%%%%%%%%%
% NEEDED GLOBAL VARIABLES
%%%%%%%%%%%%%%%%%%%%%%%%%%%%%%%%%%%%%%%%%%%%%%%
global ODEfctname TolFun tol MaxIter Delta message

%%%%%%%%%%%%%%%%%%%%%%%%%%%%%%%%%%%%%%%%%%%%%%%
% INITIALIZATION OF MESSAGE
%%%%%%%%%%%%%%%%%%%%%%%%%%%%%%%%%%%%%%%%%%%%%%%
message = '';

%%%%%%%%%%%%%%%%%%%%%%%%%%%%%%%%%%%%%%%%%%%%%%%
% CHECK IF IQMMODEL OR FILENAME
%%%%%%%%%%%%%%%%%%%%%%%%%%%%%%%%%%%%%%%%%%%%%%%
if isIQMmodel(varargin{1}),
    % IQMmodel
    iqm = varargin{1};
    % check if the model uses delays ... if yes then take care of it
    if usedelayIQM(iqm),
        disp('Delays are present in the model. They are going to be');
        disp('removed automatically for steady-state computation.');
        iqm = removedelayIQM(iqm);
    end
    % Create temporary ODE file
    [ODEfctname, ODEfilefullpath] = IQMcreateTempODEfile(iqm);    
else
    % ODEfctname of ODE file
    ODEfctname = varargin{1};
    % empty iqm variable
    iqm = [];
    % Check if file exists
    if exist(strcat(ODEfctname,'.m')) ~= 2,
        error('ODE file could not be found.');
    end
end

%%%%%%%%%%%%%%%%%%%%%%%%%%%%%%
% Set default values
%%%%%%%%%%%%%%%%%%%%%%%%%%%%%%
TolFun  = 1e-11;
tol     = 0; % standard MATLAB tolerance: s = svd(A); tol = max(size(A))*eps(max(s));
MaxIter = 1000;
Delta   = 1e-6;

%%%%%%%%%%%%%%%%%%%%%%%%%%%%%%%%%%%%%%%%%%%%%%%
% PROCESS VARIABLE INPUT ARGUMENTS
%%%%%%%%%%%%%%%%%%%%%%%%%%%%%%%%%%%%%%%%%%%%%%%
if nargin == 1,
    % handle non-numeric and numeric ICs
    initialCondition = IQMcalcICvector(ODEfctname);
    OPTIONS = []; % default options
elseif nargin == 2,
    initialCondition = varargin{2};
    OPTIONS = []; % default options
elseif nargin == 3,
    initialCondition = varargin{2};
    OPTIONS = varargin{3};
else
    error('Wrong number of input arguments');
end
% if empty initial condition => default values
if isempty(initialCondition),
    % handle non-numeric and numeric ICs
    initialCondition = IQMcalcICvector(ODEfctname);
end    
% initialCondition should be a column vector!
initialCondition = initialCondition(:); 
% if OPTIONS = [] then use default values
if ~isempty(OPTIONS),
    % TolFun
    if isfield(OPTIONS,'TolFun'),
        if ~isempty(OPTIONS.TolFun),
            TolFun = OPTIONS.TolFun;
        end
    end
    % tol
    if isfield(OPTIONS,'tol'),
        if ~isempty(OPTIONS.tol),
            tol = OPTIONS.tol;
        end
    end
    % MaxIter
    if isfield(OPTIONS,'MaxIter'),
        if ~isempty(OPTIONS.MaxIter),
            MaxIter = OPTIONS.MaxIter;
        end
    end
    % Delta
    if isfield(OPTIONS,'Delta'),
        if ~isempty(OPTIONS.Delta),
            Delta = OPTIONS.Delta;
        end
    end
end

%%%%%%%%%%%%%%%%%%%%%%%%%%%%%%%%%%%%%%%%%%%%%%%
% GET THE STEADY STATE FOR THE MODEL
%%%%%%%%%%%%%%%%%%%%%%%%%%%%%%%%%%%%%%%%%%%%%%%
steadystate = getSteadyState(initialCondition,iqm);
if ~isempty(steadystate),
    residual = norm(feval(ODEfctname,0,steadystate));
else
    messageText = 'Steady state could not be found. Try different options and/or a different starting guess.';
    message = sprintf('%s\n%s\n',message,messageText);
    residual = [];
end

%%%%%%%%%%%%%%%%%%%%%%%%%%%%%%%%%%%%%%%%%%%%%%%
% PROCESS VARIABLE OUTPUT ARGUMENTS
%%%%%%%%%%%%%%%%%%%%%%%%%%%%%%%%%%%%%%%%%%%%%%%
if nargout == 2,
    varargout{1} = steadystate;
    varargout{2} = residual;
    disp(message);
elseif nargout == 3,
    varargout{1} = steadystate;
    varargout{2} = residual;
    varargout{3} = message;
elseif nargout < 2, 
    varargout{1} = steadystate;
    disp(message);
end

%%%%%%%%%%%%%%%%%%%%%%%%%%%%%%%%%%%%%%%%%%%%%%%
% DELETE FILE IF IQMMODEL
%%%%%%%%%%%%%%%%%%%%%%%%%%%%%%%%%%%%%%%%%%%%%%%
if strcmp('IQMmodel',class(varargin{1})),
    IQMdeleteTempODEfile(ODEfilefullpath);
end
return

%%%%%%%%%%%%%%%%%%%%%%%%%%%%%%%%%%%%%%%%%%%%%%%%%%%%%%%%%%%%%%%%%%%%%%%%%%
% GET THE STEADY STATE OF THE MODEL
%%%%%%%%%%%%%%%%%%%%%%%%%%%%%%%%%%%%%%%%%%%%%%%%%%%%%%%%%%%%%%%%%%%%%%%%%%
function [steadystate] = getSteadyState(initialCondition,iqm)
% Define some global variables that are needed in other functions
global depVar factors constantValues ODEfctname TolFun tol MaxIter Delta message
EXITFLAG = 0;
% Before determining the steady-state it is necessary to check if there are
% some algebraic relationships in the model. If there are not the
% steady-state is determined directly. If there are, the algebraic
% relations are determined and the steady-state is calculated for a reduced
% system.
% check if model given as IQMmodel:
if ~isempty(iqm),
    % check if stoichiometric matrix gives full information
    N = IQMstoichiometry(iqm,0,1); % no rawFlag,  but silentFlag
    if length(initialCondition) == size(N,1),
        % yes, full information available
        useStoich = 1;
    else
        useStoich = 0;
    end
else
    useStoich = 0;
end
% handle stoich and jac differently
if useStoich == 0,
    % use Jacobian!
    % Get jacobian and determine its rankdeficiency - use default deltaX for
    % numerical differentiation
    J = IQMjacobian(ODEfctname,initialCondition);
    if tol == 0,
        d = length(J) - rank(J);        
    else
        d = length(J) - rank(J,tol);
    end
else
    % use stoichiometry
    if tol == 0,
        X = rref([N,eye(size(N,1))]);
    else
        X = rref([N,eye(size(N,1))],tol);
    end        
    Nx = X(:,1:size(N,2));
    d = length(find(sum(abs(Nx)')'==0));
end
if d == 0,
    % Jacobian has full rank => no algebraic relations => "simple"
    % computation of the steady-state
    % The systems does not contain algebraic relations
    OPTIONS = [];
    OPTIONS.MaxIter = MaxIter;
    OPTIONS.TolFun = TolFun;
    OPTIONS.Delta = Delta;
    [steadystate,FVAL,EXITFLAG] = fsolveIQM(@model,initialCondition,OPTIONS);
else
    % message
    messageText = sprintf('The system has a rank deficiency of %d. This might be due to moiety or\nother conservations in the model, to the excessive use of zero initial\nconditions and/or integrating behavior of the system. If this does\nnot seem to be correct, please check the initial conditions and/or\nset a different value for the tolerance "tol" in the options.',d);
    message = sprintf('%s\n%s\n',message,messageText);
    % Jacobian is singular => assume that algebraic relations are present
    % and solve for the steady-state by taking into account these algebraic
    % relations, by considering the singular output directions
    % Handle stoich and jac differently
    if useStoich == 0,
        message = sprintf('%s\nThe algebraic relationships are determined using the Jacobian.\nThis is numerically not very stable and you need eventually\nto change tolerance "tol" settings.\n',message);
        % Use the singular value decomposition of the Jacobian to determine
        % the algebraic relationships
        [U,S,V] = svd(J);
        U0 = U(:,end-d+1:end);
        if tol == 0,
            factors = rref(U0');
        else
            factors = rref(U0',tol);
        end
    else
        message = sprintf('%s\nThe algebraic relationships are determined using the Stoichiometric matrix.\n',message);
        % Use rref of the Stoichiometric Matrix
       factors = X(end-d+1:end,size(N,2)+1:end);
    end
    constantValues = factors*initialCondition;
    % determine the indices of the dependent variables. this is simple. 
    % just take the index of the first non zero element in each row as 
    % index for dependent a variable.
    depVar = [];
    for k = 1:size(factors,1),
        nonZeroIndices = find(factors(k,:)~=0);
        depVar = [depVar nonZeroIndices(1)];
    end
    % now do the solving for the steady-state
    OPTIONS = [];
    OPTIONS.MaxIter = MaxIter;
    OPTIONS.TolFun = TolFun;
    OPTIONS.Delta = Delta;  
    [ssVarInDep,FVAL,EXITFLAG] = fsolveIQM(@modelIndependentVariables,getIndependentVariables(initialCondition),OPTIONS);
    % Obtain the full state vector from the independent variables
    steadystate = getAllVariables(ssVarInDep);
end
% If fsolve has not converged to a solution then return an empty vector
if EXITFLAG ~= 1,
    steadystate = [];
end
return

%%%%%%%%%%%%%%%%%%%%%%%%%%%%%%%%%%%%%%%%%%%%
% REMOVE THE TIME FROM THE INPUT ARGUMENTS
%%%%%%%%%%%%%%%%%%%%%%%%%%%%%%%%%%%%%%%%%%%%
function [xdot] = model(x)
    global ODEfctname 
    xdot = feval(ODEfctname,0,x);
return

%%%%%%%%%%%%%%%%%%%%%%%%%%%%%%%%%%%%%%%%%%%%
% CALL MODEL BASED ON INDEPENDENT VARIABLES
%%%%%%%%%%%%%%%%%%%%%%%%%%%%%%%%%%%%%%%%%%%%
function [xdotVarInDep] = modelIndependentVariables(varInDep)
    global ODEfctname 
    variables = getAllVariables(varInDep);
    xdot = feval(ODEfctname,0,variables);
    xdotVarInDep = getIndependentVariables(xdot);
return

%%%%%%%%%%%%%%%%%%%%%%%%%%%%%%%%%%%%%%%%%%%%
% GET INDEPENDENT VARIABLES
%%%%%%%%%%%%%%%%%%%%%%%%%%%%%%%%%%%%%%%%%%%%
function [varInDep] = getIndependentVariables(variables)
    global depVar
    varInDep = variables(setxor([1:length(variables)],depVar));
return

%%%%%%%%%%%%%%%%%%%%%%%%%%%%%%%%%%%%%%%%%%%%
% GET ALL VARIABLES FROM INDEPENDENT
%%%%%%%%%%%%%%%%%%%%%%%%%%%%%%%%%%%%%%%%%%%%
function [variables] = getAllVariables(varInDep)
    global depVar factors constantValues
    n = size(factors,2);
    m = length(depVar);
    varInDepIndex = setxor(1:n,depVar);
    A = factors(1:m,depVar);
    B = constantValues - factors(1:m,varInDepIndex)*varInDep;
    depVarValues = linsolve(A,B);
    variables = zeros(n,1);
    variables(varInDepIndex) = varInDep;
    variables(depVar) = depVarValues;
return      
```
