## Supplementary material for "A systems biology analysis of adrenergically stimulated adiponectin exocytosis in white adipocytes": All scripts used in this work: IQMstoichiometry.m

```
function [N,componentNames,reactionNames,reversibleFlag] = IQMstoichiometry(model,varargin)
% IQMstoichiometry
% Determines the stoichiometric matrix for the given model. In order to be
% able to do that the differential equations for the components have to be
% expressed in terms of reaction rates. The stoichiometric constants need 
% to be numeric and multiplied to the reaction terms.
%
% Examples: d/dt(A) = -1*Re1+2*Re3-Re5+3.141*Re8
%
% Especially when importing models from SBML the right hand side of the
% ODEs might show a correction term that is needed for transport between two
% different compartments in the case that the species is defined in
% concentration units. In this case the ODE can look as follows:
%
%           d/dt(B): (-1*Re1+2*Re3-Re5+3.141*Re8)/compartmentsize
%
% This syntax is also accepted. In this case the stoichiometric elements
% in the parenthesis will be divided by 'compartmentsize'.
%
% The 'compartmentsize' is only allowed to be a parameter! It can not be a 
% numeric value, a state, a variable, or a function!
%
% Except the above shown pair of parentheses, no additional parentheses are
% allowed to appear in the ODE definitions. Otherwise, and in case that not
% only reaction rates are present in the ODE expression, the corresponding
% component is not taken into account for the stoichiometric matrix.
%
% USAGE:
% ======
% [N,componentNames,reactionNames,reversibleFlag] = IQMstoichiometry(model)
%
% model: IQMmodel to determine the stoichiometric matrix for
%
% DEFAULT VALUES:
% ===============
%
% Output Arguments:
% =================
% N: stoichiometric matrix
% componentNames: cell-array with the names of the components present in
%   the stoichiometric matrix
% reactionNames: cell-array with the names of the reactions present in the
%   stoichiometric matrix
% reversibleFlag: vector with same number of entries as reactions. An entry
%   of 0 indicates an irreversible reaction, an entry of 1 indicates a
%   reversible reaction. The ordering is the same as the reaction names.

% <<<COPYRIGHTSTATEMENT - IQM TOOLS LITE>>>

%%%%%%%%%%%%%%%%%%%%%%%%%%%%%%%%%%%%%%%%%%%%%%%
% CHECK IF IQMmodel
%%%%%%%%%%%%%%%%%%%%%%%%%%%%%%%%%%%%%%%%%%%%%%%
if ~strcmp('IQMmodel',class(model)),
    error('Function only defined for IQMmodels.');
end
% get the datastructure of the model
modelstruct = IQMstruct(model);

%%%%%%%%%%%%%%%%%%%%%%%%%%%%%%%%%%%%%%%%%%%%%%%
% CHECK EXTRA INPUT ARGUMENT
%%%%%%%%%%%%%%%%%%%%%%%%%%%%%%%%%%%%%%%%%%%%%%%
% the first extra input argument is used to force the IQMstoichiometry function 
% not to correct the elements of the stoichiometric matrix for the
% compartment information in cases where species are given in
% concentrations. This is needed for model construction (BC type of
% representation and for other things)
% the second is used for silent use of the function
% both flags are not further documented and should not be used by others
% unless they fully understand their meaning

rawFlag = 0;
silentFlag = 0;
if nargin == 2,
    rawFlag = varargin{1};
elseif nargin == 3,
    rawFlag = varargin{1};
    silentFlag = varargin{2};
end

%%%%%%%%%%%%%%%%%%%%%%%%%%%%%%%%%%%%%%%%%%%%%%%
% Get all reaction names
%%%%%%%%%%%%%%%%%%%%%%%%%%%%%%%%%%%%%%%%%%%%%%%
[reactionNames, dummy, reversibleFlag]  = IQMreactions(model);
if isempty(reactionNames),
    if silentFlag == 0,
        error('No reactions present in the model');
    else
        N = [];
        componentNames = {};
        reactionNames = {};
        reversibleFlag = [];
        return
    end
end

%%%%%%%%%%%%%%%%%%%%%%%%%%%%%%%%%%%%%%%%%%%%%%%
% Go trough all ODEs and check reactions ...
%%%%%%%%%%%%%%%%%%%%%%%%%%%%%%%%%%%%%%%%%%%%%%%
% go through all ODEs and check for which components the ODEs only
% consist of reaction terms - this returns already positive and negative
% terms and coefficients
componentNames = {};
[parameterNames, parameterValues] = IQMparameters(model);
N = [];
for k = 1:length(modelstruct.states),
    ODE = modelstruct.states(k).ODE;
    % check if ODE contains only reaction terms - otherwise do not 
    % consider the current state as component for the stoichiometric matrix
    Nrow = getStoichiometryInformation(ODE,reactionNames,parameterNames,parameterValues,rawFlag,silentFlag);
    if ~isempty(Nrow) && isempty(find(isnan(Nrow)==1, 1)),
        N = [N; Nrow];
        componentNames{size(N,1)} = modelstruct.states(k).name;
    end
end
% check for zero rows in N and take them away both from N and from the
% list of component names
%useIndices = find(sum(abs(N')) ~= 0);
%notUseIndices = find(sum(abs(N')) == 0);

nonReactionStates = setdiff(IQMstates(model),componentNames);
if ~isempty(nonReactionStates) && silentFlag == 0,
    disp('For the following components no stoichiometries could be determined.');
    disp('This is due to the syntax of the corresponding ODEs.');
    text = '';
    for k = 1:length(nonReactionStates),
        text = sprintf('%s %s\n',text,nonReactionStates{k});
    end
    disp(text);
end
%componentNames = componentNames(useIndices);
%N = N(useIndices,:);
return

%%%%%%%%%%%%%%%%%%%%%%%%%%%%%%%%%%%%%%%%%%%%%%%
% CHECK ODEs for reactions
%%%%%%%%%%%%%%%%%%%%%%%%%%%%%%%%%%%%%%%%%%%%%%%
% check if given ODE expression contains only reaction terms - then return
% the stoichiometry for this ODE. Otherwise if not only reactions are
% present, return an empty vector.
% in cases that the reaction rate is adjustet by the compartment volume
% (happens especially in cases of import of SBML models containing species
% with concentration rates and compartment volumes different from one) the
% adjustment term and the needed parentheses should be detected and
% neglected.
function [Nrow] = getStoichiometryInformation(ODE,reactionNames,parameterNames,parameterValues,rawFlag,silentFlag)
errorFlag = 0;
% delete all white spaces from the ODE
ODE = regexprep(ODE,'\s*','');
% first of all check if a scaling by the compartment volume is done.
% in this case the expected syntax is 
% ODE = ("reactionterms")/compartmentvolume
numberOpenParentheses = length(find(ODE == '('));
numberClosedParentheses = length(find(ODE == ')'));
compartmentSize = 1;
if numberOpenParentheses ~= numberClosedParentheses,
    % parentheses need to appear in pairs - otherwise error
    errorFlag = 1;
elseif numberOpenParentheses > 1,
    % if there is more than one pair of parentheses, then error
    errorFlag = 1;
elseif numberOpenParentheses == 0,
    errorFlag = 0;
elseif ODE(1) == '(',
    % if the first character is an open parenthesis then assume that 
    % this is due to a adjustement to compartment sizes
    % cut out the content of the parentheses
    closeParenthesis = find(ODE == ')');
    ODERHS = ODE;
    ODE = ODERHS(2:closeParenthesis-1);
    % check part outside parenthesis
    rest = ODERHS(closeParenthesis+1:end);
    % first character needs to be a '/'
    if rest(1) ~= '/',
        errorFlag = 1;
    else
        rest = rest(2:end);
        % check if this rest corresponds to a parameter name and if yes get
        % its value
        index = strmatchIQM(rest,parameterNames,'exact');
        if ~isempty(index),
            compartmentSize = parameterValues(index);
            errorFlag = 0;
        else
            if silentFlag == 0,
                error('The stoichimetric matrix can not be computed. A compartment size seems not to be given as a parameter.');
            else 
                errorFlag = 1;
            end
        end
    end
else
    % parentheses is expected as first non white space character in
    % reaction string ... therefor an error.
    errorFlag = 1;
end
% continue with the algorithm if no error occurred
if errorFlag == 0,
    
    ODE = char([double(ODE) double('+')]);  % fastest strcat
    % first explode the ODE in additive terms
    terms = [];
    termIndex = 1;
    % check the sign of the first term (first character in string)
    if ODE(1) == '-',
        signCurrent = -1;
        lastIndex = 2;
    elseif ODE(1) == '+',
        signCurrent = +1;
        lastIndex = 2;
    else
        signCurrent = +1;
        lastIndex = 1;
    end
    % explode in terms, check if term has the right format and if the
    % second term features a reactionname. then construct the row of the
    % stoichiometric matrix
    Nrow = zeros(1,length(reactionNames));
    startk = lastIndex;
    for k = startk:length(ODE),
        % check for positive term
        if ODE(k) == '+' || ODE(k) == '-',
            element = ODE(lastIndex:k-1);
            % check the element if composed of term in the right format
            multIndex = find(element == '*');
            if length(multIndex) == 0,
                stoichiometry = signCurrent*1;
                reactionterm = element;
            elseif length(multIndex) == 1,
                absStoichiometry = str2double(element(1:multIndex-1));
                if isempty(absStoichiometry),
                    errorFlag = 1;
                    break;
                end
                stoichiometry = signCurrent*absStoichiometry;
                reactionterm = element(multIndex+1:end);
            else
                % to many multiplication signs (only one allowed)
                errorFlag = 1;
                break;
            end
            % find the index of the reaction name and add the
            % stoichiometric information to Nrow
            indexReaction = strmatchIQM(reactionterm,reactionNames,'exact');
            if isempty(indexReaction),
                errorFlag = 1;
                break;
            end
            Nrow(indexReaction) = Nrow(indexReaction) + stoichiometry;
            % increment
            termIndex = termIndex + 1;
            lastIndex = k+1;
            if ODE(k) == '+',
                signCurrent = +1;
            else
                signCurrent = -1;
            end
        end
    end
end
% if an error occurred for the current ODE the Nrow is set to zero
if errorFlag == 1,
    Nrow = [];
end
% adjust stoichiometries with compartment size
if rawFlag == 0,
    Nrow = Nrow / compartmentSize;
end
return
```
