## Supplementary material for "A systems biology analysis of adrenergically stimulated adiponectin exocytosis in white adipocytes": All scripts used in this work: IQMprepredreac.m

```
function [output] = IQMprepredreac(model,timevectors,experiments,varargin)
% IQMprepredreac: prepares a model for subsequent reduction of reaction rate
% expressions. Eventual "power(x,y)" expressions are exchanged against
% "x^y". Reaction rates are expanded by substituting in variables. Ununsed
% elements (reactions, variables, and parameters) are deleted from the
% model. Furthermore reference simulations are performed based on which the
% models reactions can be reduced in following steps.
%
% USAGE:
% ======
% [output] = IQMprepredreac(model, timevectors, experiments)
% [output] = IQMprepredreac(model, timevectors, experiments, extravariables)
%
% model:            IQMmodel to consider for reduction
% timevectors:      time vectors of interest (one for each experiment)
% experiments:      cell-array with experiment definitions 
% extravariables:   cellarray with names of parameters that should be taken
%                   into account as species belonging to the M vector
%                   or parameters that are set to different values during
%                   experiments (then they need to be kept in the reduced model)
%
% Output Arguments:
% =================
% output:   structure with necessary information

% <<<COPYRIGHTSTATEMENT - IQM TOOLS LITE>>>

%%%%%%%%%%%%%%%%%%%%%%%%%%%%%%%%%%%%%%%%%%%%%%%
% HANDLE NON-NUMERIC INITIAL CONDITIONS
%%%%%%%%%%%%%%%%%%%%%%%%%%%%%%%%%%%%%%%%%%%%%%%
% just by replacing them
model = IQMconvertNonNum2NumIC(model);
if ~hasonlynumericICsIQM(model),
    disp('Non-numeric initial conditions will be removed in the reduced model. You can replace them after reduction if you like.');
end
    
output.model = [];
output.extravariables = [];
output.timevectors = timevectors;
output.reference = [];
output.reference.experiments = experiments;
output.reference.data = [];
output.reference.datavalues = [];

%%%%%%%%%%%%%%%%%%%%%%%%%%%%%%%%%%%%%%%%%%%%%%%
% check if symbolic toolbox is present
%%%%%%%%%%%%%%%%%%%%%%%%%%%%%%%%%%%%%%%%%%%%%%%
if ~isSymbolicpresentIQM,
    error('The model reduction feature requires the presence of the symbolic toolbox.');
end

%%%%%%%%%%%%%%%%%%%%%%%%%%%%%%%%%%%%%%%%%%%%%%%
% check cells
%%%%%%%%%%%%%%%%%%%%%%%%%%%%%%%%%%%%%%%%%%%%%%%
if ~iscell(timevectors),
    timevectors = {timevectors};
end
if ~iscell(experiments),
    experiments = {experiments};
end

%%%%%%%%%%%%%%%%%%%%%%%%%%%%%%%%%%%%%%%%%%%%%%%
% variable input arguments
%%%%%%%%%%%%%%%%%%%%%%%%%%%%%%%%%%%%%%%%%%%%%%%
extravariables = {};
if nargin > 3,
    extravariables = varargin{1};
end
output.extravariables = extravariables;

%%%%%%%%%%%%%%%%%%%%%%%%%%%%%%%%%%%%%%%%%%%%%%%
% check if first argument is a model
%%%%%%%%%%%%%%%%%%%%%%%%%%%%%%%%%%%%%%%%%%%%%%%
if ~isIQMmodel(model),
    error('First argument needs to be an IQMmodel.');
end

%%%%%%%%%%%%%%%%%%%%%%%%%%%%%%%%%%%%%%%%%%%%%%%
% expand reaction expressions by substituting in all variables
%%%%%%%%%%%%%%%%%%%%%%%%%%%%%%%%%%%%%%%%%%%%%%%
model = substitutevarsIQM(model);

%%%%%%%%%%%%%%%%%%%%%%%%%%%%%%%%%%%%%%%%%%%%%%%
% check and replace power expressions
%%%%%%%%%%%%%%%%%%%%%%%%%%%%%%%%%%%%%%%%%%%%%%%
[model,changeFlag] = depowerIQM(model);
% if changeFlag == 1,
% %    disp('Power operators are present. Support for Hill-type rate equations is only experimental at present.');
%     model = fixExp(model);
% end

%%%%%%%%%%%%%%%%%%%%%%%%%%%%%%%%%%%%%%%%%%%%%%%
% finally clean the model from unnecessary parameters, reactions, variables
%%%%%%%%%%%%%%%%%%%%%%%%%%%%%%%%%%%%%%%%%%%%%%%
output.model = cleanmodelIQM(model,1);

%%%%%%%%%%%%%%%%%%%%%%%%%%%%%%%%%%%%%%%%%%%%%%%%%%%%%%%%%%%%%%%%%%%%%%%%%%%
% simulate the experiments and save the data
%%%%%%%%%%%%%%%%%%%%%%%%%%%%%%%%%%%%%%%%%%%%%%%%%%%%%%%%%%%%%%%%%%%%%%%%%%%
% simulate all the experiments and collect the numeric values for states
% and reactions and additionally also the parameters (extravariables)
states = IQMstates(output.model)';
reactions = IQMreactions(output.model)';
parameters = IQMparameters(output.model);
statevalues = [];
reactionvalues = [];
parametervalues = [];
for k = 1:length(experiments),
    disp(sprintf('Simulating experiment %d/%d ...',k,length(experiments)));
    modelexp = IQMmergemodexp(output.model,experiments{k});
    % set maxstep options
    options.maxstep = max(output.timevectors{k})/length(output.timevectors{k});
    options.maxnumsteps = 10000;
    if isIQMproPresent(),
        simdata = IQMPsimulate(modelexp,timevectors{k},[],[],[],options);
    else
        simdata = IQMsimulate(modelexp,timevectors{k});
    end
    statevalues = [statevalues; simdata.statevalues];
    reactionvalues = [reactionvalues; simdata.reactionvalues];
    % get values for the extra variables that are parameters (can change
    % during experiments)
    parametervaluesscalar = IQMparameters(modelexp,extravariables)';
    parametervalues = [parametervalues; parametervaluesscalar(ones(length(timevectors{k}),1),:)];
end
% add states extravariables and reactions together
output.reference.data = {states{:} extravariables{:} reactions{:}};
output.reference.datavalues = [statevalues parametervalues reactionvalues];
return

% %%%%%%%%%%%%%%%%%%%%%%%%%%%%%%%%%%%%%%%%%%%%%%%%%%%%%%%%%%%%%%%%%%%%%%%%%%%
% % fixExp: for all exponential parameters, eg x^p1, fix them to the value
% % specified in the model => x^4 if p1=4. Do this ONLY for the reaction 
% % expressions.
% %%%%%%%%%%%%%%%%%%%%%%%%%%%%%%%%%%%%%%%%%%%%%%%%%%%%%%%%%%%%%%%%%%%%%%%%%%%
% function model = fixExp(model)
% global parameterNames parameterValues
% [parameterNames, parameterValues] = IQMparameters(model);
% iqms = struct(model);
% for k = 1:length(iqms.reactions),
%     iqms.reactions(k).formula = fixHelper(iqms.reactions(k).formula);
% end
% model = IQMmodel(iqms);    
% return
% 
% %%%%%%%%%%%%%%%%%%%%%%%%%%%%%%%%%%%%%%%%%%%%%%%
% % fixHelper: subfunction to fixExp
% %%%%%%%%%%%%%%%%%%%%%%%%%%%%%%%%%%%%%%%%%%%%%%%
% function finalString = fixHelper(string)
% global parameterNames parameterValues;
% startIndex = strfind(string, '^');
% finalString = string; index = 0;
% if ~isempty(startIndex)
%     finalString = string(1:startIndex(1)-1);
%     for i = 1:length(startIndex)
%         openParenthesis = 0;
%         for j = startIndex(i)+1:length(string)
%             if string(j) == '(',
%                 openParenthesis = openParenthesis + 1;
%             elseif string(j) == ')',
%                 openParenthesis = openParenthesis - 1;
%             end
%             if openParenthesis == 0
%                 if i == length(startIndex)
%                     if index < i
%                         selectedString = string([startIndex(i)+2:j-1]);
%                         match = findsym(sym(selectedString));
%                         [found, ia, ib] = intersect(parameterNames, match);
%                         for k = 1:length(found)
%                             selectedString = num2str(subs(selectedString, parameterNames(ia(k)), parameterValues(ia(k))));
%                         end
%                         finalString = strcat(finalString, '^(', selectedString, ')',string([j+1:end]));
%                         break;
%                     end
%                 elseif j < startIndex(i+1)
%                     if index < i
%                         selectedString = string([startIndex(i)+2:j-1]);
%                         match = findsym(sym(selectedString));
%                         [found, ia, ib] = intersect(parameterNames, match);
%                         for k = 1:length(found)
%                             selectedString = num2str(subs(selectedString, parameterNames(ia(k)), parameterValues(ia(k))));
%                         end
%                         finalString = strcat(finalString, '^(', selectedString, ')', string([j+1:startIndex(i+1)-1]));
%                         break;
%                     end
%                 else
%                     index = find(startIndex > j, 1);
%                     if isempty(index)
%                         index = length(startIndex)+1;
%                     end
%                     
%                     rep = fixHelper(string([startIndex(i)+2:j-1]));
%                     match = regexp(rep, '(.+)[,](.+)', 'tokens');
%                     finalString = strcat(finalString, '^(', rep, ')');
%                     break;
%                 end
%             end
%         end
%     end
% end
% return
```
