## Supplementary material for "A systems biology analysis of adrenergically stimulated adiponectin exocytosis in white adipocytes": All scripts used in this work: IQMredallreac.m

```
function [modelred] = IQMredallreac(model,experiments,timevectors,varargin)
% IQMredallreac: Wrapper for the IQMredreac function allowing to easily
% go through a whole model and reduce all reactions that are possible to
% reduce.
% 
% USAGE:
% ======
% [modelred] = IQMredallreac(model,experiments,timevectors)
% [modelred] = IQMredallreac(model,experiments,timevectors,OPTIONS)
% [modelred] = IQMredallreac(model,experiments,timevectors,OPTIONS,extravariables)
%
% model:            IQMmodel to consider for reduction
% timevectors:      cell-array with time vectors of interest (one for each experiment)
% experiments:      cell-array with experiment definitions 
% extravariables:   cellarray with names of parameters that should be taken
%                   into account as species belonging to the M vector
%                   or parameters that are set to different values during
%                   experiments (then they need to be kept in the reduced model)
% OPTIONS: structure containing options for the algorithm:
%        OPTIONS.tol: tolerance for singularity detection (smallest SV)
%        OPTIONS.keeporigparameters: =0: do not keep original parameters
%                                    =2: do always keep original parameters
%                                    =1: keep original parameters only if
%                                        it leads to fewer parameters
%        OPTIONS.numeratorweighting: =1: weight numerator terms and denumerator terms 
%                                    such that numerators are kept
%                                    =0: dont do any weighting
%
% DEFAULT VALUES:
% ===============
% extravariables:               {}
% OPTIONS.tol:                  1e-6
% OPTIONS.keeporigparameters:   0
% OPTIONS.numeratorweighting:   0
%
% Output Arguments:
% =================
% modelred: IQMmodel in which the defined reaction is reduced

% <<<COPYRIGHTSTATEMENT - IQM TOOLS LITE>>>

%%%%%%%%%%%%%%%%%%%%%%%%%%%%%%%%%%%%%%%%%%%%%%%
% check cell-arrays in input
%%%%%%%%%%%%%%%%%%%%%%%%%%%%%%%%%%%%%%%%%%%%%%%
if ~iscell(experiments),
    experiments = {experiments};
end
if ~iscell(timevectors),
    timevectors = {timevectors};
end

%%%%%%%%%%%%%%%%%%%%%%%%%%%%%%%%%%%%%%%%%%%%%%%
% variable input arguments
%%%%%%%%%%%%%%%%%%%%%%%%%%%%%%%%%%%%%%%%%%%%%%%
OPTIONS = [];
extravariables = {};
if nargin >= 4,
    OPTIONS = varargin{1};
end
if nargin >= 5,
    extravariables = varargin{2};
end

%%%%%%%%%%%%%%%%%%%%%%%%%%%%%%%%%%%%%%%%%%%%%%%
% prepare reduction - DOES ALSO HANDLE NON-NUMERIC ICs ... by converting
% them to NUMERIC ICs
%%%%%%%%%%%%%%%%%%%%%%%%%%%%%%%%%%%%%%%%%%%%%%%
disp('Preparing data for reduction ...');
output = IQMprepredreac(model,timevectors,experiments,extravariables);
[reactions, formulas] = IQMreactions(output.model);

%%%%%%%%%%%%%%%%%%%%%%%%%%%%%%%%%%%%%%%%%%%%%%%
% Go through all reactions
%%%%%%%%%%%%%%%%%%%%%%%%%%%%%%%%%%%%%%%%%%%%%%%
disp('Going through all reactions ...');
modelred = output.model;
for k = 1:length(reactions),
    clc;
    reaction = reactions{k};
    formula = formulas{k};
    text = sprintf('%s = %s\n',reaction,formula);
    disp(text);
    while 1,
        choose = input('Reduce this reaction (1) or not (0)? ');
        if choose == 1 || choose == 0,
            break;
        end
    end
    if choose == 1,
        % run
        modelsave = modelred;
        while 1,
            try
                [modelred] = IQMredreac(output,reaction,OPTIONS);
                output.model = modelred;
            catch
                disp(lasterr);
            end
            try
                runcomparison(model,modelred,experiments,timevectors);
            catch
                disp(lasterr);
                disp('Something went wrong ... you should press 0!');
            end
            while 1,
                check = input('Does this look ok (1) or not (0)?');
                if check == 1 || check == 0,
                    break;
                end
            end
            if check == 0,
                output.model = modelsave;
                modelred = modelsave;
            else
                break;
            end
        end
    end
end
return

%%%%%%%%%%%%%%%%%%%%%%%%%%%%%%%%%%%%%%%%%%%%%%%
% run comparison with original model
%%%%%%%%%%%%%%%%%%%%%%%%%%%%%%%%%%%%%%%%%%%%%%%
function [] = runcomparison(model,modelred,experiments,timevectors)
figure(1); clf;
for k = 1:length(experiments),
    subplot(length(experiments),1,k);
    % perform experiment
    modexp = IQMmergemodexp(model,experiments{k});
    % set maxstep options
    options.maxstep = max(timevectors{k})/length(timevectors{k});
    options.maxnumsteps = 10000;
    if isIQMproPresent(),
        expdata = IQMPsimulate(modexp,timevectors{k},[],[],[],options);
    else
        expdata = IQMsimulate(modexp,timevectors{k});
    end
    modredexp = IQMmergemodexp(modelred,experiments{k});
    if isIQMproPresent(),
        redexpdata = IQMPsimulate(modredexp,timevectors{k},[],[],[],options);
    else
        redexpdata = IQMsimulate(modredexp,timevectors{k});
    end
    % plot results
    plot(expdata.time,expdata.statevalues); hold on; 
    plot(redexpdata.time,redexpdata.statevalues,'o');
    % legend
    if k == 1,
        hlhlx = legend(IQMstates(model),'Location','NorthEastOutside');
        set(hlhlx,'Interpreter','none');
    end
    title(sprintf('Comparison for Experiment %d',k));

    % At this point the user can add whatever comparison test between the
    % original and reduced model that is necessary ...

end
return
```
