## Supplementary material for "A systems biology analysis of adrenergically stimulated adiponectin exocytosis in white adipocytes": All scripts used in this work: IQMredreac.m

```
function [modelred] = IQMredreac(output,reaction,varargin)
% IQMredreac: Reduction of single reaction expressions with the goal of reducing
% complexity and number of parameters. This function requires the prior
% execution of the IQMprepredreac.
% 
% USAGE:
% ======
% [modelred] = IQMredreac(output,reaction)
% [modelred] = IQMredreac(output,reaction,OPTIONS)
%
% output: Result returned from IQMprepredreac
% reaction: Name of the reaction that should be reduced
% OPTIONS: structure containing options for the algorithm:
%        OPTIONS.tol: tolerance for singularity detection (smallest SV)
%        OPTIONS.keeporigparameters: =0: do not keep original parameters
%                                    =2: do always keep original parameters
%                                    =1: keep original parameters only if
%                                        it leads to fewer parameters
%        OPTIONS.numeratorweighting: =1: weight numerator terms and denumerator terms 
%                                    such that numerators are kept
%                                    =0: dont do any weighting
%
% DEFAULT VALUES:
% ===============
% OPTIONS.tol:                  1e-6
% OPTIONS.keeporigparameters:   0
% OPTIONS.numeratorweighting:   0
%
% Output Arguments:
% =================
% modelred: IQMmodel in which the defined reaction is reduced

% <<<COPYRIGHTSTATEMENT - IQM TOOLS LITE>>>

% open needed figure that is to be updates during the whole reduction
% procedure of the given reaction
figH2 = figure(2); clf;
yaxismax = 0;
xplot = -1;
yplot = -1;

%%%%%%%%%%%%%%%%%%%%%%%%%%%%%%%%%%%%%%%%%%%%%%%
% check if symbolic toolbox is present
%%%%%%%%%%%%%%%%%%%%%%%%%%%%%%%%%%%%%%%%%%%%%%%
if ~isSymbolicpresentIQM,
    error('The model reduction feature requires the presence of the symbolic toolbox.');
end

%%%%%%%%%%%%%%%%%%%%%%%%%%%%%%%%%%%%%
% Handle variable input arguments (options)
%%%%%%%%%%%%%%%%%%%%%%%%%%%%%%%%%%%%%
OPTIONS = [];
if nargin == 3,
    OPTIONS = varargin{1};
end
% default values
tol = 1e-6;              % tolerance for singularity detection (smalles SV)
keeporigparameters = 0;  % do not keep original parameters
numeratorweighting = 0;  % do not weight numerators/denumerators depending on their number
% tol
if isfield(OPTIONS,'tol'),
    if ~isempty(OPTIONS.tol),
        tol = OPTIONS.tol;
    end
end
% keeporigparameters
if isfield(OPTIONS,'keeporigparameters'),
    if ~isempty(OPTIONS.keeporigparameters),
        keeporigparameters = OPTIONS.keeporigparameters;
    end
end
% numeratorweighting
if isfield(OPTIONS,'numeratorweighting'),
    if ~isempty(OPTIONS.numeratorweighting),
        numeratorweighting = OPTIONS.numeratorweighting;
    end
end

disp(' ');
disp('#####################################################################');
disp(sprintf('# Reduction of reaction:  %s', reaction));
disp('#####################################################################');


%%%%%%%%%%%%%%%%%%%%%%%%%%%%%%%%%%%%%
% Determine reduction information for specified reaction
%%%%%%%%%%%%%%%%%%%%%%%%%%%%%%%%%%%%%
output2 = reducereactionprepIQM(output,reaction);

%%%%%%%%%%%%%%%%%%%%%%%%%%%%%%%%%%%%%
% Determine which terms to reduce
%%%%%%%%%%%%%%%%%%%%%%%%%%%%%%%%%%%%%
% number terms in original equation
nrorig = size(output2.reaction_trans.A,2);
indexkeep = [1:nrorig];
indexkeep_backup = {};
while 1,
    try
        % Reduce (do this also for the case where indexkeep contains all terms)
        output3 = reducereacIQM(output2,indexkeep,keeporigparameters);
    catch
        % catch errors that can occurr in cases when no reduction is possible
        % example: reaction only defined by constant rate
        %          ...
        
        disp('************************** All nominators might have been cancelled ... try nominator weighting!');
    end
    % Optimize parameter values
    output4 = reduceoptimIQM(output3);
    % check if optimization successful (do it by looking at it!)
    orig = output3.reaction_orig.reactionvalues;
    redopt = output4.reaction_opt.reactionvalues;
    figH = figure(1); clf; 
    plot(orig); hold on; plot(redopt,'r--'); 
    
    
    
    % also plot the obtained costfunction value against the ratio between the 
    % number of parameters in the reduced and original model. Divide the
    % cost by the mean
    nrorigterms = length(output4.reaction_trans.c);
    nrredterms = length(indexkeep);
    costred_scaled = output4.reaction_opt.cost/mean(orig);
    figure(figH2);
    xoldplot = xplot;
    yoldplot = yplot;
    xplot = nrredterms/nrorigterms;
    yplot = costred_scaled;
    plot(xplot,yplot,'r*'); hold on;
    if xoldplot >= 0,
        plot(xoldplot,yoldplot,'w*'); hold on;
        plot(xoldplot,yoldplot,'bo'); hold on;
    end
    yaxismax = max(yaxismax, yplot*1.1);
    axis([0 1 0 yaxismax]);    
    xlabel('#redterms/#origterms');
    ylabel('optimal cost');
    
    
    
    % plot some info
disp(' ');
if length(indexkeep) > 1,
    disp('Linear dependency and approximation measures');
    disp('############################################');
    [Us,Ss,Vs] = svd(output3.reductioninfo.Anumscaled(:,indexkeep));
    disp(sprintf('Singular values Mscaled: %s', sprintf('%g ',diag(Ss))));
    disp(sprintf('Elements v*(Mscaled): %s',sprintf('%g ',Vs(:,end))));
    disp(' ');
    [Uc,Sc,Vc] = svd(output3.reductioninfo.AnumC(:,indexkeep));
    disp(sprintf('Singular values Mc: %s', sprintf('%g ',diag(Sc))));
    disp(sprintf('Elements v*(Mc): %s',sprintf('%g ',Vc(:,end))));
    disp(' ');
end
disp('Statistics and preliminary result ');
disp('#################################');
    text = sprintf('Nr parameters in original/reduced expression: %d/%d\n',length(output2.reaction_orig.parameters),length(output3.reaction_red.parameters));
    indexdelete = setdiff([1:nrorig],indexkeep);
    deleteText = '';
    for k = 1:length(indexdelete),
        deleteText = strcat(deleteText, ', ', output3.reaction_trans.A{indexdelete(k)});
    end
    text = sprintf('%sDeleted terms: %s\n',text,deleteText(2:end));
    text = sprintf('%s%s_red = %s\n',text,output2.reaction,output3.reaction_red.formula);
    disp(text);
    % Decide what to do next
    if length(indexkeep_backup) == 0,
        while 1,
            check = input('Continue (2), or end (0) (switch num weighting: 99): ');
            if check == 2 || check == 0,
                break;
            elseif check == 99,
                numeratorweighting = ~numeratorweighting;
                disp(sprintf('numeratorweighting = %d',numeratorweighting));
            end
        end
    else
        while 1,
            check = input('Continue (2), go back (1), end (0) (switch num weighting: 99): ');
            if check == 2 || check == 1 || check == 0,
                break;
            elseif check == 99,
                numeratorweighting = ~numeratorweighting;
                disp(sprintf('numeratorweighting = %d',numeratorweighting));
            end
        end
    end
    if check == 1,
        % go a step back
        indexkeep = indexkeep_backup{end};
        indexkeep_backup = indexkeep_backup(1:end-1);
    elseif check == 0,
        % end reduction
        break;
    else
        % continue
        indexkeep_backup{end+1} = indexkeep;
        % Select what to reduce
        Anumscaled = output2.reductioninfo.Anumscaled(:,indexkeep);
        AnumC = output2.reductioninfo.AnumC(:,indexkeep);
        [U,S,V] = svd(Anumscaled);
        SAns = diag(S)';
        VAns = V(:,end);
        [U,S,V] = svd(AnumC);
        SAnc = diag(S)';
        VAnc = V(:,end);
        V1 = abs(VAns);
        V2 = abs(VAnc);
        % do numeratorweighting (if on then set numerator elements to 0),
        % avoidng the deletion of numerators
        if numeratorweighting == 1,
            isnomterm = output2.reductioninfo.isnomterm(indexkeep);
            numbernom = sum(isnomterm);
            numberden = length(isnomterm)-numbernom;
            factor = [ones(numberden,1); 0*ones(numbernom,1)];
            V1 = V1.*factor;
            V2 = V2.*factor;
        end
        V1 = max(V1-mean(V1),0);
        V2 = max(V2-mean(V2),0);
        if SAns(end) < tol,
            disp('##########################################');
            disp('# Reduction:  Based on linear dependency #');
            disp('##########################################');
            V = V1+0.01*V2;
identFlag = 1;
        else
            disp('####################################');
            disp('# Reduction based on approximation #');
            disp('####################################');
            V = V2;
identFlag = 0;
        end

if identFlag == 0,
    [dummy,indexdeleteNEXT] = max(V);
else
% determine the k that minimizes the relative changes
    c = output4.reaction_red.parametervalues;
    testValues = zeros(1,length(c));
    for k4=1:length(c),
        ssum = 0;
        if V(k4) ~= 0,
            for i4=1:length(V),
                if k4~=i4,
                    ssum = ssum + (abs(c(k4)*V(i4)/V(k4)/c(i4)));
                end
            end
            testValues(k4) = ssum;
        else
            testValues(k4) = inf;
        end
    end
    [dummy, indexdeleteNEXT] = min(testValues);
end
       
        indexkeep(indexdeleteNEXT) = [];   % delete element
    end
end

%%%%%%%%%%%%%%%%%%%%%%%%%%%%%%%%%%%%%
% Update the model with the reduced reaction
%%%%%%%%%%%%%%%%%%%%%%%%%%%%%%%%%%%%%
modelred= redupdateIQM(output4);
```
