## Supplementary material for "A systems biology analysis of adrenergically stimulated adiponectin exocytosis in white adipocytes": All scripts used in this work: IQMinsilicoexp.m

```
function [varargout] = IQMinsilicoexp(model,experiment,timevector,varargin)
% IQMinsilicoexp runs an in-silico experiment. Simulated data can be
% returned in the IQMmeasurement format, plotted, or written in a CSV
% or Excel measurement file (the latter only under Windows).
%
% USAGE:
% ======
% [] = IQMinsilicoexp(model,experiment,timevector [,OPTIONS])         
% [] = IQMinsilicoexp(model,experiment,timevector,measurements [,OPTIONS])         
% [] = IQMinsilicoexp(model,experiment,timevector,measurements,filetypeFlag [,OPTIONS])         
% [] = IQMinsilicoexp(model,experiment,timevector,measurements,filetypeFlag,filename [,OPTIONS])         
% [output] = IQMinsilicoexp(model,experiment,timevector [,OPTIONS])         
% [output] = IQMinsilicoexp(model,experiment,timevector,measurements [,OPTIONS])         
%
% model: IQMmodel to perform the insilico experiment on
% experiment: IQMexperiment object to apply
% timevector: Timevector to be used for simulation
% measurements: cell-array with the names of the components to measure
%               (states, variables, reactions)
% filetypeFlag: 0=plot results, 1=CSV measurement file, 2=Excel measurement
%               file
% filename: Name of the measurement file (or of the measurement) to generate
% OPTIONS: structure with integrator options.
%        OPTIONS.abstol: abs tolerance
%        OPTIONS.reltol: rel tolerance
%        OPTIONS.minstep: min step-size of integrator
%        OPTIONS.maxstep: max step-size of integrator
%        OPTIONS.maxnumsteps: maximum number of steps to be
%          taken by the solver in its attempt to reach the next
%          output time 
%
% DEFAULT VALUES:
% ===============
% measurements: all states are measured
% filetypeFlag: 0 (plot the results). If output argument is specified, the 
%               setting of the filetype flag is ignored
% filename: combination of model and experiment name
% OPTIONS.abstol: 1e-6
% OPTIONS.reltol: 1e-6
% OPTIONS.minstep: 0
% OPTIONS.maxstep: inf
% OPTIONS.maxnumsteps: 500
%
% Output Arguments:
% =================
% output: An IQMmeasurement object with the resulting data

% <<<COPYRIGHTSTATEMENT - IQM TOOLS PRO>>>

%%%%%%%%%%%%%%%%%%%%%%%%%%%%%%%%%
% Check input arguments 
%%%%%%%%%%%%%%%%%%%%%%%%%%%%%%%%%
if ~isIQMmodel(model),
    error('The first input argument needs to be an IQMmodel.');
end
if ~isIQMexperiment(experiment),
    error('The first input argument needs to be an IQMexperiment.');
end
if length(timevector) <= 1,
    error('The timevector is set incorrectly.');
end
modelstruct = struct(model);
experimentstruct = struct(experiment);

%%%%%%%%%%%%%%%%%%%%%%%%%%%%%%%%%
% Handle variable input arguments
%%%%%%%%%%%%%%%%%%%%%%%%%%%%%%%%%
filetypeFlag = 0;
filename = [modelstruct.name '_' experimentstruct.name];
measurements = {};
OPTIONS = [];
if nargin == 4,
    if isstruct(varargin{1}) || (isempty(varargin{1}) && isnumeric(varargin{1})),
        OPTIONS = varargin{1};
    else
        measurements = varargin{1};
    end
elseif nargin == 5,
    measurements = varargin{1};
    if isstruct(varargin{2}) || (isempty(varargin{2}) && isnumeric(varargin{2})),
        OPTIONS = varargin{2};
    else
        filetypeFlag = varargin{2};
    end
elseif nargin == 6,
    measurements = varargin{1};
    filetypeFlag = varargin{2};
    if isstruct(varargin{3}) || (isempty(varargin{3}) && isnumeric(varargin{3})),
        OPTIONS = varargin{3};
    else
        filename = varargin{3};
    end    
elseif nargin == 7.
    measurements = varargin{1};
    filetypeFlag = varargin{2};
    filename = varargin{3};
    OPTIONS = varargin{4};
end

if isempty(measurements),
    measurements = IQMstates(model);
end
if ischar(measurements),
    measurements = {measurements};
end

%%%%%%%%%%%%%%%%%%%%%%%%%%%%%%%%%
% Merge model and experiment
%%%%%%%%%%%%%%%%%%%%%%%%%%%%%%%%%
modexp = IQMmergemodexp(model,experiment);
modelstruct = struct(modexp);

%%%%%%%%%%%%%%%%%%%%%%%%%%%%%%%%%
% Check measurements
%%%%%%%%%%%%%%%%%%%%%%%%%%%%%%%%%
meascomponenttype = [];
meascomponentindex = [];
modelStates = IQMstates(modexp);
modelVariables = IQMvariables(modexp);
modelReactions = IQMreactions(modexp);
for k=1:length(measurements),
    found = 0;
    % check states
    index = strmatchIQM(measurements{k},modelStates,'exact');
    if ~isempty(index),
        found = 1;
    end
    % check variables (if not in states)
    if found == 0,
        index = strmatchIQM(measurements{k},modelVariables,'exact');
        if ~isempty(index),
            found = 2;
        end
    end
    % check reactions (if not in states and variables)
    if found == 0,
        index = strmatchIQM(measurements{k},modelReactions,'exact');
        if ~isempty(index),
            found = 3;
        end
    end
    % check if it exists at all
    if found == 0,
        error('Measured component ''%s'' does not exist in the model.',measurements{k});
    end
    % save type and index
    meascomponenttype(end+1) = found;
    meascomponentindex(end+1) = index;
end

%%%%%%%%%%%%%%%%%%%%%%%%%%%%%%%%%
% Simulate the experiment model (Non-numeric ICs handled by not providing
% an initial condition!)
%%%%%%%%%%%%%%%%%%%%%%%%%%%%%%%%%
simdata = IQMPsimulate(modexp,timevector,[],[],[],OPTIONS);

%%%%%%%%%%%%%%%%%%%%%%%%%%%%%%%%%
% Create IQMmeasurement object
%%%%%%%%%%%%%%%%%%%%%%%%%%%%%%%%%
measstruct = struct(IQMmeasurement);
measstruct.name = filename;
measstruct.notes = sprintf('Insilico experiment measurements\n================================\nModel notes:\n------------\n%s\n\nExperiment notes:\n-----------------\n%s',modelstruct.notes,experimentstruct.notes);
measstruct.time = timevector(:);
for k=1:length(measurements),
    type = meascomponenttype(k);
    index = meascomponentindex(k);
    if type == 1, % its a state
        name = modelstruct.states(index).name;
        notes = modelstruct.states(index).notes;
        values = simdata.statevalues(:,index);
    elseif type == 2, % its a variable
        name = modelstruct.variables(index).name;
        notes = modelstruct.variables(index).notes;
        values = simdata.variablevalues(:,index);
    elseif type == 3, % its a reaction
        name = modelstruct.reactions(index).name;
        notes = modelstruct.reactions(index).notes;
        values = simdata.reactionvalues(:,index);
    end        
    maxvalues = NaN(size(values));
    minvalues = maxvalues;
    measstruct.data(k).name = name;
    measstruct.data(k).notes = notes;
    measstruct.data(k).values = values;
    measstruct.data(k).maxvalues = maxvalues;
    measstruct.data(k).minvalues = minvalues;
end
insilicomeas = IQMmeasurement(measstruct);

%%%%%%%%%%%%%%%%%%%%%%%%%%%%%%%%%
% Handle the output
%%%%%%%%%%%%%%%%%%%%%%%%%%%%%%%%%
if nargout == 0,
    if filetypeFlag == 0,
        % just plot the measurement
        IQMvisualizemeasurement(insilicomeas);
    elseif filetypeFlag == 1 || (filetypeFlag == 2 && ~ispc),
        % export to CSV (also if XLS desired but no windows PC)
        IQMexportCSVmeasurement(insilicomeas,filename);
    elseif filetypeFlag == 2 && ispc,
        % export to XLS
        IQMexportXLSmeasurement(insilicomeas,filename);
    end
elseif nargout == 1,
    varargout{1} = insilicomeas;
end
```
