## Supplementary material for "A systems biology analysis of adrenergically stimulated adiponectin exocytosis in white adipocytes": All scripts used in this work: IQMsimulate.m

```
function [varargout] = IQMsimulate(varargin)
% IQMsimulate: This function is a wrapper function for two different types of simulation:
%
% 1) Simulation of IQMmodels that are automatically converted to MATLAB ODE files
%           [output] = IQMsimulate(model)         
%           [output] = IQMsimulate(model,time)         
%           [output] = IQMsimulate(model,method,time)         
%           [output] = IQMsimulate(model,method,time,ic)         
%           [output] = IQMsimulate(model,method,time,ic,options)         
%
% 2) Simulation of IQMmodels with IQMdosing schemes
%           [output] = IQMsimulate(model,dosing)         
%           [output] = IQMsimulate(model,dosing,time)         
%           [output] = IQMsimulate(model,dosing,time,ICs)         
%           [output] = IQMsimulate(model,dosing,time,ICs,paramnames,paramvalues)         
%           [output] = IQMsimulate(model,dosing,time,ICs,paramnames,paramvalues,OPTIONS)         
%
% Both types are distinguished by the 2nd needing an IQMdosing object as second input argument.
% Full documentation of both types below:s
%
% ============================================
% Documentation of 1st type of use:
% (documentation for 2nd type below)
% ============================================
% Simulates an IQMmodel or an ODE file created by
% IQMcreateODEfile. In case the model is given as an IQMmodel, not only the 
% trajectories for the models states are determined, but also the time 
% trajectories for all variables and reaction rates in the model 
% 
% The IQMsimulate function can deal automatically with models that contain 
% state events. However, this is only possible in the case where a model is
% specified as an IQMmodel, not as an ODE file model. 
%
% USAGE:
% ======
% [output] = IQMsimulate(model)         
% [output] = IQMsimulate(model,time)         
% [output] = IQMsimulate(model,method,time)         
% [output] = IQMsimulate(model,method,time,ic)         
% [output] = IQMsimulate(model,method,time,ic,options)         
%
% model: IQMmodel or ODE file model description
% time: scalar value for simulation end time (default start time is 0),
%       vector with two elements indicating start and end time, or vector
%       with time instants for which the simulation results are to be
%       returned.
% method: name of a MATLAB integrator (ode45, ode23s, etc.) 
% ic: initial conditions of the model to be simulated
% options: standard MATLAB integrator options, set by ODESET
%
% DEFAULT VALUES:
% ===============
% time: 20 time units (=> tspan: [0 20])
% method: 'ode23s' ('ode15s' IF algebraic rules present in the model)
% ic: the initial conditions stored in the model
% options: []
%
% Output Arguments:
% =================
% If no output arguments are given, the result of the simulation is plotted
% 1) online during the simulation for monitoring purposes - the simulation 
% can here be stopped at every instant by just clicking on the "Stop"
% button. 2) after the simulation is finished the simulation data is plotted 
% using the IQMplot function, allowing to browse the data.
%
% The output of IQMsimulate is realized as a structure:
% output.time:              vector with time instants for the results
% output.states:            cell-array with state names
% output.statevalues:       matrix with state values. Each row corresponds to
%                           one time instant.
%
% The following fields are only present in case the model as been given
% as an IQMmodel:
%
% output.variables:         cell-array with variable names
% output.variablevalues:    matrix with variable values. Each row corresponds to
%                           one time instant.
% output.reactions:         cell-array with reaction names
% output.reactionvalues:    matrix with reaction values. Each row corresponds to
%                           one time instant.
%
% ============================================
% Documentation of 2nd type of use:
% ============================================
% IQMsimulatedosing: Simulates the application of a dosing schedule to an IQMmodel with relevant INPUT definitions.
% This is a wrapper for IQMsimdosing, allowing to not merge model and dosing prior to simulation. 
% Disadvantage: Only works on IQMmodels and thus requires recompilation with each simulation (if IQM Tools Pro present).
%
% USAGE:
% ======
% [output] = IQMsimulate(model,dosing)         
% [output] = IQMsimulate(model,dosing,time)         
% [output] = IQMsimulate(model,dosing,time,ICs)         
% [output] = IQMsimulate(model,dosing,time,ICs,paramnames,paramvalues)         
% [output] = IQMsimulate(model,dosing,time,ICs,paramnames,paramvalues,OPTIONS)         
%
% moddos:      IQMmodel with INPUT definitions matching the input definitions in the provided dosing scheme 
% dosing:      IQMdosing object with dosing information to simulate the model with
% time:        timevector to simulate. If scalar than the value defines the end
%              simulation time and the time vector starts at 0. 
% ICs:         vector with initial conditions
% paramnames:  string with parameter name or cell-array with parameter-names 
% paramvalues: vector with values for the parameters (in the same order as paramnames)
% OPTIONS:     Structure with optional settings for the integrator, 
%              either defined by odeset() for MATLAB simulation of by the integrator
%              options setting used in IQMPsimulate.
%
% DEFAULT VALUES:
% ===============
% time: If not defined or left empty ([]), then the timevector will be
%   chosen to cover the full dosing schedule +50%
%   If the only dosing event occurs at time = 0, then a default time of 20 is then used.
% ICs: [] (use initial conditions, stored in the model)
% paramnames: []
% paramvalues: [] (use values stored in the model)
% OPTIONS: [] (default integrator settings, defined in getDefaultIntegratorOptionsIQM.m)
%
% Output Arguments:
% =================
% If an output argument is given, this variable will contain a standard
% simulation result structure containing the time and the state, variable,
% and reactions names and values. 
% If no output argument is given, the result is plotted using the IQMplot
% function.

% <<<COPYRIGHTSTATEMENT - IQM TOOLS LITE>>>

% Check which simulation type it is
if nargin==1,
    % It is type 1
    if nargout == 1,
        varargout{1} = simulateIQM(varargin{:});
    else
        simulateIQM(varargin{:});
    end
else
    % Check second argument
    if isIQMdosing(varargin{2}),
        % It is type 2
        if nargout == 1,
            varargout{1} = simulatedosingIQM(varargin{:});
        else
            simulatedosingIQM(varargin{:});
        end        
    else
        % It is type 1
        if nargout == 1,
            varargout{1} = simulateIQM(varargin{:});
        else
            simulateIQM(varargin{:});
        end
    end
end
```
