## Supplementary material for "A systems biology analysis of adrenergically stimulated adiponectin exocytosis in white adipocytes": All scripts used in this work: IQMstochsim.m

```
function [varargout] = IQMstochsim(model,time,V,varargin)
% IQMstochsim: Stochastic simulation of IQMmodels that only contain mass
% action kinetics. The simulator is based on the paper:
% Ullah, M., Schmidt, H., Cho, K.-H., Wolkenhauer, O. (2006) Deterministic
% Modelling and Stochastic Simulation of Pathways using MATLAB, 
% IEE Proceedings - Systems Biology, 153(2), 53-60
%
% The IQMmodel needs to have a certain format, explained below.
%
% USAGE:
% ======
% [output] = IQMstochsim(model,time,V)         
% [output] = IQMstochsim(model,time,V,units)         
% [output] = IQMstochsim(model,time,V,units,runs)         
% [output] = IQMstochsim(model,time,V,units,runs,Nsample)         
%
% model:   IQMmodel 
%          There are certain limitations on an IQMmodel used for stochastic
%          simulation. Please read further below.
% time:    End time for simulation 
% V:       Volume/NumbersFlag: If a number is given then this number is in
%          interpreted as the volume of the reaction space (given in Liter).
%          This is necessary in order to do stochastic simulations if the
%          species in the model are in concentration units.
%          If the species in the model are given in molecule numbers, the 
%          volume is not important and this is indicated by setting V to
%          [].
% units:   This value is only used in the case that the species are defined
%          in concentration units. Per default nM (units=1e-9) are assumed
%          for all species. If the model uses uM the units argument needs
%          to be set to 1e-6, etc. (default: 1e-9 used only if V has a
%          numeric value) 
% runs:    number of realizations(simulation runs) (default: 1)
% Nsample: Each Nsample-th point will be used for output (to save memory) 
%          (default: 100)
%
% Output Arguments:
% =================
% If no output arguments are given, the result of the simulation is
% plotted (if runs>1 only the mean is plotted). Otherwise the output
% argument has the following structure:
%
% output.time:            cell-array with time vectors for the single runs 
% output.speciesdata:     cell-array with simulation data for the single runs
% output.runs:            number of runs
% output.timemean:        ensemble of all time instants in the single runs
% output.speciesdatamean: matrix containing the means of the simulation data
% output.species:         cell-array containing the names of the species
%
% FORMAT OF THE IQMmodel:
% ======================
% IQMmodels that can be used for stochastic simulation need to follow some
% rules:
% 1) All reaction kinetics need to be of mass action type and be defined in
%    the following syntax:     'ReactionName' = 'kineticParameter' * ...
% 2) All reactions have to be irreversible. You can use the 
%    function IQMmakeirreversible to convert your model
% 3) The reactions can at maximum have 2 substrates and 2 products.
% 4) The right hand side of the ODEs needs only to consist of reaction rate
%    terms and eventually stoichiometric coefficients. This is necessary in
%    order to be able to determine the stoichiometric matrix.
%    More information about the required syntax can be found in the help
%    text of the function IQMstoichiometry
% 5) No variables, functions, events, functionsMATLAB are allowed to be
%    present.
% 6) Initial conditions of species are assumed to be given in numbers of
%    molecules

% <<<COPYRIGHTSTATEMENT - IQM TOOLS LITE>>>

%%%%%%%%%%%%%%%%%%%%%%%%%%%%%%%%%%
% Handle variable input arguments
%%%%%%%%%%%%%%%%%%%%%%%%%%%%%%%%%%
runs = 1;
Nsample = 100;
shownr = 0;
units = 1e-9;
if nargin == 3,
elseif nargin == 4,
    units = varargin{1};
elseif nargin == 5,
    units = varargin{1};
    runs = varargin{2};
elseif nargin == 6,
    units = varargin{1};
    runs = varargin{2};
    Nsample = varargin{3};
elseif nargin == 7,
    units = varargin{1};
    runs = varargin{2};
    Nsample = varargin{3};
    shownr  = varargin{4};
else
    error('Incorrect number of input arguments.');
end

%%%%%%%%%%%%%%%%%%%%%%%%%%%%%%%%%%
% Process the model - ALSO TAKES CARE OF NON-NUMERIC ICs
%%%%%%%%%%%%%%%%%%%%%%%%%%%%%%%%%%
% Convert model to MA structure
MA = IQMconvert2MA(model);

%%%%%%%%%%%%%%%%%%%%%%%%%%%%%%%%%%
% Obtain necessary data for simulation
%%%%%%%%%%%%%%%%%%%%%%%%%%%%%%%%%%
% initial numbers of molecules (or concentration)
n0 = MA.initialConditions;
% stoichiometric matrix
D = MA.N;
% kinetic parameters
k = MA.kineticParameters';
% Stoich coeffs of the reactants (obtained via IQMconvert2MA)
L = MA.L;

if ~isempty(V),
    % Models species given in concentrations ... determine stochastic rate
    % and number of initial molecules
    % Avogadro's number times volume
    NAV = V*units*6.02214199e23;
    % convert concentration to numbers
    n0 = n0*NAV;
    % molecularity
    K = sum(L);
    % 'particle' rate constant
    kp = k./NAV.^(K-1);
    % normalise for correct units
    kp = kp./units.^(K-1);     
    % stochastic rate constant
    c = kp.*prod(factorial(L));
else
    % Models species given in numbers ... use the kinetic constants as
    % stochastic rate constants.
    c = k;
end

if shownr == 2,
    disp('kinetic constant / stochastic rate constant');
    [k(:) c(:)]
end

%%%%%%%%%%%%%%%%%%%%%%%%%%%%%%%%%%
% Stochastic simulation
%%%%%%%%%%%%%%%%%%%%%%%%%%%%%%%%%%
[Ts,Ns,TT,NBAR] = stochIQM(n0,c,D,L,time,runs,Nsample,shownr);
if nargout == 0,
    % Display mean data
    datanames = MA.species;
    IQMplot(TT,NBAR,datanames);
else
    % return results in structure
    output = [];
    output.time = Ts;
    output.speciesdata = Ns;
    output.runs = runs;
    output.timemean = TT;
    output.speciesdatamean = NBAR;
    output.species = MA.species;
    varargout{1} = output;
    datanames = MA.species;
end
return
```
