## Supplementary material for "A systems biology analysis of adrenergically stimulated adiponectin exocytosis in white adipocytes": All scripts used in this work: IQMdataset2wide.m

```
function [datawide] = IQMdataset2wide(data,paramnames,paramvalues)
% This function expands a dataset from a row based format to a column based 
% format. Records in the row based format are stored in rows, while a parameter
% column defines the type/name of the recorded parameter. In the column  based 
% format there is a column for each measured parameter with the column name as
% the parameters name. 
% The name of the columns with the parameternames and values need to be provided.
% 
% This function will return an empty table (error) if there are columns wit NaN values.
%
% [SYNTAX]
% datawide = IQMdataset2wide(data,paramnames,paramvalues)
%
% [INPUT]
% data:         MATLAB TABLE object in row format
% paramnames:   string: name of the column in which the parameter names are recorded
% paramvalues:  string: name of the column in which the parameter values are recorded
%
% [OUTPUT]
% datawide:     dataset in wide format (one column per parameter)

% <<<COPYRIGHTSTATEMENT - IQM TOOLS LITE>>>

%% Check version number (>=R2013B required)
if verLessThan('matlab','8.2.0'),
    error('The dataset import/export functions in IQM Tools Lite require at least MATLAB R2013B.');
end

datawide = unstack(data,paramvalues,paramnames);

if isempty(datawide),
    error('Please check if the dataset to convert contains NaN values.');
end
```
