## Supplementary material for "A systems biology analysis of adrenergically stimulated adiponectin exocytosis in white adipocytes": All scripts used in this work: IQMexportCSVdataset.m

```
function [] = IQMexportCSVdataset(data,filename,FLAG_QUOTE)
% This function exports a MATLAB table as standard comma separated (CSV) datafile. 
% Nothing magic, just avoiding a lengthy "export" command and allowing similar
% naming of functions as in previous versions.
%
% [SYNTAX]
% [] = IQMexportCSVdataset(data,filename)
% [] = IQMexportCSVdataset(data,filename,FLAG_QUOTE)
%
% [INPUT]
% data:         MATLAB table object
% filename:     Filename, possibly including relative or absolute path.
%               Extension does not be provided ('.csv' will always be used).
% FLAG_QUOTE:   =0 (default): do not put "quotes" around strings. =1: do it
%               This option is ignored prior to MATLAB R2015A.
%
% [OUTPUT]
% CSV file with 'filename.csv'. If the filename also includes a path to  folder
% it will be exported there. If this folder does not yet exist, it will be created.
 
% <<<COPYRIGHTSTATEMENT - IQM TOOLS LITE>>>

%% Check version number (>=R2013B required)
if verLessThan('matlab','8.2.0'),
    error('The dataset import/export functions in IQM Tools Lite require at least MATLAB R2013B.');
end

%%%%%%%%%%%%%%%%%%%%%%%%%%%%%%
% Check input argument
%%%%%%%%%%%%%%%%%%%%%%%%%%%%%%
if ~ischar(filename),
    error('Please provide a filename as second input argument.');
end
if ~istable(data),
    error('The first input argument needs to be a MATLAB TABLE object.');
end

%%%%%%%%%%%%%%%%%%%%%%%%%%%%%%
% Handle variable inpute arguments
%%%%%%%%%%%%%%%%%%%%%%%%%%%%%%
if nargin == 2,
    FLAG_QUOTE = false;
end

%%%%%%%%%%%%%%%%%%%%%%%%%%%%%%
% Create folder if needed
%%%%%%%%%%%%%%%%%%%%%%%%%%%%%%
[p,f] = fileparts(filename);
if ~isempty(p),
    warning off
    mkdir(p);
    warning on
end

%%%%%%%%%%%%%%%%%%%%%%%%%%%%%%
% Create filename with extension
%%%%%%%%%%%%%%%%%%%%%%%%%%%%%%
filename = fullfile(p,[f '.csv']);

%%%%%%%%%%%%%%%%%%%%%%%%%%%%%%
% Export dataset
%%%%%%%%%%%%%%%%%%%%%%%%%%%%%%
try
    writetable(data,filename,'Delimiter',',','FileType','text','WriteVariableNames',true,'WriteRowNames',false,'QuoteStrings',FLAG_QUOTE);
catch
    writetable(data,filename,'Delimiter',',','FileType','text','WriteVariableNames',true,'WriteRowNames',false);
end    
```
