## Supplementary material for "A systems biology analysis of adrenergically stimulated adiponectin exocytosis in white adipocytes": All scripts used in this work: IQMgetdatasetHeader.m

```
function [colnames] = IQMgetdatasetHeader(data)
% This function returns the column names of the dataset in a cell-array.
% Input dataset can be a MATLAB table or a string with the path to a
% csv file.
%
% [SYNTAX]
% colnames = IQMgetdatasetHeader(data)
%
% [INPUT]
% data:         MATLAB TABLE object or string with path to a CSV dataset
%
% [OUTPUT]
% colnames:     Column names of dataset in cell-array

% <<<COPYRIGHTSTATEMENT - IQM TOOLS LITE>>>

% Check input
if istable(data),
    colnames = data.Properties.VariableNames;
elseif ischar(data),
    fid = fopen(data,'r');
    line = fgetl(fid);
    colnames = explodePCIQM(line);
    fclose(fid);
else
    error('Unhandled data type.');
end
```
