## Supplementary material for "A systems biology analysis of adrenergically stimulated adiponectin exocytosis in white adipocytes": All scripts used in this work: IQMloadCSVdataset.m

```
function [varargout] = IQMloadCSVdataset(filename, varargin)
% This function loads a standard CSV datafile with header as a TABLE object
% into MATLAB. Nothing magic that could not be accomplished by the readtable
% command, but allowing consistent naming as in previous versions where 
% the DATASET object from the statistics toolbox was used. 
%
% Elements in numeric columns that have entries '.','NA','N/A' will be treated
% as empty and set to NaN.
%
% [SYNTAX]
% [data]    = IQMloadCSVdataset(filename)
% [header]  = IQMloadCSVdataset(filename,FLAG_HEADER_ONLY)
%
% [INPUT]
% filename:         Filename, possibly including relative or absolute path.
%                   Extension is arbitrary.
% FLAG_HEADER_ONLY: =0: loads dataset normally, =1: return cell-array with header names
%
% [OUTPUT]
% data:      a MATLAB dataset containing the contents of the CSV datafile
% header:    cell-array with header names
%
% [ASSUMPTIONS]
% The header is expected in the first row. Special characters in header names 
% are replaced by underscores.
 
% <<<COPYRIGHTSTATEMENT - IQM TOOLS LITE>>>

%% Check version number (>=R2013B required)
if verLessThan('matlab','8.2.0'),
    error('The dataset import/export functions in IQM Tools Lite require at least MATLAB R2013B.');
end

%% %%%%%%%%%%%%%%%%%%%%%%%%%%%
% Check input arguments
%%%%%%%%%%%%%%%%%%%%%%%%%%%%%%
if ~ischar(filename),
    error('Please provide a filename as input argument.');
end
if exist(filename) ~= 2,
    error(['The file ''' filename ''' does not exist.']);
end

FLAG_HEADER_ONLY = 0;
if nargin==2,
    FLAG_HEADER_ONLY = varargin{1};
end

%%%%%%%%%%%%%%%%%%%%%%%%%%%%%%
% Load header
%%%%%%%%%%%%%%%%%%%%%%%%%%%%%%
fid = fopen(filename);
header = fgetl(fid);
fclose(fid);

%%%%%%%%%%%%%%%%%%%%%%%%%%%%%%
% Process header
%%%%%%%%%%%%%%%%%%%%%%%%%%%%%%
% split into single terms (CSV)
header = explodePCIQM(header);
% replace all unwanted chars by underscore
header = regexprep(header,'\W','_');

%%%%%%%%%%%%%%%%%%%%%%%%%%%%%%
% Return only header if desired
%%%%%%%%%%%%%%%%%%%%%%%%%%%%%%
if FLAG_HEADER_ONLY,
    varargout{1} = header;
    return
end

%%%%%%%%%%%%%%%%%%%%%%%%%%%%%%
% Load table from CSV file
%%%%%%%%%%%%%%%%%%%%%%%%%%%%%%
data = readtable(filename,'FileType','text','ReadVariableNames',true,'ReadRowNames',false,'TreatAsEmpty',{'.','NA','N/A'},'Delimiter',',');

%%%%%%%%%%%%%%%%%%%%%%%%%%%%%%
% Replace the header by the cleaned header
%%%%%%%%%%%%%%%%%%%%%%%%%%%%%%
data.Properties.VariableNames = header;

%%%%%%%%%%%%%%%%%%%%%%%%%%%%%%
% Handle general and task specific dataset columns that should be strings but might be numeric and NaN if empty
%%%%%%%%%%%%%%%%%%%%%%%%%%%%%%
FLAG_HANDLE_SPECIAL_COLUMNS = 0;
try
    IQMcheckGeneralDataFormatHeader(data);
    FLAG_HANDLE_SPECIAL_COLUMNS = 1;
end
try
    IQMcheckTaskDatasetHeader(data);
    FLAG_HANDLE_SPECIAL_COLUMNS = 1;
end
try
    IQMcheckNLMEdatasetHeader(data);
    FLAG_HANDLE_SPECIAL_COLUMNS = 1;
end

if FLAG_HANDLE_SPECIAL_COLUMNS,
    %%%%%%%%%%%%%%%%%%%%%%%%%%%%%%
    % If IGNORE is numeric, convert to cell with empty
    %%%%%%%%%%%%%%%%%%%%%%%%%%%%%%
    try
        if isnumeric(data.IGNORE),
            data.IGNORE = cell(height(data),1);
            data.IGNORE(1:end) = {''};
        end
    end
    
    %%%%%%%%%%%%%%%%%%%%%%%%%%%%%%
    % If IGNORE is 'NaN' set to ''
    %%%%%%%%%%%%%%%%%%%%%%%%%%%%%%
    try
        ix = strmatchIQM('NaN',data.IGNORE,'exact');
        data.IGNORE(ix) = {''};
%         ix = strmatchIQM('.',data.IGNORE,'exact');
%         data.IGNORE(ix) = {''};
    end
    
    %%%%%%%%%%%%%%%%%%%%%%%%%%%%%%
    % If USUBJID is numeric, convert to cell with strings
    %%%%%%%%%%%%%%%%%%%%%%%%%%%%%%
    try
        if isnumeric(data.USUBJID),
            USUBJID = cell(height(data),1);
            for k=1:height(data),
                USUBJID{k} = num2str(data.USUBJID(k));
            end
            data.USUBJID = USUBJID;
        end
    end
    
    %%%%%%%%%%%%%%%%%%%%%%%%%%%%%%
    % If SUBJECT is numeric, convert to cell with strings
    %%%%%%%%%%%%%%%%%%%%%%%%%%%%%%
    try
        if isnumeric(data.SUBJECT),
            SUBJECT = cell(height(data),1);
            for k=1:height(data),
                SUBJECT{k} = num2str(data.SUBJECT(k));
            end
            data.SUBJECT = SUBJECT;
        end
    end
    
    %%%%%%%%%%%%%%%%%%%%%%%%%%%%%%
    % If PART is numeric, convert to cell with strings
    %%%%%%%%%%%%%%%%%%%%%%%%%%%%%%
    try
        if isnumeric(data.PART),
            PART = cell(height(data),1);
            for k=1:height(data),
                PART{k} = num2str(data.PART(k));
            end
            data.PART = PART;
        end
    end
    
    %%%%%%%%%%%%%%%%%%%%%%%%%%%%%%
    % If STUDY is numeric, convert to cell with strings
    %%%%%%%%%%%%%%%%%%%%%%%%%%%%%%
    try
        if isnumeric(data.STUDY),
            STUDY = cell(height(data),1);
            for k=1:height(data),
                STUDY{k} = num2str(data.STUDY(k));
            end
            data.STUDY = STUDY;
        end
    end
    
    %%%%%%%%%%%%%%%%%%%%%%%%%%%%%%
    % If VALUE_TEXT is numeric, convert to cell with empty
    %%%%%%%%%%%%%%%%%%%%%%%%%%%%%%
    try
        if isnumeric(data.VALUE_TEXT),
            data.VALUE_TEXT = cell(height(data),1);
            data.VALUE_TEXT(1:end) = {''};
        end
    end
    
    %%%%%%%%%%%%%%%%%%%%%%%%%%%%%%
    % If COMMENT is numeric, convert to cell with empty
    %%%%%%%%%%%%%%%%%%%%%%%%%%%%%%
    try
        if isnumeric(data.COMMENT),
            data.COMMENT = cell(height(data),1);
            data.COMMENT(1:end) = {''};
        end
    end
    
    %%%%%%%%%%%%%%%%%%%%%%%%%%%%%%
    % If DATEDAY is of class datetime it should be converted to cellarray
    % of strings
    %%%%%%%%%%%%%%%%%%%%%%%%%%%%%%
    try
        if strcmp(class(data.DATEDAY),'datetime'),
            data.DATEDAY = cellstr(data.DATEDAY);
        end
    end

    %%%%%%%%%%%%%%%%%%%%%%%%%%%%%%
    % If VISNAME is numeric, convert to cell with empty
    %%%%%%%%%%%%%%%%%%%%%%%%%%%%%%
    try
        if isnumeric(data.VISNAME),
            data.VISNAME = cell(height(data),1);
            data.VISNAME(1:end) = {''};
        end
    end
    
end

%%%%%%%%%%%%%%%%%%%%%%%%%%%%%%
% Handle output arguments
%%%%%%%%%%%%%%%%%%%%%%%%%%%%%%
if nargout == 1,
    varargout{1} = data;
else
    error('Incorrect number of output arguments.');
end
```
