## Supplementary material for "A systems biology analysis of adrenergically stimulated adiponectin exocytosis in white adipocytes": All scripts used in this work: IQMloadNONCSVdataset.m

```
function [data] = IQMloadNONCSVdataset(filename,varargin)
% This function loads a NON CSV datafile with header as a dataset into
% MATLAB. Nothing magic, just avoiding a lengthy "dataset" command, and 
% replacing non a-z,A-z,_,0-9 by underscores, so the dataset functionality
% in MATLAB can work with it nicely.
% Difference to loading CSV datasets is that in this function more general
% type of files is handled. They can be tab separated, space separated,
% etc. 
%
% Files that can be loaded in this way need to have numeric content (except
% the column names in the header). Typical files that can be loaded in this 
% way are: NONMEM and MONOLIX output tables.
%
% NONMEM typically has one additional "crap" line as first line and the header 
% starting in the second line. For this the number of lines to drop can be 
% passed to this function.
%
% [SYNTAX]
% [data] = IQMloadNONCSVdataset(filename)
% [data] = IQMloadNONCSVdataset(filename,droplines)
%
% [INPUT]
% filename:     Filename, possibly including relative or absolute path.
%               Extension is arbitrary.
% droplines:    Number of lines before the start of the header (default: 0)
%
% [OUTPUT]
% data:         MATLAB Table object
%
% [ASSUMPTIONS]
% It is assumed that the dataset is rectangular and either space, tab, or
% comma separated. A separation by semicolon, colon, etc. will not work. All 
% data contents need to be numeric.

% <<<COPYRIGHTSTATEMENT - IQM TOOLS LITE>>>

%% Check version number (>=R2013B required)
if verLessThan('matlab','8.2.0'),
    error('The dataset import/export functions in IQM Tools Lite require at least MATLAB R2013B.');
end

%%%%%%%%%%%%%%%%%%%%%%%%%%%%%%
% Handle varargins
%%%%%%%%%%%%%%%%%%%%%%%%%%%%%%
if nargin == 1,
    droplines = 0;
elseif nargin == 2,
    droplines = varargin{1};
else
    error('Incorrect number of input arguments.');
end 

%%%%%%%%%%%%%%%%%%%%%%%%%%%%%%
% Check input argument
%%%%%%%%%%%%%%%%%%%%%%%%%%%%%%
if ~ischar(filename),
    error('Please provide a filename as input argument.');
end
if exist(filename) ~= 2,
    error(['The file ''' filename ''' does not exist.']);
end

%%%%%%%%%%%%%%%%%%%%%%%%%%%%%%
% Load header & content
%%%%%%%%%%%%%%%%%%%%%%%%%%%%%%
fid = fopen(filename);
k = 0;
while k<droplines,
    fgetl(fid);
    k=k+1;
end
header = fgetl(fid);
content = [];
while ~feof(fid),
    aline = fgetl(fid); % get one line at the time
    [token, remain] = strtok(aline); % separate the id from the remaining columns
    [the_id, crap] = strtok(token, '#'); % separate the id from the #1, etc.
    new_line = [the_id remain]; % reconstitute the line without the crap
    content = [content char(10) new_line];
end
fclose(fid);

%%%%%%%%%%%%%%%%%%%%%%%%%%%%%%
% Process header
%%%%%%%%%%%%%%%%%%%%%%%%%%%%%%
header2 = regexprep(strtrim(header),'[\s]+',',');
% split into single terms (CSV)
header2 = explodePCIQM(header2);
% replace all unwanted chars by underscore
header2 = regexprep(header2,'\W','_');

%%%%%%%%%%%%%%%%%%%%%%%%%%%%%%
% Create table
%%%%%%%%%%%%%%%%%%%%%%%%%%%%%%
try
    values = eval(['[' content ']']);
catch
    error('The NONCSV dataset you try to load does not fulfill the assumptions on it (rectangular, numeric only, separation by comma, space or tab).');
end
data = array2table(values);
data.Properties.VariableNames = header2;
```
