## Supplementary material for "A systems biology analysis of adrenergically stimulated adiponectin exocytosis in white adipocytes": All scripts used in this work: IQMresampleDataset.m

```
function [datasampled] = IQMresampleDataset(data,IDname,groupName)
% This function resamples a dataset. The structure and the number of 
% subjects is preserved. Useful for bootstrapping. Original subjects can
% appear several times in the resampled dataset.
%
% [SYNTAX]
% [datasampled] = IQMresampleDataset(data,IDname)
% [datasampled] = IQMresampleDataset(data,IDname,groupName)
%
% [INPUT]
% data:                 Dataset (MATLAB table object) to be resampled
% IDname:               Column name defining unique subject identifier - for example "ID"
% groupName:            Column name defining structure to keep in the
%                       resampled dataset (e.g. the treatment arm identifier column). 
%                       Then resampling is done independently for each of these groups.
%                       The values in the groupName column need to be
%                       constant for each subject (IDname). Continuous
%                       covariates should not be chosen - categorical
%                       covariates are better suited. Combinations of
%                       several covariates for grouping could be done by
%                       defining a new categorical covariate that reflects
%                       the modelers wishes.
%
% [OUTPUT]
% The resampled dataset. 

% <<<COPYRIGHTSTATEMENT - IQM TOOLS LITE>>>

%% Check version number (>=R2013B required)
if verLessThan('matlab','8.2.0'),
    error('The dataset import/export functions in IQM Tools Lite require at least MATLAB R2013B.');
end

%% Handle variable input arguments
if nargin == 2,
    groupName = '';
end    

%% Check input arguments
varnames = data.Properties.VariableNames;

ix = strmatchIQM(IDname,varnames,'exact');
if isempty(ix),
    error('Selected IDname does not exist in the dataset.');
end

if ~isempty(groupName),
    ix = strmatchIQM(groupName,varnames,'exact');
    if isempty(ix),
        error('Selected groupName does not exist in the dataset.');
    end
end

%% Determine number of groups in groupName and warn if too large
if ~isempty(groupName),
    groupCount = length(unique(data.(groupName)));
    IDcount    = length(unique(data.(IDname)));
    if groupCount > IDcount/4,
        warning('Number of groupName elements larger than 25% of subjects.');
    end
end

%% Resample if groupName not defined
if isempty(groupName),
    datasampled                 = table();
    countID                     = 1;
    
    % Get all IDs
    allID                       = unique(data.(IDname));
    
    % Get number of subjects
    Nsubjects                   = length(allID);
    
    % Sample from the IDs
    newID                       = allID(ceil(Nsubjects*rand(Nsubjects,1)));
    
    for k2=1:length(newID),
        if isnumeric(allID),
            datak2          = data(data.(IDname)==newID(k2),:);
            % Update ID so it becomes unique
            datak2.(IDname) = countID*ones(height(datak2),1);
        else
            datak2          = data(strcmp(data.(IDname),newID(k2)),:);
            % Update ID so it becomes unique
            datak2.(IDname)(1:end) = {[datak2.(IDname){1} '_' num2str(countID)]};
        end
        countID             = countID+1;
        % Add to new dataset
        datasampled         = [datasampled; datak2];
    end
end

%% Resample if groupName is defined
if ~isempty(groupName),
    datasampled                 = table();
    countID                     = 1;
    allGROUP                    = unique(data.(groupName));
    
    for k=1:length(allGROUP),
        if isnumeric(allGROUP),
            datak                   = data(data.(groupName)==allGROUP(k),:);
        else
            datak                   = data(strcmp(data.(groupName),allGROUP{k}),:);
        end
        
        % Get all IDs
        allID                   = unique(datak.(IDname));
        
        % Get number of subjects
        Nsubjects               = length(allID);
        
        % Sample from the IDs
        newID                   = allID(ceil(Nsubjects*rand(Nsubjects,1)));
        
        % add new subjects to dataset
        datasampledGROUP        = table();
        
        for k2=1:length(newID),
            
            
            if isnumeric(allID),
                datak2          = datak(datak.(IDname)==newID(k2),:);
                % Update ID so it becomes unique
                datak2.(IDname) = countID*ones(height(datak2),1);
            else
                datak2          = data(strcmp(data.(IDname),newID(k2)),:);
                % Update ID so it becomes unique
                datak2.(IDname)(1:end) = {[datak2.(IDname){1} '_' num2str(countID)]};
            end
            countID             = countID+1;
            % Add to new dataset
            datasampledGROUP    = [datasampledGROUP; datak2];
        end
        
        % Add to new dataset
        datasampled             = [datasampled; datasampledGROUP];
    end
end
```
