## Supplementary material for "A systems biology analysis of adrenergically stimulated adiponectin exocytosis in white adipocytes": All scripts used in this work: IQMsas7bdat2csv.m

```
function [] = IQMsas7bdat2csv( filenameSAS7BDAT,pathCSVfile,version )
% This function converts a sas7bdat file into a CSV file. It only works if
% SAS is installed on the system and available by the command line command
% for SAS, defined in the SETUP_PATHS_TOOLS_IQMLITE.m file.
%
% [SYNTAX]
% [] = IQMsas7bdat2csv( filenameSAS7BDAT, pathCSVfile )
%
% [INPUT]
% filenameSAS7BDAT:     String with path and filename of sas7bdat file
% pathCSVfile:          Path (no filename) to where to store the CSV file. Same filename
%                       is used as for the sas7bdat file, but with .csv at the end
%
% [OUTPUT]
% exitflag:             SAS ran successfully if exitflag==0
% 
% CSV file is written in the specified folder.

%% %%%%%%%%%%%%%%%%%%%%%%%%%%%%%%%%%%%%%%%%%%%%%%%%%%%%%%%%%%%%%%%%%%%%%%%%
% Copy the original sas file to temp folder
%%%%%%%%%%%%%%%%%%%%%%%%%%%%%%%%%%%%%%%%%%%%%%%%%%%%%%%%%%%%%%%%%%%%%%%%%%%
filenameversion = filenameSAS7BDAT;
copyfile(filenameversion,fullfile(tempdirIQM,'tempsasfile.sas7bdat'),'f');

%% %%%%%%%%%%%%%%%%%%%%%%%%%%%%%%%%%%%%%%%%%%%%%%%%%%%%%%%%%%%%%%%%%%%%
% Create SAS command in temp folder
%%%%%%%%%%%%%%%%%%%%%%%%%%%%%%%%%%%%%%%%%%%%%%%%%%%%%%%%%%%%%%%%%%%%%%%
fid = fopen([tempdirIQM 'export2csv.sas'],'w');
fprintf(fid,'libname output "%s" ;\n',tempdirIQM);
fprintf(fid,'options fmterr=no;\n');
fprintf(fid,'\n');
fprintf(fid,'proc export data=output.tempsasfile\n');
fprintf(fid,'           outfile   = "%stempsasfile.csv" replace ;\n',tempdirIQM);
fprintf(fid,' 	        delimiter = ''%s'' ;\n',char(127));
fprintf(fid,'run;\n');
fclose(fid);

%% %%%%%%%%%%%%%%%%%%%%%%%%%%%%%%%%%%%%%%%%%%%%%%%%%%%%%%%%%%%%%%%%%%%%
% Run SAS command
%%%%%%%%%%%%%%%%%%%%%%%%%%%%%%%%%%%%%%%%%%%%%%%%%%%%%%%%%%%%%%%%%%%%%%%
PATH_SAS = getSAStoolInfoIQM();
if isempty(PATH_SAS),
    error('Path to SAS not defined in SETUP_PATHS_TOOLS_IQMLITE.m');
end

exitflag = system(sprintf('%s %sexport2csv.sas',PATH_SAS,tempdirIQM));
if exitflag~=0,
    error('SAS is required for conversion from sas7bdat to CSV. It might not be available on your system.');
end
    
%% %%%%%%%%%%%%%%%%%%%%%%%%%%%%%%%%%%%%%%%%%%%%%%%%%%%%%%%%%%%%%%%%%%%%
% Load file, remove commata and replace "char(127)" sign
%%%%%%%%%%%%%%%%%%%%%%%%%%%%%%%%%%%%%%%%%%%%%%%%%%%%%%%%%%%%%%%%%%%%%%%
[~,CSVfilename] = fileparts(filenameSAS7BDAT);
contents = fileread([tempdirIQM 'tempsasfile.csv']);
contents = strrep(contents,',','');
contents = strrep(contents,char(127),',');
fid = fopen(fullfile(pathCSVfile,[CSVfilename '.csv']),'w');
fprintf(fid,'%s',contents);
fclose(fid);
    
```
