## Supplementary material for "A systems biology analysis of adrenergically stimulated adiponectin exocytosis in white adipocytes": All scripts used in this work: interp0IQM.m

```
function [yi] = interp0IQM(x,y,xi)
% interp0IQM: zero order interpolation function (lookup table)
% If of limits then the extreme points in y are taken as output. 
% interp0IQM can be used together with MEX simulation files. For MEX
% simulation functions it is IMPORTANT that the elements of the x and y
% vectors are numeric and SEPARATED BY COMMATA! 
% 
% USAGE:
% ======
% [yi] = interp0IQM(x,y,xi)   
%
% x: vector of function arguments
% y: vector of function values at the points given by x
% xi: scalar value for which to determine y by zero order interpolation

% <<<COPYRIGHTSTATEMENT - IQM TOOLS LITE>>>

yi = zeros(1,length(xi));
for k=1:length(xi),
    if xi(k) < x(2),
        yi(k) = y(1);
    elseif xi(k) >= x(end),
        yi(k) = y(end);
    else
        yi(k) = y(max(find(x<=xi(k))));
    end
end
```
