## Supplementary material for "A systems biology analysis of adrenergically stimulated adiponectin exocytosis in white adipocytes": All scripts used in this work: interp1IQM.m

```
function [yi] = interp1IQM(x,y,xi)
% interp1IQM: linear interpolation function (lookup table)
% Just the same as interp1 in MATLAB (except that if of limits then the extreme
% points in y are taken as output instead of NaN). interp1IQM can be used together 
% with MEX simulation files. For MEX simulation functions it is IMPORTANT that 
% the elements of the x and y vectors are numeric and SEPARATED BY COMMATA!
%
% USAGE:
% ======
% [yi] = interp1IQM(x,y,xi)   
%
% x: vector of function arguments
% y: vector of function values at the points given by x
% xi: scalar value for which to determine y by linear interpolation

% <<<COPYRIGHTSTATEMENT - IQM TOOLS LITE>>>

yi = zeros(1,length(xi));
for k=1:length(xi),
    if xi(k) < x(1),
        yi(k) = y(1);
    elseif xi(k) > x(end),
        yi(k) = y(end);
    else
        yi(k) = interp1(x,y,xi(k));
    end
end
```
