## Supplementary material for "A systems biology analysis of adrenergically stimulated adiponectin exocytosis in white adipocytes": All scripts used in this work: interpcseIQM.c

```
/*
 * interpcseIQM.c: Cubic spline interpolation with endpoints
 *
 * MEX interface: Henning Schmidt
 * Original C-code for spline interpolation taken from:
 *   http://www.mech.uq.edu.au/staff/jacobs/nm_lib/cmathsrc/spline.c
 *
 * The syntax of this MEX functions is as follows:
 *
 * yy = interpcseIQM(x,y,xx)
 * yy = interpcseIQM(x,y,xx,e1,e2)
 *
 * x:     x-values 
 * y:     y-values
 * xx:    x-values at which to evaluate the spline function (allow multiple)
 * e1,e2: endpoint derivatives (if specified, both need to be given)
 * yy:    interpolated value
 */

/* Some includes to be sure we got everything that is needed */
#include <mex.h>
#include <stdio.h>
#include <stdlib.h>
#include <stddef.h>
#include <stdarg.h>
#include <string.h>
#include <matrix.h>
#include <math.h>

/*
 *========================================================================
 * Initialization of the spline related functions
 *========================================================================
 */
void spline (int n, int e1, int e2, double s1, double s2, double x[], double y[], double b[], double c[], double d[], int *flag);
double seval (int n, double xx, double x[], double y[], double b[], double c[], double d[], int *last);

/*
 *========================================================================
 * Definition of needed variables to be passed to the spline functions
 *========================================================================
 */
int     n;          /* The number of data points or knots (n >= 2) */
int     e1=0;       /* = 1 to specify the slopes at the end points, = 0 to obtain the default conditions */
int     e2=0;       /* = 1 to specify the slopes at the end points, = 0 to obtain the default conditions */
double  s1;         /* the slopes at the end point x[0] */
double  s2;         /* the slopes at the end point x[n-1] */
double* x;          /* the abscissas of the knots in strictly increasing order */
double* y;          /* the ordinates of the knots */
double* b=NULL;     /* arrays of spline coefficients (length n) */
double* c=NULL;     /* arrays of spline coefficients (length n) */
double* d=NULL;     /* arrays of spline coefficients (length n) */
int     flag=0;     /* status flag = 0 normal return = 1 less than two data points; cannot interpolate = 2 x[] are not in ascending order */
int     last;       /* the segment in which xx lies */
double* xx;         /* the abscissa at which the spline is to be evaluated */
double* yy=NULL;    /* the evaluated value at xx */
int     nx;         /* number of xx values to interpolate the spline function at */
int     k;          /* loop variable */
double  b2; 
double  c2; 
double  d2; 

/*
 *===================================
 * Definition of needed MEX variables
 *===================================
 */
mxArray *xMX = NULL;
mxArray *yMX = NULL;
mxArray *xxMX = NULL;
mxArray *s1MX = NULL;
mxArray *s2MX = NULL;
mxArray *yyMX = NULL;

/*
 *========================================================================
 * MEX INTERFACE FUNCTION
 *========================================================================
 */
void mexFunction(int nlhs, mxArray *plhs[], int nrhs, const mxArray *prhs[])
{
    /*************************************/
    /* Check correct number of arguments */
    /*************************************/
    /* Check correct number of input arguments */
    if (nrhs != 3 && nrhs != 5) mexErrMsgTxt("interpcseIQM: Incorrect number of input arguments.");
    /* Check correct number of output arguments */
    if (nlhs != 1) mexErrMsgTxt("interpcseIQM: Incorrect number of output arguments.");
    
    /**************************************************************************************/
    /* Get and check input arguments and extract some first info for the spline functions */
    /**************************************************************************************/
    /* First input argument needs to be a double vector with length >= 2 */
    if (!mxIsDouble(prhs[0])) mexErrMsgTxt("interpcseIQM: Check first input argument, it needs to be a vector of x-values.");
    xMX = (mxArray *) prhs[0];
    n = mxGetM(xMX)*mxGetN(xMX);
    if (n<2) mexErrMsgTxt("interpcseIQM: please provide at least n points with n>=2 (add commata as element separators in the vectors!).");
    else x = mxGetPr(xMX);    
    
    /* Second input argument needs to be a double vector with the same length as the first */
    if (!mxIsDouble(prhs[1])) mexErrMsgTxt("interpcseIQM: Check second input argument, it needs to be a vector of y-values.");
    yMX = (mxArray *) prhs[1];
    if (n != mxGetM(yMX)*mxGetN(yMX)) mexErrMsgTxt("interpcseIQM: Second input argument (y-values) needs as many elements as first argument (x-values).");
    y = mxGetPr(yMX);    
    
    /* Third input argument should be a scalar or a vector */
    if (!mxIsDouble(prhs[2])) mexErrMsgTxt("interpcseIQM: Check third input argument, it needs to be a scalar or vector of numeric values at which to evaluate the spline.");
    xxMX = (mxArray *) prhs[2];
    nx = mxGetM(xxMX)*mxGetN(xxMX);
    if (n<1) mexErrMsgTxt("interpcseIQM: please provide at least 1 points for the third input argument.");
    xx = mxGetPr(xxMX);    

    if (nrhs > 3) {
        /* Fourth input argument needs to be a scalar double */
        if (!mxIsDouble(prhs[3]) || !mxIsScalar(prhs[3])) mexErrMsgTxt("interpcseIQM: Check fourth input argument, it needs to be a scalar numeric value with the first derivative.");
        s1MX = (mxArray *) prhs[3];
        s1 = mxGetScalar(s1MX);
        e1 = 1; /* slope given */
    
        /* Fifth input argument needs to be a scalar double */
        if (!mxIsDouble(prhs[4]) || !mxIsScalar(prhs[4])) mexErrMsgTxt("interpcseIQM: Check fifth input argument, it needs to be a scalar numeric value with the last derivative.");
        s2MX = (mxArray *) prhs[4];
        s2 = mxGetScalar(s2MX);
        e2 = 1; /* slope given */
    } else {
        e1 = 0;
        e2 = 0;
        s1 = 0.0;
        s2 = 0.0;
    }
    
    /**************************/
    /* Create output argument */
    /**************************/
    yyMX = mxCreateDoubleMatrix(nx, 1, mxREAL);
    yy = mxGetPr(yyMX);

    /***********/
    /* If n==2 */
    /***********/
    if (n==2) {        
        if (e1 == 0 || e2 == 0) mexErrMsgTxt("interpcseIQM: n=2 requires the definition of slopes.");
        b2 = s1;
        c2 = -(-3.0*y[1]+3.0*y[0]+(s2+2.0*s1)*x[1]+(-s2-2.0*s1)*x[0])/(pow(x[1],2.0)-2.0*x[0]*x[1]+pow(x[0],2.0));
        d2 = (-2.0*y[1]+2.0*y[0]+(s2+s1)*x[1]+(-s2-s1)*x[0])/(pow(x[1],3.0)-3.0*x[0]*pow(x[1],2.0)+3.0*pow(x[0],2.0)*x[1]-pow(x[0],3.0));
        
        /***************************************/
        /* Call the spline evaluation function */
        /***************************************/
        for (k=0;k<nx;k++) {
            yy[k] = y[0] + b2*(xx[k]-x[0]) + c2*pow(xx[k]-x[0],2.0) + d2*pow(xx[k]-x[0],3.0);
        }
    } else {
    /***********/
    /* If n!=2 */
    /***********/        
        /********************************/
        /* Allocate memory for b,c,d,yy */
        /********************************/
        b  = (double *) mxCalloc(n, sizeof(double));
        c  = (double *) mxCalloc(n, sizeof(double));
        d  = (double *) mxCalloc(n, sizeof(double));
        
        /****************************/
        /* Call the spline function */
        /****************************/
        spline(n, e1, e2, s1, s2, x, y, b, c, d, &flag);
        
        /***************************************/
        /* Call the spline evaluation function */
        /***************************************/
        for (k=0;k<nx;k++) {
            yy[k] = seval(n, xx[k], x, y, b, c, d, &last);
        }
        
        /*************************/
        /* Free allocated memory */
        /*************************/
        mxFree(b);
        mxFree(c);
        mxFree(d);
    }

    /*******************************/
    /* Return the result to MATLAB */
    /*******************************/
    plhs[0] = yyMX;
}

/*=========================================================================
   Cubic spline coefficients
   -------------------------
   Evaluate the coefficients b[i], c[i], d[i], i = 0, 1, .. n-1 for
   a cubic interpolating spline

   S(xx) = Y[i] + b[i] * w + c[i] * w**2 + d[i] * w**3
   where w = xx - x[i]
   and   x[i] <= xx <= x[i+1]

   The n supplied data points are x[i], y[i], i = 0 ... n-1.

   Input :
   -------
   n       : The number of data points or knots (n >= 2)
   end1,
   end2    : = 1 to specify the slopes at the end points
             = 0 to obtain the default conditions
   slope1,
   slope2  : the slopes at the end points x[0] and x[n-1]
             respectively
   x[]     : the abscissas of the knots in strictly
             increasing order
   y[]     : the ordinates of the knots

   Output :
   --------
   b, c, d : arrays of spline coefficients as defined above
             (See note 2 for a definition.)
   iflag   : status flag
            = 0 normal return
            = 1 less than two data points; cannot interpolate
            = 2 x[] are not in ascending order

   This C code written by ...  Peter & Nigel,
   ----------------------      Design Software,
                               42 Gubberley St,
                               Kenmore, 4069,
                               Australia.

   Version ... 1.1, 30 September 1987
   -------     2.0, 6 April 1989    (start with zero subscript)
                                     remove ndim from parameter list
               2.1, 28 April 1989   (check on x[])
               2.2, 10 Oct   1989   change number order of matrix

   Notes ...
   -----
   (1) The accompanying function seval() may be used to evaluate the
       spline while deriv will provide the first derivative.
   (2) Using p to denote differentiation
       y[i] = S(X[i])
       b[i] = Sp(X[i])
       c[i] = Spp(X[i])/2
       d[i] = Sppp(X[i])/6  ( Derivative from the right )
   (3) Since the zero elements of the arrays ARE NOW used here,
       all arrays to be passed from the main program should be
       dimensioned at least [n].  These routines will use elements
       [0 .. n-1].
   (4) Adapted from the text
       Forsythe, G.E., Malcolm, M.A. and Moler, C.B. (1977)
       "Computer Methods for Mathematical Computations"
       Prentice Hall
   (5) Note that although there are only n-1 polynomial segments,
       n elements are requird in b, c, d.  The elements b[n-1],
       c[n-1] and d[n-1] are set to continue the last segment
       past x[n-1].
=========================================================================*/
void spline (int n, int end1, int end2,
            double slope1, double slope2,
            double x[], double y[],
            double b[], double c[], double d[],
            int *iflag)
{  /* begin procedure spline() */

int    nm1, ib, i;
double t;
int    ascend;

nm1    = n - 1;
*iflag = 0;

if (n < 2)
  {  /* no possible interpolation */
  *iflag = 1;
  return;
  }

ascend = 1;
for (i = 1; i < n; ++i) if (x[i] <= x[i-1]) ascend = 0;
if (!ascend)
   {
   *iflag = 2;
   return;
   }

if (n >= 3)
   {    /* ---- At least quadratic ---- */

   /* ---- Set up the symmetric tri-diagonal system
           b = diagonal
           d = offdiagonal
           c = right-hand-side  */
   d[0] = x[1] - x[0];
   c[1] = (y[1] - y[0]) / d[0];
   for (i = 1; i < nm1; ++i)
      {
      d[i]   = x[i+1] - x[i];
      b[i]   = 2.0 * (d[i-1] + d[i]);
      c[i+1] = (y[i+1] - y[i]) / d[i];
      c[i]   = c[i+1] - c[i];
      }

   /* ---- Default End conditions
           Third derivatives at x[0] and x[n-1] obtained
           from divided differences  */
   b[0]   = -d[0];
   b[nm1] = -d[n-2];
   c[0]   = 0.0;
   c[nm1] = 0.0;
   if (n != 3)
      {
      c[0]   = c[2] / (x[3] - x[1]) - c[1] / (x[2] - x[0]);
      c[nm1] = c[n-2] / (x[nm1] - x[n-3]) - c[n-3] / (x[n-2] - x[n-4]);
      c[0]   = c[0] * d[0] * d[0] / (x[3] - x[0]);
      c[nm1] = -c[nm1] * d[n-2] * d[n-2] / (x[nm1] - x[n-4]);
      }

   /* Alternative end conditions -- known slopes */
   if (end1 == 1)
      {
      b[0] = 2.0 * (x[1] - x[0]);
      c[0] = (y[1] - y[0]) / (x[1] - x[0]) - slope1;
      }
   if (end2 == 1)
      {
      b[nm1] = 2.0 * (x[nm1] - x[n-2]);
      c[nm1] = slope2 - (y[nm1] - y[n-2]) / (x[nm1] - x[n-2]);
      }

   /* Forward elimination */
   for (i = 1; i < n; ++i)
     {
     t    = d[i-1] / b[i-1];
     b[i] = b[i] - t * d[i-1];
     c[i] = c[i] - t * c[i-1];
     }

   /* Back substitution */
   c[nm1] = c[nm1] / b[nm1];
   for (ib = 0; ib < nm1; ++ib)
      {
      i    = n - ib - 2;
      c[i] = (c[i] - d[i] * c[i+1]) / b[i];
      }

   /* c[i] is now the sigma[i] of the text */

   /* Compute the polynomial coefficients */
   b[nm1] = (y[nm1] - y[n-2]) / d[n-2] + d[n-2] * (c[n-2] + 2.0 * c[nm1]);
   for (i = 0; i < nm1; ++i)
      {
      b[i] = (y[i+1] - y[i]) / d[i] - d[i] * (c[i+1] + 2.0 * c[i]);
      d[i] = (c[i+1] - c[i]) / d[i];
      c[i] = 3.0 * c[i];
      }
   c[nm1] = 3.0 * c[nm1];
   d[nm1] = d[n-2];

   }  /* at least quadratic */

else  /* if n >= 3 */
   {  /* linear segment only  */
   b[0] = (y[1] - y[0]) / (x[1] - x[0]);
   c[0] = 0.0;
   d[0] = 0.0;
   b[1] = b[0];
   c[1] = 0.0;
   d[1] = 0.0;
   }
}  /* end of spline() */


/*-------------------------------------------------------------------*/
/*-----------------------------------------------------------------*/
/*-----------------------------------------------------------------*/
double seval (int n, double u,
              double x[], double y[],
              double b[], double c[], double d[],
              int *last)
/*-----------------------------------------------------------------*/
/*Purpose ...
  -------
  Evaluate the cubic spline function

  S(xx) = y[i] + b[i] * w + c[i] * w**2 + d[i] * w**3
  where w = u - x[i]
  and   x[i] <= u <= x[i+1]
  Note that Horner's rule is used.
  If u < x[0]   then i = 0 is used.
  If u > x[n-1] then i = n-1 is used.

  Input :
  -------
  n       : The number of data points or knots (n >= 2)
  u       : the abscissa at which the spline is to be evaluated
  Last    : the segment that was last used to evaluate U
  x[]     : the abscissas of the knots in strictly increasing order
  y[]     : the ordinates of the knots
  b, c, d : arrays of spline coefficients computed by spline().

  Output :
  --------
  seval   : the value of the spline function at u
  Last    : the segment in which u lies

  Notes ...
  -----
  (1) If u is not in the same interval as the previous call then a
      binary search is performed to determine the proper interval.

*/
/*-------------------------------------------------------------------*/
{  /* begin function seval() */

int    i, j, k;
double w;

i = *last;
if (i >= n-1) i = 0;
if (i < 0)  i = 0;

if ((x[i] > u) || (x[i+1] < u))
  {  /* ---- perform a binary search ---- */
  i = 0;
  j = n;
  do
    {
    k = (i + j) / 2;         /* split the domain to search */
    if (u < x[k])  j = k;    /* move the upper bound */
    if (u >= x[k]) i = k;    /* move the lower bound */
    }                        /* there are no more segments to search */
  while (j > i+1);
  }
*last = i;

/* ---- Evaluate the spline ---- */
w = u - x[i];
w = y[i] + w * (b[i] + w * (c[i] + w * d[i]));
return (w);
}
/*-------------------------------------------------------------------*/
```
