## Supplementary material for "A systems biology analysis of adrenergically stimulated adiponectin exocytosis in white adipocytes": All scripts used in this work: interpcseSlopeIQM.m

```
% interpcseSlopeIQM: Cubic spline interpolation with endpoints, returning
% the derivative at the considered point.
% This is a MEX function, so this .m file only contains the documentation.
%
% USAGE:
% ======
% yy = interpcseSlopeIQM(x,y,xx)
% yy = interpcseSlopeIQM(x,y,xx,e1,e2)
% 
% x:     x-values 
% y:     y-values
% xx:    x-values at which to evaluate the derivative of the spline function (allow multiple)
% e1,e2: endpoint derivatives (if specified, both need to be given)
%
% DEFAULT VALUES:
% ===============
% e1, e2:   =0 (no endpoint slopes defined)
%
% Output Arguments:
% =================
% yy:    derivative of spline at interpolation point

% <<<COPYRIGHTSTATEMENT - IQM TOOLS LITE>>>
```
