## Supplementary material for "A systems biology analysis of adrenergically stimulated adiponectin exocytosis in white adipocytes": All scripts used in this work: interpcsexSlopeIQM.m

```
function [yy] = interpcsexSlopeIQM(x,y,xx,varargin)
% interpcsexSlopeIQM: Cubic spline interpolation with endpoints (DERIVATIVE)
% This is an interface function only to interpcseSlopeIQM. Only in the C-code
% simulation of models there is a difference. interpcseSlopeIQM does only
% evaluate the spline coefficients once in the first call. Subsequent calls
% only evaluate the spline function. This leads to a considerable speed-up.
% On the other hand, interpcsexSlopeIQM in the C-code simulation does evaluate
% the spline coefficients each time it is called. => time dependent splines
% can be implemented. However, for MATLAB simulation there is no
% difference.
%
% USAGE:
% ======
% yy = interpcsexSlopeIQM(x,y,xx)
% yy = interpcsexSlopeIQM(x,y,xx,e1,e2)
% 
% x:     x-values 
% y:     y-values
% xx:    x-values at which to evaluate the derivative of the spline function (allow multiple)
% e1,e2: endpoint derivatives (if specified, both need to be given)
%
% DEFAULT VALUES:
% ===============
% e1, e2:   =0 (no endpoint slopes defined)
%
% Output Arguments:
% =================
% yy:    spline derivative at the interpolated point (xx)

% <<<COPYRIGHTSTATEMENT - IQM TOOLS LITE>>>


if nargin < 3 || nargin > 5,
    error('interpcsexSlopeIQM: incorrect number of input arguments.');
elseif nargin == 3,
    yy = interpcseSlopeIQM(x,y,xx);
elseif nargin == 5,
    yy = interpcseSlopeIQM(x,y,xx,varargin{1},varargin{2});
end    
```
