## Supplementary material for "A systems biology analysis of adrenergically stimulated adiponectin exocytosis in white adipocytes": All scripts used in this work: interpcsIQM.m

```
function [yi] = interpcsIQM(x,y,xi)
% interpcsIQM: cubic spline interpolation function (lookup table)
% interpcsIQM can be used together with MEX simulation functions. 
% if xi is of limits of x then the extreme points in y are taken as output.
% For MEX simulation functions it is IMPORTANT that the elements of the 
% x and y vectors are numeric and SEPARATED BY COMMATA!
% 
% USAGE:
% ======
% [yi] = interpcsIQM(x,y,xi)       
%
% x: vector of function arguments
% y: vector of function values at the points given by x
% xi: scalar value for which to determine y by linear interpolation
%
% Output Arguments:
% =================
% yi: the value of the dependent variable at xi.

% <<<COPYRIGHTSTATEMENT - IQM TOOLS LITE>>>

% input check
if length(x) ~= length(y),
    error('interpcsIQM: Different number of entries in x and y');
end

% number of data points
n = length(x);

% number of interpolations to compute
ni = length(xi);

% Calculate the derivatives
d = spline_pchip_set(n, x, y);

% Calculate the output
yi = spline_pchip_val(n, x, y, d, ni, xi);

% handle off limit values
yi(find(xi<x(1))) = y(1);
yi(find(xi>x(end))) = y(end);
return


%%%%%%%%%%%%%%%%%%%%%%%%%%%%%%%%%%%%%%%%%%%%%%%%%%%%%%%%%%%%%%%%%%%%%%%%%%%
% SPLINE_PCHIP_SET
%%%%%%%%%%%%%%%%%%%%%%%%%%%%%%%%%%%%%%%%%%%%%%%%%%%%%%%%%%%%%%%%%%%%%%%%%%%

function d = spline_pchip_set ( n, x, f )
%% SPLINE_PCHIP_SET sets derivatives for a piecewise cubic Hermite interpolant.
%
%  Discussion:
%
%    This routine computes what would normally be called a Hermite 
%    interpolant.  However, the user is only required to supply function
%    values, not derivative values as well.  This routine computes
%    "suitable" derivative values, so that the resulting Hermite interpolant
%    has desirable shape and monotonicity properties.
%
%    The interpolant will have an extremum at each point where
%    monotonicity switches direction.  
%
%    The resulting piecewise cubic Hermite function may be evaluated
%    by SPLINE_PCHIP_VAL.
%
%    This routine was originally called "PCHIM".
%
%  Modified:
%
%    14 August 2005
%
%  Author:
%
%    Fred Fritsch,
%    Mathematics and Statistics Division,
%    Lawrence Livermore National Laboratory.
%
%    MATLAB translation by John Burkardt.
%
%  Reference:
%
%    Fred Fritsch and R Carlson,
%    Monotone Piecewise Cubic Interpolation,
%    SIAM Journal on Numerical Analysis,
%    Volume 17, Number 2, April 1980, pages 238-246.
%
%    Fred Fritsch and J Butland,
%    A Method for Constructing Local Monotone Piecewise Cubic Interpolants,
%    LLNL Preprint UCRL-87559, April 1982.
%
%  Parameters:
%
%    Input, integer N, the number of data points.  N must be at least 2.
%
%    Input, real X(N), the strictly increasing independent
%    variable values.
%
%    Input, real F(N), dependent variable values to be interpolated.  This
%    routine is designed for monotonic data, but it will work for any F-array.
%    It will force extrema at points where monotonicity switches direction.
%
%    Output, real D(N), the derivative values at the
%    data points.  If the data are monotonic, these values will determine
%    a monotone cubic Hermite function.
%

%
%  Check the arguments.
%
  if ( n < 2 )
    ierr = -1;
    fprintf ( 1, '\n' );
    fprintf ( 1, 'SPLINE_PCHIP_SET - Fatal error!\n' );
    fprintf ( 1, '  Number of data points less than 2.\n' );
    error ( 'SPLINE_PCHIP_SET - Fatal error!' );
  end

  for i = 2 : n
    if ( x(i) <= x(i-1) )
      ierr = -3;
      fprintf ( 1, '\n' );
      fprintf ( 1, 'SPLINE_PCHIP_SET - Fatal error!\n' );
      fprintf ( 1, '  X array not strictly increasing.\n' );
      error ( 'SPLINE_PCHIP_SET - Fatal error!' );
    end
  end

  ierr = 0;
  nless1 = n - 1;
  h1 = x(2) - x(1);
  del1 = ( f(2) - f(1) ) / h1;
  dsave = del1;
%
%  Special case N=2, use linear interpolation.
%
  if ( n == 2 )
    d(1) = del1;
    d(n) = del1;
    return
  end 
%
%  Normal case, 3 <= N.
%
  h2 = x(3) - x(2);
  del2 = ( f(3) - f(2) ) / h2;
%
%  Set D(1) via non-centered three point formula, adjusted to be
%  shape preserving.
%
  hsum = h1 + h2;
  w1 = ( h1 + hsum ) / hsum;
  w2 = -h1 / hsum;
  d(1) = w1 * del1 + w2 * del2;

  if ( pchst( d(1), del1 ) <= 0.0 )

    d(1) = 0.0;
%
%  Need do this check only if monotonicity switches.
%
  elseif ( pchst( del1, del2 ) < 0.0 )

     dmax = 3.0 * del1;

     if ( abs ( dmax ) < abs ( d(1) ) )
       d(1) = dmax;
     end

  end
%
%  Loop through interior points.
%
  for i = 2 : nless1

    if ( 2 < i )
      h1 = h2;
      h2 = x(i+1) - x(i);
      hsum = h1 + h2;
      del1 = del2;
      del2 = ( f(i+1) - f(i) ) / h2;
    end
%
%  Set D(I)=0 unless data are strictly monotonic.
%
    d(i) = 0.0;

    temp = pchst( del1, del2 );

    if ( temp < 0.0 )

      ierr = ierr + 1;
      dsave = del2;
%
%  Count number of changes in direction of monotonicity.
%
    elseif ( temp == 0.0 )

      if ( del2 ~= 0.0D+00 )
        if ( pchst( dsave, del2 ) < 0.0 )
          ierr = ierr + 1;
        end
        dsave = del2;
      end
%
%  Use Brodlie modification of Butland formula.
%
    else

      hsumt3 = 3.0 * hsum;
      w1 = ( hsum + h1 ) / hsumt3;
      w2 = ( hsum + h2 ) / hsumt3;
      dmax = max ( abs ( del1 ), abs ( del2 ) );
      dmin = min ( abs ( del1 ), abs ( del2 ) );
      drat1 = del1 / dmax;
      drat2 = del2 / dmax;
      d(i) = dmin / ( w1 * drat1 + w2 * drat2 );

    end

  end
%
%  Set D(N) via non-centered three point formula, adjusted to be
%  shape preserving.
%
  w1 = -h2 / hsum;
  w2 = ( h2 + hsum ) / hsum;
  d(n) = w1 * del1 + w2 * del2;

  if ( pchst( d(n), del2 ) <= 0.0 )
    d(n) = 0.0;
  elseif ( pchst( del1, del2 ) < 0.0 )
%
%  Need do this check only if monotonicity switches.
%
    dmax = 3.0 * del2;

    if ( abs ( dmax ) < abs ( d(n) ) )
      d(n) = dmax;
    end

  end
  
%%%%%%%%%%%%%%%%%%%%%%%%%%%%%%%%%%%%%%%%%%%%%%%%%%%%%%%%%%%%%%%%%%%%%%%%%%%
% SPLINE_PCHIP_VAL
%%%%%%%%%%%%%%%%%%%%%%%%%%%%%%%%%%%%%%%%%%%%%%%%%%%%%%%%%%%%%%%%%%%%%%%%%%%  
function fe = spline_pchip_val ( n, x, f, d, ne, xe )

%% SPLINE_PCHIP_VAL evaluates a piecewise cubic Hermite function.
%
%  Description:
%
%    This routine may be used by itself for Hermite interpolation, or as an
%    evaluator for SPLINE_PCHIP_SET.
%
%    This routine evaluates the cubic Hermite function at the points XE.
%
%    Most of the coding between the call to CHFEV and the end of
%    the IR loop could be eliminated if it were permissible to
%    assume that XE is ordered relative to X.
%
%    CHFEV does not assume that X1 is less than X2.  Thus, it would
%    be possible to write a version of SPLINE_PCHIP_VAL that assumes a strictly
%    decreasing X array by simply running the IR loop backwards
%    and reversing the order of appropriate tests.
%
%    The present code has a minor bug, which I have decided is not
%    worth the effort that would be required to fix it.
%    If XE contains points in [X(N-1),X(N)], followed by points less than
%    X(N-1), followed by points greater than X(N), the extrapolation points
%    will be counted (at least) twice in the total returned in IERR.
%
%    The evaluation will be most efficient if the elements of XE are
%    increasing relative to X; that is, for all J <= K,
%      X(I) <= XE(J)
%    implies
%      X(I) <= XE(K).
%
%    If any of the XE are outside the interval [X(1),X(N)],
%    values are extrapolated from the nearest extreme cubic,
%    and a warning error is returned.
%
%    This routine was originally named "PCHFE".
%
%  Modified:
%
%    14 August 2005
%
%  Author:
%
%    Fred Fritsch,
%    Mathematics and Statistics Division,
%    Lawrence Livermore National Laboratory.
%
%    MATLAB translation by John Burkardt.
%
%  Reference:
%
%    Fred Fritsch and R Carlson,
%    Monotone Piecewise Cubic Interpolation,
%    SIAM Journal on Numerical Analysis,
%    Volume 17, Number 2, April 1980, pages 238-246.
%
%  Parameters:
%
%    Input, integer N, the number of data points.  N must be at least 2.
%
%    Input, real X(N), the strictly increasing independent
%    variable values.
%
%    Input, real F(N), the function values.
%
%    Input, real D(N), the derivative values.
%
%    Input, integer NE, the number of evaluation points.
%
%    Input, real XE(NE), points at which the function is to
%    be evaluated.
%
%    Output, real FE(NE), the values of the cubic Hermite
%    function at XE.
%

%
%  Check arguments.
%
  if ( n < 2 )
    ierr = -1;
    fprintf ( 1, '\n' );
    fprintf ( 1, 'SPLINE_PCHIP_VAL - Fatal error!\n' );
    fprintf ( 1, '  Number of data points less than 2.\n' );
    error ( 'SPLINE_PCHIP_VAL - Fatal error!' );
  end

  for i = 2 : n
    if ( x(i) <= x(i-1) )
      ierr = -3;
      fprintf ( 1, '\n' );
      fprintf ( 1, 'SPLINE_PCHIP_VAL - Fatal error!\n' );
      fprintf ( 1, '  X array not strictly increasing.\n' );
      error ( 'SPLINE_PCHIP_VAL - Fatal error!' );
    end
  end

  if ( ne < 1 )
    ierr = -4;
    fprintf ( 1, '\n' );
    fprintf ( 1, 'SPLINE_PCHIP_VAL - Fatal error!\n' );
    fprintf ( 1, '  Number of evaluation points less than 1.\n' );
    fe = [];
    return
  end

  ierr = 0;
%
%  Loop over intervals.
%  The interval index is IL = IR-1.
%  The interval is X(IL) <= X < X(IR).
%
  j_first = 1;
  ir = 2;

  while ( 1 )
%
%  Skip out of the loop if have processed all evaluation points.
%
    if ( ne < j_first )
      break
    end
%
%  Locate all points in the interval.
%
    j_save = ne + 1;

    for j = j_first : ne
      if ( x(ir) <= xe(j) )
        j_save = j;
        if ( ir == n )
          j_save = ne + 1;
        end
        break
      end
    end
%
%  Have located first point beyond interval.
%
    j = j_save;

    nj = j - j_first;
%
%  Skip evaluation if no points in interval.
%
    if ( nj ~= 0 )
%
%  Evaluate cubic at XE(J_FIRST:J-1).
%
      [ fe(j_first:j-1), next, ierc ] = chfev ( x(ir-1), x(ir), f(ir-1), ...
        f(ir), d(ir-1), d(ir),  nj, xe(j_first:j-1) );

      if ( ierc < 0 )
        ierr = -5;
        fprintf ( 1, '\n' );
        fprintf ( 1, 'SPLINE_PCHIP_VAL - Fatal error!\n' );
        fprintf ( 1, '  Error return from CHFEV.\n' );
        error ( 'SPLINE_PCHIP_VAL - Fatal error!' );
      end
%
%  In the current set of XE points, there are NEXT(2) to the right of X(IR).
%
      if ( next(2) ~= 0 )

        if ( ir < n )
          ierr = -5;
          fprintf ( 1, '\n' );
          fprintf ( 1, 'SPLINE_PCHIP_VAL - Fatal error!\n' );
          fprintf ( 1, '  IR < N.\n' );
          error ( 'SPLINE_PCHIP_VAL - Fatal error!' );
        end
%
%  These are actually extrapolation points.
%
        ierr = ierr + next(2);

      end
%
%  In the current set of XE points, there are NEXT(1) to the left of X(IR-1).
%
      if ( next(1) ~= 0 )
%
%  These are actually extrapolation points.
%
        if ( ir <= 2 )
          ierr = ierr + next(1);
        else

          j_new = -1;

          for i = j_first : j-1
            if ( xe(i) < x(ir-1) )
              j_new = i;
              break
            end
          end

          if ( j_new == -1 )
            ierr = -5;
            fprintf ( 1, '\n' );
            fprintf ( 1, 'SPLINE_PCHIP_VAL - Fatal error!\n' );
            fprintf ( 1, '  Could not bracket the data point.\n' );
            error ( 'SPLINE_PCHIP_VAL - Fatal error!' );
          end
%
%  Reset J.  This will be the new J_FIRST.
%
          j = j_new;
%
%  Now find out how far to back up in the X array.
%
          for i = 1 : ir-1
            if ( xe(j) < x(i) )
              break
            end
          end
%
%  At this point, either XE(J) < X(1) or X(i-1) <= XE(J) < X(I) .
%
%  Reset IR, recognizing that it will be incremented before cycling.
%
          ir = max ( 1, i-1 );

        end

      end

      j_first = j;

    end

    ir = ir + 1;

    if ( n < ir )
      break
    end

  end

%%%%%%%%%%%%%%%%%%%%%%%%%%%%%%%%%%%%%%%%%%%%%%%%%%%%%%%%%%%%%%%%%%%%%%%%%%%
% CHFEV
%%%%%%%%%%%%%%%%%%%%%%%%%%%%%%%%%%%%%%%%%%%%%%%%%%%%%%%%%%%%%%%%%%%%%%%%%%% 
function [ fe, next, ierr ] = chfev ( x1, x2, f1, f2, d1, d2, ne, xe )

%% CHFEV evaluates a cubic polynomial given in Hermite form.
%
%  Discussion:
%
%    This routine evaluates a cubic polynomial given in Hermite form at an
%    array of points.  While designed for use by SPLINE_PCHIP_VAL, it may
%    be useful directly as an evaluator for a piecewise cubic
%    Hermite function in applications, such as graphing, where
%    the interval is known in advance.
%
%    The cubic polynomial is determined by function values
%    F1, F2 and derivatives D1, D2 on the interval [X1,X2].
%
%  Modified:
%
%    14 August 2005
%
%  Author:
%
%    Fred Fritsch,
%    Mathematics and Statistics Division,
%    Lawrence Livermore National Laboratory.
%
%    MATLAB translation by John Burkardt.
%
%  Reference:
%
%    Fred Fritsch and R Carlson,
%    Monotone Piecewise Cubic Interpolation,
%    SIAM Journal on Numerical Analysis,
%    Volume 17, Number 2, April 1980, pages 238-246.
%
%    David Kahaner, Clever Moler, Steven Nash,
%    Numerical Methods and Software,
%    Prentice Hall, 1988.
%
%  Parameters:
%
%    Input, real X1, X2, the endpoints of the interval of
%    definition of the cubic.  X1 and X2 must be distinct.
%
%    Input, real F1, F2, the values of the function at X1 and
%    X2, respectively.
%
%    Input, real D1, D2, the derivative values at X1 and
%    X2, respectively.
%
%    Input, integer NE, the number of evaluation points.
%
%    Input, real XE(NE), the points at which the function is to
%    be evaluated.  If any of the XE are outside the interval
%    [X1,X2], a warning error is returned in NEXT.
%
%    Output, real FE(NE), the value of the cubic function
%    at the points XE.
%
%    Output, integer NEXT(2), indicates the number of extrapolation points:
%    NEXT(1) = number of evaluation points to the left of interval.
%    NEXT(2) = number of evaluation points to the right of interval.
%
%    Output, integer IERR, error flag.
%    0, no errors.
%    -1, NE < 1.
%    -2, X1 == X2.
%
  if ( ne < 1 )
    ierr = -1;
    fprintf ( 1, '\n' );
    fprintf ( 1, 'CHFEV - Fatal error!\n' );
    fprintf ( 1, '  Number of evaluation points is less than 1.\n' );
    fprintf ( 1, '  NE = %d\n', ne );
    error ( 'CHFEV - Fatal error!' )
  end

  h = x2 - x1;

  if ( h == 0.0 )
    ierr = -2;
    fprintf ( 1, '\n' );
    fprintf ( 1, 'CHFEV - Fatal error!\n' );
    fprintf ( 1, '  The interval [X1,X2] is of zero length.\n' );
    error ( 'CHFEV - Fatal error!' )
  end
%
%  Initialize.
%
  ierr = 0;
  next(1) = 0;
  next(2) = 0;
  xmi = min ( 0.0, h );
  xma = max ( 0.0, h );
%
%  Compute cubic coefficients expanded about X1.
%
  delta = ( f2 - f1 ) / h;
  del1 = ( d1 - delta ) / h;
  del2 = ( d2 - delta ) / h;
  c2 = -( del1 + del1 + del2 );
  c3 = ( del1 + del2 ) / h;
%
%  Evaluation loop.
%
  for i = 1 : ne

    x = xe(i) - x1;
    fe(i) = f1 + x * ( d1 + x * ( c2 + x * c3 ) );
%
%  Count the extrapolation points.
%
    if ( x < xmi )
      next(1) = next(1) + 1;
    end

    if ( xma < x )
      next(2) = next(2) + 1;
    end

  end

%%%%%%%%%%%%%%%%%%%%%%%%%%%%%%%%%%%%%%%%%%%%%%%%%%%%%%%%%%%%%%%%%%%%%%%%%%%
% PCHST
%%%%%%%%%%%%%%%%%%%%%%%%%%%%%%%%%%%%%%%%%%%%%%%%%%%%%%%%%%%%%%%%%%%%%%%%%%% 
function value = pchst( arg1, arg2 )

%% PCHST: PCHIP sign-testing routine.
%
%  Discussion:
%
%    This routine essentially computes the sign of ARG1 * ARG2.
%
%    The object is to do this without multiplying ARG1 * ARG2, to avoid
%    possible over/underflow problems.
%
%  Modified:
%
%    14 August 2005
%
%  Author:
%
%    Fred Fritsch,
%    Mathematics and Statistics Division,
%    Lawrence Livermore National Laboratory.
%
%    MATLAB translation by John Burkardt.
%
%  Reference:
%
%    Fred Fritsch and R Carlson,
%    Monotone Piecewise Cubic Interpolation,
%    SIAM Journal on Numerical Analysis,
%    Volume 17, Number 2, April 1980, pages 238-246.
%
%  Parameters:
%
%    Input, real ARG1, ARG2, two values to check.
%
%    Output, real VALUE,
%    -1.0, if ARG1 and ARG2 are of opposite sign.
%     0.0, if either argument is zero.
%    +1.0, if ARG1 and ARG2 are of the same sign.
%
  if ( arg1 == 0.0 )
    value = 0.0;
  elseif ( arg1 < 0.0 )
    if ( arg2 < 0.0 )
      value = 1.0;
    elseif ( arg2 == 0.0 )
      value = 0.0;
    elseif ( 0.0 < arg2 )
      value = -1.0;
    end
  elseif ( 0.0 < arg1 )
    if ( arg2 < 0.0 )
      value = -1.0;
    elseif ( arg2 == 0.0 )
      value = 0.0;
    elseif ( 0.0 < arg2 )
      value = 1.0;
    end
  end
```
