## Supplementary material for "A systems biology analysis of adrenergically stimulated adiponectin exocytosis in white adipocytes": All scripts used in this work: fSSmIQM.m

```
function [X,FVAL,result] = fSSmIQM(varargin)
% fSSmIQM: Interface function to a Global optimization algorithm for 
% MINLP's based on Scatter Search ("fast" version). 
%
% Note that the optimization method itself is not included in 
% the toolbox but that it can be downloaded from
% http://www.iim.csic.es/~gingproc/software.html
%
%   fSSm attempts to solve problems of the form:
%       min F(x)  subject to:  ceq(x) = 0 (equality constraints)
%        x                     c_L <= c(x) <= c_U (inequality constraints)
%                              x_L <= x <= x_U (bounds on the decision variables)            
%
% If you use fSSm and publish the results, please cite the following papers:
%
% Egea, J.A., M. Rodriguez-Fernandez, J. R. Banga and R. Mart� (2007) Scatter 
% Search for chemical and bioprocess optimization. Journal of Global Optimization 
% 37(3):481-503.
%
% Rodriguez-Fernandez, M., J. A. Egea and J. R. Banga (2006) Novel Metaheuristic for Parameter 
% Estimation in Nonlinear Dynamic Biological Systems. BMC Bioinformatics 7:483.
%
% This interface function addresses the (f)SSm package:
%   Function   : SSm beta 3.3.1
%   Written by : Process Engineering Group IIM-CSIC
%   Created on : 15/06/2005
%   Last Update: 12/03/2008
%  
%   (c) CSIC, Spanish Council for Scientific Research
%
% USAGE:
% ======
% [info] = fSSmIQM()
% [X,FVAL,result] = fSSmIQM(FUN,X,OPTIONS)
%
% FUN: Function to optimize
% X: Starting Guess (Not strictly required but kept for the sake of
%    compatible optimization function calls)
% OPTIONS: structure containing options for the algorithm:
%        OPTIONS.vtr: cost function value to reach
%        OPTIONS.highbounds: vector containing upper bounds for parameters
%           Instead of a vector highbounds can also be a scalar > 1. In the
%           latter case the highbounds are determined by multiplying this
%           scalar to the initial parameter guess X. This latter case works
%           only for positive X(i) ... if negative present then the user
%           needs to specify a whole highbounds vector. 
%        OPTIONS.lowbounds: vector containing lower bounds for parameters.
%           Instead of a vector lowbounds can also be a scalar < 1. In the
%           latter case the lowbounds are determined by multiplying this
%           scalar to the initial parameter guess X. This latter case works
%           only for positive X(i) ... if negative present then the user
%           needs to specify a whole lowbounds vector. 
% User options
%        OPTIONS.maxfunevals: Maximum number of function evaluations
%        OPTIONS.maxtime: Maximum time (in minutes) for optimization
%        OPTIONS.silent: =0: output of info, =1: no output
%        OPTIONS.plot. Plots convergence curves: 
%               0-Deactivated,
%               1-Plot curves on line,
%               2-Plot final results
%        OPTIONS.weight: Weight that multiplies the penalty term added
%           to the objective function in constrained problems
%        OPTIONS.tolc: Maximum absolute violation of the constraints.
%        OPTIONS.log_var: Indexes of the variables which will be used
%           to generate diverse solutions in different orders of magnitude
% Global options
%        OPTIONS.dim_refset: Number of elements in Refset
%        OPTIONS.ndiverse: Number of solutions generated by the
%           diversificator 
%        OPTIONS.initiate: Type of Refset initialization
%               0: Take bounds, middle point and fill by euclidean distance
%               1: Evaluate all the diverse solutions,take the dim_refset/2 best
%                  solutions and fill by euclidean distance
%        OPTIONS.combination: Type of combination of Refset elements
%               1: hyper-rectangles
%               2: linear combinations
%        OPTIONS.regenerate: Type of Refset regeneration 
%               1: Regeneration by distance diversity
%               2: Regeneration by direction diversity
%               3: Randomly alternates 1 and 2
%        OPTIONS.delete: Maximum number of Refset elements
%           deleted when regenerating Refset
%               'standard': Maximum deleted elements=dim_refset/2 
%                           (half of the elements)
%               'aggressive': Delete dim_refset-1 (all of them except the 
%                           best solution found)
%        OPTIONS.intens: Iteration interval between intensifications
%        OPTIONS.tolfun: Function tolerance for joining the Refset
%        OPTIONS.diverse_criteria: Criterion for diversification in the Refset
%               1: euclidean distance
%               2: tolerances
%        OPTIONS.tolx: Variable tolerance for joining the Refset when 
%           the euclidean distance is deactivated        
% Local options
%        OPTIONS.local.solver: Choose local solver:
%               0: local search off, 'constrnew','fminsearch','nomad','solnp',
%               'n2fb','dn2fb','dhc','fsqp','ipopt', 'misqp','lsqnonlin'
%        OPTIONS.local.n1: Number of function evaluations before applying
%           local search for the 1st time
%        OPTIONS.local.n2: Minimum number of function evaluations in the 
%           global phase between 2 local calls
%        OPTIONS.local.finish: Local solver to be used in the final
%           refinement.
%        OPTIONS.local.bestx: (=1): This option ignores all the local search
%           filters and perform a local search only when the best function
%           value has been improved. This is useful for those problems in
%           which local searches are not providing satisfactory solutions
%           but still it can help a bit more than pure global search.
%           (=0): option deactivated.
%
% Output Arguments:
% =================
% info:     calling the function w/o input argument returns information about
%           the options and a flag indicating if the algorithm can handle
%           constraints or not
% X:        Found solution
% FVAL:     Value of the function FUN at X
% result:   Structure containing results
%
%       result.fbest                   = Best objective function value
%                                         found after the optimization
%       result.xbest                   = Vector providing the best
%                                         function value
%       result.cpu_time                = Time in seconds consumed in the
%                                         optimization
%       result.f                       = Vector containing the best
%                                         objective function value after each
%                                         iteration
%       result.x                       = Matrix containing the best vector
%                                         after each iteration
%       result.time                    = Vector containing the cpu time
%                                         consumed after each iteration
%       result.neval                   = Vector containing the number of
%                                         function evaluations after each
%                                         iteration
%       result.numeval                 = Number of function evaluations
%       result.local_solutions         = Local solutions found by the
%                                         local solver (in rows)
%       result.local_solutions_values  = Function values of the local
%                                         solutions
%       result.end_crit                = Criterion to finish the
%                                         optimization
%                                         1: Maximal number of function
%                                         evaluations achieved
%                                         2: Maximum allowed CPU Time
%                                         achieved
%                                         3: Value to reach achieved
%
% DEFAULT VALUES:
% ===============
% OPTIONS.lowbounds:        0.1  => lowbounds = 0.1*X 
% OPTIONS.highbounds:       10  => highbounds = 10*X 
% OPTIONS.maxfunevals:      400*ndim
% OPTIONS.maxtime:          120 (minutes)        
% OPTIONS.silent:           0          
% OPTIONS.plot              0
% OPTIONS.weight:           1e6
% OPTIONS.tolc:             1e-5
% OPTIONS.log_var:          [1:ndim] (all variables)
%
% OPTIONS.dim_refset:       'auto'
% OPTIONS.ndiverse:         10*ndim
% OPTIONS.initiate:         1
% OPTIONS.combination:      1
% OPTIONS.regenerate:       3
% OPTIONS.delete:           'standard'
% OPTIONS.intens:           10
% OPTIONS.tolfun:           1e-4
% OPTIONS.diverse_criteria: 1
% OPTIONS.tolx:             1e-3      
%
% OPTIONS.local.solver:     'dn2fb'
% OPTIONS.local.n1:         100*ndim
% OPTIONS.local.n2:         200*ndim
% OPTIONS.local.finish:     'dn2fb'
% OPTIONS.local.bestx:      0

% <<<COPYRIGHTSTATEMENT - IQM TOOLS LITE>>>

%%%%%%%%%%%%%%%%%%%%%%%%%%%%%%%%%%%%%%%%%%%%
% CHECK IF SSm IS PRESENT
%%%%%%%%%%%%%%%%%%%%%%%%%%%%%%%%%%%%%%%%%%%%
if isempty(which('ssm_kernel')),
    error(sprintf('SSm is not installed on your system! SSmIQM is only the interface function.\nYou can download SSm from: http://www.iim.csic.es/~gingproc/software.html.\nAfter installation you can use the SSmIQM interface function included in the IQM Tools Lite.'));
end

%%%%%%%%%%%%%%%%%%%%%%%%%%%%%%%%%%%%%%%%%%%%
% ASSIGNING DEFAULT OUTPUT VALUES
%%%%%%%%%%%%%%%%%%%%%%%%%%%%%%%%%%%%%%%%%%%%
X = [];
FVAL = [];
result = [];

%%%%%%%%%%%%%%%%%%%%%%%%%%%%%%%%%%%%%%%%%%%%
% VARIABLE INPUT ARGUMENTS
%%%%%%%%%%%%%%%%%%%%%%%%%%%%%%%%%%%%%%%%%%%%
if nargin == 0,
    X.name = 'fSSmIQM';
    X.constrained = 1;
    X.description = '(fast) Scatter Search based optimization (global)';
    X.defaultOptions.names = {'maxfunevals', 'maxtime'};
    X.defaultOptions.values = {'20000','500'};
    X.defaultOptions.description = {'Maximum number of function evaluations', 'Maximum time in minutes'};
    return
elseif nargin == 2,
    FUN = varargin{1};
    X = varargin{2};
    OPTIONS = [];
elseif nargin == 3,
    FUN = varargin{1};
    X = varargin{2};
    OPTIONS = varargin{3};
else
    error('Incorrect number of input arguments.');
end
% always use the fast version 
OPTIONS.fast = 1;

%%%%%%%%%%%%%%%%%%%%%%%%%%%%%%
% Set default values
%%%%%%%%%%%%%%%%%%%%%%%%%%%%%%
ndim = length(X);
lowbounds = 0.1*X;
highbounds = 10*X;

%%%%%%%%%%%%%%
% SSm options
%%%%%%%%%%%%%%
opts = [];
% user options
opts.maxeval = 400*ndim;    % Maximum number of function evaluations 
opts.maxtime = 120*60;      % Maximum CPU time in seconds
opts.iterprint = 1;         % Print each iteration on screen
opts.plot      = 0;         % Plots convergence curves: 0-Deactivated,
                            % 1-Plot curves on line, 2-Plot final results
opts.weight    = 1e6;       % Weight that multiplies the penalty term added
                            % to the objective function in constrained
                            % problems
opts.tolc      = 1e-5;                            
opts.log_var   = [];        % Indexes of the variables which will be used
                            % to generate diverse solutions in different
                            % orders of magnitude global options
% global options
opts.dim_refset = 'auto';
opts.ndiverse = 10*ndim;
opts.initiate = 1;
opts.combination = 1;
opts.regenerate = 3;
opts.delete = 'standard';
opts.intens = 10;
opts.tolf = 1e-4;           % Function tolerance for joining the Refset
opts.diverse_criteria = 1;  % Criterion for diversification in the Refset
                            %    1: euclidean distance
                            %    2: tolerances
opts.tolx = 1e-3;           % Variable tolerance for joining the Refset when 
                            % the euclidean distance is deactivated        
% local options
opts.local.solver = 'dn2fb';      % Choose local solver:
                                  % 0: local search off, 'constrnew',
                                  % 'fminsearch','nomad','solnp',
                                  % 'n2fb','dn2fb','dhc','fsqp','ipopt',
                                  % 'misqp','lsqnonlin'
opts.local.n1 = 100*ndim;   % Number of function evaluations before applying
                            % local search for the 1st time
opts.local.n2 = 200*ndim;   % Minimum number of function evaluations in the 
                            % global phase between 2 local calls         
opts.local.finish = 'dn2fb'; % Local solver for final refinement                            
opts.local.bestx = 0; 
                            
%%%%%%%%%%%%%%%%%%%%%%%%%%%%%%
% options - Structure containing optional settings
%%%%%%%%%%%%%%%%%%%%%%%%%%%%%%
% user options
try opts.maxeval            = OPTIONS.maxfunevals; catch, end; 
try opts.maxtime            = 60*OPTIONS.maxtime; catch, end; % SSm expects seconds
try opts.iterprint          = ~OPTIONS.silent; catch, end;
try opts.plot           	= OPTIONS.plot; catch, end;
try opts.weight             = OPTIONS.weight; catch, end;
try opts.tolc               = OPTIONS.tolc; catch, end;
try opts.log_var            = OPTIONS.log_var; catch, end;
% global options
try opts.dim_refset         = OPTIONS.dim_refset; catch, end; 
try opts.ndiverse           = OPTIONS.ndiverse; catch, end; 
try opts.initiate           = OPTIONS.initiate; catch, end; 
try opts.combination        = OPTIONS.combination; catch, end; 
try opts.regenerate         = OPTIONS.regenerate; catch, end; 
try opts.delete             = OPTIONS.delete; catch, end; 
try opts.intens             = OPTIONS.intens; catch, end; 
try opts.tolf               = OPTIONS.tolfun; catch, end; 
try opts.diverse_criteria   = OPTIONS.diverse_criteria; catch, end;
try opts.tolx               = OPTIONS.tolx; catch, end;
% local options
try opts.local.solver       = OPTIONS.local.solver ; catch, end;
try opts.local.n1           = OPTIONS.local.n1; catch, end;
try opts.local.n2           = OPTIONS.local.n2; catch, end;
try opts.local.finish       = OPTIONS.local.finish; catch, end;
try opts.local.bestx        = OPTIONS.local.bestx; catch, end;

%%%%%%%%%%%%%%%%%%%%%%%%%%%%%%
% problem - Structure containing problem settings
%%%%%%%%%%%%%%%%%%%%%%%%%%%%%%
[lowbounds, highbounds] = handleLowHighBoundsIQM(OPTIONS,X,lowbounds,highbounds);
if ~ischar(FUN),
    FUN = func2str(FUN);    % convert function handle to string
end
problem = [];
problem.f = FUN;            % Name of the file containing the objective function
problem.x_L = lowbounds;    % Lower bounds of decision variables
problem.x_U = highbounds;   % Upper bounds of decision variables
problem.x_0 = X;            % Initial point(s) (optional)
try problem.vtr = OPTIONS.vtr; catch, end; % value to reach

%%%%%%%%%%%%%%%%%%%%%%%%%%%%%%
% call SSm 
%%%%%%%%%%%%%%%%%%%%%%%%%%%%%%
%disp('If you use SSm and publish the results, please cite the following papers:');
%disp(' ');
%disp('Egea, J.A., M. Rodriguez-Fernandez, J. R. Banga and R. Mart� (2007) Scatter');
%disp('Search for chemical and bioprocess optimization. Journal of Global Optimization');
%disp('37(3):481-503.');
%disp(' ');
%disp('Rodriguez-Fernandez, M., J. A. Egea and J. R. Banga (2006) Novel Metaheuristic');
%disp('for Parameter Estimation in Nonlinear Dynamic Biological Systems.');
%disp('BMC Bioinformatics 7:483.');
useFAST = 0;
if isfield(OPTIONS,'fast'),
    if OPTIONS.fast == 1,
        useFAST = 1;
    end
end
if useFAST,
    result = fssm_kernel(problem,opts);
else
    result = ssm_kernel(problem,opts);
end

%%%%%%%%%%%%%%%%%%%%%%%%%%%%%%
% handle output
%%%%%%%%%%%%%%%%%%%%%%%%%%%%%%
X = result.xbest;
FVAL = result.fbest;
```
