## Supplementary material for "A systems biology analysis of adrenergically stimulated adiponectin exocytosis in white adipocytes": All scripts used in this work: isresIQM.m

```
function [X,FVAL,EXITFLAG] = isresIQM(varargin)
% isresIQM: Stochastic Ranking for Constrained Evolutionary Minimization.
%
% The algorithm is described in:
% Thomas Philip Runarsson and Xin Yao, Search Biases in Constrained 
% Evolutionary Optimization. IEEE Transactions on Systems, Man and 
% Cybernetics -- Part C: Applications and Reviews. Vol. 35, No. 2, 
% pp 233-243, May 2005.
%
% The code of this function is based on the original code written by
% Thomas Philip Runarsson
%
% If global variable stopOptimization is set to 1, the optimization is
% stopped. This allows for stopping parameter estimation functions on user
% request. 
% 
% USAGE:
% ======
% [info] = isresIQM()
% [X,FVAL,EXITFLAG] = isresIQM(FUN,X)
% [X,FVAL,EXITFLAG] = isresIQM(FUN,X,OPTIONS)
%
% FUN: Function to optimize
%      The cost function is required to have the following calling syntax:
%         [output] = FUN(X)
%      The 'output' can be a scalar defining the 'cost', or the 'output'
%      can be a cell array with two elements: {'cost', 'constraints'}.
%      The 'cost' output argument has a scalar value indicating the 'cost'
%      of a given set of parameter values X.
%      The algorithm is able to handle inequality constraints, where
%      feasible parameter sets are defined by negative values of the
%      corresponding constraint functions. The 'constraints' output
%      argument is a vector, where each entry corresponds to one
%      constraint. If the value of an element is positive the corresponding
%      contraint is active. Simple min/max bounds on parameters need not be
%      specified using constraint functions but are taken care of by the
%      isresIQM algorithm. 
%
% X: Starting Guess for parameter vector
%
% OPTIONS: structure containing options for the algorithm:
%        OPTIONS.lowbounds: vector containing upper bounds for parameters
%           Instead of a vector highbounds can also be a scalar > 1. In the
%           latter case the highbounds are determined by multiplying this
%           scalar to the initial parameter guess X. This latter case works
%           only for positive X(i) ... if negative present then the user
%           needs to specify a whole lowbounds vector. 
%        OPTIONS.higbounds: vector containing lower bounds for parameters.
%           Instead of a vector lowbounds can also be a scalar < 1. In the
%           latter case the lowbounds are determined by multiplying this
%           scalar to the initial parameter guess X. This latter case works
%           only for positive X(i) ... if negative present then the user
%           needs to specify a whole highbounds vector. 
%        OPTIONS.linlog: string, either 'linear' or 'log'. 'linear' means
%               that the parameters are optimized based on their real
%               values, while 'log' means that the parameters are optimized
%               based on their exponent to the base 10. Choose 'linear' for
%               tight min max bounds, and choose 'log' for very wide
%               bounds. The parameter bounds are always given as "real
%               values" and not as exponents!
%        OPTIONS.lambda: scalar, integer, population size (number of offspring) (100 to 400)
%        OPTIONS.maxgen: maximum number of generations
%        OPTIONS.maxtime: Maximum time (in minutes) for optimization
%        OPTIONS.mu: parent number (mu/lambda usually 1/7)
%        OPTIONS.pf: pressure on fitness in [0 0.5] try around 0.45
%        OPTIONS.varphi: expected rate of convergence (usually 1)
%        OPTIONS.outputFunction: string with output function name. If
%               not given or if empty then no output function will be used.
%               This output function can be used to display data, or to 
%               save optimization information to a file. The function needs
%               to be in the MATLAB path. Its calling syntax is:
%
%                   'outputFunction'(g,BestParameters,BestMin,Min,Mean,Std,NrFeas,Time)
%
%                   g: current generation number
%                   BestParameters: the currently best set of parameters
%                   BestMin: the currently minimal cost function value    
%                   Min: min costfunction value in current generation (only feasible parameter sets count) 
%                   Mean: mean costfunction value in current generation (only feasible parameter sets count)  
%                   Std: standard deviation of cost function in current generation  (only feasible parameter sets count) 
%                   NrFeas: number of feasible parameter sets in current generation   
%                   Time: elapsed time
%        OPTIONS.silent: =0: output of info, =1: no output
%
% DEFAULT VALUES:
% ===============
% OPTIONS.lowbounds:    0.1  => lowbounds = 0.1*X 
% OPTIONS.highbounds:   10   => highbounds = 10*X 
% OPTIONS.linlog:       'linear'
% OPTIONS.lambda:       200
% OPTIONS.maxgen:       ndim*200
% OPTIONS.maxtime:      inf
% OPTIONS.mu:           lambda*1/7
% OPTIONS.pf:           0.45
% OPTIONS.varphi:       1
% OPTIONS.outputFunction: '' (no output function)
% OPTIONS.silent:         0 (no output of info)
%
% Output Arguments:
% =================
% info: calling the function w/o input argument returns information about
%       the options and a flag indicating if the algorithm can handle
%       constraints or not
% X: Best feasible individual           ([] if EXITFLAG = 0)
% FVAL: Value of the function FUN at X  ([] if EXITFLAG = 0)
% EXITFLAG: =1 if solution is feasible, =0 if solution is infeasible

% <<<COPYRIGHTSTATEMENT - IQM TOOLS LITE>>>

global stopOptimization

%%%%%%%%%%%%%%%%%%%%%%%%%%%%%%
% Check if isrsort.c is compiled to MEX function
%%%%%%%%%%%%%%%%%%%%%%%%%%%%%%
if exist('isrsort') ~= 3,
    % Is not compiled yet, do it
    currentpath = pwd;
    cd(fileparts(which('isrsort.c')));
    mex isrsort.c;
    cd(currentpath);
end

%%%%%%%%%%%%%%%%%%%%%%%%%%%%%%%%%%%%%%%%%%%%
% VARIABLE INPUT ARGUMENTS
%%%%%%%%%%%%%%%%%%%%%%%%%%%%%%%%%%%%%%%%%%%%
if nargin == 0,
    X = [];
    X.name = 'isresIQM';
    X.constrained = 1;
    X.description = 'Stochastic Ranking for Constrained Evolutionary Minimization (global)';  
    X.defaultOptions.names = {'maxgen', 'maxtime'};
    X.defaultOptions.values = {'1000','500'};
    X.defaultOptions.description = {'Maximum number of generations', 'Maximum time in minutes'};
    FVAL = [];
    EXITFLAG = [];
    return
elseif nargin == 2,
    FUN = varargin{1};
    X = varargin{2};
    OPTIONS = [];
elseif nargin == 3,
    FUN = varargin{1};
    X = varargin{2};
    OPTIONS = varargin{3};
else
    error('Incorrect number of input arguments.');
end

%%%%%%%%%%%%%%%%%%%%%%%%%%%%%%
% Set default values
%%%%%%%%%%%%%%%%%%%%%%%%%%%%%%
ndim = length(X);
% Default parameter bounds
lowbounds = 0.1*X;
highbounds = 10*X;
% Other
lambda = 200;
maxgen = ndim*200;
mu = ceil(lambda * 1/7);
pf = 0.45;
varphi = 1;
maxtime = inf;
linlog = 0;  % 0='linear', 1='log'
silent = 0;

%%%%%%%%%%%%%%%%%%%%%%%%%%%%%%
% Handle OPTIONS
%%%%%%%%%%%%%%%%%%%%%%%%%%%%%%
% silent
if isfield(OPTIONS,'silent'),
    if ~isempty(OPTIONS.silent),
        silent = OPTIONS.silent;
    end
end
% linlog
if isfield(OPTIONS,'linlog'),
    if strcmp(OPTIONS.linlog,'linear'),
        linlog = 0;
    elseif strcmp(OPTIONS.linlog,'log'),
        linlog = 1;
    else
        error('Wrong setting for OPTIONS.linlog.');
    end
end
% lamba
if isfield(OPTIONS,'lambda'),
    if ~isempty(OPTIONS.lambda),
        lambda = OPTIONS.lambda;
    end
    if ~isfield(OPTIONS,'mu'),
        % set also mu in case mu not defined by options
        mu = ceil(lambda*1/7);
    end
end
% maxgen
if isfield(OPTIONS,'maxgen'),
    if ~isempty(OPTIONS.maxgen),
        maxgen = OPTIONS.maxgen;
    end
end
% maxtime
if isfield(OPTIONS,'maxtime'),
    if ~isempty(OPTIONS.maxtime),
        maxtime = OPTIONS.maxtime;
    end
end
% mu
if isfield(OPTIONS,'mu'),
    if ~isempty(OPTIONS.mu),
        mu = OPTIONS.mu;
    end
end
% pf
if isfield(OPTIONS,'pf'),
    if ~isempty(OPTIONS.pf),
        pf = OPTIONS.pf;
    end
end
% varphi
if isfield(OPTIONS,'varphi'),
    if ~isempty(OPTIONS.varphi),
        varphi = OPTIONS.varphi;
    end
end
% outputFunction
outputFunction = '';
if isfield(OPTIONS,'outputFunction'),
    if ~isempty(OPTIONS.outputFunction),
        outputFunction = OPTIONS.outputFunction;
    end
end
% low and highbounds:
[lowbounds, highbounds] = handleLowHighBoundsIQM(OPTIONS,X,lowbounds,highbounds);

%%%%%%%%%%%%%%%%%%%%%%%%%%%%%%
% Handle linlog
%%%%%%%%%%%%%%%%%%%%%%%%%%%%%%
if linlog == 1, % 'log'
    lowbounds = log10(lowbounds);
    highbounds = log10(highbounds);
end

%%%%%%%%%%%%%%%%%%%%%%%%%%%%%%
% Initialize population
%%%%%%%%%%%%%%%%%%%%%%%%%%%%%%
parameterPopulation = ones(lambda,1)*lowbounds+rand(lambda,ndim).*(ones(lambda,1)*(highbounds-lowbounds));

%%%%%%%%%%%%%%%%%%%%%%%%%%%%%%
% Selection index vector
%%%%%%%%%%%%%%%%%%%%%%%%%%%%%%
sI = (1:ceil(mu))'*ones(1,ceil(lambda/mu)); sI = sI(1:lambda);

%%%%%%%%%%%%%%%%%%%%%%%%%%%%%%
% Initial algorithm parameter settings
%%%%%%%%%%%%%%%%%%%%%%%%%%%%%%
eta = ones(lambda,1)*(highbounds-lowbounds)/sqrt(ndim);
gamma = 0.85;
alpha = 0.2;
chi = (1/(2*ndim)+1/(2*sqrt(ndim)));
varphi = sqrt((2/chi)*log((1/alpha)*(exp(varphi^2*chi/2)-(1-alpha))));
tau  = varphi/(sqrt(2*sqrt(ndim)));
tau_ = varphi/(sqrt(2*ndim));
ub = ones(lambda,1)*highbounds;
lb = ones(lambda,1)*lowbounds;
eta_u = eta(1,:);
nretry = 10;

%%%%%%%%%%%%%%%%%%%%%%%%%%%%%%
% Other
%%%%%%%%%%%%%%%%%%%%%%%%%%%%%%
f = zeros(lambda,1); 
phi = zeros(lambda,ndim);
BestMin = inf;
BestMean = inf;
BestStd = inf;
BestParameters = [];

%%%%%%%%%%%%%%%%%%%%%%%%%%%%%%
% Loop over 'maxgen' generations
%%%%%%%%%%%%%%%%%%%%%%%%%%%%%%
tic;
if ~silent,
    disp(sprintf('    #g        BestMin          Min(g)        Mean(g)         Std(g)     NrFeas     Time'));
end
for g=1:maxgen,
    % fitness evaluation
    for loop=1:lambda,
        if linlog == 0, % linear
            funoutput = feval(FUN,parameterPopulation(loop,:));
        else % log
            funoutput = feval(FUN,10.^parameterPopulation(loop,:));
        end
        if iscell(funoutput),
            f(loop) = funoutput{1};
            phivec = funoutput{2};
        else
            f(loop) = funoutput;
            phivec = [];
        end
        if ~isempty(phivec),
            phi(loop,:) = phivec(:)';
        end
    end
    
    FeasiblePhi = find((sum((phi>0),2)<=0));
    NonFeasibleNaN = find(isnan(f));
    NonFeasibleInf = find(isnan(f));
    Feasible = setdiff(FeasiblePhi, union(NonFeasibleNaN,NonFeasibleInf));
    
    % Performance / statistics
    if ~isempty(Feasible),
        [MinGeneration,MinInd] = min(f(Feasible));
        MinInd = Feasible(MinInd);
        MeanGeneration = mean(f(Feasible));
        StdGeneration = std(f(Feasible));
        if MinGeneration < BestMin,
            BestParameters = parameterPopulation(MinInd,:);
            BestMin = MinGeneration;
            BestMean = MeanGeneration;
            BestStd = StdGeneration;
        end    
    else
        MinGeneration = NaN; MeanGeneration = NaN;
    end
    NrFeas = length(Feasible);

    % Compute penalty function "quadratic loss function" (or any other)
    phi(find(phi<=0)) = 0;
    phi = sum(phi.^2,2);

    % Selection using stochastic ranking 
    I = isrsort(f,phi,pf);
    parameterPopulation = parameterPopulation(I(sI),:); 
    eta = eta(I(sI),:);

    % Update eta (traditional technique using exponential smoothing)
    eta_ = eta;
    eta(mu:end,:) = eta(mu:end,:).*exp(tau_*randn(lambda-mu+1,1)*ones(1,ndim)+tau*randn(lambda-mu+1,ndim));

    % Upper bound on eta (used?)
    for i=1:ndim,
        I = find(eta(:,i)>eta_u(i));
        eta(I,i) = eta_u(i)*ones(size(I));
    end

    % make a copy of the individuals for repeat ...
    x_ = parameterPopulation;

    % differential variation
    parameterPopulation(1:mu-1,:) = parameterPopulation(1:mu-1,:) + gamma*(ones(mu-1,1)*parameterPopulation(1,:) - parameterPopulation(2:mu,:));

    % Mutation
    parameterPopulation(mu:end,:) = parameterPopulation(mu:end,:) + eta(mu:end,:).*randn(lambda-mu+1,ndim);

    % If variables are out of bounds retry "nretry" times
    I = find((parameterPopulation>ub) | (parameterPopulation<lb));
    retry = 1 ;
    while ~isempty(I)
        parameterPopulation(I) = x_(I) + eta(I).*randn(length(I),1);
        I = find((parameterPopulation>ub) | (parameterPopulation<lb));
        if (retry>nretry), 
            break; 
        end
        retry = retry + 1;
    end
    % ignore failures
    if ~isempty(I),
        parameterPopulation(I) = x_(I);
    end

    % exponential smoothing
    eta(mu:end,:) = eta_(mu:end,:) + alpha*(eta(mu:end,:) - eta_(mu:end,:));

    % output
    if ~silent,
        disp(sprintf(' %5.0f   %12.6f   %12.6g   %12.6g   %12.6g       %5.0f     %1.2e sec', g, BestMin, MinGeneration, MeanGeneration, StdGeneration, NrFeas, toc));
    end
    % call output function if defined
    if ~isempty(outputFunction),
        if linlog == 0, % linear
            feval(outputFunction, g, BestParameters, BestMin, MinGeneration, MeanGeneration, StdGeneration, NrFeas, toc);
        else % log
            feval(outputFunction, g, 10.^BestParameters, BestMin, MinGeneration, MeanGeneration, StdGeneration, NrFeas, toc);
        end
    end
    
    if toc/60 > maxtime,
        % Break the loop when more than 'OPTIONS.maxtime' minutes elapsed
        break;
    end
    if stopOptimization == 1,
        disp('User Interrupt.');
        break;
    end
end

%%%%%%%%%%%%%%%%%%%%%%%%%%%%%%
% Check results and define output
%%%%%%%%%%%%%%%%%%%%%%%%%%%%%%
if isempty(BestParameters),
    % only infeasible parameter sets
    EXITFLAG = 0;
    X = [];
    FVAL = [];
else
    EXITFLAG = 1;
    if linlog == 0, % linear
        X = BestParameters;
    else % log
        X = 10.^BestParameters;
    end
    FVAL = BestMin;
end
```
