## Supplementary material for "A systems biology analysis of adrenergically stimulated adiponectin exocytosis in white adipocytes": All scripts used in this work: pswarmIQM.m

```
function [X,FVAL,RunData] = pswarmIQM(varargin)
% pswarmIQM: Particle swarm pattern search algorithm for global optimization.
%
% Algorithm developed by A. Ismael F. Vaz and L. N. Vicente.
% 04/04/2007
% More information: http://www.norg.uminho.pt/aivaz/pswarm/
%
% In case of a publication that was prepared using pswarm, users
% are asked to cite the following publication:
% A. I. F. Vaz and L. N. Vicente, A particle swarm pattern search method
% for bound constrained global optimization,
% Journal of Global Optimization, 39 (2007) 197-219.
%
% The PSwarm algorithm takes advantage of the particle swarm algorithm in
% looking for a global optimum in the search phase of the pattern search
% algorithm. The pattern search enforces convergence for a local optimum.
%
% If the global variable stopOptimization is set to 1, the optimization is
% stopped. This allows for stopping parameter estimation functions on user
% request. 
% 
% USAGE:
% ======
% [info] = pswarmIQM()
% [X,FVAL,RunData] = pswarmIQM(FUN,X)
% [X,FVAL,RunData] = pswarmIQM(FUN,X,OPTIONS)
%
% FUN:      Function to optimize
% X:        Starting Guess
% OPTIONS:  Structure containing options for the algorithm. 
%       OPTIONS.lowbounds: vector with lower bounds for variables 
%       OPTIONS.highbounds: vector with lower bounds for variables
%       OPTIONS.logFlag: Flag indicating if the parameter space should be
%               searched linearly (0) or logarithmically (1).
%       OPTIONS.Cognitial: cognitial parameter in the velocity equation
%       OPTIONS.InerciaFinalWeight: The value of the inercia parameter in the velocity
%             at last iteration.
%       OPTIONS.InerciaInitialWeight: The value of the inercia parameter in the
%             velocity equation at first iteration.
%       OPTIONS.maxfunevals: Maximum number of objective function evaluations. Since the
%             algorithm is population based this maximum number of
%             objective function evaluation may be sligtly exceeded.
%       OPTIONS.maxiter: Maximum number of iterations allowed.
%       OPTIONS.popsize: Population size.
%       OPTIONS.Social: Social parameter in the velocity equation.
%       OPTIONS.MaxVelocityFactor: Velocity will be projected in the set
%             (UB-LB)*MaxVelocityFactor.
%       OPTIONS.CPTolerance: Stopping criteria tolerance.
%       OPTIONS.InitialDelta: Initial pattern search grid step.
%       OPTIONS.DeltaIncreseFactor: Delta will be increased by this factor (on
%             successful poll steps).
%       OPTIONS.DeltaDecreaseFactor: Delta willbe decreased by this factor (on
%             unsuccessful poll steps).
%       OPTIONS.InitialDeltaFactor: The inicial Delta will be the min(UB-LB) times this
%             factor.
%       OPTIONS.silent: 0=show iterations, 1=do not show iterations.
%
% DEFAULT VALUES:
% ===============
%       OPTIONS.lowbounds:              1e-3*X
%       OPTIONS.highbounds:             1e3*X
%       OPTIONS.Cognitial:              0.5000
%       OPTIONS.InerciaFinalWeight:     0.4000
%       OPTIONS.InerciaInitialWeight:   0.9000
%       OPTIONS.maxfunevals:            5000*length(X);
%       OPTIONS.maxiter:                1000*length(X);
%       OPTIONS.popsize:                20*length(X);
%       OPTIONS.Social:                 0.5000
%       OPTIONS.MaxVelocityFactor:      0.5000
%       OPTIONS.CPTolerance:            1.0000e-005
%       OPTIONS.InitialDelta:           2
%       OPTIONS.DeltaIncreaseFactor:    2
%       OPTIONS.DeltaDecreaseFactor:    0.5000
%       OPTIONS.InitialDeltaFactor:     5
%       OPTIONS.silent:                 0
%
% Output Arguments:
% =================
% info: calling the function w/o input argument returns information about
%       the options and a flag indicating if the algorithm can handle
%       constraints or not
% X:        Found solution (The best particle obtained for the population (Leader))
% FVAL:     Value of the function FUN at X 
% RunData:  Some statistics obout the algorithm run
%
% Copyright Information:
% ======================
% Copyright (C) 2007 A. Ismael F. Vaz and L. N. Vicente.
% 04/04/2007
% http://www.norg.uminho.pt/aivaz
% http://www.mat.uc.pt/~lnv
%
% Algorithm adapted and included in the IQM TOOLS LITE by Henning Schmidt, 
% with permission of the authors.


%%%%%%%%%%%%%%%%%%%%%%%%%%%%%%%%%%%%%%%%%%%%
% GLOBAL VARIABLES
%%%%%%%%%%%%%%%%%%%%%%%%%%%%%%%%%%%%%%%%%%%%
global stopOptimization
if isempty(stopOptimization),
    stopOptimization = 0;
end

%%%%%%%%%%%%%%%%%%%%%%%%%%%%%%%%%%%%%%%%%%%%
% VARIABLE INPUT ARGUMENTS
%%%%%%%%%%%%%%%%%%%%%%%%%%%%%%%%%%%%%%%%%%%%
if nargin == 0,
    X = [];
    X.name = 'pswarmIQM';
    X.constrained = 1;
    X.description = 'Particle swarm pattern search algorithm (global)';       
    X.defaultOptions.names = {'maxfunevals', 'maxiter'};
    X.defaultOptions.values = {'100000','40000'};
    X.defaultOptions.description = {'Maximum number of function evaluations', 'Maximum number of iterations'};
    FVAL = [];
    RunData = [];
    return
elseif nargin == 2,
    FUN = varargin{1};
    X = varargin{2};
    OPTIONS = [];
elseif nargin == 3,
    FUN = varargin{1};
    X = varargin{2};
    OPTIONS = varargin{3};
else
    error('Incorrect number of input arguments.');
end

%%%%%%%%%%%%%%%%%%%%%%%%%%%%%%%%%%%%%%%%%%%%
% HANDLE OPTIONS
%%%%%%%%%%%%%%%%%%%%%%%%%%%%%%%%%%%%%%%%%%%%
DefaultOpt.lowbounds = 1e-3*X;
DefaultOpt.highbounds = 1e3*X;
DefaultOpt.Cognitial = 0.5000;
DefaultOpt.InerciaFinalWeight = 0.4000;
DefaultOpt.InerciaInitialWeight = 0.9000;
DefaultOpt.maxfunevals = 5000*length(X);
DefaultOpt.maxiter = 1000*length(X);
DefaultOpt.popsize = 20*length(X);
DefaultOpt.Social = 0.5000;
DefaultOpt.MaxVelocityFactor = 0.5000;
DefaultOpt.CPTolerance = 1.0000e-005;
DefaultOpt.InitialDelta = 2;
DefaultOpt.DeltaIncreaseFactor = 2;
DefaultOpt.DeltaDecreaseFactor = 0.5000;
DefaultOpt.InitialDeltaFactor = 5;
DefaultOpt.silent = 0;

%%%%%%%%%%%%%%%%%%%%%%%%%%%%%%%%%%%%%%%%%%%%
% CREATE INITIAL POPULATION STRUCTURE (just one entry)
%%%%%%%%%%%%%%%%%%%%%%%%%%%%%%%%%%%%%%%%%%%%
InitialPopulation(1).x = X;

%%%%%%%%%%%%%%%%%%%%%%%%%%%%%%%%%%%%%%%%%%%%
% CREATE PROBLEM STRUCTURE
%%%%%%%%%%%%%%%%%%%%%%%%%%%%%%%%%%%%%%%%%%%%
Problem = [];
Problem.ObjFunction = FUN;
Problem.LB = GetOption('lowbounds',OPTIONS,DefaultOpt);
Problem.UB = GetOption('highbounds',OPTIONS,DefaultOpt);
Problem.Variables = length(X);
if length(Problem.LB)~=length(Problem.UB)
    error('Lower bound and upper bound arrays length mismatch.');
end

% Start clock
tic;

% Initialize options. GetOption returns the user specified value, if the
% option exists. Otherwise returns the default option.

% These are local options. Not needed for the subrotines
MaxIterations=GetOption('maxiter',OPTIONS,DefaultOpt);
MaxEvals=GetOption('maxfunevals',OPTIONS,DefaultOpt);
InerciaInitial=GetOption('InerciaInitialWeight',OPTIONS,DefaultOpt);
InerciaFinal=GetOption('InerciaFinalWeight',OPTIONS,DefaultOpt);
Cognitial=GetOption('Cognitial',OPTIONS,DefaultOpt);
Social=GetOption('Social',OPTIONS,DefaultOpt);

% These are global options. Need to be available to subrotines.
Problem.Verbose=~GetOption('silent', OPTIONS, DefaultOpt);
Problem.IncreaseDelta=GetOption('DeltaIncreaseFactor',OPTIONS,DefaultOpt);
Problem.DecreaseDelta=GetOption('DeltaDecreaseFactor',OPTIONS,DefaultOpt);
Problem.Tolerance=GetOption('CPTolerance', OPTIONS, DefaultOpt);

% Compute minimum delta for pattern search
% The minimum delta available should be at least twice the Tolerance
InitialDelta=min(Problem.UB-Problem.LB);
InitialDelta = InitialDelta / GetOption('InitialDeltaFactor', ...
    OPTIONS, DefaultOpt);
if InitialDelta < 2*Problem.Tolerance
    InitialDelta = 2*Problem.Tolerance;
end


% compute the maximum allowed velocity. High velocities will be "pushing"
% particle to the border.
MaxVelocityVect=(Problem.UB(1:Problem.Variables)-Problem.LB(1:Problem.Variables))*...
    GetOption('MaxVelocityFactor',OPTIONS,DefaultOpt);

disp('pswarmIQM: In case of a publication that was prepared using the pswarm');
disp('users are asked to cite the following publication:');
disp('A. I. F. Vaz and L. N. Vicente, A particle swarm pattern search method');
disp('for bound constrained global optimization, Journal of Global Optimization,');
disp('39 (2007) 197-219.');

if Problem.Verbose
    disp(' Nr Iter    Nr Fun Eval         Min function');
end


% Initialize counters
% Iteration counter
Problem.Stats.IterCounter=0;
% How many consecutive iterations without success (progress in the leader
% particle).
IterUnsuccess=0;
% Maximum velcity among all perticles velocity
MaxVelocity=+Inf;


% Initialize statistics counters

% Number of objective function calls. This is the number of calls to
% Problem.ObjFunction. Since particle swarm relies in a infinite penalty
% function strategy the penalty number of evaluation may be different.
Problem.Stats.ObjFunCounter=0;

% Number of Poll steps taken
Problem.Stats.PollSteps=0;

% How many poll step had success
Problem.Stats.SuccPollSteps=0;

% Keep track of what direction had success in the last poll step. The Delta
% parameter is incremented only of a success occours twice in the same
% direction.
Problem.Poll.LastSuccess=[];


% Generate initial population. Include initial population provided by user
%  if available
[Problem,Population]=InitPopulation(Problem, InitialPopulation, ...
    GetOption('popsize', OPTIONS, DefaultOpt));

% Delta for the pattern search.
Population.Delta=InitialDelta;

% Initialize patter search. Initialize the coordinate search directions.
Problem=InitPatternSearch(Problem);

% Main cycle of the algorithm
% Run pattern search until no success is attained. Call for a poll step in
% the leader particle whenever no success is recorded.

% Stop if the maximum number of iterations or objective function
% evaluations is reached.
while(Problem.Stats.IterCounter<MaxIterations && Problem.Stats.ObjFunCounter<MaxEvals && stopOptimization == 0),

    % Stop also if particle swarm velocity is bellow Tolerance and pattern
    % search Delta is bellow Tolerance
    if MaxVelocity<Problem.Tolerance && Population.Delta<Problem.Tolerance
        disp('Stopping due to velocity and tolerance');
        break;
    end
    
    % Stop if the number of active particles is equal to 1 and pattern
    % search Tolerance was attained.
    if Population.ActiveParticles <=1 && Population.Delta<Problem.Tolerance
        disp('Stopping due to single particle and tolerance');
        break;
    end
    
    % Increment iteration counter.    
    Problem.Stats.IterCounter=Problem.Stats.IterCounter+1;

    % No success iteration at begining
    Success=false;
    
    % For all particles in the swarm
    for i=1:Population.popsize
        % If particle is active
        if(Population.Active(i))
            % Compute Objective function value
            [Problem,ObjValue]=...
                PenaltyEval(Problem, Population.x(i,:));
            % Was progress attained for the current particle?
            if Population.fy(i)>ObjValue
                % Yes. Update best particle position
                Population.fy(i)=ObjValue;
                Population.y(i,:)=Population.x(i,:);
                               
                % Check if new leader is available
                if Population.fy(Population.Leader)>Population.fy(i) || Population.Leader==i
                    Population.Leader=i;
                    % Particle swarm iteration declared as successful
                    Success=true;
                    % Reset last success direction for pattern search
                    Problem.Poll.LastSuccess=[];
                end
            end
        end
    end
    
    % Successful iteration?
    if ~Success
        % No success in searh phase. Proceed to a poll step if possible.
        if Population.Delta >= Problem.Tolerance
            [Problem,Population]=PollStep(Problem,Population);
            Problem.Stats.PollSteps=Problem.Stats.PollSteps+1;
            % Reset on the number of unsuccessful iterations without a poll
            % step
            IterUnsuccess=0;
        else
            % An unsuccessful iteration without poll step
            IterUnsuccess=IterUnsuccess+1;
        end
    else
        % Success
        IterUnsuccess=0;
        % Leader changed.
        % Increase Delta. Reset on the local search.
        if Population.Delta<InitialDelta
            Population.Delta=Population.Delta*Problem.IncreaseDelta;
        end
        % check for lower bounds
        if Population.Delta<Problem.Tolerance
            Population.Delta=2*Problem.Tolerance;
        end
    end
    
    
    % Compute inercia.
    Inercia = InerciaInitial - ...
        (InerciaInitial-InerciaFinal)*Problem.Stats.IterCounter/MaxIterations;

    % Update velocity and new particle positions
    % For all particles
    for i=1:Population.popsize
        % Is active?
        if Population.Active(i)
            % Update velocity for real variables
            Population.vx(i,:)=Projection(Inercia*Population.vx(i,:)+ ...
                Cognitial*unifrndIQM(0,ones(1,Problem.Variables)).*...
                  (Population.y(i,:)-Population.x(i,:))+...
                Social*unifrndIQM(0,ones(1,Problem.Variables)).*...
                  (Population.y(Population.Leader,:)-Population.x(i,:)),...
                -MaxVelocityVect,MaxVelocityVect);
            % Update particle position and check bound limits
            Population.x(i,:)=Projection(Population.x(i,:)+Population.vx(i,:),...
                Problem.LB(1:Problem.Variables), ...
                Problem.UB(1:Problem.Variables));
        end
    end

    % To compute population norm. Start with Leader.
    MaxVelocity=norm(Population.x(Population.Leader));
    
    % Reset number of active particles.
    Population.ActiveParticles=0;
    for i=1:Population.popsize
        % Check if particle is active and we do not want to remove the
        % leader.
        if Population.Active(i) && Population.Leader~=i
            % compute particle velocity norm (for the stopping criteria.
            VelocityNorm=norm(Population.vx(i,:));
            Distance=norm(Population.x(i,:)-Population.x(Population.Leader,:));
            if Distance<Population.Delta && VelocityNorm<Population.Delta
                % Is neighbour
                Population.Active(i)=false;
            else
                % Is integer, but not real neighbour
                MaxVelocity=MaxVelocity+VelocityNorm;
            end
        end
        % Account for active particles.
        if Population.Active(i)
            Population.ActiveParticles=Population.ActiveParticles+1;
        end
    end
    
    if Problem.Verbose,
        disp(sprintf(' %5.0f        %5.0f           %12.6g            %s', Problem.Stats.IterCounter, Problem.Stats.ObjFunCounter, Population.fy(Population.Leader)));
    end

end


% End of main cycle ...

% print final time
toc;

% Print if it was stopped due to the maximum of iterations or objective
% function evaluations
if Problem.Stats.IterCounter>=MaxIterations || Problem.Stats.ObjFunCounter>=MaxEvals
    disp('Maximum number of iterations or objective function evaluations reached');
end

if stopOptimization == 1,
    disp('User Interrupt.');
end

% return leader position and objective function value
BestParticle=[Population.y(Population.Leader,:)];
BestParticleObj=Population.fy(Population.Leader);
RunData=Problem.Stats;

if Problem.Verbose
    % display some statistics
    disp(Problem.Stats);
end
X = BestParticle;
FVAL = BestParticleObj;
return

%%%%%%%%%%%%%%%%%%%%%%%%%%%%%%%%%%%%%%%%%%%%%%%%%%%%%%%%%%%%%%%%%%%%%%%%%%%
%
% subrotine InitPopulation
%    Randomly initialize the population
%    Include initial guesses, if provided by the user
%
% Input:
%   Problem - problem data
%   InitialPopulation - Inicial population provided by user
%   popsize - Requested size for the population
%
% Output:
%   Problem - problem data (problem data may be update)
%   Population - population data
%
%%%%%%%%%%%%%%%%%%%%%%%%%%%%%%%%%%%%%%%%%%%%%%%%%%%%%%%%%%%%%%%%%%%%%%%%%%%
function [Problem,Population]=InitPopulation(Problem, ...
    InitialPopulation, popsize)
% Check if user provides a valid initial population
if ~isempty(InitialPopulation) && ~isstruct(InitialPopulation)
    error('pswarm:InitPopulation:InitialPopulation', 'Initial population must be defined in a structure.');
else
    % Check for size
    if length(InitialPopulation)>popsize
        % User provided an initial population greater then the population
        % size
        if Problem.Verbose
            fprintf('Initial population is greater then population size. Incresing population size from ');
            fprintf(popsize);
            fprintf(' to ');
            fprintf(length(InitialPopulation));
        end
        % Population size is increased to fit the number of initial guesses
        Population.popsize=length(InitialPopulation);
    else
        % Otherwise just accept the proposed population size
        Population.popsize=popsize;
    end 
    % Copy the initial population for the population and initialize them
    for i=1:length(InitialPopulation)
        % Particle position.
        Population.x(i,:)=Projection(InitialPopulation(i).x,...
                Problem.LB(1:Problem.Variables), ...
                Problem.UB(1:Problem.Variables));
        % Best particle position.
        Population.y(i,:)=Population.x(i,:);
        % Particle velocities.
        Population.vx(i,:)=zeros(1,Problem.Variables);
        % Particle is active at begining
        Population.Active(i)=true;
        [Problem,Population.fy(i)]=...
            PenaltyEval(Problem, Population.x(i,:));
    end
end
% Ramdomly generate the remaining population
for i=length(InitialPopulation)+1:Population.popsize
    % Particle positions.
    Population.x(i,:)=unifrndIQM(Problem.LB(1:Problem.Variables),...
        Problem.UB(1:Problem.Variables));
    % Best particle position.
    Population.y(i,:)=Population.x(i,:);
    % Particle velocities
    Population.vx(i,:)=zeros(1,Problem.Variables);
    % Particle active or inactive
    Population.Active(i)=true;
    [Problem,Population.fy(i)]=...
        PenaltyEval(Problem, Population.x(i,:));
end
Population.ActiveParticles=Population.popsize;
Population.Leader=1;
return;

%%%%%%%%%%%%%%%%%%%%%%%%%%%%%%%%%%%%%%%%%%%%%%%%%%%%%%%%%%%%%%%%%%%%%%%%%%%
%  subrotine GetOption
%
%  Input:
%    Option - option to get the value
%    OPTIONS - a list of options provided by user
%    DefaultOpt -  a list of default options
%
%  Output:
%    Value - The value specified by user for Option or the default
%
%%%%%%%%%%%%%%%%%%%%%%%%%%%%%%%%%%%%%%%%%%%%%%%%%%%%%%%%%%%%%%%%%%%%%%%%%%%
function [Value]=GetOption(Option, OPTIONS, DefaultOpt)
% Check for user provided options
if isempty(OPTIONS) || ~isstruct(OPTIONS)
    % User does not provides OPTIONS
    Value=DefaultOpt.(Option);
    return;
end
% Try the option provided by user
try
    Value=OPTIONS.(Option);
catch
    Value=[];
end
% Option not provided by user
if isempty(Value)
    Value=DefaultOpt.(Option);    
end
return
```
