## Supplementary material for "A systems biology analysis of adrenergically stimulated adiponectin exocytosis in white adipocytes": All scripts used in this work: simannealingIQM.m

```
function [X,FVAL,EXITFLAG] = simannealingIQM(varargin)
% simannealingIQM: Minimization by simulated annealing. Algorithm partly 
% based on section 10.4 in "Numerical Recipes in C", ISBN 0-521-43108-5.
% The implementation of this simmulated annealing method uses a nonlinear
% simplex search as basis. The algorithm has been modified in order to be
% able to take into account simple box constraints on parameter values.
%
% If global variable stopOptimization is set to 1, the optimization is
% stopped. This allows for stopping parameter estimation functions on user
% request. 
% 
% USAGE:
% ======
% [info] = simannealingIQM()
% [X,FVAL,EXITFLAG] = simannealingIQM(FUN,X,OPTIONS)
%
% FUN: Function to optimize
% X: Starting Guess
% OPTIONS: structure containing options for the algorithm:
%        OPTIONS.tempstart: Starting temperature (should be around the
%           order of magnitude (or higher) than the cost function at the
%           initial guess)
%        OPTIONS.tempend: Ending temperature (When performed all iterations
%           for this temperature, the temperature is set to 0 and the
%           simannealingIQM function converges to a normal simplex search)
%        OPTIONS.tempfactor: Reduction factor for temperature after running
%           through all iterations for current temperature
%        OPTIONS.maxitertemp: Number of iterations to carry put for each
%           non-zero temperature
%        OPTIONS.maxitertemp0: Number of iterations to carry out for 0
%           temperature
%        OPTIONS.maxtime: Maximum time (in minutes) for optimization
%        OPTIONS.tolx: Tolerance for max difference between the coordinates
%           of the vertices.
%        OPTIONS.tolfun: Tolerance for difference between best and worst
%           function evaluation in simplex
%        OPTIONS.highbounds: vector containing upper bounds for parameters
%           Instead of a vector highbounds can also be a scalar > 1. In the
%           latter case the highbounds are determined by multiplying this
%           scalar to the initial parameter guess X. This latter case works
%           only for positive X(i) ... if negative present then the user
%           needs to specify a whole highbounds vector. 
%        OPTIONS.lowbounds: vector containing lower bounds for parameters.
%           Instead of a vector lowbounds can also be a scalar < 1. In the
%           latter case the lowbounds are determined by multiplying this
%           scalar to the initial parameter guess X. This latter case works
%           only for positive X(i) ... if negative present then the user
%           needs to specify a whole lowbounds vector. 
%        OPTIONS.outputFunction: string with output function name. If
%           not given or if empty then no output function will be used.
%           This output function can be used to display data, or to 
%           save optimization information to a file. The function needs
%           to be in the MATLAB path. Its calling syntax is:
%                   'outputFunction'(bestparameters,bestfunctionvalue,currentsimplex)
%        OPTIONS.silent: =0: output of info, =1: no output
%
% DEFAULT VALUES:
% ===============
% OPTIONS.tempstart = 10*magnitude of function value at initial guess
% OPTIONS.tempend = 0.1
% OPTIONS.tempfactor = chosen such that 10 reductions of temperature
% OPTIONS.maxitertemp = 50*numberVariables
% OPTIONS.maxitertemp0 = 200*numberVariables
% OPTIONS.maxtime = 120
% OPTIONS.tolx = 1e-10
% OPTIONS.tolfun = 1e-10
% OPTIONS.lowbounds:    0.1  => lowbounds = 0.1*X 
% OPTIONS.highbounds:    10  => highbounds = 10*X 
% OPTIONS.outputFunction: no output function ('')
% OPTIONS.silent:         0 (no output of info)
%
% Output Arguments:
% =================
% info: calling the function w/o input argument returns information about
%       the options and a flag indicating if the algorithm can handle
%       constraints or not
% X: Found solution
% FVAL: Value of the function FUN at X
% EXITFLAG: 1=success, 0=not found

% <<<COPYRIGHTSTATEMENT - IQM TOOLS LITE>>>

%%%%%%%%%%%%%%%%%%%%%%%%%%%%%%%%%%%%%%%%%%%%
% GLOBAL VARIABLES
%%%%%%%%%%%%%%%%%%%%%%%%%%%%%%%%%%%%%%%%%%%%
global ndim nfunk Xguess lowbounds highbounds Temp ybest pbest tempstart stopOptimization

%%%%%%%%%%%%%%%%%%%%%%%%%%%%%%%%%%%%%%%%%%%%
% VARIABLE INPUT ARGUMENTS
%%%%%%%%%%%%%%%%%%%%%%%%%%%%%%%%%%%%%%%%%%%%
if nargin == 0,
    X = [];
    X.name = 'simannealingIQM';
    X.constrained = 1;
    X.description = 'Simulated annealing based on Nelder-Mead (global)';       
    X.defaultOptions.names = {'tempstart','tempend','tempfactor','maxitertemp', 'maxitertemp0', 'maxtime'};
    X.defaultOptions.values = {'1000','0.1','0.2','1000','1000','500'};
    X.defaultOptions.description = {'Starting temperature', 'Ending temperature before assuming T=0', 'Reduction factor for temperature', 'Number of iterations if T~=0', 'Number of iterations if T==0','Maximum time in minutes'};
    FVAL = [];
    EXITFLAG = [];
    return
elseif nargin == 2,
    FUN = varargin{1};
    X = varargin{2};
    OPTIONS = [];
elseif nargin == 3,
    FUN = varargin{1};
    X = varargin{2};
    OPTIONS = varargin{3};
else
    error('Incorrect number of input arguments.');
end

%%%%%%%%%%%%%%%%%%%%%%%%%%%%%%
% Set default values
%%%%%%%%%%%%%%%%%%%%%%%%%%%%%%
EXITFLAG = 1;
ndim = length(X);
tempstart = -1;     % default value calculation later
tempend = 0.1;
tempfactor = -1;    % default value calculation later
maxitertemp = 50*ndim;
maxitertemp0 = 200*ndim;
maxtime = 120;
tolx = 1e-10;
tolfun = 1e-10;
lowbounds = 0.1*X;
highbounds = 10*X;
outputFunction = '';
silent = 0;

%%%%%%%%%%%%%%%%%%%%%%%%%%%%%%%%%%%%%%%%%%%%
% HANDLE OPTIONS
%%%%%%%%%%%%%%%%%%%%%%%%%%%%%%%%%%%%%%%%%%%%
% silent
if isfield(OPTIONS,'silent'),
    if ~isempty(OPTIONS.silent),
        silent = OPTIONS.silent;
    end
end
% tolfun
if isfield(OPTIONS,'tolfun'),
    if ~isempty(OPTIONS.tolfun),
        tolfun = OPTIONS.tolfun;
    end
end
% tolx
if isfield(OPTIONS,'tolx'),
    if ~isempty(OPTIONS.tolx),
        tolx = OPTIONS.tolx;
    end
end
% maxtime
if isfield(OPTIONS,'maxtime'),
    if ~isempty(OPTIONS.maxtime),
        maxtime = OPTIONS.maxtime;
    end
end
% maxitertemp0
if isfield(OPTIONS,'maxitertemp0'),
    if ~isempty(OPTIONS.maxitertemp0),
        maxitertemp0 = OPTIONS.maxitertemp0;
    end
end
% maxitertemp
if isfield(OPTIONS,'maxitertemp'),
    if ~isempty(OPTIONS.maxitertemp),
        maxitertemp = OPTIONS.maxitertemp;
    end
end
% tempfactor
if isfield(OPTIONS,'tempfactor'),
    if ~isempty(OPTIONS.tempfactor),
        tempfactor = OPTIONS.tempfactor;
    end
end
% tempend
if isfield(OPTIONS,'tempend'),
    if ~isempty(OPTIONS.tempend),
        tempend = OPTIONS.tempend;
    end
end
% tempstart
if isfield(OPTIONS,'tempstart'),
    if ~isempty(OPTIONS.tempstart),
        tempstart = OPTIONS.tempstart;
    end
end
% outputFunction
outputFunction = '';
if isfield(OPTIONS,'outputFunction'),
    if ~isempty(OPTIONS.outputFunction),
        outputFunction = OPTIONS.outputFunction;
    end
end
% low and highbounds:
[lowbounds, highbounds] = handleLowHighBoundsIQM(OPTIONS,X,lowbounds,highbounds);

%%%%%%%%%%%%%%%%%%%%%%%%%%%%%%%%%%%%%%%%%%%%
% INITIALIZE EMPTY SIMPLEX DATA
%%%%%%%%%%%%%%%%%%%%%%%%%%%%%%%%%%%%%%%%%%%%
p = zeros(ndim+1,ndim);     % vertice vectors in rows
y = zeros(ndim+1,1);

%%%%%%%%%%%%%%%%%%%%%%%%%%%%%%%%%%%%%%%%%%%%
% EVALUATE FUNCTION IN INITIAL GUESS
%%%%%%%%%%%%%%%%%%%%%%%%%%%%%%%%%%%%%%%%%%%%
ybest = inf;
pbest = X(:)';
ybest = costFunction(FUN,pbest);

%%%%%%%%%%%%%%%%%%%%%%%%%%%%%%%%%%%%%%%%%%%%
% DETERMINE START TEMP AND TEMPFACTOR (if not given by the user)
%%%%%%%%%%%%%%%%%%%%%%%%%%%%%%%%%%%%%%%%%%%%
if tempstart < 0,
    % tempstart not given by the user => determine here a default value
    tempstart = 10*abs(ybest);
end
if tempfactor < 0,
    % tempfactor not given by the user => determine here a default value
    tempfactor = (tempstart/tempend)^(-1/9);
end

%%%%%%%%%%%%%%%%%%%%%%%%%%%%%%%%%%%%%%%%%%%%
% TEMPERATURE LOOP - STARTING FROM BEST POINT
%%%%%%%%%%%%%%%%%%%%%%%%%%%%%%%%%%%%%%%%%%%%
Temp = tempstart;
tic % start timer
nfunk = 1;
nriterations = 0;
while(1),
    if Temp < tempend,
        Temp = 0;
        MAXITERTEMP = maxitertemp0;
    else 
        MAXITERTEMP = maxitertemp;
    end
    %%%%%%%%%%%%%%%%%%%%%%%%%%%%%%%%%%%%%%%%%%%%
    % FIRST POINT OF INITIAL SIMPLEX
    %%%%%%%%%%%%%%%%%%%%%%%%%%%%%%%%%%%%%%%%%%%%
    if ~silent,
        disp(sprintf(' Current best estimation: [%s]',sprintf('%g ', pbest)));
    end
    Xguess = pbest;
    p(1,:) = Xguess;
    y(1) = costFunction(FUN,Xguess);
    if ~silent,
        disp(' Nr Iter  Nr Fun Eval    Min function       Best function       Temp      Algorithm Step');
        if Temp == tempstart,
            disp(sprintf(' %5.0f   %5.0f       %12.6g       %12.6g   %12.6g       %s', nriterations, nfunk, y(1), ybest, Temp, 'initial guess'));
        else
            disp(sprintf(' %5.0f   %5.0f       %12.6g       %12.6g   %12.6g       %s', nriterations, nfunk, y(1), ybest, Temp, 'best point'));
        end
    end
    %%%%%%%%%%%%%%%%%%%%%%%%%%%%%%%%%%%%%%%%%%%%
    % REMAINING POINTS OF INITIAL SIMPLEX
    %%%%%%%%%%%%%%%%%%%%%%%%%%%%%%%%%%%%%%%%%%%%
    % construct vertices by modifying one element each
    % relative changes in case that elements are non-zero,
    % absolute changes in case that elements are zero
    relativeDelta = 0.25;
    absoluteDelta = 0.5;
    for k = 1:ndim
        Xmodify = Xguess;
        if Xmodify(k) == 0
            % absolute change
            if highbounds(k) > absoluteDelta,
                Xmodify(k) = absoluteDelta;
            else 
                Xmodify(k) = -absoluteDelta;
            end
        else
            % relative change
            Xmodify(k) = (1 + relativeDelta)*Xmodify(k);
        end
        p(k+1,:) = Xmodify;
        y(k+1) = costFunction(FUN,Xmodify);
    end
    algostep = 'initial simplex';
    nriterations = nriterations + 1;
    nfunk = nfunk + ndim + 1;
    
    % if output function given then run output function to plot
    % intermediate result
    if length(outputFunction) ~= 0,
        feval(outputFunction,p(1,:), y(1),p);
    end
    
    %%%%%%%%%%%%%%%%%%%%%%%%%%%%%%%%%%%%%%%%%%%%
    % MAIN ALGORITHM
    %%%%%%%%%%%%%%%%%%%%%%%%%%%%%%%%%%%%%%%%%%%%
    % reorder y and p so that the first row corresponds to the
    % lowest function value
    for itertemp = 1:MAXITERTEMP,
        % add random thermal fluctuations
        yfluct = y + Temp*abs(log(rand(ndim+1,1)));
        % do sorting instead of determining the indices of the best, worst,
        % next worst
        help = sortrows([yfluct,y,p],1);
        yfluct = help(:,1);
        y = help(:,2);
        p = help(:,3:end);
        if ~silent,
            disp(sprintf(' %5.0f   %5.0f       %12.6g       %12.6g   %12.6g       %s', nriterations, nfunk, y(1), ybest, Temp, algostep));
        end
        % if output function given then run output function to plot
        % intermediate result
        if length(outputFunction) ~= 0,
            feval(outputFunction,p(1,:), y(1),p);
        end
        % end the optimization if the difference between best and worst
        % function evaluation in simplex is smaller than tolfun and the
        % max difference between the coordinates of the verttices is less than
        % tolx
        if abs(max(y)-min(y)) < tolfun && max(max(abs(p(2:ndim+1)-p(1:ndim)))) < tolx,
            break;
        end
        % check number of iterations
        if toc/60 > maxtime,
            EXITFLAG = 0;
            disp('Exceeded maximum time.');
            break;
        end
        if stopOptimization == 1,
            disp('User Interrupt.');
            break;
        end
        % Begin a new iteration. First extrapolate by a factor -1 through the face of the simplex
        % across from the high point, i.e., reflect the simplex from the high point.
        [yftry, ytry,ptry] = amotry(FUN, p, -1);
        % check the result
        if yftry <= yfluct(1),
            % Gives a result better than the best point, so try an additional
            % extrapolation by a factor 2.
            [yftryexp, ytryexp,ptryexp] = amotry(FUN, p, -2);
            if yftryexp < yftry,
                p(end,:) = ptryexp;
                y(end) = ytryexp;
                algostep = 'extrapolation';
            else
                p(end,:) = ptry;
                y(end) = ytry;
                algostep = 'reflection';
            end
        elseif yftry >= yfluct(ndim),
            % The reflected point is worse than the second-highest, so look
            % for an intermediate lower point, i.e., do a one-dimensional
            % contraction.
            [yftrycontr,ytrycontr,ptrycontr] = amotry(FUN, p, -0.5);
            if yftrycontr < yfluct(end),
                p(end,:) = ptrycontr;
                y(end) = ytrycontr;
                algostep = 'one dimensional contraction';
            else
                % Can�t seem to get rid of that high point. Better contract
                % around the lowest (best) point.
                x = ones(ndim,ndim)*diag(p(1,:));
                p(2:end,:) = 0.5*(p(2:end,:)+x);
                for k=2:ndim,
                    y(k) = costFunction(FUN,p(k,:));
                end
                algostep = 'contraction around best point';
            end
        else
            % if ytry better than second-highest point then use this point
            p(end,:) = ptry;
            y(end) = ytry;
            algostep = 'reflection';
        end
        nriterations = nriterations + 1;
    end
    % Break because of max number iterations, max time, minimum found
    % (tolerances)
    if itertemp < MAXITERTEMP,
        break;
    end
    Temp = Temp*tempfactor;
    % Break because 0 temperature has been run
    if Temp == 0,
        break;
    end
end
X = pbest;
FVAL = ybest;
return

%%%%%%%%%%%%%%%%%%%%%%%%%%%%%%%%%%%%%%%%%%%%
% AMOTRY FUNCTION
%%%%%%%%%%%%%%%%%%%%%%%%%%%%%%%%%%%%%%%%%%%%
function [yftry,ytry,ptry] = amotry(FUN, p, fac)
% Extrapolates by a factor fac through the face of the simplex across from 
% the high point, tries it, and replaces the high point if the new point is 
% better.
global ndim nfunk Temp lowbounds highbounds tempstart
psum = sum(p(1:ndim,:))/ndim;
ptry = psum*(1-fac) + p(end,:)*fac;

% Deal with low and high parameter bounds
indexXhi = find(ptry > highbounds);
indexXlo = find(ptry < lowbounds);
for k=1:length(indexXhi),
    ptry(indexXhi(k)) = highbounds(indexXhi(k))-rand(1)*(highbounds(indexXhi(k))-lowbounds(indexXhi(k)))*Temp/tempstart;
end
for k=1:length(indexXlo),
    ptry(indexXlo(k)) = lowbounds(indexXlo(k))+rand(1)*(highbounds(indexXlo(k))-lowbounds(indexXlo(k)))*Temp/tempstart;
end

% Evaluate the function at the trial point.
ytry = costFunction(FUN,ptry);
yftry = ytry - Temp*abs(log(rand(1)));
nfunk = nfunk + 1;
return

%%%%%%%%%%%%%%%%%%%%%%%%%%%%%%%%%%%%%%%%%%%%
% COST FUNCTION EVALUATION
%%%%%%%%%%%%%%%%%%%%%%%%%%%%%%%%%%%%%%%%%%%%
function [ytry] = costFunction(FUN,ptry)
global ybest pbest lowbounds highbounds
ytry = feval(FUN,ptry);
% save the best point ever (only if it is feasible, that is it fits the
% high and low bounds)
indexXhi = find(ptry > highbounds);
indexXlo = find(ptry < lowbounds);
if ytry < ybest && isempty(indexXhi) && isempty(indexXlo),
    ybest = ytry;
    pbest = ptry;
end    
return
```
