## Supplementary material for "A systems biology analysis of adrenergically stimulated adiponectin exocytosis in white adipocytes": All scripts used in this work: simplexIQM.m

```
function [X,FVAL,EXITFLAG] = simplexIQM(varargin)
% simplexIQM: Downhill Simplex Method in Multidimensions. Algorithm based on
% section 10.4 in "Numerical Recipes in C", ISBN 0-521-43108-5.
% The algorithm has been modified in order to be able to take into account
% simple box constraints on parameter values. 
%
% If global variable stopOptimization is set to 1, the optimization is
% stopped. This allows for stopping parameter estimation functions on user
% request. 
% 
% USAGE:
% ======
% [info] = simplexIQM()
% [X,FVAL,EXITFLAG] = simplexIQM(FUN,X,OPTIONS)
%
% FUN: Function to optimize
% X: Starting Guess
% OPTIONS: structure containing options for the algorithm:
%        OPTIONS.maxfunevals: Maximum number of function evaluations
%        OPTIONS.maxiter: Maximum number of iterations
%        OPTIONS.maxtime: Maximum time (in minutes) for optimization
%        OPTIONS.tolfun: Tolerance for difference between best and worst
%                        function evaluation in simplex
%        OPTIONS.tolx: Tolerance for max difference between the coordinates of
%                      the vertices.
%        OPTIONS.highbounds: vector containing upper bounds for parameters
%           Instead of a vector highbounds can also be a scalar > 1. In the
%           latter case the highbounds are determined by multiplying this
%           scalar to the initial parameter guess X. This latter case works
%           only for positive X(i) ... if negative present then the user
%           needs to specify a whole highbounds vector. 
%           If argument is empty ([]) then highbounds are set to +Inf;
%        OPTIONS.lowbounds: vector containing lower bounds for parameters.
%           Instead of a vector lowbounds can also be a scalar < 1. In the
%           latter case the lowbounds are determined by multiplying this
%           scalar to the initial parameter guess X. This latter case works
%           only for positive X(i) ... if negative present then the user
%           needs to specify a whole lowbounds vector. 
%           If argument is empty ([]) then lowbounds are set to -Inf;
%        OPTIONS.outputFunction: string with output function name. If
%               not given or if empty then no output function will be used.
%               This output function can be used to display data, or to 
%               save optimization information to a file. The function needs
%               to be in the MATLAB path. Its calling syntax is:
%                   'outputFunction'(bestparameters,bestfunctionvalue,currentsimplex)
%        OPTIONS.silent: =0: output of info, =1: no output
%
% DEFAULT VALUES:
% ===============
% OPTIONS.maxfunevals:    200*numberVariables
% OPTIONS.maxiter:        200*numberVariables
% OPTIONS.maxtime:        120
% OPTIONS.tolx:           1e-10
% OPTIONS.tolfun:         1e-10
% OPTIONS.lowbounds:      1e-3  => lowbounds = 1e-3*X 
% OPTIONS.highbounds:     1e3   => highbounds = 1e3*X 
% OPTIONS.outputFunction: no output function ('')
% OPTIONS.silent:         0 (no output of info)
%
% Output Arguments:
% =================
% info: calling the function w/o input argument returns information about
%       the options and a flag indicating if the algorithm can handle
%       constraints or not
% X: Found solution
% FVAL: Value of the function FUN at X
% EXITFLAG: 1=success, 0=not found

% <<<COPYRIGHTSTATEMENT - IQM TOOLS LITE>>>

%%%%%%%%%%%%%%%%%%%%%%%%%%%%%%%%%%%%%%%%%%%%
% GLOBAL VARIABLES
%%%%%%%%%%%%%%%%%%%%%%%%%%%%%%%%%%%%%%%%%%%%
global ndim nfunk Xguess lowbounds highbounds stopOptimization

%%%%%%%%%%%%%%%%%%%%%%%%%%%%%%%%%%%%%%%%%%%%
% VARIABLE INPUT ARGUMENTS
%%%%%%%%%%%%%%%%%%%%%%%%%%%%%%%%%%%%%%%%%%%%
if nargin == 0,
    X = [];
    X.name = 'simplexIQM';
    X.constrained = 1;
    X.description = 'Nelder-Mead nonlinear simplex (local)';    
    X.defaultOptions.names = {'maxfunevals', 'maxiter', 'tolfun', 'tolx'};
    X.defaultOptions.values = {'50000','20000','1e-10','1e-10'};
    X.defaultOptions.description = {'Maximum number of function evaluations', 'Maximum number of iterations', 'Termination tolerance on the function value', 'Termination tolerance on X'};
    FVAL = [];
    EXITFLAG = [];
    return
elseif nargin == 2,
    FUN = varargin{1};
    X = varargin{2};
    OPTIONS = [];
elseif nargin == 3,
    FUN = varargin{1};
    X = varargin{2};
    OPTIONS = varargin{3};
else
    error('Incorrect number of input arguments.');
end

%%%%%%%%%%%%%%%%%%%%%%%%%%%%%%
% Set default values
%%%%%%%%%%%%%%%%%%%%%%%%%%%%%%
EXITFLAG = 1;
ndim = length(X);
maxfunevals = 200*ndim;
maxiter = 200*ndim;
maxtime = 120;
tolx = 1e-10;
tolfun = 1e-10;
outputFunction = '';
lowbounds = 1e-3*X;
highbounds = 1e3*X;
silent = 0;

%%%%%%%%%%%%%%%%%%%%%%%%%%%%%%
% Handle OPTIONS
%%%%%%%%%%%%%%%%%%%%%%%%%%%%%%
% silent
if isfield(OPTIONS,'silent'),
    if ~isempty(OPTIONS.silent),
        silent = OPTIONS.silent;
    end
end
% tolfun
if isfield(OPTIONS,'tolfun'),
    if ~isempty(OPTIONS.tolfun),
        tolfun = OPTIONS.tolfun;
    end
end
% tolx
if isfield(OPTIONS,'tolx'),
    if ~isempty(OPTIONS.tolx),
        tolx = OPTIONS.tolx;
    end
end
% maxiter
if isfield(OPTIONS,'maxtime'),
    if ~isempty(OPTIONS.maxtime),
        maxtime = OPTIONS.maxtime;
    end
end
% maxiter
if isfield(OPTIONS,'maxiter'),
    if ~isempty(OPTIONS.maxiter),
        maxiter = OPTIONS.maxiter;
    end
end
% maxfunevals
if isfield(OPTIONS,'maxfunevals'),
    if ~isempty(OPTIONS.maxfunevals),
        maxfunevals = OPTIONS.maxfunevals;
    end
end
% outputFunction
outputFunction = '';
if isfield(OPTIONS,'outputFunction'),
    if ~isempty(OPTIONS.outputFunction),
        outputFunction = OPTIONS.outputFunction;
    end
end
% low and highbounds:
[lowbounds, highbounds] = handleLowHighBoundsIQM(OPTIONS,X,lowbounds,highbounds);

%%%%%%%%%%%%%%%%%%%%%%%%%%%%%%%%%%%%%%%%%%%%
% INITIALIZE EMPTY SIMPLEX DATA
%%%%%%%%%%%%%%%%%%%%%%%%%%%%%%%%%%%%%%%%%%%%
p = zeros(ndim+1,ndim);     % vertice vectors in rows
y = zeros(ndim+1,1);

%%%%%%%%%%%%%%%%%%%%%%%%%%%%%%%%%%%%%%%%%%%%
% FIRST POINT OF INITIAL SIMPLEX
%%%%%%%%%%%%%%%%%%%%%%%%%%%%%%%%%%%%%%%%%%%%
Xguess = X(:)'; 
p(1,:) = Xguess;    
y(1) = feval(FUN,Xguess);

if ~silent,
    disp(' Nr Iter    Nr Fun Eval         Min function          Algorithm Step');
    disp(sprintf(' %5.0f        %5.0f           %12.6g            %s', 0, 1, y(1), 'initial guess'));
end

%%%%%%%%%%%%%%%%%%%%%%%%%%%%%%%%%%%%%%%%%%%%
% RESTART MODIFICATION
%%%%%%%%%%%%%%%%%%%%%%%%%%%%%%%%%%%%%%%%%%%%
finished = 0;
ndimmult = 20;  % factor for ndim to get the number of iterations to do until restart (adapted later)
OLDmeanimprovement = 1/eps;
nriterations = 1;
nfunk = ndim+1;

while ~finished,
    %%%%%%%%%%%%%%%%%%%%%%%%%%%%%%%%%%%%%%%%%%%%
    % REMAINING POINTS OF INITIAL SIMPLEX
    %%%%%%%%%%%%%%%%%%%%%%%%%%%%%%%%%%%%%%%%%%%%
    % construct vertices by modifying one element each
    % relative changes in case that elements are non-zero,
    % absolute changes in case that elements are zero
    relativeDelta = 0.05;
    absoluteDelta = 0.00025;
    for k = 1:ndim
        Xmodify = p(1,:);
        if Xmodify(k) == 0
            % absolute change
            if highbounds(k) > absoluteDelta,
                Xmodify(k) = absoluteDelta;
            else
                Xmodify(k) = -absoluteDelta;
            end
        else
            % relative change
            Xmodify(k) = (1 + relativeDelta)*Xmodify(k);
        end
        p(k+1,:) = Xmodify;
        y(k+1) = feval(FUN,Xmodify);
    end
    algostep = 'initial simplex';

    % if output function given then run output function to plot
    % intermediate result
    if ~isempty(outputFunction),
        feval(outputFunction,p(1,:), y(1),p);
    end

    %%%%%%%%%%%%%%%%%%%%%%%%%%%%%%%%%%%%%%%%%%%%
    % MAIN ALGORITHM
    %%%%%%%%%%%%%%%%%%%%%%%%%%%%%%%%%%%%%%%%%%%%
    % reorder y and p so that the first row corresponds to the
    % lowest function value
    tic % start timer
counter = 1;    
fvalues_history = [];    
    while(1),
        % do sorting instead of determining the indices of the best, worst,
        % next worst
        help = sortrows([y,p],1);
        y = help(:,1);
        p = help(:,2:end);
        if ~silent,
            disp(sprintf(' %5.0f        %5.0f           %12.6g            %s', nriterations, nfunk, y(1), algostep));
        end
        
%%%%%%%%%%%%%%%%%%%%%%%%%%%%%%%%%%%%%%%%%%%%%%%%%
% HANDLE THE RESTART OF THE SIMPLEX
%%%%%%%%%%%%%%%%%%%%%%%%%%%%%%%%%%%%%%%%%%%%%%%%%
% collect information about the improvement in the last 2*ndim iterations
fvalues_history(end+1) = y(1);
if counter > ndimmult*ndim,
    meanimprovement = mean(fvalues_history(end-2*ndim:end-1)-fvalues_history(end-2*ndim+1:end));
    if OLDmeanimprovement-100*eps(OLDmeanimprovement) >= meanimprovement,
        ndimmult = ndimmult*1.5;
%        disp('Restart later')
    else
        ndimmul = ndimmult*0.9;
%        disp('Restart earlier');
    end   
    OLDmeanimprovement = meanimprovement;
    finished = 0; % make a restart at current optimum
    break;
end
counter = counter + 1;
%%%%%%%%%%%%%%%%%%%%%%%%%%%%%%%%%%%%%%%%%%%%%%%%%
% HANDLE THE RESTART OF THE SIMPLEX - END
%%%%%%%%%%%%%%%%%%%%%%%%%%%%%%%%%%%%%%%%%%%%%%%%%
        
        % if output function given then run output function to plot
        % intermediate result
        if ~isempty(outputFunction),
            feval(outputFunction,p(1,:), y(1),p);
        end
        % end the optimization if the difference between best and worst
        % function evaluation in simplex is smaller than tolfun and the
        % max difference between the coordinates of the verttices is less than
        % tolx
        if abs(y(end)-y(1)) < tolfun && max(max(abs(p(2:ndim+1)-p(1:ndim)))) < tolx,
            finished = 1; % exit also the restart loop
            break;
        end
        % check number of function evaluations
        if nfunk >= maxfunevals,
            EXITFLAG = 0;
            if ~silent,
                disp('Exceeded maximum number of function evaluations.');
            end
            finished = 1; % exit also the restart loop
            break;
        end
        % check number of iterations
        if nriterations >= maxiter,
            EXITFLAG = 0;
            if ~silent,
                disp('Exceeded maximum number of iterations.');
            end
            finished = 1; % exit also the restart loop
            break;
        end
        if toc/60 > maxtime,
            EXITFLAG = 0;
            if ~silent,
                disp('Exceeded maximum time.');
            end
            finished = 1; % exit also the restart loop
            break;
        end
        if stopOptimization == 1,
            EXITFLAG = 0;
            disp('User Interrupt.');
            finished = 1; % exit also the restart loop
            break;
        end
        % Begin a new iteration. First extrapolate by a factor -1 through the face of the simplex
        % across from the high point, i.e., reflect the simplex from the high point.
        [ytry,ptry] = amotry(FUN, p, y, -1);
        % check the result
        if ytry <= y(1),
            % Gives a result better than the best point, so try an additional
            % extrapolation by a factor 2.
            [ytryexp,ptryexp] = amotry(FUN, p, y, -2);
            if ytryexp < ytry,
                p(end,:) = ptryexp;
                y(end) = ytryexp;
                algostep = 'extrapolation';
            else
                p(end,:) = ptry;
                y(end) = ytry;
                algostep = 'reflection';
            end
        elseif ytry >= y(ndim),
            % The reflected point is worse than the second-highest, so look
            % for an intermediate lower point, i.e., do a one-dimensional
            % contraction.
            [ytrycontr,ptrycontr] = amotry(FUN, p, y, -0.5);
            if ytrycontr < y(end),
                p(end,:) = ptrycontr;
                y(end) = ytrycontr;
                algostep = 'one dimensional contraction';
            else
                % Can�t seem to get rid of that high point. Better contract
                % around the lowest (best) point.
                x = ones(ndim,ndim)*diag(p(1,:));
                p(2:end,:) = 0.5*(p(2:end,:)+x);
                for k=2:ndim,
                    y(k) = feval(FUN,p(k,:));
                    nfunk = nfunk+1;
                end
                algostep = 'contraction around best point';
            end
        else
            % if ytry better than second-highest point then use this point
            p(end,:) = ptry;
            y(end) = ytry;
            algostep = 'reflection';
        end
        nriterations = nriterations + 1;
    end
    % do the sorting a last time
    help = sortrows([y,p],1);
    y = help(:,1);
    p = help(:,2:end);
end
%%%%%%%%%%%%%%%%%%%%%%%%%%%%%%%%%%%%%%%%%%%%
% RESTART MODIFICATION END
%%%%%%%%%%%%%%%%%%%%%%%%%%%%%%%%%%%%%%%%%%%%

X = p(1,:);
FVAL = y(1);
return

%%%%%%%%%%%%%%%%%%%%%%%%%%%%%%%%%%%%%%%%%%%%
% AMOTRY FUNCTION
%%%%%%%%%%%%%%%%%%%%%%%%%%%%%%%%%%%%%%%%%%%%
function [ytry,ptry] = amotry(FUN, p, y, fac)
% Extrapolates by a factor fac through the face of the simplex across from 
% the high point, tries it, and replaces the high point if the new point is 
% better.
global ndim nfunk lowbounds highbounds
psum = sum(p(1:ndim,:))/ndim;
ptry = psum*(1-fac) + p(end,:)*fac;

% Deal with low and high parameter bounds
indexXhi = find(ptry > highbounds);
indexXlo = find(ptry < lowbounds);
ptry(indexXhi) = highbounds(indexXhi);
ptry(indexXlo) = lowbounds(indexXlo);

% Evaluate the function at the trial point.
ytry = feval(FUN,ptry);
nfunk = nfunk + 1;
return
```
