## Supplementary material for "A systems biology analysis of adrenergically stimulated adiponectin exocytosis in white adipocytes": All scripts used in this work: IQMbarplotErrors.m

```
function [bar_xtick,hb,he]=IQMbarplotErrors(bar_input,errorbar_input,varargin)
% IQMbarplotErrors: grouped bar plot with error bars, customizable.
%
% IQMbarplotErrors(bar_input,errorbar_input) produces a bar plot of matrix
% bar_input. Each column corresponds to one group and each element in a
% column to one bar. The bars are
% grouped together, similar to the plot produced by BAR(Y,'grouped').
%
% If a row vector is passed, it is converted to a column vector to avoid
% grouping => single group then also used.
%
% It then overlays an error bar plot of errorbar_input, the size of which
% must match that of bar_input. Error bars are assumed to be symmetric
% around the values of the bar plot, similar to the plot produced by
% ERRORBAR(Y,E). Note that, if all the values of errorbar_input are set to
% zero, the function skips plotting the error bars.
%
% IQMbarplotErrors(bar_input,errorbar_lower,errorbar_upper) allows
% the lower and upper bounds of the error bars to be asymmetric around the
% values of the bar plot, similar to the plot produced by ERRORBAR(Y,L,U).
%
% Note that it is impossible to input the X coordinates of the bars using
% this function.
%
% bar_xtick = IQMbarplotErrors(...) returns the X coordinates for the center
% of each group of bars.
%
% [...,hb,he] = IQMbarplotErrors(...) returns the handles to the bars
% (produced by BAR) and the error bars (produced by ERRORBAR).
%
% IQMbarplotErrors(...,'ParameterName',ParameterValue) allows customizing
% the display.
%
% 'bar_width': scalar determining bar width. Must be between 0 and 1. A
% bar_width value of 1 causes all the bars to touch each other. This
% parameter is identical to the optional width parameter of BAR. Default
% value: 0.9.
%
% 'errorbar_width': scalar determining error bar width, expressed as a
% fraction of the width of the bars themselves. Default value: 0.75.
%
% 'bar_colors': N-by-3 matrix determing the RGB values for bar colors. You
% must provide at least as many colors as there are groups in your plot.
% Default values: from IQMgetcolors()
%
% 'errorbar_colors': N-by-3 matrix determining the RGB values for error bar
% colors. You must provide at least as many colors as there are groups in
% your plot. Default value: the error bar colors are set to black.
%
% 'optional_bar_arguments': cell array containing any 'PropertyName' -
% 'PropertyValue' input argument pair that you would like to pass on to
% BAR. Default value: no further input arguments are passed to BAR.
%
% 'optional_errorbar_arguments': cell array containing any 'PropertyName' -
% 'PropertyValue' input argument pair that you would like to pass on to
% ERRORBAR. Default value: {'LineStyle','none','Marker','none'}
% (causes ERRORBAR to plot no line between error bars, and no marker at the
% center of each error bar)
%
% 'bar_names': cell string array containing labels to apply to each group
% of bars. You must provide at least as many labels as there are bars in
% your plot. Default value: the bars are numbered consecutively, starting
% from 1.
%
% Examples:
%
% Basic usage:
%   bar_input=rand(3,8)/2+0.5;
%   errorbar_input=rand(3,8)/8;
%   IQMbarplotErrors(bar_input,errorbar_input);
%
% Set the lower bound of the error bars to 0, effectively plotting only the
% upper bound:
%   bar_input=rand(4,6)/2+0.5;
%   errorbar_lower=zeros(size(bar_input));
%   errorbar_upper=rand(4,6)/8;
%   IQMbarplotErrors(bar_input,errorbar_lower,errorbar_upper);
%
% When plotting fewer groups and bars, the plot might look better with
% thinner bars and error bars. This also shows how to input custom names
% for the groups of bars:
%   bar_input=rand(2,4)/2+0.5;
%   errorbar_input=rand(2,4)/8;
%   IQMbarplotErrors(bar_input,errorbar_input, ...
%       'bar_width',0.75,'errorbar_width',0.5, ...
%       'bar_names',{'A','B','C','D'});
%
% Here is how to pass optional input arguments to BAR and ERRORBAR:
%   bar_input=rand(2,4)/2+0.5;
%   errorbar_input=rand(2,4)/8;
%   IQMbarplotErrors(bar_input,errorbar_input, ...
%       'bar_width',0.75,'errorbar_width',0.5, ...
%       'optional_bar_arguments',{'LineWidth',1.5}, ...
%       'optional_errorbar_arguments',{'LineStyle','none','Marker','none','LineWidth',1.5});

% Author of this function:
% Changes to work with R2014B by Henning Schmidt
%
% change log:
% 2014/07/16 -- first uploaded to the MATLAB File Exchange. Does not
% do anything with the 'grp_names' optional input yet.
% 2014/07/18 -- The x axis is now adjusted to fit the plot.
% 2014/07/28 -- With David Groppe's help, added optional figure and axes
% handles.
% 2014/08/19 -- Simplified how default parameters are defined. Added option
% to skip plotting the error bars altogether.

% Make column vector if row vectors
if isvector(bar_input),
    bar_input = bar_input(:);
end
if isvector(errorbar_input),
    errorbar_input = errorbar_input(:);
end

% Interface to function by transposing
bar_input = bar_input';
errorbar_input = errorbar_input';

% init defaults for parameters
[N_grps,N_bars]             = size(bar_input);
bar_width                   = 0.8;
errorbar_width              = 0.75;
bar_colors                  = IQMgetcolors();

% get some distinguishable colors for the bars!
errorbar_colors             = zeros(N_grps,3); % default errorbar color is black
optional_bar_arguments      = {}; % there are no default optional arguments for bar
optional_errorbar_arguments = {'LineStyle','none','Marker','none'}; % default optional arguments for errorbar
bar_names                   = strtrim(cellstr(num2str((1:N_bars)')));
grp_names                   = strtrim(cellstr(num2str((1:N_grps)')));

% deal with the input arguments
if nargin<2 % the indispensable input arguments are not provided
    error('You need to provide at least ''bar_input'' and ''errorbar_input''.');
else
    errorbar_lower=errorbar_input;
    errorbar_upper=errorbar_input;
    if any(size(bar_input)~=size(errorbar_input)) % the indispensable input arguments must have the exact same size
        error('The size of ''bar_input'' and ''errorbar_input'' must be the same.');
    else
        if numel(varargin)>0 % optional input arguments are provided
            if ~ischar(varargin{1}) % if the first optional input argument is not a character array, then by design it must be the upper bound of the error bars, or an error is thrown
                errorbar_lower=errorbar_input;
                errorbar_upper=varargin{1};
                % Adjust errorbar_upper as the other two input arguments above
                if isvector(errorbar_upper),
                    errorbar_upper = errorbar_upper(:);
                end
                % Interface to function by transposing
                errorbar_upper = errorbar_upper';
                if any(size(errorbar_lower)~=size(errorbar_upper))
                    error('The size of ''errorbar_input_low'' and ''errorbar_input_high'' must be the same.');
                end
                varargin(1)=[]; % the first optional argument has been dealt with -- remove...
            end
            while ~isempty(varargin)
                if numel(varargin)<2
                    error('You need to provide optional input arguments as ''ParameterName''-''ParameterValue'' pairs.');
                end
                switch varargin{1}
                    case 'bar_width'
                        bar_width=varargin{2};
                    case 'errorbar_width'
                        errorbar_width=varargin{2};
                    case 'bar_colors'
                        bar_colors=varargin{2};
                    case 'errorbar_colors'
                        errorbar_colors=varargin{2};
                    case 'optional_bar_arguments'
                        optional_bar_arguments=varargin{2};
                    case 'optional_errorbar_arguments'
                        optional_errorbar_arguments=varargin{2};
                    case 'bar_names'
                        bar_names=varargin{2};
                    case 'grp_names'
                        grp_names=varargin{2};
                    otherwise
                        error('Unexpected ''ParameterName'' input: %s\n',varargin{1});
                end
                varargin(1:2)=[]; % this pair of optional input arguments has been dealt with -- remove...
            end
        end
    end
end

% init group width and bar shift
shift_span                  = (1-bar_width)*(N_grps-1);
bar_shift                   = linspace(-shift_span/2,+shift_span/2,N_grps);

% compute position of group x ticks
bar_xtick                   = N_grps/2+0.5:N_grps:N_bars*N_grps-N_grps/2+0.5;

% init handles vectors
hb                          = zeros(N_grps,1);
he                          = zeros(N_grps,1);

% Clear current axes ... or create new figure
cla
hold on;

% plot the bars themselves
for grp=1:N_grps
    hb(grp)=bar( ...
        (grp:N_grps:N_bars*N_grps-(N_grps-grp))-bar_shift(grp), ... % this is the x position for each bar
        bar_input(grp,:),  ... % this is the y position for each bar
        bar_width/N_grps, ... % this is the width of each bar
        'FaceColor',bar_colors(grp,:), ... % color parameter
        optional_bar_arguments{:}); % extra parameters
end

% plot the error bars
if ~all(all(errorbar_lower==0))&&~all(all(errorbar_upper==0))
    
    for grp=1:N_grps
        he(grp)=errorbar( ...
            (grp:N_grps:N_bars*N_grps-(N_grps-grp))-bar_shift(grp), ... % this is the x position for each bar
            bar_input(grp,:),  ... % this is the y position for each bar
            errorbar_lower(grp,:), ... % this is the error low value for each bar
            errorbar_upper(grp,:), ... % this is the error high value for each bar
            'Color',errorbar_colors(grp,:), ... % color parameter
            optional_errorbar_arguments{:}); % extra parameters
    end
        
    % Set the errorbar widths    
    if verLessThan('matlab', '8.4'),
        % Before R2014B
        he_c=get(he,'Children');
        if ~iscell(he_c)
            temp=he_c;
            he_c=cell(1,1);
            he_c{1}=temp;
            clear temp;
        end
        for grp=1:N_grps
            he_xdata=get(he_c{grp}(2),'XData');
            he_xdata(4:9:end)=he_xdata(1:9:end)-errorbar_width*bar_width/2;
            he_xdata(7:9:end)=he_xdata(1:9:end)-errorbar_width*bar_width/2;
            he_xdata(5:9:end)=he_xdata(1:9:end)+errorbar_width*bar_width/2;
            he_xdata(8:9:end)=he_xdata(1:9:end)+errorbar_width*bar_width/2;
            set(he_c{grp}(2),'XData',he_xdata);
        end
    else
        % From R2014B
        % Keep as is for now ... can not find in the objects
    end
end

% set the x tick labels
set(gca,'XTick',bar_xtick,'XTickLabel',bar_names);

% cosmetic fine-tuning of the figure
set(gca,'XLim',[0 bar_xtick(end)+bar_xtick(1)]); % adjusts the x axis to the plot
hold off;
```
