## Supplementary material for "A systems biology analysis of adrenergically stimulated adiponectin exocytosis in white adipocytes": All scripts used in this work: IQMplotCatCat.m

```
function [] = IQMplotCatCat(data,catNames,compareNames,options)
% This function plots a "bubble" plot for categorical data. This allows to
% at least graphically explore correlations between different categorical
% covariates. Areas of circles are proportional to the fraction.
%
% There are 2 cases. Either all categorical covariates are compared to each
% other (only catNames provided). Or categorical covariates are compared to
% categorical covariates, defined by compareNames.
%
% [SYNTAX]
% [] = IQMplotCatCat(data,catNames)
% [] = IQMplotCatCat(data,catNames,compareNames)
% [] = IQMplotCatCat(data,catNames,compareNames,options)
%
% [INPUT]
% data:         Matlab dataset. Each column corresponds to a variable
%               and each row to a sample. The columns with the names
%               defined in "catNames" need to be present in
%               the dataset and contain numerical categorical data.
% catNames:     Cell-array with names of categorical variables
% compareNames: Cell-array with names of categorical variables to compare
%               with the categorical covariates in catNames
% options:      Matlab structure with additional options
%       options.percent: Show percentages instead of numbers. Percentages
%                        are calculated relative to the number in the bins
%                        of the catNames and only displayed if compared to
%                        compareNames. =0 do show numbers and percent. =1 show
%                        percent (default) =2 show numbers and percent.
%       options.bubbleSize: Factor to change size of bubbles (default: 500)
%       options.fontSizeText: Fontsize for the number and percent text
%                             (default: 10)
%       options.hideFirstRow: Hides first row with bars in case
%                             compareNames is used. =0: not hide (default),
%                             =1: hide (default)
%       options.color:  Color of bubbles default: 0.4*[1 1 1]
%
% [OUTPUT]
% Plot

% <<<COPYRIGHTSTATEMENT - IQM TOOLS LITE>>>

% Check if data is table
if ~istable(data),
    error('Input data needs to be provided as MATLAB table.');
end

%%%%%%%%%%%%%%%%%%%%%%%%%%%%%%%%%%%%%%%%%%%%%%%%%%%%%%%%%%%%%%%%%%%%%%%%%%%
% Check which case it is and handle each case completely separately
%%%%%%%%%%%%%%%%%%%%%%%%%%%%%%%%%%%%%%%%%%%%%%%%%%%%%%%%%%%%%%%%%%%%%%%%%%%
if nargin==2,
    compareNames = {};
    options = [];
end
if nargin==3,
    options = [];
end

try percentFlag = options.percent; catch, percentFlag = 1; end
try bubbleSize  = options.bubbleSize; catch, bubbleSize  = 500; end
try fontSizeText  = options.fontSizeText; catch, fontSizeText  = 10; end
try hideFirstRow  = options.hideFirstRow; catch, hideFirstRow  = 0; end
try color  = options.color; catch, color = 0.4*[1 1 1]; end

% Subindex properties
Spacing = 0.0005;
Padding = 0.001;
Margin  = .1;

% Define fontsize
if length(catNames)<=6,
    FONTSIZE = 10;
else
    FONTSIZE = 10;
end

if isempty(compareNames),
    
    %%%%%%%%%%%%%%%%%%%%%%%%%%%%%%%%%%%%%%%%%%%%%%%%%%%%%%%%%%%%%%%%%%%%%%%%%%%
    % Handle first case (compare all catNames with each other)
    %%%%%%%%%%%%%%%%%%%%%%%%%%%%%%%%%%%%%%%%%%%%%%%%%%%%%%%%%%%%%%%%%%%%%%%%%%%
    
    %%%%%%%%%%%%%%%%%%%%%%%%%%%%%%%%%%%%%%%%
    % Check if dataset contains defined columns
    %%%%%%%%%%%%%%%%%%%%%%%%%%%%%%%%%%%%%%%%
    for k=1:length(catNames),
        try
            data.(catNames{k});
        catch
            error(sprintf('Please check if "%s" is a column in the dataset!',catNames{k}));
        end
    end
    
    %%%%%%%%%%%%%%%%%%%%%%%%%%%%%%%%%%%%%%%%
    % Keep only selected columns
    %%%%%%%%%%%%%%%%%%%%%%%%%%%%%%%%%%%%%%%%
    data = data(:,catNames);
    
    %%%%%%%%%%%%%%%%%%%%%%%%%%%%%%%%%%%%%%%%
    % If is table then convert to double
    %%%%%%%%%%%%%%%%%%%%%%%%%%%%%%%%%%%%%%%%
    if istable(data),
        data = table2array(data);
    end
    
    %%%%%%%%%%%%%%%%%%%%%%%%%%%%%%%%%%%%%%%%
    % Keep only selected columns
    %%%%%%%%%%%%%%%%%%%%%%%%%%%%%%%%%%%%%%%%
       
    clf;
    n = length(catNames);
    for ir=1:n
        ystr = catNames{ir};
        y = data(:,ir);
        for ic=1:ir
            xstr = catNames{ic};
            x = data(:,ic);
            ip = (ir-1)*n+ic;
            subaxis(n,n,ip,'Spacing',Spacing,'Padding',Padding,'Margin',Margin);
            if ir==ic % plot "histogram"
                
                xu = unique(x);
                xu = xu(~isnan(xu));
                b  = zeros(size(xu));
                nu = length(xu);
                for i=1:nu
                    b(i) = sum(x==xu(i));
                end
                h = bar(1:nu,b,0.5);
                
                % Write out numbers
                for knr=1:length(b),
                    aaa = get(gca,'YLim');
                    text(knr,b(knr)+max(aaa)*0.02,sprintf('%d',b(knr)),'VerticalAlign','bottom','HorizontalAlign','center','FontSize',fontSizeText+2);
                end
                
                set(h,'FaceColor',color)
                set(gca,'XLim',[0.5 nu+.5]);
                title(xstr,'Interpreter','none')
                set(gca,'YTick',[]);
                set(gca,'XTick',[]);
                set(gca,'YLim',[0 max(b)*1.3]);
                set(gca,'FontSize',fontSizeText);
                              
                if ir==1,
                    ylabel('#','FontSize',8);
                end
                
                if ir==n
                    xlabel(xstr,'Interpreter','none','FontSize',fontSizeText);
                    set(gca,'XTick',1:length(xu))
                    set(gca,'XTickLabel',xu);
                end
                
            else % plot bubble plot
                % Exchange x categories for numbers 1:N
                xu = unique(x);
                iu = {};
                for k=1:length(xu),
                    iu{k} = find(x==xu(k));
                end
                xn = NaN(size(x));
                for k=1:length(iu),
                    xn(iu{k}) = k;
                end
                
                % Exchange y categories for numbers 1:N
                yu = unique(y);
                iu = {};
                for k=1:length(yu),
                    iu{k} = find(y==yu(k));
                end
                yn = NaN(size(y));
                for k=1:length(iu),
                    yn(iu{k}) = k;
                end
                
                % Find unique pairs of xn and yn (these will be plotted in the
                % bubble plot
                [pairs,~,b] = unique([xn yn],'rows');
                
                % Determine number of rows in [xn yn] with these pairs
                nr_pairs = NaN(size(pairs,1),1);
                
                for k=1:size(pairs,1),
                    nr_pairs(k) = sum(b==k);
                end
                
                % Determine relative number of pairs in fraction
                nr_pairs_fraction = nr_pairs/sum(nr_pairs);
                
                % Determine size based on fraction
                use_size = bubbleSize*nr_pairs_fraction;
                
                % Determine default color
                color_use = color(ones(1,length(nr_pairs)),:);
                
                % Plot
                scatter(pairs(:,1),pairs(:,2), use_size, color_use, 'filled', 'MarkerEdgeColor', 'none');
                
                % Annotate
                set(gca,'XLim',[min(xn)-0.5 max(xn)+0.5])
                set(gca,'YLim',[min(yn)-0.5 max(yn)+0.5])
                
                % Write out percentage as text
                for knr=1:size(pairs,1),
                    text(pairs(knr,1)+0.025,pairs(knr,2)+0.2,sprintf('%d',nr_pairs(knr)),'FontSize',FONTSIZE);
                end
                
                set(gca,'XTick',1:length(xu))
                set(gca,'YTick',1:length(yu))
                set(gca,'FontSize',fontSizeText);
                
                if ic==1
                    ylabel(ystr,'Interpreter','none','FontSize',fontSizeText);
                    set(gca,'YTickLabel',yu);
                else
                    set(gca,'YTickLabel',[]);
                end
                
                if ir==n
                    xlabel(xstr,'Interpreter','none','FontSize',fontSizeText);
                    set(gca,'XTickLabel',xu);
                else
                    set(gca,'XTickLabel',[]);
                end
                
                grid on;
                
            end
        end
    end
    
else
    
    %%%%%%%%%%%%%%%%%%%%%%%%%%%%%%%%%%%%%%%%%%%%%%%%%%%%%%%%%%%%%%%%%%%%%%%%%%%
    % Handle second case (compare all catNames with compareNames)
    %%%%%%%%%%%%%%%%%%%%%%%%%%%%%%%%%%%%%%%%%%%%%%%%%%%%%%%%%%%%%%%%%%%%%%%%%%%
    
    %%%%%%%%%%%%%%%%%%%%%%%%%%%%%%%%%%%%%%%%
    % Check if dataset contains defined columns
    %%%%%%%%%%%%%%%%%%%%%%%%%%%%%%%%%%%%%%%%
    for k=1:length(catNames),
        try
            data.(catNames{k});
        catch
            error(sprintf('Please check if "%s" is a column in the dataset!',catNames{k}));
        end
    end
    
    for k=1:length(compareNames),
        try
            data.(compareNames{k});
        catch
            error(sprintf('Please check if "%s" is a column in the dataset!',compareNames{k}));
        end
    end
    
    %%%%%%%%%%%%%%%%%%%%%%%%%%%%%%%%%%%%%%%%
    % Keep only selected columns
    %%%%%%%%%%%%%%%%%%%%%%%%%%%%%%%%%%%%%%%%
    dataCompare = data(:,compareNames);
    data        = data(:,catNames);
    
    %%%%%%%%%%%%%%%%%%%%%%%%%%%%%%%%%%%%%%%%
    % If is table then convert to double
    %%%%%%%%%%%%%%%%%%%%%%%%%%%%%%%%%%%%%%%%
    if istable(data),
        data = table2array(data);
    end
    if istable(dataCompare),
        dataCompare = table2array(dataCompare);
    end
    
    %%%%%%%%%%%%%%%%%%%%%%%%%%%%%%%%%%%%%%%%
    % Options / fontsizes
    %%%%%%%%%%%%%%%%%%%%%%%%%%%%%%%%%%%%%%%%
       
    clf;
    
    %%%%%%%%%%%%%%%%%%%%%%%%%%%%%%%%%%%%%%%%
    % First row is frequencies of catNames
    %%%%%%%%%%%%%%%%%%%%%%%%%%%%%%%%%%%%%%%%
    if hideFirstRow==0,
        ncols = length(catNames);
        nrows = 1+length(compareNames);
        for ir=1:ncols
            ystr = catNames{ir};
            y = data(:,ir);
            xstr = catNames{ir};
            x = data(:,ir);
            ip = ir;
            subaxis(nrows,ncols,ip,'Spacing',Spacing,'Padding',Padding,'Margin',Margin);
            % Plot frequencies
            xu = unique(x);
            xu = xu(~isnan(xu));
            b  = zeros(size(xu));
            nu = length(xu);
            for i=1:nu
                b(i) = sum(x==xu(i));
            end
            h = bar(1:nu,b,0.5);
            grid on
            set(gca,'FontSize',fontSizeText);
            
            % Write out numbers
            for knr=1:length(b),
                aaa = get(gca,'YLim');
                text(knr,b(knr)+max(aaa)*0.02,sprintf('%d',b(knr)),'VerticalAlign','bottom','HorizontalAlign','center','FontSize',fontSizeText+2);
            end
            
            set(h,'FaceColor',color)
            set(gca,'XLim',[0.5 nu+.5]);
            title(xstr,'Interpreter','none')
%             set(gca,'YTick',[]);
            if ir>1,
                set(gca,'YTickLabel','');
            end
            set(gca,'XTick',[]);
            set(gca,'YLim',[0 max(b)*1.3]);
            
            if ir==1,
                ylabel('#','FontSize',8);
            end
        end
        offset = 1;
    else
        ncols = length(catNames);
        nrows = length(compareNames);
        offset = 0;
    end
    
    %%%%%%%%%%%%%%%%%%%%%%%%%%%%%%%%%%%%%%%%
    % Next rows is correlation of catNames with compareNames
    %%%%%%%%%%%%%%%%%%%%%%%%%%%%%%%%%%%%%%%%
    % Cycle through compareNames
    for kcompare=1:length(compareNames),
        ir = kcompare+offset;
        ystr = compareNames{kcompare};
        y = dataCompare(:,kcompare);
        
        % Cycle through covNames
        for kcov=1:length(catNames),
            ic = kcov;
            xstr = catNames{ic};
            x = data(:,ic);
            ip = (ir-1)*ncols+ic;
            subaxis(nrows,ncols,ip,'Spacing',Spacing,'Padding',Padding,'Margin',Margin);
            
            % plot bubble plot
            % Exchange x categories for numbers 1:N
            xu = unique(x);
            iu = {};
            for k=1:length(xu),
                iu{k} = find(x==xu(k));
            end
            xn = NaN(size(x));
            for k=1:length(iu),
                xn(iu{k}) = k;
            end
            
            % Exchange y categories for numbers 1:N
            yu = unique(y);
            iu = {};
            for k=1:length(yu),
                iu{k} = find(y==yu(k));
            end
            yn = NaN(size(y));
            for k=1:length(iu),
                yn(iu{k}) = k;
            end
            
            % Find unique pairs of xn and yn (these will be plotted in the
            % bubble plot
            [pairs,~,b] = unique([xn yn],'rows');
            
            % Determine number of rows in [xn yn] with these pairs
            nr_pairs = NaN(size(pairs,1),1);
            
            for k=1:size(pairs,1),
                nr_pairs(k) = sum(b==k);
            end
            
            % Determine relative number of pairs in fraction
            nr_pairs_fraction = nr_pairs/sum(nr_pairs);
            
            % Determine size based on fraction
            use_size = bubbleSize*nr_pairs_fraction;
            
            % Determine default color
            color_use = color(ones(1,length(nr_pairs)),:);
            
            % Plot
            scatter(pairs(:,1),pairs(:,2), use_size, color_use, 'filled', 'MarkerEdgeColor', 'none');
            
            % Annotate
            set(gca,'XLim',[min(xn)-0.5 max(xn)+0.5])
            set(gca,'YLim',[min(yn)-0.5 max(yn)+0.5])
            
            % Write out numbers as text 
            for knr=1:size(pairs,1),
                textShow = sprintf('%d',nr_pairs(knr));
                text(pairs(knr,1)+0.025,pairs(knr,2)+0.2,textShow,'FontSize',FONTSIZE);
            end
            
            set(gca,'XTick',1:length(xu))
            set(gca,'YTick',1:length(yu))
            set(gca,'FontSize',fontSizeText);
            
            if ic==1
                ylabel(ystr,'Interpreter','none','FontSize',10);
                set(gca,'YTickLabel',yu);
            else
                set(gca,'YTickLabel',[]);
            end
            
            if ir==nrows
                xlabel(xstr,'Interpreter','none','FontSize',fontSizeText);
                set(gca,'XTickLabel',xu);
            else
                set(gca,'XTickLabel',[]);
            end
            
            grid on;
            
        end
    end
    
    % Adjust YLim on each row to same values
    if length(catNames) > 1,
        for krow=1:length(compareNames)+1,
            YLimALL = [];
            for kcol=1:length(catNames),
                subaxis(length(compareNames)+1,length(catNames),(krow-1)*length(catNames)+kcol);
                YLimALL = [YLimALL; get(gca,'YLim')];
            end
            YLimMin = min(YLimALL);
            YLimMax = max(YLimALL);
            for kcol=1:length(catNames),
                subaxis(length(compareNames)+1,length(catNames),(krow-1)*length(catNames)+kcol);
                set(gca,'YLim',[YLimMin(1) YLimMax(2)]);
            end
        end
    end
end
```
