## Supplementary material for "A systems biology analysis of adrenergically stimulated adiponectin exocytosis in white adipocytes": All scripts used in this work: IQMplotCovarianceCat.m

```
function [] = IQMplotCovarianceCat(data,contNames,catNames,options)
% This function plots the covariance relationship between a list of
% continuous variables (contNames) and a list of categorical variables
% (catNames), passed in "data".
%
% [SYNTAX]
% [] = IQMplotCovarianceCat(data,contNames,catNames)
% [] = IQMplotCovarianceCat(data,contNames,catNames,options)
%
% [INPUT]
% data:         Matlab dataset. Each column corresponds to a variable
%               and each row to a sample. The columns with the names
%               defined in "contNames" and "catNames" need to be present in
%               the dataset.
% contNames:    Cell-array with names of continuous variables
% catNames:     Cell-array with names of categorical variables
% options:      MATLAB structure with optional arguments
%
%                   options.LogFlag:   =1 do log transform the variables,
%                                      =0 do not transform (default: 0)
%                   options.fontSizeText: Fontsize for the number and percent text
%                                         (default: 10)
%
% [OUTPUT]
% Plot

% <<<COPYRIGHTSTATEMENT - IQM TOOLS LITE>>>

LogFlag = 0;
try LogFlag = options.LogFlag; catch, end
try fontSizeText = options.fontSizeText; catch, fontSizeText = 10; end

%%%%%%%%%%%%%%%%%%%%%%%%%%%%%%%%%%%%%%%%%%%%%%%%%%%%%%%%%%%%%%%%%%%%%%%%%%%
% Check if dataset contains defined columns
%%%%%%%%%%%%%%%%%%%%%%%%%%%%%%%%%%%%%%%%%%%%%%%%%%%%%%%%%%%%%%%%%%%%%%%%%%%
for k=1:length(contNames),
    try
        data.(contNames{k});
    catch,
        error(sprintf('Please check if "%s" is a column in the dataset!',contNames{k}));
    end
end
for k=1:length(catNames),
    try
        data.(catNames{k});
    catch,
        error(sprintf('Please check if "%s" is a column in the dataset!',catNames{k}));
    end
end

%%%%%%%%%%%%%%%%%%%%%%%%%%%%%%%%%%%%%%%%%%%%%%%%%%%%%%%%%%%%%%%%%%%%%%%%%%%
% Settings for subaxis
%%%%%%%%%%%%%%%%%%%%%%%%%%%%%%%%%%%%%%%%%%%%%%%%%%%%%%%%%%%%%%%%%%%%%%%%%%%
Spacing = 0;
Padding = 0;
Margin  = .1;

%%%%%%%%%%%%%%%%%%%%%%%%%%%%%%%%%%%%%%%%%%%%%%%%%%%%%%%%%%%%%%%%%%%%%%%%%%%
% Do the plotting
%%%%%%%%%%%%%%%%%%%%%%%%%%%%%%%%%%%%%%%%%%%%%%%%%%%%%%%%%%%%%%%%%%%%%%%%%%%
ncat = length(catNames);  %rows are the categorical covariates
ncts = length(contNames);
for icat=1:ncat
    xstr = catNames{icat};
    x = data.(xstr);
    
    if ~isnumeric(x),
        error('Covariate "%s" not of numeric type.',xstr);
    end
    
    % Only plot if x not only NaN
    if ~isempty(find(isnan(x)==0)),
        
        ip=icat;
        subaxis(ncts+1,ncat,ip,'Spacing',Spacing,'Padding',Padding,'Margin',Margin);
        grid on
        
        xu = unique(x);
        xu = xu(~isnan(xu));
        b  = zeros(size(xu));
        nu = length(xu);
        for i=1:nu
            b(i) = sum(x==xu(i));
        end
        h = bar(1:nu,b,0.5);
        set(h,'FaceColor',0.6*[1 1 1])
        set(gca,'XLim',[0.5 nu+.5]);
        set(gca,'FontSize',fontSizeText);
        
        grid on
        
        % Write out numbers
        for knr=1:length(b),
            aaa = get(gca,'YLim');
            text(knr,b(knr)+max(aaa)*0.02,sprintf('%d',b(knr)),'VerticalAlign','bottom','HorizontalAlign','center','FontSize',fontSizeText+2);
        end
        
        title(xstr,'Interpreter','none')
        if icat==1
            ylabel('#','Interpreter','none');
        else
            %             set(gca,'YTick',[]);
            set(gca,'YTickLabel','');
        end
        
        set(gca,'XTick',[]);
        set(gca,'YLim',[0 max(b)*1.3]);
        
        for icts=1:ncts
            ystr = contNames{icts};
            y = data.(ystr);
            
            % Only plot if y not only NaN
            if ~isempty(find(isnan(y)==0)),
                if LogFlag==1
                    y = log(y);
                    ystr = {'log',ystr};
                end
                ip = icat + ncat + (icts-1)*ncat;
                
                subaxis(ncts+1,ncat,ip,'Spacing',Spacing,'Padding',Padding,'Margin',Margin);
                
                xx = unique(x);
                xx = xx(~isnan(xx));
                M  = NaN(length(y),length(xx));
                for i=1:length(xx)
                    M(x==xx(i),i) = y(x==xx(i));
                end
                OPTIONSbox.NumFlag = 0;
                OPTIONSbox.BoxColor = 0.6*[1 1 1];
                OPTIONSbox.BoxWidth = .5;
                OPTIONSbox.MedianWidth = .7;
                OPTIONSbox.OutlierColor = 0.4*[1 1 1];
                OPTIONSbox.OutlierSize  = 5;
                OPTIONSbox.MedianColor = [0 0 0];
                plotboxIQM(M,1:length(xx),OPTIONSbox);
                set(gca,'XLim',[.5 length(xx)+.5]);
                grid on
                set(gca,'FontSize',fontSizeText);
                
                set(gca,'XTick',1:length(xx));
                if icts<ncts
                    set(gca,'XTick',[]);
                else
                    xxx = unique(x);
                    xxx = xxx(~isnan(xxx));
                    set(gca,'XTickLabel',xxx)
                    xlabel(xstr,'Interpreter','none');
                end
                if icat==1
                    ylabel(ystr,'Interpreter','none');
                else
                    %                     set(gca,'YTick',[]);
                    set(gca,'YTickLabel','');
                end
                %                 set(gca,'YTick',[]);
                if min(y)~=max(y),
                    set(gca,'YLim',[min(y) max(y)]);
                else
                    set(gca,'YLim',[min(y)-1 max(y)+1]);
                end
            end
        end
        %         set(gca,'YTick',[]);
    end
end

% Adjust YLim on each row to same values
for krow=1:ncts+1,
    YLimALL = [];
    for kcol=1:ncat,
        subaxis(ncts+1,ncat,(krow-1)*ncat+kcol);
        YLimALL = [YLimALL; get(gca,'YLim')];
    end
    YLimMin = min(YLimALL);
    YLimMax = max(YLimALL);
    for kcol=1:ncat,
        subaxis(ncts+1,ncat,(krow-1)*ncat+kcol);
        set(gca,'YLim',[min(YLimMin) max(YLimMax)]);
    end    
end
```
