## Supplementary material for "A systems biology analysis of adrenergically stimulated adiponectin exocytosis in white adipocytes": All scripts used in this work: IQMplotfacetgrid.m

```
function [] = IQMplotfacetgrid(data,nameX,nameY,nameGroupX,nameGroupY,options)
% This function plots a matrix version of a trellis plot.
%
% [SYNTAX]
% [] = IQMplotfacetgrid(data,nameX,nameY,nameGroupX,nameGroupY)
% [] = IQMplotfacetgrid(data,nameX,nameY,nameGroupX,nameGroupY,options)
%
% [INPUT]
% data:         Matlab dataset
% nameX:        Column name in dataset to plot on X axis
% nameY:        Column name in dataset to plot on Y axis
% nameGroupX:   Column name in dataset to use as group variable for each column. 
% nameGroupY:   Column name in dataset to use as group variable for each row. 
% options:      Structure with optional settings as follows:
%   options.xlabelText          = String with xlabel text. Default: nameX
%   options.ylabelText          = Cell-array with as many entries as groups in nameGroupY
%   options.nameSubGroup        = Column name in dataset for subgrouping within a group/subplot (nicer line drawing and used for moving median line);
%   options.nameColorGroup      = Column name in dataset for coloring a certain group differently
%   options.logX                = 0 if lin, 1 if log X axis (default: 0)
%   options.logY                = 0 or 1, alternatively a vector with as many 0 or 1 as elements in nameGroupY - allows to set axes differentlty for different nameGroupY elements
%   options.linetype            = String with MATLAB linetype (default: '.-')
%   options.linecolor           = Color for lines (default: [0.2 0.2 0.2])
%   options.showmarkers         = 1 shows different linestyles with markers, =0 uses linstyle setting
%   options.markersize          = numeric value for markersizes (default: 10)
%   options.linecolorsCustom    = Matrix with 3 comlumns and arbitrary rows. Defines custom color settings to use for color grouping
%   options.linetypesCustom     = Cell-array with linestyle strings (e.g.: {'x','-.','--'}). If defined it overrides the standard lines from IQMgetcolors (only active if color group selected)
%                                 Only active if "options.showmarkers=1".
%   options.linewidth           = Numeric value 1-5 (default: 1)
%   options.sameaxes            = If 0 (default) all subplot axes will be scaled to best fit the data, if 1 all axes will have same scaling 
%                                 All rows will always be on same Y axes, as will be all columns on same X axes
%   options.showgrid            = 0: no grid, 1: grid (no minor grid lines) (default: 1)
%   options.colortitlebar       = Vector with three values between 0 and 1 to set he titlebar color (default: [1 1 0.8])
%   options.heighttitlebarX     = Height of the titlebar on top in fraction of subplot (default: determined automatically)
%   options.widthtitlebarY      = Width of the titlebar on the right in fraction of subplot (default: determined automatically)
%   options.showmedian           = 0 (default): do not show a moving median line per group, 1: do show it
%   options.showmean            = 0 (default): do not show a moving mean line per group, 1: do show it
%   options.NbinsMedian          = value between 0 and 100 defining the range of data to take into account (default: 15)
%   options.showlegend          = 0: do not show a legend, 1 (default): show a legend for the color grouping
%   options.labeltextsize       = Fontsize for x and y labels (default: 10)
%   options.maxlegendentries    = If more elements in colorgroup then do not show legend (becomes messy). Default: 10
%   options.ticklabeltextsize   = Fontsize for axes number (default: 10)
%   options.legendtextsize      = Fontsize for legend (default: 10)
%   options.nameText            = Column name in dataset (text in column) to display next to datapoints
%   options.nameTextLines       = 1: show lines additional to text (if nameText is defined). 0: do not show lines (if nameText is defined).
%   options.textFontsize        = Fontsize for additional text (default: 10)
%   options.axescolor           = Sets the color of axes and grid (default: [0.2 0.2 0.2])
%   options.windowcolor         = color definition for outside frame of window (default: [1 1 1])
%   options.titlefontsize       = Fontsize for title bars (default: 8)
% 
% [OUTPUT]
% This function creates a new figure. 
%
% [ASSUMPTIONS]
% Data for X and Y axes and all groups needs to be numeric. 
 
% <<<COPYRIGHTSTATEMENT - IQM TOOLS LITE>>>

%%%%%%%%%%%%%%%%%%%%%%%%%%%%%%%%%%%%%%%%%%%%%%%%%%%%%%%%%%%%%%%%%%%%%%%%%%%
% Get default colors and linestyles
%%%%%%%%%%%%%%%%%%%%%%%%%%%%%%%%%%%%%%%%%%%%%%%%%%%%%%%%%%%%%%%%%%%%%%%%%%%
[colorsDefault,linesDefault] = IQMgetcolors();

%%%%%%%%%%%%%%%%%%%%%%%%%%%%%%%%%%%%%%%%%%%%%%%%%%%%%%%%%%%%%%%%%%%%%%%%%%%
% Get optional variables and set defaults if undefined
%%%%%%%%%%%%%%%%%%%%%%%%%%%%%%%%%%%%%%%%%%%%%%%%%%%%%%%%%%%%%%%%%%%%%%%%%%%
try options             = options;                          catch, options              = [];               end;
try xlabelText          = options.xlabelText;               catch, xlabelText           = nameX;            end; %#ok<*CTCH>
try ylabelText          = options.ylabelText;               catch, ylabelText           = NaN;              end; % handled later
try nameSubGroup        = options.nameSubGroup;             catch, nameSubGroup         = '';               end;
try nameColorGroup      = options.nameColorGroup;           catch, nameColorGroup       = '';               end;
try logX                = options.logX;                     catch, logX                 = 0;                end;
try logY                = options.logY;                     catch, logY                 = 0;                end; % Can be vector
try linetype            = options.linetype;                 catch, linetype             = '.-';             end; %#ok<*NASGU>
try linecolor           = options.linecolor;                catch, linecolor            = [0.2 0.2 0.2];    end;
try showmarkers         = options.showmarkers;              catch, showmarkers          = 0;                end;
try markersize          = options.markersize;               catch, markersize           = 10;               end;
try linecolorsCustom    = options.linecolorsCustom;         catch, linecolorsCustom     = colorsDefault;    end;
try linetypesCustom     = options.linetypesCustom;          catch, linetypesCustom      = linesDefault;     end;
try linewidth           = options.linewidth;                catch, linewidth            = 1;                end;
try sameaxes            = options.sameaxes;                 catch, sameaxes             = 1;                end;
try showgrid            = options.showgrid;                 catch, showgrid             = 1;                end;
try colortitlebar       = options.colortitlebar;            catch, colortitlebar        = [1 1 0.8];        end;
try heighttitlebarX     = options.heighttitlebarX;          catch, heighttitlebarX      = NaN;              end;
try widthtitlebarY      = options.widthtitlebarY;           catch, widthtitlebarY       = NaN;              end;
try showmedian          = options.showmedian;               catch, showmedian           = 0;                end;
try showmean            = options.showmean;                 catch, showmean             = 0;                end;
try NbinsMedian         = options.NbinsMedian;              catch, NbinsMedian           = 15;              end;
try showlegend          = options.showlegend;               catch, showlegend           = 1;                end;
try labeltextsize       = options.labeltextsize;            catch, labeltextsize        = 10;               end;
try ticklabeltextsize   = options.ticklabeltextsize;        catch, ticklabeltextsize    = 10;               end;
try legendtextsize      = options.legendtextsize;           catch, legendtextsize       = 10;               end;
try maxlegendentries    = options.maxlegendentries;         catch, maxlegendentries     = 10;               end;
try nameText            = options.nameText;                 catch, nameText             = '';               end;
try nameTextLines       = options.nameTextLines;            catch, nameTextLines        = 1;                end;
try textFontsize        = options.textFontsize;             catch, textFontsize         = 10;               end;
try axescolor           = options.axescolor;                catch, axescolor            = 0.2*[1 1 1];      end;
try windowcolor         = options.windowcolor;              catch, windowcolor          = 1*[1 1 1];        end;
try titlefontsize       = options.titlefontsize;            catch, titlefontsize        = 8;               end

%%%%%%%%%%%%%%%%%%%%%%%%%%%%%%%%%%%%%%%%%%%%%%%%%%%%%%%%%%%%%%%%%%%%%%%%%%%
% Settings for making plots nicer when same axes are chosen
%%%%%%%%%%%%%%%%%%%%%%%%%%%%%%%%%%%%%%%%%%%%%%%%%%%%%%%%%%%%%%%%%%%%%%%%%%%
SpacingHoriz = 0.002;
SpacingVert  = 0.003;
Padding      = 0;
Margin       = 0.1;

%%%%%%%%%%%%%%%%%%%%%%%%%%%%%%%%%%%%%%%%%%%%%%%%%%%%%%%%%%%%%%%%%%%%%%%%%%%
% Get group information
%%%%%%%%%%%%%%%%%%%%%%%%%%%%%%%%%%%%%%%%%%%%%%%%%%%%%%%%%%%%%%%%%%%%%%%%%%%
if isempty(nameGroupX),
    allGroupX = -1;
else
    allGroupX = unique(data.(nameGroupX));
end
if isempty(nameGroupY),
    allGroupY = -1;
else
    allGroupY = unique(data.(nameGroupY));
end
if ~isempty(nameColorGroup),
    allGROUPcolor   = unique(data.(nameColorGroup));
    data.colorgroup = data.(nameColorGroup);
else
    allGROUPcolor   = -1;
    data.colorgroup = -1*ones(height(data),1);
end

%%%%%%%%%%%%%%%%%%%%%%%%%%%%%%%%%%%%%%%%%%%%%%%%%%%%%%%%%%%%%%%%%%%%%%%%%%%
% Adjust logY settings
%%%%%%%%%%%%%%%%%%%%%%%%%%%%%%%%%%%%%%%%%%%%%%%%%%%%%%%%%%%%%%%%%%%%%%%%%%%
if length(logY)==1,
    logY = logY*ones(1,length(allGroupY));
elseif length(logY)~=length(allGroupY),
    error('options.logY needs to be 0 or 1, or a vector with 0 or 1 of same length as number of Y groups.');
end
% If sameaxes, use first setting of logY
if sameaxes,
    logY = logY(1)*ones(1,length(allGroupY));
end

%%%%%%%%%%%%%%%%%%%%%%%%%%%%%%%%%%%%%%%%%%%%%%%%%%%%%%%%%%%%%%%%%%%%%%%%%%%
% Define ylabeltext if not given as option
%%%%%%%%%%%%%%%%%%%%%%%%%%%%%%%%%%%%%%%%%%%%%%%%%%%%%%%%%%%%%%%%%%%%%%%%%%%
if ~iscell(ylabelText),
    if isnan(ylabelText),
        ylabelText = {};
        for k=1:length(allGroupY),
            ylabelText{k} = sprintf('%s (%s=%g)',nameY,nameGroupY,allGroupY(k));
        end
    end
end

%%%%%%%%%%%%%%%%%%%%%%%%%%%%%%%%%%%%%%%%%%%%%%%%%%%%%%%%%%%%%%%%%%%%%%%%%%%
% Define subplot information
%%%%%%%%%%%%%%%%%%%%%%%%%%%%%%%%%%%%%%%%%%%%%%%%%%%%%%%%%%%%%%%%%%%%%%%%%%%
ncols       = length(allGroupX);
nrows       = length(allGroupY);
nSubplots   = nrows*ncols;

%%%%%%%%%%%%%%%%%%%%%%%%%%%%%%%%%%%%%%%%%%%%%%%%%%%%%%%%%%%%%%%%%%%%%%%%%%%
% Handle title bar sizes
%%%%%%%%%%%%%%%%%%%%%%%%%%%%%%%%%%%%%%%%%%%%%%%%%%%%%%%%%%%%%%%%%%%%%%%%%%%
if isnan(heighttitlebarX),
    heighttitlebarX = 0.04+0.02*nrows;
end
if isnan(widthtitlebarY),
    widthtitlebarY = 3/4*heighttitlebarX/nrows*ncols;
    if widthtitlebarY > 0.5,
        widthtitlebarY = 0.5;
    end
end

%%%%%%%%%%%%%%%%%%%%%%%%%%%%%%%%%%%%%%%%%%%%%%%%%%%%%%%%%%%%%%%%%%%%%%%%%%%
% Run through the group X and Y, define subplot settings, color settings
%%%%%%%%%%%%%%%%%%%%%%%%%%%%%%%%%%%%%%%%%%%%%%%%%%%%%%%%%%%%%%%%%%%%%%%%%%%
dataPlot            = table();
for kX=1:length(allGroupX),
    if ~isempty(nameGroupX),
        dataX = subsetIQM(data,nameGroupX,allGroupX(kX));
    else
        dataX = data;
    end
    
    for kY=1:length(allGroupY),
        if ~isempty(nameGroupY),
            dataXY = subsetIQM(dataX,nameGroupY,allGroupY(kY));
        else
            dataXY = dataX;
        end
        
        %%%%%%%%%%%%%%%%%%%%%%%%%%%%%%%%%%%%%%%%%%%%%%%%%%%%%%%%%%%%%%%%%%%%%%%%%%%
        % If log X or Y, remove <=0 data from respective parts
        %%%%%%%%%%%%%%%%%%%%%%%%%%%%%%%%%%%%%%%%%%%%%%%%%%%%%%%%%%%%%%%%%%%%%%%%%%%
        if ~isempty(dataXY),
            if logX == 1,
                dataXY(dataXY.(nameX)<=0,:) = [];
            end
            if logY(kY) == 1,
                dataXY(dataXY.(nameY)<=0,:) = [];
            end
            
            for kC=1:length(allGROUPcolor),
                dataXYC = subsetIQM(dataXY,'colorgroup',allGROUPcolor(kC));
                
                if ~isempty(dataXYC),
                    datatemp = table();
                    datatemp.(nameX)        = dataXYC.(nameX);
                    datatemp.(nameY)        = dataXYC.(nameY);
                    if isnumeric(allGroupX(kX)),
                        datatemp.groupX         = allGroupX(kX)*ones(height(datatemp),1);
                    else
                        datatemp.groupX         = cell(height(datatemp),1);
                        datatemp.groupX(1:end)  = {allGroupX{kX}};
                    end
                    if isnumeric(allGroupY(kY)),
                        datatemp.groupY         = allGroupY(kY)*ones(height(datatemp),1);
                    else
                        datatemp.groupY         = cell(height(datatemp),1);
                        datatemp.groupY(1:end)  = {allGroupY{kY}};
                    end
                    datatemp.plotSubplot    = ( (kY-1)*ncols + kX )*ones(height(datatemp),1);
                    datatemp.colorgroup     = dataXYC.colorgroup;
                    if ~isempty(nameSubGroup),
                        datatemp.subgroup   = dataXYC.(nameSubGroup);
                    else
                        datatemp.subgroup   = ones(height(datatemp),1);
                    end
                    if isnumeric(dataXYC.colorgroup(1)),
                        if dataXYC.colorgroup(1) == -1,
                            datatemp.color      = -1*ones(height(datatemp),1);
                        else
                            datatemp.color      = find(allGROUPcolor==dataXYC.colorgroup(1))*ones(height(datatemp),1);
                        end
                    else
                        datatemp.color      = find(strcmp(allGROUPcolor,dataXYC.colorgroup{1}))*ones(height(datatemp),1);
                    end
                    if ~isempty(nameText),
                        datatemp.(nameText) = dataXYC.(nameText);
                    end
                    dataPlot = [dataPlot; datatemp];
                end
            end
        end
    end
end

%%%%%%%%%%%%%%%%%%%%%%%%%%%%%%%%%%%%%%%%%%%%%%%%%%%%%%%%%%%%%%%%%%%%%%%%%%%
% Determine the x and y axes ranges if sameaxes selected and
% adjust the maxY and maxX part of the axes settings to allow for title bar
% on the X and Y axes.
%%%%%%%%%%%%%%%%%%%%%%%%%%%%%%%%%%%%%%%%%%%%%%%%%%%%%%%%%%%%%%%%%%%%%%%%%%%
minX = min(dataPlot.(nameX));
maxX = max(dataPlot.(nameX));
minY = min(dataPlot.(nameY));
maxY = max(dataPlot.(nameY));
% Adjust maxY for titlebarX
if logY(1)==1,
    rangenew =  1/(1-heighttitlebarX-0.01)*(log10(maxY)-log10(minY));
    maxY     = 10.^(log10(minY) + rangenew);
else
    rangenew =  1/(1-heighttitlebarX-0.01)*(maxY-minY);
    maxY     = minY + rangenew;
end
% Adjust maxX for titlebarY
if logX==1,
    rangenew =  1/(1-widthtitlebarY-0.01)*(log10(maxX)-log10(minX));
    maxX     = 10.^(log10(maxX) + rangenew);
else
    rangenew =  1/(1-widthtitlebarY-0.01)*(maxX-minX);
    maxX     = minX + rangenew;
end
sameAxesLimits = [minX maxX minY maxY];

% Determine the axes limits for each row (for each row it should at
% least be the same) Only for Y axis. X the same as under sameaxes
rowAxesLimits = [];
for krow=1:length(allGroupY),
    datarow = subsetIQM(dataPlot,'groupY',allGroupY(krow));
    if ~isempty(datarow),
        minY = min(datarow.(nameY));
        maxY = max(datarow.(nameY));
    else
        minY = eps;
        maxY = 1;
    end
    if minY==maxY,
        % could happen if all data 0 for a row
        maxY = minY+1;
    end
    % Adjust maxY for titlebarX
    if logY(krow)==1,
        rangenew =  1/(1-heighttitlebarX-0.01)*(log10(maxY)-log10(minY));
        maxY     = 10.^(log10(minY) + rangenew);
    else
        rangenew =  1/(1-heighttitlebarX-0.01)*(maxY-minY);
        maxY     = minY + rangenew;
    end
    rowAxesLimits = [rowAxesLimits; sameAxesLimits(1:2) minY maxY];
end

%%%%%%%%%%%%%%%%%%%%%%%%%%%%%%%%%%%%%%%%%%%%%%%%%%%%%%%%%%%%%%%%%%%%%%%%%%%
% Open new figure
%%%%%%%%%%%%%%%%%%%%%%%%%%%%%%%%%%%%%%%%%%%%%%%%%%%%%%%%%%%%%%%%%%%%%%%%%%%
figure;
set(gcf,'Color',windowcolor);

%%%%%%%%%%%%%%%%%%%%%%%%%%%%%%%%%%%%%%%%%%%%%%%%%%%%%%%%%%%%%%%%%%%%%%%%%%%
% Plot the data in the order of subplots
%%%%%%%%%%%%%%%%%%%%%%%%%%%%%%%%%%%%%%%%%%%%%%%%%%%%%%%%%%%%%%%%%%%%%%%%%%%

for kSubplot=1:nSubplots,
    
    %%%%%%%%%%%%%%%%%%%%%%%%%
    % Choose subplot
    %%%%%%%%%%%%%%%%%%%%%%%%%
    
    subaxis(nrows,ncols,kSubplot,'SpacingVert',SpacingVert,'SpacingHoriz',SpacingHoriz,'Padding',Padding,'Margin',Margin);

    %%%%%%%%%%%%%%%%%%%%%%%%%
    % Get subplot data
    %%%%%%%%%%%%%%%%%%%%%%%%%
        
    dataSk = dataPlot(dataPlot.plotSubplot==kSubplot,:);
    
    %%%%%%%%%%%%%%%%%%%%%%%%%
    % Plot single plot using general auxiliary function
    %%%%%%%%%%%%%%%%%%%%%%%%%
            
    if ~isempty(dataSk),
        
        optionsplotgeneral = options;
        if ~isempty(nameGroupX),
            if isnumeric(dataSk.groupX(1)),
                optionsplotgeneral.titleText = sprintf('%s=%s',nameGroupX,num2str(dataSk.groupX(1)));
            else
                optionsplotgeneral.titleText = sprintf('%s=%s',nameGroupX,dataSk.groupX{1});
            end
        else
            optionsplotgeneral.titleText = 'All Data';
        end
        if sameaxes,
            optionsplotgeneral.axesLimits = sameAxesLimits;
        else
            subplotrow = ceil(kSubplot/ncols);
            optionsplotgeneral.axesLimits = rowAxesLimits(subplotrow,:);
        end
        optionsplotgeneral.showtitlebar = 0;
        optionsplotgeneral.sameaxes     = 1;
        optionsplotgeneral.logY = logY(ceil(kSubplot/ncols));
        
        plotgeneralIQM(dataSk,nameX,nameY,optionsplotgeneral);

    else
        if sameaxes, axis(sameAxesLimits);
        else subplotrow = ceil(kSubplot/ncols); axis(rowAxesLimits(subplotrow,:)); end
        hold on
        if logX, set(gca,'XScale','log'); end
        if logY(ceil(kSubplot/ncols)), set(gca,'YScale','log'); end
        if showgrid, grid on; end
        set(gca,'XMinorGrid','off')
        set(gca,'YMinorGrid','off')
        set(gca,'XColor',axescolor);
        set(gca,'YColor',axescolor);
        set(gca,'FontUnits','points','FontSize',ticklabeltextsize);    
    end
    
    %%%%%%%%%%%%%%%%%%%%%%%%%
    % Switch off xtick and ytick 
    % labels if sameaxes is chosen
    %%%%%%%%%%%%%%%%%%%%%%%%%   
    
    if mod(kSubplot-1,ncols)+1 > 1,
        set(gca,'YtickLabel','');
    end
    
    if kSubplot <= nSubplots-ncols,
        set(gca,'XtickLabel','');
    end
    
    %%%%%%%%%%%%%%%%%%%%%%%%%
    % Handle x labels
    %%%%%%%%%%%%%%%%%%%%%%%%%
    
    if kSubplot > nSubplots-ncols,
        xlabel(xlabelText,'FontUnits','points','FontSize',labeltextsize,'Interpreter','none');
    end
    
    %%%%%%%%%%%%%%%%%%%%%%%%%
    % Handle y labels
    %%%%%%%%%%%%%%%%%%%%%%%%%
    
    if mod(kSubplot-1,ncols)+1 == 1,
        nrow = ceil(kSubplot/ncols);
        ylabel(ylabelText{nrow},'FontUnits','points','FontSize',labeltextsize,'Interpreter','none');       
    end
    
    %%%%%%%%%%%%%%%%%%%%%%%%%
    % Handle titlebarX
    %%%%%%%%%%%%%%%%%%%%%%%%%
    
    if kSubplot<=ncols,
        % Get axes limits
        XLim = get(gca,'XLim');
        YLim = get(gca,'YLim');
        % Get X position for text
        if logX==1,
            X = 10^(log10(XLim(1)) + 0.5*(log10(XLim(2))-log10(XLim(1))));
        else
            X = XLim(1) + 0.5*(XLim(2)-XLim(1));
        end
        % Get Y position for text and title bar
        if logY(ceil(kSubplot/ncols))==1,
            Y = 10^(log10(YLim(1)) + (1-heighttitlebarX/2)*(log10(YLim(2))-log10(YLim(1))));
            YtitlebarDN = 10^(log10(YLim(1)) + (1-heighttitlebarX)*(log10(YLim(2))-log10(YLim(1))));
            YtitlebarUP = 10^(log10(YLim(1)) + 1*(log10(YLim(2))-log10(YLim(1))));
        else
            Y = YLim(1) + (1-heighttitlebarX/2)*(YLim(2)-YLim(1));
            YtitlebarDN = YLim(1) + (1-heighttitlebarX)*(YLim(2)-YLim(1));
            YtitlebarUP = YLim(1) + 1*(YLim(2)-YLim(1));
        end
        % Create a title bar background
        filled = [YtitlebarUP*[1 1],YtitlebarDN*[1 1]];
        xpoints=[XLim,fliplr(XLim)];
        h = fill(xpoints,filled,colortitlebar);
        set(h,'EdgeColor',axescolor);
        % Print the title text
        if length(allGroupX)==1,
            text(X,Y,'All Data','FontWeight','bold','FontSize',titlefontsize,'HorizontalAlignment','Center');
        else
            if isnumeric(allGroupX(1)),
                text(X,Y,sprintf('%s\n%g',nameGroupX,allGroupX(kSubplot)),'FontWeight','bold','FontSize',titlefontsize,'HorizontalAlignment','Center');
            else
                text(X,Y,sprintf('%s\n%s',nameGroupX,allGroupX{kSubplot}),'FontWeight','bold','FontSize',titlefontsize,'HorizontalAlignment','Center','Interpreter','none');
            end
        end
    end
    
    %%%%%%%%%%%%%%%%%%%%%%%%%
    % Handle titlebarY
    %%%%%%%%%%%%%%%%%%%%%%%%%
    
    if  mod(kSubplot-1,ncols)+1==ncols,
        % Get axes limits
        XLim = get(gca,'XLim');
        YLim = get(gca,'YLim');
        % Get Y position for text
        if logY(ceil(kSubplot/ncols)) == 1,
            Y = 10^(log10(YLim(1)) + 0.5*(log10(YLim(2))-log10(YLim(1))));
        else
            Y = YLim(1) + 0.5*(YLim(2)-YLim(1));
        end
        % Get X position for text and title bar
        if logX==1,
            X = 10^(log10(XLim(1)) + (1-widthtitlebarY/2)*(log10(XLim(2))-log10(XLim(1))));
            XtitlebarDN = 10^(log10(XLim(1)) + (1-widthtitlebarY)*(log10(XLim(2))-log10(XLim(1))));
            XtitlebarUP = 10^(log10(XLim(1)) + 1*(log10(XLim(2))-log10(XLim(1))));
        else
            X = XLim(1) + (1-widthtitlebarY/2)*(XLim(2)-XLim(1));
            XtitlebarDN = XLim(1) + (1-widthtitlebarY)*(XLim(2)-XLim(1));
            XtitlebarUP = XLim(1) + 1*(XLim(2)-XLim(1));
        end
        % Create a title bar background
        xpoints = [XtitlebarUP*[1 1],XtitlebarDN*[1 1]];
        filled=[YLim,fliplr(YLim)];
        h = fill(xpoints,filled,colortitlebar);
        set(h,'EdgeColor',axescolor);
        % Print the title text
        if length(allGroupY)==1,
            h = text(X,Y,'All Data','FontWeight','bold','FontSize',titlefontsize,'HorizontalAlignment','Center','Rotation',270);
        else
            if isnumeric(allGroupY(1)),
                h = text(X,Y,sprintf('%s\n%g',nameGroupY,allGroupY(ceil(kSubplot/ncols))),'FontWeight','bold','FontSize',titlefontsize,'HorizontalAlignment','Center','Rotation',270);
            else
                h = text(X,Y,sprintf('%s\n%s',nameGroupY,allGroupY{ceil(kSubplot/ncols)}),'FontWeight','bold','FontSize',titlefontsize,'HorizontalAlignment','Center','Rotation',270,'Interpreter','none');
            end
        end
    end
     
    %%%%%%%%%%%%%%%%%%%%%%%%%
    % Handle legend - simplistic one
    % Only do so for color group and median plots
    % Only show legend if less than maxlegendentries
    %%%%%%%%%%%%%%%%%%%%%%%%%
    
    if showlegend && kSubplot==1 && length(allGROUPcolor)<=maxlegendentries,
        legendText = {};
        legendColor = {};
        % Check if color group chosen
        if allGROUPcolor(1) ~= -1,
            for kleg=1:length(allGROUPcolor),
                % Get text part
                legendText{kleg} = sprintf('%s=%g',nameColorGroup,allGROUPcolor(kleg));
                % Get color part
                legendColor{kleg} = linecolorsCustom(mod(kleg-1,length(linecolorsCustom))+1,:);
                % Get marker part
                if showmarkers,
                    legendText{kleg} = sprintf('%s %s',linetypesCustom{   mod(kleg-1,length(linetypesCustom))+1},legendText{kleg});
                end
            end
        end
        % Check if moving median lines are shown
        if showmedian,
            legendText{end+1} = 'Binned median'; %#ok<*AGROW>
            legendColor{end+1} = [0 0 0];
        end

        if showmean,
            legendText{end+1} = 'Binned mean'; %#ok<*AGROW>
            legendColor{end+1} = [0 0 0];
        end
        
        % Determine y locations
        nlegend = length(legendText);
        YLim = get(gca,'YLim');
        XLim = get(gca,'XLim');
        
        if logY==1,
            textY = 10.^(log10(YLim(1))+((maxlegendentries:-1:1)/(maxlegendentries+1)-heighttitlebarX)*(log10(YLim(2))-log10(YLim(1))));
        else
            textY = YLim(1)+((maxlegendentries:-1:1)/(maxlegendentries+1)-heighttitlebarX)*(YLim(2)-YLim(1));
        end
        if logX==1,
            textX = 10.^(log10(XLim(1))+0.98*(log10(XLim(2))-log10(XLim(1))));
        else
            textX = XLim(1)+0.98*(XLim(2)-XLim(1));
        end
        for klegend=1:nlegend,
            text(textX,textY(klegend),legendText{klegend},'Color',legendColor{klegend},'FontWeight','bold','FontUnits','points','FontSize',legendtextsize,'HorizontalAlignment','Right','Interpreter','none');
        end
    end
    
end
    
  
```
