## Supplementary material for "A systems biology analysis of adrenergically stimulated adiponectin exocytosis in white adipocytes": All scripts used in this work: IQMplotfill.m

```
function [fillhandle] = IQMplotfill(x,lower,upper,color,transparency,logY,logX,edgeColor)
% This function will fill a region with a color between the two vectors 
% provided using the Matlab fill command. With the logY and logX flags
% correct handling of the data will be done, allowing to correctly plot the
% filled area. 
%
% [SYNTAX]
% [fillhandle] = IQMplotfill(x,lower,upper)
% [fillhandle] = IQMplotfill(x,lower,upper,color)
% [fillhandle] = IQMplotfill(x,lower,upper,color,transparency)
% [fillhandle] = IQMplotfill(x,lower,upper,color,transparency,logY)
% [fillhandle] = IQMplotfill(x,lower,upper,color,transparency,logY,logX)
% [fillhandle] = IQMplotfill(x,lower,upper,color,transparency,logY,logX,edgeColor)
%
% [INPUT]
% x             = The horizontal data points. Note: length(upper)
%                 must equal length(lower)and must equal length(x)!
% upper         = the upper curve values 
% lower         = the lower curve values 
% color         = the color of the filled area (default: [0.8 0.8 0.8])
% transparency  = is a value ranging from 1 for opaque to 0 for invisible for
%                 the filled color only (default: 1)
% logY:         = 1: set log y-axis, =0: set lin y-axis (Default: 0 if no figure, otherwise take settings from current axes)
% logX:         = 1: set log x-axis, =0: set lin x-axis (Default: 0 if no figure, otherwise take settings from current axes)
% edgeColor:    = the color around the edge of the filled area (default: none)
%
% [OUTPUT]
% fillhandle is the returned handle to the filled region in the plot.

% <<<COPYRIGHTSTATEMENT - IQM TOOLS LITE>>>

% Handle variable input arguments
if nargin<4,
    color = [0.8 0.8 0.8];
end
if nargin<5,
    transparency = 1;
end
if nargin<6,
    logY = 0;
    % Set YScale according to current plot
    if strcmp(get(gca,'YScale'),'log'),
        logY = 1;
    end
end
if nargin<7,
    logX = 0;
    % Set XScale according to current plot
    if strcmp(get(gca,'XScale'),'log'),
        logX = 1;
    end
end
if nargin<8,
    edgeColor = [];
end

% Check that inputs are vectors
if ~isvector(x),
    error('x needs t be a vector.');
end
if ~isvector(lower),
    error('lower needs t be a vector.');
end
if ~isvector(upper),
    error('upper needs t be a vector.');
end

% Force inputs to row vectors
x               = x(:)';
upper           = upper(:)';
lower           = lower(:)';

% Find NaN values and remove
ix              = find(isnan(upper)); x(ix) = []; upper(ix) = []; lower(ix) = [];
ix              = find(isnan(lower)); x(ix) = []; upper(ix) = []; lower(ix) = [];
ix              = find(isnan(x));     x(ix) = []; upper(ix) = []; lower(ix) = [];

% Find <=0 values in x and remove (only remove if logX==1)
if logX,
    ix              = find(x<=0); x(ix) = []; upper(ix) = []; lower(ix) = [];
end

% Find <=0 values in upper and lower remove (only remove if logY==1)
if logY,
    ix              = find(upper<=0); x(ix) = []; upper(ix) = []; lower(ix) = [];
    ix              = find(lower<=0); x(ix) = []; upper(ix) = []; lower(ix) = [];
end

% Plot the filled area
fillhandle = fill( [x fliplr(x)],  [upper fliplr(lower)], color);

% Set transparency
set(fillhandle,'FaceAlpha',transparency);

% Set edge color if defined
if ~isempty(edgeColor),
    set(fillhandle,'EdgeColor',edgeColor,'EdgeAlpha',transparency);
else
    set(fillhandle,'EdgeColor',color,'EdgeAlpha',transparency);
end    

% Set axes transformation
if logX,
    set(gca,'XScale','log');
else
    set(gca,'XScale','linear');
end
if logY,
    set(gca,'YScale','log');
else
    set(gca,'YScale','linear');
end
```
