## Supplementary material for "A systems biology analysis of adrenergically stimulated adiponectin exocytosis in white adipocytes": All scripts used in this work: IQMplotHistogram.m

```
function [] = IQMplotHistogram(Xdata,varargin)
% This function plots histograms of the data provided in X. X can be a vector
% or a matrix. If a matrix, then histograms are plotted for each column in 
% separate box-plots. Data can be grouped, resulting in several histograms in the 
% same plot. The grouping vector needs to have same length as X and consist 
% of some discrete values. Additionally, standard normal curves can be added,
% -2,2 range lines also, the number of samples in % outside the -2,2 interval can 
% be displayed. 
%
% [SYNTAX]
% [] = IQMplotHistogram(X)
% [] = IQMplotHistogram(X,options)
%
% [INPUT]
% X:            Vector or Matrix or MATLAB dataset containing the values to plot the 
%               histograms for (In columns).
% options:      MATLAB structure with optional arguments
%
%                   options.group:    Vector or MATLAB dataset containing discrete information.
%                                     For each value in it the analysis is grouped and the plots
%                                     are done in different colors.
%                   options.groupName: Name of the group variable
%                   options.Nbins:    Number of bins for the histogram (default: 20)
%                   options.show2lines: =1: do show the lines at -2 and 2, =0: dont (default: 0)
%                                     Disabled if groups selected
%                   options.stdNorm:  =1: show standard normal curve, =0: dont (default: 0)
%                   options.names:    Cell-array with names of the variables to be plotted.
%                   options.color:    =1: color, 0: black and white (default: 1)
%
% [OUTPUT]
% Histogram plots

% <<<COPYRIGHTSTATEMENT - IQM TOOLS LITE>>>

%%%%%%%%%%%%%%%%%%%%%%%%%%%%%%
% colors
%%%%%%%%%%%%%%%%%%%%%%%%%%%%%%
[colors,lines,dots,bwcolors] = IQMgetcolors();

%%%%%%%%%%%%%%%%%%%%%%%%%%%%%%
% Handle varargins
%%%%%%%%%%%%%%%%%%%%%%%%%%%%%%
options = [];
if nargin == 1,
elseif nargin == 2,
    options = varargin{1};
else
    error('Incorrect number of input arguments.');
end

%%%%%%%%%%%%%%%%%%%%%%%%%%%%%%
% Handle options
%%%%%%%%%%%%%%%%%%%%%%%%%%%%%%
group = [];
Nbins = 20;
show2lines = 0;
stdNorm = 0;
names = {};
color = 1;
groupName = 'GROUP';

try group = options.group; catch, end
try Nbins = options.Nbins; catch, end
try show2lines = options.show2lines; catch, end
try stdNorm = options.stdNorm; catch, end
try names = options.names; catch, end
try color = options.color; catch, end
try groupName = options.groupName; catch, end

%%%%%%%%%%%%%%%%%%%%%%%%%%%%%%
% Handle names
%%%%%%%%%%%%%%%%%%%%%%%%%%%%%%
if ischar(names),
    names = {names};
end

%%%%%%%%%%%%%%%%%%%%%%%%%%%%%%
% Convert possible datasets to double
%%%%%%%%%%%%%%%%%%%%%%%%%%%%%%
Xdata = double(Xdata); 
group = double(group); 

%%%%%%%%%%%%%%%%%%%%%%%%%%%%%%
% Find size of Xdata to determine need for number of subplots
%%%%%%%%%%%%%%%%%%%%%%%%%%%%
[nRows,nCols] = size(Xdata);
% Determine subplot structure
nTotal = nCols;
nsubplotCols = ceil(sqrt(nTotal));
nsubplotRows = ceil(nTotal/nsubplotCols);

%%%%%%%%%%%%%%%%%%%%%%%%%%%%%%
% Open new Figure
%%%%%%%%%%%%%%%%%%%%%%%%%%%%
figure; clf;

%%%%%%%%%%%%%%%%%%%%%%%%%%%%%%
% Cycle through all columns and do the plotting
%%%%%%%%%%%%%%%%%%%%%%%%%%%%%%
for kk = 1:nCols,
    subplot(nsubplotRows, nsubplotCols,kk);
    
    XdataColk = Xdata(:,kk);
    
    %%%%%%%%%%%%%%%%%%%%%%%%%%%%%%
    % Do the plot if group not defined
    %%%%%%%%%%%%%%%%%%%%%%%%%%%%%%
    if isempty(group),
%        NbinsUse = max(Nbins,ceil(Nbins/3*((max(XdataColk)-min(XdataColk)))));
        NbinsUse = Nbins;
        [h,x] = hist(XdataColk,NbinsUse);
        if color == 1,
            bar(x,h,1,'FaceColor',colors(1,:),'EdgeColor',colors(1,:)); hold on;
        else
            bar(x,h,1,'FaceColor',bwcolors(1,:),'EdgeColor',bwcolors(1,:)); hold on;
        end
        % Add standard normal curve
        if stdNorm,
            maxHIST = max(hist(XdataColk,NbinsUse));
            XLim = get(gca,'XLim');
            if XLim(1) > -5,
                XLim(1) = -5;
            end
            if XLim(2) < 5,
                XLim(2) = 5;
            end
            delta = (XLim(2)-XLim(1))/1000;
            x = [XLim(1):delta:XLim(2)];
            y = normpdfIQM(x,0,1)/0.4*maxHIST;
            plot(x,y,'--','Linewidth',2,'Color','black');
        end
    else
        %%%%%%%%%%%%%%%%%%%%%%%%%%%%%%
        % Do the plot if group is defined
        %%%%%%%%%%%%%%%%%%%%%%%%%%%%%%
        % Get unique group values
        groupall = unique(group);
        maxHIST = [];
        % Cycle through all group values
        for k=1:length(groupall),
            % Get group value to be handled
            groupk = groupall(k);
            % Get all Xdata in this group
            XdataColkk = XdataColk(group==groupk);
            % Calculate the histogram
%             NbinsUse = max(Nbins,ceil(Nbins/3*((max(XdataColkk)-min(XdataColkk)))));
            NbinsUse = Nbins;            [h,x] = hist(XdataColkk,NbinsUse);
            % Plot the histogram
            if color == 1,
                bar(x,h,1,'FaceColor',colors(k,:),'EdgeColor',colors(k,:)); hold on;
            else
                bar(x,h,1,'FaceColor',bwcolors(k,:),'EdgeColor',bwcolors(k,:)); hold on;
            end                
            h = findobj(gca,'Type','Patch');
            if k>1,
                set(h(1),'FaceAlpha',0.5)
            end
            % Determine for std norm curve:
            maxHIST(end+1) = max(hist(XdataColkk,NbinsUse));
        end
        % Add standard normal curve
        if stdNorm,
            for k=1:length(groupall),
                % Add standard normal curve
                XLim = get(gca,'XLim');
                if XLim(1) > -5,
                    XLim(1) = -5;
                end
                if XLim(2) < 5,
                    XLim(2) = 5;
                end
                delta = (XLim(2)-XLim(1))/1000;
                x = [XLim(1):delta:XLim(2)];
                y = normpdfIQM(x,0,1)/0.4*maxHIST(k);
                if color == 1,
                    plot(x,y,'--','Linewidth',2,'Color',colors(k,:));
                else
                    plot(x,y,'--','Linewidth',2,'Color',bwcolors(k,:));
                end
            end
        end
    end
    
    % Title etc.
    name = ['Xdata #' num2str(kk)];
    if ~isempty(names),
        try
            name = names{kk};
        catch
        end
    end        
    title(['Histogram of ' name],'FontSize',14,'FontWeight','bold','Interpreter','none');
    xlabel(name,'FontSize',14,'Interpreter','none');
    ylabel('Number per bin','FontSize',14,'Interpreter','none');
    set(gca,'FontSize',12) 
        
    % -2/2 lines - only if no groups defined
    if show2lines && isempty(group),
        YLim = get(gca,'YLim');
        plot([-2 -2],YLim,'k--');
        plot([2 2],YLim,'k--');
        % Get info about % samples outside -2,2 interval and show in plot
        XLim = get(gca,'XLim');
        percentXdatakOutside2 = round(length(find(abs(XdataColk) > 2))./length(find(abs(XdataColk) <= 2))*100*10)/10;
        text(XLim(1)+0.1*(XLim(2)-XLim(1)),0.9*YLim(2),sprintf('Samples outside\n[-2,2] interval:\n%g %%',percentXdatakOutside2),'FontSize',12);
    end

    % Legend if groups
    if ~isempty(group),
        legendtext = {};
        groupall = unique(group);
        for k=1:length(groupall),
            legendtext{end+1} = sprintf('%s: %g',groupName,groupall(k));
        end
        legend(legendtext{:},'Location','NorthWest');
    end
    
end
```
