## Supplementary material for "A systems biology analysis of adrenergically stimulated adiponectin exocytosis in white adipocytes": All scripts used in this work: IQMplotKM.m

```
function [ST50_output_cellTable,ST50_output] = IQMplotKM(TIME,CENS,color,line,type,marksFlag,CIflag)
% Plots a Kaplan Meier curve.
%
% [SYNTAX]
% [ST50_output] = IQMplotKM(TIME,CENS)
% [ST50_output] = IQMplotKM(TIME,CENS,color)
% [ST50_output] = IQMplotKM(TIME,CENS,color,line)
% [ST50_output] = IQMplotKM(TIME,CENS,color,line,type)
% [ST50_output] = IQMplotKM(TIME,CENS,color,line,type,marksFlag)
% [ST50_output] = IQMplotKM(TIME,CENS,color,line,type,marksFlag,CIflag)
%
% [INPUT]
% TIME:         vector with time information for time to event
% CENS:         vector with censoring information (0: uncensored, 1: right censored)
% color:        [r g b] color default: black
% line:         Line style (default: "-")
% type:         "Survivor" (default) or "cumulative hazard" or "cdf"
% marksFlag:    =1: censoring marks are plotted (default). =0 not plotted
% CIflag:       =1: plots 95% CI. =0 not plotted (default)
%
% [OUTPUT]
% ST50_output: [50% survival time, 95% CI - lower and upper bound]

% <<<COPYRIGHTSTATEMENT - IQM TOOLS LITE>>>

if nargin<3,
    color = [0 0 0];
end
if nargin<4,
    line = '-';
end
if nargin<5,
    type = 'survivor';
end
if nargin<6,
    marksFlag = 1;
end
if nargin<7,
    CIflag = 0;
end

% Remove NaN values from TIME and CENS (paired)
ixNAN           = find(isnan(TIME));
TIME(ixNAN)     = [];
CENS(ixNAN)     = [];

% Plot KM curve
[f,x,flo,fup]   = ecdf(TIME,'censoring',CENS,'function',type);
stairs(x,f,line,'LineWidth',1,'Color',color,'LineWidth',2); hold on;

% Plot marks for censored data
if marksFlag,
    xi = TIME(logical(CENS));
    [~,~,step] = histcounts(xi,x);
    step(xi>max(TIME)) = length(x);
    ixstep0 = find(step==0);
    xi(ixstep0) = [];
    step(ixstep0) = [];
    fi = f(step);
    plot(xi,fi, 's','MarkerEdgeColor','white','MarkerFaceColor','white','MarkerSize',8)
    plot(xi,fi, 'b+','color',color,'MarkerSize',8,'LineWidth',1)
end

% Plot 95% confidence interval
if CIflag,
    IQMplotfill(x,flo,fup,color,0.1); hold on
end

% Annotate
grid on
set(gca,'FontSize',12);
xlabel('Time','FontSize',14);
ylabel('Probability','FontSize',14);
set(gca,'YLim',[-0.05 1.05]);

% Determine 50% survival time and 95% CI
ix = find(f-0.5<0);
if isempty(ix),
    ST50 = NaN;
else
    ST50 = x(ix(1));
end

ix = find(flo-0.5<0);
if isempty(ix),
    ST50_CI_lo = NaN;
else
    ST50_CI_lo = x(ix(1));
end

ix = find(fup-0.5<0);
if isempty(ix),
    ST50_CI_up = NaN;
else
    ST50_CI_up = x(ix(1));
end

ST50_output = [ST50, ST50_CI_lo, ST50_CI_up];

ST50_output_cellTable(1,1:3) = {'<TH>' '50% Survival Time' '95% CI'};
ST50_output_cellTable(2,1:3) = {'<TR>' sprintf('%1.3g',ST50) sprintf('[%1.3g, %1.3g]',ST50_CI_lo,ST50_CI_up)};
```
