## Supplementary material for "A systems biology analysis of adrenergically stimulated adiponectin exocytosis in white adipocytes": All scripts used in this work: IQMplotpairwiseCorr.m

```
function [] = IQMplotpairwiseCorr(data,OPTIONS)
% This function plots the pairwise correlation between variables passed in
% columns of a matrix or passed as a dataset.
%
% [SYNTAX]
% [] = IQMplotpairwiseCorr(data)
% [] = IQMplotpairwiseCorr(data,OPTIONS)
%
% [INPUT]
% data:         Matrix or dataset. Each column corresponds to a variable
%               and each row to a sample
% OPTIONS:      MATLAB structure with optional arguments
%
%                   options.names:     cell-array with variable names. In
%                       case of data as dataset names will be taken from the
%                       header but can be overwritten with this option. If
%                       variable values are provided in a matrix, it is better
%                       to provide the names using this option
%                   options.LogFlag:   =1 do log transform the variables,
%                                      =0 do not transform (default: 0)
%                   options.CorrThres:   Value between 0 and 1 indicating the
%                       threshold for the Pearson correlation coefficient from which on a
%                       different color as background should be used (default:
%                       0.3)
%                   options.AxisColor:  [r g b] numeric values to use as color 
%                       for background if Corr>CorrThres =1 do log transform
%                       the variables (default: [1 0.2 0.2])
%
% [OUTPUT]
% Pairwise correlation plots.

% <<<COPYRIGHTSTATEMENT - IQM TOOLS LITE>>>

% Check if dataset or if matrix
% If dataset ten use columnnames as default values for names
if istable(data),
    names = data.Properties.VariableNames;
    % Make a matrix out of it
    data = table2array(data);
elseif strcmp(class(data),'double'),
    names = {};
    for k=1:size(data,2),
        names{k} = sprintf('Data#%d',k);
    end
end

% Default settings
% Flag to take log of values
LogFlag = 0;
% Coloring
CorrThres = 0.3; % Pearson correlation coefficient threshold for different background color
AxisColor = [1 .4 .4]; %color of axis when 

% Get optional values if defined
try names = OPTIONS.names; catch, end;
try LogFlag = OPTIONS.LogFlag; catch, end;
try CorrThres = OPTIONS.CorrThres; catch, end;
try AxisColor = OPTIONS.AxisColor; catch, end;

% Subindex properties
Spacing = 0;
Padding = 0;
Margin  = .1;
    
clf;
n = length(names);
for ir=1:n
    ystr = names{ir};
    y = data(:,ir);
    if LogFlag==1
        y = log(y);
        ystr = {'log',ystr}; %#ok<*AGROW>
    end
    for ic=1:ir
        xstr = names{ic};
        x = data(:,ic);
        if LogFlag==1
            x = log(x);
            xstr = {'log',xstr};
        end
        ip = (ir-1)*n+ic;
        subaxis(n,n,ip,'Spacing',Spacing,'Padding',Padding,'Margin',Margin);
        if ir==ic %plot histogram 
            % Only do this if not only NaN values are present
            if ~isempty(find(isnan(x)==0)),
                [b xbin] = hist(x,20);
                h = bar(xbin,b,1);
                if min(x)~=max(x),
                    set(gca,'XLim',[min(x) max(x)]);
                else
                    set(gca,'XLim',[min(x)-1 max(x)+1]);
                end                    
            end
            set(h,'FaceColor',0.4*[1 1 1],'LineStyle','none')
            title(xstr,'Interpreter','none')
        else %plot correlation
            % Remove pairs if at least one is NaN
            xuse = x;
            yuse = y;
            ixx = find(isnan(xuse));
            xuse(ixx) = []; yuse(ixx) = [];
            ixx = find(isnan(yuse));
            xuse(ixx) = []; yuse(ixx) = [];
            if ~isempty(xuse),
                % Determine pearsons coefficient of correlation
                [rho,pval]  = corrcoef(xuse,yuse);
                rho         = rho(1,2);
                pval        = pval(1,2);

                % Plot
                if abs(rho)>CorrThres
                    optcorr.Color     = AxisColor;
                else
                    optcorr.Color     = 0.6*[1 1 1];
                end
                optcorr.LineColor = [0 0 0];
                optcorr.TitleType = 'none';
                optcorr.LineStyle = '-';
                optcorr.LineWidth = 2;
                optcorr.MarkerSize = 10;
                plotcorrIQM(xuse,yuse,optcorr);
                
                xt = (min(xuse)+max(xuse))/2;
                yt = min(yuse)+.6*(max(yuse)-min(yuse));
                if abs(rho)>=0.01
                    if pval>=0.01,
                        str = sprintf('corr=%1.2f\np=%1.2f',rho,pval);
                    else
                        str = sprintf('corr=%1.2f\np<0.01',rho);
                    end
                else
                    str = '|corr|<0.01';
                end
                text(xt,yt,str,'Color',[0 0 0],'Hor','Right','Ver','Middle','FontWeight','Bold','Interpreter','none');
                
                if min(xuse)~=max(xuse),
                set(gca,'XLim',[min(xuse) max(xuse)]);
                else
                set(gca,'XLim',[min(xuse)-1 max(xuse)+1]);
                end
                if min(yuse)~=max(yuse),
                    set(gca,'YLim',[min(yuse) max(yuse)]);
                else
                    set(gca,'YLim',[min(yuse)-1 max(yuse)+1]);
                end
            end
        end
        
        % Define fontsize
        if length(names)<=6,
            FONTSIZE = 10;
        else
            FONTSIZE = 8;
        end
        
        if ic==1
            ylabel(ystr,'Interpreter','none','FontSize',FONTSIZE);
        end
        set(gca,'YTick',[]);
        
        if ir==n
            xlabel(xstr,'Interpreter','none','FontSize',FONTSIZE);
        end
        set(gca,'XTick',[]);
        
    end
end
```
