## Supplementary material for "A systems biology analysis of adrenergically stimulated adiponectin exocytosis in white adipocytes": All scripts used in this work: IQMplotPercentStratified.m

```
function [yn_all_percent] = IQMplotPercentStratified(data,catNames,compareNames,options)
% This function plots column plots / stacked bar plots for categorical outcomes (percent
% of occurrence) stratified by categorical compare variables in
% compareNames.
%
% Works for one catName and one compareName for now.
%
% [SYNTAX]
% [] = IQMplotPercentStratified(data,catNames,compareNames)
%
% [INPUT]
% data:         Matlab dataset. Each column corresponds to a variable
%               and each row to a sample. The columns with the names
%               defined in "catNames" need to be present in
%               the dataset and contain numerical categorical data.
% catNames:     Cell-array with names of categorical variables
% compareNames: Cell-array with names of categorical variables to compare
%               with the categorical covariates in catNames
% options:      Matlab structure with additional options
%       options.percent: Show percentages instead of numbers. Percentages
%                        are calculated relative to the number in the bins
%                        of the catNames and only displayed if compared to
%                        compareNames. =0 do show numbers and percent. =1 show
%                        percent (default) =2 show numbers and percent.
%       options.fontSizeText: Fontsize for the number and percent text
%                             (default: 10)
%       options.color:  Color of bubbles default: 0.4*[1 1 1]
%
% [OUTPUT]
% Plot

% <<<COPYRIGHTSTATEMENT - IQM TOOLS LITE>>>

% Check if data is table
if ~istable(data),
    error('Input data needs to be provided as MATLAB table.');
end

%%%%%%%%%%%%%%%%%%%%%%%%%%%%%%%%%%%%%%%%%%%%%%%%%%%%%%%%%%%%%%%%%%%%%%%%%%%
% Check which case it is and handle each case completely separately
%%%%%%%%%%%%%%%%%%%%%%%%%%%%%%%%%%%%%%%%%%%%%%%%%%%%%%%%%%%%%%%%%%%%%%%%%%%
if nargin==3,
    options = [];
end

try fontSizeText  = options.fontSizeText; catch, fontSizeText  = 10; end
try hideFirstRow  = options.hideFirstRow; catch, hideFirstRow  = 0; end
try color  = options.color; catch, color = 0.4*[1 1 1]; end

%%%%%%%%%%%%%%%%%%%%%%%%%%%%%%%%%%%%%%%%
% Check if dataset contains defined columns
%%%%%%%%%%%%%%%%%%%%%%%%%%%%%%%%%%%%%%%%
for k=1:length(catNames),
    try
        data.(catNames{k});
    catch
        error(sprintf('Please check if "%s" is a column in the dataset!',catNames{k}));
    end
end

for k=1:length(compareNames),
    try
        data.(compareNames{k});
    catch
        error(sprintf('Please check if "%s" is a column in the dataset!',compareNames{k}));
    end
end

if length(catNames)>1 || length(compareNames)>1,
    error('Only single entry in catNames and compareNames allowed.');
end

%%%%%%%%%%%%%%%%%%%%%%%%%%%%%%%%%%%%%%%%%%%%%%%%%%%%%%%%%%%%%%%%%%%%%%%%%%%
% Generate the plot
%%%%%%%%%%%%%%%%%%%%%%%%%%%%%%%%%%%%%%%%%%%%%%%%%%%%%%%%%%%%%%%%%%%%%%%%%%%

%%%%%%%%%%%%%%%%%%%%%%%%%%%%%%%%%%%%%%%%
% Keep only selected columns
%%%%%%%%%%%%%%%%%%%%%%%%%%%%%%%%%%%%%%%%
dataCompare = data(:,compareNames);
data        = data(:,catNames);

%%%%%%%%%%%%%%%%%%%%%%%%%%%%%%%%%%%%%%%%
% If is table then convert to double
%%%%%%%%%%%%%%%%%%%%%%%%%%%%%%%%%%%%%%%%
if istable(data),
    data = table2array(data);
end
if istable(dataCompare),
    dataCompare = table2array(dataCompare);
end

%%%%%%%%%%%%%%%%%%%%%%%%%%%%%%%%%%%%%%%%
% Options / fontsizes
%%%%%%%%%%%%%%%%%%%%%%%%%%%%%%%%%%%%%%%%

% Subindex properties
Spacing = 0.00;
Padding = 0.0005;
Margin  = .1;

% Define fontsize
if length(catNames)<=6,
    FONTSIZE = 10;
else
    FONTSIZE = 8;
end

% Define ncols and 
ncols = length(catNames);
nrows = length(compareNames);

% Cycle through compareNames
for kcompare=1:length(compareNames),
    ir = kcompare;
    ystr = compareNames{kcompare};
    y = dataCompare(:,kcompare);
    
    % Cycle through covNames
    for kcov=1:length(catNames),
        ic = kcov;
        xstr = catNames{ic};
        x = data(:,ic);
        ip = (ir-1)*ncols+ic;
%         subaxis(nrows,ncols,ip,'Spacing',Spacing,'Padding',Padding,'Margin',Margin);
        
        % Exchange x categories for numbers 1:N
        xu = unique(x);
        % Find positions where the unique items are listed (to obtain
        % number 1...N) ... stored in xn
        iu = {};
        for k=1:length(xu),
            iu{k} = find(x==xu(k));
        end
        xn = NaN(size(x));
        for k=1:length(iu),
            xn(iu{k}) = k;
        end
        
        % Exchange y categories for numbers 1:N
        yu = unique(y);
        iu = {};
        for k=1:length(yu),
            iu{k} = find(y==yu(k));
        end
        yn = NaN(size(y));
        for k=1:length(iu),
            yn(iu{k}) = k;
        end
        
        % Find numbers of unique yn in unique xn
        yn_all_percent = [];
        for k=1:max(xn),
            % Get number of xn values in bin k
            NR_xnk = length(xn(xn==k));
            % Get number of different yn values in bin k
            yn_all = [];
            for k2=1:max(yn),
                ynk = sum(xn==k & yn==k2);
                yn_all(k2) = ynk;
            end
            % Determine percent
            yn_all_percent = [yn_all_percent; 100*yn_all/NR_xnk];
        end
        
        % Plot
        hp = bar(1:max(xn),yn_all_percent,'stacked');
        
%         hatch_combinations = {
%         'fill'       0
%         'single'     0
%         'single'     45
%         'single'     90
%         'cross'      0
%         'cross'      45
%         };
%     
%         for k=1:min(length(hp),6),
%             hatchfill2(hp(k),hatch_combinations{k,1},'HatchAngle',hatch_combinations{k,2});
%         end
        
        % Annotate
        if ic==1
            ylabel('Percentage of x-bin','Interpreter','none','FontSize',12);
        else
            set(gca,'YTickLabel',[]);
        end
        
        if ir==nrows
            xlabel(xstr,'Interpreter','none','FontSize',FONTSIZE);
            set(gca,'XTickLabel',xu);
        else
            set(gca,'XTickLabel',[]);
        end
        
        set(gca,'YLim',[0 100]);
        grid on;
        
    end
end
```
