## Supplementary material for "A systems biology analysis of adrenergically stimulated adiponectin exocytosis in white adipocytes": All scripts used in this work: IQMplotQQ.m

```
function [] = IQMplotQQ(Xdata,varargin)
% QQ plot for provided input data.
%
% [SYNTAX]
% [] = IQMplotQQ(X)
% [] = IQMplotQQ(X,options)
%
% [INPUT]
% X:            Vector or Matrix or MATLAB dataset containing the values to plot the 
%               QQplot for (In columns).
% options:      MATLAB structure with optional arguments
%
%                   options.names:    Cell-array with names of the variables to be plotted.
%
% [OUTPUT]
% QQ plot
 
% <<<COPYRIGHTSTATEMENT - IQM TOOLS LITE>>>

%%%%%%%%%%%%%%%%%%%%%%%%%%%%%%
% Handle varargins
%%%%%%%%%%%%%%%%%%%%%%%%%%%%%%
options = [];
if nargin == 1,
elseif nargin == 2,
    options = varargin{1};
else
    error('Incorrect number of input arguments.');
end

%%%%%%%%%%%%%%%%%%%%%%%%%%%%%%
% Handle options
%%%%%%%%%%%%%%%%%%%%%%%%%%%%%%
names = {};
try names = options.names; catch, end

%%%%%%%%%%%%%%%%%%%%%%%%%%%%%%
% Handle names
%%%%%%%%%%%%%%%%%%%%%%%%%%%%%%
if ischar(names),
    names = {names};
end

%%%%%%%%%%%%%%%%%%%%%%%%%%%%%%
% Convert possible datasets to double
%%%%%%%%%%%%%%%%%%%%%%%%%%%%%%
Xdata = double(Xdata); 

%%%%%%%%%%%%%%%%%%%%%%%%%%%%%%
% Find size of Xdata to determine need for number of subplots
%%%%%%%%%%%%%%%%%%%%%%%%%%%%
[nRows,nCols] = size(Xdata);
% Determine subplot structure
nTotal = nCols;
nsubplotCols = ceil(sqrt(nTotal));
nsubplotRows = ceil(nTotal/nsubplotCols);

%%%%%%%%%%%%%%%%%%%%%%%%%%%%%%
% Open new Figure
%%%%%%%%%%%%%%%%%%%%%%%%%%%%
figure; clf;

%%%%%%%%%%%%%%%%%%%%%%%%%%%%%%
% Cycle through all columns and do the plotting
%%%%%%%%%%%%%%%%%%%%%%%%%%%%%%
for kk = 1:nCols,
    subplot(nsubplotRows, nsubplotCols,kk);
    
    XdataColk = Xdata(:,kk); 
    % Do the plot 
    qqplotIQM(XdataColk);

    % Title etc.
    name = ['Xdata #' num2str(kk)];
    if ~isempty(names),
        try
            name = names{kk};
        catch
        end
    end
    
    title(['QQplot of ' name],'FontSize',14,'FontWeight','bold','Interpreter','none');
    xlabel('Standard Normal Quantiles','FontSize',14,'Interpreter','none');
    ylabel(['Quantiles of ' name],'FontSize',14,'Interpreter','none');
    set(gca,'FontSize',12)
end
```
