## Supplementary material for "A systems biology analysis of adrenergically stimulated adiponectin exocytosis in white adipocytes": All scripts used in this work: IQMplotselected.m

```
function IQMplotselected(simdata,plotcomponents,varargin)
% IQMplotselected: Takes a simulation result as input. Additionally, the
% names of states, variables, and reactions can be defined for which the
% values should be plotted. 
%
% USAGE:
% ======
% IQMplotselected(simdata,plotcomponents)
% IQMplotselected(simdata,plotcomponents,headers)
%
% simdata: Simulation results returned, e.g. from IQMsimulate
% plotcomponents: cell-array with component names to plot (states,
%   variables and/or reactions). Alternatively, the cell-array can contain 
%   only cell-arrays of component names. This allows to define
%   plot-subgroups, which can be selected using the pulldown menu in the
%   upper left corner of the IQMplot window.
% headers: if plotcomponents contains only cell-arrays, then this input
%   argument contains the names to be displayed in the pull-down menu.
%
% DEFAULT VALUES:
% ===============
% headers: {'plot 1', 'plot 2', ...}

% <<<COPYRIGHTSTATEMENT - IQM TOOLS LITE>>>

%%%%%%%%%%%%%%%%%%%%%%%%%%%%%%%%%%%%%%%%%%%%%%%
% Check plotcomponents argument
%%%%%%%%%%%%%%%%%%%%%%%%%%%%%%%%%%%%%%%%%%%%%%%
if ~iscell(plotcomponents),
    error('IQMplotselected:componentsNotCellArray','Second input argument needs to be a cell-array.');
end
charfound = 0;
cellfound = 0;
for k=1:length(plotcomponents),
    if ischar(plotcomponents{k}),
        charfound = 1;
    elseif iscell(plotcomponents{k}),
        cellfound = 1;
    end
end
if charfound == 1 && cellfound == 1,
    error('IQMplotselected:charCellMix','Second input argument needs to contain either chars or cells. Not both!');
end
% convert char to cell model
if charfound == 1,
    plotcomponents = {plotcomponents};
end

%%%%%%%%%%%%%%%%%%%%%%%%%%%%%%%%%%%%%%%%%%%%%%%
% HANDLE VARIABLE INPUT ARGUMENTS
%%%%%%%%%%%%%%%%%%%%%%%%%%%%%%%%%%%%%%%%%%%%%%%
headers = {};
for k=1:length(plotcomponents),
    headers{k} = sprintf('plot %d',k);
end
if nargin == 3,
    headers = varargin{1};
end
if length(headers) ~= length(plotcomponents),
    error('IQMplotselected:wrongHeadersLength','Wrong number of elements in third input argument.');
end

%%%%%%%%%%%%%%%%%%%%%%%%%%%%%%%%%%%%%%%%%%%%%%%
% Prepare the different plot structures
%%%%%%%%%%%%%%%%%%%%%%%%%%%%%%%%%%%%%%%%%%%%%%%
plotstructures = {};
states = simdata.states; sind = 1:length(states);
variables = simdata.variables; vind = 1:length(variables);
reactions = simdata.reactions; rind = 1:length(reactions);
for k=1:length(plotcomponents),
    comp = plotcomponents{k};
    simdatak = simdata;
    sind_keep = [];
    vind_keep = [];
    rind_keep = [];
    for k2=1:length(comp),
        % reduce simulation results to only contain the components in comp
        sind_keep = [sind_keep strmatchIQM(comp{k2},states,'exact')];
        vind_keep = [vind_keep strmatchIQM(comp{k2},variables,'exact')];
        rind_keep = [rind_keep strmatchIQM(comp{k2},reactions,'exact')];
    end
    simdatak.states = simdatak.states(sind_keep);
    simdatak.statevalues = simdatak.statevalues(:,sind_keep);
    simdatak.variables = simdatak.variables(vind_keep);
    simdatak.variablevalues = simdatak.variablevalues(:,vind_keep);
    simdatak.reactions = simdatak.reactions(rind_keep);
    simdatak.reactionvalues = simdatak.reactionvalues(:,rind_keep);
    % convert to plot structure 
    plotstructures{k} = createdatastruct2IQMplotIQM(simdatak,headers{k}); 
end

%%%%%%%%%%%%%%%%%%%%%%%%%%%%%%%%%%%%%%%%%%%%%%%
% Construct IQMplot command and plot
%%%%%%%%%%%%%%%%%%%%%%%%%%%%%%%%%%%%%%%%%%%%%%%
plotcommand = 'IQMplot(';
for k=1:length(plotstructures),
    plotcommand = sprintf('%splotstructures{%d},',plotcommand,k);
end
plotcommand = [plotcommand(1:end-1) ')'];
eval(plotcommand)
```
