## Supplementary material for "A systems biology analysis of adrenergically stimulated adiponectin exocytosis in white adipocytes": All scripts used in this work: IQMplottrellis.m

```
function [] = IQMplottrellis(data,nameGroup,nameX,nameY,options)
% This function plots a Trellis plot. Plenty of options available.
%
% [SYNTAX]
% [] = IQMplottrellis(data,nameGroup,nameX,nameY)
% [] = IQMplottrellis(data,nameGroup,nameX,nameY,options)
%
% [INPUT]
% data:         Matlab table
% nameGroup:    Column name in dataset to use as group variable. If empty,
%               then no grouping is done.
% nameX:        Column name in dataset to plot on X axis
% nameY:        Column name in dataset to plot on Y axis
% options:      Structure with optional settings as follows:
%   options.xlabelText          = String with xlabel text. Default: nameX
%   options.ylabelText          = String with ylabel text. Default: nameY
%   options.nameSubGroup        = Column name in dataset for subgrouping within a group/subplot (nicer line drawing and used for moving median line);
%   options.nameColorGroup      = Column name in dataset for coloring a certain group differently
%   options.logX                = 0 if lin, 1 if log X axis (default: 0)
%   options.logY                = 0 if lin, 1 if log Y axis (default: 0)
%   options.linetype            = String with MATLAB linetype (default: '.-')
%   options.linecolor           = Color for lines (default: [0.2 0.2 0.2])
%   options.showmarkers         = 1 shows different linestyles with markers, =0 uses linstyle setting
%   options.markersize          = numeric value for markersizes (default: 10)
%   options.linecolorsCustom    = Matrix with 3 comlumns and arbitrary rows. Defines custom color settings to use for color grouping
%   options.linetypesCustom     = Cell-array with linestyle strings (e.g.: {'x','-.','--'}). If defined it overrides the standard lines from IQMgetcolors (only active if color group selected)
%                                 Only active if "options.showmarkers=1".
%   options.linewidth           = Numeric value 1-5 (default: 1)
%   options.sameaxes            = If 0 (default) subplot axes will be scaled to best fit the data, if 1 all axes will have same scaling
%   options.showgrid            = 0: no grid, 1: grid (no minor grid lines) (default: 1)
%   options.colortitlebar       = Vector with three values between 0 and 1 to set he titlebar color (default: [1 1 0.8])
%   options.heighttitlebar      = Height of the titlebar in fraction of subplot (default: 0.08)
%   options.showtitlebar        = 0: do not show titlebar, 1: show titlebar (default)
%   options.showmedian           = 0 (default): do not show a moving median line per group, 1: do show it
%   options.NbinsMedian          = value between 0 and 100 defining the range of data to take into account (default: 15)
%   options.showlegend          = 0: do not show a legend, 1 (default): show a legend for the color grouping
%   options.labeltextsize       = Fontsize for x and y labels (default: 10)
%   options.maxlegendentries    = If more elements in colorgroup then do not show legend (becomes messy). Default: 10
%   options.ticklabeltextsize   = Fontsize for axes number (default: 10)
%   options.legendtextsize      = Fontsize for legend (default: 10)
%   options.nrows               = Number of rows per subplot (default: all groups in one figure)
%   options.ncols               = Number of columns per subplot (default: all groups in one figure)
%   options.ylabelfirstonly     = 1: show the ylabel only for first subplot in figure, 0: show for all first in row subplots
%   options.nameText            = Column name in dataset (text in column) to display next to datapoints
%   options.nameTextLines       = 1: show lines additional to text (if nameText is defined). 0: do not show lines (if nameText is defined).
%   options.textFontsize        = Fontsize for additional text (default: 10)
%   options.filename            = Filename for PS (windows) or PDF (unix) output
%   options.axescolor           = Sets the color of axes and grid (default: [0.2 0.2 0.2])
%   options.windowcolor         = color definition for outside frame of window (default: [1 1 1])
%   options.titlefontsize       = Fontsize for title bars (default: 8)
% 
%   The "Vertical Line" option allows to draw vertical lines in subplots in
%   certain places. Practical, e.g., when plotting PK data and wanting to
%   plot the dosing times additionally. Several lines can be plotted in a
%   single subplot. A numeric value needs to be provided for each such
%   line. The maximum value will correspond to a line covering the whole
%   range. Smaller values will lead to a shorter line. It is possible to
%   plot the numeric value next to the top of each line.
%   options.verticallines.linestyle = linestyle string for the vertical line (default: '--')
%   options.verticallines.linecolor = color vector for the vertical line (default: [0 0 0])
%   options.verticallines.data      = dataset with info for vertical lines.
%       This dataset needs:
%           - same nameGroup column as main data
%           - same nameX column as main data
%           - additionally a nameDataVertical to be used to determine the height of the line
%   options.verticallines.nameDataVertical = string, specifying the column to use to determine the height of the line
%   options.verticallines.showtext  = 1: shows the value of "nameDataVertical" column next to the top of the line, =1: does not
%   options.verticallines.textsize  = text size for the printed value (default: 10)
%   options.verticallines.shownameDataVertical = 1: prints also the name of "nameDataVertical", =0 does not
%
% [OUTPUT]
% This function creates a new figure. If a filename is provided it also can
% export plots in a PS (windows) or PDF (unix) document.
%
% [ASSUMPTIONS]
% Data for X and Y axes and all groups needs to be numeric. 
 
% <<<COPYRIGHTSTATEMENT - IQM TOOLS LITE>>>

%%%%%%%%%%%%%%%%%%%%%%%%%%%%%%%%%%%%%%%%%%%%%%%%%%%%%%%%%%%%%%%%%%%%%%%%%%%
% Get default colors and linestyles
%%%%%%%%%%%%%%%%%%%%%%%%%%%%%%%%%%%%%%%%%%%%%%%%%%%%%%%%%%%%%%%%%%%%%%%%%%%
[colorsDefault,linesDefault] = IQMgetcolors();

%%%%%%%%%%%%%%%%%%%%%%%%%%%%%%%%%%%%%%%%%%%%%%%%%%%%%%%%%%%%%%%%%%%%%%%%%%%
% Get optional variables and set defaults if undefined
%%%%%%%%%%%%%%%%%%%%%%%%%%%%%%%%%%%%%%%%%%%%%%%%%%%%%%%%%%%%%%%%%%%%%%%%%%%
try xlabelText          = options.xlabelText;               catch, xlabelText           = nameX;            end; %#ok<*CTCH>
try ylabelText          = options.ylabelText;               catch, ylabelText           = nameY;            end;
try nameSubGroup        = options.nameSubGroup;             catch, nameSubGroup         = '';               end;
try nameColorGroup      = options.nameColorGroup;           catch, nameColorGroup       = '';               end;
try logX                = options.logX;                     catch, logX                 = 0;                end;
try logY                = options.logY;                     catch, logY                 = 0;                end;
try linetype            = options.linetype;                 catch, linetype             = '.-';             end; %#ok<*NASGU>
try linecolor           = options.linecolor;                catch, linecolor            = [0.2 0.2 0.2];    end;
try showmarkers         = options.showmarkers;              catch, showmarkers          = 0;                end;
try markersize          = options.markersize;               catch, markersize           = 10;                end;
try linecolorsCustom    = options.linecolorsCustom;         catch, linecolorsCustom     = colorsDefault;    end;
try linetypesCustom     = options.linetypesCustom;          catch, linetypesCustom      = linesDefault;     end;
try linewidth           = options.linewidth;                catch, linewidth            = 1;                end;
try sameaxes            = options.sameaxes;                 catch, sameaxes             = 1;                end;
try showgrid            = options.showgrid;                 catch, showgrid             = 1;                end;
try colortitlebar       = options.colortitlebar;            catch, colortitlebar        = [1 1 0.8];        end;
try heighttitlebar      = options.heighttitlebar;           catch, heighttitlebar       = 0.08;             end;
try showtitlebar        = options.showtitlebar;             catch, showtitlebar         = 1;                end;
try showmedian          = options.showmedian;               catch, showmedian           = 0;                end;
try NbinsMedian         = options.NbinsMedian;              catch, NbinsMedian          = 15;               end;
try showlegend          = options.showlegend;               catch, showlegend           = 1;                end;
try labeltextsize       = options.labeltextsize;            catch, labeltextsize        = 10;               end;
try ticklabeltextsize   = options.ticklabeltextsize;        catch, ticklabeltextsize    = 10;               end;
try legendtextsize      = options.legendtextsize;           catch, legendtextsize       = 10;               end;
try nrows               = options.nrows;                    catch, nrows                = NaN;              end;
try ncols               = options.ncols;                    catch, ncols                = NaN;              end;
try ylabelfirstonly     = options.ylabelfirstonly;          catch, ylabelfirstonly      = 0;                end;
try maxlegendentries    = options.maxlegendentries;         catch, maxlegendentries     = 10;               end;
try nameText            = options.nameText;                 catch, nameText             = '';               end;
try nameTextLines       = options.nameTextLines;            catch, nameTextLines        = 1;                end;
try textFontsize        = options.textFontsize;             catch, textFontsize         = 10;               end;
try filename            = options.filename;                 catch, filename             = '';               end;
try axescolor           = options.axescolor;                catch, axescolor            = 0.2*[1 1 1];      end;
try windowcolor         = options.windowcolor;              catch, windowcolor          = 1*[1 1 1];        end;
try verticallines_linestyle             = options.verticallines.linestyle;              catch, verticallines_linestyle              = '--';             end;
try verticallines_linecolor             = options.verticallines.linecolor;              catch, verticallines_linecolor              = [0 0 0];          end;
try verticallines_data                  = options.verticallines.data;                   catch, verticallines_data                   = [];               end;
try verticallines_nameDataVertical      = options.verticallines.nameDataVertical;       catch, verticallines_nameDataVertical       = [];               end;
try verticallines_showtext              = options.verticallines.showtext;               catch, verticallines_showtext               = 1;                end;
try verticallines_textsize              = options.verticallines.textsize;               catch, verticallines_textsize               = 10;               end;
try verticallines_shownameDataVertical  = options.verticallines.shownameDataVertical;   catch, verticallines_shownameDataVertical   = 0;                end;
try titlefontsize                       = options.titlefontsize;            catch, titlefontsize        = 8;               end

%%%%%%%%%%%%%%%%%%%%%%%%%%%%%%%%%%%%%%%%%%%%%%%%%%%%%%%%%%%%%%%%%%%%%%%%%%%
% If logY chosen then remove all <=0 data
% Same for logX
%%%%%%%%%%%%%%%%%%%%%%%%%%%%%%%%%%%%%%%%%%%%%%%%%%%%%%%%%%%%%%%%%%%%%%%%%%%
if logY==1,
    data(data.(nameY)<=0,:) = [];
end
if logX==1,
    data(data.(nameX)<=0,:) = [];
end
 
%%%%%%%%%%%%%%%%%%%%%%%%%%%%%%%%%%%%%%%%%%%%%%%%%%%%%%%%%%%%%%%%%%%%%%%%%%%
% If NaN in nameX then remove
%%%%%%%%%%%%%%%%%%%%%%%%%%%%%%%%%%%%%%%%%%%%%%%%%%%%%%%%%%%%%%%%%%%%%%%%%%%
data(isnan(data.(nameY)),:) = [];
data(isnan(data.(nameX)),:) = [];

%%%%%%%%%%%%%%%%%%%%%%%%%%%%%%%%%%%%%%%%%%%%%%%%%%%%%%%%%%%%%%%%%%%%%%%%%%%
% Settings for making plots nicer when same axes are chosen
%%%%%%%%%%%%%%%%%%%%%%%%%%%%%%%%%%%%%%%%%%%%%%%%%%%%%%%%%%%%%%%%%%%%%%%%%%%
if sameaxes,
    SpacingHoriz = 0.002;
    SpacingVert  = 0.003;
    Padding      = 0;
    Margin       = 0.1;
else
    SpacingHoriz = 0.05;
    SpacingVert  = 0.05;
    Padding      = 0.0;
    Margin       = 0.1;
end

%%%%%%%%%%%%%%%%%%%%%%%%%%%%%%%%%%%%%%%%%%%%%%%%%%%%%%%%%%%%%%%%%%%%%%%%%%%
% Format the data to match the grouping
%%%%%%%%%%%%%%%%%%%%%%%%%%%%%%%%%%%%%%%%%%%%%%%%%%%%%%%%%%%%%%%%%%%%%%%%%%%
% Get plot data alone
dataPlot            = table();
dataPlot.(nameX)    = data.(nameX);
dataPlot.(nameY)    = data.(nameY);
if ~isempty(nameText),
    dataPlot.(nameText) = data.(nameText);
end
if ~isempty(nameGroup)
    dataPlot.group  = data.(nameGroup);
else
    dataPlot.group  = ones(length(dataPlot),1);
end
if ~isempty(nameSubGroup),
    dataPlot.subgroup = data.(nameSubGroup);
else
    dataPlot.subgroup = ones(length(dataPlot),1);
end
if ~isempty(nameColorGroup),
    dataPlot.colorgroup = data.(nameColorGroup);
    dataPlot.color      = -1*ones(height(dataPlot),1);
    allGROUPcolor       = unique(dataPlot.colorgroup);    
    for k=1:length(allGROUPcolor),
        dataPlot.color(ixdataIQM(dataPlot,'colorgroup',allGROUPcolor(k))) = k;
    end
else
    dataPlot.colorgroup = -1*ones(height(dataPlot),1);
    dataPlot.color      = -1*ones(height(dataPlot),1);
end

%%%%%%%%%%%%%%%%%%%%%%%%%%%%%%%%%%%%%%%%%%%%%%%%%%%%%%%%%%%%%%%%%%%%%%%%%%%
% Determine the x and y axes ranges if sameaxes selected and
% adjust the maxY part of the axes settings to allow for title bar
% Goal is to increase the range of minY to maxY by a fraction of
% heighttitlebar
%%%%%%%%%%%%%%%%%%%%%%%%%%%%%%%%%%%%%%%%%%%%%%%%%%%%%%%%%%%%%%%%%%%%%%%%%%%
if sameaxes,
	minX = min(dataPlot.(nameX));
	maxX = max(dataPlot.(nameX));
	minY = min(dataPlot.(nameY));
	maxY = max(dataPlot.(nameY));
    % Adjust maxY for titlebar
    if logY==1,
        rangenew =  1/(1-heighttitlebar-0.01)*(log10(maxY)-log10(minY));
        maxY     = 10.^(log10(minY) + rangenew);
    else
        rangenew =  1/(1-heighttitlebar-0.01)*(maxY-minY);
        maxY     = minY + rangenew;
    end
    sameAxesLimits = [minX maxX minY maxY];
end

%%%%%%%%%%%%%%%%%%%%%%%%%%%%%%%%%%%%%%%%%%%%%%%%%%%%%%%%%%%%%%%%%%%%%%%%%%%
% Get different group elements - for each subplot
%%%%%%%%%%%%%%%%%%%%%%%%%%%%%%%%%%%%%%%%%%%%%%%%%%%%%%%%%%%%%%%%%%%%%%%%%%%
allGROUP = unique(dataPlot.group);

%%%%%%%%%%%%%%%%%%%%%%%%%%%%%%%%%%%%%%%%%%%%%%%%%%%%%%%%%%%%%%%%%%%%%%%%%%%
% Get number of possible elements in sub group
%%%%%%%%%%%%%%%%%%%%%%%%%%%%%%%%%%%%%%%%%%%%%%%%%%%%%%%%%%%%%%%%%%%%%%%%%%%
allGROUPsub = unique(dataPlot.subgroup);    

%%%%%%%%%%%%%%%%%%%%%%%%%%%%%%%%%%%%%%%%%%%%%%%%%%%%%%%%%%%%%%%%%%%%%%%%%%%
% Get number of possible elements in color group
%%%%%%%%%%%%%%%%%%%%%%%%%%%%%%%%%%%%%%%%%%%%%%%%%%%%%%%%%%%%%%%%%%%%%%%%%%%
allGROUPcolor = unique(dataPlot.colorgroup);    

%%%%%%%%%%%%%%%%%%%%%%%%%%%%%%%%%%%%%%%%%%%%%%%%%%%%%%%%%%%%%%%%%%%%%%%%%%%
% Determine number of subplot rows/cols per figure
%%%%%%%%%%%%%%%%%%%%%%%%%%%%%%%%%%%%%%%%%%%%%%%%%%%%%%%%%%%%%%%%%%%%%%%%%%%
nAllGroups = length(allGROUP);
if isnan(ncols) || isnan(nrows),
    ncols = ceil(sqrt(nAllGroups));
    nrows = ceil(nAllGroups/ncols);
end
nSubplots = ncols*nrows;

%%%%%%%%%%%%%%%%%%%%%%%%%%%%%%%%%%%%%%%%%%%%%%%%%%%%%%%%%%%%%%%%%%%%%%%%%%%
% Determine number of figures needed for the plotting
%%%%%%%%%%%%%%%%%%%%%%%%%%%%%%%%%%%%%%%%%%%%%%%%%%%%%%%%%%%%%%%%%%%%%%%%%%%
nFigures = ceil(nAllGroups/nSubplots);

%%%%%%%%%%%%%%%%%%%%%%%%%%%%%%%%%%%%%%%%%%%%%%%%%%%%%%%%%%%%%%%%%%%%%%%%%%%
% If output to file desired then start here the file
%%%%%%%%%%%%%%%%%%%%%%%%%%%%%%%%%%%%%%%%%%%%%%%%%%%%%%%%%%%%%%%%%%%%%%%%%%%
if ~isempty(filename),
    IQMstartNewPrintFigure(filename);
end

%%%%%%%%%%%%%%%%%%%%%%%%%%%%%%%%%%%%%%%%%%%%%%%%%%%%%%%%%%%%%%%%%%%%%%%%%%%
% Plot the data
% Plot moving median line if desired in each subplot for data within the same color group
% Subgroup is not considered
%%%%%%%%%%%%%%%%%%%%%%%%%%%%%%%%%%%%%%%%%%%%%%%%%%%%%%%%%%%%%%%%%%%%%%%%%%%
for k=1:length(allGROUP),

    %%%%%%%%%%%%%%%%%%%%%%%%%
    % Handle multiple figures and subplots
    %%%%%%%%%%%%%%%%%%%%%%%%%
    
    plotFigure  = ceil(k/nSubplots);
    plotSubplot = mod(k-1,nSubplots)+1;  

    % Clear figure if first subplot going to be plotted and set windowcolor
    if plotSubplot==1, 
        try close(plotFigure); catch, end
        figure(plotFigure); 
        set(gcf,'Color',windowcolor);
    else
        figure(plotFigure); 
    end
    % Choose subplot
    subaxis(nrows,ncols,plotSubplot,'SpacingVert',SpacingVert,'SpacingHoriz',SpacingHoriz,'Padding',Padding,'Margin',Margin);
    
    %%%%%%%%%%%%%%%%%%%%%%%%%
    % Get group data for each subplot
    %%%%%%%%%%%%%%%%%%%%%%%%%
    datak = subsetIQM(dataPlot,'group',allGROUP(k));
        
    %%%%%%%%%%%%%%%%%%%%%%%%%
    % Handle correct XLim in case of vertical lines
    %%%%%%%%%%%%%%%%%%%%%%%%%
    if ~isempty(verticallines_data),
        dataVLk = subsetIQM(verticallines_data,nameGroup,allGROUP(k));
        
        if ~isempty(dataVLk),
            % Get data - check if available in verticallines_data
            try
                XVLk = dataVLk.(nameX);
            catch
                error(sprintf('Please check settings for vertical lines - nameX and options.verticallines.nameDataVertical\nneed to be set correctly and be present in options.verticallines.data.')); %#ok<SPERR>
            end
            % Get X axis if needed because of vertical lines
            XLimMin = [min(XVLk) max(XVLk)];
        else
            XLimMin = [];
        end
    else
        XLimMin = [];
    end        
    
    %%%%%%%%%%%%%%%%%%%%%%%%%
    % Plot single plot using general auxiliary function
    %%%%%%%%%%%%%%%%%%%%%%%%%
    if ~isempty(nameGroup),
        if isnumeric(allGROUP),
            options.titleText = sprintf('%s=%s',nameGroup,num2str(allGROUP(k)));
        else
            options.titleText = sprintf('%s',allGROUP{k});
        end
    else
        options.titleText = 'All Data';
    end    
    if sameaxes,
        options.axesLimits = sameAxesLimits;
    else
        options.axesLimits = [];
    end
    % Set minimum X-axes in options
    options.XLimMin = XLimMin;
    
    % Set axes color
    options.axescolor = axescolor;
    
    options.titlefontsize = titlefontsize;
    
    plotgeneralIQM(datak,nameX,nameY,options);
 
    %%%%%%%%%%%%%%%%%%%%%%%%%
    % Switch off xtick and ytick 
    % labels if sameaxes is chosen
    %%%%%%%%%%%%%%%%%%%%%%%%%
    if sameaxes,
        % xticklabels
        if plotFigure < nFigures,
            % All subplots used => remove xticklabels in all but last row
            if plotSubplot <= nSubplots-ncols,
                set(gca,'XtickLabel','');
            end
        else
            % Figure only partially filled with subplots - only labels for
            % the last ncols subplots
            if k <= nAllGroups-ncols,
                set(gca,'XtickLabel','');
            end
        end
        % yticklabels
        if mod(plotSubplot-1,ncols)+1 > 1,
            set(gca,'YtickLabel','');
        end
    end
    
    %%%%%%%%%%%%%%%%%%%%%%%%%
    % Handle x labels
    %%%%%%%%%%%%%%%%%%%%%%%%%
    
    if plotFigure < nFigures,
        % All subplots used => xlabel for the last ncols subplots
        if plotSubplot > nSubplots-ncols,
            xlabel(xlabelText,'FontUnits','points','FontSize',labeltextsize,'Interpreter','none','Color',axescolor);
        end
    else
        % Figure only partially filled with subplots - only labels for
        % the last ncols subplots
        if k > nAllGroups-ncols,
            xlabel(xlabelText,'FontUnits','points','FontSize',labeltextsize,'Interpreter','none','Color',axescolor);
        end
    end
        
    %%%%%%%%%%%%%%%%%%%%%%%%%
    % Handle y labels
    %%%%%%%%%%%%%%%%%%%%%%%%%
    
    if ylabelfirstonly && plotSubplot==1 && nrows > 1,
        ylabel(ylabelText,'FontUnits','points','FontSize',labeltextsize,'HorizontalAlignment','Right','Interpreter','none','Color',axescolor);
    elseif nrows == 1,
        if plotSubplot==1,
            ylabel(ylabelText,'FontUnits','points','FontSize',labeltextsize,'Interpreter','none','Color',axescolor);     
        end
    elseif ~ylabelfirstonly && mod(plotSubplot-1,ncols)+1==1,
        ylabel(ylabelText,'FontUnits','points','FontSize',labeltextsize,'Interpreter','none','Color',axescolor);       
    end
    
    %%%%%%%%%%%%%%%%%%%%%%%%%
    % Handle legend - simplistic one
    % Only do so for color group and median plots
    % Only show legend if less than maxlegendentries
    %%%%%%%%%%%%%%%%%%%%%%%%%
    
    if showlegend && plotSubplot==1 && length(allGROUPcolor)<=maxlegendentries,
        legendText = {};
        legendColor = {};
        % Check if color group chosen
        if isnumeric(allGROUPcolor(1)),
            if allGROUPcolor(1)~= -1,
                for kleg=1:length(allGROUPcolor),
                    % Get text part
                    legendText{kleg} = sprintf('%s=%g',nameColorGroup,allGROUPcolor(kleg));
                    % Get color part
                    legendColor{kleg} = linecolorsCustom(mod(kleg-1,length(linecolorsCustom))+1,:);
                    % Get marker part
                    if showmarkers,
                        legendText{kleg} = sprintf('%s %s',linetypesCustom{   mod(kleg-1,length(linetypesCustom))+1},legendText{kleg});
                    end
                end
            end
        else
            for kleg=1:length(allGROUPcolor),
                % Get text part
                legendText{kleg} = sprintf('%s=%s',nameColorGroup,allGROUPcolor{kleg});
                % Get color part
                legendColor{kleg} = linecolorsCustom(mod(kleg-1,length(linecolorsCustom))+1,:);
                % Get marker part
                if showmarkers,
                    legendText{kleg} = sprintf('%s %s',linetypesCustom{   mod(kleg-1,length(linetypesCustom))+1},legendText{kleg});
                end
            end
        end
                
        % Check if moving median lines are shown
        if showmedian,
            legendText{end+1} = 'Binned median'; %#ok<*AGROW>
            legendColor{end+1} = [0 0 0];
        end
        
        % Determine y locations
        nlegend = length(legendText);
        YLim = get(gca,'YLim');
        XLim = get(gca,'XLim');
        
        if logY==1,
            textY = 10.^(log10(YLim(1))+((maxlegendentries:-1:1)/(maxlegendentries+1)-heighttitlebar)*(log10(YLim(2))-log10(YLim(1))));
        else
            textY = YLim(1)+((maxlegendentries:-1:1)/(maxlegendentries+1)-heighttitlebar)*(YLim(2)-YLim(1));
        end
        if logX==1,
            textX = 10.^(log10(XLim(1))+0.98*(log10(XLim(2))-log10(XLim(1))));
        else
            textX = XLim(1)+0.98*(XLim(2)-XLim(1));
        end
        for klegend=1:nlegend,
            text(textX,textY(klegend),legendText{klegend},'Color',legendColor{klegend},'FontWeight','bold','FontUnits','points','FontSize',legendtextsize,'HorizontalAlignment','Right','Interpreter','none');
        end
    end
    
    %%%%%%%%%%%%%%%%%%%%%%%%%
    % Handle and plot vertical lines if specified
    %%%%%%%%%%%%%%%%%%%%%%%%%
    
    if ~isempty(verticallines_data),
        dataVLk = subsetIQM(verticallines_data,nameGroup,allGROUP(k));
        if ~isempty(dataVLk),
            
            % Get data - check if available in verticallines_data
            try
                XVLk = dataVLk.(nameX);
                DVLk = dataVLk.(verticallines_nameDataVertical);
            catch
                error(sprintf('Please check settings for vertical lines - nameX and options.verticallines.nameDataVertical\nneed to be set correctly and be present in options.verticallines.data.')); %#ok<SPERR>
            end
            % Normalize DVLk - and handle also titlebar
            DVLknorm = (1-heighttitlebar-0.01)* DVLk/max(DVLk);
            % Determine Y range for plot
            YLim = get(gca,'YLim');
            Ydn = YLim(1);
            if logY==1,
                Yup = 10.^(log10(YLim(1))+DVLknorm*(log10(YLim(2))-log10(YLim(1))));
                Ytext = 10.^(log10(YLim(1))+(DVLknorm-0.05)*(log10(YLim(2))-log10(YLim(1))));
            else
                Yup = YLim(1)+DVLknorm*(YLim(2)-YLim(1));
                Ytext = YLim(1)+(DVLknorm-0.05)*(YLim(2)-YLim(1));
            end
            % Plot the lines
            for kVL=1:length(Yup),
                plot([XVLk(kVL),XVLk(kVL)],[Ydn,Yup(kVL)],verticallines_linestyle,'Color',verticallines_linecolor);
            end
            % Show text if desired - but only first and then when changes
            if verticallines_showtext,
                for kVL=1:length(Yup),
                    if kVL==1,
                        if verticallines_shownameDataVertical,
                            text(XVLk(kVL),Ytext(kVL),[' ' verticallines_nameDataVertical '=' num2str(DVLk(kVL))],'Color',verticallines_linecolor,'FontSize',verticallines_textsize,'BackgroundColor',windowcolor);
                        else
                            text(XVLk(kVL),Ytext(kVL),[' ' num2str(DVLk(kVL))],'Color',verticallines_linecolor,'FontSize',verticallines_textsize,'BackgroundColor',windowcolor);
                        end
                    elseif Ytext(kVL)~=Ytext(kVL-1),
                        if verticallines_shownameDataVertical,
                            text(XVLk(kVL),Ytext(kVL),[' ' verticallines_nameDataVertical '=' num2str(DVLk(kVL))],'Color',verticallines_linecolor,'FontSize',verticallines_textsize,'BackgroundColor',windowcolor);
                        else
                            text(XVLk(kVL),Ytext(kVL),[' ' num2str(DVLk(kVL))],'Color',verticallines_linecolor,'FontSize',verticallines_textsize,'BackgroundColor',windowcolor);
                        end
                    end
                end
            end
        end
    end
    
    %%%%%%%%%%%%%%%%%%%%%%%%%
    % If figure finished and output to file desired then do that
    %%%%%%%%%%%%%%%%%%%%%%%%%
    if ~isempty(filename),
        if plotFigure < nFigures,
            if plotSubplot == nSubplots,
                IQMprintFigure(gcf,filename);
                close(plotFigure);
            end
        else
            if k == nAllGroups,
                IQMprintFigure(gcf,filename);
                close(plotFigure);
            end
        end
    end
end

%%%%%%%%%%%%%%%%%%%%%%%%%
% If output to file desired then convert to pdf
%%%%%%%%%%%%%%%%%%%%%%%%%
IQMconvert2pdf(filename);
```
