## Supplementary material for "A systems biology analysis of adrenergically stimulated adiponectin exocytosis in white adipocytes": All scripts used in this work: IQMplotXY.m

```
function [] = IQMplotXY(data,nameX,nameY,options)
% This function plots pairwise Ydata vs. Xdata. Several optional settings
% are possible (see below)
%
% [SYNTAX]
% [] = IQMplotXY(data,nameX,nameY)
% [] = IQMplotXY(data,nameX,nameY,options)
%
% [INPUT]
% data:         Matlab dataset
% nameX:        Column name in dataset to plot on X axis
% nameY:        Cell-array with column names in dataset to plot on Y axis (if more than one name, then do subplots)
% options:      Structure with optional settings as follows:
%   options.xlabelText          = String with xlabel text. Default: nameX
%   options.ylabelText          = Cell-array with names of ylabels instead of nameY
%   options.nameColorGroup      = Column name in dataset for coloring a certain group differently
%   options.nameSubGroup        = Column name in dataset for subgrouping
%   options.logX                = 0 if lin, 1 if log X axis (default: 0)
%   options.logY                = 0 if lin, 1 if log Y axis (default: 0)
%                               Can be vector for different nameYs
%   options.linetype            = String with MATLAB linetype (default: '.')
%   options.linecolor           = Color for lines (default: [0.6 0.6 0.6])
%   options.showmarkers         = 1 shows different linestyles with markers, =0 uses linstyle setting
%   options.markersize          = numeric value for markersizes (default: 10)
%   options.linecolorsCustom    = Matrix with 3 comlumns and arbitrary rows. Defines custom color settings to use for color grouping
%   options.linetypesCustom     = Cell-array with linestyle strings (e.g.: {'x','-.','--'}). If defined it overrides the standard lines from IQMgetcolors (only active if color group selected)
%                                 Only active if "options.showmarkers=1".
%   options.linewidth           = Numeric value 1-5 (default: 1)
%   options.sameaxes            = If 0 (default) subplot axes will be scaled to best fit the data, if 1 all axes will have same scaling
%   options.showgrid            = 0: no grid, 1: grid (no minor grid lines) (default: 1)
%   options.colortitlebar       = Vector with three values between 0 and 1 to set he titlebar color (default: [1 1 0.8])
%   options.heighttitlebar      = Height of the titlebar in fraction of subplot (default: 0.04)
%   options.showtitlebar        = 0: do not show titlebar, 1: show titlebar (default)
%   options.showmedian           = 0 (default): do not show a median line per group, 1: do show it
%   options.NbinsMedian          = value between 0 and 100% defining the range of data to take into account (default: 15)
%   options.showregressionline  = 1: shwows a linear regression line, 0 does not (default)
%
%   options.showloessline       = 1: shwows a loess line (10% range), 0 does not (default)
%
%   options.showslope1line      = 1: shows a slope 1 line, 0 (default) does not
%   options.slope1linecolor     = color of slope1line (default: [0.4 0.4 0.4]) 
%   options.slope1linestyle     = style of slope1line (default: '--')
%   options.squareaxes          = sets X and Y limits to same values 
%   options.showzeroLines       = 1: shows lines at 0X and 0Y, 0 (default) does not   
%   options.zeroLinescolor      = color of zero lines (default: [0.4 0.4 0.4])  
%   options.zeroLinesstyle      = style of zero lines (default: '--'))  
%   options.showlegend          = 0: do not show a legend, 1 (default): show a legend for the color grouping
%   options.labeltextsize       = Fontsize for x and y labels (default: 10)
%   options.maxlegendentries    = If more elements in colorgroup then do not show legend (becomes messy). Default: 10
%   options.ticklabeltextsize   = Fontsize for axes number (default: 10)
%   options.legendtextsize      = Fontsize for legend (default: 10)
%   options.nrows               = Number of rows per subplot (default: all groups in one figure)
%   options.ncols               = Number of columns per subplot (default: all groups in one figure)
%   options.ylabelfirstonly     = 1: show the ylabel only for first subplot in figure, 0: show for all first in row subplots
%   options.nameText            = Column name in dataset (text in column) to display next to datapoints
%   options.nameTextLines       = 1: show lines additional to text (if nameText is defined). 0: do not show lines (if nameText is defined).
%   options.textFontsize        = Fontsize for additional text (default: 10)
%   options.filename            = Filename for PS (windows) or PDF (unix) output
%   options.axescolor           = Sets the color of axes and grid (default: [0.2 0.2 0.2])
%   options.windowcolor         = color definition for outside frame of window (default: [1 1 1])
%   options.removeNaNs          = 1: no breaks in lines due to NaN. =0: breaks in lines (default)
%   options.titleText           = String with title text (default: '')

% [OUTPUT]
% Figure with Y vs. X plots
 
% <<<COPYRIGHTSTATEMENT - IQM TOOLS LITE>>>

%% %%%%%%%%%%%%%%%%%%%%%%%%%%%%%%%%%%%%%%%%%%%%%%%%%%%%%%%%%%%%%%%%%%%%%%%%%
% Check inputs
%%%%%%%%%%%%%%%%%%%%%%%%%%%%%%%%%%%%%%%%%%%%%%%%%%%%%%%%%%%%%%%%%%%%%%%%%%%
if ischar(nameY),
    nameY = {nameY};
end

%%%%%%%%%%%%%%%%%%%%%%%%%%%%%%%%%%%%%%%%%%%%%%%%%%%%%%%%%%%%%%%%%%%%%%%%%%%
% Get default colors and linestyles
%%%%%%%%%%%%%%%%%%%%%%%%%%%%%%%%%%%%%%%%%%%%%%%%%%%%%%%%%%%%%%%%%%%%%%%%%%%
[colorsDefault,notused,linesDefault] = IQMgetcolors();

%%%%%%%%%%%%%%%%%%%%%%%%%%%%%%%%%%%%%%%%%%%%%%%%%%%%%%%%%%%%%%%%%%%%%%%%%%%
% Get optional variables and set defaults if undefined
%%%%%%%%%%%%%%%%%%%%%%%%%%%%%%%%%%%%%%%%%%%%%%%%%%%%%%%%%%%%%%%%%%%%%%%%%%%
try xlabelText          = options.xlabelText;               catch, xlabelText           = nameX;            end; %#ok<*CTCH>
try ylabelText          = options.ylabelText;               catch, ylabelText           = nameY;            end;
try nameSubGroup        = options.nameSubGroup;             catch, nameSubGroup        = '';                end; 
try nameColorGroup      = options.nameColorGroup;           catch, nameColorGroup       = '';               end; % also used for coloring
try logX                = options.logX;                     catch, logX                 = 0;                end;
try logY                = options.logY;                     catch, logY                 = [];               end;
try linetype            = options.linetype;                 catch, linetype             = '.';              end; 
try linecolor           = options.linecolor;                catch, linecolor            = [0.6 0.6 0.6];    end;
try showmarkers         = options.showmarkers;              catch, showmarkers          = 0;                end;
try markersize          = options.markersize;               catch, markersize          = 20;                end;
try linecolorsCustom    = options.linecolorsCustom;         catch, linecolorsCustom     = colorsDefault;    end;
try linetypesCustom     = options.linetypesCustom;          catch, linetypesCustom      = linesDefault;     end;
try linewidth           = options.linewidth;                catch, linewidth            = 1;                end;
try sameaxes            = options.sameaxes;                 catch, sameaxes             = 0;                end;
try showgrid            = options.showgrid;                 catch, showgrid             = 1;                end;
try colortitlebar       = options.colortitlebar;            catch, colortitlebar        = [1 1 0.8];        end;
try heighttitlebar      = options.heighttitlebar;           catch, heighttitlebar       = 0.04;             end;
try showtitlebar        = options.showtitlebar;             catch, showtitlebar         = 1;                end;
try showmedian          = options.showmedian;               catch, showmedian            = 0;               end;
try NbinsMedian         = options.NbinsMedian;              catch, NbinsMedian           = 15;              end;
try showregressionline  = options.showregressionline;       catch, showregressionline   = 0;                end;

try showloessline       = options.showloessline;            catch, showloessline        = 0;                end;

try showlegend          = options.showlegend;               catch, showlegend           = 1;                end;
try labeltextsize       = options.labeltextsize;            catch, labeltextsize        = 10;               end;
try ticklabeltextsize   = options.ticklabeltextsize;        catch, ticklabeltextsize    = 10;               end;
try legendtextsize      = options.legendtextsize;           catch, legendtextsize       = 10;               end;
try nrows               = options.nrows;                    catch, nrows                = NaN;              end;
try ncols               = options.ncols;                    catch, ncols                = NaN;              end;
try maxlegendentries    = options.maxlegendentries;         catch, maxlegendentries     = 15;               end;
try nameText            = options.nameText;                 catch, nameText             = '';               end;
try nameTextLines       = options.nameTextLines;            catch, nameTextLines        = 1;                end;
try textFontsize        = options.textFontsize;             catch, textFontsize         = 10;               end;
try axescolor           = options.axescolor;                catch, axescolor            = 0.2*[1 1 1];      end;
try windowcolor         = options.windowcolor;              catch, windowcolor          = 1*[1 1 1];        end;
try showslope1line      = options.showslope1line;           catch, showslope1line       = 0;                end;
try slope1linecolor     = options.slope1linecolor;          catch, slope1linecolor      = 0.4*[1 1 1];      end;
try slope1linestyle     = options.slope1linestyle;          catch, slope1linestyle      = '--';             end;
try squareaxes          = options.squareaxes;               catch, squareaxes           = 0;                end;
try showzeroLines       = options.showzeroLines;            catch, showzeroLines        = 0;                end;
try zeroLinescolor      = options.zeroLinescolor;           catch, zeroLinescolor       = 0.4*[1 1 1];      end;
try zeroLinesstyle      = options.zeroLinesstyle;           catch, zeroLinesstyle       = '--';             end;
try removeNaNs          = options.removeNaNs;               catch, removeNaNs           = 0;                end;
try titleText           = options.titleText;                catch, titleText            = '';               end;

%%%%%%%%%%%%%%%%%%%%%%%%%%%%%%%%%%%%%%%%%%%%%%%%%%%%%%%%%%%%%%%%%%%%%%%%%%%
% Handle nameY as char def
%%%%%%%%%%%%%%%%%%%%%%%%%%%%%%%%%%%%%%%%%%%%%%%%%%%%%%%%%%%%%%%%%%%%%%%%%%%
if ischar(nameY),
    nameY = {nameY};
end

% Handle logY
if isempty(logY),
    logY = zeros(1,length(nameY));
end
if length(logY) == 1,
    logY = logY*ones(1,length(nameY));
end
if length(logY)~=length(nameY),
    error('logY has wrong length');
end

%%%%%%%%%%%%%%%%%%%%%%%%%%%%%%%%%%%%%%%%%%%%%%%%%%%%%%%%%%%%%%%%%%%%%%%%%%%
% If logX remove negative X values
%%%%%%%%%%%%%%%%%%%%%%%%%%%%%%%%%%%%%%%%%%%%%%%%%%%%%%%%%%%%%%%%%%%%%%%%%%%
if logX,
    % Remove entries in data with negative X values
    data(data.(nameX)<=0,:) = [];
end

%%%%%%%%%%%%%%%%%%%%%%%%%%%%%%%%%%%%%%%%%%%%%%%%%%%%%%%%%%%%%%%%%%%%%%%%%%%
% Other options
%%%%%%%%%%%%%%%%%%%%%%%%%%%%%%%%%%%%%%%%%%%%%%%%%%%%%%%%%%%%%%%%%%%%%%%%%%%
ylabelfirstonly = 0;

%%%%%%%%%%%%%%%%%%%%%%%%%%%%%%%%%%%%%%%%%%%%%%%%%%%%%%%%%%%%%%%%%%%%%%%%%%%
% Settings for making plots nicer when same axes are chosen
%%%%%%%%%%%%%%%%%%%%%%%%%%%%%%%%%%%%%%%%%%%%%%%%%%%%%%%%%%%%%%%%%%%%%%%%%%%
if sameaxes,
    SpacingHoriz = 0.05;
    SpacingVert  = 0.05;
    Padding      = 0.0;
    Margin       = 0.1;
else
    SpacingHoriz = 0.05;
    SpacingVert  = 0.05;
    Padding      = 0.0;
    Margin       = 0.1;
end

%%%%%%%%%%%%%%%%%%%%%%%%%%%%%%%%%%%%%%%%%%%%%%%%%%%%%%%%%%%%%%%%%%%%%%%%%%%
% Format the data to match the grouping
% Each nameY is one element in the group
%%%%%%%%%%%%%%%%%%%%%%%%%%%%%%%%%%%%%%%%%%%%%%%%%%%%%%%%%%%%%%%%%%%%%%%%%%%
% Get plot data alone
dataPlot            = table();
for k=1:length(nameY),
    dataPlotk = table();
    dataPlotk.X          = data.(nameX);
    dataPlotk.Y          = data.(nameY{k});
   
    if ~isempty(nameText),
        dataPlotk.(nameText) = data.(nameText);
    end
    % No group name definition - it is via Yname definition
    dataPlotk.group  = k*ones(height(dataPlotk),1);
    % Handle subgrouping
    if isempty(nameSubGroup),
        dataPlotk.subgroup = ones(height(dataPlotk),1);
    else
        dataPlotk.subgroup = data.USUBJID;
    end
    % Handle color group
    if ~isempty(nameColorGroup),
        dataPlotk.colorgroup = data.(nameColorGroup);
        dataPlotk.color      = -1*ones(height(dataPlotk),1);
        allGROUPcolor       = unique(dataPlotk.colorgroup);
        for k=1:length(allGROUPcolor),
            dataPlotk.color(ixdataIQM(dataPlotk,'colorgroup',allGROUPcolor(k))) = k;
        end
    else
        % colorgroup same as subgroup
        dataPlotk.colorgroup = -1*ones(height(dataPlotk),1);
        dataPlotk.color      = -1*ones(height(dataPlotk),1);
    end
    
    % If logY remove negative Y values
    if logY,
        % Remove entries in data with negative X values
        dataPlotk(dataPlotk.Y<=0,:) = [];
    end
    
    dataPlot = [dataPlot; dataPlotk];
end

%%%%%%%%%%%%%%%%%%%%%%%%%%%%%%%%%%%%%%%%%%%%%%%%%%%%%%%%%%%%%%%%%%%%%%%%%%%
% Determine the x and y axes ranges if sameaxes selected and
% adjust the maxY part of the axes settings to allow for title bar
% Goal is to increase the range of minY to maxY by a fraction of
% heighttitlebar
%%%%%%%%%%%%%%%%%%%%%%%%%%%%%%%%%%%%%%%%%%%%%%%%%%%%%%%%%%%%%%%%%%%%%%%%%%%
if sameaxes,
    % Adjust for logY
    if logY,
        ix0 = find(dataPlot.X==0);
        dataPlot(ix0,:) = [];
        ix0 = find(dataPlot.Y==0);
        dataPlot(ix0,:) = [];
    end
    
	minX = min(dataPlot.X);
	maxX = max(dataPlot.X);
    minY = min(dataPlot.Y);
    maxY = max(dataPlot.Y);
        
    % Adjust maxY for titlebar
    if logY==1,
        rangenew =  1/(1-heighttitlebar-0.01)*(log10(maxY)-log10(minY));
        maxY     = 10.^(log10(minY) + rangenew);
    else
        rangenew =  1/(1-heighttitlebar-0.01)*(maxY-minY);
        maxY     = minY + rangenew;
    end
    % handle squareaxes
    if squareaxes,
        minX = min(minX,minY);
        minY = minX;
        maxX = max(maxX,maxY);
        maxY = maxX;        
    end
    % Define axes limits
    sameAxesLimits = [minX maxX minY maxY];
end

%%%%%%%%%%%%%%%%%%%%%%%%%%%%%%%%%%%%%%%%%%%%%%%%%%%%%%%%%%%%%%%%%%%%%%%%%%%
% Get different group elements - for each subplot
%%%%%%%%%%%%%%%%%%%%%%%%%%%%%%%%%%%%%%%%%%%%%%%%%%%%%%%%%%%%%%%%%%%%%%%%%%%
allGROUP = unique(dataPlot.group);

%%%%%%%%%%%%%%%%%%%%%%%%%%%%%%%%%%%%%%%%%%%%%%%%%%%%%%%%%%%%%%%%%%%%%%%%%%%
% Get number of possible elements in sub group
%%%%%%%%%%%%%%%%%%%%%%%%%%%%%%%%%%%%%%%%%%%%%%%%%%%%%%%%%%%%%%%%%%%%%%%%%%%
allGROUPsub = unique(dataPlot.subgroup);    

%%%%%%%%%%%%%%%%%%%%%%%%%%%%%%%%%%%%%%%%%%%%%%%%%%%%%%%%%%%%%%%%%%%%%%%%%%%
% Get number of possible elements in color group
%%%%%%%%%%%%%%%%%%%%%%%%%%%%%%%%%%%%%%%%%%%%%%%%%%%%%%%%%%%%%%%%%%%%%%%%%%%
allGROUPcolor = unique(dataPlot.colorgroup);    

%%%%%%%%%%%%%%%%%%%%%%%%%%%%%%%%%%%%%%%%%%%%%%%%%%%%%%%%%%%%%%%%%%%%%%%%%%%
% Determine number of subplot rows/cols per figure
%%%%%%%%%%%%%%%%%%%%%%%%%%%%%%%%%%%%%%%%%%%%%%%%%%%%%%%%%%%%%%%%%%%%%%%%%%%
nAllGroups = length(allGROUP);
if isnan(ncols) || isnan(nrows),
    ncols = ceil(sqrt(nAllGroups));
    nrows = ceil(nAllGroups/ncols);
end
nSubplots = ncols*nrows;

%%%%%%%%%%%%%%%%%%%%%%%%%%%%%%%%%%%%%%%%%%%%%%%%%%%%%%%%%%%%%%%%%%%%%%%%%%%
% Determine number of figures needed for the plotting
%%%%%%%%%%%%%%%%%%%%%%%%%%%%%%%%%%%%%%%%%%%%%%%%%%%%%%%%%%%%%%%%%%%%%%%%%%%
nFigures = ceil(nAllGroups/nSubplots);

%%%%%%%%%%%%%%%%%%%%%%%%%%%%%%%%%%%%%%%%%%%%%%%%%%%%%%%%%%%%%%%%%%%%%%%%%%%
% Plot the data
% Plot moving median line if desired
% moving median is done in each subplot for data within the same color group
% Subgroup is not considered
%%%%%%%%%%%%%%%%%%%%%%%%%%%%%%%%%%%%%%%%%%%%%%%%%%%%%%%%%%%%%%%%%%%%%%%%%%%
for k=1:length(allGROUP),

    %%%%%%%%%%%%%%%%%%%%%%%%%
    % Handle multiple figures and subplots
    %%%%%%%%%%%%%%%%%%%%%%%%%
    
    plotFigure  = ceil(k/nSubplots);
    plotSubplot = mod(k-1,nSubplots)+1;  

    % Clear figure if first subplot going to be plotted and set windowcolor
    if plotSubplot==1, 
        hfig = figure(); 
        set(gcf,'Color',windowcolor);
    else
        figure(hfig); 
    end
    % Choose subplot
    subaxis(nrows,ncols,plotSubplot,'SpacingVert',SpacingVert,'SpacingHoriz',SpacingHoriz,'Padding',Padding,'Margin',Margin);
    
    %%%%%%%%%%%%%%%%%%%%%%%%%
    % Get group data for each subplot
    %%%%%%%%%%%%%%%%%%%%%%%%%
    
    datak = dataPlot(dataPlot.group==allGROUP(k),:);
    
    %%%%%%%%%%%%%%%%%%%%%%%%%
    % Handle logY  
    %%%%%%%%%%%%%%%%%%%%%%%%%
    options.logY = logY(plotSubplot);
    
    % If logY than remove negative Y values
    if options.logY,
        datak(datak.Y<=0,:) = [];     
    end
    
    %%%%%%%%%%%%%%%%%%%%%%%%%
    % Plot single plot using general auxiliary function
    %%%%%%%%%%%%%%%%%%%%%%%%%
    if isempty(titleText),
        options.titleText = sprintf('%s vs %s',nameY{plotSubplot},nameX);
    else
        options.titleText = sprintf('%s (%s)',titleText,nameY{plotSubplot});
    end
    
    if sameaxes,
        options.axesLimits = sameAxesLimits;
    else
        minX = min(datak.X);
        maxX = max(datak.X);
        minY = min(datak.Y);
        maxY = max(datak.Y);
        
        % handle squareaxes
        if squareaxes,
            minX = min(minX,minY);
            minY = minX;
            maxX = max(maxX,maxY);
            maxY = maxX;     
        end
        % Adjust maxY for titlebar
        if options.logY==1,
            rangenew =  1/(1-heighttitlebar-0.01)*(log10(maxY)-log10(minY));
            maxY     = 10.^(log10(minY) + rangenew);
        else
            rangenew =  1/(1-heighttitlebar-0.01)*(maxY-minY);
            maxY     = minY + rangenew;
        end
        options.axesLimits = [minX maxX minY maxY];
    end
    options.linetype            = linetype;
    options.markersize          = markersize;
    options.linetypesCustom     = linetypesCustom;
    options.heighttitlebar      = heighttitlebar;
    options.NbinsMedian         = NbinsMedian;
    options.sameaxes            = sameaxes;
    options.axescolor           = axescolor;
    
    if removeNaNs,
        datak(isnan(datak.X),:) = [];
        datak(isnan(datak.Y),:) = [];        
    end
       
    plotgeneralIQM(datak,'X','Y',options);     
 
    %%%%%%%%%%%%%%%%%%%%%%%%%
    % Handle x labels
    %%%%%%%%%%%%%%%%%%%%%%%%%
    
    if plotFigure < nFigures,
        % All subplots used => xlabel for the last ncols subplots
        if plotSubplot > nSubplots-ncols,
            xlabel(xlabelText,'FontUnits','points','FontSize',labeltextsize,'Interpreter','none');
        end
    else
        % Figure only partially filled with subplots - only labels for
        % the last ncols subplots
        if k > nAllGroups-ncols,
            xlabel(xlabelText,'FontUnits','points','FontSize',labeltextsize,'Interpreter','none');
        end
    end
        
    %%%%%%%%%%%%%%%%%%%%%%%%%
    % Handle y labels
    %%%%%%%%%%%%%%%%%%%%%%%%%
    
    ylabel(ylabelText{k},'FontUnits','points','FontSize',labeltextsize,'Interpreter','none');
    
    %%%%%%%%%%%%%%%%%%%%%%%%%
    % Handle legend - simplistic one
    % Only do so for color group and median plots
    % Only show legend if less than maxlegendentries
    %%%%%%%%%%%%%%%%%%%%%%%%%
    
    if showlegend && plotSubplot==1 && length(allGROUPcolor)<=maxlegendentries,
        legendText = {};
        legendColor = {};
        % Check if color group chosen
        if allGROUPcolor(1) ~= -1,
            for kleg=1:length(allGROUPcolor),
                % Get text part
                legendText{kleg} = sprintf('%s=%g',nameColorGroup,allGROUPcolor(kleg));
                % Get color part
                legendColor{kleg} = linecolorsCustom(mod(kleg-1,length(linecolorsCustom))+1,:);
                % Get marker part
                if showmarkers,
                    legendText{kleg} = sprintf('%s %s',linetypesCustom{   mod(kleg-1,length(linetypesCustom))+1},legendText{kleg});
                end
            end
        end
        % Check if moving median lines are shown
        if showmedian,
            legendText{end+1} = 'Binned Median'; %#ok<*AGROW>
            legendColor{end+1} = [0 0 0];
        end
        % Check if regression lines are shown
        if showregressionline,
            legendText{end+1} = 'Regression line'; %#ok<*AGROW>
            legendColor{end+1} = [0 0 0];
        end
        % Check if loess lines are shown
        if showloessline,
            legendText{end+1} = 'Loess line'; %#ok<*AGROW>
            legendColor{end+1} = [0 0 0];
        end
        
        % Determine y locations
        nlegend = length(legendText);
        YLim = get(gca,'YLim');
        XLim = get(gca,'XLim');
        
        if options.logY==1,
            textY = 10.^(log10(YLim(1))+((maxlegendentries:-1:1)/(maxlegendentries+1)-heighttitlebar)*(log10(YLim(2))-log10(YLim(1))));
        else
            textY = YLim(1)+((maxlegendentries:-1:1)/(maxlegendentries+1)-heighttitlebar)*(YLim(2)-YLim(1));
        end
        if logX==1,
            textX = 10.^(log10(XLim(1))+0.02*(log10(XLim(2))-log10(XLim(1))));
        else
            textX = XLim(1)+0.02*(XLim(2)-XLim(1));
        end
        for klegend=1:nlegend,
            text(textX,textY(klegend),legendText{klegend},'Color',legendColor{klegend},'FontWeight','bold','FontUnits','points','FontSize',legendtextsize,'HorizontalAlignment','Left','Interpreter','none');
        end
    end
    
    %%%%%%%%%%%%%%%%%%%%%%%%%
    % Handle showslope1line
    %%%%%%%%%%%%%%%%%%%%%%%%%
    
    if showslope1line,
        XLim = get(gca,'XLim');
        YLim = get(gca,'YLim');
        % Adjust maxY for titlebar
        maxY = YLim(2);
        minY = YLim(1);
        if options.logY==1,
            rangenew =  (1-heighttitlebar)*(log10(maxY)-log10(minY));
            maxY     = 10.^(log10(minY) + rangenew);
        else
            rangenew =  (1-heighttitlebar)*(maxY-minY);
            maxY     = minY + rangenew;
        end         
        plot([min(XLim(1),YLim(1)) min(max(XLim(2),maxY),maxY)], [min(XLim(1),YLim(1)) min(max(XLim(2),maxY),maxY)],slope1linestyle,'Color',slope1linecolor,'LineWidth',2);
    end
    
    %%%%%%%%%%%%%%%%%%%%%%%%%
    % Handle showzeroLines
    %%%%%%%%%%%%%%%%%%%%%%%%%

    if showzeroLines,
        XLim = get(gca,'XLim');
        YLim = get(gca,'YLim');
        % Adjust maxY for titlebar
        maxY = YLim(2);
        minY = YLim(1);
        if options.logY==1,
            rangenew =  (1-heighttitlebar)*(log10(maxY)-log10(minY));
            maxY     = 10.^(log10(minY) + rangenew);
        else
            rangenew =  (1-heighttitlebar)*(maxY-minY);
            maxY     = minY + rangenew;
        end 
        plot([0 0],[minY maxY],zeroLinesstyle,'Color',zeroLinescolor,'LineWidth',1)
        plot(XLim,[0 0],zeroLinesstyle,'Color',zeroLinescolor,'LineWidth',1)
    end
    
end
```
