## Supplementary material for "A systems biology analysis of adrenergically stimulated adiponectin exocytosis in white adipocytes": All scripts used in this work: centeredfftIQM.m

```
function [X,freq]=centeredfftIQM(x,Fs)
% centeredfftIQM: Uses the fft function of MATLAB to determine a two sided
% spectrum of x, where te data in x have been sampled with the frequency 
% Fs. 
%
% [X,freq] = centeredfftIQM(x,Fs)

% <<<COPYRIGHTSTATEMENT - IQM TOOLS LITE>>>

% Get the length of the data vector x 
N=length(x);

% Determine the frequency axis
if mod(N,2)==0
    k = -N/2:N/2-1; % N even
else
    k = -(N-1)/2:(N-1)/2; % N odd
end
T = N/Fs;
freq = k/T;  % Frequency axis

% Determine the fft and normalize the result
X = fft(x)/N; 
% Shift the data in order to center it
X = fftshift(X); 
return
```
