## Supplementary material for "A systems biology analysis of adrenergically stimulated adiponectin exocytosis in white adipocytes": All scripts used in this work: positivefftIQM.m

```
function [X,freq] = positivefftIQM(x,Fs)
% positivefftIQM: Uses the fft function of MATLAB to determine a one sided
% spectrum of x, where te data in x have been sampled with the frequency 
% Fs. 
%
% [X,freq] = positivefftIQM(x,Fs)

% <<<COPYRIGHTSTATEMENT - IQM TOOLS LITE>>>

% Get the length of the data vector x 
N=length(x);

% Create the frequency axis
k=0:N-1;     
T=N/Fs;      
freq=k/T;    

% Determine the fft and normalize it
X=fft(x)/N; 
 
% Remove the negative frequency part and double the result (one sided)
cutOff = ceil(N/2); 
X = 2*X(1:cutOff);
freq = freq(1:cutOff);
return
```
