## Supplementary material for "A systems biology analysis of adrenergically stimulated adiponectin exocytosis in white adipocytes": All scripts used in this work: prepadIQM.m

```
function y = prepadIQM(x, l, c, dim)
% y = prepadIQM(x, l)
% y = prepadIQM(x, l, c)
% y = prepadIQM(x, l, c, dim)
% prepadIQM: prepends value c (default: 0) until to extend vector x to a
% length of l. Same with matrices x, where the dimension to extend is given
% by dim (default: 1). 

% <<<COPYRIGHTSTATEMENT - IQM TOOLS LITE>>>

if (nargin < 2 || nargin > 4)
    print_usage ();
end

if (nargin < 3 || isempty (c))
    c = 0;
else
    if (~isscalar (c))
        error ('prepadIQM: third argument must be empty or a scalar');
    end
end

nd = ndims (x);
sz = size (x);
if (nargin < 4),
    %% Find the first non-singleton dimension
    dim  = 1;
    while (dim < nd + 1 && sz (dim) == 1)
        dim = dim + 1;
    end
    if (dim > nd)
        dim = 1;
    end
else
    if (~(isscalar (dim) && dim == round (dim)) && dim > 0 && dim < (nd + 1))
        error ('prepadIQM: dim must be an integer and valid dimension');
    end
end

if (~ isscalar (l) || l < 0)
    error ('prepadIQM: second argument must be a positive scaler');
end

if (dim > nd)
    sz(nd+1:dim) = 1;
end

d = sz (dim);

if (d >= l)
    idx = cell ();
    for i = 1:nd
        idx{i} = 1:sz(i);
    end
    idx{dim} = d-l+1:d;
    y = x(idx{:});
else
    sz (dim) = l - d;
    y = cat (dim, c * ones (sz), x);
end
return
```
