## Supplementary material for "A systems biology analysis of adrenergically stimulated adiponectin exocytosis in white adipocytes": All scripts used in this work: resampleIQM.m

```
function [x2,t2] = resampleIQM(t1,x1,t2,method)
% resampleIQM: resamples time series x1, which is sampled at the time
% instances t1 to time series x2 using a sampling defined by t2.
%
% t2: scalar representing sampling interval or vector of sampling instances
% method: 'zoh', 'linear', 'cubic'. The use of 'method' is optional.
%         (default: 'linear')
%
% The output t2 is the vector that has been used for the resampling.
%
% [x2,t2] = resampleIQM(t1,x1,t2,method)

% <<<COPYRIGHTSTATEMENT - IQM TOOLS LITE>>>

if nargin < 3 || nargin > 4,
    error('Incorrect number of input arguments.');
end

if nargin == 3,
    method = 'linear';
end

if length(t2) == 1,
    t2 = [0:t2:t1(end)];
end

% Handle NaN data values
t1end = t1(end);
indnan = find(isnan(x1));
x1(indnan) = [];
t1(indnan) = [];
% Handle the case when the last value is NaN
if t1(end) ~= t1end,
    t1(end) = t1end;
end

% Do the resampling
x2 = zeros(1,length(t2));
if strcmp(method,'linear'),
    for k=1:length(t2),
        x2(k) = interp1IQM(t1,x1,t2(k));
    end
elseif strcmp(method,'zoh'),
    for k=1:length(t2),
        x2(k) = interp0IQM(t1,x1,t2(k));
    end    
elseif strcmp(method,'cubic'),
    for k=1:length(t2),
        x2(k) = interpcsIQM(t1,x1,t2(k));
    end    
else
    error('Wrong definition for ''method'' input argument.');
end

return
    
```
