## Supplementary material for "A systems biology analysis of adrenergically stimulated adiponectin exocytosis in white adipocytes": All scripts used in this work: xcorrIQM.m

```
function [R, lags] = xcorrIQM(X, varargin)
% Compute correlation R_xy of X and Y for various lags k:  
%
%    R_xy(k) = sum_{i=1}^{N-k}{x_i y_{i-k}}/(N-k),  for k >= 0
%    R_xy(k) = R_yx(-k),  for k <= 0
%
% usage: [R, lag] = xcorrIQM(X)
% usage: [R, lag] = xcorrIQM(X, Y)
% usage: [R, lag] = xcorrIQM(X, Y, maxlag)
% usage: [R, lag] = xcorrIQM(X, Y, maxlag, scale)
%
% Returns R(k+maxlag+1)=Rxy(k) for lag k=[-maxlag:maxlag].
% Scale is one of:
%    'biased'   for correlation=raw/N, 
%    'unbiased' for correlation=raw/(N-|lag|), 
%    'coeff'    for correlation=raw/(correlation at lag 0),
%    'none'     for correlation=raw
% If Y is omitted, compute autocorrelation.  
% If maxlag is omitted, use N-1 where N=max(length(X),length(Y)).
% If scale is omitted, use 'none'.
%
% If X is a matrix, computes the cross correlation of each column
% against every other column for every lag.  The resulting matrix has
% 2*maxlag+1 rows and P^2 columns where P is columns(X). That is,
%    R(k+maxlag+1,P*(i-1)+j) == Rij(k) for lag k=[-maxlag:maxlag],
% so
%    R(:,P*(i-1)+j) == xcorr(X(:,i),X(:,j))
% and
%    reshape(R(k,:),P,P) is the cross-correlation matrix for X(k,:).
%
% xcorr computes the cross correlation using an FFT, so the cost is
% dependent on the length N of the vectors and independent of the
% number of lags k that you need.  If you only need lags 0:k-1 for 
% vectors x and y, then the direct sum may be faster:
%
% unbiased:
%  ( hankel(x(1:k),[x(k:N); zeros(k-1,1)]) * y ) ./ [N:-1:N-k+1](:)
% biased:
%  ( hankel(x(1:k),[x(k:N); zeros(k-1,1)]) * y ) ./ N
%
% If length(x) == length(y) + k, then you can use the simpler
%    ( hankel(x(1:k),x(k:N-k)) * y ) ./ N
%
% Ref: Stearns, SD and David, RA (1988). Signal Processing Algorithms.
%      New Jersey: Prentice-Hall.

% <<<COPYRIGHTSTATEMENT - IQM TOOLS LITE>>>

if (nargin < 1 || nargin > 4)
    error('Incorrect number of input arguments.');
end

% assign arguments from list
Y = [];
maxlag = [];
scale = 'none';
N = length(X);
if nargin==2
    Y = varargin{1};
    N = max(length(X),length(Y));
elseif nargin==3
    Y = varargin{1};
    maxlag = varargin{2};
elseif nargin == 4,
    Y = varargin{1};
    maxlag = varargin{2};
    scale = varargin{3};
end
if isempty(maxlag),
    maxlag = N-1;
end

% check argument values
if isscalar(X) || ischar(X) || isempty(X),
    error('xcorrIQM: X must be a vector or matrix');
end
if isscalar(Y) || ischar(Y) || (~isempty(Y) && ~isvector(Y)),
    error('xcorrIQM: Y must be a vector');
end
if ~isvector(X) && ~isempty(Y),
    error('xcorrIQM: X must be a vector if Y is specified');
end
if ~isscalar(maxlag) && ~isempty(maxlag)
    error('xcorrIQM: maxlag must be a scalar');
end
if maxlag>N-1,
    error('xcorrIQM: maxlag must be less than length(X)');
end
if isvector(X) && isvector(Y) && length(X) ~= length(Y) && ~strcmp(scale,'none')
    error('xcorrIQM: scale must be ''none'' if length(X) ~= length(Y)')
end

P = size(X,2);
M = 2^nextpow2(N + maxlag);
if ~isvector(X)
    % For matrix X, compute cross-correlation of all columns
    R = zeros(2*maxlag+1,P^2);

    % Precompute the padded and transformed `X' vectors
    pre = fft (postpadIQM (prepadIQM (X, N+maxlag), M) );
    post = conj (fft (postpadIQM (X, M)));

    % For diagonal (i==j)
    cor = ifft (post .* pre);
    R(:, 1:P+1:P^2) = conj (cor (1:2*maxlag+1,:));

    % For remaining i,j generate xcorr(i,j) and by symmetry xcorr(j,i).
    for i=1:P-1
        j = i+1:P;
        cor = ifft (pre(:,i*ones(length(j),1)) .* post(:,j));
        R(:,(i-1)*P+j) = conj (cor (1:2*maxlag+1, :));
        R(:,(j-1)*P+i) = flipud (cor (1:2*maxlag+1, :));
    end
elseif isempty(Y)
    % compute autocorrelation of a single vector
    post = fft (postpadIQM(X,M));
    cor = ifft (conj(post(:)) .* post(:));
    R = [ conj(cor(maxlag+1:-1:2)) ; cor(1:maxlag+1) ];
else
    % compute cross-correlation of X and Y
    post = fft (postpadIQM(Y,M));
    pre = fft (postpadIQM(prepadIQM(X,N+maxlag),M));
    cor = conj (ifft (conj(post(:)) .* pre(:)));
    R = cor(1:2*maxlag+1);
end

% if inputs are real, outputs should be real, so ignore the
% insignificant complex portion left over from the FFT
if isreal(X) && (isempty(Y) || isreal(Y))
    R=real(R);
end

% correct for bias
if strcmp(scale, 'biased')
    R = R ./ N;
elseif strcmp(scale, 'unbiased')
    R = R ./ ( [ N-maxlag:N-1, N, N-1:-1:N-maxlag ]' * ones(1,columns(R)) );
elseif strcmp(scale, 'coeff')
    R = R ./ ( ones(size(R,1),1) * R(maxlag+1, :) );
elseif ~strcmp(scale, 'none')
    error('xcorr: scale must be ''biased'', ''unbiased'', ''coeff'' or ''none''');
end

% correct the shape so that it is the same as the input vector
if isvector(X) && P > 1
    R = R';
end

% return the lag indices if desired
if nargout == 2
    lags = [-maxlag:maxlag];
end

return
```
