## Supplementary material for "A systems biology analysis of adrenergically stimulated adiponectin exocytosis in white adipocytes": All scripts used in this work: binnedmedianIQM.m

```
function [xbin,ybinmedian] = binnedmedianIQM(x,y,numbins,FLAGlogX)
% Simple function to calcuate binned medians. Number of bins can be provided and 
% it is possible to tell the function to do the binning on a log x axis.
% The binning results might not be very nice. 
%
% [SYNTAX]
% [xbin,ybinmedian] = binnedmedianIQM(x,y)
% [xbin,ybinmedian] = binnedmedianIQM(x,y,numbins)
% [xbin,ybinmedian] = binnedmedianIQM(x,y,numbins,FLAGlogX)
%
% [INPUT]
% x:           x - values
% y:           y - values
% numbins:     number of bins (default: 15)
% FLAGlogX:    0: bin on linear axis (default), 1: bin on log axis
%
% [OUTPUT]
% Bins in xbin and binned medians in ybinmedian

% <<<COPYRIGHTSTATEMENT - IQM TOOLS LITE>>>

if nargin <= 2,
    numbins = 15;
    FLAGlogX = 0;
elseif nargin<=3,
    FLAGlogX = 0;
end    

if length(numbins)==1,
    if FLAGlogX,
        bins = logspace(log10(min(x)), log10(max(x)), numbins);
    else
        bins = linspace(min(x), max(x), numbins);
    end
else
    bins = numbins;
    numbins = length(bins);
end

[n,bin] = histc(x, bins); %#ok<ASGLU>
mu = NaN*zeros(size(bins));
for k = [1:numbins], %#ok<NBRAK>
  ind = find(bin==k);
  if (~isempty(ind))
    mu(k) = median(y(ind));
  end
end

% Remove NaNs
Z = [bins(:) mu(:)];
Z(isnan(Z(:,2)),:) = [];

% Assign outputs
xbin = Z(:,1);
ybinmedian = Z(:,2);
```
