## Supplementary material for "A systems biology analysis of adrenergically stimulated adiponectin exocytosis in white adipocytes": All scripts used in this work: binnedquantilesIQM.m

```
function [xbin,ybinquantile] = binnedquantilesIQM(x,y,quantile_value,binningInfo,FLAGlogX)
% Simple function to calcuate binned quantiles. Number of bins can be provided and 
% it is possible to tell the function to do the binning on a log x axis. Instead
% of providing the number of bins it is also possible to provide the bins driectly along with 
% some range around these values to use for binning.
% 
% [SYNTAX]
% [xbin,ybinquantile] = binnedquantilesIQM(x,y,quantile_value)
% [xbin,ybinquantile] = binnedquantilesIQM(x,y,quantile_value,numbins)
% [xbin,ybinquantile] = binnedquantilesIQM(x,y,quantile_value,numbins,FLAGlogX)
% [xbin,ybinquantile] = binnedquantilesIQM(x,y,quantile_value,binningInfo)
% [xbin,ybinquantile] = binnedquantilesIQM(x,y,quantile_value,binningInfo,FLAGlogX)
%
% [INPUT]
% x:           x - values
% y:           y - values
% quantileIQM:    quantileIQM to calculate
% numbins:     number of bins (default: 15)
% FLAGlogX:    0: bin on linear axis (default), 1: bin on log axis
% binningInfo: Cell-array. The first element is a vector with centers of chosen bins. 
%              The second element is a vector with ranges to look around left and right 
%              from the chosen centers.
%
% [OUTPUT]
% Bins in xbin and binned quantiles in ybinquantile

% <<<COPYRIGHTSTATEMENT - IQM TOOLS LITE>>>

if nargin <= 3,
    binningInfo = 15;  % number bins
    FLAGlogX = 0;
elseif nargin<=4,
    FLAGlogX = 0;
end    

if length(binningInfo) == 1,
    % Bin equidistantly
    numbins = binningInfo;
    if FLAGlogX,
        bins = logspace(log10(min(x)), log10(max(x)), numbins);
    else
        bins = linspace(min(x), max(x), numbins);
    end
    
    try
        [n,bin] = histc(x, bins); %#ok<ASGLU>
    catch
        'error'
    end
    
    mu = NaN*zeros(size(bins));
    for k = [1:numbins], %#ok<NBRAK>
        ind = find(bin==k);
        if (~isempty(ind))
            mu(k) = quantileIQM(y(ind),quantile_value);
        end
    end
    
    % Remove NaNs
    Z = [bins(:) mu(:)];
    Z(isnan(Z(:,2)),:) = [];
    
    % Assign outputs
    xbin = Z(:,1);
    ybinquantile = Z(:,2);
else
    % bin my mean binning value and look around range
    bins_mean = binningInfo{1};
    bins_lookaround = binningInfo{2};
    
    ybinquantile = [];
    for k=1:length(bins_mean),
        ix = find(x>=bins_mean(k)-bins_lookaround(k) & x<bins_mean(k)+bins_lookaround(k));
        ybinquantile(k) = quantileIQM(y(ix),quantile_value);
    end
    xbin = bins_mean;
    ix = find(isnan(ybinquantile));
    xbin(ix) = [];
    ybinquantile(ix) = [];
end
```
