## Supplementary material for "A systems biology analysis of adrenergically stimulated adiponectin exocytosis in white adipocytes": All scripts used in this work: movingAverageIQM.m

```
function [y]=movingAverageIQM(x,range,fun)
% movingAverageIQM will compute moving averages over a range, defined by the user.
%
% Usage: y = movingAverageIQM(x,range[,fun])
% where 
%      x:       is the input vector (or matrix) to be smoothed. 
%      range: 	percentage of number of points in x to be used to average over
%      y:       is output vector of same length as x
%      fun:     (optional) is a custom function rather than moving averages
%
% Note:if x is a matrix then the smoothing will be done 'vertically'.
% 
%
% Example:
%
% x=randn(300,1);
% plot(x,'g.'); 
% hold on;
% plot(movingAverageIQM(x,10),'k'); 
% plot(movingAverageIQM(x,30,'median'),'c');
% plot(movingAverageIQM(x,7,@(x)max(x)),'b'); 
% legend('x','10% range moving mean','30% range moving median','7% range moving max','location','best')

% <<<COPYRIGHTSTATEMENT - IQM TOOLS LITE>>>

% Determine m based on range (in percent)
m = length(x)*range/100;
% make m odd, since it seems to be better
m = 2*round(m/2)+1;

if m==1
    y=x;
    return
end
if size(x,1)==1
    x=x';
end

if nargin<3
    fun=[];
elseif ischar(fun)
    fun=eval(['@(x)' fun '(x)']);
end

if isempty(fun)

    f=zeros(m,1)+1/m;
    n=size(x,1);
    isodd=bitand(m,1);
    m2=floor(m/2);


    if (size(x,2)==1)
        y=filter(f,1,x);
        y=y([zeros(1,m2-1+isodd)+m,m:n,zeros(1,m2)+n]);
    else
        y=filter2(f,x);
        y(1:(m2-~isodd),:)=y(m2+isodd+zeros(m2-~isodd,1),:);
        y((n-m2+1):end,:)=y(n-m2+zeros(m2,1),:);
    end

else
    y=zeros(size(x));
    sx=size(x,2);
    x=[nan(floor(m*.5),sx);x;nan(floor(m*.5),sx)];
    m1=m-1;
    for ii=1:size(y,1);
        y(ii,:)=fun(x(ii+(0:m1),:));
    end
    
end

return
```
