## Supplementary material for "A systems biology analysis of adrenergically stimulated adiponectin exocytosis in white adipocytes": All scripts used in this work: movingMedianIQM.m

```
function [y]=movingMedianIQM(x,range)
% movingMedianIQM will compute moving medians over a range, defined by the user.
%
% Usage: y = movingMedianIQM(x,range)
% where 
%      x:       is the input vector (or matrix) to be smoothed. 
%      range: 	percentage of number of points in x to be used to average over
%      y:       is output vector of same length as x
%
% Note:if x is a matrix then the smoothing will be done 'vertically'.
%
% This function is a wrapper for the movingAverageIQM function.
% 
% Examples:
%   x=randn(300,1);
%   plot(x,'g.'); 
%   hold on;
%   plot(movingMedianIQM(x,30),'k-');
%   plot(movingQuantileIQM(x,30,0.5),'r--');

% <<<COPYRIGHTSTATEMENT - IQM TOOLS LITE>>>

[y] = movingAverageIQM(x,range,'nanmedianIQM');
```
