## Supplementary material for "A systems biology analysis of adrenergically stimulated adiponectin exocytosis in white adipocytes": All scripts used in this work: smoothIQM.m

```
function c = smoothIQM(x,y,span,method,iter,weighting)
% Smoothes data using Robust or Non-robust Lowess smoother or with the
% Savitzky-Golay smoother. 
%
% Usage: Z = smoothIQM(X,Y,span,method,iter,weighting)
%        Z = smoothIQM(X,Y,span,'sgolay',degree)
%
% X,Y          input data
% span         number of points used to compute each element in Z, default
%              is 5 - smaller 1 its the fraction in the window
% method       'loess', 'lowess' (default), 'mean', 'rloess', 'rlowess', or
%              'rmean', 'sgolay'   
% iter         number of robust iterations, default is 5
% weighting    'tricubic' (default), 'gaussian' or 'linear'
% degree       degree of the polynomial fit for the Savitzky-Golay smoother
%
% Note: The difference between 'lowess' and 'loess' is that 'lowess' uses a
% linear model to do the local fitting (order = 1) whereas 'loess' uses a
% quadratic model to do the local fitting (order = 2). 'mean' is just a
% weighted local mean estimation (order = 0).

outputAsRow = diff(size(y))>0;
runSgolay = false;
xIsOwnEnumeration = false;

% Set x, y, and t
y = y(:);
t = numel(y);
if isempty(x)
    x = (1:t)';
    xIsOwnEnumeration = true;
elseif numel(x) == t
    x = x(:)+min(x)+1; % for better conditioning
else
    error('X and Y must be the same length.');
end

% Set span
if nargin<3 || isempty(span)
    span = 5;     % default span
elseif span <= 0  % span must be positive
    error('SPAN must be positive.'); 
elseif span < 1   % percent convention
    span = ceil(span*t); 
else              % span is given in samples, then round
    span = round(span);
end 

% Set method
if nargin<4 || isempty(method)
    order = 1; % default method
    robust = false; %default
else
    robust = method(1)=='r';
    switch method
        case {'loess','rloess'}
            order = 2;
        case {'lowess','rlowess'}
            order = 1;
        case {'mean','rmean'}  
            order = 0;
        case 'sgolay'
            runSgolay = true;
        otherwise
            error('Unknown smoothing method.')
    end
end

if runSgolay
    if nargin<5 || isempty(iter)
        degree = 2; %default degree for sgolay method
    else
        degree = iter;
    end
else % then it is any lowess method
    % Set number of iterations for robust
    if nargin<5 || isempty(iter)
        iter = 5;
    end
    if robust
        robust = iter;
    else
        robust = 0;
    end
end

if runSgolay
    if nargin>5
        error('Too many input arguments for SGOLAY method.')
    end
else % the it is any lowess method
    % Set weighting
    if nargin<6 || isempty(weighting)
        weighting = 3; % tricubic
    else
        switch weighting
            case 'tricubic'
                weighting = 3;
            case 'gaussian'
                weighting = 2;
            case 'linear'
                weighting = 1;
            otherwise
                error('Unknown weighting method.')
        end
    end
end
    
% is x sorted ?
if any(diff(x(~isnan(x)))<0)
    [x,idx] = sort(x);
    y = y(idx);
    unSort = true;
else
    unSort = false;
end

c = NaN(size(y),class(y));
xnotNaN = ~isnan(x);

% high limit for span
span = min(span,sum(xnotNaN));

% does x have repeated values ?
if xIsOwnEnumeration
    repeatedX = false;
else
    diffX = diff(x(xnotNaN));
    repeatedX = any(~diffX);
end

% are there NaN's in y ?
yNaNs = any(isnan(y(xnotNaN)));

% can we run 'Fast' Lowess/Sgolay instead ?
runFAlg = ~repeatedX && ~yNaNs && (span>4);
  % the fast alg 1) does not support repeated x values (otherwise the size
  % of the sliding window changes), 2) does not supports NaNs in y
  % (otherwise we need to check for NaN's in every window), and 3) does not
  % allow span <= 4 (otherwise we would need to check for likely square
  % matrices that generate singular solutions)
  
% can we run 'SuperFast' Lowess/Sgolay instead ?
runSFAlg = runFAlg && (xIsOwnEnumeration||((max(diffX)-min(diffX))<(max(x(xnotNaN))*eps)));
  % the super fast alg. is only for all above + equally distribited samples
        
if span == 1
    c(xnotNaN) = y(xnotNaN);
elseif runSgolay
    if runSFAlg
        c(xnotNaN) = sfsgolay(x(xnotNaN),y(xnotNaN),span,degree);
    elseif runFAlg
        c(xnotNaN) = fsgolay(x(xnotNaN),y(xnotNaN),span,degree);
    else
        c(xnotNaN) = sgolay(x(xnotNaN),y(xnotNaN),span,degree);
    end
else % then running Lowess Alg
    if runSFAlg
        c(xnotNaN) = sflowess(x(xnotNaN),y(xnotNaN),span,order,robust,weighting);
    elseif runFAlg
        c(xnotNaN) = flowess(x(xnotNaN),y(xnotNaN),span,order,robust,weighting);
    else
        c(xnotNaN) = lowess(x(xnotNaN),y(xnotNaN),span,order,robust,weighting);
    end
end

if unSort
    c(idx) = c;
end

if outputAsRow
    c = c';
end

%--------------------------------------------------------------------
function c = lowess(x,y,span,order,robust,weighting)
% LOWESS  Smoothes data using Robust Lowess method.
% 
% x,y             input data
% span            number of points used to compute each element in c
% method          2 = 'loess', 1 = 'lowess', or 0 = 'mean'
% robust          number of robust iterations, 0 means non-robust smoothing
% weighting       3' = 'tricubic',  2 = 'gaussian', or 1 = 'linear'
%
% The 'slow' version of LOWESS can handle missing values in y (i.e. NaNs),
% not evenly spaced x vector and repeated x values.

n = length(y);
c = zeros(size(y),class(y));

if (weighting == 2) && (span-1 <= order)
    span = order+2;
    warning('Rank deficiency when SPAN-1 <= ORDER, changing SPAN to %d.',span)    
elseif span-2 <= order
    span = order+3;
    warning('Rank deficiency when SPAN-2 <= ORDER, changing SPAN to %d.',span)
end
   
ws = warning;
warning('off');

% pre-allocate space for lower and upper indices for each fit
if robust>0
    lbound = zeros(n,1);
    rbound = zeros(n,1);
    dmaxv = zeros(n,1);
end

% compute some constants out of the loop
ynan = isnan(y);
anyNans = any(ynan(:));
seps = sqrt(eps);
theDiffs = [1; diff(x);1];

ll = 1;    % first window left point
rr = span; % first window right point

for i=1:n
    % if x(i) and x(i-1) are equal we just use the old value.
    if theDiffs(i) == 0
        c(i) = c(i-1);
        if robust>0
            lbound(i) = lbound(i-1);
            rbound(i) = rbound(i-1);
            dmaxv(i) = dmaxv(i-1);
        end    
        continue;
    end
    
    mx = x(i);                     % center
   
    while (rr<n) && (x(rr+1)-mx) < (mx-x(ll))
        rr = rr + 1;
        ll = ll + 1;
    end

    dmax = max(x(rr)-mx,mx-x(ll)); % maximum distance
        
    rrr=rr;                        % extend right limit ?
    while (rrr<n) && dmax==(x(rrr+1)-mx)
        rrr = rrr + 1;
    end
    lll=ll;                        % extend left limit ?
    while (lll>1) && dmax==(mx-x(lll-1))
        lll = lll - 1;
    end
    
    idx = lll:rrr;                 % indices for this window
   
    if anyNans                     % are there NaN's in yl
        idx = idx(~ynan(idx));     % remove them from my idx
        if isempty(idx)            
            c(i) = NaN;
            continue
        end
    end
    
    if dmax==0, dmax = 1; end      % in case all x's are the same 
       
    if robust
        lbound(i) = idx(1);
        rbound(i) = idx(end);
        dmaxv(i) = dmax;
    end
    
    % setting the current window
    xw = x(idx)-mx;     % centering to improve conditioning
    
    % outweight far samples
    switch weighting
        case 3 % tri-cubic weight
            weight = (1 - (abs(xw)/dmax).^3).^1.5; 
        case 2 % gaussian weight
            weight = exp(-(abs(xw)/dmax*2).^2); 
        case 1 % linear weight
            weight = 1 - abs(xw)/dmax;
    end
    if all(weight<seps)
        weight(:) = 1;    % if all weights are 0, just skip weighting
    end
    
    % least squares regression
    lw = length(xw);
    switch order
       case 2 % loess
           v = weight(:,[1 1 1]).*[ones(lw,1) xw xw.*xw];
       case 1 % lowess
           v = weight(:,[1 1]).*[ones(lw,1) xw];
       case 0 % local mean estimator
           c(i) = (weight'*y(idx)) / sum(weight);
           continue
    end
    
    if lw==order+1  % Square v may give infs in the \ solution ...
        b = [v;zeros(1,lw)]\[weight.*y(idx);0]; % ... so force least squares
    else
        b = v \ (weight.*y(idx));
    end
    c(i) = b(1);
end

% now that we have a non-robust fit, we can compute the residual and do
% the robust fit if required
maxabsyXeps = max(abs(y))*eps;
for k = 1:robust
    r = y-c;
    for i=1:n
        if theDiffs(i) == 0
            c(i) = c(i-1);
            continue;
        end
        if isnan(c(i)), continue; end
        idx = lbound(i):rbound(i);
        if anyNans
            idx = idx(~ynan(idx));
        end
        mx = x(i);
        xw = x(idx)-mx;  % centering to improve conditioning
        rw = r(idx);

        % outweight far samples
        switch weighting
            case 3 % tri-cubic weight
                weight = (1 - (abs(xw)/dmaxv(i)).^3).^1.5;
            case 2 % gaussian weight
                weight = exp(-(abs(xw)/dmaxv(i)*2).^2);
            case 1 % linear weight
                weight = 1 - abs(xw)/dmax;
        end
        if all(weight<seps)
            weight(:) = 1;    % if all weights 0, just skip weighting
        end

        % outweight outliers
        rw = abs(rw-median(rw));
        mad = median(rw);
        if mad > maxabsyXeps
            rw = 1-(rw./(6*mad)).^2;
            rw(rw<0) = 0;
            weight = weight .* rw; 
        end
        
        % least squares regression
        lw = length(xw);
        % correct low rank problems
        norder = min(order,sum(weight>0)-1);
        switch norder
           case 2 % loess
               v = weight(:,[1 1 1]).*[ones(lw,1) xw xw.*xw];
           case 1 % lowess
               v = weight(:,[1 1]).*[ones(lw,1) xw];
           case 0 % local mean estimator
               c(i) = (weight'*y(idx)) / sum(weight);
               continue
        end
    
        if lw==norder+1  % Square v may give infs in the \ solution ...
            b = [v;zeros(1,lw)]\[weight.*y(idx);0]; % ... so force least squares
        else
            b = v \ (weight.*y(idx));
        end
        c(i) = b(1);
    end
end

warning(ws);

%--------------------------------------------------------------------
function c = flowess(x,y,span,order,robust,weighting)
% FLOWESS  Smoothes data using Robust Lowess method.
%
% x,y             input data
% span            number of points used to compute each element in c
% method          2 = 'loess', 1 = 'lowess', or 0 = 'mean'
% robust          number of robust iterations, 0 means non-robust smoothing
% weighting       3' = 'tricubic',  2 = 'gaussian', or 1 = 'linear'
%
% The 'Fast' version of LOWESS assumes:
%
% 1) there are not repeated values in x, therefore the sliding window is
% size fixed and its limits can be computed by updating the limits of the
% previous window with a simple while-loop. 
%
% 2) there are not NaNs in the y vector, therefore there is not need to
% check for NaNs in y and take appropiate measures.
%
% 3) span>4, therefore there is not need to check for for likely square
% matrices that generate singular solutions.

n = length(y);

ws = warning('off');

c = zeros(size(y),class(y));                 % allocate output
               
ll = 1;    % first window left point
rr = span; % first window right point

for i = 1:n
    % update limits of the window
    mx = x(i);
    while (rr<n) && (x(rr+1)-mx) < (mx-x(ll))
        rr = rr + 1;
        ll = ll + 1;
    end
    if (rr<n) && (x(rr+1)-mx) == (mx-x(ll))
        rrr = rr + 1;
    else
        rrr = rr;
    end
    
    dmax = max(x(rrr)-mx,mx-x(ll));     % maximum value 
    xw = x(ll:rrr) - mx;  % center around x(i) to improve conditioning 
    
    % outweight far samples
    switch weighting
        case 3 % tri-cubic weight
            weight = (1 - (abs(xw)/dmax).^3).^1.5; 
        case 2 % gaussian weight
            weight = exp(-(xw/dmax*2).^2); 
        case 1 % linear weight
            weight = 1 - abs(xw)/dmax;
    end
    
    % least squares regression
    switch order
        case 2 % Loess 
            b = [weight weight.*xw weight.*xw.*xw] \ (weight.*y(ll:rrr));
            c(i) = b(1); % put estimated point into output vector
        case 1 % Lowess
            b = [weight weight.*xw] \ (weight.*y(ll:rrr));
            c(i) = b(1); % put estimated point into output vector
        case 0 % Local mean estimator
            c(i) = (weight'*y(ll:rrr)) / sum(weight);
    end
    
end
  
% now that we have a non-robust fit, we can compute the residual and do
% the robust fit if required
maxabsyXeps = max(abs(y))*eps;
for k = 1:robust
    r = y-c;
    ll = 1;
    rr = span;
    for i=1:n
        mx = x(i);
        while (rr<n) && (x(rr+1)-mx) < (mx-x(ll))
            rr = rr + 1;
            ll = ll + 1;
        end
        if (rr<n) && (x(rr+1)-mx) == (mx-x(ll))
            rrr = rr + 1;
        else
            rrr = rr;
        end
        dmax = max(x(rrr)-mx,mx-x(ll));
        xw = x(ll:rrr)-mx;  % centering for conditioning
        rw = r(ll:rrr);

        % outweight far samples
        switch weighting
            case 3 % tri-cubic weight
                weight = (1 - (abs(xw)/dmax).^3).^1.5;
            case 2 % gaussian weight
                weight = exp(-(xw/dmax*2).^2);
            case 1 % linear weight
                weight = 1 - abs(xw)/dmax;
        end

        % outweight outliers
        rw = abs(rw-median(rw));
        mad = median(rw);
        if mad > maxabsyXeps
            rw = 1-(rw./(6*mad)).^2;
            rw(rw<0) = 0;
            weight = weight .* rw;
        end

        % correct low rank problems
        norder = min(order,sum(weight>0)-1);
        % least squares regression
        switch norder
            case 2 % Loess
                b = [weight weight.*xw weight.*xw.*xw] \ (weight.*y(ll:rrr));
                c(i) = b(1); % put estimated point into output vector
            case 1 % Lowess
                b = [weight weight.*xw] \ (weight.*y(ll:rrr));
                c(i) = b(1); % put estimated point into output vector
            case 0 % Local mean estimator
                c(i) = (weight'*y(ll:rrr)) / sum(weight);
        end
        
    end
end

warning(ws);

%--------------------------------------------------------------------
function c = sflowess(x,y,span,order,robust,weighting)
% SFLOWESS  Smoothes data using Robust Lowess method.
%
% x,y             input data
% span            number of points used to compute each element in c
% method          2 = 'loess', 1 = 'lowess', or 0 = 'mean'
% robust          number of robust iterations, 0 means non-robust smoothing
% weighting       3' = 'tricubic',  2 = 'gaussian', or 1 = 'linear'
%
% The 'SuperFast' version of LOWESS assumes:
%
% 1) x is uniformely spaced, therefore a linear filter can be used.
%
% 2) there are not NaNs in the y vector, therefore there is not need to
% check for NaNs in y and take appropiate measures.
%
% 3) span>4, therefore there is not need to check for for likely square
% matrices that generate singular solutions.

n = length(y);

ws = warning('off');

% For uniform data, an even span actually covers an odd number of
% points.  For example, the four closest points to 5 in the
% sequence 1:10 are {3,4,5,6}, but 7 is as close as 3.
% Therfore force an odd span.
halfw = floor(span/2);
halfwp1 = halfw + 1;
span = 2*halfw + 1;
x1 = 1-halfw:halfw-1;

switch weighting
    case 3 % tri-cubic weight
        weight = (1 - (abs(x1)/halfw).^3).^1.5;
    case 2 % gaussian weight
        weight = exp(-(x1/halfw*2).^2);
    case 1 % linear weight
        weight = 1 - abs(x1)/halfw;
end
% Set up weighted Vandermonde matrix using equally spaced X values
switch order
    case 2
        V = [weight;weight.*x1;weight.*x1.*x1]';
    case 1
        V = [weight;weight.*x1]';
    case 0
        V = weight';
end

% Do QR decomposition
[Q,R] = qr(V,0); %#ok

% The projection matrix is Q*Q'. 
alpha = Q(halfw,:)*Q';

% Incorporate the weights into the coefficients of the linear combination,
% then apply filter. 
c = filter(alpha.*weight,1,y);

% We need to slide the values into the center of the array.
c(halfw+1:end-halfw) = c(span-1:end-1);

% Now we have taken care of everything except the end effects.  Loop over
% the points where we don't have a complete span.  Now the Vandermonde
% matrix has span-1 points, because only 1 has zero weight.
x1 = 1:span-1;
switch order
    case 2
        V = [ones(1,span-1);x1;x1.*x1]';
    case 1
        V = [ones(1,span-1);x1]';
    case 0
        V = ones(span-1,1);
end

for j=1:halfw
    % Compute weights based on deviations from the jth point,
    % then compute weights and apply them as above.
    switch weighting
        case 3 % tri-cubic weight
            weight = (1 - (abs((1:span-1) - j)/(span-j)).^3).^1.5;
        case 2 % gaussian weight
            weight = exp(-(abs((1:span-1) - j)/(span-j)*2).^2);
        case 1 % linear weight
            weight = 1 - abs((1:span-1) - j)/(span-j);
    end
    [Q,R] = qr(V.*repmat(weight(:),1,order+1),0);%#ok
    alpha = (Q(j,:)*Q') .* weight;
    c(j) = alpha * y(1:span-1);
    c(end+1-j) = alpha * y(end:-1:end-span+2);
end

% now that we have a non-robust fit, we can compute the residual and do
% the robust fit if required
maxabsyXeps = max(abs(y))*eps;
for k = 1:robust
    r = y-c;
    ll = 1;
    rr = span;
    for i=1:n
        
        if i>halfwp1 && rr<n
            rr = rr + 1;
            ll = ll + 1;
        end
        
        % update windows and weights only close to the edges where xw
        % changes, the weights at i==halfwp1 will be the ones used for all
        % the center samples
        if i<=halfwp1 || rr==n
            mx = x(i);
            dmax = max(x(rr)-mx,mx-x(ll));
            xw = x(ll:rr)-mx;  % centering for conditioning
            % outweight far samples
            switch weighting
                case 3 % tri-cubic weight
                    weight = (1 - (abs(xw)/dmax).^3).^1.5;
                case 2 % gaussian weight
                    weight = exp(-(xw/dmax*2).^2);
                case 1 % linear weight
                    weight = 1 - abs(xw)/dmax;
            end
            if (order == 2)  
                xws = xw .* xw;  
            end
        end

        % outweight outliers
        rw = r(ll:rr);
        rw = abs(rw-median(rw));
        mad = median(rw);
        if mad > maxabsyXeps
            rw = 1-(rw./(6*mad)).^2;
            rw(rw<0) = 0;
            rw = weight.*rw;
        else
            rw = weight;
        end

        % correct low rank problems
        norder = min(order,sum(weight>0)-1);
        % least squares regression
        switch norder
            case 2 % Loess
                b = [rw rw.*xw rw.*xws] \ (rw.*y(ll:rr));
                c(i) = b(1); % put estimated point into output vector
            case 1 % Lowess
                b = [rw rw.*xw] \ (rw.*y(ll:rr));
                c(i) = b(1); % put estimated point into output vector
            case 0 % Local mean estimator
                c(i) = sum(rw.*y(ll:rr)) / sum(rw);
        end
    end
end

warning(ws);

%--------------------------------------------------------------------
function c = sgolay(x,y,span,degree)
% SGOLAY  Smoothes data using Savitsky-Golay method.
% 
% x,y             input data
% span            number of points used to compute each element in c
% degree          degree of the polynomial fit
%
% The 'slow' version of SGOLAY can handle missing values in y (i.e. NaNs),
% not evenly spaced x vector and repeated x values.

n = length(y);
c = zeros(size(y),class(y));
ws = warning;
warning('off');
% compute some constants out of the loop
ynan = isnan(y);
anyNans = any(ynan(:));
theDiffs = [1; diff(x);1];
ll = 1;    % first window left point
rr = span; % first window right point
for i=1:n
    % if x(i) and x(i-1) are equal we just use the old value.
    if theDiffs(i) == 0
        c(i) = c(i-1);
        continue;
    end
    mx = x(i);                     % center
    while (rr<n) && (x(rr+1)-mx) < (mx-x(ll))
        rr = rr + 1;
        ll = ll + 1;
    end
    dmax = max(x(rr)-mx,mx-x(ll)); % maximum distance
    rrr=rr;                        % extend right limit ?
    while (rrr<n) && dmax==(x(rrr+1)-mx)
        rrr = rrr + 1;
    end
    lll=ll;                        % extend left limit ?
    while (lll>1) && dmax==(mx-x(lll-1))
        lll = lll - 1;
    end
    idx = lll:rrr;                 % indices for this window
    if anyNans                     % are there NaN's in yl
        idx = idx(~ynan(idx));     % remove them from my idx
        if isempty(idx)            
            c(i) = NaN;
            continue
        end
    end
    % centering and scaling to improve conditioning
    xw = (x(idx)./mx-1);       
    % least squares regression
    vrank = 1 + sum(diff(xw)>0);
    ncols = min(degree+1, vrank);
    lw = length(xw);
    v = ones(lw,ncols);
    for j = 1:ncols-1
        v(:,j+1) = xw.^j;
    end
    if lw==ncols
        % Square v may give infs in the \ solution, so force least squares
        b = [v;zeros(1,size(v,2))]\[y(idx);0];
    else
        b = v\y(idx);
    end
    c(i) = b(1);
end
warning(ws);

%--------------------------------------------------------------------
function c = fsgolay(x,y,span,degree)
% FSGOLAY  Smoothes data using Savitsky-Golay method.
% 
% x,y             input data
% span            number of points used to compute each element in c
% degree          degree of the polynomial fit
%
% The 'Fast' version of SGOLAY assumes:
%
% 1) there are not repeated values in x, therefore the sliding window is
% size fixed and its limits can be computed by updating the limits of the
% previous window with a simple while-loop. 
%
% 2) there are not NaNs in the y vector, therefore there is not need to
% check for NaNs in y and take appropiate measures.

n = length(y);
c = zeros(size(y),class(y));
ncols = min(degree+1, span);
ws = warning;
warning('off');
ll = 1;    % first window left point
rr = span; % first window right point

for i=1:n
    % update the limits of the window
    mx = x(i);                     % center
    while (rr<n) && (x(rr+1)-mx) < (mx-x(ll))
        rr = rr + 1;
        ll = ll + 1;
    end
    if (rr<n) && (x(rr+1)-mx) == (mx-x(ll))
        rrr = rr + 1;
        lw = span + 1;
    else
        rrr = rr;
        lw = span;
    end
    % centering and scaling to improve conditioning
    xw = (x(ll:rrr)./mx-1);     
    % least squares regression
    v = ones(lw,ncols);
    for j = 1:ncols-1
        v(:,j+1) = xw.^j;
    end
    if lw==ncols
        % Square v may give infs in the \ solution, so force least squares
        b = [v;zeros(1,size(v,2))]\[y(ll:rrr);0];
    else
        b = v\y(ll:rrr);
    end
    c(i) = b(1);
end
warning(ws);

%--------------------------------------------------------------------
function c = sfsgolay(x,y,span,degree)  %#ok
% SFSGOLAY  Smoothes data using Savitsky-Golay method.
% 
% x,y             input data
% span            number of points used to compute each element in c
% degree          degree of the polynomial fit
%
% The 'SuperFast' version of SGOLAY assumes:
%
% 1) x is uniformely spaced, therefore a linear filter can be used.
%
% 2) there are not NaNs in the y vector, therefore there is not need to
% check for NaNs in y and take appropiate measures.

n = length(y);
span = min(n-1,span);      % set upper limit for span
span = span+rem(span-1,2); % will add 1 if frame is even.
halfw = (span-1)/2;

V = ones(span,degree+1);
xw=(-halfw:halfw)';
for i=1:degree
    V(:,i+1)=xw.^i;
end
[Q,R]=qr(V,0); %#ok
ymid = filter(Q*Q(halfw+1,:)',1,y);
ybegin = Q(1:halfw,:)*Q'*y(1:span);
yend = Q((halfw+2):end,:)*Q'*y(n-span+1:n);
c = [ybegin;ymid(span:end);yend];
```
