## Supplementary material for "A systems biology analysis of adrenergically stimulated adiponectin exocytosis in white adipocytes": All scripts used in this work: fsolveIQM.m

```
function [X,FVAL,EXITFLAG] = fsolveIQM(FUN,X,varargin)
% fsolveIQM: attempts to solve equations of the form FUN(X)=0    
% where FUN and X may be vectors. Newton iteration.
% 
% USAGE:
% ======
% [X,FVAL,EXITFLAG] = fsolveIQM(FUN,X)
% [X,FVAL,EXITFLAG] = fsolveIQM(FUN,X,OPTIONS)
%
% FUN: Sunction to minimize/solve
% X: Starting Guess
% OPTIONS: structure containing options 
%          OPTIONS.MaxIter: Maximum number of iterations
%          OPTIONS.TolFun:  Tolerance for max element in function evaluation
%          OPTIONS.Delta:   Step length for numerical differentiation to 
%                           obtain the Jacobian
%
% DEFAULT VALUES:
% ===============
% OPTIONS.MaxIter: 1000
% OPTIONS.TolFun: 1e-11
% OPTIONS.Delta: 1e-8
%
% Output Arguments:
% =================
% X: Found solution
% FVAL: Value of the equations FUN at X
% EXITFLAG: 1=success, 0=not found

% <<<COPYRIGHTSTATEMENT - IQM TOOLS LITE>>>


MaxIter = 1000;
TolFun = 1e-11; 
Delta = 1e-8;

% Handle OPTIONS if given
if nargin == 3,
    OPTIONS = varargin{1};
    % TolFun
    if isfield(OPTIONS,'TolFun'),
        if ~isempty(OPTIONS.TolFun),
            TolFun = OPTIONS.TolFun;
        end
    end
    % MaxIter
    if isfield(OPTIONS,'MaxIter'),
        if ~isempty(OPTIONS.MaxIter),
            MaxIter = OPTIONS.MaxIter;
        end
    end
    % Delta
    if isfield(OPTIONS,'Delta'),
        if ~isempty(OPTIONS.Delta),
            Delta = OPTIONS.Delta;
        end
    end
end

EXITFLAG = 0;
for k = 1:MaxIter,
    FVAL = feval(FUN,X);
    Jacobian = getJacobian(FUN,X,Delta);
    X = X - 0.5*pinv(Jacobian)*FVAL;
    if norm(FVAL) < TolFun,
        EXITFLAG = 1;
        break
    end
end

if nargout < 3 && EXITFLAG == 0,
    disp('IQMsteadystate/fsolveIQM: Exceeded maximum number of iterations - check options');
    disp(sprintf('Last residual: %g',max(abs(FVAL))));
end
return

%%%%%%%%%%%%%%%%%%%%%%%%%%%%%%%%%%%%%%%%%%%%
% GET JACOBIAN AT GIVEN STATE
%%%%%%%%%%%%%%%%%%%%%%%%%%%%%%%%%%%%%%%%%%%%
function [Jacobian] = getJacobian(FUN,X,DELTA)
% size of system
n = length(X);          
% initialize jacobian variable
Jacobian = zeros(n,n);  
% determine the Jacobian by numerical differentiation
for k = 1:n,                
    Xup = X;
    Xup(k) = Xup(k)+DELTA;
    Jacobian(:,k) = (FUN(Xup)'-FUN(X)')/DELTA;
end
return
```
