## Supplementary material for "A systems biology analysis of adrenergically stimulated adiponectin exocytosis in white adipocytes": All scripts used in this work: boxplotIQM.m

```
function boxplotIQM(data,varargin)
% Plot a boxplot for given data.
%
% USAGE:
% ======
% boxplotIQM(data)        
% boxplotIQM(data,group)        
% boxplotIQM(data,OPTIONS)        
% boxplotIQM(data,group,OPTIONS)        
%
% data:  Matrix where each column contains data for which one box is plotted
%        Alternatively, data can also be a column vector. Depending on the
%        group argument either one box is plotted or several, depending on
%        the group information.
% group: Column vector, defining the grouping (by distinct numerical
%        values) of the data. In this case dara needs to be a column vector
%        of the same size as group.
%        
% OPTIONS: structure containing options for the function
%       OPTIONS.samplenames:        Cell-array with names or other information of the samples
%       OPTIONS.boxWidth:           Width of the boxes to be drawn
%       OPTIONS.filled:             Will fill the box (good for small boxWidth)
%       OPTIONS.verticalFlag:       Flag determining if the boxes are oriented
%                                   vertically (=1) or horizontally (=0)
%       OPTIONS.outliers:           0: do not plot them, 1: do plot them
%       OPTIONS.whiskerLength:      Whisker length relative to the length of the box
%       OPTIONS.whiskerPercentiles: If this option is defined then the
%                                   whiskers are not plotted based on
%                                   whiskerLength but based on the provided
%                                   percentiles in this option. Example:
%                                   whiskerPercentiles = [2.5 97.5] will
%                                   plot the whiskers at 2.5 and 97.5
%                                   percentiles of the data.
%       OPTIONS.axisij:             If plot is postprocessed with axisih
%                                   then this option needs to be set to 1.
%                                   Otherwise 0. 
%
% DEFAULT VALUES:
% ===============
% OPTIONS.samplenames:          Group/sample numbers
% OPTIONS.boxWidth:             0.5
% OPTIONS.filled:               0
% OPTIONS.verticalFlag:         0 (plot horizontally)
% OPTIONS.outliers:             1 (plot outliers)
% OPTIONS.whiskerLength:        1.5 (standard)
% OPTIONS.whiskerPercentiles:   [] (use whiskerLength)
% OPTIONS.axisij:               0

% <<<COPYRIGHTSTATEMENT - IQM TOOLS LITE>>>

% GLOBAL VARIABLES
global verticalFlag boxWidth whiskerLength outliers whiskerPercentiles filled

% Handle variable input arguments
if nargin==1,
    group   = [];
    OPTIONS = [];
elseif nargin==2,
    if isstruct(varargin{1}),
        OPTIONS = varargin{1};
        group   = [];
    else
        OPTIONS = [];
        group   = varargin{1};
    end
elseif nargin==3,
    group   = varargin{1};
    OPTIONS = varargin{2};
else
    error('Incorrect number of input arguments.');
end

if ~isempty(group),
    % Handle definition of group
    % If group defined then data needs to be a vector etc.
    if ~isvector(data),
        error('If group defined then data needs to be a vector of same size.');
    end
    % group nees to be a column vector
    if ~isvector(group),
        error('group needs to be a column vector.');
    end
    % data and group need to have same lengths
    if length(data) ~= length(group),
        error('data and group arguments need to have same length.');
    end
    % Enforce column vectors
    data    = data(:);
    group   = group(:);
else
    % Handle if group is not defined (then data can be vector or matrix)
    data_stacked    = [];
    group_stacked   = [];
    for k=1:size(data,2),
        data_stacked    = [data_stacked; data(:,k)];
        group_stacked   = [group_stacked; k*ones(length(data(:,k)),1)];
    end
    data    = data_stacked;
    group   = group_stacked;
end

% Determine number of samples (number groups) 
ALLgroups = unique(group);
nrgroups  = length(ALLgroups);

% Handle options
try samplenames             = OPTIONS.samplenames;          catch, samplenames          = {};   end
try boxWidth                = OPTIONS.boxWidth;             catch, boxWidth             = 0.5;  end
try filled                  = OPTIONS.filled;               catch, filled               = 0;    end
try verticalFlag            = OPTIONS.verticalFlag;         catch, verticalFlag         = 0;    end
try outliers                = OPTIONS.outliers;             catch, outliers             = 1;    end
try whiskerLength           = OPTIONS.whiskerLength;        catch, whiskerLength        = 1.5;  end
try whiskerPercentiles      = OPTIONS.whiskerPercentiles;   catch, whiskerPercentiles   = [];   end
try FLAG_axisij             = OPTIONS.axisij;               catch, FLAG_axisij          = 0;    end

% Option check
if ~isempty(samplenames) && length(samplenames) ~= nrgroups,
    error('Number of samplenames does not fit the number of samples.');
end

% Cycle through the groups and plot
hold on
for k=1:nrgroups
    dataPlot = data(group==ALLgroups(k));
    doplot(dataPlot,k);
end

% Annotate
Ymin = min(min(data)); Ymax = max(max(data));
DeltaY = 0.025*(Ymax-Ymin); % just a little bit of space on limits
if verticalFlag
    axis([[1-boxWidth, nrgroups+boxWidth] [(Ymin-DeltaY) (Ymax+DeltaY)]]);
    set(gca,'XTick',[1:nrgroups]);
    if ~isempty(samplenames), 
        set(gca, 'XTickLabel',samplenames); 
        try
            set(gca,'XTickLabelRotation',45);
        catch
        end
    else
        set(gca, 'XTickLabel',ALLgroups)
    end   
    set(gca,'XLim',[1-0.5 nrgroups+0.5]);    
else
    axis([[(Ymin-DeltaY) (Ymax+DeltaY)] [1-boxWidth, nrgroups+boxWidth]]);
    set(gca,'YTick',[1:nrgroups]);
    if ~isempty(samplenames), 
        set(gca, 'YTickLabel',samplenames); 
    else
        set(gca, 'YTickLabel',ALLgroups)
    end
    set(gca,'YLim',[1-0.5 nrgroups+0.5]);
end
% Whisker annotation if percentiles
if ~isempty(whiskerPercentiles),
    textWhiskers = sprintf('Whiskers: [%g%%, %g%%] of range',min(whiskerPercentiles),max(whiskerPercentiles));
    XLim = get(gca,'XLim');
    YLim = get(gca,'YLim');
    if ~FLAG_axisij,
        text(XLim(2),YLim(2),textWhiskers,'HorizontalAlign','Right','VerticalAlign','Top','FontSize',10);
    else
        text(XLim(2),YLim(1),textWhiskers,'HorizontalAlign','Right','VerticalAlign','Top','FontSize',10);
    end
end

return

%%%%%%%%%%%%%%%%%%%%%%%%%%%%%%%%%%%%%%%%%%%%%%%%%%%%%%%%%%%%%%%%%%%%%%%%%%
% PLOT FUNCTION FOR PLOTTING A SINGLE BOX WITH WHISKERS AND OUTLIERS
% THE FUNCTION ALSO NEEDS TO DETERMINE THE PERCENTILES
%%%%%%%%%%%%%%%%%%%%%%%%%%%%%%%%%%%%%%%%%%%%%%%%%%%%%%%%%%%%%%%%%%%%%%%%%%
function [] = doplot(sampledata,k)
global verticalFlag boxWidth whiskerLength outliers whiskerPercentiles filled
% Determine the percentiles
prctiles = prctileIQM(sampledata,[25 50 75]);
p25 = prctiles(1,:); p50 = prctiles(2,:); p75 = prctiles(3,:);
% Determine the whiskers
if isempty(whiskerPercentiles),
    % Determine the upper whisker position (needs to be on a sampledata point)
    index = find(sampledata <= p75+whiskerLength*(p75-p25));
    if isempty(index),
        upperWhisker = p75; % if no larger data points then use the 75 percentile
    else
        upperWhisker = max(sampledata(index)); % get the max sampledata point smaller than the 75% percentile
    end
    % Determine the upper whisker position (needs to be on a sampledata point)
    index = find(sampledata >= p25-whiskerLength*(p75-p25));
    if isempty(index),
        lowerWhisker = p25; % if no smaller data points then use the 25 percentile
    else
        lowerWhisker = min(sampledata(index)); % get the min sampledata point larger than the 25% percentile
    end
else
    % Determine whiskers based on percentiles
    upperWhisker = prctileIQM(sampledata,max(whiskerPercentiles));
    lowerWhisker = prctileIQM(sampledata,min(whiskerPercentiles));
end
% Determine the outliers
% Outliers are all the data points that lie outside the whiskers
outlier = sampledata([find(sampledata<lowerWhisker); find(sampledata > upperWhisker)]);
% Determine the right and the left (in a vertical sense) values
% for the box
rightValue = k+0.5*boxWidth;
leftValue = k-0.5*boxWidth;
% All things are determined so now the messy plot :)
if verticalFlag
    % plot the whiskers
    if ~filled,
        plot([k-0.25*boxWidth k+0.25*boxWidth],[upperWhisker upperWhisker],'b'); % plot end half as wide as the box
        plot([k-0.25*boxWidth k+0.25*boxWidth],[lowerWhisker lowerWhisker],'b'); % plot end half as wide as the box
        plot([k k],[lowerWhisker p25],'k--'); 
        plot([k k],[p75 upperWhisker],'k--');
    else
        plot([k k],[lowerWhisker p25],'k'); 
        plot([k k],[p75 upperWhisker],'k');
    end
    % plot the median
    if ~filled,
        plot([leftValue rightValue],[p50 p50],'r');
    else
        plot([k-min(0.5*boxWidth*6,0.5) k+min(0.5*boxWidth*6,0.5)],[p50 p50],'r');
    end
    % plot the outliers
    if outliers,
        plot(k*ones(1,length(outlier)),outlier,'k.');
    end
    % plot the 4 sides of the box
    if ~filled,
        plot([leftValue rightValue],[p75 p75],'b');
        plot([rightValue rightValue],[p75 p25],'b');
        plot([rightValue leftValue],[p25 p25],'b');
        plot([leftValue leftValue],[p25 p75],'b');
    else
         IQMplotfill([leftValue rightValue],[p25 p25],[p75 p75],[0 0 0],1); hold on;
    end
else
    % plot the whiskers
    if ~filled,
        plot([upperWhisker upperWhisker],[k-0.25*boxWidth k+0.25*boxWidth],'b'); % plot end half as wide as the box
        plot([lowerWhisker lowerWhisker],[k-0.25*boxWidth k+0.25*boxWidth],'b'); % plot end half as wide as the box
        plot([lowerWhisker p25],[k k],'k--'); 
        plot([p75 upperWhisker],[k k],'k--'); 
    else
        plot([lowerWhisker p25],[k k],'k'); 
        plot([p75 upperWhisker],[k k],'k'); 
    end
    % plot the median
    if ~filled,
        plot([p50 p50],[leftValue rightValue],'r');
    else
        plot([p50 p50],[k-min(0.5*boxWidth*6,0.5) k+min(0.5*boxWidth*6,0.5)],'r');
    end
    
    
    % plot the outliers
    if outliers,
        plot(outlier,k*ones(1,length(outlier)),'k.');
    end
    % plot the 4 sides of the box
    if ~filled,
        plot([p75 p75],[leftValue rightValue],'b');
        plot([p75 p25],[rightValue rightValue],'b');
        plot([p25 p25],[rightValue leftValue],'b');
        plot([p25 p75],[leftValue leftValue],'b');
    else
        IQMplotfill([p25 p75],[leftValue leftValue],[rightValue rightValue],[0 0 0],1); hold on
    end
    
end
return
```
