## Supplementary material for "A systems biology analysis of adrenergically stimulated adiponectin exocytosis in white adipocytes": All scripts used in this work: centerIQM.m

```
function [retval] = centerIQM(x, varargin)
% Center by subtracting means
%
% USAGE:
% ======
% retval = centerIQM(x)
% retval = centerIQM(x,dim)
%
% If x is a vector, subtract its mean. If x is a matrix, do the above for
% each column. If the optional argument dim is given, perform the above
% operation along this dimension 

% <<<COPYRIGHTSTATEMENT - IQM TOOLS LITE>>>

if (nargin ~= 1 && nargin ~= 2)
    print_usage ();
end

if (isvector (x))
    retval = x - mean (x, varargin{:});
elseif (ismatrixIQM (x))
    if nargin < 2
        dim = find (size (x) > 1, 1);
        if isempty (dim),
            dim=1;
        end;
    else
        dim = varargin{1};
    end
    sz = ones (1, ndims (x));
    sz (dim) = size (x, dim);
    retval = x - repmat (mean (x, dim), sz);
elseif (isempty (x))
    retval = x;
else
    error ('centerIQM: x must be a vector or a matrix');
end

return
```
