## Supplementary material for "A systems biology analysis of adrenergically stimulated adiponectin exocytosis in white adipocytes": All scripts used in this work: clusteringIQM.m

```
function [varargout] = clusteringIQM(dist,varargin)
% clusteringIQM: Performs UPGMA on distance matrix and produces a
% denddrogram plot.
% 
% Initially, each object is in its own cluster. At each step, the nearest 2
% clusters are combined into a higher-level cluster. The distance between 
% any 2 clusters A and B is taken to be the average of all distances
% between pairs of objects "a" in A and "b" in B. 
%
% USAGE:
% ======
% [] = clusteringIQM(dist,labels,fontsize)       
% [topology] = clusteringIQM(dist,labels,fontsize)
%
% dist:     [n x n] symmetric distance matrix
% labels:   optional cell-array with label names
% fontsize: optional font size for labels [default = 10]

% Output Arguments:
% =================
% If no output argument is specified the dendrogram is plotted.
%
%    topology   - [(n-1) x 4] matrix summarizing dendrogram topology:
%                 col 1 = 1st OTU/cluster being grouped at current step
%                 col 2 = 2nd OTU/cluster
%                 col 3 = ID of cluster being produced
%                 col 4 = distance at node

% RE Strauss, 5/27/96
%   9/7/99 - miscellaneous changes for Matlab v5.
%   9/24/01 - check diagonal elements against eps rather than zero.
%
% Adapted for the Systems Biology Toolbox 2 for MATLAB by
% Henning Schmidt, 01. January 2008

%%%%%%%%%%%%%%%%%%%%%%%%%%%%%%%%%%%%%%%%%%%%%%%
% Handle variable input arguments
%%%%%%%%%%%%%%%%%%%%%%%%%%%%%%%%%%%%%%%%%%%%%%%
labels = [];
fontsize = [];
if nargin < 1 || nargin > 3,
    error('Incorrect number of input arguments.');
elseif nargin == 2,
    labels = varargin{1};
elseif nargin == 3,
    labels = varargin{1};
    fontsize = varargin{2};
end
    
%%%%%%%%%%%%%%%%%%%%%%%%%%%%%%%%%%%%%%%%%%%%%%%
% Some checks
%%%%%%%%%%%%%%%%%%%%%%%%%%%%%%%%%%%%%%%%%%%%%%%
[n,p] = size(dist);
if n~=p || any(diag(dist)>eps),
    error('Input matrix is not a distance matrix - consider function "pdistIQM" to calculate it.');
end
if ~isempty(labels),
    if size(labels,1) ~= n,
        error('Numbers of taxa and taxon labels do not match');
    end
end

%%%%%%%%%%%%%%%%%%%%%%%%%%%%%%%%%%%%%%%%%%%%%%%
% Do the clustering
%%%%%%%%%%%%%%%%%%%%%%%%%%%%%%%%%%%%%%%%%%%%%%%
clstsize = ones(1,n);               % Number of elements in clusters/otus
id = 1:n;                           % Cluster IDs
topology = zeros(n-1,4);            % Output dendrogram-topology matrix

plug = 10e6;
dist = dist + eye(n)*plug;          % Replace diagonal with plugs

for step = 1:(n-1),                 % Clustering steps
    min_dist = min(dist(:));            % Find minimum pairwise distance
    [ii,jj] = find(dist==min_dist);     % Find location of minimum
    k = 1;                              % Use first identified minimum
    while (ii(k)>jj(k)),                 %   for which i<j
        k = k+1;
    end
    i = ii(k);
    j = jj(k);
    if (id(i)<id(j)),
        topology(step,:) = [id(i) id(j) n+step min_dist];
    else
        topology(step,:) = [id(j) id(i) n+step min_dist];
    end
    id(i) = n+step;
    dist(i,j) = plug;
    dist(j,i) = plug;

    new_clstsize = clstsize(i) + clstsize(j);
    alpha_i = clstsize(i) / new_clstsize;
    alpha_j = clstsize(j) / new_clstsize;
    clstsize(i) = new_clstsize;

    for k = 1:n,                         % For all other clusters/OTUs,
        if (k~=i && k~=j),                %   adjust distances to new cluster
            dist(k,i) = alpha_i * dist(k,i) + alpha_j * dist(k,j);
            dist(i,k) = alpha_i * dist(i,k) + alpha_j * dist(j,k);
            dist(k,j) = plug;
            dist(j,k) = plug;
        end
    end
end

%%%%%%%%%%%%%%%%%%%%%%%%%%%%%%%%%%%%%%%%%%%%%%%
% Handle output
%%%%%%%%%%%%%%%%%%%%%%%%%%%%%%%%%%%%%%%%%%%%%%%
if nargout == 1,
    varargout{1} = topology;
else
    figure; clf;
    plot_dendrogram(topology,labels,fontsize);
end    
return
```
