## Supplementary material for "A systems biology analysis of adrenergically stimulated adiponectin exocytosis in white adipocytes": All scripts used in this work: kurtosisIQM.m

```
function retval = kurtosisIQM(x, dim)
% If 'x' is a vector of length 'N', return the kurtosis
% of 'x'.  If 'x' is a matrix, return the kurtosis over the
% first non-singleton dimension. The optional argument 'dim'
% can be given to force the kurtosis to be given over that 
% dimension.
%
% USAGE:
% ======
% retval = kurtosisIQM(x, dim)

% <<<COPYRIGHTSTATEMENT - IQM TOOLS LITE>>>

  if (nargin ~= 1 && nargin ~= 2)
    error('Incorrect number of input arguments'); 
  end

  nd = ndims (x);
  sz = size (x);
  if (nargin ~= 2)
    % Find the first non-singleton dimension.
    dim  = 1;
    while (dim < nd + 1 && sz(dim) == 1)
      dim = dim + 1;
    end
    if (dim > nd)
      dim = 1;
    end
  else
    if (~(isscalar (dim) && dim == round (dim)) && dim > 0	&& dim < (nd + 1))
      error ('dim must be an integer and valid dimension');
    end
  end
  
  if (~ismatrixIQM(x) && ~isvector(x))
    error ('x has to be a matrix or a vector');
  end

  c = sz(dim);
  sz(dim) = 1;
  idx = ones (1, nd);
  idx(dim) = c;
  x = x - repmat (mean (x, dim), idx);
  retval = zeros (sz);
  s = std (x, [], dim);
  x = sum(x.^4, dim);
  ind = find (s > 0);
  retval(ind) = x(ind) / (c*s(ind) .^ 4);

return
```
