## Supplementary material for "A systems biology analysis of adrenergically stimulated adiponectin exocytosis in white adipocytes": All scripts used in this work: mvnrndIQM.m

```
function [s] = mvnrndIQM(mu,Sigma,K)
% Draw n random d-dimensional vectors from a multivariate Gaussian distribution
% with mean mu and covariance matrix Sigma.
%
% USAGE:
% ======
% [s] = mvnrndIQM(mu,Sigma)
% [s] = mvnrndIQM(mu,Sigma,K)
%
% K: number of samples

% <<<COPYRIGHTSTATEMENT - IQM TOOLS LITE>>>

% If mu is column vector and Sigma not a scalar then assume user didn't read
% help but let them off and flip mu. Don't be more liberal than this or it will
% encourage errors (eg what should you do if mu is square?).
if ((size(mu,2)==1) && (size(Sigma)~=[1,1])),
	mu=mu';
end

% May 2004 take a third arg, cases. Makes it more compatible with Matlab's.
if nargin==3
	mu=repmat(mu,K,1);
end

[n,d]=size(mu);

if (size(Sigma)~=[d,d])
	error('Sigma must have dimensions dxd where mu is nxd.');
end

try
	U=chol(Sigma);
catch
	[E,Lambda]=eig(Sigma);
% 	if (min(diag(Lambda))<0),error('Sigma must be positive semi-definite.'),end
	U = sqrt(Lambda)*E';
end

s = randn(n,d)*U + mu;

% {{{ END OF CODE --- Guess I should provide an explanation:
% 
% We can draw from axis aligned unit Gaussians with randn(d)
% 	x ~ A*exp(-0.5*x'*x)
% We can then rotate this distribution using
% 	y = U'*x
% Note that
% 	x = inv(U')*y
% Our new variable y is distributed according to:
% 	y ~ B*exp(-0.5*y'*inv(U'*U)*y)
% or
% 	y ~ N(0,Sigma)
% where
% 	Sigma = U'*U
% For a given Sigma we can use the chol function to find the corresponding U,
% draw x and find y. We can adjust for a non-zero mean by just adding it on.
% 
% But the Cholsky decomposition function doesn't always work...
% Consider Sigma=[1 1;1 1]. Now inv(Sigma) doesn't actually exist, but Matlab's
% mvnrnd provides samples with this covariance st x(1)~N(0,1) x(2)=x(1). The
% fast way to deal with this would do something similar to chol but be clever
% when the rows aren't linearly independent. However, I can't be bothered, so
% another way of doing the decomposition is by diagonalising Sigma (which is
% slower but works).
% if
% 	[E,Lambda]=eig(Sigma)
% then
% 	Sigma = E*Lambda*E'
% so
% 	U = sqrt(Lambda)*E'
% If any Lambdas are negative then Sigma just isn't even positive semi-definite
% so we can give up.
% }}}
```
