## Supplementary material for "A systems biology analysis of adrenergically stimulated adiponectin exocytosis in white adipocytes": All scripts used in this work: nanmeanIQM.m

```
function [v] = nanmeanIQM(X,varargin)
% Determines the mean of x along the dimension dim by treating NaNs as
% missing values.
%
% USAGE:
% ======
% output = nanmeanIQM(x)
% output = nanmeanIQM(x,dim)
%
% If x is a matrix and dim not defined then determine mean across rows
% (dim=1). 

% <<<COPYRIGHTSTATEMENT - IQM TOOLS LITE>>>

n           = sum (~isnan(X), varargin{:});
n(n == 0)   = NaN;
X(isnan(X)) = 0;
v           = sum (X, varargin{:}) ./ n;
```
