## Supplementary material for "A systems biology analysis of adrenergically stimulated adiponectin exocytosis in white adipocytes": All scripts used in this work: nanmedianIQM.m

```
function v = nanmedianIQM(X,varargin)
% Determines the median of x along the dimension dim by treating NaNs as
% missing values.
%
% USAGE:
% ======
% output = nanmedianIQM(x)
% output = nanmedianIQM(x,dim)
%
% If x is a matrix and dim not defined then determine mean across rows
% (dim=1). 

% <<<COPYRIGHTSTATEMENT - IQM TOOLS LITE>>>

if nargin < 1 || nargin > 2
    error('Incorrect number of input arguments');
end

if nargin < 2
    dim = min(find(size(X)>1));
    if isempty(dim), dim=1; end;
else
    dim = varargin{:};
end

sz = size (X);
if (prod (sz) > 1)
    % Find lengths of datasets after excluding NaNs; valid datasets
    % are those that are not empty after you remove all the NaNs
    n = sz(dim) - sum (isnan(X),varargin{:});
    
    % When n is equal to zero, force it to one, so that median
    % picks up a NaN value below
    n (n==0) = 1;
    
    % Sort the datasets, with the NaN going to the end of the data
    X = sort (X, varargin{:});
    
    % Determine the offset for each column in single index mode
    colidx = reshape((0:(prod(sz) / sz(dim) - 1)), size(n));
    colidx = floor(colidx / prod(sz(1:dim-1))) * prod(sz(1:dim)) + mod(colidx,prod(sz(1:dim-1)));
    stride = prod(sz(1:dim-1));
    
    % Average the two central values of the sorted list to compute
    % the median, but only do so for valid rows.  If the dataset
    % is odd length, the single central value will be used twice.
    % E.g.,
    %   for n==5, ceil(2.5+0.5) is 3 and floor(2.5+0.5) is also 3
    %   for n==6, ceil(3.0+0.5) is 4 and floor(3.0+0.5) is 3
    % correction made for stride of data "stride*ceil(2.5-0.5)+1"
    v = (X(colidx + stride*ceil(n./2-0.5) + 1)  + X(colidx + stride*floor(n./2-0.5) + 1)) ./ 2;
elseif (prod (sz) == 1)
    v = X;
else
    error ('nanmedianIQM: invalid matrix argument');
end
```
