## Supplementary material for "A systems biology analysis of adrenergically stimulated adiponectin exocytosis in white adipocytes": All scripts used in this work: nanstdIQM.m

```
function v = nanstdIQM(X, varargin)
% Determines the std of x along the dimension dim by treating NaNs as
% missing values.
%
% USAGE:
% ======
% output = nanstdIQM(x)
% output = nanstdIQM(x,dim)
%
% If x is a matrix and dim not defined then determine mean across rows
% (dim=1). 

% <<<COPYRIGHTSTATEMENT - IQM TOOLS LITE>>>

if nargin < 1 || nargin > 2
    error('Incorrect number of input arguments');
end

if nargin < 2
    dim = min(find(size(X)>1));
    if isempty(dim), dim=1; end;
else
    dim = varargin{1};
end
opt = 0;

% determine the number of non-missing points in each data set
n = sum (~isnan(X), varargin{:});

% replace missing data with zero and compute the mean
X(isnan(X)) = 0;
meanX = sum (X, varargin{:}) ./ n;

% subtract the mean from the data and compute the sum squared
sz = ones(1,length(size(X)));
sz(dim) = size(X,dim);
v = sum((X - repmat(meanX,sz)).^2, varargin{:});

% because the missing data was set to zero each missing data
% point will contribute (-meanX)^2 to sumsq, so remove these
v = v - (meanX .^ 2) .* (size(X,dim) - n);

if (opt == 0)
    % compute the standard deviation from the corrected sumsq using
    % max(n-1,1) in the denominator so that the std for a single point is 0
    v = sqrt ( v ./ max(n - 1, 1) );
elseif (opt == 1)
    % compute the standard deviation from the corrected sumsq
    v = sqrt ( v ./ n );
else
    error ('nanstdIQM: unrecognized normalization type');
end

% make sure that we return a real number
v = real (v);
```
