## Supplementary material for "A systems biology analysis of adrenergically stimulated adiponectin exocytosis in white adipocytes": All scripts used in this work: pdistIQM.m

```
function [D] = pdistIQM(datamatrix,varargin)
% pdistIQM: Determines the distance matrix for a set of points whose
% coordinates are given as row-vectors in the data matrix.
%
% USAGE:
% ======
% D = pdistIQM(datamatrix) 
% D = pdistIQM(datamatrix, method) 
%
% datamatrix: datamatrix is an NxP matrix of coordinates for N points in P dimensions
% method: is an integer between 1 and 3 representing the chosen
%         method for computing the distance matrix (see note below)
%
% DEFAULT VALUES:
% ===============
% method: method chosen problem size dependend if not specified
%
% Output Arguments:
% =================
% D: An NxN matrix, where the value of DMAT(i,j) corresponds to
%           the distance from datamatrix(i,:) to datamatrix(j,:)
% Note:
% =====
% method=1: Usually fastest for small inputs. Takes advantage of the symmetric
%           property of distance matrices to perform half as many calculations
% method=2: Usually fastest for medium inputs. Uses a fully vectorized method
% method=3: Usually fastest for large inputs. Uses a partially vectorized
%           method with relatively small memory requirement

% Author: Joseph Kirk
% Email: jdkirk630 at gmail dot com
% Release: 1.0
% Release Date: 5/29/07
%
% Adapted for the Systems Biology Toolbox 2 for MATLAB by
% Henning Schmidt, 01. January 2008

% Handle variable input arguments and select the method
if nargin < 1 || nargin > 2,
    error('Incorrect number of input arguments.');
end
if nargin == 2,
    method = varargin{1};
else
    [n,dims] = size(datamatrix);
    numels = n*n*dims;
    method = 2; 
    if numels > 5e4, 
        method = 3; 
    elseif n < 20, 
        method = 1; 
    end
end

% distance matrix calculation options
switch method
    case 1 % half as many computations (symmetric upper triangular property)
        [k,kk] = find(triu(ones(n),1));
        D = zeros(n);
        D(k+n*(kk-1)) = sqrt(sum((datamatrix(k,:) - datamatrix(kk,:)).^2,2));
        D(kk+n*(k-1)) = D(k+n*(kk-1));
    case 2 % fully vectorized calculation (very fast for medium inputs)
        a = reshape(datamatrix,1,n,dims);
        b = reshape(datamatrix,n,1,dims);
        D = sqrt(sum((a(ones(n,1),:,:) - b(:,ones(n,1),:)).^2,3));
    case 3 % partially vectorized (smaller memory requirement for large inputs)
        D = zeros(n,n);
        for k = 1:n
            D(k,:) = sqrt(sum((datamatrix(k*ones(n,1),:) - datamatrix).^2,2));
        end
end
```
