## Supplementary material for "A systems biology analysis of adrenergically stimulated adiponectin exocytosis in white adipocytes": All scripts used in this work: prctileIQM.m

```
function [Q] = prctileIQM(Y,q,DIM)
% prctileIQM calculates the percentiles of histograms and sample arrays.  
%
% USAGE:
% ======
% Q = prctileIQM(Y,q)
% Q = prctileIQM(Y,q,DIM)
%
% Returns the q-th percentile along dimension DIM of sample array Y.
% size(Q) is equal size(Y) except for dimension DIM which is size(Q,DIM)=length(Q)

% <<<COPYRIGHTSTATEMENT - IQM TOOLS LITE>>>

if nargin==2,
	Q = quantileIQM(Y,q/100); 
elseif nargin==3,
	Q = quantileIQM(Y,q/100,DIM); 
else
    error('Incorrect number of input arguments.');
end
```
