## Supplementary material for "A systems biology analysis of adrenergically stimulated adiponectin exocytosis in white adipocytes": All scripts used in this work: princompIQM.m

```
function [pc,z,w,Tsq] = princompIQM(X)
% princompIQM: Compute principal components of X
%
% USAGE:
% ======
% [pc,z,w,Tsq] = princomp(X)
%
% pc:  the principal components
% z:   the transformed data
% w:   the eigenvalues of the covariance matrix
% Tsq: Hotelling's T^2 statistic for the transformed data

% <<<COPYRIGHTSTATEMENT - IQM TOOLS LITE>>>

C = cov(X);
[U,D,pc] = svd(C);
z = centerIQM(X)*pc;
w = diag(D);
Tsq = sumsqIQM(zscoreIQM(z),2);
```
