## Supplementary material for "A systems biology analysis of adrenergically stimulated adiponectin exocytosis in white adipocytes": All scripts used in this work: qqplotIQM.m

```
function [q,s] = qqplotIQM(x)
% Perform a QQ-plot (quantile plot). Only for comparison to standard normal
% distribution.
%
% USAGE:
% ======
% [] = qqplotIQM(x)

% <<<COPYRIGHTSTATEMENT - IQM TOOLS LITE>>>

if ~isvector(x),
    error ('qqplotIQM: x must be a vector');
end

s = sort (x);
n = length (x);
t = ((1 : n)' - .5) / n;
if (nargin == 1)
    f = 'stdnormalinvIQM';
else
    f = str2func (sprintf ('%s_inv', dist));
end

if (nargin <= 2)
    q = feval (f, t);
    q_label = f;
else
    param_string = sprintf ('%g', varargin{1});
    for k = 2 : (nargin - 2);
        param_string = sprintf ('%s, %g', param_string, varargin{k});
    end
    q = eval (sprintf ('%s (t, %s);', f, param_string));
    q_label = sprintf ('%s with parameter(s) %s', func2str (f), param_string);
end

% colors
colors = IQMgetcolors();

% Compute the line the y=x line
dx = prctileIQM(q, 75) - prctileIQM(q, 25);
dy = prctileIQM(s, 75) - prctileIQM(s, 25);
b = dy./dx;                                % slope
xc = (prctileIQM(q, 25) + prctileIQM(q, 75))/2;  % center points
yc = (prctileIQM(s, 25) + prctileIQM(s, 75))/2;  % ...
ymax = yc + b.*(max(q)-xc);
ymin = yc - b.*(xc-min(q));

% Plot the qqplot
plot (q,s,'x','Color',colors(1,:),'MarkerSize',12,'LineWidth',2); hold on
plot([min(q); max(q)], [ymin; ymax],'k--','LineWidth',2); 

% Annotate
xlabel (q_label);
ylabel ('Sample points');
grid on
```
