## Supplementary material for "A systems biology analysis of adrenergically stimulated adiponectin exocytosis in white adipocytes": All scripts used in this work: getMONOLIXversionIQM.m

```
function [MLXversion] = getMONOLIXversionIQM()
% Function tries to determine the MONOLIX version that is used with IQM
% Tools. Since Lixoft does not understand the need of some users to access
% the version number by a script we try to infer the version from the path
% in which MONOLIX is installed.
%
% MLXversion: output argument with the following possibilities
%       '432': Version 4.3.2
%       '433': Version 4.3.3
%       'unknown': Not identified

% Get the MONOLIX path
PATH_MONOLIX = getNLMEtoolInfoIQM();

% Remove special signs
test = regexprep(PATH_MONOLIX,'\W','');

% Check 4.3.2
ix432 = strfind(test,'432');

% Check 4.3.3
ix433 = strfind(test,'433');

% Assign output argument
if isempty(ix432) && isempty(ix433),
    MLXversion = 'unknown';
end

if ~isempty(ix432) && isempty(ix433),
    MLXversion = '432';
end
 
if isempty(ix432) && ~isempty(ix433),
    MLXversion = '433';
end
```
