## Supplementary material for "A systems biology analysis of adrenergically stimulated adiponectin exocytosis in white adipocytes": All scripts used in this work: getNLMEtoolInfoIQM.m

```
function [PATH_MONOLIX,PATH_MONOLIX_PAR,PATH_NONMEM,PATH_NONMEM_PAR] = getNLMEtoolInfoIQM()
% Function loads and returns the paths to MONOLIX and NONMEM executables, set-up in the 
% SETUP_PATHS_TOOLS_IQMPRO file

% Run the SETUP_NLME_TOOLS script
SETUP_PATHS_TOOLS_IQMPRO

if isunix,
    PATH_MONOLIX        = PATH_SYSTEM_MONOLIX_UNIX;
    PATH_MONOLIX_PAR    = PATH_SYSTEM_MONOLIX_PARALLEL_UNIX;
    PATH_NONMEM         = PATH_SYSTEM_NONMEM_UNIX;
    PATH_NONMEM_PAR     = PATH_SYSTEM_NONMEM_PARALLEL_UNIX;
else
    PATH_MONOLIX        = PATH_SYSTEM_MONOLIX_WINDOWS;
    PATH_MONOLIX_PAR    = PATH_SYSTEM_MONOLIX_PARALLEL_WINDOWS;
    PATH_NONMEM         = PATH_SYSTEM_NONMEM_WINDOWS;
    PATH_NONMEM_PAR     = PATH_SYSTEM_NONMEM_PARALLEL_WINDOWS;
end
```
