## Supplementary material for "A systems biology analysis of adrenergically stimulated adiponectin exocytosis in white adipocytes": All scripts used in this work: Contents.m

```
% IQM Tools Pro
% Version 1.2.2 (R2015B) 02.01.2017
% 
% IQMpro
% =======
% 	Contents                                               - This help text
% 	SETUP_PATHS_TOOLS_IQMPRO                               - Setup script for IQMpro - please customize according to your needs
% 	installIQMpro                                          - This script installs the IQMpro package of the IQM Suite of modeling tools
% 
% Project ceation and handling (Sysbio/syspharm)
% ==============================================
% 	IQMprojectSB                                           - Creates a project object containing models, experiments, measurements, and information needed for parameterestimation
% 	IQMexportproject                                       - Exports a given project to a projectfolder
% 	IQMsaveproject                                         - Save an IQMprojectSB object as a binary MATLAB MAT fil with .iqmp as extension
% 
% projectSBhandling
% 	IQManalyzeresiduals                                    - Determines and analyzes the residuals for a given project
% 	IQMcomparemeasurements                                 - Simulates the experiments in the project for the models in the project and compares the simulated results to the  measurements
% 	IQMcreaterunestimationscript                           - Creates a run estimation script for the  given project and eventually given modelindex
% 	IQMexportestimation                                    - Exports the selected estimation settings in an IQMprojectSB into a flat text file
% 	IQMgetexperiment                                       - Get experiment(s) from the given project
% 	IQMgetmeasurement                                      - Get measurement(s) from the given project and  experiment
% 	IQMgetmodel                                            - Get model(s) from the given project
% 	IQMidentifiability                                     - Performs parameter identifiability analysis for a  given project
% 	IQMinfo                                                - Quick dump of contents of an IQMprojectSB
% 	IQMinsilicoexpproj                                     - Runs an in-silico experiment directly on a model and  experiment within a IQMprojectSB
% 	IQMplotmeasurements                                    - Plots all measurements in the given IQMprojectSB using IQMplot
% 	IQMreducerateexpressionsProject                        - Function allowing the interactive and iterative reduction of complex kinetic rate expressions
% 	IQMupdateexperiment                                    - Update or add a experiment in a project
% 	IQMupdatemeasurement                                   - Update or add a measurement in an experiment of a project
% 	IQMupdatemodel                                         - Update or add a model in a project
% 	isIQMprojectSB                                         - Checks if provided input argument is an IQMprojectSB
% 
% High performace simulation, using MEX models
% ============================================
% 	IQMPsimulate                                           - Function allowing to simulate IQMmodels and MEX simulation functions
% 	IQMsensitivity                                         - Determine the sensitivities of states and variables with respect to initial conditions and parameters, using a finite difference approach.
% 	
%   IQMmakeMEXmodel                                        - Converts an IQMmodel to an executable C-code MEX model
% 	IQMmakeTempMEXmodel                                    - Used to create a temporary MEX simulation function
% 
% Systems Modeling Tools
% ======================
% 	IQMparametercorrelation                                - Function for determining parameter correlations based on parametric sensitivities
% 	IQMmanualtuning                                        - Allows to compare simulated experiments data to measurements and perform manual parameter tuning
% 	IQMmodeltuning                                         - Allows to compare and tune a model to one or more sets of data
% 	IQMparamestGUI                                         - Graphical user-interface for running parameter estimation tasks
% 
% 	plotIQMP                                               - Allows to compare simulated experiments data to measurements
% 
%   IQMparameterestimation                                 - This function performs parameter estimation for  a given IQM project
% 	defaultcostparameterestimationIQM                      - Default cost function used for parameter estimation. Can be customized and passed to IQMparameterestimation
% 
% 	IQMfaboxplot                                           - Plots a box-and-whisker diagram for the estimation data
% 	IQMfaclustering                                        - Performs hierarchical clustering based on Euclidean distance of the estimated parameter sets
% 	IQMfacorr                                              - Determines the correlation matrix for the parameter sets determined with the IQMparameterfitanalysis function
% 	IQMfahist                                              - Plots histograms for the parameter values that have been estimated using the IQMparameterfitanalysis function
% 	IQMfasigncorr                                          - A matrix of p-values for testing the hypothesis of no significant correlation is computed 
% 	IQMparameterfitanalysis                                - Parameter estimation for the given project is repeated several times from randomly perturbed starting conditions
% 
% 	IQMfadetcorr                                           - Plots detailed pairwise correlations between parameters
%     
% Clinical Data Tools
% ===================
% IQM Tools use a standard format for clinical data. This format is documented in the function IQMcheckGeneralDataFormat.
% 
% GeneralDataFormat
% 	IQMaddNLMEinfo2data                                    - Adds some analysis and NLME tool relevant information into the provided dataset
% 	IQMcheckGeneralDataFormat                              - Check the format of a dataset against the requirements of the IQM Tools General Dataset format for clinical data
% 	IQMcheckGeneralDataFormatHeader                        - Similar to IQMcheckGeneralDataFormat but check only the header names but not internal consistency
% 	IQMcreateGeneralDataset                                - Create an empty general dataset
% 	IQMdataAddTimeDependentCovariate                       - Adds user defined time dependent continuous and  categorical covariates to a dataset in the general data format.
% 	IQMdataAddTimeIndependentCovariate                     - Adds user defined time independent continuous and  categorical covariates to a dataset in the general data format.
% 	IQMgenerateVALUEfromVALUE_TEXT                         - Generate numerical values in VALUE for VALUETXT definitions
% 	isGeneralDatasetIQM                                    - Checks if the dataset "data" is in the general dataset format
% 
% ProcessData
% 	IQMdataGetBaselineValues                               - Calculate the baseline values of specified  readouts
% 	IQMdataGetValues                                       - Extract values for each individual for event NAME from the dataset 
% 	IQMselectDataEvents                                    - Allows to select events to retain in the dataset
% 	IQMsubsetData                                          - Allows to subset individual subjects in clinical datasets
% 
% TaskDataset
% 	IQMcheckTaskDataset                                    - Check if dataset in task specific dataset format
% 	IQMcheckTaskDatasetHeader                              - Similar to IQMcheckTaskDataset but check only the header names but not internal consistency
% 	IQMconvertGeneral2TaskDataset                          - Augments a dataset in the general data format, used in IQM Tools, with information needed for analysis
% 	isTaskDatasetIQM                                       - Checks if the dataset "data" is in the general task dataset format
% 
% DataExploration
% 	IQMdataInfoValues                                      - Function to get information about the link between VALUETXT and VALUE
% 	IQMexploreBLLOQdata                                    - Displays a table with information about the number of BLOQ data per NAME
% 	IQMexploreCovariateCorrelations                        - Graphical exploration of covariates 
% 	IQMexploreDataContents                                 - Produces a table, showing the study numbers, the study description, the contained treatment groups in the studies, etc.
% 	IQMexploreDataMedian                                   - Plot data with respect to the medians of selected readouts
% 	IQMexploreDataMedianCovariates                         - Plot data with respect to the medians of selected readouts with covariate information
% 	IQMexploreDataVariability                              - Explore varibility of readouts in the data
% 	IQMexploreDistribution                                 - Explore distribution of readouts in the data
% 	IQMexploreDistributionCorrelation                      - Explore distribution of readouts in the data with covariate information
% 	IQMexploreIndivData                                    - Plot individual data for on readout from the task specific dataset used in IQM Tools
% 	IQMexploreIndivDataRelation                            - Plot individual data for multiple readouts from the task specific dataset used in IQM Tools
% 	IQMexploreMissingEventRate                             - Rate of missing events over NT by group
% 	IQMexploreMissingObservationGraphics                   - Missing observations in the dataset
% 	IQMexploreNTvsTIME                                     - NT vs TIME
% 	IQMexplorePDdataWrapper                                - Function generating typical plots for PD data - bot continuous and categorical
% 	IQMexplorePKdataWrapper                                - Function generating typical standard plots for dose and PK data
% 	IQMexploreSummaryStats                                 - Produces summary statistics for the provided dataset
% 
% NLME Dataset Preparation
% ========================
% DataCleaning
% 	IQMcleanImputeCovariates                               - Imputation of missing covariates
% 	IQMcleanRemoveIGNOREDrecords                           - Removes all records from the dataset that have non empty (or NaN) entries in the IGNORE column
% 	IQMcleanRemovePlaceboSubjects                          - Removes all subjects which only received 0 doses or  no doses at all
% 	IQMcleanRemoveRecordsSUBJECTs                          - Removes defined subjects (by USUBJID) and records (by index of  the row in which the record is located)
% 	IQMcleanRemoveSubjectsNoObservations                   - Removes all subjects which do not have any  observations (defined by EVID=0 and MDV=0)
% 	IQMcleanRemoveZeroDoses                                - Removes all dose records with 0 amount
% 
% BLOQhandling
% 	IQMhandleBLOQdata                                      - Allows handling of M1,3,4,5,6,7 methods for BLOQ
% 
% DataNLMEConversion
% 	IQMaddIndivRegressionParamFromNLMEproject              - Allows to add columns to dataset to be used as regression parameters in the model
% 	IQMcheckNLMEdatasetHeader                              - Checks if the minimal required elements are present in an NLME dataset that is going to be used for fitting
% 	IQMconvertTask2NLMEdataset                             - Converts a task specific general dataset into a dataset that is suitable for NLME analysis in NONMEM and MONOLIX
% 	IQMgetNLMEdataHeader                                   - Generates information about the dataset header needed for NLME model creation
% 	IQMgetRegressionParameters                             - Returns information about regression parameters in an IQMmodel
% 
% DataNLMEInfo
% 	IQMinfoNLMEdata                                        - Provides information mapping ADM and doses, YTYPE and readouts
% 
%     
% NLME Parameter Estimation Tools Interfaces
% ==========================================
% MONOLIX
% 	IQMcreateMLXTRANfile                                   - Creates an MLXTRAN structural model file based on the IQMmodel and the dosing information
% 	IQMcreateMONOLIXproject                                - Creates a Monolix/MLXTRAN project from an IQMmodel and an IQMdosing scheme
% 	IQMrunMONOLIXproject                                   - Runs a specified Monolix project
% 	IQMrunMONOLIXprojectFolder                             - Runs all the Monolix projects in the specified folder
% 	IQMsampleMONOLIXparam                                  - Samples parameters from both uncertainty and variability distributions from a Monolix fit
% 	isMONOLIXprojectIQM                                    - Checks if given project path is a Monolix project
% 
% NONMEM
% 	IQMcreateNONMEMproject                                 - Creates a NONMEM project from an IQMmodel and an IQMdosing scheme
% 	IQMcreateNONMEMresultsTable                            - Parses the results of a NONMEM run and reports them in a similar manner as in the MONOLIX pop_parameters.txt file
% 	IQMplotConvergenceNONMEM                               - Plots the convergence plots for NONMEM 
% 	IQMrunNONMEMproject                                    - Runs a specified NONMEM project
% 	IQMrunNONMEMprojectFolder                              - Runs all the NONMEM projects in the specified folder
% 	IQMsampleNONMEMparam                                   - Samples parameters from both uncertainty and variability distributions from a NONMEM fit
% 	isNONMEMprojectIQM                                     - Checks if given project path is a NONMEM project
% 
% NLME Modeling Tools
% ===================
% NMLEprojectHandling
% 	IQMcreateNLMEmodelGENcode                              - Generates code that can be used to generate a NONMEM or MONOLIX NLME  project
% 	IQMcreateNLMEproject                                   - Creates a NONMEM or MONOLIX roject from an IQMmodel and an IQMdosing scheme
%   IQMcreateGeneralLinearNLMEproject                      - This function allows to generate a general linear model using NONMEM or MONOLIX
% 	IQMgetNLMEfitIndivPopMeanParam                         - Returns the individual population mean parameters for all subjects in the NLME fit
% 	IQMgetNLMEfitIndivparam                                - Returns the individual parameters from an NLME fit (NONMEM or MONOLIX)
% 	IQMgetNLMEparameterResults                             - Parses an NLME project folder and returns the parameter estimates and additional information
% 	IQMrunNLMEproject                                      - Runs a specified NLME project (NONMEM or MONOLIX)
% 	IQMrunNLMEprojectFolder                                - Runs all NLME projects (NONMEM and MONOLIX) in the specified folder
% 	IQMsampleNLMEfitParam                                  - Samples parameters from both uncertainty and variability distributions from a NLME fit
% 	isNLMEprojectIQM                                       - Checks if given project path is an NMLE project folder (NONMEM or MONOLIX)
% 
% NLMEfitAnalysis
% 	IQMfitanalysisETAvsCOV                                 - Plot the individual variations over covariates and categorical covariates
% 	IQMfitanalysisGOFplots                                 - Produces several plots that can be used for checking the goodness of fit
% 	IQMfitanalysisGeneralPlots                             - Wrapper for different fit analysis functions that plot things that are independent of a specific output of the model
% 	IQMfitanalysisIndividualFits                           - Plots individual fits and population prediction against observed data over time
% 	IQMfitanalysisOutlierDetection                         - Searches for outliers and displays info about them
% 	IQMfitanalysisOutputPlots                              - Wrapper for different fit analysis functions that plot things that are dependent of a specific output of the model
% 	IQMfitanalysisRandomEffects                            - Plots information about the random effects 
% 
% NLMEfitSummary
% 	IQMfitsummaryAll                                       - Wrapper for IQMfitsummaryMetrics, IQMfitsummaryParameters, IQMfitsummaryCovariances, IQMfitsummaryCovariates 
% 	IQMfitsummaryCovariances                               - Generates information about covariance estimates of all projects in the same folder
% 	IQMfitsummaryCovariates                                - Generates information about covariate estimates of all projects in the same folder
% 	IQMfitsummaryMetrics                                   - Generates information about model metrics of all projects in the same folder
% 	IQMfitsummaryParameters                                - Generates information about parameter estimates of all projects in the same folder
% 	IQMfitsummaryTable                                     - Creates a typical report-type NLME model parameter table
% 
% NLMEtools
% 	IQMassessInformationContent                            - Allows to predict the information content in data of future studies, given the planned dosing and observation schedule
% 	IQMbootstrap                                           - Runs a bootstrap analysis on the provided NLME project folder
% 	IQMcompareModels                                       - Allows to compare the structural models for different estimation results from NLME models
% 	IQMcovariateAssessmentUncertainty                      - Assesses the changes that a covariates introduces on the model parameters, relative to a reference individual
% 	IQMcreateVPC                                           - Generates a VPC for a given NLMEproject
% 	IQMcreateVPCplot                                       - Plots a VPC without the need to re-run the simulations
% 	IQMcreateVPCstratified                                 - Generates a stratified VPC for a given NLMEproject
% 	IQMduplicateNLMEmodel                                  - Duplicates a NLME model, specified by the path to its project folder (modelSource) to a new path (modelDestination)
% 	IQMscm                                                 - Stepwise covariate search, using forward inclusion / backward  elimination
% 	IQMtrialGroupSimulation                                - Performs a trial simulation for a single treatment arm
% 
% IQM Tools Workflow Tools
% ========================
% PopPKmodel
% 	IQMbuildPopPKModelSpace                                - Assess a user defined popPK model subspace
% 	IQMcleanPopPKdataWrapper                               - Wrapper for different cleaning functions for standard popPK datasets
% 	IQMcomparePopPKmodels                                  - Allows to compare different popPK models created with the popPK workflow in IQM Tools
% 	IQMcreatePopPKstratifiedVPC                            - Creates a stratified VPC for a given model on a given dataset
% 	IQMscmPopPK                                            - Stepwise covariate search, using forward inclusion / backward  elimination
% 	IQMsimulatePopPKmodel                                  - Allows to simulate PK models for user defined dosing schemes
% 
% Simplified PopPK workflow
%   IQMsimplifiedPopPKinit 								   - Initialize simplified popPK workflow
% 	IQMsimplifiedPopPKcheckData 						   - Check dataset format and do some preparations
%   IQMsimplifiedPopPKexploreData 						   - Explore data 
%   IQMsimplifiedPopPKgetNLMEdata 						   - Get NLME dataset and header info
%   IQMsimplifiedPopPKmodel 							   - Build popPK model functions
%
% PopPDmodel
% 	IQMcleanPopPDdataWrapper                               - Wrapper for different cleaning functions for standard popPD datasets
% 
% Auxiliary
% =========
% 	lookforIQMP                                            - Allows to search IQM Pro folders recursively for a given text

% <<<COPYRIGHTSTATEMENT - IQM TOOLS PRO>>>
```
