## Supplementary material for "A systems biology analysis of adrenergically stimulated adiponectin exocytosis in white adipocytes": All scripts used in this work: installation.txt

Installation of IQM Tools Pro
=============================

When you are reading this document, you already have successfully obtained IQM Tools Pro from somewhere.
Please perform the following steps to install it:

- Move the IQMpro folder to the same folder in which IQMlite is located
- Start MATLAB
- Install first IQMlite
- Change into the IQMpro folder
- Open the file "SETUP_PATHS_TOOLS_IQMPRO.m" in the editor, read its documentation and modify setup accordingly (if needed). Then save and close the file
- Run "installIQMpro.m" script
- If you have the parallel toolbox for MATLAB, please do the following setting
    - Home->parallel->parallel preferences deselect the checkbox "Automatically create a parallel pool" 

Note: The installation of IQM Tools Pro does not save the MATLAB path. This means that everytime you start MATLAB you will need to 
      execute the "installIQMpro" script. This procedure has been chosen on purpose, for compliance and reproducibility reasons.

Requirements:
- MATLAB >=R2013b is required. 
- MATLAB >=R2015b is suggested.
If using MATLAB R2014b or R2015a, the export of figures to files is VERY slow. Please avoid using these versions.
