## Supplementary material for "A systems biology analysis of adrenergically stimulated adiponectin exocytosis in white adipocytes": All scripts used in this work: installIQMpro.m

```
function [] = installIQMpro()
% installIQMpro
% This script installs the PRO version of the IQM Suite of modeling tools.
%
%       installIQMpro
%
% The function will check for an installation of IQM Tools Lite. If not 
% present, and error will be shown. It also will check for a previous
% installation of IQM Tools Pro and in this case issue an error as well.
% The reasons for this are: IQM Tools Lite needs to be present for the
% functions in the PRO part. MATLAB allows simultaneous installations of
% the same toolboxes, which might lead to issues with different versions.

% <<<COPYRIGHTSTATEMENT - IQM TOOLS PRO>>>

% Check correct starting folder
currentDir = pwd;
installIQMproDir = fileparts(which('installIQMpro.m'));
if ~strcmp(currentDir,installIQMproDir),
    error('Run the ''installIQMpro'' script from the folder where it is located.');
end

% Check that IQM Tools Lite is installed
IGMLiteVer = ver('IQMlite');
if isempty(IGMLiteVer),
    error('Please install the "IQM Tools Lite" first.');
end

% Check that correct local path (network paths are not allowed)
if strcmp(currentDir(1:2),'\\'),
    error(sprintf('The installation can not be run from a network path (\\\\...).\nPlease run the installation from a local path.'));
end

% Check if IQM Pro already installed
IGMProVer = ver('IQMpro');
if length(IGMProVer) >= 1,
    error('"IQM Tools Pro" already installed. Please use "restoredefaultpath" before installing a different version.');
end

% Add IQM Tools Pro to the path 
addpath(genpath(pwd));

% Compile and install the needed packages 
PATH_IQMPRO = pwd();
if ~ispc,
    cd(fileparts(which('buildCVODElibunix.m')));
    buildCVODElibunix
    cd(PATH_IQMPRO);
else
    cd(fileparts(which('buildCVODElib.m')));
    buildCVODElib
    cd(PATH_IQMPRO);
end

% Message
disp(' ');
disp('IQM Tools Pro');
disp('	- Developer: IntiQuan GmbH');
disp('	- Installation completed');
disp(' ');
disp(' ');
disp('License IQM Tools Pro:');
disp(' ');
disp(regexprep(fileread('license.txt'),'\r',''));


try
    % Check version number (>=R2013B required)
    if verLessThan('matlab','8.2.0'),
        warning('All Pharmacometrics related functionality in IQM Tools Pro require >=MATLAB R2013B.');
    end
end
```
