## Supplementary material for "A systems biology analysis of adrenergically stimulated adiponectin exocytosis in white adipocytes": All scripts used in this work: release.txt

Release Notes IQM Tools Pro
===========================

Version 1.2.2 (02.01.2017)
--------------------------
- Minor big fixes
- Improved graphical output

Version 1.2.1 (30.04.2016)
--------------------------
- Minor bug fixes
- Error message when MONOLIX 2006R1 is used - will be handled in V1.3
- Use of the IGNORE column for data cleaning (without removal of records from the dataset)
- Allow an NLME tool not to be present and still run IQMrunNLMEprojectFolder without error (not running the tool obviously)

Version 1.2 (13.04.2016)
------------------------
- Minor bug fixes
- Added handling of sequential 0th/1st order absorption 
- Added handling of non-numeric lag time definitions and their handling in IQMdosing schemes
- Support to convert NRDOSES and INTERVAL to ADDL and II in NLME dataset
- $SIZES in NONMEM control files is now set automatically
- Added support for automatic generation of general linear models in both NONMEM (ADVAN5/7) and MONOLIX
- Update to also be compatible with MONOLIX 4.3.3
- Added support for handling of time dependent categorical covariates (limited for now to NONMEM and analytic models in the popPK workflow)
- Added loess smoothing to some GOF plots

Version 1.1 (10.02.2016)
------------------------
- Minor bug fixes
- Allow covariate centering in popPK workflowAdded
- Allow zero order absorption time to be estimated
- Included possibility for zero and mixed order absorption into the popPK workflow
- Added simplified popPK workflow

Version 1.0 (09.11.2015)
------------------------
- First full release of IQM Pro - the follow-up of SBPOP and SBTOOLBOX2
- Many improvements with respect to SBPOP (stability, usability, documentation, functionality)
- Where SBPOP used the "dataset" type, IQM Tools uses the "table" type
- IQM Tools does not require the presence of the stats toolbox 
- Parallel computation is used in several functions if the parallel toolbox is available
- Recommended use of MATLAB >= R2015b (>=R2013B is required)
- Implemented "compliance mode" for time and information stamps on figures, tables, text
- Pre Monolix 4.3.2 not supported anymore
- Support for NONMEM >= 7.2
- Improved support for pharmacometric work with a new general dataset format
- Many new tutorials available from the www.intiquan.com website, documenting the
  use of IQM Tools Pro for pharmacometric work

  
