## Supplementary material for "A systems biology analysis of adrenergically stimulated adiponectin exocytosis in white adipocytes": All scripts used in this work: SETUP_PATHS_TOOLS_IQMPRO.m

```
% In this file you need to provide the names and potentially the paths to
% the executables for tools, such as NONMEM and MONOLIX.
% 
% If executables in the system path, then only the names of the executables are needed.
%
% It is possible to define paths for Windows and Unix (use Unix for Mac) independently,
% allowing to use the package on different systems without the need to
% re-edit the paths. If a tool is not available on a system, then just
% leave the path empty.

% In the case that a queuing system is used (only under unix),
% then please provide here the command that reads out the queue. It is 
% assumed that the command that runs a NONMEM or MONOLIX job via the queue
% returns the jobID as a number. It is also assumed that this number
% appears when the queue status command is called:
PATH_SYSTEM_QUEUE_STATUS_UNIX               = '';
% If no queuing system used then keep this variable empty ('')

% NONMEM (currently tested version: 7.2 and 7.3)
PATH_SYSTEM_NONMEM_WINDOWS                  = 'nmfe73';
PATH_SYSTEM_NONMEM_UNIX                     = 'nmfe73';                            

% NONMEM PARALLEL
PATH_SYSTEM_NONMEM_PARALLEL_WINDOWS         = 'nmfe73par';
PATH_SYSTEM_NONMEM_PARALLEL_UNIX            = 'nmfe73par';                         

% nmfe73par is assumed to be a shell script with the following calling
% syntax:   "nmfe73par NRNODES controlfile outputfile". If you do not have
% one available, please ask your sysadmin to generate on for you

% MONOLIX STANDALONE (ONLY VERSION 4.3.2 and 4.3.3 supported at the moment)
PATH_SYSTEM_MONOLIX_WINDOWS                 = 'C:\INSTALL\Monolix\Monolix432s\bin\Monolix.bat';
PATH_SYSTEM_MONOLIX_UNIX                    = '.../bin/Monolix.sh';                % Define UNIX path for Mac
% MONOLIX 4.3.3 on Mac typically installs in the following folder. If you have got a Mac,
% and have Monolix 4.3.3 installed, uncomment the following line:
% PATH_SYSTEM_MONOLIX_UNIX = '/Applications/Monolix-4.3.3s.app/Contents/MacOS/run_monolix.sh';

% MONOLIX STANDALONE PARALLEL (version >= 4.3.2)
PATH_SYSTEM_MONOLIX_PARALLEL_WINDOWS        = '';
PATH_SYSTEM_MONOLIX_PARALLEL_UNIX           = '';               				% Define UNIX path for Mac

% Name of MATLAB parallel toolbox profile
MATLABPOOL_PROFILE                          = 'local';

% Preferred ordering criterion for tables of NLME estimates
% Only impacts dislay (ordering) in output tables)
NLME_ORDER_CRITERION                        = 'BIC'; % Alternative: 'AIC', 'BIC', or 'OBJ'

% Define default number of processors to use in case that parallel toolbox available
N_PROCESSORS_PAR                            = 4;
```
