## Supplementary material for "A systems biology analysis of adrenergically stimulated adiponectin exocytosis in white adipocytes": All scripts used in this work: natsortfiles_doc.html

NATSORTFILES Examples 

### NATSORTFILES Examples

The function NATSORTFILES sorts a cell array of filenames or filepaths (1xN char), taking into account any number values within the strings. This is known as a *natural order sort* or an *alphanumeric sort*. Note that MATLAB's inbuilt SORT function sorts the character codes only (as does SORT in most programming languages).

NATSORTFILES is not a naive natural-order sort, but splits and sorts filenames and file extensions separately, which means that NATSORTFILES sorts shorter filenames before longer ones: this is known as a *dictionary sort*. For the same reason filepaths are split at every path-separator character (either '\' or '/'), and each directory level is sorted separately. See the "Explanation" sections below for more details.

For sorting the rows of a cell array of strings use NATSORTROWS.

For sorting a cell array of strings use NATSORT.

#### Contents

- Basic Usage
- Output 2: Sort Index
- Output 3: Debugging Array
- Example with DIR and a Cell Array
- Example with DIR and a Structure
- Explanation: Dictionary Sort
- Explanation: Filenames
- Explanation: Filepaths
- Regular Expression: Decimal Numbers, E-notation, +/- Sign
- Regular Expression: Interactive Regular Expression Tool

#### Basic Usage

By default NATSORTFILES interprets consecutive digits as being part of a single integer, each number is considered to be as wide as one letter:

```
A = {'a2.txt', 'a10.txt', 'a1.txt'};
sort(A)
natsortfiles(A)
```

```
ans = 
    'a1.txt'    'a10.txt'    'a2.txt'
ans = 
    'a1.txt'    'a2.txt'    'a10.txt'
```

#### Output 2: Sort Index

The second output argument is a numeric array of the sort indices ndx, such that Y = X(ndx) where Y = natsortfiles(X):

```
[~,ndx] = natsortfiles(A)
```

```
ndx =
     3     1     2
```

#### Output 3: Debugging Array

The third output is a cell vector of cell arrays, where each cell array contains individual characters and numbers (after converting to numeric). This is useful for confirming that the numbers are being correctly identified by the regular expression. The cells of the cell vector correspond to the split directories, filenames, and file extensions. Note that the rows of the debugging cell arrays are linearly indexed from the input cell array.

```
[~,~,dbg] = natsortfiles(A);
dbg{:}
```

```
ans = 
    'a'    [ 2]
    'a'    [10]
    'a'    [ 1]
ans = 
    '.'    't'    'x'    't'
    '.'    't'    'x'    't'
    '.'    't'    'x'    't'
```

#### Example with DIR and a Cell Array

One common situation is to use DIR to identify files in a folder, sort them into the correct order, and then loop over them: below is an example of how to do this. Remember to preallocate all output arrays before the loop!

```
D = 'natsortfiles_test'; % directory path
S = dir(fullfile(D,'*.txt')); % get list of files in directory
N = natsortfiles({S.name}); % sort file names into order
for k = 1:numel(N)
	disp(fullfile(D,N{k}))
end
```

```
natsortfiles_test\A_1.txt
natsortfiles_test\A_1-new.txt
natsortfiles_test\A_1_new.txt
natsortfiles_test\A_2.txt
natsortfiles_test\A_3.txt
natsortfiles_test\A_10.txt
natsortfiles_test\A_100.txt
natsortfiles_test\A_200.txt
```

#### Example with DIR and a Structure

Users who need to access the DIR structure fields can use NATSORTFILE's second output to sort DIR's output structure into the correct order:

```
D = 'natsortfiles_test'; % directory path
S = dir(fullfile(D,'*.txt')); % get list of files in directory
[~,ndx] = natsortfiles({S.name}); % indices of correct order
S = S(ndx); % sort structure using indices
for k = 1:numel(S)
	fprintf('%-13s%s\n',S(k).name,S(k).date)
end
```

```
A_1.txt      22-Jul-2017 09:13:23
A_1-new.txt  22-Jul-2017 09:13:23
A_1_new.txt  22-Jul-2017 09:13:23
A_2.txt      22-Jul-2017 09:13:23
A_3.txt      22-Jul-2017 09:13:23
A_10.txt     22-Jul-2017 09:13:23
A_100.txt    22-Jul-2017 09:13:23
A_200.txt    22-Jul-2017 09:13:23
```

#### Explanation: Dictionary Sort

Filenames and file extensions are separated by the extension separator: the period character '.'. Using a normal SORT the period gets sorted *after* all of the characters from 0 to 45 (including !"#$%&'()\*+,-, the space character, and all of the control characters, e.g. newlines, tabs, etc). This means that a naive SORT or natural-order sort will sort some short filenames after longer filenames. In order to provide the correct dictionary sort, with shorter filenames first, NATSORTFILES splits and sorts filenames and file extensions separately:

```
B = {'test_ccc.m'; 'test-aaa.m'; 'test.m'; 'test.bbb.m'};
sort(B) % '-' sorts before '.'
natsort(B) % '-' sorts before '.'
natsortfiles(B) % correct dictionary sort
```

```
ans = 
    'test-aaa.m'
    'test.bbb.m'
    'test.m'
    'test_ccc.m'
ans = 
    'test-aaa.m'
    'test.bbb.m'
    'test.m'
    'test_ccc.m'
ans = 
    'test.m'
    'test-aaa.m'
    'test.bbb.m'
    'test_ccc.m'
```

#### Explanation: Filenames

NATSORTFILES combines a dictionary sort with a natural-order sort, so that the number values within the filenames are taken into consideration:

```
C = {'test2.m'; 'test10-old.m'; 'test.m'; 'test10.m'; 'test1.m'};
sort(C) % Wrong numeric order.
natsort(C) % Correct numeric order, but longer before shorter.
natsortfiles(C) % Correct numeric order and dictionary sort.
```

```
ans = 
    'test.m'
    'test1.m'
    'test10-old.m'
    'test10.m'
    'test2.m'
ans = 
    'test.m'
    'test1.m'
    'test2.m'
    'test10-old.m'
    'test10.m'
ans = 
    'test.m'
    'test1.m'
    'test2.m'
    'test10.m'
    'test10-old.m'
```

#### Explanation: Filepaths

For the same reason, filepaths are split at each file path separator character (both '/' and '\' are considered to be file path separators) and every level of directory names are sorted separately. This ensures that the directory names are sorted with a dictionary sort and that any numbers are taken into consideration:

```
D = {'A2-old\test.m';'A10\test.m';'A2\test.m';'AXarchive.zip';'A1\test.m'};
sort(D) % Wrong numeric order, and '-' sorts before '\':
natsort(D) % correct numeric order, but longer before shorter.
natsortfiles(D) % correct numeric order and dictionary sort.
```

```
ans = 
    'A10\test.m'
    'A1\test.m'
    'A2-old\test.m'
    'A2\test.m'
    'AXarchive.zip'
ans = 
    'A1\test.m'
    'A2-old\test.m'
    'A2\test.m'
    'A10\test.m'
    'AXarchive.zip'
ans = 
    'AXarchive.zip'
    'A1\test.m'
    'A2\test.m'
    'A2-old\test.m'
    'A10\test.m'
```

#### Regular Expression: Decimal Numbers, E-notation, +/- Sign

NATSORTFILES is a wrapper for NATSORT, which means all of NATSORT's options are also supported. In particular the number recognition can be customized to detect numbers with decimal digits, E-notation, a +/- sign, or other specific features. This detection is defined by providing an appropriate regular expression: see NATSORT for details and examples.

```
E = {'test24.csv','test1.8.csv','test5.csv','test3.3.csv','test12.csv'};
natsortfiles(E,'\d+\.?\d*')
```

```
ans = 
    'test1.8.csv'    'test3.3.csv'    'test5.csv'    'test12.csv'    'test24.csv'
```

#### Regular Expression: Interactive Regular Expression Tool

Regular expressions are powerful and compact, but getting them right is not always easy. One assistance is to download my interactive tool IREGEXP, which lets you quickly try different regular expressions and see all of REGEXP's outputs displayed and updated as you type.

Published with MATLAB® 7.11
