## Supplementary material for "A systems biology analysis of adrenergically stimulated adiponectin exocytosis in white adipocytes": All scripts used in this work: natsort.m

```
function [X,ndx,dbg] = natsort(X,xpr,varargin) %#ok<*SPERR>
% Alphanumeric / Natural-Order sort the strings in a cell array of strings (1xN char).
%
% (c) 2012 Stephen Cobeldick
%
% Alphanumeric sort of a cell array of strings: sorts by character order
% and also by the values of any numbers that are within the strings. The
% default is case-insensitive ascending with integer number substrings:
% optional inputs control the sort direction, case sensitivity, and number
% matching (see the section "Number Substrings" below).
%
%%% Example:
% X = {'x2', 'x10', 'x1'};
% sort(X)
%  ans =  'x1'  'x10'  'x2'
% natsort(X)
%  ans =  'x1'  'x2'  'x10'
%
%%% Syntax:
%  Y = natsort(X)
%  Y = natsort(X,xpr)
%  Y = natsort(X,xpr,<options>)
% [Y,ndx] = natsort(X,...)
% [Y,ndx,dbg] = natsort(X,...)
%
% To sort filenames or filepaths use NATSORTFILES (File Exchange 47434).
% To sort the rows of a cell array of strings use NATSORTROWS (File Exchange 47433).
%
% See also NATSORTFILES NATSORTROWS SORT CELLSTR IREGEXP REGEXP SSCANF INTMAX
%
%% Number Substrings %%
%
% By default consecutive digit characters are interpreted as an integer.
% The optional regular expression pattern <xpr> permits the numbers to also
% include a +/- sign, decimal digits, exponent E-notation, or any literal
% characters, quantifiers, or look-around requirements. For more information:
% http://www.mathworks.com/help/matlab/matlab_prog/regular-expressions.html
%
% The substrings are then parsed by SSCANF into numeric variables, using
% either the *default format '%f' or the user-supplied format specifier.
%
% This table shows some example regular expression patterns for some common
% notations and ways of writing numbers (see section "Examples" for more):
%
% <xpr> Regular | Number Substring | Number Substring              | SSCANF
% Expression:   | Match Examples:  | Match Description:            | Format Specifier:
% ==============|==================|===============================|==================
% *         \d+ | 0, 1, 234, 56789 | unsigned integer              | %f  %u  %lu  %i
% --------------|------------------|-------------------------------|------------------
%     (-|+)?\d+ | -1, 23, +45, 678 | integer with optional +/- sign| %f  %d  %ld  %i
% --------------|------------------|-------------------------------|------------------
%     \d+\.?\d* | 012, 3.45, 678.9 | integer or decimal            | %f
% --------------|------------------|-------------------------------|------------------
%   \d+|Inf|NaN | 123, 4, Inf, NaN | integer, infinite or NaN value| %f
% --------------|------------------|-------------------------------|------------------
%  \d+\.\d+e\d+ | 0.123e4, 5.67e08 | exponential notation          | %f
% --------------|------------------|-------------------------------|------------------
%  0[0-7]+      | 012, 03456, 0700 | octal prefix & notation       | %o  %i
% --------------|------------------|-------------------------------|------------------
%  0X[0-9A-F]+  | 0X0, 0XFF, 0X7C4 | hexadecimal prefix & notation | %x  %i
% --------------|------------------|-------------------------------|------------------
%  0B[01]+      | 0B101, 0B0010111 | binary prefix & notation      | %b   (not SSCANF)
% --------------|------------------|-------------------------------|------------------
%
% The SSCANF format specifier (including %b) can include literal characters
% and skipped fields. The octal, hexadecimal and binary prefixes are optional.
% For more information: http://www.mathworks.com/help/matlab/ref/sscanf.html
%
%% Debugging Output Array %%
%
% The third output is a cell array <dbg>, to check if the numbers have
% been matched by the regular expression <rgx> and converted to numeric
% by the SSCANF format. The rows of <dbg> are linearly indexed from <X>:
%
% [~,~,dbg] = natsort(X)
% dbg = 
%    'x'    [ 2]
%    'x'    [10]
%    'x'    [ 1]
%
%% Relative Sort Order %%
%
% The sort order of the number substrings relative to the characters
% can be controlled by providing one of the following string options:
%
% Option Token:| Relative Sort Order:                 | Example:
% =============|======================================|====================
% 'beforechar' | numbers < char(0:end)                | '1' < '#' < 'A'
% -------------|--------------------------------------|--------------------
% 'afterchar'  | char(0:end) < numbers                | '#' < 'A' < '1'
% -------------|--------------------------------------|--------------------
% 'asdigit'   *| char(0:47) < numbers < char(48:end)  | '#' < '1' < 'A'
% -------------|--------------------------------------|--------------------
%
% Note that the digit characters have character values 48 to 57, inclusive.
%
%% Examples %%
%
%%% Multiple integer substrings (e.g. release version numbers):
% B = {'v10.6', 'v9.10', 'v9.5', 'v10.10', 'v9.10.20', 'v9.10.8'};
% sort(B)
%  ans =  'v10.10'  'v10.6'  'v9.10'  'v9.10.20'  'v9.10.8'  'v9.5'
% natsort(B)
%  ans =  'v9.5'  'v9.10'  'v9.10.8'  'v9.10.20'  'v10.6'  'v10.10'
%
%%% Integer, decimal or Inf number substrings, possibly with +/- signs:
% C = {'test+Inf', 'test11.5', 'test-1.4', 'test', 'test-Inf', 'test+0.3'};
% sort(C)
%  ans =  'test'  'test+0.3'  'test+Inf'  'test-1.4'  'test-Inf'  'test11.5'
% natsort(C, '(-|+)?(Inf|\d+\.?\d*)')
%  ans =  'test'  'test-Inf'  'test-1.4'  'test+0.3'  'test11.5'  'test+Inf'
%
%%% Integer or decimal number substrings, possibly with an exponent:
% D = {'0.56e007', '', '4.3E-2', '10000', '9.8'};
% sort(D)
%  ans =  ''  '0.56e007'  '10000'  '4.3E-2'  '9.8'
% natsort(D, '\d+\.?\d*(E(+|-)?\d+)?')
%  ans =  ''  '4.3E-2'  '9.8'  '10000'  '0.56e007'
%
%%% Hexadecimal number substrings (possibly with '0X' prefix):
% E = {'a0X7C4z', 'a0X5z', 'a0X18z', 'aFz'};
% sort(E)
%  ans =  'a0X18z'  'a0X5z'  'a0X7C4z'  'aFz'
% natsort(E, '(?<=a)(0X)?[0-9A-F]+', '%x')
%  ans =  'a0X5z'  'aFz'  'a0X18z'  'a0X7C4z'
%
%%% Binary number substrings (possibly with '0B' prefix):
% F = {'a11111000100z', 'a0B101z', 'a0B000000000011000z', 'a1111z'};
% sort(F)
%  ans =  'a0B000000000011000z'  'a0B101z'  'a11111000100z'  'a1111z'
% natsort(F, '(0B)?[01]+', '%b')
%  ans =  'a0B101z'  'a1111z'  'a0B000000000011000z'  'a11111000100z'
%
%%% UINT64 number substrings (with full precision!):
% natsort({'a18446744073709551615z', 'a18446744073709551614z'}, [], '%lu')
%  ans =   'a18446744073709551614z'  'a18446744073709551615z'
%
%%% Case sensitivity:
% G = {'a2', 'A20', 'A1', 'a10', 'A2', 'a1'};
% natsort(G, [], 'ignorecase') % default
%  ans =   'A1'  'a1'  'a2'  'A2'  'a10'  'A20'
% natsort(G, [], 'matchcase')
%  ans =   'A1'  'A2'  'A20'  'a1'  'a2'  'a10'
%
%%% Sort direction:
% H = {'2', 'a', '3', 'B', '1'};
% natsort(H, [], 'ascend') % default
%  ans =   '1'  '2'  '3'  'a'  'B'
% natsort(H, [], 'descend')
%  ans =   'B'  'a'  '3'  '2'  '1'
%
%%% Relative sort-order of number substrings compared to characters:
% V = num2cell(char(32+randperm(63)));
% cell2mat(natsort(V, [], 'asdigit')) % default
%  ans = '!"#$%&'()*+,-./0123456789:;<=>?@ABCDEFGHIJKLMNOPQRSTUVWXYZ[\]^_'
% cell2mat(natsort(V, [], 'beforechar'))
%  ans = '0123456789!"#$%&'()*+,-./:;<=>?@ABCDEFGHIJKLMNOPQRSTUVWXYZ[\]^_'
% cell2mat(natsort(V, [], 'afterchar'))
%  ans = '!"#$%&'()*+,-./:;<=>?@ABCDEFGHIJKLMNOPQRSTUVWXYZ[\]^_0123456789'
%
%% Input and Output Arguments %%
%
%%% Inputs (*=default):
%   X   = CellArrayOfCharRowVectors, to be sorted into natural-order.
%   xpr = CharRowVector, regular expression for number substrings, '\d+'*.
% <options> tokens can be entered in any order, as many as required:
%   - Sort direction: 'descend'/'ascend'*.
%   - Case sensitive/insensitive matching: 'matchcase'/'ignorecase'*.
%   - Relative sort of numbers: 'beforechar'/'afterchar'/'asdigit'*.
%   - The SSCANF number conversion format, e.g.: '%x', '%i', '%f'*, etc.
%
%%% Outputs:
%   Y   = CellArrayOfCharRowVectors, <X> sorted into natural-order.
%   ndx = NumericArray, such that Y = X(ndx). The same size as <X>.
%   dbg = CellArray of the parsed characters and number values. Each row is
%         one input char vector, linear-indexed from <X>. To help debug <xpr>.
%
% [X,ndx,dbg] = natsort(X,xpr*,<options>)

%% Input Wrangling %%
%
assert(iscell(X),'First input <X> must be a cell array.')
tmp = cellfun('isclass',X,'char') & cellfun('size',X,1)<2 & cellfun('ndims',X)<3;
assert(all(tmp(:)),'First input <X> must be a cell array of char row vectors (1xN char).')
%
% Regular expression:
if nargin<2 || isnumeric(xpr)&&isempty(xpr)
	xpr = '\d+';
else
	assert(ischar(xpr)&&isrow(xpr),'Second input <xpr> must be a regular expression (char row vector).')
end
%
% Optional arguments:
tmp = cellfun('isclass',varargin,'char') & 1==cellfun('size',varargin,1) & 2==cellfun('ndims',varargin);
assert(all(tmp(:)),'All optional arguments must be char row vectors (1xN char).')
% Character case matching:
ChrM = strcmpi(varargin,'matchcase');
ChrX = strcmpi(varargin,'ignorecase')|ChrM;
% Sort direction:
DrnD = strcmpi(varargin,'descend');
DrnX = strcmpi(varargin,'ascend')|DrnD;
% Relative sort-order of numbers compared to characters:
RsoB = strcmpi(varargin,'beforechar');
RsoA = strcmpi(varargin,'afterchar');
RsoX = strcmpi(varargin,'asdigit')|RsoB|RsoA;
% SSCANF conversion format:
FmtX = ~(ChrX|DrnX|RsoX);
%
if nnz(FmtX)>1
	tmp = sprintf(', ''%s''',varargin{FmtX});
	error('Overspecified optional arguments:%s.',tmp(2:end))
end
if nnz(DrnX)>1
	tmp = sprintf(', ''%s''',varargin{DrnX});
	error('Sort direction is overspecified:%s.',tmp(2:end))
end
if nnz(RsoX)>1
	tmp = sprintf(', ''%s''',varargin{RsoX});
	error('Relative sort-order is overspecified:%s.',tmp(2:end))
end
%
%% Split Strings %%
%
% Split strings into number and remaining substrings:
[MtS,MtE,MtC,SpC] = regexpi(X(:),xpr,'start','end','match','split',varargin{ChrX});
%
% Determine lengths:
MtcD = cellfun(@minus,MtE,MtS,'UniformOutput',false);
LenZ = cellfun('length',X(:))-cellfun(@sum,MtcD);
LenY = max(LenZ);
LenX = numel(MtC);
%
dbg = cell(LenX,LenY);
NuI = false(LenX,LenY);
ChI = false(LenX,LenY);
ChA = char(double(ChI));
%
ndx = 1:LenX;
for k = ndx(LenZ>0)
	% Determine indices of numbers and characters:
	ChI(k,1:LenZ(k)) = true;
	if ~isempty(MtS{k})
		tmp = MtE{k} - cumsum(MtcD{k});
		dbg(k,tmp) = MtC{k};
		NuI(k,tmp) = true;
		ChI(k,tmp) = false;
	end
	% Transfer characters into char array:
	if any(ChI(k,:))
		tmp = SpC{k};
		ChA(k,ChI(k,:)) = [tmp{:}];
	end
end
%
%% Convert Number Substrings %%
%
if nnz(FmtX) % One format specifier
	fmt = varargin{FmtX};
	err = ['The supplied format results in an empty output from sscanf: ''',fmt,''''];
	pct = '(?<!%)(%%)*%'; % match an odd number of % characters.
	[T,S] = regexp(fmt,[pct,'(\d*)([bdiuoxfeg]|l[diuox])'],'tokens','split');
	assert(isscalar(T),'Unsupported optional argument: ''%s''',fmt)
	assert(isempty(T{1}{2}),'Format specifier cannot include field-width: ''%s''',fmt)
	switch T{1}{3}(1)
		case 'b' % binary
			fmt = regexprep(fmt,[pct,'(\*?)b'],'$1%$2[01]');
			val = dbg(NuI);
			if numel(S{1})<2 || ~strcmpi('0B',S{1}(end-1:end))
				% Remove '0B' if not specified in the format string:
				val = regexprep(val,'(0B)?([01]+)','$2','ignorecase');
			end
			val = cellfun(@(s)sscanf(s,fmt),val, 'UniformOutput',false);
			assert(~any(cellfun('isempty',val)),err)
			NuA(NuI) = cellfun(@(s)sum(pow2(s-'0',numel(s)-1:-1:0)),val);
		case 'l' % 64-bit
			NuA(NuI) = cellfun(@(s)sscanf(s,fmt),dbg(NuI)); %slow!
		otherwise % double
			NuA(NuI) = sscanf(sprintf('%s\v',dbg{NuI}),[fmt,'\v']); % fast!
	end
else % No format specifier -> double
	NuA(NuI) = sscanf(sprintf('%s\v',dbg{NuI}),'%f\v');
end
% Note: NuA's class is determined by SSCANF or the custom binary parser.
NuA(~NuI) = 0;
NuA = reshape(NuA,LenX,LenY);
%
%% Debugging Array %%
%
if nargout>2
	dbg(:) = {''};
	for k = reshape(find(NuI),1,[])
		dbg{k} = NuA(k);
	end
	for k = reshape(find(ChI),1,[])
		dbg{k} = ChA(k);
	end
end
%
%% Sort Columns %%
%
if ~any(ChrM) % ignorecase
	ChA = upper(ChA);
end
%
ide = ndx.';
% From the last column to the first...
for n = LenY:-1:1
	% ...sort the characters and number values:
	[C,idc] = sort(ChA(ndx,n),1,varargin{DrnX});
	[~,idn] = sort(NuA(ndx,n),1,varargin{DrnX});
	% ...keep only relevant indices:
	jdc = ChI(ndx(idc),n); % character
	jdn = NuI(ndx(idn),n); % number
	jde = ~ChI(ndx,n)&~NuI(ndx,n); % empty
	% ...define the sort-order of numbers and characters:
	jdo = any(RsoA)|(~any(RsoB)&C<'0');
	% ...then combine these indices in the requested direction:
	if any(DrnD) % descending
		idx = [idc(jdc&~jdo);idn(jdn);idc(jdc&jdo);ide(jde)];
	else % ascending
		idx = [ide(jde);idc(jdc&jdo);idn(jdn);idc(jdc&~jdo)];
	end
	ndx = ndx(idx);
end
%
ndx  = reshape(ndx,size(X));
X = X(ndx);
%
end
%%%%%%%%%%%%%%%%%%%%%%%%%%%%%%%%%%%%%%%%%%%%%%%%%%%%%%%%%%%%%%%%%%%%%%%%%%%natsort
```
