## Supplementary material for "A systems biology analysis of adrenergically stimulated adiponectin exocytosis in white adipocytes": All scripts used in this work: natsortfiles.m

```
function [X,ndx,dbg] = natsortfiles(X,varargin)
% Alphanumeric / Natural-Order sort of a cell array of filename/filepath strings (1xN char).
%
% (c) 2012 Stephen Cobeldick
%
% Alphanumeric sort of a cell array of filenames or filepaths: sorts by
% character order and also by the values of any numbers that are within
% the names. Filenames, file-extensions, and directories (if supplied)
% are split apart and are sorted separately: this ensures that shorter
% filenames sort before longer ones (i.e. thus giving a dictionary sort).
%
%%% Example:
% D = 'C:\Test';
% S = dir(fullfile(D,'*.txt'));
% N = natsortfiles({S.name});
% for k = 1:numel(N)
%     fullfile(D,N{k})
% end
%
%%% Syntax:
%  Y = natsortfiles(X)
%  Y = natsortfiles(X,xpr)
%  Y = natsortfiles(X,xpr,<options>)
% [Y,ndx] = natsortfiles(X,...)
% [Y,ndx,dbg] = natsortfiles(X,...)
%
% To sort all of the strings in a cell array use NATSORT (File Exchange 34464).
% To sort the rows of a cell array of strings use NATSORTROWS (File Exchange 47433).
%
% See also NATSORT NATSORTROWS SORT CELLSTR IREGEXP REGEXP SSCANF DIR FILEPARTS FULLFILE
%
%% File Dependency %%
%
% NATSORTFILES requires the function NATSORT (File Exchange 34464). The inputs
% <xpr> and <options> are passed directly to NATSORT: see NATSORT for case
% sensitivity, sort direction, numeric substring matching, and other options.
%
%% Explanation %%
%
% Using SORT on filenames will sort any of char(0:45), including the printing
% characters ' !"#$%&''()*+,-', before the file extension separator character '.'.
% Therefore this function splits the name and extension and sorts them separately.
%
% Similarly the file separator character within filepaths can cause longer
% directory names to sort before shorter ones, as char(0:46)<'/' and char(0:91)<'\'.
% NATSORTFILES splits filepaths at each file separator character and sorts
% every level of the directory hierarchy separately, ensuring that shorter
% directory names sort before longer, regardless of the characters in the names.
%
%% Examples %%
%
% A = {'a2.txt', 'a10.txt', 'a1.txt'};
% sort(A)
%  ans = 'a1.txt'  'a10.txt'  'a2.txt'
% natsortfiles(A)
%  ans = 'a1.txt'  'a2.txt'  'a10.txt'
%
% B = {'test_new.m'; 'test-old.m'; 'test.m'};
% sort(B)         % Note '-' sorts before '.':
%  ans =
%    'test-old.m'
%    'test.m'
%    'test_new.m'
% natsortfiles(B) % Shorter names before longer (dictionary sort):
%  ans =
%    'test.m'
%    'test-old.m'
%    'test_new.m'
%
% C = {'test2.m'; 'test10-old.m'; 'test.m'; 'test10.m'; 'test1.m'};
% sort(C)         % Wrong numeric order:
%  ans =
%    'test.m'
%    'test1.m'
%    'test10-old.m'
%    'test10.m'
%    'test2.m'
% natsortfiles(C) % Correct numeric order, shorter names before longer:
%  ans =
%    'test.m'
%    'test1.m'
%    'test2.m'
%    'test10.m'
%    'test10-old.m'
%
%%% Directory Names:
% D = {'A2-old\test.m';'A10\test.m';'A2\test.m';'A1archive.zip';'A1\test.m'};
% sort(D)         % Wrong numeric order, and '-' sorts before '\':
%  ans =
%    'A10\test.m'
%    'A1\test.m'
%    'A1archive.zip'
%    'A2-old\test.m'
%    'A2\test.m'
% natsortfiles(D) % Shorter names before longer (dictionary sort):
%  ans =
%    'A1archive.zip'
%    'A1\test.m'
%    'A2\test.m'
%    'A2-old\test.m'
%    'A10\test.m'
%
%% Input and Output Arguments %%
%
% See NATSORT for a full description of <xpr> and the <options>.
%
%%% Inputs (*=default):
%  X   = CellArrayOfCharRowVectors, with filenames or filepaths to be sorted.
%  xpr = CharRowVector, regular expression to detect numeric substrings, '\d+'*.
%  <options> can be supplied in any order and are passed directly to NATSORT.
%
%%% Outputs:
%  Y   = CellArrayOfCharRowVectors, filenames of <X> sorted into natural-order.
%  ndx = NumericMatrix, same size as <X>. Indices such that Y = X(ndx).
%  dbg = CellVectorOfCellArrays, size 1xMAX(2+NumberOfDirectoryLevels).
%        Each cell contains the debug cell array for directory names,
%        filenames, or file extensions. To help debug <xpr>. See NATSORT.
%
% [Y,ndx,dbg] = natsortfiles(X,*xpr,<options>)

%% Input Wrangling %%
%
assert(iscell(X),'First input <X> must be a cell array.')
tmp = cellfun('isclass',X,'char') & cellfun('size',X,1)<2 & cellfun('ndims',X)<3;
assert(all(tmp(:)),'First input <X> must be a cell array of strings (1xN character).')
%
%% Split and Sort File Names/Paths %%
%
% Split full filepaths into file [path,name,extension]:
[pth,fnm,ext] = cellfun(@fileparts,X(:),'UniformOutput',false);
% Split path into {dir,subdir,subsubdir,...}:
pth = regexp(pth,'[^/\\]+','match'); % either / or \ as filesep.
len = cellfun('length',pth);
num = max(len);
vec{numel(len)} = [];
%
% Natural-order sort of the file extensions and filenames:
if nargout<3 % faster:
	[~,ndx] = natsort(ext,varargin{:});
	[~,ids] = natsort(fnm(ndx),varargin{:});
else % for debugging:
	[~,ndx,dbg{num+2}] = natsort(ext,varargin{:});
	[~,ids,tmp] = natsort(fnm(ndx),varargin{:});
	[~,idd] = sort(ndx);
	dbg{num+1} = tmp(idd,:);
end
ndx = ndx(ids);
%
% Natural-order sort of the directory names:
for k = num:-1:1
	idx = len>=k;
	vec(:) = {''};
	vec(idx) = cellfun(@(c)c(k),pth(idx));
	if nargout<3 % faster:
		[~,ids] = natsort(vec(ndx),varargin{:});
	else % for debugging:
		[~,ids,tmp] = natsort(vec(ndx),varargin{:});
		[~,idd] = sort(ndx);
		dbg{k} = tmp(idd,:);
	end
	ndx = ndx(ids);
end
%
% Return the sorted array and indices:
ndx = reshape(ndx,size(X));
X = X(ndx);
%
end
%%%%%%%%%%%%%%%%%%%%%%%%%%%%%%%%%%%%%%%%%%%%%%%%%%%%%%%%%%%%%%%%%%%%%%%%%%%natsortfiles
```
