## Supplementary material for "A systems biology analysis of adrenergically stimulated adiponectin exocytosis in white adipocytes": All scripts used in this work: SaveFile.m

```
function [] = SaveFile(fileName,variable, variableName)
%SAVEFILE Saves a file with "fileName" with the content of a "variable" with
%name "variableName"

S.(variableName)=variable;
save(fileName,'-struct', 'S')

end
```
